# ΔSCOPE: A new method to quantify 3D biological structures and identify differences in zebrafish forebrain development

**DOI:** 10.1101/715698

**Authors:** Morgan S Schwartz, Jake Schnabl, Mackenzie P.H. Litz, Benjamin S Baumer, Michael Barresi

## Abstract

Research in the life sciences has traditionally relied on the analysis of clear morphological phenotypes, which are often revealed using increasingly powerful microscopy techniques analyzed as maximum intensity projections (MIPs). However, as biology turns towards the analysis of more subtle phenotypes, MIPs and qualitative approaches are failing to adequately describe these phenotypes. To address these limitations and quantitatively analyze the three-dimensional (3D) spatial relationships of biological structures, we developed the computational method and program called ΔSCOPE (Changes in Spatial Cylindrical Coordinate Orientation using PCA Examination). Our approach uses the fluorescent signal distribution within a 3D data set and reorients the fluorescent signal to a relative biological reference structure. This approach enables quantification and statistical analysis of spatial relationships and signal density in 3D multichannel signals that are positioned around a well-defined structure contained in a reference channel. We validated the application of ΔSCOPE by analyzing normal axon and glial cell guidance in the zebrafish forebrain and by quantifying the commissural phenotypes associated with abnormal Slit guidance cue expression in the forebrain. Despite commissural phenotypes which display disruptions to the reference structure, ΔSCOPE was able to detect subtle, previously uncharacterized changes in zebrafish forebrain midline crossing axons and glia. This method has been developed as a user-friendly, open source program. We propose that ΔSCOPE is an innovative approach to advancing the state of image quantification in the field of high resolution microscopy, and that the techniques presented here are of broad applications to the life science field.

## Introduction

Since Robert Hooke identified cells in a piece of cork, biologists’ search for patterns has been informed by qualitative observations. As microscopy and imaging techniques have advanced and generated larger and more complex data, our qualitative abilities are proving to be inadequate to extract all the information these data may hold [1, 2, 3]. The field of biology now faces a problem in which the complexity and granularity of current data collection methods has surpassed the ability of researchers to fully conceptualize all of the data collected. Moreover, the challenges of many of the phenotypes being studied in the modern era, whether slight changes in neuronal positioning in an autism spectrum disease model or the significant perturbations in the size of the brains of children infected with Zika, require greater statistical rigor to detect and quantify [4, 5, 6, 7]. To overcome these challenges, we need new computational tools to process, quantify, and statistically analyze complex 3D image-based data.

There are several prominent obstacles to analyzing 3D image datasets that need to be overcome to facilitate the acquisition of quantitative and statistical metrics. First, biological specimens within the same species and age group exhibit morphological variation [3]. Second, all image data contains a subset of positive pixels, such as background noise or off target labeling that can create ambiguity in isolating the true signal of the sample [1]. Additionally, 3D image data has historically been visualized using maximum intensity projections (MIPs), which collapse the third dimension of the data in order to present the image in a form that is easier to visualize and conceptualize. Unfortunately, this compression leads to a loss of information that may be critical to detecting both coarse and subtle phenotypes. Advances in data visualization software and computational power have begun to enable a shift away from MIPs and towards analyzing the whole 3D data sets. However, these techniques often rely on either machine learning that lacks descriptive ability or on a process that warps the data to fit a model, both of which have the potential to introduce new errors [8, 3, 1, 2]. Finally, experimental variability during image collection can further complicate phenotype interpretations [3]. Taken together, the variation contributed by both the natural biological and experimental preparations paired with the loss of data from image projections has traditionally made obtaining meaningful statistical metrics of biological phenomena intractable.

Since the emergence of the light microscope, a range of advances in biological imaging have occurred, enabling the acquisition of high resolution data sets of 3D biological structures. These advances however, have intensified the need for more powerful methods to measure changes within and between samples. Currently there are three main classes of image-based analysis: visualization, filter-based analysis and machine learning classification [9]. Tools for visualization have been critical to render 3D-volumetric data; however, 3D visualization tools, such as Amira [10] and Vaa3D [11, 12], rarely extend beyond rendering the data and lack tools for sample comparisons. Filter-based analysis has been dominated by the open-source program Fiji due to its ease of use and applicability to a wide variety of data types [9, 13]. While Fiji provides a variety of tools and plugins for processing and enhancing features of image data, it fails to provide rigorous options to compare images between samples. Importantly, machine learning based programs, such as ilastik [14], have excelled at classifying objects and signal within individual images, but unfortunately are unable to classify signals of whole images across an entire sample set. Attempts to overcome some of these challenges have included manually assigning each image a score that corresponds to qualitative assessments of phenotypic variation [15], yet this approach is limited by the error and bias inherent to the human eye. Moreover, such classification approaches are often performed on the MIP as opposed to considering the whole image stack, thus these approaches rarely discern subtle changes present in the data.

One such field that has come to rely heavily on image-based data is neuroscience. In particular, attempts to map the circuitry of the adult human brain have fostered the creation of some novel methods for 3D analysis. For instance, applying linear and non-linear data transformations enabled the superimposition of multiple samples of myelin histology stained sections with digitized reference brain reconstructions from Magnetic Resonance Imaging (MRI) [16]. Moreover, such MRI data has been analyzed more recently with probabilistic tractography to segment and compare the visual pathway affected in the brains of individuals with multiple sclerosis [17].

It is however a different type of challenge to investigate the embryonic origins of these neuronal pathways during development of the central nervous system (CNS). We are specifically interested in commissure development, where tightly bundled fascicles of axons cross the midline to form commissures, which offers a model to study how the two halves of the CNS of bilaterally symmetric organisms become connected [18, 19, 20, 15]. Development of these stereotypical structures is pioneered by pathfinding axons that grow towards and cross the midline in response to local and global signaling cues. Overall, commissure development represents a dynamic event with visible degrees of variation [15, 20, 18]. Unfortunately, the expected degree of biological variation paired with the additional experimental and image analysis challenges detailed above have hampered the application of robust quantification and statistical approaches to the study of commissure development.

We have taken advantage of the accessible embryonic brain of the zebrafish model system to characterize the first forming commissures during forebrain development [15, 20, 18]. The post-optic commissure (POC) is the first commissure to form, with pioneering axons projecting across the diencephalic midline as early as 23 hours post fertilization (hpf) [21], and the POC becomes tightly bundled by 30 hpf (Fig 1). The navigation of the first midline crossing axons is mediated by precisely positioned extracellular guidance cues as well as essential axon to glial cell guidance interactions [15, 19, 22, 23]. This complex array of long- and short-range factors functions to combinatorially guide pathfinding commissural axons across the midline and onward to their final synaptic target cells.

**Figure 1:**
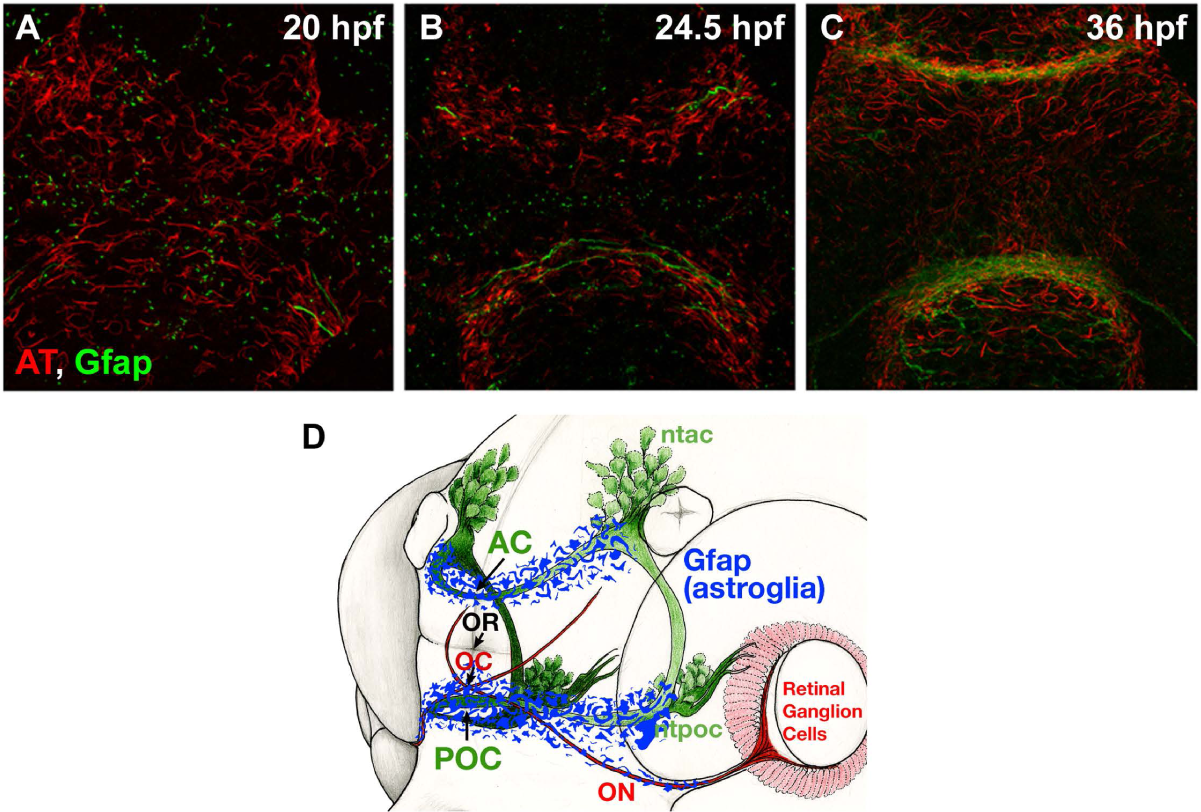
Post-optic commissure formation in zebrafish embryos. The post-optic commissure (POC) is formed by midline crossing axons (AT) in concert with a structure of glial cells called the glial bridge (Gfap). A) Frontal MIP of the zebrafish forebrain at 20 hpf labeled with anti-acetylated tubulin (AT) (green) and anti-Gfap (red). Gfap signal is distributed across the whole forebrain with the glial bridge beginning to condense in both the telencephalon (top half) and diencephalon (bottom). The first pioneering axons are visible in the diencephalon, where they will construct POC. B) Frontal MIP of the zebrafish forebrain at 24.5 hpf labeled with AT and Gfap. Axons (green) are observed pioneering the diencephalic midline, forming the POC, in concert with the glial bridge which has condensed around the forming commissure. C) Frontal MIP of the zebrafish forebrain at 36 hpf labeled with AT and Gfap. Both the diencephalic POC and telencephalic anterior commissure have been successfully constructed and positioned at the midline in concert with their respective glial bridges. D) Model of the commissure (green) and glial bridge (blue) positioning in the zebrafish forebrain with respect to the eye and dorsal and ventral clusters.

Reminiscent of the corpus collossum in the mammalian brain, the wild type zebrafish POC is composed of many tightly adhered axons that form a fascicle spanning the midline of the forebrain [21]. In addition to axon-to-axon adhesion, fasciculation of the POC is achieved in part by both the actions of repellent guidance cues, such as Slit 2 or Slit 3 ligands, that serve to limit the region of allowable space for axon exploration [24, 25, 15]. At 30hpf, the final shape of the POC resembles a curving band of axon fascicles that is 2-3 microns thick as it stretches from one side of the diencephalon to the other. Developing prior to and concomitantly with the POC is a midline spanning swath of astroglial cells, termed the “glial bridge” [15]. These glial cells are most commonly identified by their expression of Glial fibrillary acidic protein (Gfap), which is an intermediate filament found broadly in astroglial cells [26, 27, 28, 29] (Fig 1). These glial cells are known to function as both the main stem cells of the developing nervous system (known as radial glia cells) and as a supportive cellular substrate for migrating cells and pathfinding axons [21, 30, 15, 20, 18]. Although researchers have begun to identify the factors important for the guidance of commissural axons, little is known about how these factors may influence glial bridge development nor what may be required for axon-glial cell interactions during commissure formation.

In order to study the development of and relationship between POC axons and the cells of the glial bridge, we have employed immunocytochemistry (ICC) to label the two structures using antibodies against acetylated tubulin (anti-AT, axons) and Gfap (anti-Gfap, astroglia) (Fig 1). Imaging with confocal microscopy enabled us to collect high resolution 3D data sets of the labeled structures within the zebrafish forebrain. Quantification of these imaged structures has previously been limited due to wide degrees of variation in the elaboration of their final forms as well as the inconsistencies inherent to the methodology. Further complicating this analysis was the amorphous distribution of Gfap labeling, which defied confidence in any qualitative inspection. To overcome these challenges we have created a new computational method we call ΔSCOPE (Spatial Cylindrical Coordinate Orientation with PCA Examination). This method aligns the gross morphology of each biological sample in 3D space before assigning a new set of relational coordinates. This new coordinate system then serves to directly represent the biological data contained within an image relative to a model of the imaged structure, consequently enabling statistical comparisons to be performed. We validate the use of ΔSCOPE as a tool to quantify biological structures by quantitatively describing the relationship of POC axons and glial bridge cells during commissure formation, and differences between wild type commissures and mutants lacking normal axon guidance cue expression. Lastly, we show how ΔSCOPE analysis revealed a subtle role for Slit1a in facilitating POC axon-glial interactions during commissure formation. We conclude that ΔSCOPE provides a novel approach for the quantification of biological structures that we propose will be of broad application across the life sciences.

## Materials and Methods

### Zebrafish husbandry

Fish lines were maintained in the Smith College Animal Quarters according to Smith College Institutional Animal Care and Use Committee (IACUC) and AAALAC regulations. Groups of 12-15 fish were housed in 1 L tanks on an Aquaneering engineered fish facility with recycling water at a standard conditions including 1300 *µS*, pH of 7.2, a temperature of 28.5–30.0°C and a 12 h light-dark cycle with a 1 h 50% transition period before each light change. Adult zebrafish were maintained on a diet of dry fish food (Gemma micro 300; skretting) and live brine shrimp (Artemia International, Fairview, TX).

Embryos were maintained in embryo media (EM) (5 mM NaCl, 0.17 mM KCl, 0.33 mM *CaCl*_2_, 0.33 mM *MgSO*_4_ and 0.00003% methylene blue) at 28.5°C and under a 12 h light-dark cycle according to standard procedures [31]. The following genetic strains were used: wild type (AB and TU; ZIRC) and *you-too* (*gli2-DR, yot*) [32]. Homozygotic *yot* embryos were identified based on tail curvature, chevron shaped somites, and unresponsiveness to touch, which was then confirmed with genotyping as was previously described [15].

### Immunocytochemistry

Immunocytochemistry procedures were carried out as previously described [15, 33]. Briefly, embryos were fixed at 27.5–28 hpf (hours post-fertilization) with 4% formaldehyde diluted in 0.025 M phosphate buffer (PB) for 2 h or 16 h at room temperature or 4°C, respectively. Tissue penetration steps included treatment with 100% acetone for 4 minutes with a rehydration methanol series. Embryos were washed and buffered with 2% v/v triton x-100 (PBS-Tx). Embryos were blocked for 1 h at room temperature in PBS-Tx with 2% w/v bovine serum albumin fraction V, 1% v/v dimethyl sulfoxide, and 10% v/v normal goat serum (block), and then followed by primary and secondary antibody incubations for 2 h at room temperature or overnight at 4°C. Primary antibodies used included anti-rabbit glial fibrillary acidic protein (GFAP, Sigma, 1:400), mouse anti-Zrf1 (Gfap; IgG1; ZIRC 1:20) and mouse anti-acetylated tubulin (AT; IgG2b; Sigma 1:800). Secondary were all raised in goat and included anti-rabbit conjugated to Alexa 488 (Invitrogen, 1:200), anti-mouse IgG1 conjugated to Alexa 488 and anti-mouse IgG2b conjugated to Alexa 647. Labeled embryos were stored and imaged in 70% glycerol made up in 30% PBS. Samples used for comparative experiments were collected from the same clutch of embryos and immunocytochemistry was performed on all embryos at the same time. Immuno batch effects which lead to differential labeling of structures have been noted in anti-AT immunocytochemistry, which may lead to artificial detection of differences in signal due to real changes in immunolabeling. We have specifically noted that changing antibody concentrations or the length of incubation can result in significant changes in POC signal amount and distribution.

### Confocal microscopy

To visualize the POC, immunolabeled embryos were decapitated and heads mounted in 70% glycerol with the ventral forebrain oriented closest to the glass coverslip. Appropriate and consistent mounting was critical to prevent anisotropy in pixel resolution from influencing the pixel count in bins around the commissure. Samples were imaged on a Leica SP5 scanning confocal microscope at leica HC apochromat (CS2) 63X oil objective (0FN25/E) with a numerical aperture of 1.4 with a 1.5 optical zoom. Each image was collected at a 1024 by 1024 pixel resolution with an additional line averaging of 4. Z-stacks of the POC region were collected for each embryo with an optical step size of 0.21 *µm*, resulting in stacks ranging in thickness from 20 to 35 *µm*. Z step size was chosen to minimize anisotropy, with x and y pixel dimensions set at 0.169 *µm*, 0.21 *µm* was chosen approach isotropic voxel size without oversampling in the z dimension. Laser power was maintained at the following percentages for all experiments: the argon laser at 25% with a 12% intensity, the 594 nm laser at 80%, and the 633 nm laser at 20%. Image acquisition was captured bidirectionally at 600 hz.

### Pre-analysis data processing

Following imaging on the confocal microscope, images were saved in LIF files. Each sample was opened in Fiji [34] using the Bio-Formats plugin [35], cropped to eliminate background in *X* and *Y*, and rotated around the *Z* axis to position anterior as up. Each channel was isolated and saved as an individual HDF5 (.h5) file using the HDF5 plugin for Fiji [36].

Image analysis was determined to require image pre-processing to reduce noise, both biological, from AT labeled cilia, and experimental, from sample collection. We first have evaluated the use of simple thresholding of the signal intensity in raw images of 28 hpf wild type embryos labeled with anti-acetylated tubulin to remove background signal and improve the signal to noise ratio. We tested intensity thresholds at the 0, 25, 50, 75, and 100th percentiles of the intensity observed in each image to evaluate how the removal of background signal could improve the overall signal to noise ratio. Using the python SciPy modules labels (pixel distance = 3) and region props, we evaluated the number of discrete objects and the area of those objects at each intensity threshold (Fig 2). The 100th percentile signal intensity threshold dramatically reduced small area objects, like cilia, but also reduced large objects like portions of axons. In fact this thresholding approach eliminated a majority of the structure of the POC and many points of spurious non-axonal signal still remained (Fig 2 A, C-G). The inability to reduce unwanted labeling prompted us to turn to machine learning methods, like the program ilastik [14], to perform image pre-processing.

**Figure 2:**
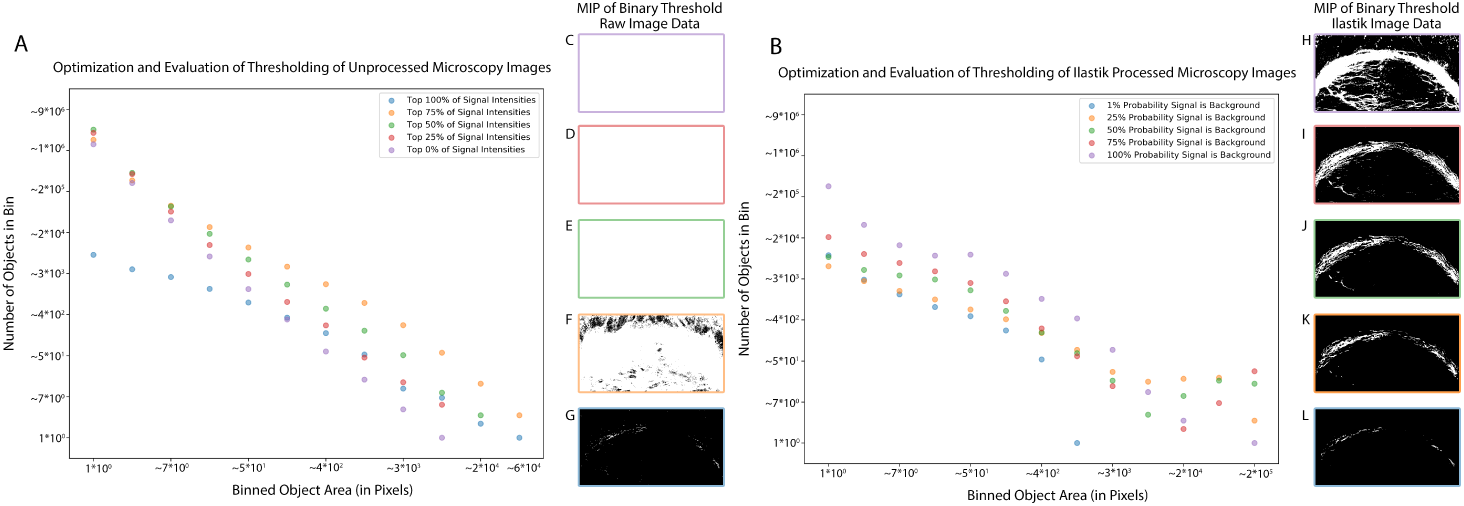
Comparison of image processing techniques and optimal threshold. ilastik processing reduces small area objects while preserving large area structures when compared to different pixel intensity thresholds of raw microscopy data. A) Results of thresholding raw microscopy data using the percentile of the pixel intensity of the image. Using the SciPy module label and region props, the areas of objects and the number of objects at each area for wild type data sets at different thresholds was plotted. B) Results of thresholding the probability that the pixel is background signal from ilastik processed microscopy data. Using the SciPy module label and region props, the areas of objects and the number of objects at each area for wild type data sets at different thresholds are plotted. Thresholds of intensities: 0% (All signal) Purple, 25% Orange, 50% Green, 25% Red, 100% (top percentile signal) Blue. C-G) Maximum Intensity Projections of thresholded raw microscopy images bordered by corresponding graph colors with C:Purple (0%)- G:Blue (100%). C-G) Maximum Intensity Projections of thresholded by p-value ilastik microscopy images bordered by corresponding graph colors with C:Purple (No Statistically significant probability (P ¡ 1) that pixels are signal)- G:Blue (P <0.01 probability that signal is background).

ilastik combines user input and a machine learning algorithm to assign a probability to each pixel that the signal observed at that point was signal of interest. For processing, an ilastik file (see S1 Data and S2 Data) was generated for either AT or Gfap processing. Training was performed using two labels, one for signal, and one for background, on a subset of samples contained within the data set of images (for samples used in training see Table 1) until the output of both labels at 80% confidence generated separation (background with no signal) between axonal filaments, signal in fine and faint axonal processes, and eliminated the majority of cilia. AT and Gfap experiment samples for all ages were processed using batch processing provided by the 28 hpf wild type ilastik AT and Gfap files. This was done to limit alternate training effects and make signal intensity comparisons comparable. Additional training for AT was required on *you-too* sample sets as the sparse labeling in these samples resulted in the inability of the ilastik program to detect any remaining axon signal. To enable visualization of this data and enable computation of the defasciculation metric, individual AT ilastik files were generated for data sets in a *you-too* background and trained on a subset of samples (for samples used in training see Table 1). This likely results in an over-reporting of signal in these data sets, but as the loss of signal was so dramatically apparent in these samples, quantification of loss of signal was deemed to be less critical than quantification of defasciculation and axon patterning. Probability images were exported and compared to the originals to ensure fidelity of signal and success in noise removal.

**Table 1:** List of samples used in ilastik training. ilastik training was performed manually by marking on a training image to denote where definitive signal and noise within the image was contained. Sample names of images used in ilastik training as they are found in the supplementary data section are included.

| AT wild type | AT *you-too* | AT tg(*slit1a yot*) | Gfap wild type |
| --- | --- | --- | --- |
| 120 | 335 | 16 | 109 |
| 112 | 332 | 12 | 102 |
| 109 | 21 | 11 | 101 |
| 101 | 3 | 8 | 114 |
| - | 1 | 6 | - |
| - | 340 | 5 | - |
| - | 337 | 2 | - |

ilastik distinguishes signal from noise by generating a probability value ranging from 0 to 1 for each pixel indicating the probability that the pixel is not background. (Fig 7 A). This method was chosen over utilization of a simple threshold of the collected images for several reasons: 1) The application of a simple threshold to the data resulted in significantly more points being included in the computation when compared to ilastik (Fig 2 A.), and also resulted in increased processing time. 2) Thresholding the data tended to linearly decrease all structural signal, resulting in the loss of real axonal signal (structures with large area), cilia (structures with small area), and background (Fig 2 B). In contrast, ilastik preferentially reduced the number of very small objects while preserving larger objects, and further, when 0.25¡p¡0.75, it reduced the number of small objects several fold while preserving more large area structures, in comparison to raw thresholding. 3) Visual inspection of the processed images shows that severe thresholding of raw data tends to eliminate large portions of the POC while not removing the most intense cilia, while ilastik processing at p=0.5 preserved POC structure while exhibiting appreciable visual reductions of ciliary labeling, consistent with the small area reduction observed in Fig 2. Based on these metrics, we elected to apply a 0.5 probability cutoff to select a set of points representative of true signal for each channel, as 0.5 was intermediate between and roughly equivalent with the bounds of 0.25 to 0.75 in terms of reducing small area labels while preserving larger ones. This probability-based threshold enabled confident selection of points by relying on statistical significance as opposed to intensity thresholds (which can exclude real signal in fainter images).

**Figure 3:**
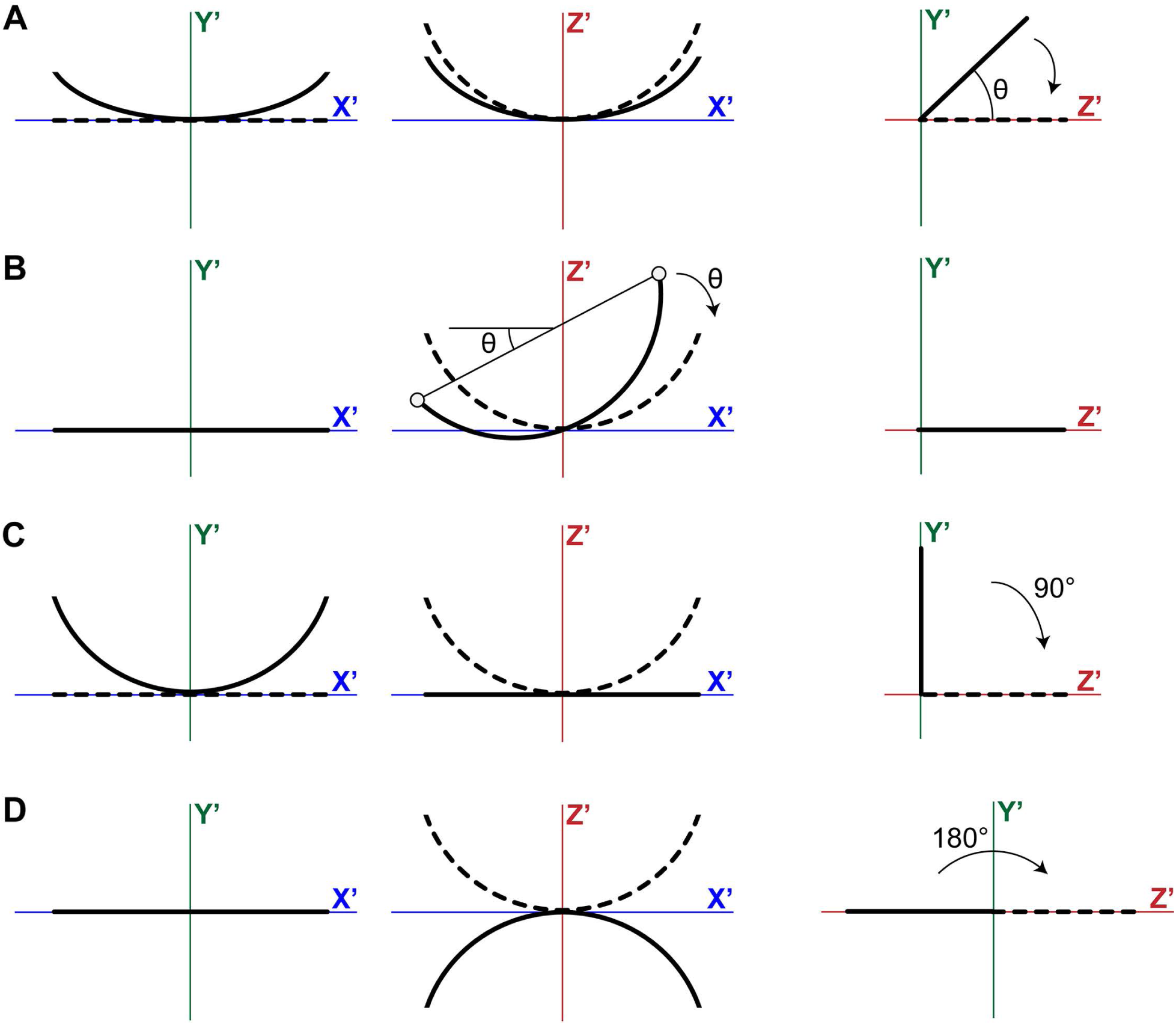
Correction of alignment errors. After PCA, images can be misaligned in four different ways. Subfigures A-D illustrate the relevant corrections. A) Rotation around the *X*′-axis. B) Rotation around the *Y*′-axis. C) Incorrect assignment of the *Y*′- and *Z*′-axis. D) Inverted orientation around the *Z*′-axis.

**Figure 4:**
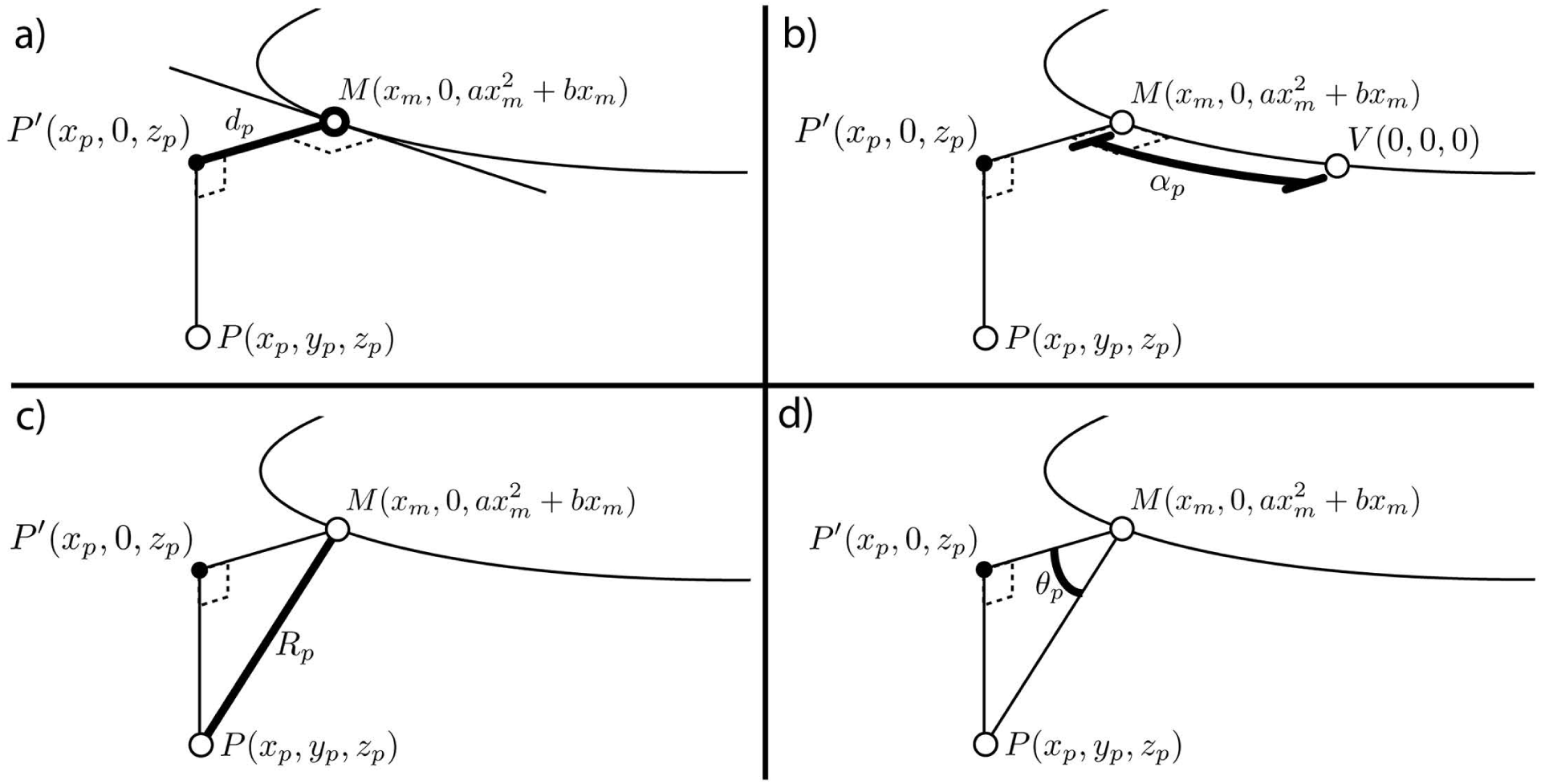
Calculation of cylindrical coordinates. Given a data point *P* (*xp, yp, zp*) and a model of the commissure 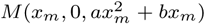, we calculated a cylindrical coordinate system that defined the position of *P* relative to *M*. A) First, we must identify the *xm* value which defines the point *M* (*xm*) that was closest to *P*. Since *M* lies in the *X*′*Z*′ plane where all *y* = 0, we can consider *P* as it lies in the *X*′*Z*′-plane, *P*′(*xp*, 0, *zp*). The distance between *M* and *P*′ (*d*_*p*_) was calculated by finding the Euclidean distance between the two points. We then used an optimization function to find the value of *x*_*m*_ that minimizes *d*_*p*_. B) Given *M* (*x*_*m*_) that minimizes *d*_*p*_, the arclength distance between *M* and *V* (0, 0, 0) at the midline (*α*_*p*_) was found by calculating the integral of *M* between *M* (*x*_*m*_) and *V*. C) Given *M* (*x*_*m*_) that minimizes *d*_*p*_, the Euclidean distance (*R*_*p*_) between *M* and *P* could be calculated in 3 dimensions. D) Finally, given *d*_*p*_ and *R*_*p*_, the angle of *P* (*θ*_*p*_) could be calculated according to trigonometric rules.

**Figure 5:**
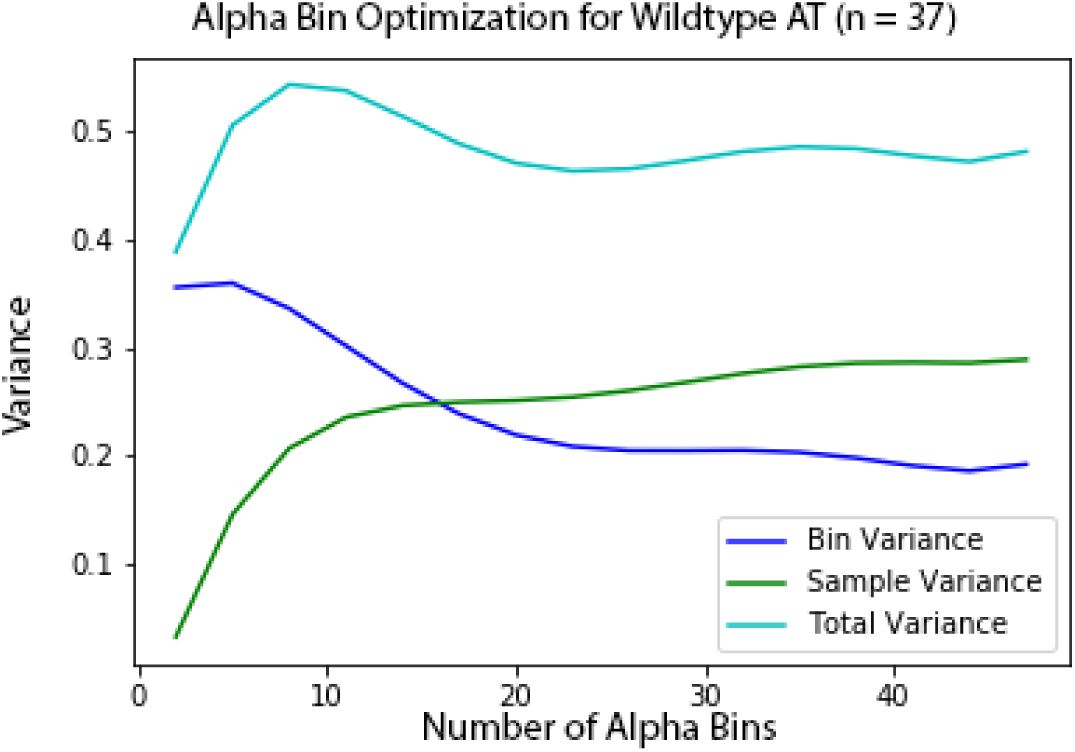
Alpha bin size calculation for landmark generation. Within a sample, the smaller the distance between any two points, the more likely that those two points would be similar and exhibit lower variance. Calculations of variance of R between adjacent bins using differing bin sizes between 2 bins per sample and 50 bins per sample for all samples were calculated and averaged (dark blue). Between samples of the same type, the larger the area queried, the more likely they were to be similar and exhibit lower variance. R distance of each bin in each sample was calculated and variance between sample bins for bin sizes 2 to 50 were graphed (green). These two metrics were then optimized to minimize these two sources of variance (light blue).

**Figure 6:**
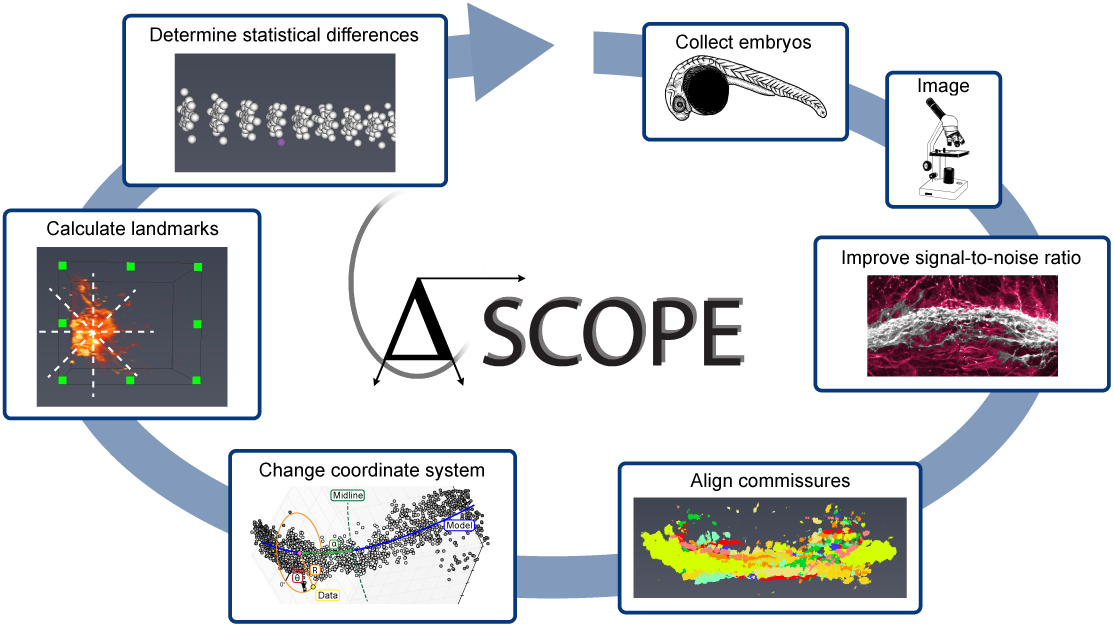
ΔSCOPE workflow for biological structural analysis and quantification. In a clockwise manner, ΔSCOPE processing involves: 1) collecting and immunostaining samples, 2) generating confocal stacks of immunostained samples with close to isotropic voxels, 3) processing confocal stacks with ilastik, 4) performing principal component analysis on the resultant data, 5) changing the coordinate system of the data to be biologically appropriate, 6) calculating bin sizes and then binning signal into landmarks, and 7) performing statistical tests.

**Figure 7:**
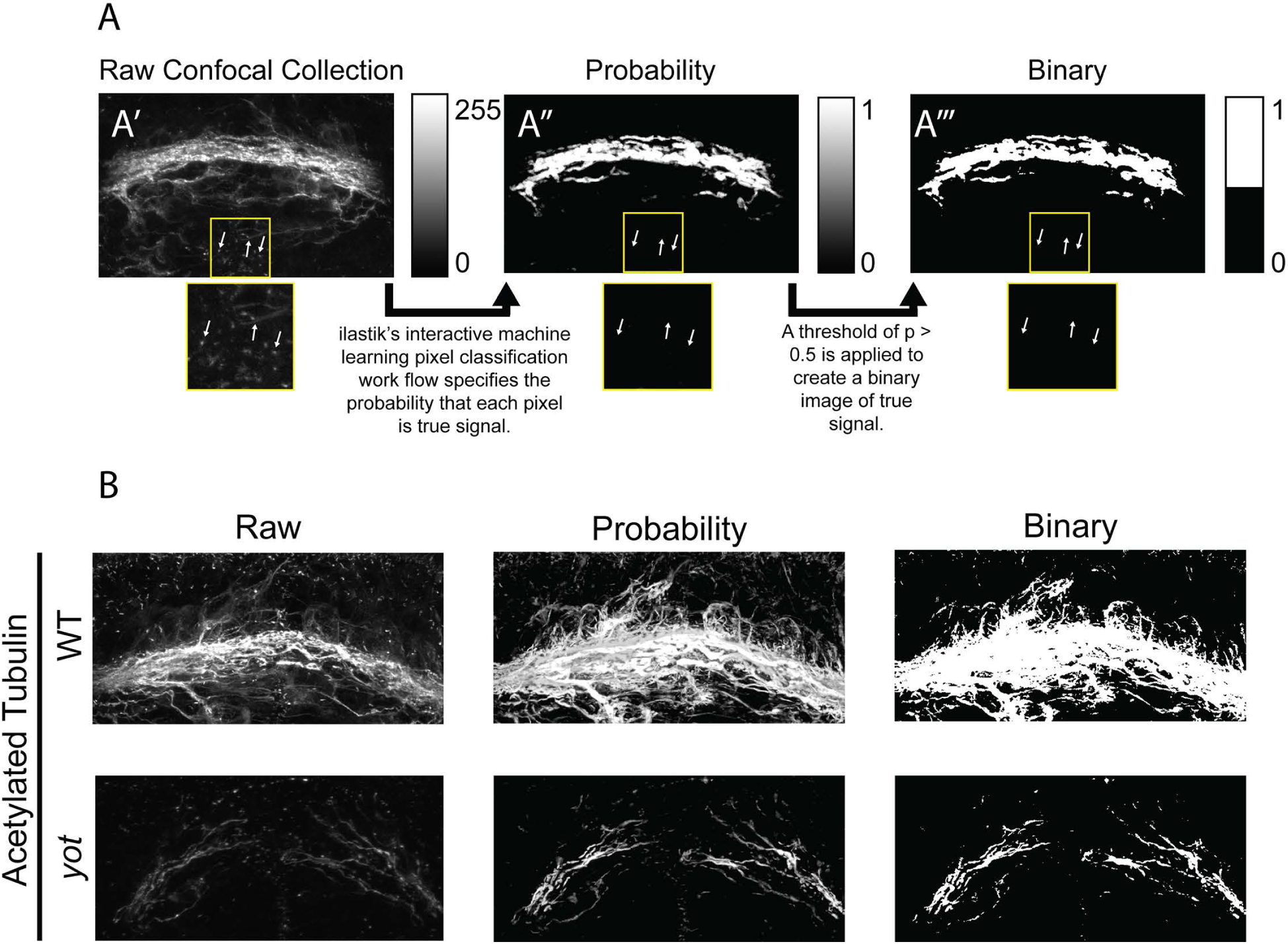
ilastik workflow to reduce noise from 2D and 3D images. (A) ilastik’s machine learning pixel classification work flow enables the selection of a binary set of points that represent true signal after eliminating noise and variable intensity. (B) ilastik processes images to reduce contamination of secondary signal from cilia while retaining axonal labeling.

Each probability file was read in Python by H5Py and saved as a NumPy 4D array ([`zyxc`]) with the fourth dimension containing two channels: signal and background [37, 38]. In order to distinguish between channels, we assumed that the channel that contains more points with a probability of greater than 0.1 would represent the true signal channel. Each point was then saved to a Pandas data frame as a row with *x, y*, and *z* values obtained from the point’s position in the array [39]. At this time, each point’s *xyz* position was scaled to account for the size of the voxel collected by the microscope, typically 0.16 × 0.16 × 0.21*µm*. Finally, a threshold was applied to the data frame to select only points with a probability of less than 0.5. This final set of points served as the representative data of the POC structure. Wild type AT samples generally contained less than half a million points, while the corresponding Gfap sample ranged from 300,000 to less than 100,000 points (Fig 13).

**Figure 8:**
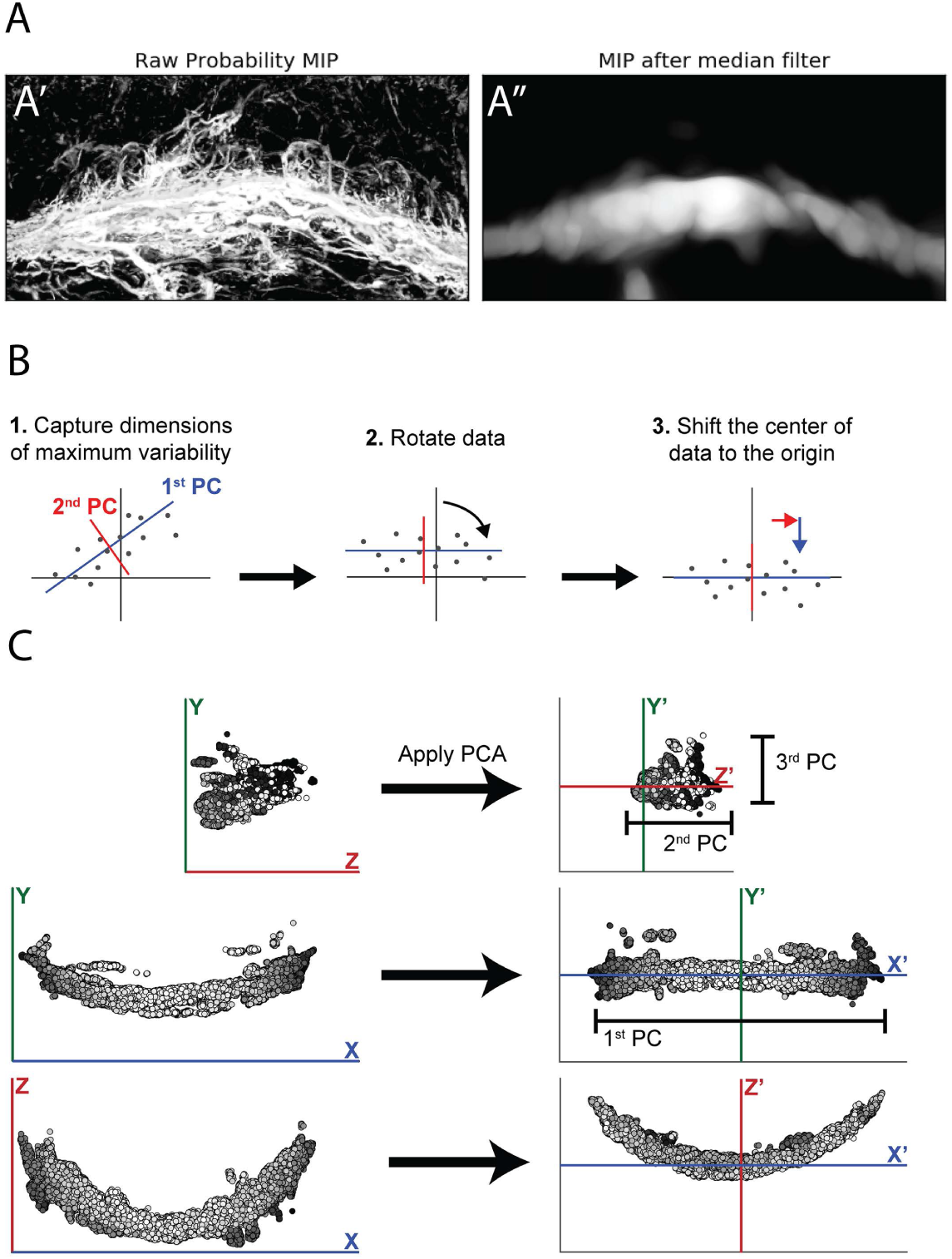
Median filter processing and principal component analysis for sample alignment. (A)Axons of the POC tend to wander and deviate from the central core of the commissure. While axons preserved in the probability MIP on the left were biologically relevant, they could interfere with the alignment process. On the right, duplicate application of a median filter to the axon data extracted the core structure of the commissure while eliminating fine processes that could interfere with alignment. (B) PCA was applied to n-dimensional data to identify axes that captured the most variability in the data. 1) In this two dimensional example, the first principal component (PC) was identified and the second PC is oriented perpendicular to the first. 2) The data was rotated so that the first PC was horizontal and the second PC was vertical. 3) Finally, the data was shifted so that the center of the data was at the origin. (C) After applying PCA to a single POC sample, the arc of the commissure lied in the new XZ plane with the midline positioned at the origin. In each 2D projection, the intensity of each point corresponded to depth in the third dimension.

**Figure 9:**
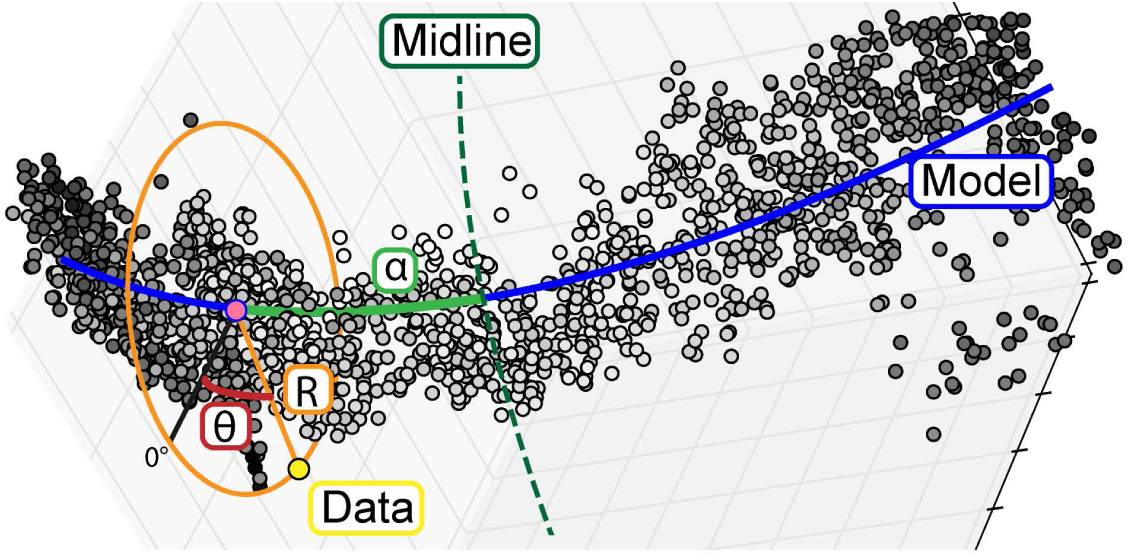
Biological cylindrical coordinate system for biological structure realignment and quantification. Creating a cylindrical coordinate system around the POC. To enable analysis of data points relative to a biological structure, points were transformed from a Cartesian coordinate system (x, y, z) into a cylindrical coordinate system (*α,θ*,R) defined relative to the structure of the commissure. This new coordinate system provided biologically relevant metrics of position.

**Figure 10:**
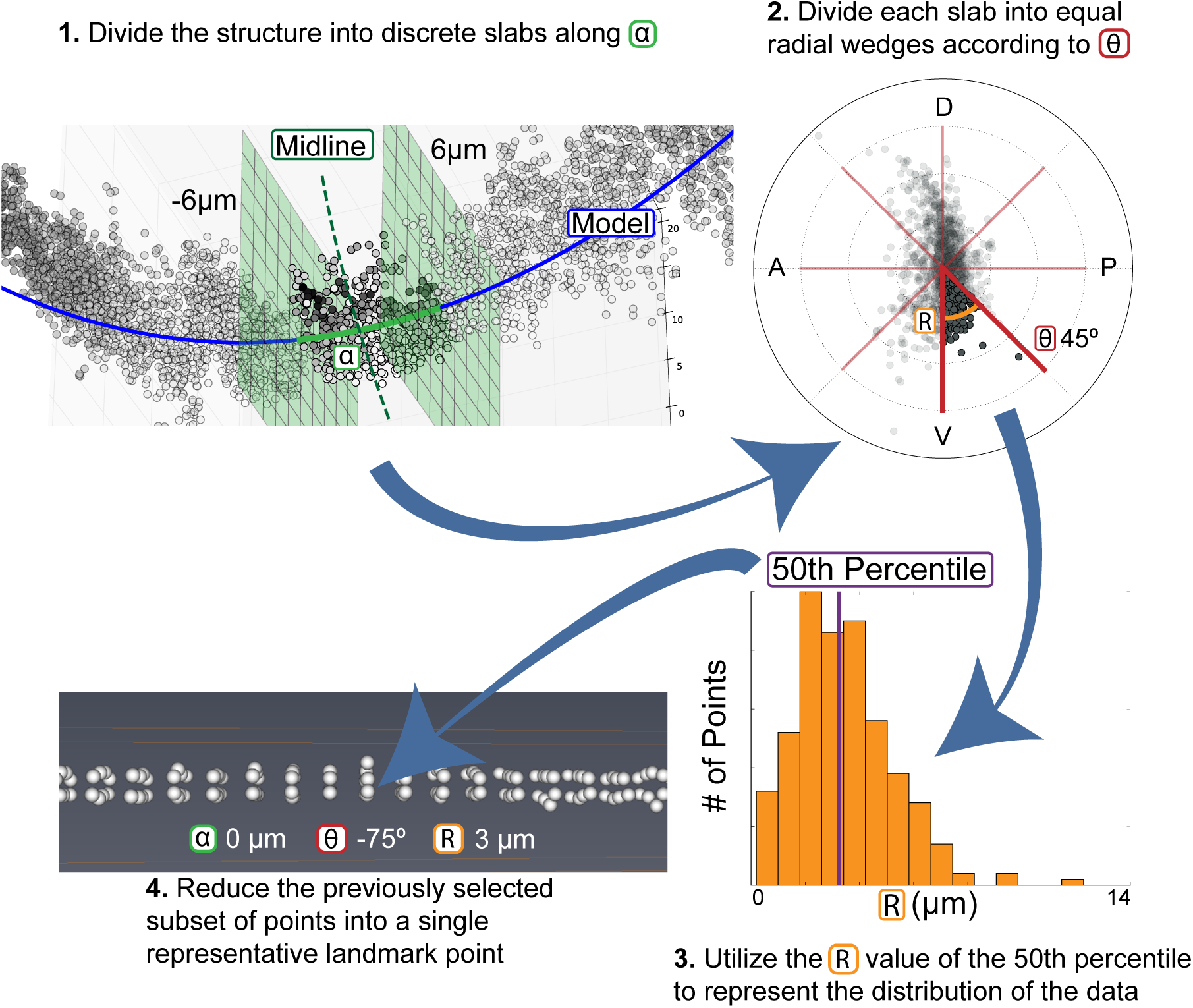
Calculation of biological landmarks. Calculation of biological landmarks. In order to facilitate direct comparisons between samples, we subdivide the data into a set of representative landmarks. 1) The POC was divided into equal slabs along *α*. 2) Each *α* slab was divided into eight *θ* wedges of 45°. 3) The set of points in each wedge consists of a set of R values that can be visualized in a histogram. In order to reduce a set of points to one representative point, we calculated the median R values in the wedge. 4) Each landmark point could be plotted and visualized according to the average *α* and *θ* values for the wedge and the median of the R values in the wedge

**Figure 11:**
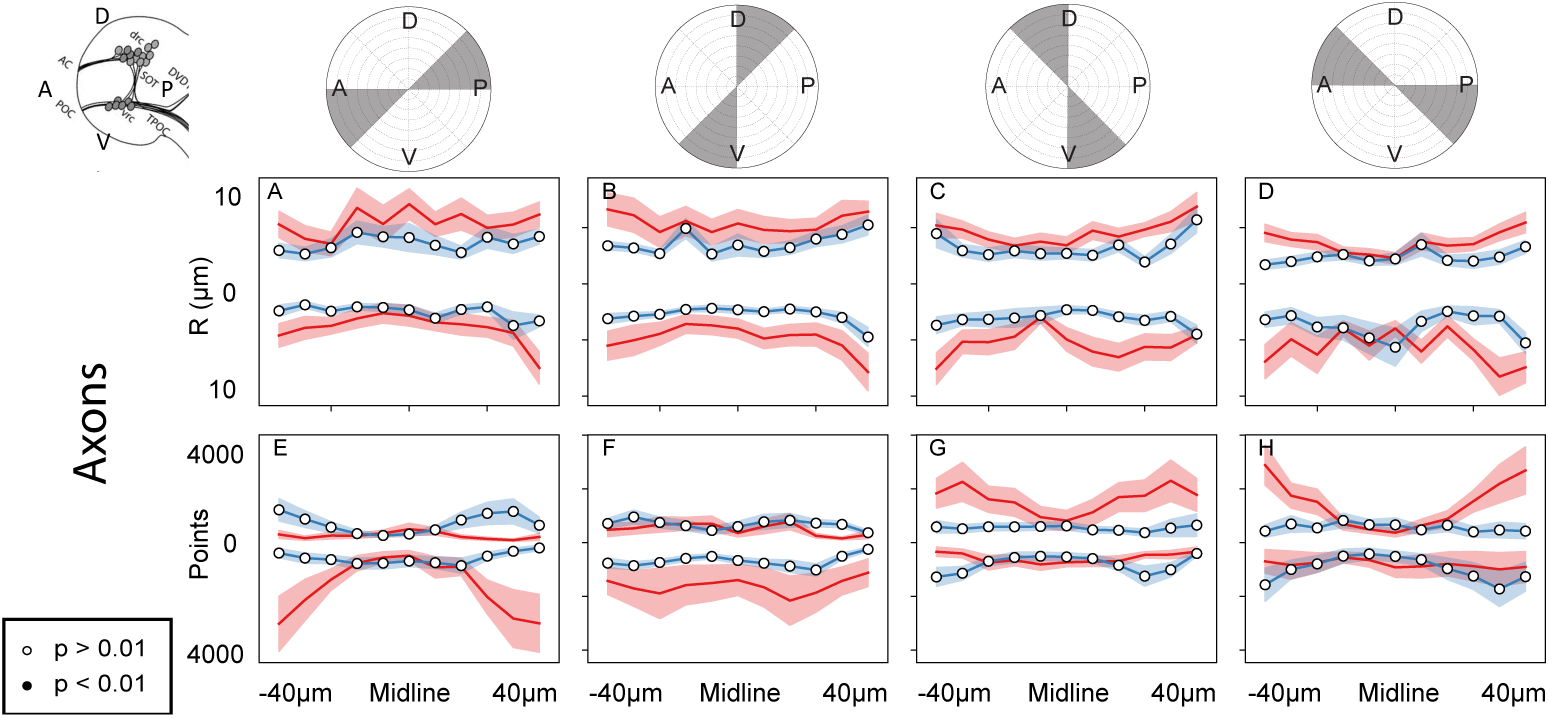
ΔSCOPE does not detect significant differences in similarly processed experimental groups. Wild type samples were collected and randomly assigned to either the control or out group to determine whether ΔSCOPE would detect spurious differences between sample sets. The top of the data plots correspond to the top half of the corresponding radial plot, and bottom to the bottom. Significant differences of p< 0.01 are denoted by black filled circles. A-D) No significant increases in the radial distance of axon signal were observed in any of the *α* or R positions E-H) No significant changes in AT signal were observed in the dorsal posterior axis at the midline though some non-significant variation was observed at the periphery. Despite this variation, ΔSCOPE did not consider this variation to be a significant deviation in signal amount.

**Figure 12:**
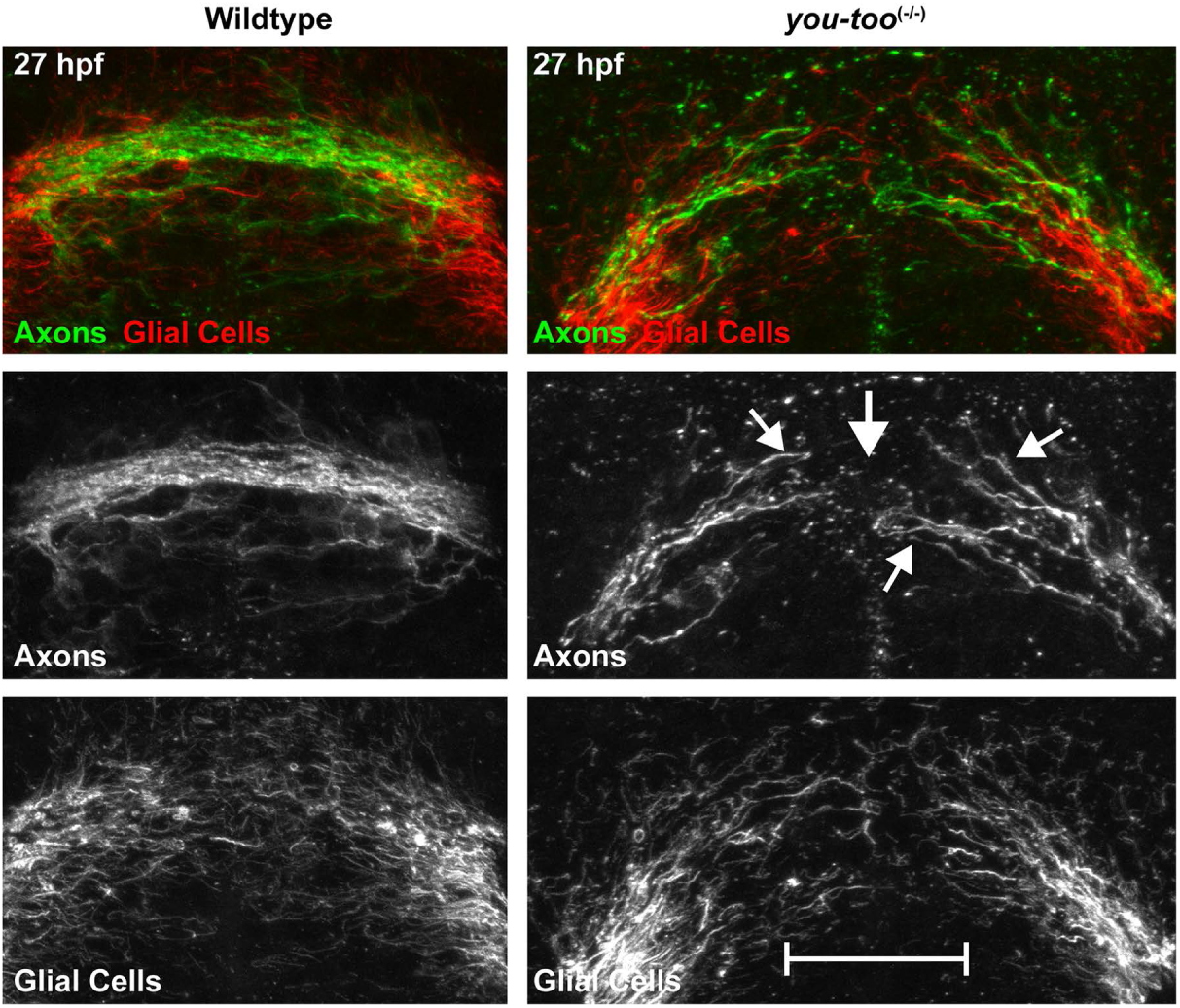
The *you-too* mutant POC exhibits a loss of commissure formation. The *you-too* mutant *(gli2-DR)* experiences a loss of commissure formation (AT) (Green) as compared to WT and disruption to the glial bridge (Gfap) (Red).

**Figure 13:**
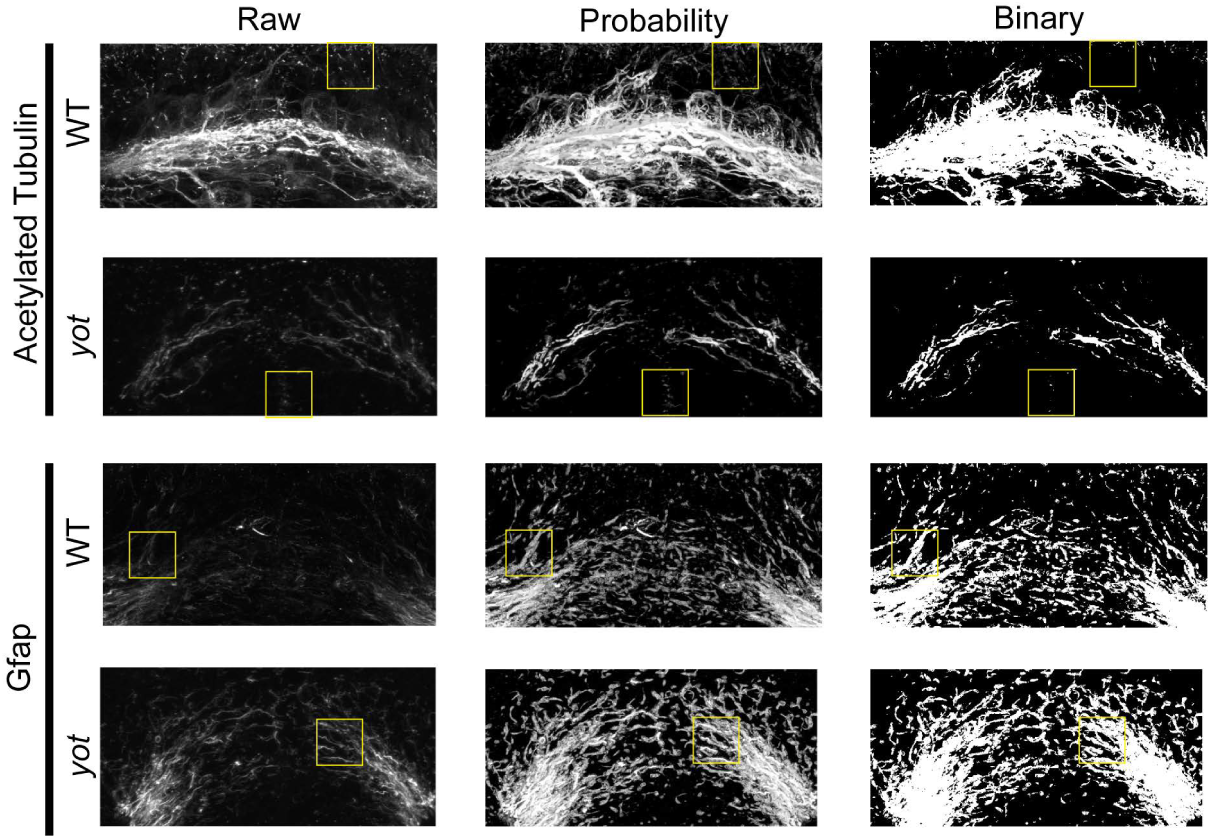
Representative ilastik results after processing wild type and *yot* anti-AT and anti-Gfap signal. Raw collections from the confocal microscope were processed by ilastik following training based on 5-10 representative samples. The resulting probability images contain the probability that each pixel was signal or background. Finally a threshold was applied to the probability image in order to extract a set of binary points. Following ilastik processing, faint axons were enhanced and preserved while points of ciliary labeling were eliminated. Faint Gfap signal was greatly enhanced in both wild type and mutant samples.

### Sample alignment

The position of each point in the ilastik-generated binary dataset of true signal was scaled by the dimensions (um) of the voxel. In order to prevent fine structures, such as wandering axons, from interfering with the core morphology of the commissure (and primary structural channel), we applied a median filter twice to smooth the structure and remove fine processes (Fig 8 A). PCA identified the orthogonal set of axes in the dataset that captured the widest range of variability in the data (Fig 8 B); therefore, the median filtering we applied serves to smooth out outlier signal and convolve individual fascicles into a singular structure. Selection of median filter size was dependent on the thickness of the structural signal, and thus required adjustment to best suit the sample set.

Since biological structures frequently maintain consistent proportions, PCA can use the median filtered data to isolate three orthogonal axes that are consistent between samples (Fig 8 C). Importantly, we only used the median filtered data to align channels but did not use it for data analysis. After PCA, each image was visually inspected to ensure that the structure was appropriately fit to each axis. ΔSCOPE comes complete with a set of alignment tools to make informed corrections to the PCA alignment.

The relative X-axis of our raw microscopy images of the POC consistently spanned the medial-lateral dimension of the embryo and consequently contained more variability than the other dimensions, therefore PCA identified the original Cartesian X-axis as the first principal component, though small adjustments to this alignment were achieved by PCA to compensate for errors in collection or sample orientation (Fig 8 C). For samples in which most of the signal in the microscopy collected X-axis was lost however, errors PCA axis assignment were evident, which was overcome by manually assigning the image X-axis as the first PCA component. Additionally, the anatomical dorsal to ventral axis of the forebrain commissures at the embryonic stages examined were collected in the relative Z-axis, which typically had a greater range of values as compared to the anatomical anterior to posterior axis that was assigned to the Y-axis, however, this was not definitive, and did not affect PCA axis assignment.

In order to ensure that all samples were in the same position following the alignment process, we then fit a polynomial model to the data and identify a centerpoint for translation to the origin. We mathematically described the shape of the POC with a parabola, such that the vertex of the parabola signifies both the center of the data and the position of the origin. Following PCA alignment, the commissure lies entirely in the XZ plane with the midline of the commissure positioned at the origin (Fig 8 C). The same transformation and translation completed on the structural channel was then applied to the secondary channel.

Sample alignment was performed only on the fluorescent channels that describe the basic structure (structural channels). In our data, acetylated tubulin (AT) served to label the basic structure of the POC. Any subsequent fluorescent channels, such as Gfap, were aligned according to the transformation of the structural channel. An additional two pre-processing steps were conducted on the structural channel in order to ensure that sample alignment was not negatively impacted by sub-cellular structures or remaining noise. First, a new data frame was created as described above, yet with a more stringent threshold of 0.25 in order to select points with the highest probability of being true signal. Second, the Scikit-Image median filter (radius 20) was applied to the thresholded data twice in order to smooth out noise on the surface of the structure [40]. For wild type samples, principal component analysis (PCA) from Scikit-Learn was applied to the processed data using all three original dimensions (*X*′, *Y*′, and *Z*′) [41]. Following transformation of the structural channel, the first principal component was assigned to a new *X* axis, the second to *Z* and the third to *Y*. The same transformation matrix that was calculated for the structural channel was also applied to any secondary channels. In contrast, the components used to align *yot* mutants were different due to the nature of the severity of POC phenotypes in this mutant. To reduce error in alignment, the *X*′ axis was held constant while PCA was applied to the *Y*′ and *Z*′ axes, which were reassigned from the first and second principal components, respectively.

### Alignment Correction

After PCA alignment, some samples contained minor errors in orientation, which prevented direct comparison across multiple samples. The four error types are produced by rotation of the sample around each axis. Some samples experience rotation around the *X*′ axis, which means that the parabola of the commissure no longer lies exclusively in the *X*′*Z*′ plane (Fig 3 A). In order to correct this error, a line was fit to the data in the *Y*′*Z*′ plane and its slope (*m*) was used to calculate the necessary angle of rotation

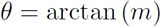

around the *X*′ axis.

In order to correct rotation around the *Y*′ axis, we identified two points in the dataset that marked the extremes of the commissure (Fig 3 B). The first endpoint was assigned based on the maximum or minimum *z* value in the sample. In order to determine if the maximum or minimum should be used, the concavity of the commissure was calculated based on a best fit parabola *z* = *ax*^2^ + *bx* + *c*. A concave up commissure (*a* > 0) uses the maximum *z* value, while a concave down commissure (*a* < 0) uses the minimum *z* value. After the *z* value of the endpoint was identified, its corresponding *x* value could be found in the dataset. To identify the second anchor point, we calculated the distance between the first anchor point and the minimum or maximum *x* value in in the commissure. The *x* value that maximizes the distance to the first anchor point was set as the second anchor point and its corresponding *z* value was found in the dataset. Given two anchor points (*x*_1_, *z*_1_) and (*x*_2_, *z*_2_), the slope of the line between the two points was used to calculate an angle of rotation

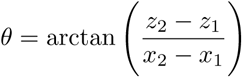

around the *Y*′ axis that will position both anchor points at equal *Z*′ values.

In some samples, the principal components were incorrectly assigned such that the *Y*′ and *Z*′ axes were exchanged (Fig 3 C). This error was corrected by applying a 90° rotation around the *X*′ axis. Finally, the last error occurs when the commissure was flipped upside down in the *X*′*Z*′ plane (Fig 3 D). This error was corrected by rotating the data by 180° around the *X*′ axis.

### Sample centering

PCA alignment results in a consistent image orientation between samples, however it failed to account for the position of the POC in 3D-space. In order to center each sample accurately at a common origin, a polynomial model was used to represent the underlying shape of the data and to identify a consistent center point. After alignment, the POC lied in the *X*′*Z*′ plane and formed a parabolic structure. We fit a quadratic ordinary least squares regression model,

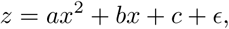

to the structural channel by minimizing the squared error, where *ϵ* ∼ *N* (0, *σ*_*ϵ*_) and *σ*_*ϵ*_ is a fixed constant [38]. The *x*-coordinate (in the *X*′*Z*′ plane) of the parabola’s vertex is 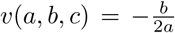. Letting 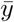 denote the average value of *y*, we translated the coordinates of the data such that the point 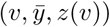 was moved to the origin. The necessary translation was calculated based on the structural channel and applied to all secondary channels.

### Cylindrical coordinates

Calculating a parabola as a representative model of the POC provided the foundation of a cylindrical coordinate system. We converted the *xyz* coordinates of each point into a cylindrical coordinate system defined by *α, θ* and *R*. For any point *P* (*x*_*p*_, *y*_*p*_, *z*_*p*_), we identified its projection in the *X*′*Z*′-plane *P*′(*x*_*p*_, 0, *z*_*p*_), and a point *M*′(*x*_*m*_, 0, *z*(*x*_*m*_)), where the Euclidean distance between *P*′ and *M*′,

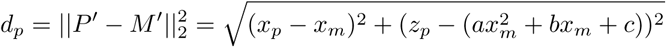

was minimized (Fig 4 A).

Each point *P* (*x*_*p*_, *y*_*p*_, *z*_*p*_) was considered in the XZ plane *P*′(*x*_*p*_, 0, *z*_*p*_) and a secondary point 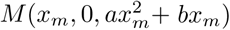 was identified on the parabolic model that minimizes the euclidean distance between the point and the model

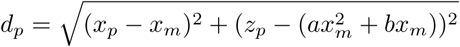

(Fig 4 A). In order to identify the value of *x*_*m*_, the optimization function minimize from SciPy was used to minimize *d*_*p*_ as a function of *x*_*m*_ [42]. *In other words, given a point P* (*x, y, z*), find a value *x*_*m*_ such that the distance between the projection of *P* to the *X*′*Z*′-plane and the point *M* (*x*_*m*_, 0, *z*(*x*_*m*_)) was minimized. *α*_*p*_ was defined by calculating the distance along the model between *M* and the vertex *V* (0, 0, 0),

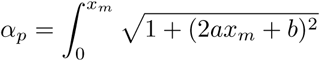

(Fig 4 B). Given the minimal Euclidean distance between *P* and *M* along the curve, *R* was then calculated as the Euclidean distance between *P* and *M* in 3D space,

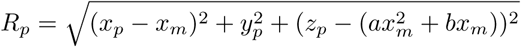

(Fig 4 C). *θ*_*p*_ was finally calculated as follows:

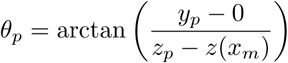

(Fig 4 D).

After transforming the data to cylindrical coordinates, the dataset was saved to a PSI file according to PSI Format 1.0. Each point was assigned an ID number and the following values were saved: *x, y, z, α, R, θ*. The Cartesian coordinates correspond to the position of the data following sample alignment.

### Landmark calculation

In order to perform statistical comparisons between samples, we reduced the sample data to a set of representative landmark points. The POC was divided into eight wedges around *θ*, each spanning 45 degrees, and into *n*_*α*_ slices along *α*. In particular, *θ* bins were calculated relative to the plane of the parabola and assigned individually the same way to each sample, so that the data for all samples was similarly oriented and binned in the *θ* dimension. This however requires that during acquisition of subsequent data processing, that the data be transformed so that the anterior-posterior and dorsal-ventral dimensions were roughly aligned in 3D space, prior to PCA, so that PCA may a perform fine tuning of alignment. For each sample 1 ≤ *s* ≤ *n*, the number of points in each wedge (after calculation of *α*) for each individual sample was calculated. The 50^*th*^ percentile (median) of the *R* values for all data points within a sample *αθ* wedge, for all such wedges within a sample, and across all samples, was then determined.

Let *w*_*ij*_ be a wedge, for integers 1 ≤ *i* ≤ *n*_*α*_ and 1 ≤ *j* ≤ 8. Then *n*_*s*_(*w*_*ij*_) was the number of points in the *ij*^*th*^ wedge in sample *s*, and 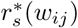 was the median value of *R* among all points in wedge *w*_*ij*_ in sample *s*.

In order to identify the size of an *α* bin that maximized the amount of data gleaned from the analysis while minimizing sample noise, we calculated two measures of variance. First, the variance was calculated for each bin and its adjacent neighbors. This type of variance between bins decreased as the number of *α* bins increased. In order to counter this trend, we calculated the variance between samples for each bin, which increased as the number of *α* bins increased (Fig 5). By selecting the number of *α* bins that minimized both types of variance, we were able to identify the appropriate number of bins for the sample. This number was dependent on the type of signal under examination, the number of samples being tested, the resolution of the data, and the success of the alignment steps.

The number of bins *n*_*α*_ was optimized by minimizing both the variance in median *R* values across samples (i.e., global variance) and across adjacent bins (i.e., local variance).

For a given wedge *w*_*ij*_ in a particular sample *s*, we define the local variance in median radius across the neighboring wedges as:

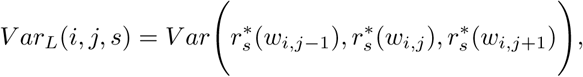

where the *j*’s were taken modulo 8. The average local variance for a particular choice of *n*_*α*_ was thus,

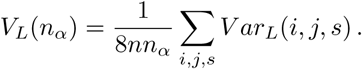

At the same time, we defined the global variance in median radius across samples for a particular wedge *w*_*ij*_ to be:

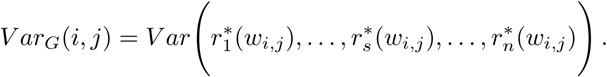

The average global variance for a particular choice of *n*_*α*_ was thus,

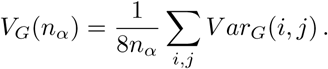

The optimal value for *n*_*α*_ was identified as the value that minimizes *V*_*L*_(*n*_*α*_) and *V*_*G*_(*n*_*α*_) (Fig 5)

We observed that the optimal number of *α* bins, located at the local minimum of the optimization curve of was slightly above 21, depending on experimental group (24 for *you-too* experiments, 21 for glial bridge experiment). We then rounded to the next highest odd number to capture the midline.

The calculation of biological landmarks was done on a sample by sample basis, with the midline landmark assigned the vertex of the parabola, which was assumed to be the midline of the commissure. The ability of subsequent statistical tests to detect variation from the mean depended on the appropriate assignment of the midline, and hence, samples which cannot be reliably assigned this center point were removed from testing, as they would poorly influence the results. The micron distances between the central landmark and more lateral landmarks were calculated by dividing the largest alpha value in the landmark calculation data set by one less than half the alpha bins. All other data from all other samples in the data set were then binned by this alpha micron distance. Rather than scaling the data by a relative alpha bin number, so that all data from all samples was evenly distributed across all alpha bins we elected to use absolute micron distances. As not all commissures were imaged to the same maximal z-depth, and thus the furthest extent of the commissure was not always captured, using relative alpha distances would tend to introduce data distortion artifacts. Moreover, relative alpha bin values would only benefit our calculations if significant differences in brain or commissure size were 1) evident and common and 2) were likely to result from or be experimentally related to a treatment under consideration. While implementation of relative alpha bin assignment was possible, the overall size of the brain has been observed to be neither evident nor common and did not appear to vary in lateral extent significantly during the period of development (22 hpf-30 hpf) or in response to Slit perturbation. We anticipated that small variances in size which were expected to occur in equal proportion in all experimental sample sets, may increase the variance of the data sets, but that this variance was likely to be limited to more lateral landmarks. In contrast, scaling of all embryos was determined to be more likely to result in the creation of artifacts, especially since not all samples were equally easy to image to their full extent, and scaling could result in the artificial detection of these differences as differences in signal distribution. We further limited our calculations and examinations to the middle third of landmarks in *you-too* analysis to further reduce the impact size variance might have on landmark evaluation, to focus our investigation on midline commissure formation and glial bridge development, and to limit the number of statistical tests and conclusions drawn from sample sets.

In order to identify landmarks that exhibited statistically significant differences in median radius experimental data sets, we tested whether the means of similar bins between experimental groups had the same mean using the non-parametric Kruskal-Wallis H-test analysis of variance, using the stats.kruskal function offered by within the python SciPy module, which does not assume equal variances or normality around the group mean. While this test may lack statistical power for experimental groups which are normally distributed, it was determined that implementation of a non-parametric test which did not make assumptions about the data distribution would be more extensible. The results of the set of statistical differences between sample sets were then evaluated by the multipletest module of the python statsmodels module, using a two stage Benjamini-Hochberg correction at an *α* of 0.01 for *you-too* and *slit1a* analysis and at 0.05 for glial bridge comparisons where fewer samples were able to be collected due to the difficulty of mounting. Bins were then evaluated using the adjusted p values provided by the multitest module, and were then considered statistically significant at the same *α* value. In this way, the likelihood of a false positive bin appearing was kept to 1-5% probability, and as we have evaluated only structural features with multiple adjacent statistically different bins, the likelihood of false positive features should significantly less than 1%.

A table of key resources used in generation of microscopy data and ΔSCOPE program, including repositories of the data and code base generated in the development of ΔSCOPE is provided as a supplemental table (KRT table).

## Results

### Development of ΔSCOPE – analysis of biological structures

Quantitative image analysis of commissure formation in the vertebrate brain has been impeded by four major challenges that include image noise, experimental variation due to biological and sample based misalignment, loss of sample dimensionality, and a lack of statistical power when comparing between images. We developed ΔSCOPE, a new 3D image analysis program designed to overcome these challenges. The premise of ΔSCOPE centers on the registration of the data around a common biological structure. We focused our analysis on the structural components of the POC during embryonic forebrain development in zebrafish.

The POC is structurally composed of bundled axons derived in part from neurons of the ventral rostral cluster in the diencephalon [43, 44, 20, 21]. POC axons were visualized with immunocytochemical procedures using antibodies that recognize Acetylated Tubulin (AT), which establishes the primary structural channel for our analysis. Secondary structural markers, such as anti-Gfap antibodies used to label astroglial cells, were registered and processed relative to the primary structural channel (the POC in our case). ΔSCOPE was designed to 1) take in confocal image stacks of the POC and secondary structures, 2) output quantitative comparisons between sample types of both primary and secondary structures, and 3) quantify and identify statistically significant biological differences and distributions between the primary and secondary structures between sample sets (Fig 6).

### Image pre-processing

Before ΔSCOPE analysis could be performed, image pre-processing was necessary to isolate true structural signal from experimental and biological noise. A further complication in the isolation of the POC as the primary structural signal, was the additional labeling of tubulin found within cilia of cells that densely line the ventricle walls of the brain [45]. To segment the labeling of axons from cilia, we first determined that simple thresholding of signal intensity would not be feasible. Although thresholding dramatically reduced small area objects, like cilia, it also reduced large objects like portions of axons in the POC (Fig 2; data not shown). Therefore, we next tested an existing interactive machine learning program called ilastik to identify an isolate axonal labeling in our images. In brief, ilastik relies on user input to create a training dataset consisting of images labeled for signal and background, and outputs an image composed of probability values for each pixel being true intensity value, as opposed to background noise [14]. We hypothesized that ilastik, which features user guided adaptive image processing features, would reduce the number of data points while improving the resolution of the interrogated structures better than simple thresholding of the data (see ilastik section in Methods).

We trained ilastik on 28 hpf wild type samples (see ilastik section in Methods for training procedure), and evaluated the ability of ilastik to reduce the count of small area objects while preserving large scale structures. Ilastik image processing is based upon probabilities instead of intensities, and we determined that p values between p = 0.25 and 0.75 were equally effective at reducing small objects, while preserving objects larger than individual axons at 20 pixels. The midpoint probability of p=0.5 was chosen for all ilastik image pre-processing, which preserved the qualitative structure of the POC while significantly reducing ciliary labeling (Fig 7).

### Principal component analysis aligns samples on biological axes

For any comparison of biological samples to be possible, the structures being analyzed must be consistently oriented in 3D space. The uniform alignment of samples enables direct measurements of structural differences between samples. However, the position and shape of biological structures, like the POC, exhibit variability due to natural variation in biological samples and inconsistencies in experimental preparations and microscopic image acquisition. These varied inconsistencies can cause differences in how an individual sample is oriented in 3D space, which presents a problem for accurate sample to sample alignment and any possibility for robust statistical comparisons between experiments. To overcome these problems, mathematical transformations of the image data to fit a wild type reference brain have been used in past studies, however this approach also captures normal variation in a structure between individuals as significant differences [16]. To surmount this limitation, we chose an alignment process that centered on the analysis of a single discrete structure and its associated components, which enabled analysis on a sample-by-sample basis before computing averages of a sample set. This enabled detection of significant shifts in signal distribution and intensity across all imaged axes in a sample set without distorting the data to fit an arbitrary reference sample.

We applied principal component analysis (PCA) to isolate consistent and unbiased sets of biologically meaningful axes from anisotropic 3D samples (where each dimension is proportionally different in size). Our use of PCA in ΔSCOPE relied on a primary structural channel to calculate the PCA transformation matrix, which in our study was the axon labeled channel (anti-acetylated tubulin). Any secondary channels such as astroglia (anti-Gfap) were similarly transformed according to the matrix calculated for the structural channel. A median filter was applied to smooth the ilastik pre-processed data, which served to eliminate outlier signal and convolve the structure of the POC. PCA was then applied to this data to identify 3 new Cartesian axes across all sample sets. A center point focused at the POC midline and a parabola were fit to the data using a best fit parabolic curve to define the model of each individual POC (Please see PCA section in Methods for more details).

### Cylindrical coordinates define signal position

Following PCA alignment, each point was still described by x, y, and z Cartesian coordinates; however, these coordinates were not directly related to the structure itself and thus limited in their applicability to the analysis of the structure in question. In order to facilitate direct comparison of specific structures in 3-dimensional space, we implemented a cylindrical coordinate system that was defined relative to the biological components of the structure itself. Due to the parabolic structure of the POC relative to the image stack itself, conversion of the image stack to a polar or cylindrical coordinate system by choosing an axis and origin point is inherently faulty, because wrapping the data around that point failed to capture the structure or relationship of the POC with its environment at all points other than where the POC and origin coincidentally overlap. Thus, capturing the structure of the POC required inventing a coordinate system that was inherently dependent on the fluctuating position of the POC in the image stack and then using that position to calculate the positions of signal relative to the POC itself. In this way, our use of an adaptable cylindrical coordinate system reduced the detection of biological variation while making it more sensitive to changes in the actual structure of the POC.

To calculate these new coordinates required an assumption that the primary structure being analyzed had a stereotypical shape that was consistent between samples, and in our case, could be represented with a parabola fit to anti-AT labeling, though other models could be used. We defined a set of relative cylindrical coordinates oriented around a single POC parabolic axis in 3D space. Each point in the data was assigned three new values that described its position relative to the parabola as the central axis: distance from the model (R), angular position around the model (*θ*), and distance from the midline along the model (*α*) (Fig 9). These new coordinates served as parameters that contained biologically meaningful information about the shape and composition of the structure. *α* described the position of points relative to the midline or periphery of the commissure. Depending on the embryonic stage being analyzed, *θ* captured the dorsal-ventral or anterior-posterior position of the point. Finally, R described how far a point was from the commissure, which was informative to the degree of axon fasciculation observed in the commissure or the distribution of secondary channel signal around the commissure (glia in this study). By re-defining how the data was organized, ΔSCOPE was able to convert all image data to a 3D cylindrical coordinate point cloud system. We observed that anti-AT signal (S1 Movie) and anti-Gfap signal (S2 Movie) were consistent between the raw data and the 3D system, and also observed that the relationship between AT labeling (Green) and Gfap labeling (Red) (S3 Movie) was preserved without warping or manipulating the data.

### Building Landmarks to compare samples

The conversion of our image data into cylindrical coordinates enabled each point to encode biological information relevant to the development of the POC. We next binned the data by establishing regularly spaced landmarks – a method of analysis commonly used in morphological studies, which permits comparisons and statistical tests to be performed between similarly positioned bins [46]. Although historically landmarks were often assigned by an expert with domain knowledge of the structure, we took an alternative approach that eliminated human decision making by mathematically calculating a set of regularly distributed landmarks across the structure. To do this, we divided the commissure into a set of bins in the *α* and *θ* axes (Fig 6), within which two representative metrics were calculated to describe the nature of the signal: 1) the median R distance of the signal and 2) the total number of points of signal. We divided the *θ* dimension into eight evenly spaced bins, each of *π*/4 radians in order to enable visualization of signal around the entire commissure without oversampling. The number of bins along the *α* dimension was empirically determined (Fig 5) to minimize sample-to-sample and bin-to-bin variance. These landmarks were then computed for each independent experiment (See Landmark section in Methods for details on calculation of bin sizes and statistical methods). Upon successful conversion of raw data (e.g. S1 Movie-S3 Movie) into landmark representations of the data, we observed that landmarks were tightly clustered where AT signal was most abundant and more dispersed at the periphery where AT signal was less fasciculated (S1 Movie vs S4 Movie). This trend also held true with Gfap signal, where we also observed greater apparent R distances, consistent with Gfap data (S2 Movie vs S5 Movie). This observation is even more apparent when comparing landmarks of AT and Gfap data (S6 Movie), for which the greater R distance of Gfap over AT was clearly different. This data suggested that both the cylindrical coordinate conversion and subsequent landmark assignment was working as intended.

### Interpreting ΔSCOPE landmark results

ΔSCOPE was designed to leverage the use of biological replicates, which serves to reduce overall noise while enabling detection of changes in signal morphology. The position and density of the post-optic commissure was defined by the pathfinding fidelity and quantity of axons spanning the diencephalon. Our interpretations of ΔSCOPE analysis of POC development were based on two key assumptions. 1) The more axons present within a set landmark wedge would represent a higher positive pixel count at that landmark. 2) A highly fasciculated commissure would be represented by lower radial distance values at the midline, which consequently would also show more pixels found nearest to the modeled POC parabola. Using these metrics, we evaluated the utility of ΔSCOPE as a new methodology for image quantification, as applied to the comparative analysis of post-optic commissural development over time and for the assessment of both gross and subtle phenotypic differences.

### Validation of ΔSCOPE processing

In order for ΔSCOPE to serve as a reliable computational method to detect differences between experimental data sets, it was necessary to first validate that sample-to-sample variation within identically collected sample sets would not detect false significant differences. To this end, we collected wild type data sets (n = 32), and randomly assigned 16 images to the control group and 16 images to the out group. These two groups were then compared to one another using ΔSCOPE to evaluate whether false positive detections arose in the workflow. As predicted, no significant differences were detected between these two groups, demonstrating that ΔSCOPE was not overly sensitive to minor variation between groups (11). Although, multiple metric and intra-group comparisons were possible, we determined they were not appropriate analyses due to the presence of non-biologically relevant variation like differential embryo depth, size and collected orientation, and the inability to conduct 1D statistical tests on such metrics. Instead, ΔSCOPE affords the user the ability to select an R percentile, as well as limiting the sampling of pixel number to a given R distance, based on knowledge of the data. After confirming ΔSCOPE was not overly sensitive, we next sought to determine whether ΔSCOPE could detect significant differences in commissure formation in mutants and transgenic fish lines that have previously been shown to exhibit POC defects.

### Validation of ΔSCOPE through analysis of known mutant phenotypes

We approached the validation of ΔSCOPE by analyzing several different degrees of commissure manipulation from a severe loss of midline axon crossing to more subtle errors in axon and astroglial cell positioning. We first tested the accuracy of ΔSCOPE to quantitatively describe the severe POC and glial bridge defects previously reported in the *you-too (gli2-DR; yot*) mutant [15]. Homozygous *yot* mutants express a dominant repressive form of the Gli2 transcription factor, which functions to repress the Hedgehog signaling pathway in those cells normally expressing the *gli2* gene [47]. We have previously shown that loss of Hedgehog signaling in the *yot* mutant causes the misexpression of the Slit family of guidance molecules that play essential roles in directing POC axons across the diencephalic midline of the zebrafish forebrain. More specifically, *yot* mutants exhibit expanded expression of *slit2* and *slit3* throughout midline regions where they were normally absent. In contrast, *slit1a* was found to be reduced in the forebrain of *yot* mutants compared to controls [15]. Slit2 and Slit3 are accepted to function as axon guidance repellents, thus their expanded expression was hypothesized to be responsible for the reduced midline crossing seen in *yot* mutants [15]. The function of Slit1a is currently unresolved. The consensus on the qualitative severity of POC loss in the *yot* mutant provided a valuable reference phenotype to validate the utility of ΔSCOPE to accurately describe this known POC defect. In addition, we sought to extend our understanding of commissure formation in *yot* mutants by analyzing glial cell position in combination with POC axons, and together this analysis presents the first quantitative assessment of *yot* commissural phenotypes.

Prior to our comparative assessment of POC formation in *yot* mutants and wild type siblings, we had to tailor our ΔSCOPE pre-processing for the type of labeling exhibited in *yot* mutants. Due to the severe lack of POC axons in *yot* mutants, the secondary labeling of cilia and deeper axons recognized by the anti-AT antibody posed a potential source of artificially signal contamination of the signal representing the structural axon channel (Fig 12). Therefore, we first trained the machine learning pre-processing of ilastik on *yot* mutants to reduce the influence of the ciliary tubulin or deeper axon labeling.

We show that applying ilastik’s pixel classification algorithm as part of the ΔSCOPE methodology effectively reduced ciliary labeling and significantly reduced contributions from signal derived from deeper positioned non-commissural axons (Fig 13). Visual observation of MIP’s from pre- and post-ilastik processed wild type and *you-too* embryos revealed clear reductions in non-POC signal. With this successful pre-processing and the isolated true POC axonal signal we next commenced with ΔSCOPE’s analysis of PCA-based sample alignments and landmark calculations.

To conduct PCA alignment we first pre-processed both wild type and *yot* mutant samples with a median filter of 20 pixels. Although this filtering level was sufficient for PCA alignment of wild type samples, it resulted in a loss of most of the data at the midline of *yot* mutant samples. This reduced labeling was due to the known reduction of midline crossing of axons in *yot* mutants. ΔSCOPE requires a minimum amount of signal for the PCA alignment and proper parabola modeling; therefore, we empirically determined that a median filter of 10 pixels was sufficient to reduce noise while still enabling alignment of the sparse axonal commissures seen in the *yot* mutant.

### ΔSCOPE analysis of POC axons in you-too mutants

Landmarks were next assigned to the aligned samples of wild type and *yot* mutant POC axons, which are graphically represented by two opposing wedges of landmarks along the modeled parabola (Fig 14). To quantify whether POC axons were significantly mis-positioned in *yot* mutants, we analyzed the defasciculation metric (the median radial distance (R)) of axons from the modeled POC parabola. Our analysis revealed significant increases in the radial distance of signal in homozygous *yot* mutants at the midline and along the whole tract of the commissure as compared to homozygous wild type siblings (Fig 14). More specifically, we observed that signal lies between 3-5 um from the calculated model in all radial axes of wild type commissures (Fig 14), light green line, top panel). In contrast, anti-AT signal in *yot* mutants ranged in distance from 5 to 15 um from the modeled POC parabola, with the greatest deviations observed at the midline (Fig 14), dark green line, top panel). Furthermore, the greatest magnitude of difference was observed along the ventral anterior (Fig 14 A) and dorsal anterior axes (Fig 14 C,D). The average distance of anti-AT signal in *yot* mutants from the midline was 15 um (*yot*, R = 15 um) as compared to the tighter fasciculated wild type commissure (wt, R = 3 um). Little divergence was observed along the dimensions of the posterior axes (dorsal-posterior (Fig 14 A,B):WT R = 2 um; *yot* R = 8 um; ventral-posterior (Fig 14 C,D); WT R = 2 um; *yot* R = 14 um). Biologically, these data suggest that commissural axons in *yot* mutants are distributed more dorso-ventrally (towards the pre-optic area or yolk sac respectively) and that wandering axons in *yot* are preferentially located more superficially near the pial surface of the forebrain as opposed to pathfinding deeper into the forebrain.

**Figure 14:**
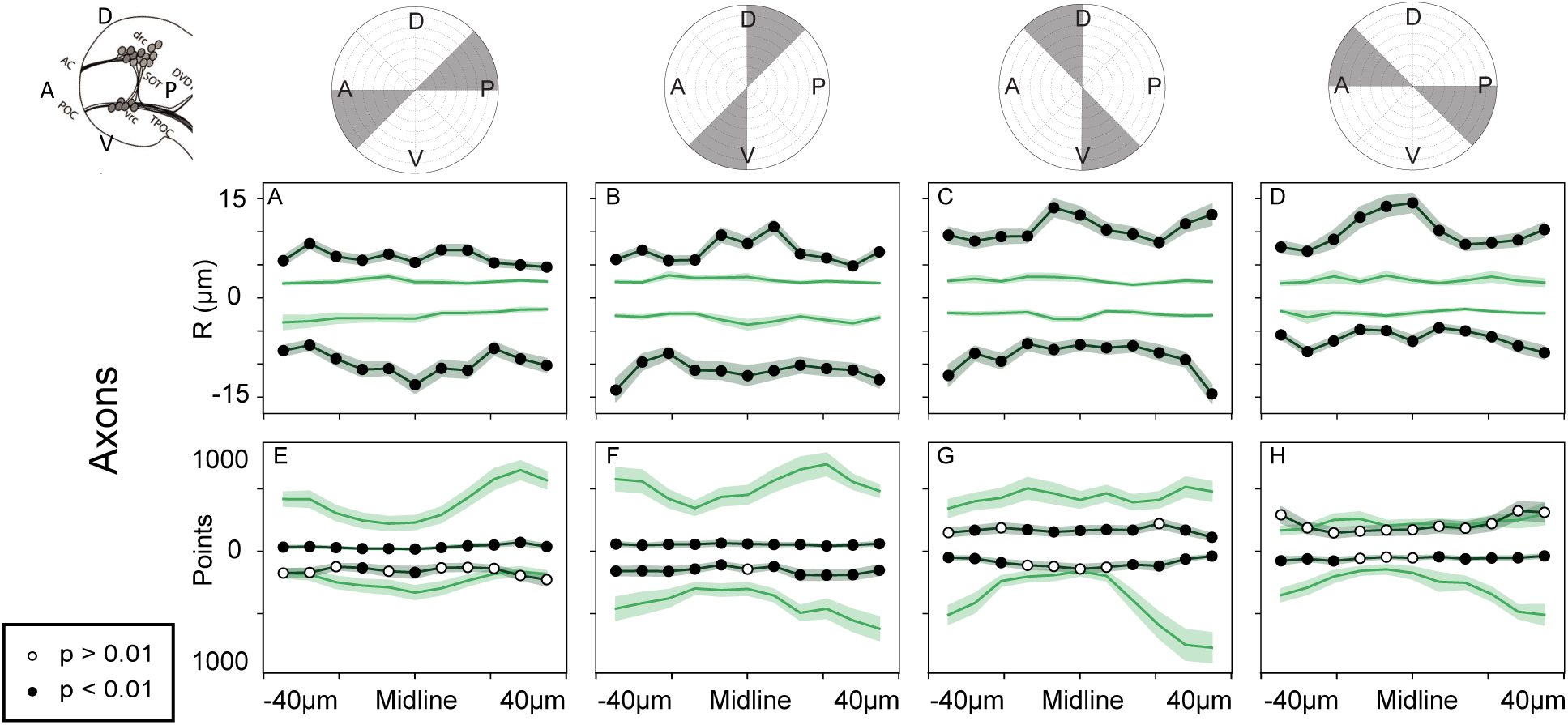
Landmark analysis comparing the structure and distribution of wild type and *yot* POCs. Analysis of changes in amount of axon signal (A-D) and distance of axon signal from POC model (E-H) in WT (n=37) and *yot* (n=33) 28 hpf embryos. Top of data plots correspond to the top half of the corresponding radial plot, and bottom to the bottom. Significant differences p< 0.01 are denoted by black filled circles. A-D) Significant increases in the radial distance of axon signal were observed in all *α* and R positions in *yot* embryos. We observed an increase in average R distance of 3-5 um in wild type embryos to 5-15 um in *yot* embryos. E-G) Significant reductions in AT signal were observed in the dorsal posterior axis at the midline. F-H) Significant reductions in the ventral-posterior axis were observed in *yot* embryos.

To confirm that fewer POC axons were crossing the midline in *yot* mutants as previously described in Barresi et al. 2005 [15], we used ΔSCOPE to quantify the number of anti-AT pixels at and around the midline. We observed significant reductions in positive pixel number at the midline in *yot* mutants, which were most pronounced along the dorsal and dorsal-posterior axes (Fig 14 E-G). ΔSCOPE also detected significant but intermittent reductions in the number of anti-AT pixels (axons) in the ventral and ventral-posterior axes along the commissures’ periphery (Fig 14 F-H). Reductions in the amount of signal in the periphery of *yot* mutants could be due to fewer axons projecting towards the midline (ispilateral side) and/or fewer axons extending away from the midline following crossing (pathfinding on the contralateral side).

### ΔSCOPE analysis of the glial bridge in you-too mutants

Qualitative characterization of the diencephalic glial bridge has suggested that loss of *gli2* leads to a spreading out of these cells along the dorsal/anterior to ventral/posterior dimensions of the forebrain [15]. However, the fibrous and compositionally amorphous pattern of anti-Gfap labeling of astroglial cells in the zebrafish forebrain has made quantitative characterization of their positioning difficult and inadequate to date. Continuing to use the modeled POC parabola as a structural anchor for sample comparisons, we applied ΔSCOPE to the analysis of the secondary anti-Gfap channel to uncover whether the more obscure phenotypes in the glial bridge could be detected and quantified. In contrast to our qualitatively deduced expansion of the glial bridge in *yot-/-*, we found that ΔSCOPE detected sporadic Gfap signal closer to the POC model in *yot* mutant embryos as compared to wild type embryos. These reductions in the distance of Gfap signal from the POC model were found in both the ventral and dorsal posterior axes (Fig 15 A-D). Concomitantly, we also observed reductions in the amount of Gfap signal detected in *yot* mutants along these same posterior axes, with the most significant reductions found in the commissure periphery (Fig 15 E-G). Intriguingly, these same posterior axes correlate with the locations which also lacked significant anti-AT (axon) signal in both WT and *yot* mutant commissures (Fig 14). These data taken together suggest that in *yot* mutants, Gfap+ astroglial cells or their cell processes are reduced in the same locations where commissural axons are normally found in wild type embryos, namely the dorsal posterior axes (compare (Fig 14 A,B with (Fig 15 A,B) and in the periphery of the ventral-posterior axes (compare (Fig 14 G,H with (Fig 15 G,H). No significant reductions were observed in Gfap signal along the ventral/anterior axes. We note that both the amount and positioning of Gfap signal was not significantly different in anterior or anterior ventral positions (Fig 17 A, B, D-F, H). When also considering the significant dorsal/posterior reductions of signal underneath the commissure, we interpret these data to suggest that the only remaining midline signal lies primarily in the ventral/anterior axis. We propose that this results in the apparent redistribution of the remainder of the Gfap signal appearing in the ventral axis, consistent with previous reports [15].

**Figure 15:**
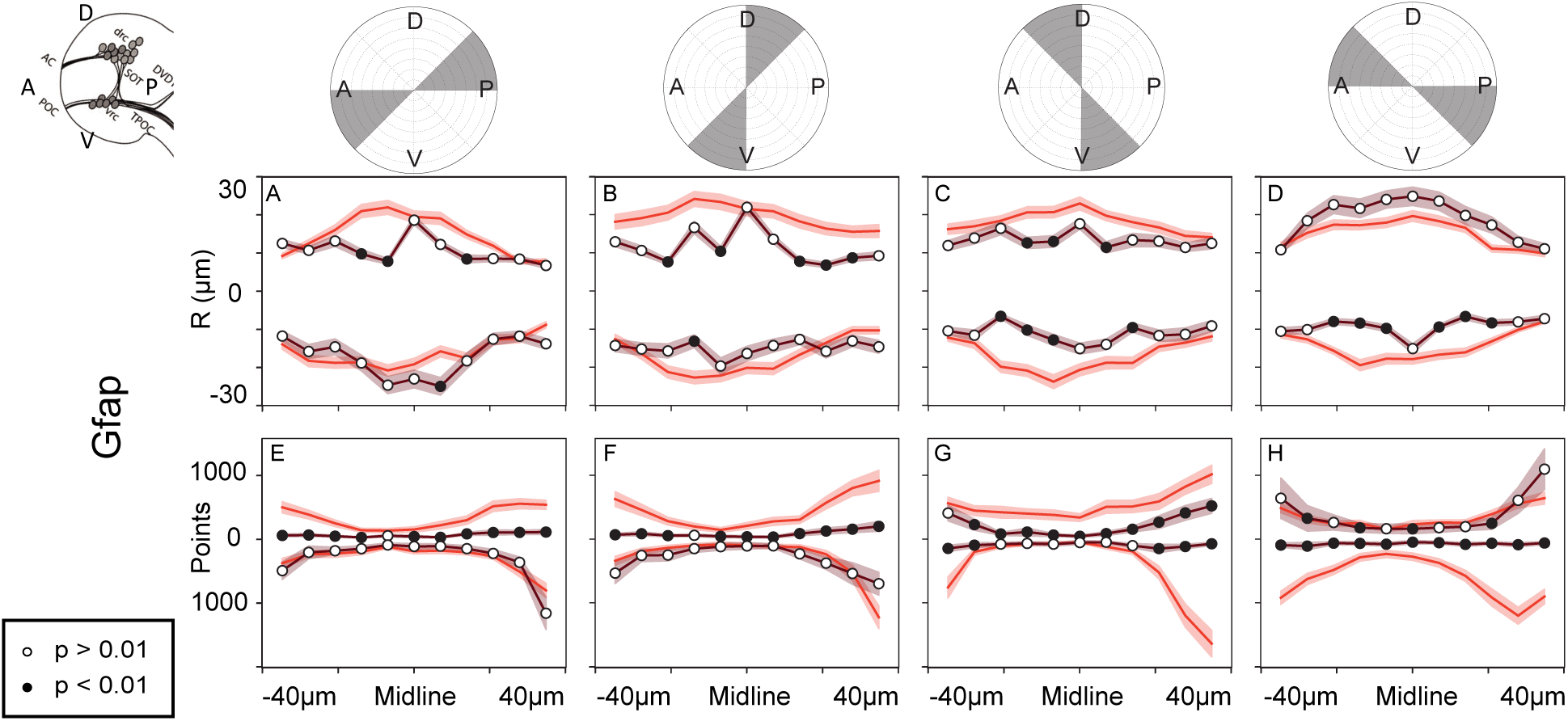
Landmark analysis comparing the structure and distribution of Gfap signal in wild type and *yot*. Analysis of changes in the amount of Gfap signal (A-D) and distance of the Gfap signal from the POC model (E-H) in WT (n=37) and *yot* (n=33) 28 hpf embryos. The top of the data plots correspond to the top half of the corresponding radial plot, and the bottom to the bottom. Significant differences p< 0.01 are denoted by black filled circles. A-D) Significant reductions in the radial distance of Gfap signal were observed in midline dorsal posterior *α* and R positions bins in *yot* embryos. We observed a reduction in average R distance from 10-25 um in wild type embryos to 10-20 um in *yot* embryos. E-H) Significant reductions in Gfap signal were observed in the dorsal posterior axis at the midline in *yot* embryos.

**Figure 16:**
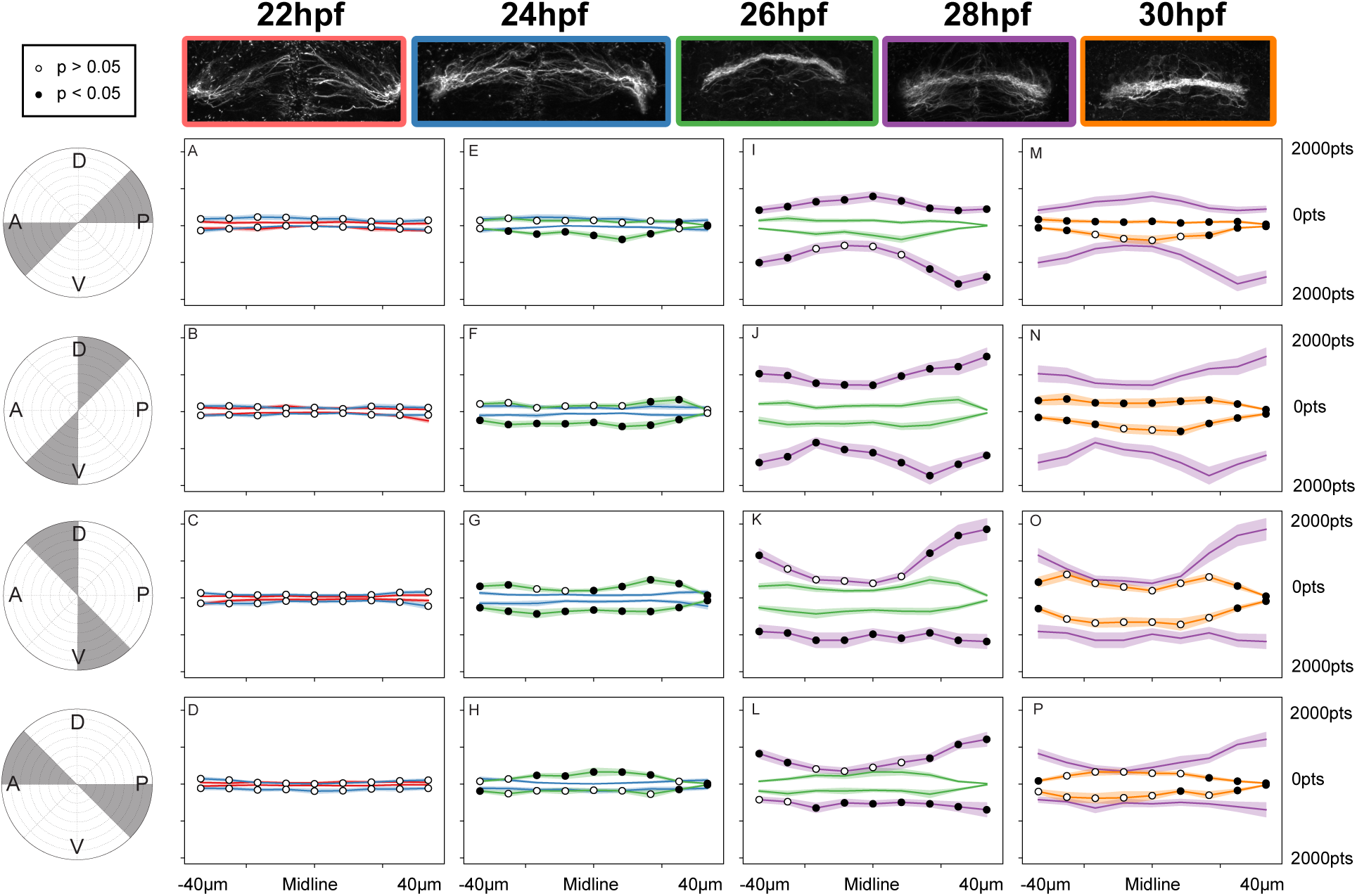
Landmark point analysis of POC development and anti-AT signal. A-D) Statistical comparison of the number of positive pixels in 22 hpf (n=13) and 24 hpf (n=20) POCs. No statistical differences were observed. E-H) Statistical comparison of axon positive pixels between 24 hpf (n=20) and 26 hpf (n=18). Significant changes observed in anterior ventral positions. I-L) Statistical comparison of axon positive pixels between 26 hpf (n=18) to 28 hpf (n=17) POCs. Increases in AT signal observed in all but dorsal-anterior bins. M-P) Statistical comparison of axon positive pixels between 28 hpf (n=17) and 30 hpf (n=17). Significant changes observed in dorsal posterior bins.

**Figure 17:**
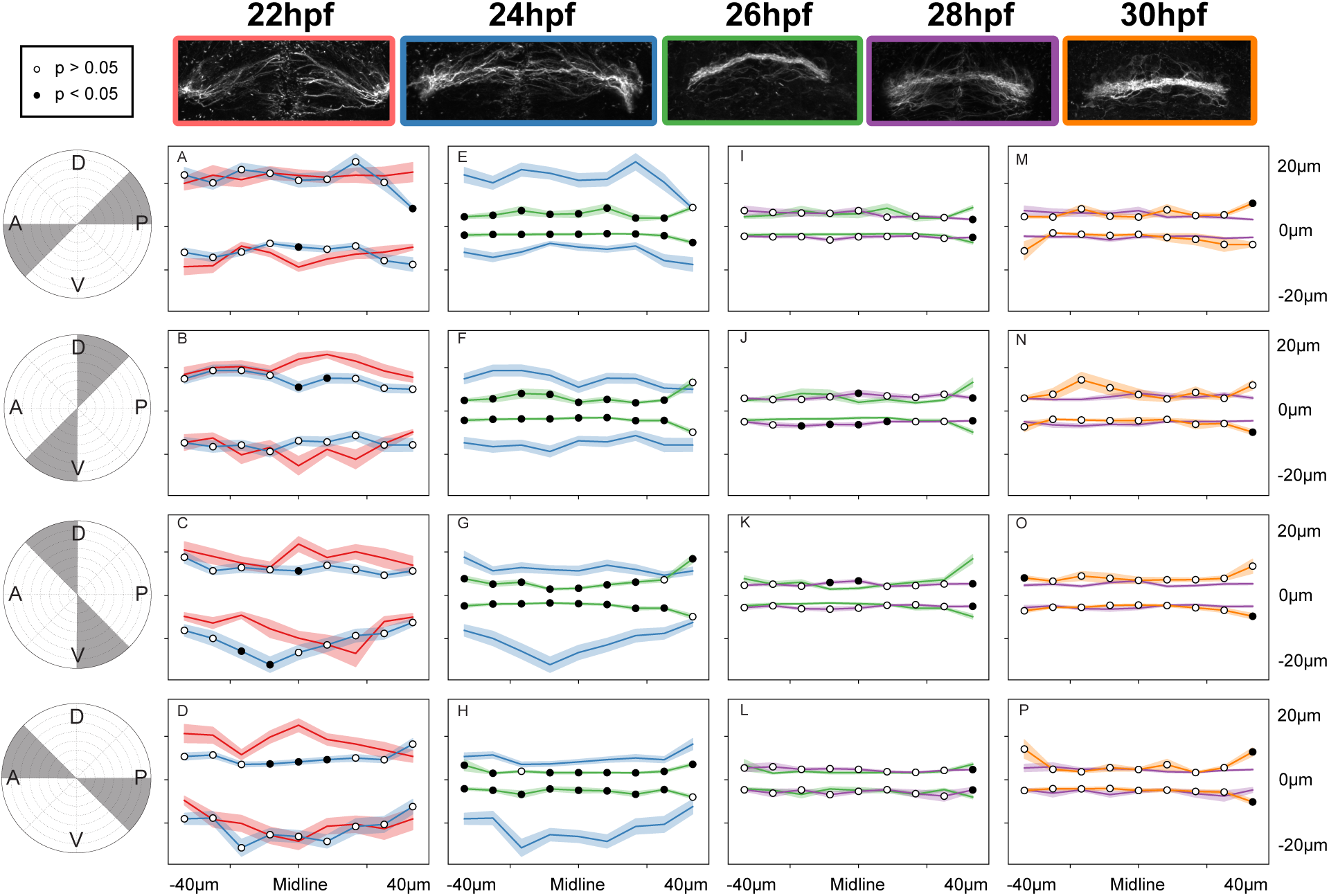
Landmark median radial distance analysis of POC development and fasciculation. A-D) Statistical comparison of the radial distance of axon signal from the POC model in 22 hpf (n=13) and 24 hpf (n=20) POCs. Statistical reduction in radial distance was observed in dorsal-anterior positions. E-H) Statistical comparison of the radial distance of axon signal from the POC model in 24 hpf (n=20) and 26 hpf (n=18). Significant reductions in radial distance was observed in all positions. I-L) Statistical comparison of the radial distance of axon signal from the POC model in 26 hpf (n=18) to 28 hpf (n=17) POCs. No statistical significance observed. M-P) Statistical comparison of the radial distance of axon signal from the POC model in 28 hpf (n=17) and 30 hpf (n=17). No significant changes observed.

### Quantification of the development of the POC

Demonstration of ΔSCOPE to successfully quantify both axon and glial phenotypes in the *yot* mutant led us to next ask whether ΔSCOPE can also quantitatively describe how axon-glial interactions change over the course of commissure development. However, the earliest embryonic time point when sufficient commissural axon signal (anti-AT) in zebrafish for ΔSCOPE structural alignment to work is at 22 hpf. Thus we can only start to use ΔSCOPE to study POC formation just after its POC initiation.

Using antibodies that recognize AT and Gfap, we analyzed POC and glial bridge development specifically between 22 and 30 hpf, which captured both the pioneering midline crossing events as well as the fasciculation and thickening of the POC. For each time point, we quantified the number of positive pixels and their median radius from the modeled POC parabola. Between 22 and 30 hpf, we observed significant increases in the number of positive pixels of anti-AT signal at the midline, which was indicative of progressive axon fasciculation over time (Fig 16). Interestingly, between 24 and 26 hpf we observed a large increase in anti-AT signal occurred between (Fig 16 E-P), which was also paired with significant reductions in the distance of signal from the modeled POC parabola in all radial bins (Fig 17 E-P). This comprehensive axonal condensation around the modeled POC parabola was preceded by an earlier statistically significant reduction in the radial distance of the anti-AT signal along the dorsal-anterior axis between 22 and 24 hpf (Fig 17 D). Importantly, during this same early time period (22-24 hpf) the amount of axon signal did not change; however, by 26 hpf, asymmetric increases in anti-AT signal were quantified along the anterior-posterior and ventral axes (Fig 16 E-H). We interpret the biological relevance of these data to suggest that the early reductions in median R distance with no corresponding change to the number of points represents axons undergoing a period of error correction as they pathfind across the midline. However, soon after this first period of midline crossing, the number of points increased as median R distance continued to decrease, suggesting a period of mounting commissural fasciculation, consistent with previous descriptions of POC development [15].

### Quantification of glial bridge development

Having quantitatively described the development of POC axons during commissure formation, we next sought to determine how the cells of the glial bridge may be changing relative to the assembly of the POC. Using the modeled POC parabola as the primary structural channel, we applied ΔSCOPE to the Gfap labeling of the secondary channel. This analysis detected significant reductions in Gfap radial distance in ventral-posterior bins at the beginning moments of commissure development (24hpf) (Fig 18 E-H). During this same time period, we also observed significant increases in Gfap signal within ventral-anterior bins (Fig 19 E-H). Although over the remaining time course, we observed a cyclical behavior in Gfap radial distance, which showed a change from the condensed configuration at 24hpf to a visibly and statistically significant expansion in Gfap position from the parabola at 28hpf that then returned to shorter radial positions at 30hpf. These movements were largely equivalent around the entire parabola. Interestingly, this cyclical pattern of radial moment correlated with different changes in Gfap signal (pts) that showed an inverse spatial relationship between the dorsal/anterior and ventral/posterior quadrants of Gfap signal (Fig 19 E-H;M-P vs Fig 18 E-H;M-P). More specifically, the moments of radial distance reduction during these cycles were correlated with significant increases in the amount of Gfap signal along the dorsal/anterior domains yet also paired with stable (24 - 26 hpf) to reduced (28 - 30 hpf) signal within the ventral/posterior domains. In contrast, during the period (26 - 28hpf) when radial distance positions increased for Gfap, the opposite but similarly inverse spatial changes in the amount of Gfap signal was quantified as compared to the previous and later periods (Fig 19 vs Fig 18). It is relevant to recall that these cycles in Gfap signal and radial distance occurred in the absence of any detected changes in axon positioning between 26 hpf and 30 hpf, which argues against these quantified changes in Gfap labeling as artifacts of poor positioning of the modeled parabola (Fig 17 I-P). When considering these data together we were able to detect a behavioral relationship between the amount and position of POC axons with astroglial cells over the course of commissure formation. POC axons quickly established a condensed commissure by 26 hpf, during which time glia showed a similar condensing movement (Fig 16 E-H vs Fig 19 E-H and Fig 17 E-H vs Fig 18 E-H). In addition, as the amount of POC axons (AT points) changed over time, there appeared to be a similar response in glial cell movement. For instance, the increased AT signal observed between 26 - 28 hpf was paired with a dramatic spreading of Gfap position, and yet as the AT signal dropped by 30 hpf glial cells were detected closer to the parabola (Fig 16 I-P vs Fig 19 I-P and Fig 17 I-P vs Fig 18 I-P). These data of glial bridge dynamics taken in consideration with POC axon behaviors largely supports remarkable correspondence in position and quantities between these two structural labels. These data demonstrate that the glial bridge is both present and similarly reactive to POC axons during development of the post-optic commissure, which provides the first quantitative analysis to suggest the glial bridge may provide contact-mediated structural guidance support to POC pathfinding axons.

**Figure 18:**
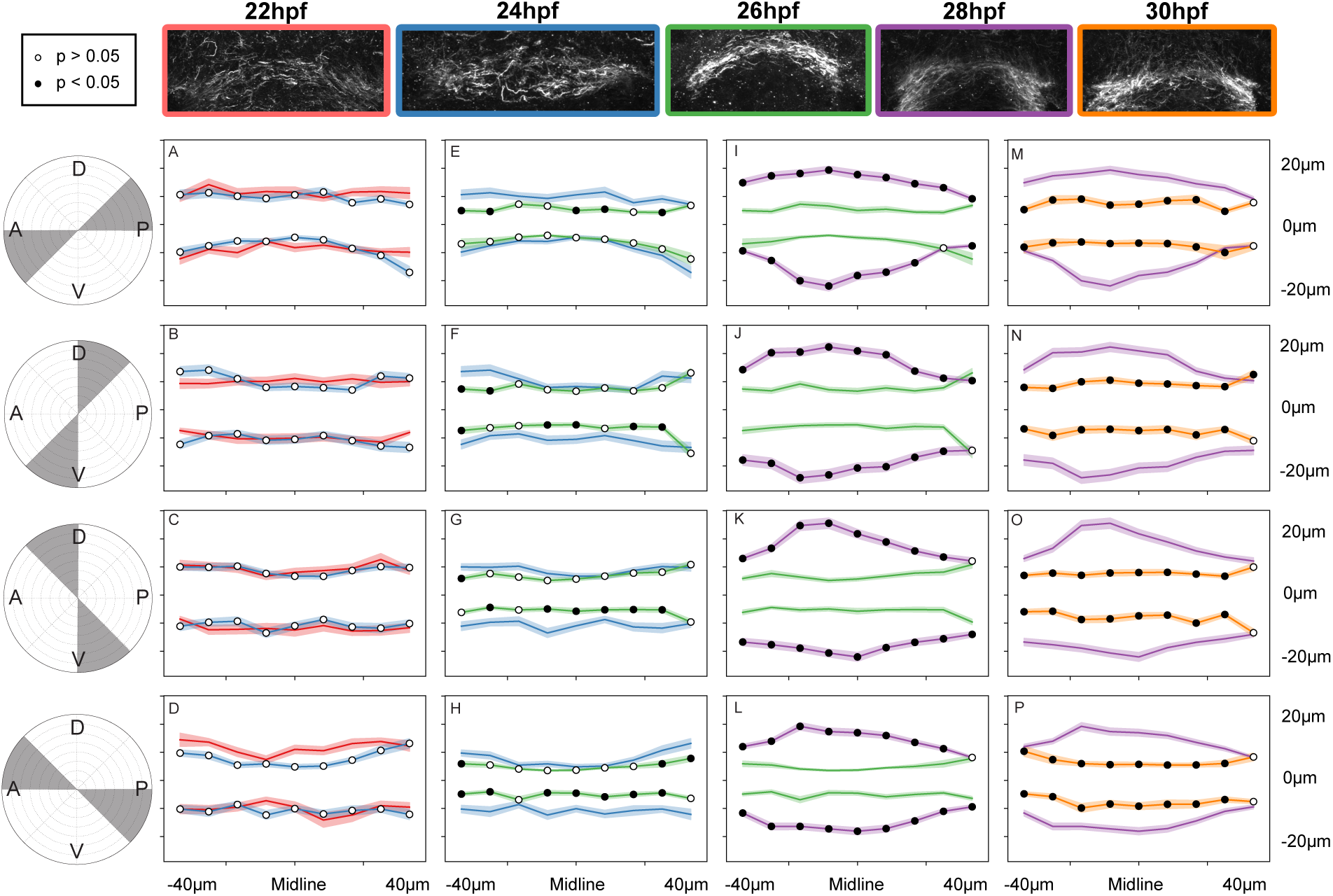
Landmark median radial distance analysis of glial bridge development and distribution. A-D) Statistical comparison of the radial distance of Gfap signal from the POC model in 22 hpf (n=13) and 24 hpf (n=20) POCs. No statistical difference observed. E-H) Statistical comparison of Gfap signal from the the POC model in 24 hpf (n=20) and 26 hpf (n=18). Significant reductions in radial distance was observed in all ventral posterior locations. I-L) Statistical comparison of the radial distance of Gfap signal from the POC model in 26 hpf (n=18) to 28 hpf (n=17) POCs. Significant increases in radial distance noted in all radial bins. M-P) Statistical comparison of the radial distance of Gfap signal from the POC model in 28 hpf (n=17) and 30 hpf (n=17). Significant reductions in radial distance observed in all radial bins.

**Figure 19:**
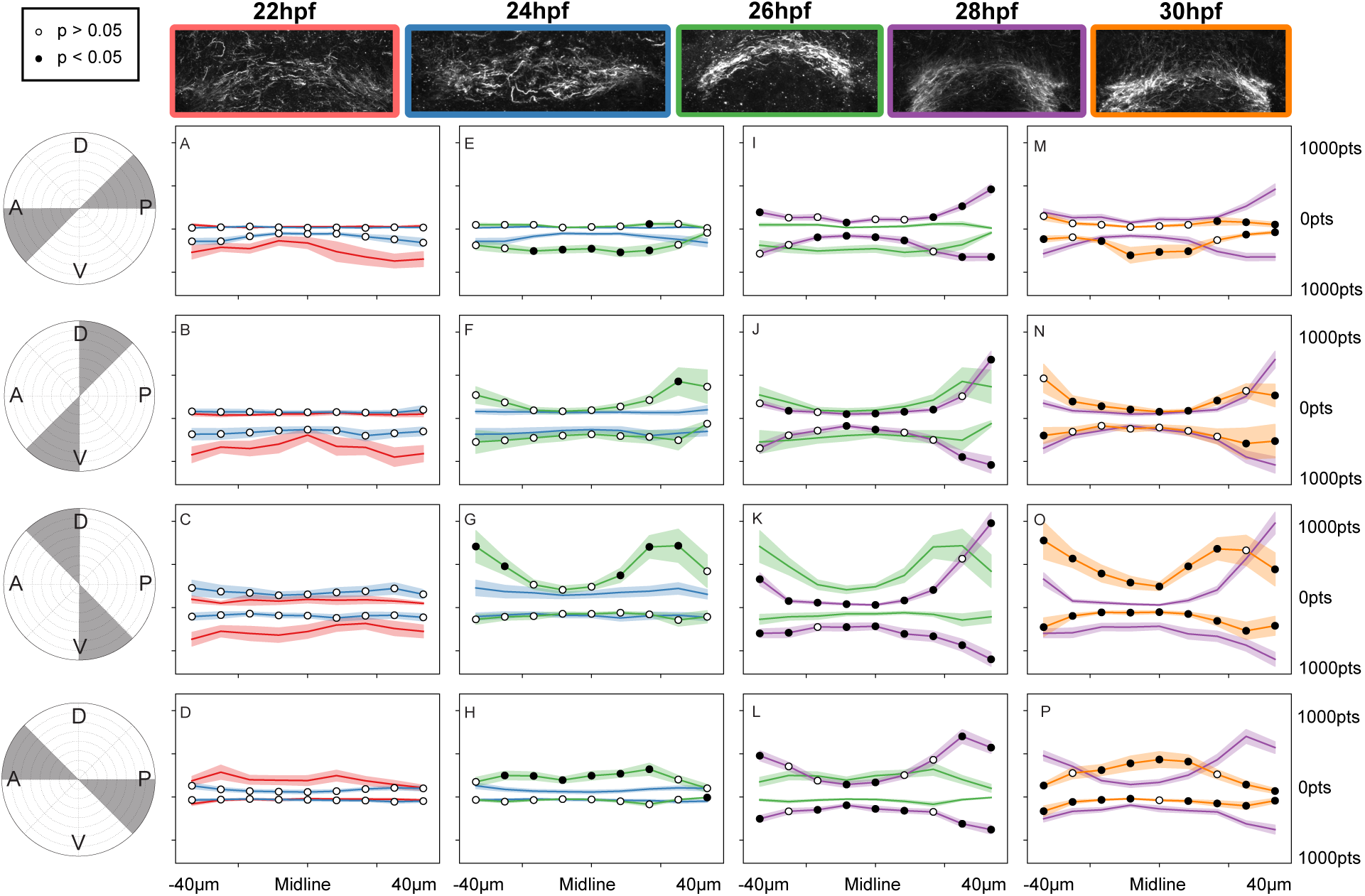
Landmark point analysis of anti-Gfap signal and glial bridge development. A-D) Statistical comparison of the number of Gfap positive pixels in 22 hpf (n=13) and 24 hpf (n=20) POCs. No statistical increase in Gfap signal. E-H) Statistical comparison of Gfap positive pixels between 24 hpf (n=20) and 26 hpf (n=18). Significant changes observed in ventral-anterior positions. I-L) Statistical comparison of Gfap positive pixels between 26 hpf (n=18) to 28 hpf (n=17) POCs. Statistical increase in Gfap in ventral-posterior locations and a decrease in ventral-anterior. M-P) Statistical comparison of Gfap positive pixels between 28 hpf (n=17) and 30 hpf (n=17). Significant increase in Gfap signal changes observed in anterior bins with a slight reduction in ventral-posterior locations.

### ΔSCOPE detects subtle commissural phenotypes

We have shown that ΔSCOPE can be used to describe and characterize both normal commissure development and severely disrupted commissural phenotypes in the *yot* mutant. However, one of the greatest challenges in studying the development of the nervous system is being able to uncover subtle phenotypes that may escape qualitative identification but still could have profound functional and behavioral deficits for the adult organism. We next tested the detection sensitivity of ΔSCOPE by applying its analysis to the effects of a misexpression of *slit1a* on POC development, for which we show here for the first time, represents a subtle commissural phenotype that has previously defied our ability to accurately characterize.

Slit1a is a member of the Slit family of axon guidance cues [25, 24], and was previously speculated that Slit1a may function distinctly from its other known repellent family members, Slit2 and Slit3 [15]. Knockdown of *slit2* and/or *slit3* results in defasciculation of the POC and expansion of the glial bridge in wild type embryos and was remarkably capable of restoring midline crossing of POC axons in the *yot* mutant background. In contrast, knockdown of *slit1a* results in a loss of commissure formation and demonstrating that *slit1a* is not capable of rescuing commissure formation in the *yot* mutant background. These and other expression data have led to the hypothesis that Slit1a may function as an attractant for midline crossing by POC axons [15]. If this were true, then over-expression of Slit1a in the *slit1a* depleted *yot* mutant background might be sufficient to rescue commissure formation.

To test this hypothesis, we used two transgenic lines to overexpress *slit1a-mCherry* and a control transcript of mCherry alone via the heatshock inducible promoter (tg*(hsp70:slit1a-mcherry)* and tg*(hsp70:mcherry)*) in both wild type and *yot* backgrounds. Qualitative examination of MIPs of heatshock-induced mCherry controls showed no difference between commissures of heatshocked and non-heatshocked embryos (Fig 20 A-D). In contrast, heatshock induction of Slit1a-mCherry caused apparent defasciculation of the POC in wild type embryos that was accompanied with a potential disruption in glial bridge positioning (Fig 20 E-H). Interestingly, overexpression of Slit1a-mCherry in the *yot* background resulted in a significant proportion of the embryos exhibiting an increase in axons projecting to and across the midline (compare Fig 20 I,J,M,N with Fig 20 K,L,O,P). In addition, there appeared to be similar disruptions in glial bridge organization in *yot* mutants as qualitatively observed in the wild type embryos following Slit1a-mCherry overexpression (Fig 20). However, in both of these experimental groups, sporadic axon wandering and defasciculation were observed, which made our interpretations of these qualitative results challenging. To more objectively analyze the results of these experiments, we took advantage of ΔSCOPE using the POC as the anchoring structural channel for the quantification of both POC axon and glial bridge comparisons across all conditions.

**Figure 20:**
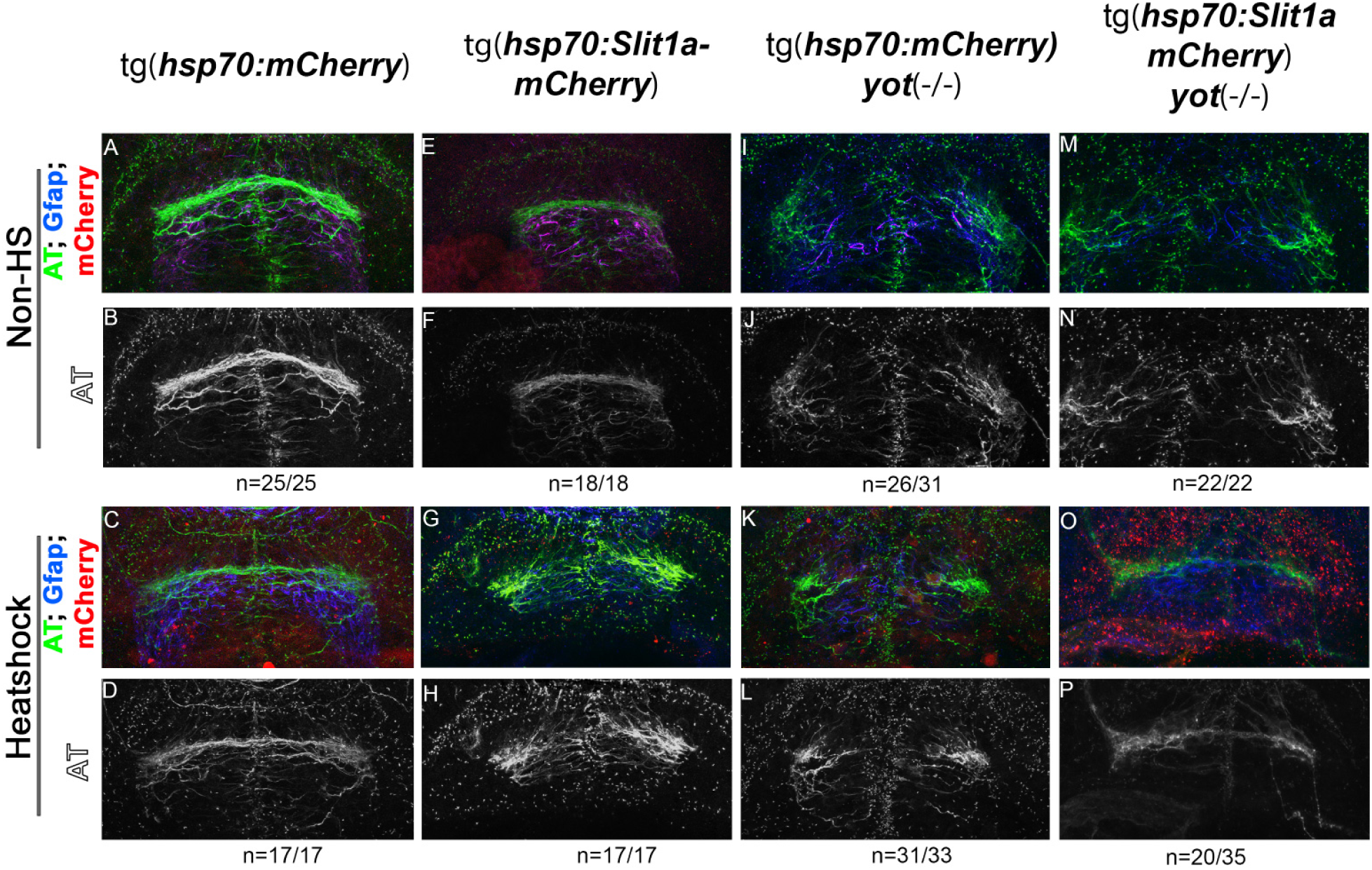
*slit1a* over-expression affects POC formation in wild type and homozygous *yot* embryos. A,B) Non-heatshock control of *hsp70:mcherry* embryo. A) Color composite MIP of frontal zebrafish forebrain showing POC and glial bridge, with cherry (red), Gfap (blue), AT (green), showing no red expression, and coincidence of the glial bridge (blue) with the POC (green). B) Single channel MIP of AT showing normal commissure formation. C,D) Heatshock control of *hsp70:mcherry embryo*. C) Color composite MIP of frontal zebrafish forebrain showing POC and glial bridge, with cherry (red), Gfap (blue), AT (green), showing red expression, and coincidence of the glial bridge (blue) with the POC (green). D) Single channel MIP of AT showing normal commissure formation. E,F) Non-heatshock control of *hsp70:slit1a-mcherry* embryo. E) Color composite MIP of frontal zebrafish forebrain showing POC and glial bridge, with cherry (red), Gfap (blue), AT (green), showing no red expression, and coincidence of the glial bridge (blue) with the POC (green). F) Single channel MIP of AT showing normal commissure formation. G,H) Heatshock *hsp70:slit1a-mcherry* embryo. G) Color composite MIP of frontal zebrafish forebrain showing POC and glial bridge, with cherry (red), Gfap (blue), AT (green), showing red expression, and disturbed glial bridge (blue) with a defasciculated POC (green). H) Single channel MIP of AT showing aberrant and defasciculated commissure formation. I,J) Non-heatshock control of *you-too* homozygous *hsp70:mcherry* embryo. I) Color composite MIP of frontal zebrafish forebrain showing POC and glial bridge, with cherry (red), Gfap (blue), AT (green), showing no red expression, and disturbed glial bridge formation (blue) and loss of commissure formation (green). J) Single channel MIP of AT showing loss of commissure formation. K,L) Heatshock control of *you-too* homozygous *hsp70:mcherry* embryo. J) Color composite MIP of frontal zebrafish forebrain showing POC and glial bridge, with cherry (red), Gfap (blue), AT (green), showing red expression, and disturbed glial bridge formation (blue) and loss of commissure formation (green). L) Single channel MIP of AT showing loss of commissure formation. M,N) Non-heatshock control of *you-too* homozygous *hsp70:slit1a-mcherry* embryo. M) Color composite MIP of frontal zebrafish forebrain showing POC and glial bridge, with cherry (red), Gfap (blue), AT (green), showing no red expression, and disturbed glial bridge formation (blue) and loss of commissure formation (green). N) Single channel MIP of AT showing loss of commissure formation. O,P) Heatshock *you-too* homozygous *hsp70:slit1a-mcherry* embryo. O) Color composite MIP of frontal zebrafish forebrain showing POC and glial bridge, with cherry (red), Gfap (blue), AT (green), showing red expression, and disturbed glial bridge formation (blue) and some commissure formation (green). P) Single channel MIP of AT showing partial commissure formation.

ΔSCOPE examination of Slit1a-mCherry overexpression in wild type showed a significant reduction in the number of anti-AT positive pixels (axons) at the midline in both the anterior (Fig 21 E, F) and dorsal-ventral axes (Fig 21 F, G). We further noted that all radial bins in Slit1a-mCherry overexpressed embryos exhibited significant expansion in radial distances from the modeled POC parabola, suggesting significant defasciculation (Fig 22 I-L). When compared to wild type embryos, Slit1a-mCherry overexpression in *yot* mutant embryos exhibited both significant reductions in anti-AT signal as well as in the R distance in all radial bins (Fig 21 I-L, Fig 22 I-L). Importantly, we acknowledge that this quantification of axon patterning in *yot* mutants in response to *slit1a* was not consistent with our earlier qualitative assessment that suggested a potential POC rescue (Fig 20 O,P). In fact, our ΔSCOPE analysis indicates that *slit1a-mCherry* overexpression results in a greater loss of anti-AT signal in all posterior axes when compared to non-heatshocked *yot* controls (Fig 22 Q-T). Likewise, more severe defasciculation was quantified in *yot* mutants following Slit1a-mCherry overexpression as compared to *yot* mutant controls (Fig 21 Q,S). Indeed, overexpression of Slit1a-mCherry in wild type resulted in phenotypes that were not statistically different from *yot* mutants alone (Fig 21 M-P, Fig 22 M-P). Our prior *slit1a* morphant loss of function data suggested that Slit1a was required for midline crossing, however we demonstrated here using ΔSCOPE that the misexpression of *slit1a* alone was insufficient to rescue midline crossing in *yot* mutants and rather caused even more deleterious effects on POC development.

**Figure 21:**
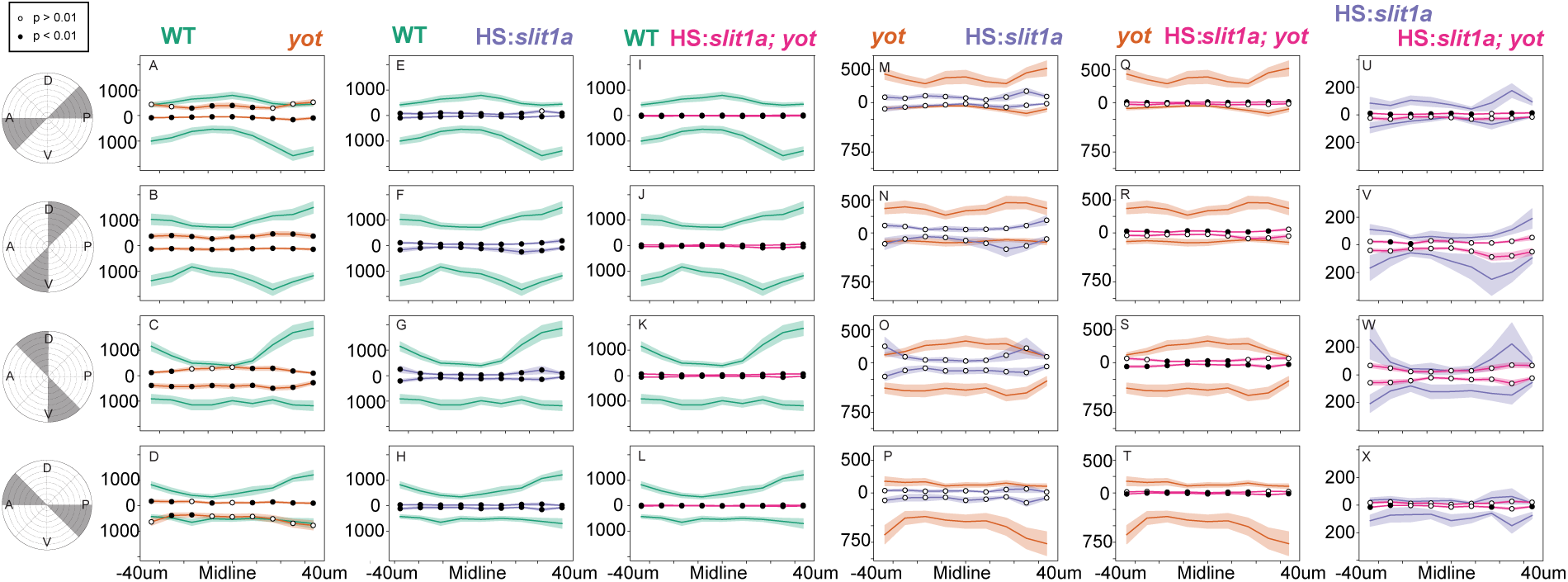
The effect of *slit1a* overexpression on commissure formation. Comparisons of the number of AT positive points in radial bins. A-D) Comparison of WT (n=37) (green) and *you-too* (n=33) (orange) homozygous POC. Loss of AT signal in *you-too* was observed in all octants except the posterior quadrant. E-H) Comparison of WT (n=37) (green) and heatshock *slit1a* (n=10) (purple) embryos. Significant reductions in AT signal in heatshock *slit1a* commissures was observed in dorsal ventral octants. I-L) Comparison of WT (n=37) (green) and heatshock *slit1a you-too* (n=17) (red) homozygous embryos. Significant reductions were observed in all octants of *slit1a you-too* commissures. M-P) Comparison of *you-too* homozygous (n=33) (orange) and heatshock *slit1a* (n=10) (purple) embryos. No significant differences observed. Q-T) Comparison of *you-too* homozygous (n=33) (orange) and heatshock *slit1a you-too* (n=17) (red) embryos. Significant reductions in the number of points in heatshock *slit1a you-too* embryos in both dorsal posterior and ventral posterior positions. U-X) Comparison of heatshock *slit1a* (n=10) (purple) and heatshock *slit1a you-too* (n=17) (red) embryos. Significant reductions in the number of AT points in heatshock *slit1a you-too* embryos were observed in the posterior quadrant.

**Figure 22:**
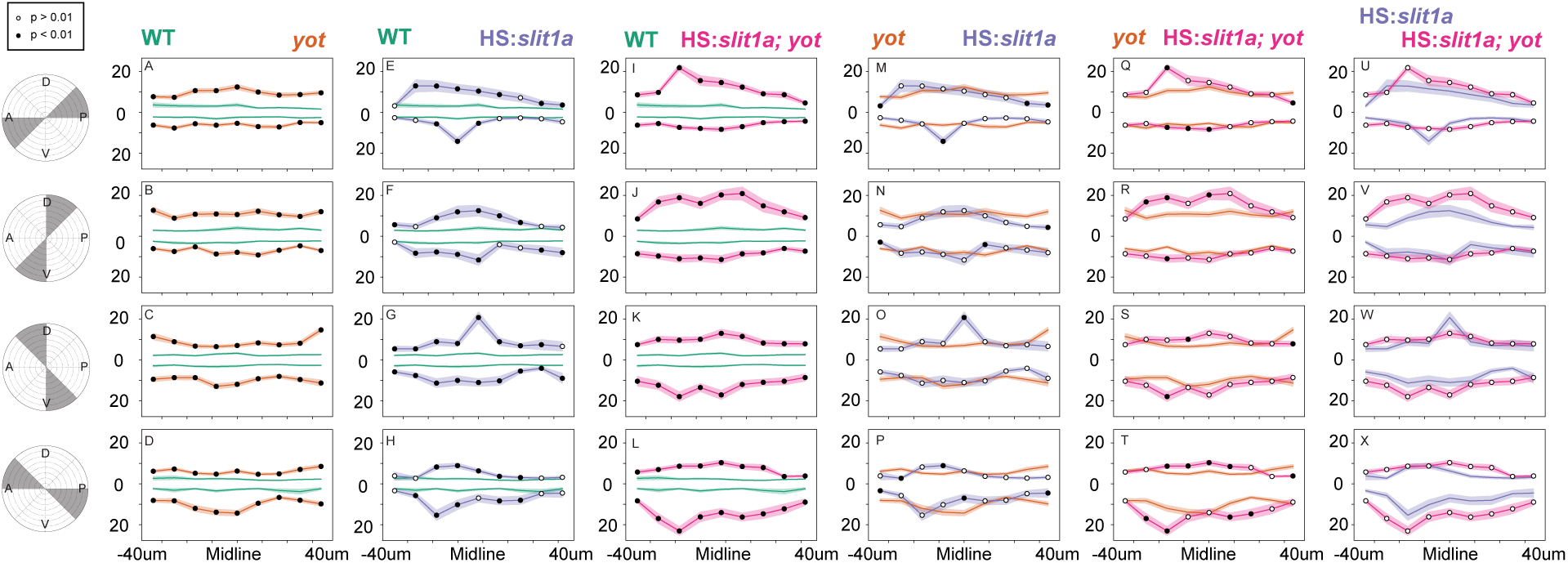
The effect of *slit1a* overexpression on commissure distribution and fasciculation. Comparisons of the radial distance of axon signal from the POC model. A-D) Comparison of WT (n=37) (green) and *you-too* (n=33) (orange) homozygous POC. Significant expansion of the radial distance in *you-too* was observed in all octants. E-H) Comparison of WT (n=37) (green) and heatshock *slit1a* (n=10) (purple) embryos. Significant expansion of the radial distance in heatshock *slit1a* was observed in all octants. I-L) Comparison of WT (n=37) (green) and heatshock *slit1a you-too* (n=17) (red) homozygous embryos. Significant expansions of the radial distance of AT signal in heatshock *slit1a you-too* was observed in all octants. M-P) Comparison of *you-too* homozygous (n=33) (orange) and heatshock *slit1a* (n=10) (purple) embryos. No significant differences observed. Q-T) Comparison of *you-too* homozygous (n=33) (orange) and heatshock *slit1a you-too* (n=17) (red) embryos. Significant expansions in radial distance of AT signal in heatshock *slit1a you-too* were observed in dorsal and dorsal anterior positions. U-X) Comparison of heatshock *slit1a* (n=10) (purple) and heatshock *slit1a you-too* (n=17) (red) embryos. No significant changes observed.

The disagreement of this new quantitative analysis of Slit1a function by ΔSCOPE with our previous qualitative assessments, suggests the guidance mechanisms of Slit1a may be more complex than our original model posited. Interestingly, *slit1a* has been shown to be expressed by cells of the glial bridge [15]. Moreover, we showed here for the first time that the development of the glial bridge and POC axons are inextricably linked, therefore we hypothesize that the primary role of Slit1a may be in the positioning of astroglial cells. We found through ΔSCOPE quantification that *slit1a-mCherry* misexpression has a significant effect on the distribution of Gfap in the forebrain. Although we detected only slight reductions in Gfap signal in response to *slit1a-mCherry* overexpression (Fig 23), there was a positive correlation with reductions in median R distance(Fig 24). Furthermore, we previously noted significant reductions of both median R distance and signal of Gfap in *you-too* embryos. As such, we note that both increases and decreases in Slit1a result in disruptions to the appropriate positioning of the glial bridge ((Fig 23; [15]). Our data suggests that Slit1a may first function to condense the cells of the glial bridge, and that disruptions to the proper patterning of the glial bridge results in the weakening of a key supportive structure to the guidance of POC axons across the midline.

**Figure 23:**
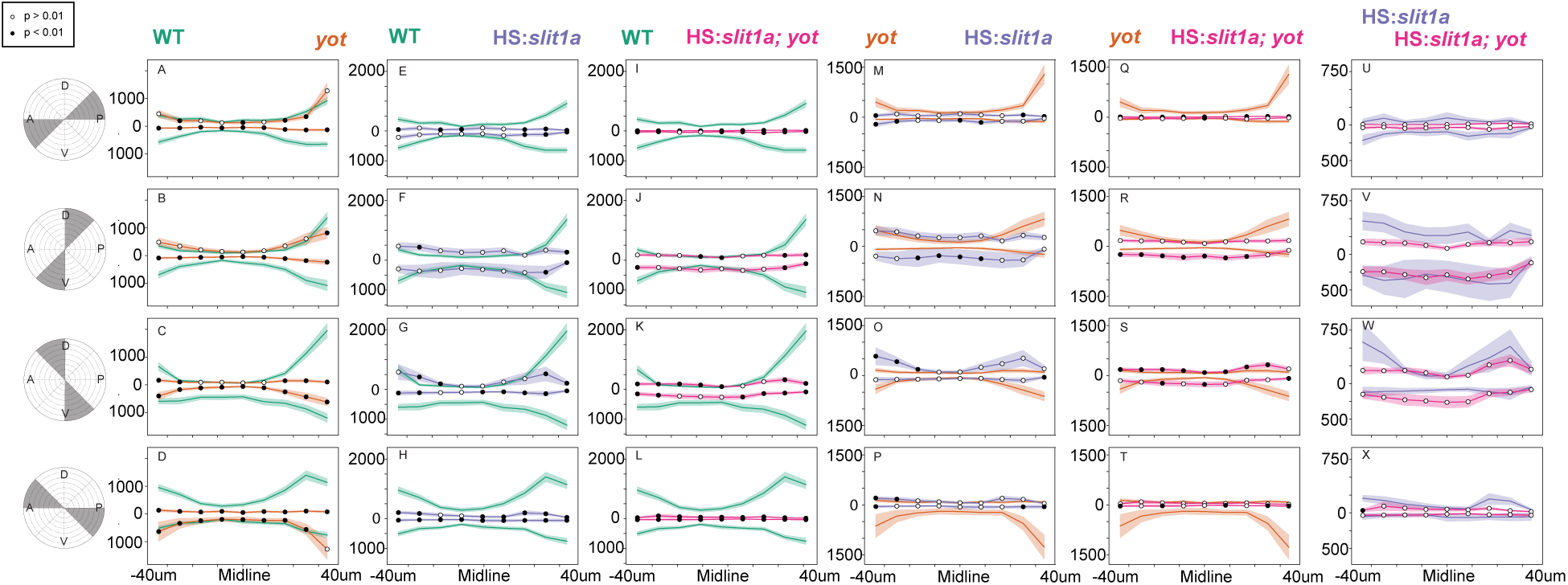
The effect of *slit1a* overexpression on glial bridge formation. Comparisons of the number of Gfap positive points in radial bins. A-D) Comparison of WT (n=37) (green) and *you-too* (n=33) (orange) homozygous glial bridge. Loss of Gfap signal in *you-too* was observed in dorsal anterior and octants and the lateral dorsal anterior octant. E-H) Comparison of WT (n=37) (green) and heatshock *slit1a* (n=10) (purple) embryos. Significant reductions in Gfap signal in heatshock *slit1a* the glial bridge was observed in the lateral-anterior and posterior octants. I-L) Comparison of WT (n=37) (green) and heatshock *slit1a you-too* (n=17) (red) homozygous embryos. Significant reductions in Gfap signal were observed in anterior and posterior octants of *slit1a you-too* glial bridges. M-P) Comparison of *you-too* homozygous (n=33) (orange) and heatshock *slit1a* (n=10) (purple) embryos. No significant differences observed. Q-T) Comparison of *you-too* homozygous (n=33) (orange) and heatshock *slit1a you-too* (n=17) (red) embryos. A significant increase in the number of Gfap positive points in heatshock *slit1a you-too* embryos was observed in the ventral octant of the glial bridge. U-X) Comparison of heatshock *slit1a* (n=10) (purple) and heatshock *slit1a you-too* (n=17) (red) embryos. No significant differences were observed.

**Figure 24:**
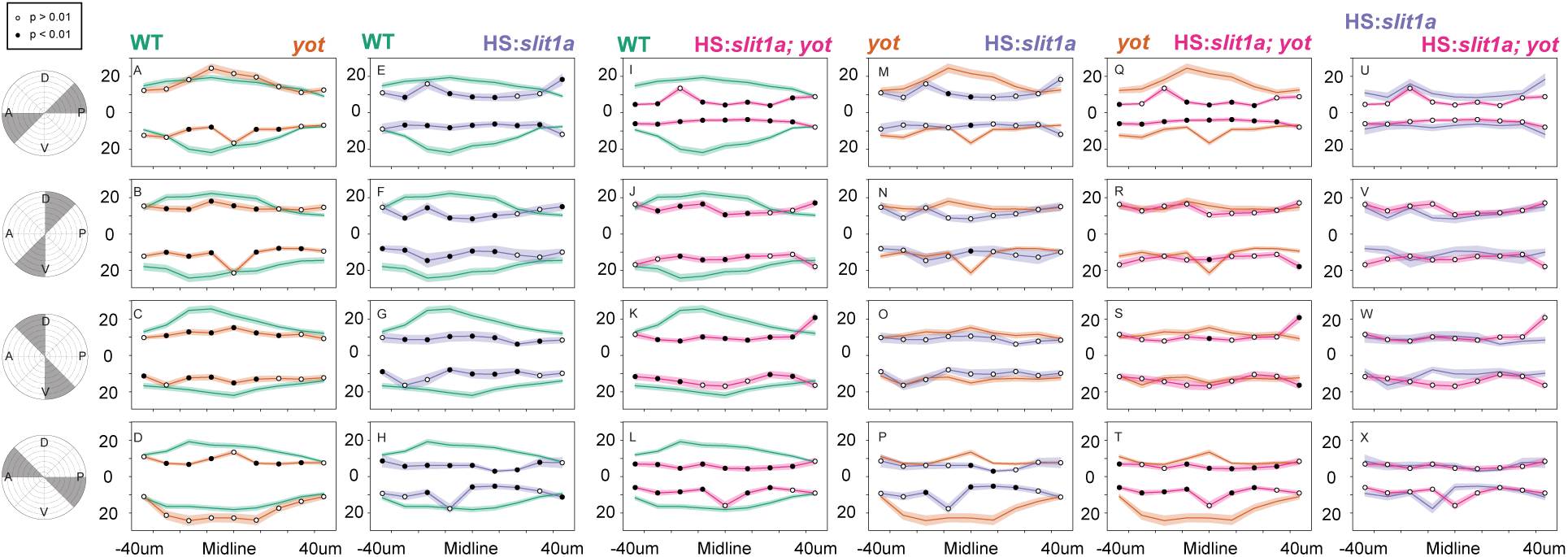
The effect of *slit1a* overexpression on glial bridge condensation. The effect of *slit1a* overexpression on glial bridge formation. Comparisons of the radial distance of Gfap signal from the POC model. A-D) Comparison of WT (n=37) (green) and *you-too* (n=33) (orange) homozygous POC. Significant reduction of the radial distance of Gfap signal in *you-too* glial bridges were observed in all anterior octants. E-H) Comparison of WT (n=37) (green) and heatshock *slit1a* (n=10) (purple) embryos. Significant reduction of the radial distance of Gfap signal in heatshock *slit1a* embryos was observed in all octants. I-L) Comparison of WT (n=37) (green) and heatshock *slit1a you-too* (n=17) (red) homozygous embryos. Significant reductions of the radial distance of Gfap signal in heatshock *slit1a you-too* was observed in all octants but the ventral and dorsal posterior octants. M-P) Comparison of *you-too* homozygous (n=33) (orange) and heatshock *slit1a* (n=10) (purple) embryos. Significant reductions in the radial distance of Gfap signal in heatshock *slit1a* glial bridges were observed in the posterior quadrant. Q-T) Comparison of *you-too* homozygous (n=33) (orange) and heatshock *slit1a you-too* (n=17) (red) embryos. Significant reductions in the radial distance of Gfap signal in heatshock *slit1a you-too* glial bridges were observed in anterior and posterior quadrants. U-X) Comparison of heatshock *slit1a* (n=10) (purple) and heatshock *slit1a you-too* (n=17) (red) embryos. No significant changes observed.

## Discussion

Unbiased tools to objectively analyze image-based data have not kept pace with advances in microscopy. The development of analytical tools which might enable image analysis have been largely hampered by the inherent challenges associated with 3D image data: image noise, sample variability, loss of dimensionality, and challenges in generating an average structure from multiple samples. To overcome these challenges, we built ΔSCOPE a new computational method to quantify 3D image-based data, which we have applied to the study of axon-glial interactions during commissure development in zebrafish. We demonstrated that ΔSCOPE’s innovative structural anchoring and use of principal component analysis enabled automated sample alignment to register the 3D pixel data of all samples being tested into a cylindrical coordinate system. With the data organized along this new coordinate system, we further showed how the integration of landmark analysis could enable not only the statistical quantification of two discrete biological but also associated structures between several different comparative conditions. Our application of ΔSCOPE to the analysis of axon and glial cell behaviors during commissure formation in the zebrafish forebrain has quantitatively proved for the first time the direct association of pathfinding POC axons with astroglial cell positioning during midline crossing. Moreover, we validated the sensitivity of ΔSCOPE by examining a zebrafish mutant and transgenic embryos with known severe and subtle phenotypes in POC pathfinding and glial cell positioning, respectively. Lastly, ΔSCOPE provided us the opportunity to report here for the first time that the Slit1a guidance cue was sufficient to cause modest but statistically significant commissural pathfinding errors as well as exert a direct influence on glial bridge development. Our results taken together suggest a model in which Slit1a may function to guide the organization of the glial bridge that then provides the necessary structural support for midline crossing POC axons.

### ΔSCOPE provides a structure-based method for adaptable analyzes

In order to achieve sample-to-sample alignment for comparative and statistical analyzes, we anchored ΔSCOPE around a common structural element. Although this approach solved the problem of image registration it also introduced a restriction of requiring a relatively consistent structure amongst all images being quantified. Thus, while ΔSCOPE was successful in defining the parameters of glial bridge development during POC formation (Fig 18 and Fig 19), it could not characterize the positioning of these same glial cells prior to the emergence of POC axons that served as the primary structural channel. Within the scope of this limitation, we predict that ΔSCOPE’s use of PCA will be applicable to any definite biological structure that exhibits consistently different signal distributions along the three dimensions. In other words, we anticipate that the PCA function of ΔSCOPE will not perform well on an alignment channel where signal is either randomly or evenly distributed across the field of view, such that two axes share similar signal distributions. However, we have also implemented manual PCA correction tools, including tools to hold known axes from image acquisition as constant. When paired with consistent image and structure acquisition, these tools can overcome limitations in PCA alignment resulting from poor axis variability. Moreover, image registration is currently of great interest to the field of microscopy, and we anticipate that advances in machine learning methods will provide new alternatives to PCA to ameliorate ΔSCOPE’s dependence on a consistent anisotropic structure.

ΔSCOPE also successfully retains the three-dimensionality of our data, which is rooted in the creation of a new cylindrical (biologically oriented) coordinate system. This coordinate system is based on a model of the biological structure of interest and transforms a Cartesian coordinate signal into a signal that is registered to the structure of interest. This approach reduced the impact of variation in sample positioning or microscopy and instead isolated the actual biological variation. We suggest that this new structure centric technique of evaluating data provides the user with the opportunity to see, evaluate, and test biological data in ways that have not been achieved in other coordinate or alignment paradigms. In particular, as two biological specimens are rarely structurally identical, Cartesian transformation tools often have to warp microscopy data to fit a reference structure, a process which may lead to the introduction of artificial variation. Of particular concern, transformations to a reference structure can obscure subtle biological differences between samples, which greatly limits the sensitivity of such techniques for the accurate assessment of subtle phenotypes. We have demonstrated here that ΔSCOPE is able to detect, quantify and statistically analyze even subtle axonal pathfinding errors (Fig 21). We propose that using the biological structure itself to describe the distribution of data affords a powerful new way to evaluate biological structures. We specifically recommend that the application of this cylindrical coordinate system would be similarly informative in the analysis of other 3D structures where the signal can be mathematically described by a simple algebraic equation and where a consistent centering point can be identified. For example, neuro-developmental structures, such as the spinal cord and optic cup, could respectively be described with a line that runs parallel or perpendicular to the structure.

ΔSCOPE leverages the program ilastik to remove noise while preserving the signal best representative of the experimental data. We acknowledge that this pre-processing step involved two aspects of data alteration, of which one could introduce human bias. Although the objectivity of the machine learning methods of ilastik reduced the influence of human biases, this approach first relied on expertly annotated training sets of data with a wide range of images and image intensities. We have observed that errors in training or the use of disparate image collection parameters can result in processed images with significantly higher levels of signal noise or alternatively completely blank data sets. From a statistical rigor perspective, it is fortunate that in both cases this tends to result in greater image variation and thus decreased statistical power, as opposed to a higher false positive rate. It is important to also acknowledge that we must apply a signal threshold to this post-processed data to generate a binary dataset for ΔSCOPE analysis. While we have observed that a wide range of thresholds were both tolerated by ΔSCOPE and anecdotally noted that varied thresholds do not significantly influence the ultimate results, the same may not be true of all potential future structural images analyzed with ΔSCOPE. We further note however that ΔSCOPE was sensitive to variations in sample labeling and intensity, as noted by its ability to detect changes in commissure signal and distribution (Fig 16 and 17). This sensitivity requires that comparative samples must be properly age matched and processed equivalently, which is a requirement that should fall within the norm for most experimental paradigms. Failure to account for experimental differences may result in the detection of significant differences between experiments that are due to real changes in label quality. Additionally, to assist future users of ΔSCOPE, we provide code designed to evaluate and determine the optimal threshold for the elimination of background or non-structure real signal and safeguard against these potential introduced biases. We conclude that ilastik processing generally results in images with less noise and greater preservation of signal as compared to using methods of raw intensity thresholding (Fig 2).

### ΔSCOPE reveals the complexities of axon and glial cell guidance

The wiring of the vertebrate brain is based upon a stereotypical pattern of neuron-to-neuron connections that are laid down during embryonic development through a process called axon guidance. Because of the foundational role axon guidance plays in building the nervous system, as well as repairing it during neural regeneration, there has been a long history of research into the signaling mechanisms governing the extracellular guidance information and the intracellular machinery required to interpret those environmental cues [18, 48, 49, 50, 51]. This research has traditionally relied upon the analysis of visual representations of axonal anatomy. However, while a species may exhibit conserved patterns of axon pathways, no two embryos of that same species are identical. Therefore, it has been a longstanding challenge of the field to generate significant confidence in the interpretation of axonal phenotypes, particularly when those changes may be subtle in appearance. We present here, ΔSCOPE, a new method to produce an unbiased, objective assessment of changes in axonal anatomy. To demonstrate its utility, we have leveraged ΔSCOPE to describe and quantify the development of the post optic commissure in the zebrafish forebrain.

It has been known that the loss of hedgehog signaling, in particular through the dominant repressive effects of the *you-too (gli2DR)* mutation, causes changes in gene expression of the slit family of guidance cues, resulting in both axon pathfinding and astroglial cell positioning errors [15]. This previous work by ourselves and colleagues was dependent on obvious midline crossing defect phenotypes, and it therefore lacked the analytical approaches to characterize the three-dimensionality of these phenotypes. We leveraged the current knowledge of axon and glial cell phenotypes in the *yot* mutant as means to first validate the sensitivity of ΔSCOPE, and then went further, to analyze these defects in much greater detail. Through this more quantitative approach we were able to show that ΔSCOPE was capable of detecting and identifying a statistically significant loss of midline crossing axons in the same exact locations where astroglial cells were similarly reduced particularly along the dorsal-posterior axes (compare (Fig 14) A,B) with (Fig 15 A,B)). In fact, our analysis suggested a much more refined understanding of POC axon phenotypes in *yot*, such that those axons that were found to be wandering did so preferentially closer to the pial surface as opposed to pathfinding deeper into the forebrain.

By applying this now validated ΔSCOPE analysis to POC axon and glial cell positioning over embryological time, we were able to generate the first quantitative description of pathfinding axons and glial bridge formation during commissure development. Our comparative measurements for changes in pixel position and quantity relative to the modeled commissure and midline revealed an initial period of POC axon remodeling as pathfinding axons approach and cross the midline followed by a period of increasing commissure fasciculation. Most fascinating was the paired quantification of glial bridge condensation about the modeled parabola, strongly suggesting a tight interaction between pathfinding POC axons and midline spanning astroglial cells (Figs 17, 18, 16, and 19).

Lastly, we extended our ΔSCOPE analysis of axon guidance to shed light upon the role that Slit1a may play during commissure formation. Based on differing responses, it was previously suggested that Slit1a may function distinctly from its known axon repellent family members Slit2 and Slit3 [15]. We present here the first demonstration of a temporally controlled misexpression of Slit1a just prior to commissure formation. Interestingly, while the widespread misexpression of Slit1a caused subtle but statistically significant indications of POC axon defasciculation (Fig 22), astroglial cell labeling appeared to respond differently by more tightly condensing around the modeled commissure (Fig 24). Such different axon and glial cell responses to the same guidance cue in the same context suggests the existence of different intracellular machinery to mediate the guidance of POC axons and glial cells to Slit1a. We interpret our data to suggest a model in which Slit1a first functions to condense the cells of the glial bridge, which then serves a more permissive role in the physical growth of POC axons across the midline.

As the amount and complexity of image-based data continues to grow, so will the need for improved methods that can bridge the gap between 3D visual inspection of data and quantitative analysis of that data. ΔSCOPE provides a new option for the quantification and statistical analysis of 3D visual data. We purport that ΔSCOPE presents a new paradigm for image analysis, representing a shift away from the use of reference atlases or evaluation of MIPS. In particular ΔSCOPE enables the use of multiple embryos for generation of an averaged structure that fosters quantitative and statistical analyses of the biology while reducing the impact of normal biological variation which is frequently emphasized in atlases and MIPs. Further, we present ΔSCOPE as an extensible, open source software, which others can use, edit, and adapt to suit their needs and biological question, allowing it to adapt to meet the “dimensions” of future systems and questions.

### Distribution and accessibility

The code base of ΔSCOPE is available online as a Python package hosted on the https://pypi.org/project/deltascope/. The raw code repository is available on https://github.com/msschwartz21/deltascope, which records changes to the code and logs issues encountered by users of the code. In order to facilitate ease-of-use, extensive documentation of ΔSCOPE and its associated workflows is available on https://deltascope.readthedocs.io/en/latest/.

## Supporting information

An archived copy of the DeltaSCOPE code repository.

A pdf copy of documentation that accompanies the DeltaSCOPE code repository.

Cylindrical coordinate point cloud rotation of single AT data

Cylindrical coordinate point cloud rotation of single Gfap data

Cylindrical coordinate point cloud rotation of combined AT (green)/Gfap (red) data

Landmark rotation movie of a single sample of AT data

Landmark rotation movie of a single sample of Gfap data

Landmark rotation movie of a single sample of combined AT (green)/Gfap (red) data

## Supporting information

*S1 File*. **An archived copy of the** Δ**SCOPE code repository**. The most up-to-date version of the code is available on https://github.com/msschwartz21/deltascope or for installation via https://pypi.org/project/deltascope/Pyt Package Index.

*S2 File*. **A pdf copy of documentation that accompanies the** Δ**SCOPE code repository**. Also available on https://deltascope.readthedocs.io/en/latest/.

*S1 Data*. **Data associated with the developmental timecourse analysis of the glial bridge and post-optic commissure**. https://doi.org/10.35482/bld.003.2019

*S2 Data*. **Data associated with the analysis of the role of S1a**. https://doi.org/10.35482/bld.002.2019

*S1 Table*. **Table of key resources used in generation of microscopy data**, Δ**SCOPE code base, and locations of data repositories**.

*S1 Movie*. **Cylindrical coordinate point cloud rotation of single AT data**.

*S2 Movie*. **Cylindrical coordinate point cloud rotation of single Gfap data**

*S3 Movie*. **Cylindrical coordinate point cloud rotation of combined AT (green)/Gfap (red) data**

*S4 Movie*. **Landmark rotation movie of a single sample of AT data**.

*S5 Movie*.. **Landmark rotation movie of a single sample of Gfap data**

*S6 Movie*. **Landmark rotation movie of a single sample of combined AT (green)/Gfap (red) data**

## Acknowledgments

We would like to thank the generous contributions of Melissa Hardy and the late Chi-Bin Chien to this research in their development and donation of the heatshock (hsp70) responsive *slit1a* tg*hsp70:slit1a:mcherry* and tg*hsp70:mcherry* lines. We would like to Nadia PenkoffLiedbeck, Cassie Kemmler, and Jin-Sook Park for their substantial contributions to data collection and project assistance. We would also like to thank Margaret Perry, Audrey Bertin, Kalynn Kosyka, Crystal Zang, Emma Ning for their assistance reviewing the code base and providing technical assistance weeding out errors. This work would not have been possible without the generous contributions of time and effort by the Smith College Center for Microscopy and its director, Judith Wopereis, and the Smith Animal Care facility. We are very thankful for all of the technical support provided by Alicia Famiglietti and all constructive discourse by the entire Barresi lab throughout this work.

## Contribution

MJFB, JS, and MSS conceived the project. BSB, JS, and MSS developed the methodology. MPHL, MS wrote the code. MPHL and JS performed experiments and collected data. JS annotated training data for ilastik pixel classification. MPHL, JS, and MS corrected commissure alignments. JS and MS wrote the original draft. MJFB, MPHL, and JS revised the final draft. MJFB and BSB supervised the project. MJFB funded the project.

