## Supplementary material for "ΔSCOPE: A new method to quantify 3D biological structures and identify differences in zebrafish forebrain development": An archived copy of the DeltaSCOPE code repository.: CHANGELOG.md

# Change Log

## [1.0.0] - 2019-07-08
Official publication release.
### Additions
- Added experimental files associated with publication

## [0.2.6] - 2019-02-11
### Changes
- Restore read the docs build functionality for autodocs

## [0.2.5] - 2019-02-11
### Additions
- Created template notebooks in the experiments directory

## [0.2.4] - 2018-08-23
### Changes
- Add sys path to sphinx conf file
- Add all modules as mock modules for read the docs

## [0.2.3] - 2018-08-23
### Changes
- Correct unittest.mock import to mock in conf.py file

## [0.2.2] 2018-08-26
### Additions
- Created a new class, `anumSelect`, which determines the optimum number of bins along alpha
- Created two new glasses, `graphData` and `graphSet` to handle plotting landmark data
- Created a new class `treeClassifier` which handles running the random forest classifier
- Added functions to facilitate corrections to pca alignment
- Added utility.py which contains functions that assist with manual alignment correction
### Changes
- In lm.calc_bins, input is a dictionary of dfs as opposed to a list of df values from dfs.values()
- `convert_to_arr` added option for selecting r or pts datatype (`DT`)
- Added the datatype parameter required for `convert_to_arr` to the `anumSelect` functions
- `raw_data` funtion changed to select channel so that background = 1 and signal = 0 in line with expectation that pixel values function like a p value
- Correct `calc_variance` to use `np.nanvar` instead of `np.var`
- Change names to deltascope in github and folders
- Median threshold value is rescaled to accommodate 0-255 image data that is output by skimage.filters.median
- `find_r` was only calculating distance in Y and Z so added X as well
- median filter threshold in `calculate_pca_median` is set to select all points that do not have a value of 255 (background). User input signal is not used.
- Move content from old_cranium back into __init__.py
- Change documentation references to cranium to deltascope

## [0.2.1] - 2018-01-14
### Changes
- Correct unittest.mock import to mock in conf.py file

## [0.2.0] - 2018-01-14
### Additions
- Json module implemented for config data structure
- Added data folder with two samples and a config file for testing
- Added 2D transformation option to mp-transformation.py
- Placing mp-transformation script in the cranium directory to function as a module
- Added try/except statements to each processing step in mp-transformation to allow the script to continue running if a single sample failed
- `calc_variance` function calculates bin variance and sample variance for a particular landmark size
- `anumSelect` class for finding the optimum value of anum in landmark calculation
### Changes
- Changed `convert_to_arr` to accommodate a main array and a list of additional arrays
- `read_psi_to_dict` now uses regular expression to find sample numbers instead of splitting and indexing the file path
- `convert_to_arr`  uses the minimum bin value for cartesian as opposed to the mean so that it is compatible with lm.acbins and lm.tbins


## [0.1.8] - 2018-01-10
### Changes
- Update matplotlib requirement from 1.5 to 2.0 to avoid installation problems with matplotlib dependencies for freetype and pngg

## [0.1.7] - 2018-01-10
### Changes
- Pip installing and importing mock library in place of unittest.mock

## [0.1.6] - 2018-01-10
### Changes
- Changed pytz requirement from 2017 to 2017.3 in response to build fail on RTD

## [0.1.5] - 2018-01-10
### Changes
- Allowed any numpy package >1.0 and <2.0
- Softened other package requirements to allow any patch number
### Added
- Returned the matplotlib dependency

## [0.1.4] - 2018-01-09
### Changes
- Temporarily removing the matplotlib dependency while working on beta testing

## [0.1.3] - 2018-01-08
### Changes
- Manually added package requirements to setup.py install-requires

## [0.1.2] - 2018-01-08
### Changes
- Implemented mock shielding for c dependent modules (numpy,scipy,pandas) in conf.py

## [0.1.1] - 2018-01-08
### Changed
- Removed `import cranium` from conf.py because it was causing error with read the docs builds

## [0.1.0] - 2018-01-08
### Added
- Apply double median filter to primary channel to calculate PCA fit; Removes noise to create a cleaner dataset which fits the POC into the xz plane
- Landmark code analysis
- Implemented autodoc system for sphinx
### Changed
- Fit model in the xz plane as opposed to the xy plane to match natural position of the parabolic commissure in approximately xz
- Upside down samples are fliped using a 180 degree rotation matrix as opposed to multiplying the z axis by -1
- Corrected coordinate transform where y and z were mixed up from when model was assumed to lie in xy plane
### Deprecated
- pca_transform
- calculate_pca
- add_pca
- pca_double_transform

## [0.0.6] - 2017-06-02
### Added
- Implemented PCA to align samples along consistent axes
- Vertex of data/math model is centered at the origin in the XY plane
- Checking sign of a coefficient in model and multiplying y coordinates by -1 as necessary
- Created embryo class to manage multiple channels associated with a single sample
### Changed
- Initialiization of brain object does not automatically run `read_data` so the command needs to be called seperately by the user
### Deprecated
- Eliminated the plane intersection method used originally to find the math model

## [O.0.5] - 2017-04-23
### Changed
- Using arclength from vertex instead of alpha due to alpha's uneven point distribution

[Unreleased]: https://github.com/msschwartz21/craniumPy/compare/v0.0.6...HEAD
[0.0.6]: https://github.com/msschwartz21/craniumPy/compare/v0.0.5...v0.0.6
