## Supplementary material for "ΔSCOPE: A new method to quantify 3D biological structures and identify differences in zebrafish forebrain development": An archived copy of the DeltaSCOPE code repository.: __init__.py

```
import os
import json
import numpy as np
import h5py ######### Updated recently and now module can't be started
#import skimage.io as io
import matplotlib.pyplot as plt
import time
import itertools as it
import matplotlib as mpl
import multiprocessing as mp
from functools import partial
#import statsmodels.formula.api as smf
#from mpl_toolkits.mplot3d import Axes3D
import pandas as pd
#import plotly.plotly as py
#import plotly.graph_objs as go
from scipy.optimize import minimize
from sklearn.preprocessing import normalize,scale
import scipy
from sklearn.decomposition import PCA
from skimage.filters import median
from skimage.morphology import disk
from sklearn.metrics import mean_squared_error
from scipy.integrate import simps
import scipy.stats as stats
import re
from sklearn.ensemble import RandomForestClassifier
import tqdm
import glob

class brain:
	''' Object to manage biological data and associated functions. '''

	def __init__(self):
		'''Initialize brain object'''

	def setup_test_data(self,size=None,gthresh=0.5,scale=[1,1,1],microns=[0.16,0.16,0.21],mthresh=0.2,radius=20,comp_order=[0,2,1],fit_dim=['x','z'],deg=2):
		'''Setup a test dataset to use for testing transform coordinates
		:param int size: Number of points to sample for the test dataset
		'''

		self.read_data(os.path.abspath('..\data\C1\AT_01_Probabilities.h5'))
		self.preprocess_data(gthresh,scale,microns)
		self.calculate_pca_median(self.raw_data,mthresh,radius,microns)
		self.pca_transform_3d(self.df_thresh,self.pcamed,comp_order,fit_dim,deg=deg)
		self.mm = self.fit_model(self.df_align,2,['x','z'])
		if size!=None:
			self.df_align = self.df_align.sample(size)

	def read_data(self,filepath):
		'''
		Reads 3D data from file and selects appropriate channel based on the
		assumption that the channel with the most zeros has zero as the value for no signal

		:param str filepath: Filepath to hdf5 probability file
		:return: Creates the variable :attr:`brain.raw_data`

		.. py:attribute:: brain.raw_data

			Array of shape [z,y,x] containing raw probability data
		'''

		#Read h5 file and extract probability data
		f = h5py.File(filepath,'r')

		#Select either channe0/1 or exported_data
		d = f.get('exported_data')
		if d != None:
			c1 = np.array(d[:,:,:,0])
			c2 = np.array(d[:,:,:,1])
		elif d == None:
			c1 = np.array(f.get('channel0'))
			c2 = np.array(f.get('channel1'))

		#Figure out which channels has more zeros and therefore is background
		if np.count_nonzero(c1<0.1) > np.count_nonzero(c1>0.9):
			#: Array of shape [z,y,x] containing raw probability data
			self.raw_data = c2
		else:
			#: Array of shape [z,y,x] containing raw probability data
			self.raw_data = c1

	def create_dataframe(self,data,scale):
		'''
		Creates a pandas dataframe containing the x,y,z and signal/probability value for each point in the :py:attr:`brain.raw_data` array

		:param array data: Raw probability data in 3D array
		:param array scale: Array of length three containing the micron values for [x,y,z]
		:return: Pandas DataFrame with xyz and probability value for each point
		'''

		#NB: scale variable actually contains microns dimensions

		dim = data.shape
		xyz = np.zeros((dim[0],dim[1],dim[2],4))

		#Generate array with xyz values for each point
		for x in range(dim[2]):

			xyz[:,:,x,2] = x
			xyz[:,:,x,3] = data[:,:,x]

			zindex = np.arange(0,dim[0],1)
			yindex = np.arange(0,dim[1],1)
			gy,gz = np.meshgrid(yindex,zindex)

			xyz[:,:,x,0] = gz
			xyz[:,:,x,1] = gy

		flat = np.reshape(xyz,(-1,4))

		#Create dataframe of points and scale to microns
		df = pd.DataFrame({'x':flat[:,2]*scale[0],'y':flat[:,1]*scale[1],'z':flat[:,0]*scale[2],'value':flat[:,3]})
		return(df)

	def plot_projections(self,df,subset):
		'''
		Plots the x, y, and z projections of the input dataframe in a matplotlib plot

		:param pd.DataFrame df: Dataframe with columns: 'x','y','z'
		:param float subset: Value between 0 and 1 indicating what percentage of the df to subsample
		:returns: Matplotlib figure with three labeled scatterplots
		'''

		df = df.sample(frac=subset)

    	#Create figure and subplots
		fig = plt.figure(figsize=(12,6))
		ax = fig.add_subplot(131)
		ay = fig.add_subplot(132)
		az = fig.add_subplot(133)

		#Create scatter plot for each projection
		ax.scatter(df.x,df.z)
		ay.scatter(df.x,df.y)
		az.scatter(df.z,df.y)

		#Plot model
		xvalues = np.arange(np.min(df.x),np.max(df.x))
		ax.plot(xvalues,self.mm.p(xvalues),c='y')

		#Add labels
		ax.set_title('Y projection')
		ay.set_title('Z projection')
		az.set_title('X projection')
		ax.set_xlabel('X')
		ax.set_ylabel('Z')
		ay.set_xlabel('X')
		ay.set_ylabel('Y')
		az.set_xlabel('Z')
		az.set_ylabel('Y')

		#Adjust spacing and show plot
		plt.subplots_adjust(wspace=0.4)

		return(fig)

	def preprocess_data(self,threshold,scale,microns):
		'''
		Thresholds and scales data prior to PCA

		Creates :py:attr:`brain.threshold`, :py:attr:`brain.df_thresh`, and :py:attr:`brain.df_scl`

		:param float threshold: Value between 0 and 1 to use as a cutoff for minimum pixel value
		:param array scale: Array with three values representing the constant by which to multiply x,y,z respectively
		:param array microns: Array with three values representing the x,y,z micron dimensions of the voxel

		.. py:attribute:: brain.threshold

			Value used to threshold the data prior to calculating the model

		.. py:attribute:: brain.df_thresh

			Dataframe containing only points with values above the specified threshold

		.. py:attribute:: brain.df_scl

			Dataframe containing data from :py:attr:`brain.df_thresh` after a scaling value has been applied
		'''

		#: Dataframe with four columns: x,y,z,value with all points in :py:attr:`brain.raw_data`
		self.df = self.create_dataframe(self.raw_data,microns)

		#Create new dataframe with values above threshold
		self.threshold = threshold
		self.df_thresh = self.df[self.df.value < self.threshold]

		#Scale xyz by value in scale array to force PCA axis selection
		self.scale = scale
		self.df_scl = pd.DataFrame({
			'x':self.df_thresh.x * self.scale[0],
			'y':self.df_thresh.y * self.scale[1],
			'z':self.df_thresh.z * self.scale[2]})

	def process_alignment_data(self,data,threshold,radius,microns):
		'''
		Applies a median filter twice to the data which is used for alignment

		Ensures than any noise in the structural data does not interfere with alignment

		:param array data: Raw data imported by the function :py:func:`brain.read_data`
		:param float threshold: Value between 0 and 1 to use as a cutoff for minimum pixel value
		:param int radius: Integer that determines the radius of the circle used for the median filter
		:param array microns: Array with three values representing the x,y,z micron dimensions of the voxel
		:returns: Dataframe containing data processed with the median filter and threshold
		'''

		#Iterate over each plane and apply median filter twice
		self.out = np.zeros(data.shape)
		for z in range(data.shape[0]):
			self.out[z] = median(median(data[z],disk(radius)),disk(radius))

		outdf = self.create_dataframe(self.out,microns)
		thresh = outdf[outdf.value != 255]
		return(thresh)

	def calculate_pca_median(self,data,threshold,radius,microns):
		'''
		Calculate PCA transformation matrix, :py:attr:`brain.pcamed`, based on data (:py:attr:`brain.pcamed`) after applying median filter and threshold

		:param array data: 3D array containing raw probability data
		:param float threshold: Value between 0 and 1 indicating the lower cutoff for positive signal
		:param int radius: Radius of neighborhood that should be considered for the median filter
		:param array microns: Array with three values representing the x,y,z micron dimensions of the voxel

		.. py:attribute:: brain.median

			Pandas dataframe containing data that has been processed with a median filter twice and thresholded

		.. py:attribute:: brain.pcamed

			PCA object managing the transformation matrix and any resulting transformations

		'''

		self.median = self.process_alignment_data(data,threshold,radius,microns)

		self.pcamed = PCA()
		self.pcamed.fit(self.median[['x','y','z']])

	def calculate_pca_median_2d(self,data,threshold,radius,microns):
		'''
		Calculate PCA transformation matrix for 2 dimensions of data, :py:attr:`brain.pcamed`, based on data after applying median filter and threshold

		.. warning:: `fit_dim` is not used to determine which dimensions to fit. Defaults to x and z

		:param array data: 3D array containing raw probability data
		:param float threshold: Value between 0 and 1 indicating the lower cutoff for positive signal
		:param int radius: Radius of neighborhood that should be considered for the median filter
		:param array microns: Array with three values representing the x,y,z micron dimensions of the voxel
		'''

		self.median = self.process_alignment_data(data,threshold,radius,microns)

		self.pcamed = PCA()
		self.pcamed.fit(self.median[['y','z']])

	def pca_transform_2d(self,df,pca,comp_order,fit_dim,deg=2,mm=None,vertex=None,flip=None):
		'''
		Transforms `df` in 2D based on the PCA object, `pca`, whose transformation matrix has already been calculated

		Calling :py:func:`brain.align_data` creates :py:attr:`brain.df_align`

		.. warning:: `fit_dim` is not used to determine which dimensions to fit. Defaults to x and z

		:param pd.DataFrame df: Dataframe containing thresholded xyz data
		:param pca_object pca: A pca object containing a transformation object, e.g. :py:attr:`brain.pcamed`
		:param array comp_order: Array specifies the assignment of components to x,y,z. Form [x component index, y component index, z component index], e.g. [0,2,1]
		:param array fit_dim: Array of length two containing two strings describing the first and second axis for fitting the model, e.g. ['x','z']
		:param int deg: (or None) Degree of the function that should be fit to the model. deg=2 by default
		:param mm: (:py:class:`math_model` or None) Math model for primary channel
		:param array vertex: (or None) Array of type [vx,vy,vz] (:py:attr:`brain.vertex`) indicating the translation values
		:param Bool flip: (or None) Boolean value to determine if the data should be rotated by 180 degrees
		'''

		fit = pca.transform(df[['y','z']])
		df_fit = pd.DataFrame({
			'x':df.x,
			'y':fit[:,comp_order[1]-1],
			'z':fit[:,comp_order[2]-1]
			})

		self.align_data(df_fit,fit_dim,deg=2,mm=None,vertex=None,flip=None)

	def pca_transform_3d(self,df,pca,comp_order,fit_dim,deg=2,mm=None,vertex=None,flip=None):
		'''
		Transforms `df` in 3D based on the PCA object, `pca`, whose transformation matrix has already been calculated

		:param pd.DataFrame df: Dataframe containing thresholded xyz data
		:param pca_object pca: A pca object containing a transformation object, e.g. :py:attr:`brain.pcamed`
		:param array comp_order: Array specifies the assignment of components to x,y,z. Form [x component index, y component index, z component index], e.g. [0,2,1]
		:param array fit_dim: Array of length two containing two strings describing the first and second axis for fitting the model, e.g. ['x','z']
		:param int deg: (or None) Degree of the function that should be fit to the model. deg=2 by default
		:param mm: (:py:class:`math_model` or None) Math model for primary channel
		:param array vertex: (or None) Array of type [vx,vy,vz] (:py:attr:`brain.vertex`) indicating the translation values
		:param Bool flip: (or None) Boolean value to determine if the data should be rotated by 180 degrees
		'''

		fit = pca.transform(df[['x','y','z']])
		df_fit = pd.DataFrame({
			'x':fit[:,comp_order[0]],
			'y':fit[:,comp_order[1]],
			'z':fit[:,comp_order[2]]
			})

		self.align_data(df_fit,fit_dim,deg=deg,mm=mm,vertex=vertex,flip=flip)

	def align_data(self,df_fit,fit_dim,deg=2,mm=None,vertex=None,flip=None):
		'''
		Apply PCA transformation matrix and align data so that the vertex is at the origin

		Creates :py:attr:`brain.df_align` and :py:attr:`brain.mm`

		:param pd.DataFrame df: dataframe containing thresholded xyz data
		:param array comp_order: Array specifies the assignment of components to x,y,z. Form [x component index, y component index, z component index], e.g. [0,2,1]
		:param array fit_dim: Array of length two containing two strings describing the first and second axis for fitting the model, e.g. ['x','z']
		:param int deg: (or None) Degree of the function that should be fit to the model. deg=2 by default
		:param mm: (:py:class:`math_model` or None) Math model for primary channel
		:param array vertex: (or None) Array of type [vx,vy,vz] (:py:attr:`brain.vertex`) indicating the translation values
		:param Bool flip: (or None) Boolean value to determine if the data should be rotated by 180 degrees

		.. py:attribute:: brain.df_align

			Dataframe containing point data aligned using PCA

		.. py:attribute:: brain.mm

			Math model object fit to data in brain object
		'''

		#If vertex for translation is not included
		if vertex == None and deg==2:
			#Calculate model
			model = np.polyfit(df_fit[fit_dim[0]],df_fit[fit_dim[1]],deg=deg)
			p = np.poly1d(model)

			#Find vertex
			a = -model[1]/(2*model[0])
			if fit_dim[0] == 'x':
				vx = a
				if fit_dim[1] == 'y':
					vy = p(vx)
					vz = df_fit.z.mean()
				else:
					vz = p(vx)
					vy = df_fit.y.mean()
			elif fit_dim[0] == 'y':
				vy = a
				if fit_dim[1] == 'x':
					vx = p(a)
					vz = df_fit.z.mean()
				else:
					vz = p(a)
					vx = df_fit.x.mean()
			elif fit_dim[0] == 'z':
				vz = a
				if fit_dim[1] == 'x':
					vx = p(a)
					vy = df_fit.y.mean()
				else:
					vy = p(a)
					vx = df_fit.x.mean()
			self.vertex = [vx,vy,vz]
		elif deg == 1:
			#Calculate model
			model = np.polyfit(df_fit[fit_dim[0]],df_fit[fit_dim[1]],deg=deg)
			p = np.poly1d(model)

			if fit_dim[0] == 'x':
				vx = df_fit.x.mean()
				if fit_dim[1] == 'y':
					vy = p(vx)
					vz = df_fit.z.mean()
				else:
					vz = p(vx)
					vy = df_fit.y.mean()
			elif fit_dim[0] == 'y':
				vy = df_fit.y.mean()
				if fit_dim[1] == 'x':
					vx = p(vy)
					vz = df_fit.z.mean()
				else:
					vz = p(vy)
					vx = df_fit.x.mean()
			elif fit_dim[0] == 'z':
				vz = df_fit.z.mean()
				if fit_dim[1] == 'x':
					vx = p(vz)
					vy = df_fit.y.mean()
				else:
					vy = p(vz)
					vx = df_fit.x.mean()
			self.vertex = [vx,vy,vz]
		else:
			self.vertex = vertex

		#Translate data so that the vertex is at the origin
		self.df_align = pd.DataFrame({
			'x': df_fit.x - self.vertex[0],
			'y': df_fit.y - self.vertex[1],
			'z': df_fit.z - self.vertex[2]
			})

		#Rotate data by 180 degrees if necessary
		if flip == None or flip == False:
			#Calculate model
			model = np.polyfit(df_fit[fit_dim[0]],df_fit[fit_dim[1]],deg=deg)
			p = np.poly1d(model)

			#If a is less than 0, rotate data
			if model[0] < 0:
				self.df_align = self.flip_data(self.df_align)
		elif flip == True:
			self.df_align = self.flip_data(self.df_align)

		#Calculate final math model
		if mm == None:
			self.mm = self.fit_model(self.df_align,deg,fit_dim)
		else:
			self.mm = mm

	def flip_data(self,df):
		'''
		Rotate data by 180 degrees

		:param dataframe df: Pandas dataframe containing x,y,z data
		:returns: Rotated dataframe
		'''

		r = np.array([[np.cos(np.pi),0,np.sin(np.pi)],
			[0,1,0],
			[-np.sin(np.pi),0,np.cos(np.pi)]])

		rot = np.dot(np.array(df),r)

		dfr = pd.DataFrame({'x':rot[:,0],'y':rot[:,1],'z':rot[:,2]})
		return(dfr)

	def fit_model(self,df,deg,fit_dim):
		'''Fit model to dataframe

		:param pd.DataFrame df: Dataframe containing at least x,y,z
		:param int deg: Degree of the function that should be fit to the model
		:param array fit_dim: Array of length two containing two strings describing the first and second axis for fitting the model, e.g. ['x','z']
		:returns: math model
		:rtype: :py:class:`math_model`
		'''

		mm = math_model(np.polyfit(df[fit_dim[0]], df[fit_dim[1]], deg=deg))
		return(mm)

	###### Functions associated with alpha, r, theta coordinate system ######

	def find_distance(self,t,point):
		'''
		Find euclidean distance between math model(t) and data point in the xy plane

		:param float t: float value defining point on the line
		:param array point: array [x,y] defining data point
		:returns: distance between the two points
		:rtype: float
		'''

		x = float(t)
		z = self.mm.p(x)

		#Calculate distance between two points passed as array
		dist = np.linalg.norm(point - np.array([x,z]))

		return(dist)

	def find_min_distance(self,row):
		'''
		Find the point on the curve that produces the minimum distance between the point and the data point using scipy.optimize.minimize(:py:func:`brain.find_distance`)

		:param pd.Series row: row from dataframe in the form of a pandas Series
		:returns: point in the curve (xc, yc, zc) and r
		:rtype: floats
		'''

		dpoint = np.array([row.x,row.z])

		#Use scipy.optimize.minimize to find minimum solution of brain.find_distance,
		#dpoint[0] is starting guess for x value
		result = minimize(self.find_distance, dpoint[0], args=(dpoint))

		x = result['x'][0]
		y = 0
		z = self.mm.p(x)
		#r = result['fun']

		return(x,y,z)
		#return(pd.Series({'xc':x, 'yc':y, 'zc':z, 'r':r}))

	def integrand(self,x):
		'''
		Function to integrate to calculate arclength

		:param float x: integer value for x
		:returns: arclength value for integrating
		:rtype: float
		'''

		y_prime = self.mm.cf[0]*2*x + self.mm.cf[1]

		arclength = np.sqrt(1 + y_prime**2)
		return(arclength)

	def find_arclength(self,xc):
		'''
		Calculate arclength by integrating the derivative of the math model in xy plane

		.. math::

			\int_{vertex}^{point} \sqrt{1 + (2ax + b)^2}

		:param float row: Postion in the x axis along the curve
		:returns: Length of the arc along the curve between the row and the vertex
		:rtype: float
		'''

		ac,err = scipy.integrate.quad(self.integrand,xc,0)
		return(ac)

	def find_theta(self,row,zc,yc):
		'''
		Calculate theta for a row containing data point in relationship to the xz plane

		:param pd.Series row: row from dataframe in the form of a pandas Series
		:param float yc: Y position of the closest point in the curve to the data point
		:param float zc: Z position of the closest point in the curve to the data point
		:returns: theta, angle between point and the model plane
		:rtype: float
		'''

		theta = np.arctan2(row.y-yc,row.z-zc)
		return(theta)

	def find_r(self,row,zc,yc,xc):
		'''
		Calculate r using the Pythagorean theorem

		:param pd.Series row: row from dataframe in the form of a pandas Series
		:param float yc: Y position of the closest point in the curve to the data point
		:param float zc: Z position of the closest point in the curve to the data point
		:param float xc: X position of hte closest point in the curve to the data point
		:returns: r, distance between the point and the model
		:rtype: float
		'''

		r = np.sqrt((row.z-zc)**2 + (row.y-yc)**2 + (row.x-xc)**2)
		return(r)

	def calc_coord(self,row):
		'''
		Calculate alpah, r, theta for a particular row

		:param pd.Series row: row from dataframe in the form of a pandas Series
		:returns: pd.Series populated with coordinate of closest point on the math model, r, theta, and ac (arclength)
		'''

		xc,yc,zc = self.find_min_distance(row)
		ac = self.find_arclength(xc)
		theta = self.find_theta(row,zc,yc)
		r = self.find_r(row,zc,yc,xc)

		return(pd.Series({'x':row.x,'y':row.y,'z':row.z,'xc':xc, 'yc':yc, 'zc':zc,
					'r':r, 'ac':ac, 'theta':theta}))

	def transform_coordinates(self):
		'''
		Transform coordinate system so that each point is defined relative to math model by (alpha,theta,r) (only applied to :py:attr:`brain.df_align`)

		:returns: appends columns r, xc, yc, zc, ac, theta to :py:attr:`brain.df_align`
		'''

		#Calculate alpha, theta, r for each row in dataset
		self.df_align = self.df_align.merge(self.df_align.apply((lambda row: self.calc_coord(row)), axis=1))

	def subset_data(self,df,sample_frac=0.5):
		'''
		Takes a random sample of the data based on the value between 0 and 1 defined for sample_frac

		Creates the variable :py:attr:`brain.subset`

		:param pd.DataFrame: Dataframe which will be sampled
		:param float sample_frac: (or None) Value between 0 and 1 specifying proportion of the dataset that should be randomly sampled for plotting

		.. py:attribute:: brain.subset

			Random sample of the input dataframe
		'''

		self.subset = df.sample(frac=sample_frac)

	def add_thresh_df(self,df):
		'''
		Adds dataframe of thresholded and transformed data to :py:attr:`brain.df_thresh`

		:param pd.DataFrame df: dataframe of thesholded and transformed data
		:returns: :py:attr:`brain.df_thresh`
		'''

		self.df_thresh = df

	def add_aligned_df(self,df):
		'''
		Adds dataframe of aligned data

		.. warning:: Calculates model, but assumes that the dimensions of the fit are x and z

		:param pd.DataFrame df: Dataframe of aligned data
		:returns: :py:attr:`brain.df_align`
		'''

		self.df_align = df
		self.mm = self.fit_model(self.df_align,2,['x','z'])

class embryo:
	'''
	Class to managed multiple brain objects in a multichannel sample

	:param str name: Name of this sample set
	:param str number: Sample number corresponding to this embryo
	:param str outdir: Path to directory for output files

	.. py:attribute:: embryo.chnls

		Dictionary containing the :py:class:`brain` object for each channel

	.. py:attribute:: embryo.outdir

		Path to directory for output files

	.. py:attribute:: embryo.name

		Name of this sample set

	.. py:attribute:: embryo.number

		Sample number corresponding to this embryo
	'''

	def __init__(self,name,number,outdir):
		'''Initialize embryo object'''

		self.chnls = {}
		self.outdir = outdir
		self.name = name
		self.number = number

	def add_channel(self,filepath,key):
		'''
		Add channel to :py:attr:`embryo.chnls` dictionary

		:param str filepath: Complete filepath to image
		:param str key: Name of the channel
		'''

		s = brain()
		s.read_data(filepath)

		self.chnls[key] = s

	def process_channels(self,mthresh,gthresh,radius,scale,microns,deg,primary_key,comp_order,fit_dim):
		'''
		Process all channels through the production of the :py:attr:`brain.df_align` dataframe

		:param float mthresh: Value between 0 and 1 to use as a cutoff for minimum pixel value for median data
		:param float gthresh: Value between 0 and 1 to use as a cutoff for minimum pixel value for general data
		:param int radius: Size of the neighborhood area to examine with median filter
		:param array scale: Array with three values representing the constant by which to multiply x,y,z respectively
		:param array microns: Array with three values representing the x,y,z micron dimensions of the voxel
		:param int deg: Degree of the function that should be fit to the model
		:param str primary_key: Key for the primary structural channel which PCA and the model should be fit too
		:param array comp_order: Array specifies the assignment of components to x,y,z. Form [x component index, y component index, z component index], e.g. [0,2,1]
		:param array fit_dim: Array of length two containing two strings describing the first and second axis for fitting the model, e.g. ['x','z']
		'''

		#Process primary channel
		self.chnls[primary_key].preprocess_data(gthresh,scale,microns)

		self.chnls[primary_key].calculate_pca_median(self.chnls[primary_key].raw_data,
			mthresh,radius,microns)
		self.pca = self.chnls[primary_key].pcamed

		self.chnls[primary_key].align_data(self.chnls[primary_key].df_thresh,
			self.pca,comp_order,fit_dim,deg=deg)
		self.mm = self.chnls[primary_key].mm
		self.vertex = self.chnls[primary_key].vertex

		self.chnls[primary_key].transform_coordinates()

		print('Primary channel',primary_key,'processing complete')

		for ch in self.chnls.keys():
			if ch != primary_key:
				self.chnls[ch].preprocess_data(gthresh,scale,microns)

				self.chnls[ch].align_data(self.chnls[ch].df_thresh,
					self.pca,comp_order,fit_dim,deg=deg,
					mm = self.mm, vertex = self.vertex)

				self.chnls[ch].transform_coordinates()
				print(ch,'processed')

	def save_projections(self,subset):
		'''
		Save projections of both channels into png files in :py:attr:`embryo.outdir` following the naming scheme [:py:attr:`embryo.name`]_[:py:attr:`embryo.number`]_[`channel name`]_MIP.png

		:param float subset: Value between 0 and 1 to specify the fraction of the data to randomly sample for plotting
		'''

		for ch in self.chnls.keys():
			fig = self.chnls[ch].plot_projections(self.chnls[ch].df_align,subset)
			fig.savefig(os.path.join(self.outdir,
				self.name+'_'+self.number+'_'+ch+'_MIP.png'))

		print('Projections generated')

	def save_psi(self):
		'''
		Save all channels into psi files following the naming scheme [:py:attr:`embryo.name`]_[:py:attr:`embryo.number`]_[`channel name`].psi
		'''

		columns = ['x','y','z','ac','r','theta']

		for ch in self.chnls.keys():
			write_data(os.path.join(self.outdir,
				self.name+'_'+str(self.number)+'_'+ch+'.psi'),
				self.chnls[ch].df_align[columns])

		print('PSIs generated')

	def add_psi_data(self,filepath,key):
		'''
		Read psi data into a channel dataframe

		:param str filepath: Complete filepath to data
		:param str key: Descriptive key for channel dataframe in dictionary
		'''

		self.chnls[key] = read_psi(filepath)

class math_model:
	'''
	Object to contain attributes associated with the math model of a sample

	:param array model: Array of coefficients calculated by np.polyfit

	.. py:attribute:: math_model.cf

		Array of coefficients for the math model

	.. py:attribute:: math_model.p

		Poly1d function for the math model to allow calculation and plotting of the model
	'''

	def __init__(self,model):

		self.cf = model
		self.p = np.poly1d(model)

####### Alignment correction functions#########

def find_anchors(df,dim):
	''':param str dim: either y or z'''
	mm = math_model(np.polyfit(df['x'],df[dim],2))

	pt1 = {}
	if mm.cf[0] > 0:
		pt1[dim] = np.max(df[dim])
	else:
		pt1[dim] = np.min(df[dim])
	pt1['x'] = df[df[dim] == pt1[dim]].x.values[0]

	pt2 = {}
	if abs(pt1['x']-np.min(df.x)) > abs(pt1['x']-np.max(df.x)):
		pt2['x'] = np.min(df.x)
	else:
		pt2['x'] = np.max(df.x)
	pt2[dim] = df[df.x == pt2['x']][dim].values[0]
	return(pt1,pt2)

def calc_rotation(pts,d):
	dx = (pts.iloc[0].x - pts.iloc[1].x)/2
	dy = (pts.iloc[0][d] - pts.iloc[1][d])/2

	mp = [pts.iloc[1].x + dx, pts.iloc[1][d] + dy]

	phi = np.arctan2(dy,dx)
	if d == 'y':
			A = np.array([[np.cos(phi),-np.sin(phi),0],
						[np.sin(phi),np.cos(phi),0],
						[0,0,1]])
	else:
		A = np.array([[np.cos(phi),0,-np.sin(phi)],
						[0,1,0],
						[np.sin(phi),0,np.cos(phi)]])

	return(mp,A)

def cant_correction(df,d):
    pts = find_anchors(df,d)
    mp,A = calc_rotation(pts,d)
    rot = np.dot(np.array(df[['x','y','z']]),A)
    rdf = pd.DataFrame({'x':rot[:,0],'y':rot[:,1],'z':rot[:,2]})
    return(rdf)

class landmarks:
	'''
	Class to handle calculation of landmarks to describe structural data

	:param list percbins: (or None) Must be a list of integers between 0 and 100
	:param int rnull: (or None) When the r value cannot be calculated it will be set to this value

	.. py:attribute:: brain.lm_wt_rf

		pd.DataFrame, which wildtype landmarks will be added to

	.. py:attribute:: brain.lm_mt_rf

		pd.DataFrame, which mutant landmarks will be added to

	.. py:attribute:: brain.rnull

		Integer specifying the value which null landmark calculations will be set to

	.. py:attribute:: brain.percbins

		Integer specifying the percentiles which will be used to calculate landmarks
	'''

	def __init__(self,percbins=[10,50,90],rnull=15):

		self.lm_wt_rf = pd.DataFrame()
		self.lm_mt_rf = pd.DataFrame()

		self.rnull = rnull
		self.percbins = percbins

	def calc_bins(self,Ldf,ac_num,tstep):
		'''
		Calculates alpha and theta bins based on ac_num and tstep

		Creates :py:attr:`landmarks.acbins` and :py:attr:`landmarks.tbins`

		.. warning:: `tstep` does not handle scenarios where 2pi is not evenly divisible by tstep

		:param dict Ldf: Dict dataframes that are being used for the analysis
		:param int ac_num: Integer indicating the number of divisions that should be made along alpha
		:param float tstep: The size of each bin used for alpha

		.. py:attribute:: landmarks.acbins

			List containing the boundaries of each bin along alpha based on `ac_num`

		.. py:attribute:: landmarks.tbins

			List containing the boundaries of each bin along theta based on `tstep`
		'''

		Ldf = Ldf.values()

		#Find the minimum and maximum values of alpha in the dataset
		acmin,acmax = 0,0
		for df in Ldf:
			if df.ac.min() < acmin:
				acmin = df.ac.min()
			if df.ac.max() > acmax:
				acmax = df.ac.max()

		#Correct min and max values so that arclength is equal on each side
		if abs(acmin) > acmax:
			self.acbins = np.linspace(acmin,abs(acmin),ac_num)
		else:
			self.acbins = np.linspace(-acmax,acmax,ac_num)

		#Calculate tbins divisions based on tstep
		self.tbins = np.arange(-np.pi,np.pi+tstep,tstep)

	def calc_perc(self,df,snum,dtype,out):
		'''
		Calculate landmarks for a dataframe based on the bins and percentiles that have been previously defined

		:param pd.DataFrame df: Dataframe containing columns x,y,z,alpha,r,theta
		:param str snum: String containing a sample identifier that can be converted to an integer
		:param str dtype: String describing the sample group to which the sample belongs, e.g. control or experimental
		:returns: pd.DataFrame with new landmarks appended
		'''

		D = {'stype':dtype}

		#Go through each arclength bins
		for a in range(len(self.acbins)):
			if a+1 < len(self.acbins):
				arange = [self.acbins[a],self.acbins[a+1]]

				#Go through each theta bin
				for t in range(len(self.tbins)):
					if t+1 < len(self.tbins):
						trange = [self.tbins[t],self.tbins[t+1]]

						#Go through each percentile
						for p in self.percbins:
							d = df[(df.ac > arange[0]) & (df.ac < arange[1])]
							d = d[(d.theta > trange[0]) & (d.theta < trange[1])]

							#Try to calculate percentile, but set to null if it fails
							try:
								r = np.percentile(d.r,p)
								pts = d[d.r < r].count()['i']
							except:
								r = self.rnull
								pts = 0

							#Create name of column based on bins
							L = []
							for s in [arange[0],arange[1],trange[0],trange[1]]:
								L.append(str(np.around(s,decimals=2)))
							name = '_'.join(L)

							D[name+'_'+str(p)+'_pts'] = pts
							D[name+'_'+str(p)+'_r'] = r

		out = out.append(pd.Series(D,name=int(snum)))
		return(out)

	def calc_wt_reformat(self,df,snum):
		'''
		.. warning:: Deprecated function, but includes code pertaining to calculating point based data
		'''

		D = {'stype':'wildtype'}

		for a in range(len(self.acbins)):
			if a+1 < len(self.acbins):
				arange = [self.acbins[a],self.acbins[a+1]]

				for t in range(len(self.tbins)):
					if t+1 < len(self.tbins):
						trange = [self.tbins[t],self.tbins[t+1]]

						for p in self.percbins:
							d = df[(df.ac > arange[0]) & (df.ac < arange[1])]
							d = d[(d.theta > trange[0]) & (d.theta < trange[1])]

							try:
								r = np.percentile(d.r,p)
								pts = d[d.r < r].count()['i']
							except:
								r = self.rnull
								pts = 0

							L = []
							for s in [arange[0],arange[1],trange[0],trange[1]]:
								L.append(str(np.around(s,decimals=2)))
							name = '_'.join(L)

							D[name+'_'+str(p)+'_pts-pts'] = pts
							D[name+'_'+str(p)+'_perc-pts'] = pts
							D[name+'_'+str(p)+'_pts-r'] = r
							D[name+'_'+str(p)+'_perc-r'] = r

		self.lm_wt_rf = self.lm_wt_rf.append(pd.Series(D,name=int(snum)))

	def calc_mt_landmarks(self,df,snum,wt):
		'''
		.. warning:: Deprecated function, but attempted to calculate mutant landmarks based on the number of points found in the wildtype standard
		'''

		D = {'stype':'mutant'}

		for c in wt.columns:
			if len(c.split('_')) == 6:
				amn,amx,tmn,tmx,p,dtype = c.split('_')
				p = int(p)

				# print(c)
				# print(df.count()['i'])
				d = df[(df.ac > float(amn))&(df.ac < float(amx))]
				# print(d.count()['i'])
				d = d[(d.theta > float(tmn))&(d.theta < float(tmx))]
				# print(d.count()['i'])

				if dtype.split('-')[0] == 'perc':
					try:
						perc_r = np.percentile(d.r,p)
						perc_pts = d[d.r < perc_r]
					except:
						perc_r = self.rnull
						perc_pts = 0

					if dtype.split('-')[1] == 'r':
						D[c] = perc_r
					else:
						D[c] = perc_pts

				else:
					if dtype.split('-')[1] == 'pts':
						#Calculate precent of points by dividing pts by total pts
						p = wt[c].mean()/d.count()['i']
						try:
							pt_r = np.percentile(d.r,p)
						except:
							pt_r = self.rnull
						D[c] = pt_r
					else:
						D[c] = np.nan

		self.lm_mt_rf = self.lm_mt_rf.append(pd.Series(D,name=int(snum)))

def reformat_to_cart(df):
	'''
	Take a dataframe in which columns contain the bin parameters and convert to a cartesian coordinate system

	:param pd.DataFrame df: Dataframe containing columns with string names that contain the bin parameter
	:returns: pd.DataFrame with each landmark as a row and columns: x,y,z,r,r_std,t,pts
	'''

	ndf = pd.DataFrame()
	for c in df.columns:
		if len(c.split('_')) == 6:
			amn,amx,tmn,tmx,p,dtype = c.split('_')
			x = np.mean([float(amn),float(amx)])
			t = np.mean([float(tmn),float(tmx)])

			if dtype == 'r':
				r = np.mean(df[c])
				r_std = stats.sem(df[c])
				y = np.sin(t)*r
				z = np.cos(t)*r

				pts = np.mean(df['_'.join([amn,amx,tmn,tmx,p,'pts'])])

				D = pd.Series({'x':x,'y':y,'z':z,'r':r,'r_sem':r_std,'t':t,'pts':pts})

				ndf = ndf.append(pd.Series(D),ignore_index=True)

	return(ndf)

def convert_to_arr(xarr,tarr,DT,mdf,Ldf=[]):
	'''
	Convert a pandas dataframe containing landmarks as columns and samples as rows into a 3D numpy array

	The columns of `mdf` determine which landmarks will be saved into the array. Any additional dataframes that need to be converted can be included in Ldf

	:param np.array xarr: Array containing all unique x values of landmarks in the dataset
	:param np.array tarr: Array containing all unique t values of landmarks in the dataset
	:param str DT: Either ``r`` or ``pts`` indicating which data type should be saved to the array
	:param pd.DataFrame mdf: Main landmark dataframe containing landmarks as columns and samples as rows
	:param list Ldf: List of additional pd.DataFrames that should also be converted to arrays
	:returns: Array of the main dataframe and list of arrays converted from Ldf
	'''
	marr = np.zeros((len(xarr),len(tarr),len(mdf.index)))
	xarr = np.round(xarr,2)
	tarr = np.round(tarr,2)
	Larr = []
	for df in Ldf:
		Larr.append(np.zeros((len(xarr),len(tarr),len(df.index))))

	for c in mdf.columns:
		if len(c.split('_')) == 6:
			amn,amx,tmn,tmx,p,dtype = c.split('_')
			x = float(amn)#np.mean([float(amn),float(amx)])
			t = float(tmn) #np.mean([float(tmn),float(tmx)])

			if dtype == DT:
				marr[np.where(xarr==x)[0],np.where(tarr==t)[0]] = mdf[c]
				for arr,df in zip(Larr,Ldf):
					arr[np.where(xarr==x)[0],np.where(tarr==t)[0]] = df[c]

	return(marr,Larr)

def calc_variance(anum,dfs):
	'''
	Calculate the variance between samples according to bin position and variance between adjacent bins

	:param int anum: Number of bins which the arclength axis should be divided into
	:param dict dfs: Dictionary of dfs which are going to be processed
	:returns: Two arrays: svar (anum,tnum) and bvar (anum*tnum,snum)
	:rtype: np.array
	'''

	#Set up bins
	lm = landmarks(percbins=[50],rnull=15)
	lm.calc_bins(dfs.values(),anum,np.pi/4)

	 #Calculate landmarks
	outlm = pd.DataFrame()
	for k in dfs.keys():
		outlm = lm.calc_perc(dfs[k],k,'wt',outlm)

	#Convert to arr for variance calculation
	lmarr,arr = convert_to_arr(lm.acbins,lm.tbins,outlm)

	svar = np.var(lmarr,axis=2)

	Lvar = []
	for i in range(1,len(lm.acbins)-2):
		for t in range(0,len(lm.tbins)):
			#Save the variance of a particular bin and adjacent neighbors across a set of samples
			Lvar.append(np.var(lmarr[i-1:i+2,t],axis=0))

	return(svar,np.array(Lvar))

class anumSelect:

	def __init__(self,dfs):
		'''
		A class that assists in selecting the optimum value of anum

		:param dict dfs: Dictionary of pd.DataFrames with samples to use for optimization

		.. attribute:: anumSelect.dfs

			Dictionary of dataframes that will be used for the parameter sweep

		.. attribute:: anumSelect.Lsv

			List of sample variance arrays for each anum in the sweep

		.. attribute:: anumSelect.Lbv

			List of bin variance arrays for each anum in the sweep

		.. attribute:: anumSelect.Msv

			List of values of the average sample variance for each anum in the sweep

		.. attribute:: anumSelect.Mbv

			List of values of the average bin variance for each anum in the sweep

		.. attribute:: anumSelect.Llm

			List of landmark arrays for each anum in the sweep

		'''

		self.dfs = dfs
		self.Lsv,self.Lbv = [],[]
		self.Msv,self.Mbv = [],[]
		self.Llm = []

	def calc_variance(self,anum,tstep,percbins,rnull,DT):
		'''
		Calculate the variance between samples according to bin position and variance between adjacent bins

		:param int anum: Number of bins which the arclength axis should be divided into
		:param float tstep: The size of each bin used for alpha
		:param dict dfs: Dictionary of dfs which are going to be processed
		:param list percbins: Must be a list of integers between 0 and 100
		:param int rnull: When the r value cannot be calculated it will be set to this value
		:param str DT: Data type for which variance is measured, e.g. ``r`` or ``pts``
		'''

		#Set up bins
		lm = landmarks(percbins=percbins,rnull=rnull)
		lm.calc_bins(self.dfs,anum,tstep)

		#Calculate landmarks
		outlm = pd.DataFrame()
		for k in self.dfs.keys():
			outlm = lm.calc_perc(self.dfs[k],k,'s',outlm)

		#Convert to arr for variance calculations
		lmarr,arr = convert_to_arr(lm.acbins,lm.tbins,DT,outlm)
		self.Llm.append(lmarr)

		#Calculate variance between samples
		svar = np.nanvar(lmarr,axis=2)

		#Calculate variance between bins
		Lvar = []
		for i in range(1,len(lm.acbins-2)):
			for t in range(0,len(lm.tbins)):
				Lvar.append(np.nanvar(lmarr[i-1:i+2,t],axis=0))

		self.Lsv.append(svar)
		self.Msv.append(np.nanmean(svar))
		self.Lbv.append(np.array(Lvar))
		self.Mbv.append(np.nanmean(np.array(Lvar)))

		print(anum,'calculation complete')

	def param_sweep(self,tstep,amn=2,amx=50,astep=1,percbins=[50],rnull=15,DT='pts'):
		'''
		Calculate landmarks for each value of anum specified in input range

		:param float tstep: The size of each theta wedge in radians
		:param int amn: The minimum number of alpha bins that should be considered
		:param int amx: The maximum number of alpha bins that should be considered
		:param int astep: The step size in the range of amn to amx
		:param list percbins: (or None) Must be a list of integers between 0 and 100
		:param int rnull: (or None) When the r value cannot be calculated it will be set to this value
		:param str DT: Default=``pts`` Data type for which variance is measured, e.g. ``r`` or ``pts``

		.. attribute:: anumSelect.amn

			User defined minimum number of alpha bins considered in sweep

		.. attribute:: anumSelect.amx

			User defined maximum number of alpha bins considered in sweep

		.. attribute:: anumSelect.astep

			The step size in the range of amn to amx
		'''
		self.amn,self.amx,self.astep = amn,amx,astep

		for a in np.arange(self.amn,self.amx,self.astep):
			self.calc_variance(a,tstep,percbins,rnull,DT)

		print('Parameter sweep complete')

	def plot_rawdata(self):
		'''
		Plot raw data from parameter sweep
		'''
		x = np.arange(self.amn,self.amx,self.astep)
		fig,ax = plt.subplots()

		ax.plot(x,self.Mbv[2:],c='b',ls='--',label='Raw Bin Variance')
		ax.plot(x,self.Msv[2:],c='g',ls='--',label='Raw Sample Variance')

		ax.legend()
		ax.set_xlabel('Number of Alpha Bins')
		ax.set_ylabel('Relative Variance')

	def plot_fitted(self,dof):
		'''
		Plot polynomial fits over the raw data

		:param int dof: Degree of the polynomial function to be fit
		'''

		x = np.arange(self.amn,self.amx,self.astep)
		fig,ax = plt.subplots()

		ax.plot(x,self.Mbv[2:],c='b',ls='--',label='Raw Bin Variance')
		ax.plot(x,self.Msv[2:],c='g',ls='--',label='Raw Sample Variance')

		pbv = np.polyfit(x,self.Mbv[2:],dof)
		fbv = np.poly1d(pbv)
		ax.plot(x,fbv(x),c='b',label='Fit Bin Variance')

		psv = np.polyfit(x,self.Msv[2:],dof)
		fsv = np.poly1d(psv)
		ax.plot(x,fsv(x),c='g',label='Fit Sample Variance')

		ax.legend()
		ax.set_xlabel('Number of Alpha Bins')
		ax.set_ylabel('Relative Variance')

	def find_optimum_anum(self,dof,guess):
		'''
		Calculate optimum anum and plot

		:param int dof: Degree of the polynomial function to be fit
		:param int guess: Best guess of the optimum anum, which cannot be less than the maximum bin variance
		'''

		x = np.arange(self.amn,self.amx,self.astep)
		fig,ax = plt.subplots()

		pbv = np.polyfit(x,normalize(self.Mbv[2:])[0],dof)
		fbv = np.poly1d(pbv)
		ax.plot(x,fbv(x),c='b',label='Bin Variance')

		psv = np.polyfit(x,normalize(self.Msv[2:])[0],dof)
		fsv = np.poly1d(psv)
		ax.plot(x,fsv(x),c='g',label='Sample Variance')

		opt = minimize(fbv+fsv,guess)
		ax.axvline(opt.x,c='r',label='Optimum: '+str(np.round(opt.x[0],2)))

		ax.legend()
		ax.set_xlabel('Number of Alpha Bins')
		ax.set_ylabel('Relative Variance')

class graphSet:

	def __init__(self,tpairs,xarr,tarr,sharey=True):
		'''
		This object manages subobjects containing sample data, :class:`graphData`,
		and generates the appropriate landmark graph

		:param np.array tpairs: A list that specifies which theta bins should be paired together for graphing
		:param np.array xarr: An array listing the bin division points along alpha
		:param np.array tarr: Array listing the bin division points along theta

		.. attribute:: graphSet.tpairs

			A list that specifies which theta bins should be paired together for graphing

		.. attribute:: graphSet.xarr

			An array listing the bin division points along alpha

		.. attribute:: graphSet.tarr

			Array listing the bin division points along theta
		'''

		self.tpairs,self.xarr,self.tarr = tpairs,xarr,tarr
		self.sharey = sharey

		self.Ls,self.Lc = [],[]
		self.Ds, self.Dc = {},{}

	def add_data(self,gD,stype,ctype,dtype):
		'''
		Add :class:`graphData` object with descriptive tags of stype,ctype,color

		:param obj gD: graphData object initialized for one sample set
		:param str stype: String describing the sample type
		:param str ctype: String describing channel type
		:param str dtype: Either ``pts`` or ``r`` specifies which type of landmark data will be used
		'''
		gD.prepare_data(self.xarr,self.tarr,dtype)

		self.add_to_dict(self.Ds,stype,ctype,gD)
		self.add_to_dict(self.Dc,ctype,stype,gD)

		#self.Ds.setdefault(stype).append({ctype:gD})
		#self.Dc.setdefault(ctype).append({stype:gD})
		self.Ls.append(stype)
		self.Lc.append(ctype)

	def add_to_dict(self,D,k1,k2,item):
		'''Add :class:`graphData` object to a dictionary checking for exisiting keys

		:param dict D: the dictionary which the data will be added to
		:param str k1: Key for the first index into the Dictionary
		:param str k2: Key for the second internal dictionary
		:param obj item: :class:`graphData` object
		'''

		if k1 in D.keys():
			D[k1][k2] = item
		else:
			D[k1] = {k2:item}

	def start_figure(self,figsize=(10,8)):

		self.LsUn = np.unique(self.Ls)
		self.LcUn = np.unique(self.Lc)

		self.fig,self.axr = plt.subplots(4,len(self.LsUn),figsize=figsize,sharex=True,sharey=self.sharey)

	def add_plot(self,go,c,s,j,a=0.3):

		# go = self.Dc[c][s]

		for i,p in enumerate(self.tpairs):
			ti1 = np.where(self.tarr==p[0])[0][0]
			ti2 = np.where(self.tarr==p[1])[0][0]

			self.axr[i,j].fill_between(self.xarr,go.avg[:,ti1]+go.sem[:,ti1],go.avg[:,ti1]-go.sem[:,ti1],alpha=a,color=go.c,zorder=1)
			self.axr[i,j].fill_between(self.xarr,-go.avg[:,ti2]+go.sem[:,ti2],-go.avg[:,ti2]-go.sem[:,ti2],alpha=a,color=go.c,zorder=1)

			self.axr[i,j].plot(self.xarr,go.avg[:,ti1],c=go.c,zorder=2,label=c+s)
			self.axr[i,j].plot(self.xarr,-go.avg[:,ti2],c=go.c,zorder=2)

	def add_legends(self,I,J):

		for i in range(I):
			for j in range(J):
				self.axr[i,j].legend()

	def make_figure(self,a,mt_key,figsize=(10,8),P=True,pthresh=None,cbar=False):
		'''
		Creates a figure showing four theta slices and as many columns as ctypes

		.. todo:: P value scatter plot is broken

		.. todo:: Control p val for multiple testing

		:param float a: Alpha value for fill_between ribbons
		:param str mt_key: String indicating the data type of the experimental group
		:param tuple figsize: Tuple specifying the height and width of the figure
		:param bool P: True if pvalue should be plotted
		:param float pthresh: None or float value for p-value threshold
		:param bool cbar: Default false, if true plots colorbar for p values

		.. attribute:: graphSet.fig

			Figure object created by :func:`graphSet.make_figure`

		.. attribute:: graphSet.axr

			Subplot axis array created by :func:`graphSet.make_figure`
		'''

		self.LsUn = np.unique(self.Ls)
		self.LcUn = np.unique(self.Lc)

		self.fig,self.axr = plt.subplots(4,len(self.LsUn),figsize=figsize,sharey=True,sharex=True)

		for j,c in enumerate(self.LcUn):
			dc = self.Dc[c]
			parr = stats.ttest_ind(dc[self.LsUn[0]].arr_masked,dc[self.LsUn[1]].arr_masked,axis=2,nan_policy='omit')[1]

			if pthresh != None:
				bhthresh = pthresh/(len(self.xarr)*len(self.tarr))
				parr[parr<bhthresh] = 0
				parr[parr>bhthresh] = 1
				print(bhthresh)

			for i,p in enumerate(self.tpairs):
				for s in self.LsUn:
					go = dc[s]

					ti1 = np.where(self.tarr==p[0])[0][0]
					ti2 = np.where(self.tarr==p[1])[0][0]

					self.axr[i,j].fill_between(self.xarr,go.avg[:,ti1]+go.sem[:,ti1],go.avg[:,ti1]-go.sem[:,ti1],alpha=a,color=go.c,zorder=1)
					self.axr[i,j].fill_between(self.xarr,-go.avg[:,ti2]+go.sem[:,ti2],-go.avg[:,ti2]-go.sem[:,ti2],alpha=a,color=go.c,zorder=1)

					self.axr[i,j].plot(self.xarr,go.avg[:,ti1],c=go.c,zorder=2,label=c+s)
					self.axr[i,j].plot(self.xarr,-go.avg[:,ti2],c=go.c,zorder=2)

					if (s == mt_key) & (P==True):
						cax = self.axr[i,j].scatter(self.xarr,go.avg[:,ti1],c=parr[:,ti1],cmap='Greys_r',zorder=3,vmin=0,vmax=1,edgecolors='k')
						self.axr[i,j].scatter(self.xarr,-go.avg[:,ti2],c=parr[:,ti2],cmap='Greys_r',zorder=3,vmin=0,vmax=1,edgecolors='k')
						# print('plot pval')
						if cbar==True:
							plt.colorbar(cax,ax=self.axr[i,j])

				self.axr[i,j].legend()

class graphData:

	def __init__(self,rawdf,color):
		'''
		Object to contain data and attributes needed for graphing landmark dataset

		:param pd.DataFrame raw: Landmark dataframe with each column as a landmarks
		:param str color: Hexcode or single letter color code

		.. attribute:: graphData.c

			Color code for this dataset

		.. attribute:: graphData.rawdf

			Dataframe containing each landmark as a column
		'''
		self.c = color
		self.rawdf = rawdf

	def prepare_data(self,xarr,tarr,dtype):
		'''
		Convert to array and calculate average and sem

		:param arr xarr: List of min and max borders of the alpha bins
		:param arr tarr: List of min and max borders of the theta bins
		:param str dtype: Either ``pts`` or ``r`` specifies which data to save to array

		.. attribute:: graphData.arr

			Array containing the landmark data for this particular sample

		.. attribute:: graphData.avg

			Average of :attr:`graphData.arr`

		.. attribute:: graphData.sem

			Standard error of the mean for :attr:`graphData.arr`
		'''
		self.arr,L = convert_to_arr(xarr,tarr,dtype,self.rawdf)

		nanmask = np.nanmean(self.arr,axis=2)[:,:,np.newaxis] + np.ones((1,1,self.arr.shape[-1]))
		nanmask = 2*nanmask

		self.arr_masked = self.arr
		self.arr_masked[np.isnan(self.arr)] = nanmask[np.isnan(self.arr)]

		self.avg = np.nanmean(self.arr_masked,axis=2)
		self.sem = stats.sem(self.arr_masked,axis=2,nan_policy='omit')

class treeClassifier:

	def __init__(self,df):
		'''A class to manage classifying a landmark dataset using a random forest classifier

		:param pd.DataFrame df: Landmark dataframe containing both sample types with different keys in the stype column

		.. attribute:: treeClassifier.df

			Landmark dataframe containing both sample types with different keys in the stype column
		'''

		self.df = df
		self.Y = self.df['stype']
		print('Sample types:',np.unique(self.Y))
		try:
			self.Xraw = self.df.drop(['stype','Unnamed: 0'],axis=1)
		except:
			self.Xraw = self.df.drop(['stype'],axis=1)

		self.Xnan = self.Xraw.dropna(axis=1,how='all').fillna(self.Xraw.mean())
		self.Xsc = scale(self.Xnan)

	def apply_pca(self,plot=False):
		'''Find optimal number of components and transform data

		:param bool plot: If true, a plot showing the pca parameter sweep will be generated

		.. attribute:: treeClassifier.Xtr

			:attr:`treeClassifier.Xsc` after transformation by PCA

		.. attribute:: treeClassifier.n

			The number of components that captures all of the variability in the data

		.. attribute:: treeClassifier.comp

			pd.DataFrame containing :attr:`treeClassifier.n` rows with the weight of each landmark in the column
		'''

		#Parameter sweep for optimum component number
		evar = []
		for i in range(1,100):
			tpca = PCA(n_components=i)
			tpca.fit(self.Xsc)
			evar.append(tpca.explained_variance_ratio_.sum())

		#Find first n that is close to equaling 1
		self.n = np.nonzero(np.array(evar)>=0.9999)[0][0] +1 #Accounts for list starting at 1
		print(str(self.n),'components accounts for 100 percent of the variability in the data')

		pca = PCA(n_components=self.n)
		pca.fit(self.Xsc)
		self.Xtr = pca.transform(self.Xsc)
		self.comp = pd.DataFrame(pca.components_,columns=self.Xnan.columns)

		if plot == True:
			fig,ax = plt.subplots()
			ax.plot(range(1,200),np.array(evar)*100)
			ax.set_ylabel('Explained Variance (%)')
			ax.set_xlabel('Number of Components')
			ax.axvline(self.n,c='r',label='N_components = '+str(n))
			ax.legend()

	def fit_classifier(self,max_features='sqrt',oob_score=True,n_estimators=500):
		'''Fit a random forest classifer to the transformed data to the trained data

		:param str max_features: Default=``'sqrt'``. See `sklearn.ensemble.RandomForestClassifier`_ for more info.
		:param bool oob_score: Default=``True``. See `sklearn.ensemble.RandomForestClassifier`_ for more info.
		:param int n_estimators: Default=``500``. See `sklearn.ensemble.RandomForestClassifier`_ for more info.

		.. attribute:: treeClassifier.rftree

			Random Forest Classifier generated by `sklearn.ensemble.RandomForestClassifier`_

		.. attributes:: treeClassifier.Importance

			pd.DataFrame containing the relative importance value for each component

		.. _sklearn.ensemble.RandomForestClassifier: <http://scikit-learn.org/stable/modules/generated/sklearn.ensemble.RandomForestClassifier.html
		'''
		self.rftree = RandomForestClassifier(max_features=max_features,
											oob_score=oob_score,
											n_estimators=n_estimators)
		self.rftree.fit(self.Xtr,self.Y)

		self.Importance = pd.DataFrame({'Importance':self.rftree.feature_importances_*100})
		self.Importance = self.Importance.sort_values(by='Importance',axis=0,ascending=False)

		print('Out of Bag score:',str(self.rftree.oob_score_))

	def print_top_components(self,index=None,thresh=None):
		'''Prints a list of top components, which can be limited by index or threshold

		:param int index: Limits how many components will be shown
		:param str thresh: Show only components above the minimum threshold set by thresh
		'''

		if type(index)==int:
			print(self.Importance[self.Importance.index < index])
		elif (type(thresh)==int) | (type(thresh)==str):
			print(self.Importance[self.Importance.Importance > thresh])
		else:
			print(self.Importance)

	def comp_to_arr(self):
		'''Converts dataframe of component landmarks to cartesian array

		.. attribute:: treeClassifier.xarr

			An array listing the bin division points along alpha

		.. attribute:: treeClassifier.tarr

			Array listing the bin division points along theta

		.. attribute:: treeClassifier.cParr

			A cartesian array of the dimensions [len(:attr:`treeClassifier.xarr`),len(:attr:`treeClassifier.tarr`),:attr:`treeClassifier.n`]
			containing the number of points per landmark

		.. attribute:: treeClassifier.cRarr

			A cartesian array of the dimensions [len(:attr:`treeClassifier.xarr`),len(:attr:`treeClassifier.tarr`),:attr:`treeClassifier.n`]
			containing the r distance of the percentile per landmark
		'''

		dfc = reformat_to_cart(self.comp)
		self.xarr = np.round(np.unique(dfc.x),2)
		self.tarr = np.round(np.unique(dfc.t),2)

		self.cParr = np.zeros((len(self.xarr),len(self.tarr),self.n))
		self.cRarr =  np.zeros((len(self.xarr),len(self.tarr),self.n))

		for c in self.comp.columns:
			if len(c.split('_')) == 6:
				amn,amx,tmn,tmx,p,dtype = c.split('_')
				x = np.mean([float(amn),float(amx)])
				t = np.mean([float(tmn),float(tmx)])

				if dtype=='r':
					self.cRarr[np.where(self.xarr==np.round(x,2))[0],np.where(self.tarr==np.round(t,2))[0]] = self.comp[c]
				elif dtype =='pts':
					self.cParr[np.where(self.xarr==np.round(x,2))[0],np.where(self.tarr==np.round(t,2))[0]] = self.comp[c]

	def plot_top_components(self,index=None,thresh=None,path=None):
		'''Plots the r heatmap and pts heatmap according to importance
		:param int index: Limits how many components will be shown
		:param str thresh: Show only components above the minimum threshold set by thresh
		:param str path: Optionally include a path to a directory to save images if desired
		'''
		if type(index)==int:
			L = self.Importance[self.Importance.index < index]
		elif (type(thresh)==int) | (type(thresh)==str):
			L = self.Importance[self.Importance.Importance > thresh]
		else:
			L = self.Importance

		if not hasattr(self,'cParr'):
			self.comp_to_arr()

		for i in L.index:
			fig,axr = plt.subplots(2,2,figsize=(15,4),gridspec_kw={'height_ratios':[1,0.05]})
			plt.suptitle('Importance = '+str(L.Importance.iloc[i]),fontsize=16)

			cb1 = axr[0,0].imshow(np.abs(self.cRarr[:,:,i].T),extent=[self.xarr[0],self.xarr[-1],self.tarr[0],self.tarr[-1]],aspect=10,cmap='viridis')
			cb2 = axr[0,1].imshow(np.abs(self.cParr[:,:,i].T),extent=[self.xarr[0],self.xarr[-1],self.tarr[0],self.tarr[-1]],aspect=10,cmap='viridis')

			axr[0,0].set_title(str(i)+'r')
			plt.colorbar(cb1,cax=axr[1,0],orientation='horizontal')
			axr[0,1].set_title(str(i)+'pts')
			plt.colorbar(cb2,cax=axr[1,1],orientation='horizontal')

			if path != None:
				if os.path.isdir(path):
					fig.savefig(os.path.join(path,'Imp_'+str(L.Importance.iloc[i])+'.jpg'))


P = {
	'zln':2,'zpt':3,'zfb':1,
	'wtc':'b','mtc':'r',
	'alpha':0.3,
	'cmap':'Greys_r',
	'xarr':None,
	'tarr':None
}

def subplot_lmk(ax,p,avg,sem,parr,xarr,tarr,dtype,Pn=P):
	'''
	Plot a ribbon of average and standard error of the mean onto the subplot, `ax`

	:param plt.Subplot ax: Matplotlib subplot onto which the data should be plotted
	:param list p: List of two theta values that should be plotted
	:param np.array avg: Array of shape (xvalues,tvalues) containing the average values of the data
	:param np.array sem: Array of shape (xvalues,tvalues) containing the standard error of the mean values of the data
	:param np.array parr: Array of shape (xvalues,tvalues) containing the p values for the data
	:param str dtype: String describing sample type
	:param Pn: Dictionary containing the following values: 'zln':2,'zpt':3,'zfb':1,'wtc':'b','mtc':'r','alpha':0.3,'cmap':'Greys_r'
	:type: dict or None
	'''

	for k in Pn.keys():
		P[k] = Pn[k]

	if dtype=='wt':
		c = P['wtc']
	else:
		c = P['mtc']

	ti1 = np.where(tarr==p[0])[0][0]
	ti2 = np.where(tarr==p[1])[0][0]

	ax.fill_between(xarr,avg[:,ti1]+sem[:,ti1],avg[:,ti1]-sem[:,ti1],alpha=P['alpha'],color=c,zorder=P['zfb'])
	ax.fill_between(xarr,-avg[:,ti2]+sem[:,ti2],-avg[:,ti2]-sem[:,ti2],alpha=P['alpha'],color=c,zorder=P['zfb'])

	ax.plot(xarr,avg[:,ti1],c=c,zorder=P['zln'])
	ax.plot(xarr,-avg[:,ti2],c=c,zorder=P['zln'])

	if dtype=='mt':
		ax.scatter(xarr,avg[:,ti1],c=parr[:,ti1],cmap=P['cmap'],zorder=P['zpt'])
		ax.scatter(xarr,-avg[:,ti2],c=parr[:,ti2],cmap=P['cmap'],zorder=P['zpt'])

##### PSI file processing ############

def write_header(f):
	'''
	Writes header for PSI file with columns Id,x,y,z,ac,r,theta

	:param file f: file object created by 'open(filename,'w')`
	'''

	contents = [
		'PSI Format 1.0',
		'',
		'column[0] = "Id"',
		'column[1] = "x"',
		'column[2] = "y"',
		'column[3] = "z"',
		'column[4] = "ac"',
		'symbol[4] = "A"',
		'type[4] = float',
		'column[5] = "r"',
		'symbol[5] = "R"',
		'type[5] = float',
		'column[6] = "theta"',
		'symbol[6] = "T"',
		'type[6] = float'
	]

	for line in contents:
		f.write('# '+ line + '\n')

def write_data(filepath,df):
	'''
	Writes data in PSI format to file after writing header using :py:func:`write_header`. Closes file at the conclusion of writing data.

	:param str filepath: Complete filepath to output file
	:param pd.DataFrame df: dataframe containing columns x,y,z,ac,r,theta
	'''

	#Open new file at given filepath
	f = open(filepath,'w')

	#Write header contents to file
	write_header(f)

	n = df.count()['x']

	#Write line with sample number
	f.write('\n'+str(n)+' 0 0\n')

	#Write translation matrix
	f.write('1 0 0\n'+
			'0 1 0\n'+
			'0 0 1\n\n')

	#Write dataframe to file using pandas to_csv function to format
	try:
		f.write(df[['x','y','z','ac','theta','r']].to_csv(sep=' ', index=True, header=False))
	except:
		f.write(df[['x','y','z']].to_csv(sep=' ', index=True, header=False))

	f.close()

	print('Write to',filepath,'complete')

def read_psi(filepath):
	'''
	Reads psi file at the given filepath and returns data in a pandas DataFrame

	:param str filepath: Complete filepath to file
	:returns: pd.Dataframe containing data
	'''

	df = pd.read_csv(filepath,
		sep=' ',
		header=19)

	if len(df.columns) <= 4:
		df.columns = ['i','x','y','z']
	elif len(df.columns) >= 6:
		df.columns=['i','x','y','z','ac','theta','r']

	return(df)

def read_psi_to_dict(directory,dtype):
	'''
	Read psis from directory into dictionary of dfs with filtering based on dtype

	:param str directory: Directory to get psis from
	:param str dtype: Usually 'AT' or 'ZRF1'
	:returns: Dictionary of pd.DataFrame
	'''

	dfs = {}
	for f in tqdm.tqdm(os.listdir(directory)):
		if (dtype in f)&('psi' in f):
			df = read_psi(os.path.join(directory,f))
			num = re.findall(r'\d+',f.split('.')[0])[0]
			dfs[num] = df

	return(dfs)

###### Stand alone functions

def process_sample(num,root,outdir,name,chs,prefixes,threshold,scale,deg,primary_key,comp_order,fit_dim,flip_dim):
	'''
	Process single sample through :py:class:`brain` class and saves df to csv

	.. warning:: Out of date and will probably fail

	:param str num: Sample number
	:param str root: Complete path to the root directory for this sample set
	:param str name: Name describing this sample set
	:param str outdir: Complete path to output directory
	:param array chs: Array containing strings specifying the directories for each channel
	:param array prefixes: Array containing strings specifying the file prefix for each channel
	:param float threshold: Value between 0 and 1 to use as a cutoff for minimum pixel value
	:param array scale: Array with three values representing the constant by which to multiply x,y,z respectively
	:param int deg: Degree of the function that should be fit to the model
	:param str primary_key: Key for the primary structural channel which PCA and the model should be fit too
	'''

	tic = time.time()

	print(num,'Starting',name)

	e = embryo(name,num,outdir)
	e.add_channel(os.path.join(root,chs[0],prefixes[0]+'_'+num+'_Probabilities.h5'),
		'at')
	e.add_channel(os.path.join(root,chs[1],prefixes[1]+'_'+num+'_Probabilities.h5'),'zrf1')
	e.process_channels(threshold,scale,deg,primary_key,[0,2,1],['x','z'],'z')

	print(num,'Data processing complete',name)

	e.save_projections(0.1)
	e.save_psi()

	toc = time.time()
	print(name,'complete',toc-tic)

def calculate_models(Ldf):
	'''
	Calculate model for each dataframe in list and add to new dataframe

	:param list Ldf: List of dataframes containing aligned data
	:returns: pd.Dataframe with a,b,c values for parabolic model
	'''

	modeldf = pd.DataFrame({'a':[],'b':[],'c':[]})

	for df in Ldf:
		s = cranium.brain()
		s.add_aligned_df(df)
		s.fit_model(s.df_align,2)

		modeldf = modeldf.append(pd.DataFrame({'a':[s.mm.cf[0]],'b':[s.mm.cf[1]],'c':[s.mm.cf[2]]}))

	return(modeldf)

def generate_kde(data,var,x,absv=False):
	'''
	Generate list of KDEs from either dictionary or list of data

	:param data: pd.DataFrames to convert
	:type: dict or list
	:param str var: Name of column to select from df
	:param array x: Array of datapoints to evaluate KDE on
	:param bool absv: (or None) Set to True to use absolute value of selected data for KDE calculation
	:returns: List of KDE arrays
	'''

	L = []

	#Dictionary workflow
	if type(data) ==  type({}):
		for key in data.keys():
			if absv == False:
				y = data[key][var].sample(frac=0.1)
			elif absv == True:
				y = np.abs(data[key][var].sample(frac=0.1))
			kde = scipy.stats.gaussian_kde(y).evaluate(x)
			L.append(kde)
	#List workflow
	elif type(data) == type([]):
		for df in data:
			if absv == False:
				y = df[var].sample(frac=0.1)
			elif absv == True:
				y = np.abs(df[var].sample(frac=0.1))
			kde = scipy.stats.gaussian_kde(y).evaluate(x)
			L.append(kde)

	return(L)

def calculate_area_error(pdf,Lkde,x):
	'''
	Calculate area between PDF and each kde in Lkde

	:param array pdf: Array of probability distribution function that is the same shape as kdes in Lkde
	:param list Lkde: List of arrays of Kdes
	:param array x: Array of datapoints used to generate pdf and kdes
	:returns: List of error values for each kde in Lkde
	'''

	L = []

	for kde in Lkde:
		L.append(simps(np.abs(pdf-kde),x))

	return(L)

def rescale_variable(Ddfs,var,newvar):
	'''
	Rescale variable from -1 to 1 and save in newvar column on original dataframe

	:param dict Ddfs: Dictionary of pd.DataFrames
	:param str var: Name of column to select from dfs
	:param str newvar: Name to use for new data in appended column
	:returns: Dictionary of dataframes containing column of rescaled data
	'''

	Dout = {}

	for key in Ddfs.keys():
		df = Ddfs[key]
		y = df[var]

		#Find min and max values
		mi,ma = y.min(),y.max()

		#Divide dataframe
		dfmin = df[y < 0]
		dfmax = df[y >= 0]

		#Add rescaled variable to dataframe
		dfmin = dfmin.join(pd.DataFrame({newvar:y/np.abs(mi)}))
		dfmax = dfmax.join(pd.DataFrame({newvar:y/np.abs(ma)}))

		dfout = dfmin.append(dfmax)
		Dout[key] = dfout

	return(Dout)

class paramsClass:
	'''
	A class to read and validate parameters for multiprocessing transformation.
	Validated parameters can be read as attributes of the object
	'''

	def __init__(self,path=None,dparams=None):
		'''
		Read json data in config file and validate that parameter inputs are correct

		:param str path: optional, Complete path to the config file
		:param dict dparams: optional, Dictionary with configuration parameters
		'''

		
		if dparams!=None:
			params = dparams
		elif path!=None:
			# Read file containing config data and parse json data
			config_data = open(path).read()
			params = json.loads(config_data)
		else:
			print('''No valid input in form of path to config file
				or existing parameter dictionary''')

		self.check_config(params,path)

	def add_outdir(self,path):
		'''
		Add out directory as an attribute of the class

		:param str path: Complete path to the output directory
		'''

		self.outdir = path

	def check_config(self,D,path):
		'''
		Check that each parameter in the config file is correct and raise an error if it isn't

		:param dict D: Dictionary containing parameters from the config file
		:param str path: Complete filepath to the config file
		'''

		#Check that the rootdir is valid
		if D['rootdir'] == '':
			print('Root directory path (rootdir) is not defined. Modify in',path)
			raise
		elif os.path.isdir(D['rootdir']):
			self.rootdir = D['rootdir']
		else:
			print('Root directory path (rootdir) must specify an existing directory. Modify in',path)
			raise
		#Check c1 key
		if D['c1-key'] == '':
			print('C1 directory key (c1-key) is not defined. Modify in',path)
			raise
		else:
			self.c1_key = D['c1-key']
		
		#Check that c1, structural channel, is defined
		if D['c1-dir'] == '':
			print('C1 directory path (c1-dir) is not defined. Modify in',path)
			raise
		elif os.path.isdir(D['c1-dir']):
			self.c1_dir = D['c1-dir']
			self.c1_files = glob.glob(os.path.join(self.c1_dir,D['c1-key']+'*.psi'))
		else:
			print('C1 directory path (c1-dir) is not defined. Modify in',path)
			raise

		#Check other channels and add to list if valid
		self.Lcdir = []
		self.Lcfiles = []
		self.Lckey = []
		for d,k in zip(['c2-dir','c3-dir','c4-dir'],['c2-key','c3-key','c4-key']):
			if D[d] != '':
				if os.path.isdir(D[d]):
					self.Lcdir.append(D[d])
					self.Lcfiles.append(glob.glob(os.path.join(D[d],D['c1-key']+'*.psi')))
					if D[k] == '':
						print('Channel key (',k,') is not defined. Modify in',path)
						raise
					else:
						self.Lckey.append(D[k])
				else:
					print('Channel directory path (',d,') is not defined. Modify in',path)
					raise

		#Validate that experiment name is defined
		if D['expname'] == '':
			print('Experiment name (expname) is not defined in',path)
			raise
		else:
			self.expname = D['expname']

		#Check median threshold value
		if D['medthresh'] <= 1 and D['medthresh'] >= 0:
			self.medthresh = D['medthresh']
		else:
			print('Median threshold (medthresh) must be a number between 0 and 1. Modify in',path)
			raise

		#Check that radius is an integer
		if type(D['radius']) == int:
			self.radius = D['radius']
		else:
			print('Radius input must be an integer. Modify in',path)
			raise

		#Check genthresh float between 0 and 1
		if D['genthresh'] <= 1 and D['genthresh'] >= 0:
			self.genthresh = D['genthresh']
		else:
			print('General threshold input must be a number between 0 and 1. Modify in',path)
			raise

		#Check format of microns
		if type(D['microns']) == list:
			for i in D['microns']:
				#If it is a float or int
				if not (type(i) == float) | (type(i) == int):
					print('Micron inputs must be numeric values. Modify in',path)
					raise
			self.microns = D['microns']
		else:
			print('Micron input must be a list of numeric values. Modify in',path)
			raise

		#Check degree
		if type(D['deg']) != int:
			print('Degree input must be an integer. Modify in',path)
			raise
		else:
			self.deg = D['deg']

		#Check that comporder is a list of integers between 0 and 2
		if type(D['comporder']) == list:
			for i in D['comporder']:
				#Check that it is an int between 0 and 2
				if i not in [0,1,2]:
					print('Comporder input must be a list of integers between 0 and 2. Modify in',path)
					raise
			self.comporder = D['comporder']
		else:
			print('Comporder input must be a list of integers between 0 and 2. Modify in',path)
			raise

		#Check fit dimensions
		if type(D['fitdim']) == list:
			for i in D['fitdim']:
				if i not in ['x','y','z']:
					print('Fitdim input must be a list of \'x\', \'y\', or \'z\'. Modify in',path)
					raise
			self.fitdim = D['fitdim']
		else:
			print('Fitdim input must be a list of \'x\', \'y\', or \'z\'. Modify in',path)
			raise

		if type(D['twoD']) == bool:
			self.twoD = D['twoD']
		else:
			print('Specification for 2D transformation must be boolean. Modify in',path)
			raise

		self.scale = [1,1,1]

		print('All parameter inputs are correct')
```
