## Supplementary material for "ΔSCOPE: A new method to quantify 3D biological structures and identify differences in zebrafish forebrain development": An archived copy of the DeltaSCOPE code repository.: alignment.py

```
import numpy as np
import pandas as pd
import matplotlib.pyplot as plt
import os
import re
import h5py
import deltascope as cranium

def read_h5prob_to_dict(fdir):
    '''
    Read in hdf5 array from probability file into dictionary

    :param str fdir: Filepath to directory
    :return: Dictionary with filenumber as key and raw data as item
    '''
    D = {}
    for f in os.listdir(fdir):
        if 'h5' in f:
            num = re.findall(r'\d+',f.split('.')[0])[1]
            file = h5py.File(os.path.join(fdir,f))
            D[num] = file.get('exported_data')

    return(D)

def preprocess(fpath,p,stop=None,pca=None,mm=None,vertex=None):
    '''
    Generate brain object and process until stop is triggered

    :param str stop: 'df_thresh' 'median' 'df_align'
    :return: Cranium brain object
    '''

    b = cranium.brain()
    b.read_data(fpath)
    b.preprocess_data(p['gthresh'],p['scale'],p['microns'])
    if stop == 'df_thresh':
        return(b)

    if pca == None:
        b.calculate_pca_median(b.raw_data,p['mthresh'],p['radius'],p['microns'])
        pca = b.pcamed
    if stop == 'median':
        return(b)

    b.pca_transform_3d(b.df_thresh,pca,p['comp_order'],p['fit_dim'],deg=p['deg'],mm=mm,vertex=vertex)
    if (stop == 'df_align') | (stop == None):
        return(b)

def plot_lines(ax,i):
    '''
    Plots origin lines on graph

    :param plt.subplots ax: Subplot array with minimum dimension [2,3]
    :param int i: Index of row that should be labeled
    '''

    ax[i,0].axhline(0,c='y')
    ax[i,0].axvline(0,c='r')
    ax[i,1].axhline(0,c='y')
    ax[i,1].axvline(0,c='c')
    ax[i,2].axhline(0,c='c')
    ax[i,2].axvline(0,c='r')

def scatter_df(ax,i,df):
    '''
    Scatter plot of df with projections in all 3 axes

    :param plt.subplots ax: Subplot array with minimum dimension [2,3]
    :param int i: Index of row that should be labeled
    :param pd.DataFrame df: Dataframe containing 'x','y','z' columns preferably downsampled for faster plotting
    '''

    ax[i,0].scatter(df.x,df.y,s=10)
    ax[i,1].scatter(df.x,df.z,s=10)
    ax[i,2].scatter(df.z,df.y,s=10)

def imshow_arr(ax,i,arr,f):
    '''
    Scatter plot of df with projections in all 3 axes according to supplied function f

    :param plt.subplots ax: Subplot array with minimum dimension [2,3]
    :param int i: Index of row that should be labeled
    :param np.array arr: 3D image array
    :param np.fxn f: Numpy function that takes an array and an axis value as parameters, usually np.min or np.max
    '''

    ax[i,0].imshow(f(arr,axis=0),cmap='Greys')
    ax[i,1].imshow(f(arr,axis=1),cmap='Greys')
    ax[i,2].imshow(f(arr,axis=2),cmap='Greys')

def rotate(df,A):
    '''
    Transform dataframe according to matrix A

    :param pd.DataFrame df: Dataframe containing 'x','y','z'
    :param np.array A: 3x3 array containing transformation matrix
    :return: New dataframe with transformed data
    '''

    rt = np.dot(np.array(df[['x','y','z']]),A)
    dfo = pd.DataFrame({'x':rt[:,0],'y':rt[:,1],'z':rt[:,2]})
    return(dfo)

def save_at(k,df,outdir,expname):
    '''
    Save dataframe as psi file in outdir

    :param str k: Sample number
    :param pd.DataFrame df: Dataframe containing 'x','y','z'
    :param str outdir: Filepath to destination directory for psi
    :param str expname: Descriptive name for experiment to be used in filename
    '''
    cranium.write_data(os.path.join(outdir,'AT_'+k+'_'+expname+'.psi'),df)

def save_zrf(k,df,outdir,expname):
    '''
    Save dataframe as psi file in outdir

    :param str k: Sample number
    :param pd.DataFrame df: Dataframe containing 'x','y','z'
    :param str outdir: Filepath to destination directory for psi
    :param str expname: Descriptive name for experiment to be used in filename
    '''
    cranium.write_data(os.path.join(outdir,'ZRF_'+k+'_'+expname+'.psi'),df)

def save_both(k,dfa,dfz,outdir,expname):
    '''
    Save dataframe as psi file in outdir

    :param str k: Sample number
    :param pd.DataFrame dfa: AT dataframe containing 'x','y','z'
    :param pd.DataFrame dfa: Zrf dataframe containing 'x','y','z'
    :param str outdir: Filepath to destination directory for psi
    :param str expname: Descriptive name for experiment to be used in filename
    '''
    save_at(k,dfa,outdir,expname)
    save_zrf(k,dfz,outdir,expname)

def save_caax(k,df,outdir,expname):

    cranium.write_data(os.path.join(outdir,'caax_'+k+'_'+expname+'.psi'),df)

def save_all(k,dfa,dfz,dfc,outdir,expname):
    '''
    Save dataframe as psi file in outdir

    :param str k: Sample number
    :param pd.DataFrame dfa: AT dataframe containing 'x','y','z'
    :param pd.DataFrame dfa: Zrf dataframe containing 'x','y','z'
    :param str outdir: Filepath to destination directory for psi
    :param str expname: Descriptive name for experiment to be used in filename
    '''
    save_at(k,dfa,outdir,expname)
    save_zrf(k,dfz,outdir,expname)
    save_caax(k,dfc,outdir,expname)

######### Alignment functions ###############

def calc_rotation(pts,d):
    '''
    Calculate a rotation matrix A based on two anchor points

    :param pd.DataFrame pts: Two points in x and the dimension d
    :param str d: Second dimension of the two points: 'z' or 'y'
    :return: midpoint, rotation matrix
    '''
    dx = (pts.iloc[0].x - pts.iloc[1].x)/2
    dy = (pts.iloc[0][d] - pts.iloc[1][d])/2

    mp = [pts.iloc[1].x + dx, pts.iloc[1][d] + dy]

    phi = np.arctan2(dy,dx)
    if phi > np.pi/2:
        phi = -(np.pi - phi)
    if d == 'y':
        A = np.array([[np.cos(phi),-np.sin(phi),0],
        			[np.sin(phi),np.cos(phi),0],
        			[0,0,1]])
    else:
        A = np.array([[np.cos(phi),0,-np.sin(phi)],
                    [0,1,0],
                    [np.sin(phi),0,np.cos(phi)]])

    return(mp,A)

def xrotate(df,Ldf,d,pts=None):
    '''
    Define two anchor points and rotate data so that the anchor points are level

    :param pd.DataFrame df: Dataframe that will be used to calculate the anchor points and rotation
    :param list Ldf: List of additionally dataframes that should be rotated based on the calculation from `df`
    :param str d: Second dimension of the plane that needs to be rotated: 'z' or 'y'
    :param pd.DataFrame pts: Two points in x and the dimension d
    :return: `df` after rotation, a list of rotated dataframes from `Ldf`, `pts`
    '''

    if type(pts) == type(None):
        pt1,pt2 = cranium.find_anchors(df,d)
        pts = pd.DataFrame({'x':[pt1['x'],pt2['x']],d:[pt1[d],pt2[d]]})

    mp,A = calc_rotation(pts,d)

    out = rotate(df,A)
    Lout = []
    for d in Ldf:
        Lout.append(rotate(d,A))

    return(out,Lout,pts)

def zyswitch(df,Ldf):
    '''
    Switch z and y axes by reassigning z to y and vice versa

    :param pd.DataFrame df: Dataframe that will be used to calculate the anchor points and rotation
    :param list Ldf: List of additionally dataframes that should be rotated based on the calculation from `df`
    :return: `df` and `Ldf` after exchanging z and y
    '''

    out = pd.DataFrame({'x':df.x,'y':df.z,'z':df.y})
    Lout = []
    for d in Ldf:
        Lout.append(pd.DataFrame({'x':d.x,'y':d.z,'z':d.y}))

    return(out,Lout)

def vertex(df,Ldf,pts=None):
    '''
    Calculate the vertex in the XZ plane and shift the vertex to the origin

    :param pd.DataFrame df: Dataframe that will be used to calculate the anchor points and rotation
    :param list Ldf: List of additionally dataframes that should be rotated based on the calculation from `df`
    :return: `df` and `Ldf` after translation of the vertex to the origin
    '''

    if type(pts)!=type(None):
        cf = np.polyfit(pts.x,pts.z,2)
    else:
        cf = np.polyfit(df.x,df.z,2)
    x = -cf[1]/(2*cf[0])
    z = np.poly1d(cf)(x)
    y = np.mean(df.y)

    out = pd.DataFrame({
        'x':df.x-x,
        'y':df.y-y,
        'z':df.z-z
    })
    Lout = []
    for d in Ldf:
        Lout.append(pd.DataFrame({
            'x':d.x-x,
            'y':d.y-y,
            'z':d.z-z
        }))

    if type(pts)!=type(None):
        cfout = np.polyfit(pts.x-x,pts.z-z,2)
    else:
        cfout = np.polyfit(out.x,out.z,2)

    return(out,Lout,cfout)

def flip(df,Ldf):
    '''
    Rotate by 180 degrees around the x axis

    :param pd.DataFrame df: Dataframe that will be used to calculate the anchor points and rotation
    :param list Ldf: List of additionally dataframes that should be rotated based on the calculation from `df`
    :return: `df` and `Ldf` after rotation
    '''

    A = np.array([[1,0,0],
                [0,np.cos(np.pi),-np.sin(np.pi)],
                [0,np.sin(np.pi),np.cos(np.pi)]])

    out = rotate(df,A)
    Lout = []
    for d in Ldf:
        Lout.append(rotate(d,A))

    return(out,Lout)

def yzrotate(df,Ldf,mm=None):
    '''
    Fit a line in YZ and rotate so the line is horizontal

    :param pd.DataFrame df: Dataframe that will be used to calculate the anchor points and rotation
    :param list Ldf: List of additionally dataframes that should be rotated based on the calculation from `df`
    :return: `df` and `Ldf` after rotation to correct the lines, 1d line function, range of z values for the line
    '''

    if mm == None:
        mm = np.polyfit(df.z,df.y,1)
    p = np.poly1d(mm)
    xrange = np.arange(1.1*np.min(df.z), 1.1*np.max(df.z))
    phi = -np.arctan(mm[0])
    A = np.array([[1,0,0],
                [0,np.cos(phi),-np.sin(phi)],
                [0,np.sin(phi),np.cos(phi)]])

    out = rotate(df,A)
    Lout = []
    for d in Ldf:
        Lout.append(rotate(d,A))

    return(out,Lout,p,xrange)

def check_yz(df,Ldf,mm=None):
    '''
    Fit line in YZ and rotate line to be horizontal using `yzrotate` then display graph

    :param pd.DataFrame df: Dataframe that will be used to calculate the anchor points and rotation
    :param list Ldf: List of additionally dataframes that should be rotated based on the calculation from `df`
    :return: `df` and `Ldf` after rotation to correct the lines, array of graph subplots, 1d line function
    '''

    out,Lout,p,xrange = yzrotate(df,Ldf,mm=mm)
    ax = make_graph([df,out]+Lout)
    ax[0,2].plot(xrange,p(xrange),c='m')

    return(out,Lout,ax,p)

def make_graph(Ldf,Lim=[]):
    '''
    Plot the three projections of each dataframe in `Ldf`

    :param list Ldf: List of additionally dataframes that should be rotated based on the calculation from `df`
    :return: Array of subplots with minimum dimension of 2x3
    '''

    n = len(Ldf)+len(Lim)
    fig,ax = plt.subplots(n,3,subplot_kw={'aspect':'equal','adjustable':'datalim'},figsize=(12,n*4))

    for i,d in enumerate(Ldf):
        scatter_df(ax,i,d.sample(frac=0.1))
        plot_lines(ax,i)

    for i,im in enumerate(Lim):
        imshow_arr(ax,len(Ldf)+i,im,np.min)

    return(ax)

def start(k,Da,LD,im=False):
    '''
    Extract `df_align` dataframes from `cranium.brain` objects and plots using `make_graph`

    :param str k: Key corresponding to the relevant sample
    :param dict Da: Dictionary containing brain objects for each sample with key `k`
    :param list LD: List of any additional dictionaries from which `df_align` should be extracted
    :returns: key, primary dataframe, list of additional dataframes, subplot array
    '''
    Lim = []
    Lim.append(Da[k].raw_data)
    df = Da[k].df_align
    Ldf = []
    for D in LD:
        Ldf.append(D[k].df_align)
        Lim.append(D[k].raw_data)

    if im==True:
        ax = make_graph([df]+Ldf,Lim)
    else:
        ax = make_graph([df]+Ldf,[])
    return(k,df,Ldf,ax)

def check_pts(df,Ldf,d):
    '''
    Calculates anchor points and plots before and after rotation

    :param pd.DataFrame df: Dataframe that will be used to calculate the anchor points and rotation
    :param list Ldf: List of additionally dataframes that should be rotated based on the calculation from `df`
    :param str d: Second dimension of the plane that needs to be rotated: 'z' or 'y'
    :returns: Rotated `df`, list of rotated dfs from `Ldf`, anchor points, subplot array
    '''
    df1,Ldf1,pts = xrotate(df,Ldf,d)
    ax = make_graph([df,df1]+Ldf1)
    if d=='z':
        ax[0,1].scatter(pts.x,pts[d],c='r')
    if d=='y':
        ax[0,0].scatter(pts.x,pts[d],c='r')
    return(df1,Ldf1,pts,ax)

def revise_pts(df,Ldf,d,pts):
    '''
    Rotate and plot data based on `pts` parameter

    :param pd.DataFrame df: Dataframe that will be used to calculate the anchor points and rotation
    :param list Ldf: List of additionally dataframes that should be rotated based on the calculation from `df`
    :param str d: Second dimension of the plane that needs to be rotated: 'z' or 'y'
    :param pd.DataFrame pts: Two points in x and the dimension d
    :returns: Rotated `df`, list of rotated dfs from `Ldf`, subplot array
    '''
    df1,Ldf1,pts2 = xrotate(df,Ldf,d,pts=pts)
    ax = make_graph([df,df1]+Ldf1)
    if d=='z':
        ax[0,1].scatter(pts2.x,pts2[d],c='r')
    if d=='y':
        ax[0,0].scatter(pts2.x,pts2[d],c='r')
    return(df1,Ldf1,ax)

def ch_vertex(df,Ldf,pts=None):
    '''
    Calculate vertex based on `df` and translate all dataframes to shift vertex to origin

    :param pd.DataFrame df: Dataframe that will be used to calculate the anchor points and rotation
    :param list Ldf: List of additionally dataframes that should be rotated based on the calculation from `df`
    :returns: Translated `df`, list of translated dataframes from `Ldf`, subplot array
    '''
    df1,Ldf1,mm = vertex(df,Ldf,pts=pts)
    ax = make_graph([df1]+Ldf1)
    xrange = np.arange(df1.x.min(),df1.x.max())
    ax[0,1].plot(xrange,np.poly1d(mm)(xrange),c='m')
    return(df1,Ldf1,mm,ax)
```
