## Supplementary material for "ΔSCOPE: A new method to quantify 3D biological structures and identify differences in zebrafish forebrain development": An archived copy of the DeltaSCOPE code repository.: mpTransformation.py

```
import deltascope as cranium
import multiprocessing as mp
import time
import os
from functools import partial
from sys import argv
from sys import exc_info
import re
import json

class paramsClass:
	'''
	A class to read and validate parameters for multiprocessing transformation.
	Validated parameters can be read as attributes of the object
	'''

	def __init__(self,path):
		'''
		Read json data in config file and validate that parameter inputs are correct

		:param str path: Complete path to the config file
		'''

		# Read file containing config data and parse json data
		config_data = open(path).read()
		params = json.loads(config_data)

		self.check_config(params,path)

	def add_outdir(self,path):
		'''
		Add out directory as an attribute of the class

		:param str path: Complete path to the output directory
		'''

		self.outdir = path

	def check_config(self,D,path):
		'''
		Check that each parameter in the config file is correct and raise an error if it isn't

		:param dict D: Dictionary containing parameters from the config file
		:param str path: Complete filepath to the config file
		'''

		#Check that the rootdir is valid
		if D['rootdir'] == '':
			print('Root directory path (rootdir) is not defined. Modify in',path)
			raise
		elif os.path.isdir(D['rootdir']):
			self.rootdir = D['rootdir']
		else:
			print('Root directory path (rootdir) must specify an existing directory. Modify in',path)
			raise

		#Check that c1, structural channel, is defined
		if D['c1-dir'] == '':
			print('C1 directory path (c1-dir) is not defined. Modify in',path)
			raise
		elif os.path.isdir(D['c1-dir']):
			self.c1_dir = D['c1-dir']
			self.c1_files = os.listdir(self.c1_dir)
		else:
			print('C1 directory path (c1-dir) is not defined. Modify in',path)
			raise

		#Check c1 key
		if D['c1-key'] == '':
			print('C1 directory key (c1-key) is not defined. Modify in',path)
			raise
		else:
			self.c1_key = D['c1-key']

		#Check other channels and add to list if valid
		self.Lcdir = []
		self.Lcfiles = []
		self.Lckey = []
		for d,k in zip(['c2-dir','c3-dir','c4-dir'],['c2-key','c3-key','c4-key']):
			if D[d] != '':
				if os.path.isdir(D[d]):
					self.Lcdir.append(D[d])
					self.Lcfiles.append(os.listdir(D[d]))
					if D[k] == '':
						print('Channel key (',k,') is not defined. Modify in',path)
						raise
					else:
						self.Lckey.append(D[k])
				else:
					print('Channel directory path (',d,') is not defined. Modify in',path)
					raise

		#Validate that experiment name is defined
		if D['expname'] == '':
			print('Experiment name (expname) is not defined in',path)
			raise
		else:
			self.expname = D['expname']

		#Check median threshold value
		if D['medthresh'] <= 1 and D['medthresh'] >= 0:
			self.medthresh = D['medthresh']
		else:
			print('Median threshold (medthresh) must be a number between 0 and 1. Modify in',path)
			raise

		#Check that radius is an integer
		if type(D['radius']) == int:
			self.radius = D['radius']
		else:
			print('Radius input must be an integer. Modify in',path)
			raise

		#Check genthresh float between 0 and 1
		if D['genthresh'] <= 1 and D['genthresh'] >= 0:
			self.genthresh = D['genthresh']
		else:
			print('General threshold input must be a number between 0 and 1. Modify in',path)
			raise

		#Check format of microns
		if type(D['microns']) == list:
			for i in D['microns']:
				#If it is a float or int
				if not (type(i) == float) | (type(i) == int):
					print('Micron inputs must be numeric values. Modify in',path)
					raise
			self.microns = D['microns']
		else:
			print('Micron input must be a list of numeric values. Modify in',path)
			raise

		#Check degree
		if type(D['deg']) != int:
			print('Degree input must be an integer. Modify in',path)
			raise
		else:
			self.deg = D['deg']

		#Check that comporder is a list of integers between 0 and 2
		if type(D['comporder']) == list:
			for i in D['comporder']:
				#Check that it is an int between 0 and 2
				if i not in [0,1,2]:
					print('Comporder input must be a list of integers between 0 and 2. Modify in',path)
					raise
			self.comporder = D['comporder']
		else:
			print('Comporder input must be a list of integers between 0 and 2. Modify in',path)
			raise

		#Check fit dimensions
		if type(D['fitdim']) == list:
			for i in D['fitdim']:
				if i not in ['x','y','z']:
					print('Fitdim input must be a list of \'x\', \'y\', or \'z\'. Modify in',path)
					raise
			self.fitdim = D['fitdim']
		else:
			print('Fitdim input must be a list of \'x\', \'y\', or \'z\'. Modify in',path)
			raise

		if type(D['twoD']) == bool:
			self.twoD = D['twoD']
		else:
			print('Specification for 2D transformation must be boolean. Modify in',path)
			raise

		self.scale = [1,1,1]

		print('All parameter inputs are correct')

def check_nums(P):
	'''
	Check that the numbers of files selected by the same list index match
	Raises an error if file numbers are mismatched

	:param :class:`paramClass` P: Object containing all variables from config file
	'''
	Lnums = []

	print('hello')

	for i,f in enumerate(P.c1_files):
		print(i,f)
		if 'h5' in f:
			n = re.findall(r'\d+',f.split('.')[0])[0]
			print('n',n)
			for Lf in P.Lcfiles:
				if n not in Lf[i]:
					print('File numbers are mismatched between channel directories.')
					raise
			Lnums.append(i)

	return(Lnums)

def process(num,P=None):
	'''
	Run through the processing steps for a single sample through saving psi files

	:param int num: Index of the file that is currently being processed
	:param :class:`paramClass` P: Object containing all variables from config file
	'''

	tic = time.time()
	print(num,'Starting sample')
	try:
		e = cranium.embryo(P.expname,num,P.outdir)

		#Add channels and preprocess data
		e.add_channel(os.path.join(P.c1_dir,P.c1_files[num]),P.c1_key)
		e.chnls[P.c1_key].preprocess_data(P.genthresh,P.scale,P.microns)
		for i in range(len(P.Lcdir)):
			e.add_channel(os.path.join(P.Lcdir[i],P.Lcfiles[i][num]),P.Lckey[i])
			e.chnls[P.Lckey[i]].preprocess_data(P.genthresh,P.scale,P.microns)

		#Calculate PCA transformation for structural channel, c1
		if P.twoD == True:
			e.chnls[P.c1_key].calculate_pca_median_2d(e.chnls[P.c1_key].raw_data,P.medthresh,P.radius,P.microns)
			pca = e.chnls[P.c1_key].pcamed
			e.chnls[P.c1_key].pca_transform_2d(e.chnls[P.c1_key].df_thresh,pca,P.comporder,P.fitdim,deg=P.deg)

			#Transform additional channels
			for i in range(len(P.Lcdir)):
				e.chnls[P.Lckey[i]].pca_transform_2d(e.chnls[P.Lckey[i]].df_thresh,pca,P.comporder,P.fitdim,deg=P.deg)

		else:
			e.chnls[P.c1_key].calculate_pca_median(e.chnls[P.c1_key].raw_data,P.medthresh,P.radius,P.microns)
			pca = e.chnls[P.c1_key].pcamed
			e.chnls[P.c1_key].pca_transform_3d(e.chnls[P.c1_key].df_thresh,pca,P.comporder,P.fitdim,deg=P.deg)

			#Transform additional channels
			for i in range(len(P.Lcdir)):
				e.chnls[P.Lckey[i]].pca_transform_3d(e.chnls[P.Lckey[i]].df_thresh,pca,P.comporder,P.fitdim,deg=P.deg)

		# print(num,'Starting coordinate transformation')
		# e.chnls[P.c1_key].transform_coordinates()
		# for i in range(len(P.Lcdir)):
		# 	e.chnls[P.Lckey[i]].transform_coordinates()

		#e.save_psi()
		columns = ['x','y','z']

		for ch in e.chnls.keys():
			cranium.write_data(os.path.join(e.outdir,
				e.name+'_'+str(e.number)+'_'+ch+'.psi'),
				e.chnls[ch].df_align[columns])

		toc = time.time()
		print(num,'Complete',toc-tic)

	except:
		toc = time.time()
		print(num,'Failed',toc-tic,exc_info())


if __name__=='__main__':

	f,config_path = argv
	P = paramsClass(config_path)

	#Create out directory stamped with current date and time
	outdir = os.path.join(P.rootdir,'Output'+time.strftime("%m-%d-%H-%M",time.localtime()))
	os.mkdir(outdir)
	P.add_outdir(outdir)
	print('outdir',outdir)

	processfxn = partial(process,P=P)

	Lnums = check_nums(P)
	n = len(Lnums)
	# Initiate map pools in sets of 5 samples
	for i in range(0,n,5):
		if i+5>n:
			L = Lnums[i:n]
		else:
			L = Lnums[i:i+5]

		pool = mp.Pool()
		pool.map(processfxn,L)
		pool.close()
		pool.join()
```
