## Supplementary material for "ΔSCOPE: A new method to quantify 3D biological structures and identify differences in zebrafish forebrain development": An archived copy of the DeltaSCOPE code repository.: new.py

```
import os
import numpy as np
import h5py
import pandas as pd
from sklearn.decomposition import PCA
from skimage.filters import median
from skimage.morphology import disk
import sympy as sp
from scipy.spatial import KDTree
import time

class brain:
	''' Object to manage biological data and associated functions. '''

	def __init__(self):
		'''Initialize brain object'''

	def setup_test_data(self,size=None,gthresh=0.5,scale=[1,1,1],microns=[0.16,0.16,0.21],mthresh=0.2,radius=20,comp_order=[0,2,1],fit_dim=['x','z'],deg=2):
		'''Setup a test dataset to use for testing transform coordinates
		:param int size: Number of points to sample for the test dataset
		'''

		self.read_data(os.path.abspath('..\data\C1\AT_01_Probabilities.h5'))
		self.preprocess_data(gthresh,scale,microns)
		self.calculate_pca_median(self.raw_data,mthresh,radius,microns)
		self.pca_transform_3d(self.df_thresh,self.pcamed,comp_order,fit_dim,deg=deg)
		self.mm = self.fit_model(self.df_align,2,['x','z'])
		if size!= None:
			self.df_align = self.df_align.sample(size)

	def read_data(self,filepath):
		'''
		Reads 3D data from file and selects appropriate channel based on the
		assumption that the channel with the most zeros has zero as the value for no signal

		:param str filepath: Filepath to hdf5 probability file
		:return: Creates the variable :attr:`brain.raw_data`

		.. py:attribute:: brain.raw_data

			Array of shape [z,y,x] containing raw probability data
		'''

		#Read h5 file and extract probability data
		f = h5py.File(filepath,'r')

		#Select either channe0/1 or exported_data
		d = f.get('exported_data')
		if d != None:
			c1 = np.array(d[:,:,:,0])
			c2 = np.array(d[:,:,:,1])
		elif d == None:
			c1 = np.array(f.get('channel0'))
			c2 = np.array(f.get('channel1'))

		#Figure out which channels has more zeros and therefore is background
		if np.count_nonzero(c1<0.1) > np.count_nonzero(c1>0.9):
			#: Array of shape [z,y,x] containing raw probability data
			self.raw_data = c2
		else:
			#: Array of shape [z,y,x] containing raw probability data
			self.raw_data = c1

	def create_dataframe(self,data,scale):
		'''
		Creates a pandas dataframe containing the x,y,z and signal/probability value for each point in the :py:attr:`brain.raw_data` array

		:param array data: Raw probability data in 3D array
		:param array scale: Array of length three containing the micron values for [x,y,z]
		:return: Pandas DataFrame with xyz and probability value for each point
		'''

		#NB: scale variable actually contains microns dimensions

		dim = data.shape
		xyz = np.zeros((dim[0],dim[1],dim[2],4))

		#Generate array with xyz values for each point
		for x in range(dim[2]):

			xyz[:,:,x,2] = x
			xyz[:,:,x,3] = data[:,:,x]

			zindex = np.arange(0,dim[0],1)
			yindex = np.arange(0,dim[1],1)
			gy,gz = np.meshgrid(yindex,zindex)

			xyz[:,:,x,0] = gz
			xyz[:,:,x,1] = gy

		flat = np.reshape(xyz,(-1,4))

		#Create dataframe of points and scale to microns
		df = pd.DataFrame({'x':flat[:,2]*scale[0],'y':flat[:,1]*scale[1],'z':flat[:,0]*scale[2],'value':flat[:,3]})
		return(df)

	def plot_projections(self,df,subset):
		'''
		Plots the x, y, and z projections of the input dataframe in a matplotlib plot

		:param pd.DataFrame df: Dataframe with columns: 'x','y','z'
		:param float subset: Value between 0 and 1 indicating what percentage of the df to subsample
		:returns: Matplotlib figure with three labeled scatterplots
		'''

		df = df.sample(frac=subset)

    	#Create figure and subplots
		fig = plt.figure(figsize=(12,6))
		ax = fig.add_subplot(131)
		ay = fig.add_subplot(132)
		az = fig.add_subplot(133)

		#Create scatter plot for each projection
		ax.scatter(df.x,df.z)
		ay.scatter(df.x,df.y)
		az.scatter(df.z,df.y)

		#Plot model
		xvalues = np.arange(np.min(df.x),np.max(df.x))
		ax.plot(xvalues,self.mm.p(xvalues),c='y')

		#Add labels
		ax.set_title('Y projection')
		ay.set_title('Z projection')
		az.set_title('X projection')
		ax.set_xlabel('X')
		ax.set_ylabel('Z')
		ay.set_xlabel('X')
		ay.set_ylabel('Y')
		az.set_xlabel('Z')
		az.set_ylabel('Y')

		#Adjust spacing and show plot
		plt.subplots_adjust(wspace=0.4)

		return(fig)

	def preprocess_data(self,threshold,scale,microns):
		'''
		Thresholds and scales data prior to PCA

		Creates :py:attr:`brain.threshold`, :py:attr:`brain.df_thresh`, and :py:attr:`brain.df_scl`

		:param float threshold: Value between 0 and 1 to use as a cutoff for minimum pixel value
		:param array scale: Array with three values representing the constant by which to multiply x,y,z respectively
		:param array microns: Array with three values representing the x,y,z micron dimensions of the voxel

		.. py:attribute:: brain.threshold

			Value used to threshold the data prior to calculating the model

		.. py:attribute:: brain.df_thresh

			Dataframe containing only points with values above the specified threshold

		.. py:attribute:: brain.df_scl

			Dataframe containing data from :py:attr:`brain.df_thresh` after a scaling value has been applied
		'''

		#: Dataframe with four columns: x,y,z,value with all points in :py:attr:`brain.raw_data`
		self.df = self.create_dataframe(self.raw_data,microns)

		#Create new dataframe with values above threshold
		self.threshold = threshold
		self.df_thresh = self.df[self.df.value < self.threshold]

		#Scale xyz by value in scale array to force PCA axis selection
		self.scale = scale
		self.df_scl = pd.DataFrame({
			'x':self.df_thresh.x * self.scale[0],
			'y':self.df_thresh.y * self.scale[1],
			'z':self.df_thresh.z * self.scale[2]})

	def process_alignment_data(self,data,threshold,radius,microns):
		'''
		Applies a median filter twice to the data which is used for alignment

		Ensures than any noise in the structural data does not interfere with alignment

		:param array data: Raw data imported by the function :py:func:`brain.read_data`
		:param float threshold: Value between 0 and 1 to use as a cutoff for minimum pixel value
		:param int radius: Integer that determines the radius of the circle used for the median filter
		:param array microns: Array with three values representing the x,y,z micron dimensions of the voxel
		:returns: Dataframe containing data processed with the median filter and threshold
		'''

		#Iterate over each plane and apply median filter twice
		out = np.zeros(data.shape)
		for z in range(data.shape[0]):
			out[z] = median(median(data[z],disk(radius)),disk(radius))

		outdf = self.create_dataframe(out,microns)

		# Changed 7/5/18
		# Skimage filter changes datatype to 0-255 range from 0-1
		# Modification to threshold allows consistent use of threshold based on 0-1 range
		thresh = outdf[outdf.value < 255*(1-threshold)]
		return(thresh)

	def calculate_pca_median(self,data,threshold,radius,microns):
		'''
		Calculate PCA transformation matrix, :py:attr:`brain.pcamed`, based on data (:py:attr:`brain.pcamed`) after applying median filter and threshold

		:param array data: 3D array containing raw probability data
		:param float threshold: Value between 0 and 1 indicating the lower cutoff for positive signal
		:param int radius: Radius of neighborhood that should be considered for the median filter
		:param array microns: Array with three values representing the x,y,z micron dimensions of the voxel

		.. py:attribute:: brain.median

			Pandas dataframe containing data that has been processed with a median filter twice and thresholded

		.. py:attribute:: brain.pcamed

			PCA object managing the transformation matrix and any resulting transformations

		'''

		self.median = self.process_alignment_data(data,threshold,radius,microns)

		self.pcamed = PCA()
		self.pcamed.fit(self.median[['x','y','z']])

	def calculate_pca_median_2d(self,data,threshold,radius,microns):
		'''
		Calculate PCA transformation matrix for 2 dimensions of data, :py:attr:`brain.pcamed`, based on data after applying median filter and threshold

		.. warning:: `fit_dim` is not used to determine which dimensions to fit. Defaults to x and z

		:param array data: 3D array containing raw probability data
		:param float threshold: Value between 0 and 1 indicating the lower cutoff for positive signal
		:param int radius: Radius of neighborhood that should be considered for the median filter
		:param array microns: Array with three values representing the x,y,z micron dimensions of the voxel
		'''

		self.median = self.process_alignment_data(data,threshold,radius,microns)

		self.pcamed = PCA()
		self.pcamed.fit(self.median[['y','z']])

	def pca_transform_2d(self,df,pca,comp_order,fit_dim,deg=2,mm=None,vertex=None,flip=None):
		'''
		Transforms `df` in 2D based on the PCA object, `pca`, whose transformation matrix has already been calculated

		Calling :py:func:`brain.align_data` creates :py:attr:`brain.df_align`

		.. warning:: `fit_dim` is not used to determine which dimensions to fit. Defaults to x and z

		:param pd.DataFrame df: Dataframe containing thresholded xyz data
		:param pca_object pca: A pca object containing a transformation object, e.g. :py:attr:`brain.pcamed`
		:param array comp_order: Array specifies the assignment of components to x,y,z. Form [x component index, y component index, z component index], e.g. [0,2,1]
		:param array fit_dim: Array of length two containing two strings describing the first and second axis for fitting the model, e.g. ['x','z']
		:param int deg: (or None) Degree of the function that should be fit to the model. deg=2 by default
		:param mm: (:py:class:`math_model` or None) Math model for primary channel
		:param array vertex: (or None) Array of type [vx,vy,vz] (:py:attr:`brain.vertex`) indicating the translation values
		:param Bool flip: (or None) Boolean value to determine if the data should be rotated by 180 degrees
		'''

		fit = pca.transform(df[['y','z']])
		df_fit = pd.DataFrame({
			'x':df.x,
			'y':fit[:,comp_order[1]-1],
			'z':fit[:,comp_order[2]-1]
			})

		self.align_data(df_fit,fit_dim,deg=2,mm=None,vertex=None,flip=None)

	def pca_transform_3d(self,df,pca,comp_order,fit_dim,deg=2,mm=None,vertex=None,flip=None):
		'''
		Transforms `df` in 3D based on the PCA object, `pca`, whose transformation matrix has already been calculated

		:param pd.DataFrame df: Dataframe containing thresholded xyz data
		:param pca_object pca: A pca object containing a transformation object, e.g. :py:attr:`brain.pcamed`
		:param array comp_order: Array specifies the assignment of components to x,y,z. Form [x component index, y component index, z component index], e.g. [0,2,1]
		:param array fit_dim: Array of length two containing two strings describing the first and second axis for fitting the model, e.g. ['x','z']
		:param int deg: (or None) Degree of the function that should be fit to the model. deg=2 by default
		:param mm: (:py:class:`math_model` or None) Math model for primary channel
		:param array vertex: (or None) Array of type [vx,vy,vz] (:py:attr:`brain.vertex`) indicating the translation values
		:param Bool flip: (or None) Boolean value to determine if the data should be rotated by 180 degrees
		'''

		fit = pca.transform(df[['x','y','z']])
		df_fit = pd.DataFrame({
			'x':fit[:,comp_order[0]],
			'y':fit[:,comp_order[1]],
			'z':fit[:,comp_order[2]]
			})

		self.align_data(df_fit,fit_dim,deg=2,mm=None,vertex=None,flip=None)

	def align_data(self,df_fit,fit_dim,deg=2,mm=None,vertex=None,flip=None):
		'''
		Apply PCA transformation matrix and align data so that the vertex is at the origin

		Creates :py:attr:`brain.df_align` and :py:attr:`brain.mm`

		:param pd.DataFrame df: dataframe containing thresholded xyz data
		:param array comp_order: Array specifies the assignment of components to x,y,z. Form [x component index, y component index, z component index], e.g. [0,2,1]
		:param array fit_dim: Array of length two containing two strings describing the first and second axis for fitting the model, e.g. ['x','z']
		:param int deg: (or None) Degree of the function that should be fit to the model. deg=2 by default
		:param mm: (:py:class:`math_model` or None) Math model for primary channel
		:param array vertex: (or None) Array of type [vx,vy,vz] (:py:attr:`brain.vertex`) indicating the translation values
		:param Bool flip: (or None) Boolean value to determine if the data should be rotated by 180 degrees

		.. py:attribute:: brain.df_align

			Dataframe containing point data aligned using PCA

		.. py:attribute:: brain.mm

			Math model object fit to data in brain object
		'''

		#If vertex for translation is not included
		if vertex == None and deg==2:
			#Calculate model
			model = np.polyfit(df_fit[fit_dim[0]],df_fit[fit_dim[1]],deg=deg)
			p = np.poly1d(model)

			#Find vertex
			a = -model[1]/(2*model[0])
			if fit_dim[0] == 'x':
				vx = a
				if fit_dim[1] == 'y':
					vy = p(vx)
					vz = df_fit.z.mean()
				else:
					vz = p(vx)
					vy = df_fit.y.mean()
			elif fit_dim[0] == 'y':
				vy = a
				if fit_dim[1] == 'x':
					vx = p(a)
					vz = df_fit.z.mean()
				else:
					vz = p(a)
					vx = df_fit.x.mean()
			elif fit_dim[0] == 'z':
				vz = a
				if fit_dim[1] == 'x':
					vx = p(a)
					vy = df_fit.y.mean()
				else:
					vy = p(a)
					vx = df_fit.x.mean()
			self.vertex = [vx,vy,vz]
		elif deg == 1:
			#Calculate model
			model = np.polyfit(df_fit[fit_dim[0]],df_fit[fit_dim[1]],deg=deg)
			p = np.poly1d(model)

			if fit_dim[0] == 'x':
				vx = df_fit.x.mean()
				if fit_dim[1] == 'y':
					vy = p(vx)
					vz = df_fit.z.mean()
				else:
					vz = p(vx)
					vy = df_fit.y.mean()
			elif fit_dim[0] == 'y':
				vy = df_fit.y.mean()
				if fit_dim[1] == 'x':
					vx = p(vy)
					vz = df_fit.z.mean()
				else:
					vz = p(vy)
					vx = df_fit.x.mean()
			elif fit_dim[0] == 'z':
				vz = df_fit.z.mean()
				if fit_dim[1] == 'x':
					vx = p(vz)
					vy = df_fit.y.mean()
				else:
					vy = p(vz)
					vx = df_fit.x.mean()
			self.vertex = [vx,vy,vz]
		else:
			self.vertex = vertex

		#Translate data so that the vertex is at the origin
		self.df_align = pd.DataFrame({
			'x': df_fit.x - self.vertex[0],
			'y': df_fit.y - self.vertex[1],
			'z': df_fit.z - self.vertex[2]
			})

		#Rotate data by 180 degrees if necessary
		if flip == None or flip == False:
			#Calculate model
			model = np.polyfit(df_fit[fit_dim[0]],df_fit[fit_dim[1]],deg=deg)
			p = np.poly1d(model)

			#If a is less than 0, rotate data
			if model[0] < 0:
				self.df_align = self.flip_data(self.df_align)
		elif flip == True:
			self.df_align = self.flip_data(self.df_align)

		#Calculate final math model
		if mm == None:
			self.mm = self.fit_model(self.df_align,deg,fit_dim)
		else:
			self.mm = mm

	def flip_data(self,df):
		'''
		Rotate data by 180 degrees

		:param dataframe df: Pandas dataframe containing x,y,z data
		:returns: Rotated dataframe
		'''

		r = np.array([[np.cos(np.pi),0,np.sin(np.pi)],
			[0,1,0],
			[-np.sin(np.pi),0,np.cos(np.pi)]])

		rot = np.dot(np.array(df),r)

		dfr = pd.DataFrame({'x':rot[:,0],'y':rot[:,1],'z':rot[:,2]})
		return(dfr)

	def fit_model(self,df,deg,fit_dim):
		'''Fit model to dataframe

		:param pd.DataFrame df: Dataframe containing at least x,y,z
		:param int deg: Degree of the function that should be fit to the model
		:param array fit_dim: Array of length two containing two strings describing the first and second axis for fitting the model, e.g. ['x','z']
		:returns: math model
		:rtype: :py:class:`math_model`
		'''

		mm = math_model(np.polyfit(df[fit_dim[0]], df[fit_dim[1]], deg=deg))
		return(mm)

	def setup_tree(self,xmin,xmax,xstep,rmax,rnum,acf):

		from sympy.abc import alpha,rho
		a,drho = sp.symbols('a drho')
		# s position as a function of alpha and rho
		sp_s = alpha + rho
		F_s = sp.lambdify((alpha,rho),sp_s,'numpy')
		# t position as a function of alpha and rho
		sp_t = a*alpha**2 - rho/(2*a*alpha)
		F_t = sp.lambdify((a,alpha,rho),sp_t,'numpy')
		#Rho step as a function of drho
		sp_rstep = sp.sqrt( drho*4*a**2*alpha**2/(1+4*a**2*alpha**2) )
		F_rstep = sp.lambdify((a,drho,alpha),sp_rstep,'numpy')

		alphas = np.arange(float(xmin),float(xmax),float(xstep))
		alphas[alphas == 0] = 0.1

		self.grid = pd.DataFrame({'alpha':[],'r':[],'s':[],'t':[]})
		for alph in alphas:
		    rst = F_rstep(acf,5,alph)
		    Lrho = np.array([0-sp.sign(alph)*rst*i for i in range(rmax) if rst*i <= abs(alph)]+[
		        sp.sign(alph)*rst*i for i in range(1,rmax) if rst*i <= rnum])
		    Lalpha = np.array([alph]*Lrho.shape[0])

		    Ls = F_s(Lalpha,Lrho)
		    Lt = F_t(acf,Lalpha,Lrho)

		    self.grid = self.grid.append(pd.DataFrame({'alpha':Lalpha,'r':Lrho,'s':Ls,'t':Lt}),ignore_index=True)

		self.tree = KDTree(self.grid[['s','t']])
		# return(self.grid,self.tree)

	def transform_cylindrical(self,df,xstep,rmax,rnum,mm):

		tic = time.time()
		from sympy.abc import alpha,rho
		a,ds,dt,x = sp.symbols('a ds dt x')
		sp_dalpha = alpha*(ds + 2*a*dt*alpha)/(alpha+4*a**2*alpha**3+rho)
		F_dalpha = sp.lambdify((a,alpha,rho,ds,dt),sp_dalpha,'numpy')

		self.setup_tree(df.x.min(),df.x.max(),xstep,rmax,rnum,mm.cf[0])
		print(time.time()-tic,'Tree setup complete')

		tic = time.time()
		# Find alpha position on parabola
		nbd,nbi = self.tree.query(df[['x','z']])
		tpt = df.loc[df.index]
		npt = self.grid.loc[nbi]
		ds = tpt.x - np.array(npt.s)
		dt = tpt.z - np.array(npt.t)
		alpha = F_dalpha(mm.cf[0],np.array(npt.alpha),np.array(npt.r),ds,dt) + np.array(npt.alpha)
		alpha = alpha.astype('float64')
		print(tic-time.time(),'Alpha query complete')

		tic = time.time()
		# Calculate arclength
		x1,x2 = sp.symbols('x1 x2')
		sp_integral = (1/(2*a))*( a*x*sp.sqrt((1 + 4*a**2*x**2)) + (1/2)*sp.log(2*a*x + sp.sqrt((1 + 4*a**2*x**2))) )
		sp_arclength = sp_integral.subs(x,x2) - sp_integral.subs(x,x1)
		F_arclength = sp.lambdify((a,x1,x2),sp_arclength,'numpy')
		df['ac'] = F_arclength(np.array([mm.cf[0]]*alpha.shape[0]),np.zeros(alpha.shape[0]),alpha)
		print(tic-time.time(),'Arclength complete')

		tic = time.time()
		# Calculate theta
		df['theta'] = np.arctan2(df.y-0,df.z-mm.cf[0]*np.power(alpha,2))
		print(tic-time.time(),'theta')

		tic = time.time()
		#Calculate r
		df['r'] = np.sqrt( (df.z-mm.cf[0]*np.power(alpha,2))**2 + (df.y-0)**2 + (df.x-alpha)**2 )
		print(tic-time.time(),'r')

		return(df)

class math_model:
	'''
	Object to contain attributes associated with the math model of a sample

	:param array model: Array of coefficients calculated by np.polyfit

	.. py:attribute:: math_model.cf

		Array of coefficients for the math model

	.. py:attribute:: math_model.p

		Poly1d function for the math model to allow calculation and plotting of the model
	'''

	def __init__(self,model):

		self.cf = model
		self.p = np.poly1d(model)
```
