## Supplementary material for "ΔSCOPE: A new method to quantify 3D biological structures and identify differences in zebrafish forebrain development": An archived copy of the DeltaSCOPE code repository.: process_after_gui.py

```
import deltascope as cranium
import multiprocessing as mp
import time
import os
import sys
from functools import partial
from random import randint
import numpy as np
import pandas as pd
import re

def transform_file(f,model=None,test=False,tri=False):

	print(f, 'starting')

	tic = time.time()

	s = cranium.brain()
	df = cranium.read_psi(f)
	if 'ac' not in df.columns:

		# if tri==True:
		# 	s.df_align = df[['x','y','z']]
		# 	pt1x = np.max(s.df_align.x)
		# 	pt1z = s.df_align[s.df_align.x == pt1x].z.values[0]
		# 	pt2x = np.min(s.df_align.x)
		# 	pt2z = s.df_align[s.df_align.x == pt2x].z.values[0]
		#
		# 	s.mm = cranium.math_model(np.polyfit([pt1x,0,pt2x],[pt1z,0,pt2z],2))
		#
		# 	if test==True:
		# 		return(s,[pt1x,pt2x],[pt1z,pt2z])

		# if tri==False:
		s.df_align = df[['x','y','z']]
		snum  = re.findall(r'\d+',f.split('.')[0])[0]
		s.mm = cranium.math_model(model.loc[int(snum)].values)
		num = int(snum)
			# return(s)

		# if test==False:
		s.transform_coordinates()

		cranium.write_data(f,s.df_align)

	print(f,'complete',time.time()-tic)


if __name__=='__main__':

	outdirs = ['D:\\HSs1a_yot_expt\\zrf1\\hss1a\\Output-08-13']

	for outdir in outdirs:

		files = os.listdir(outdir)
		files.remove('model.csv')

		os.chdir(outdir)

		model = pd.read_csv(os.path.join(outdir,'model.csv'),index_col='Unnamed: 0')
		transform = partial(transform_file,model=model)

		n = len(files)
		for i in range(0,n,5):
			if i+5>n:
				L = files[i:n]
			else:
				L = files[i:i+5]

			pool = mp.Pool()
			pool.map(transform,L)
			pool.close()
			pool.join()

		print('Processing complete')
```
