## Supplementary material for "ΔSCOPE: A new method to quantify 3D biological structures and identify differences in zebrafish forebrain development": An archived copy of the DeltaSCOPE code repository.: README.md

# Deltascope scripts and modules

`__init__.py`  
This module can be called directly by importing deltascope. It contains the bulk of the functions needed for transforming and analyzing the data.

`mpTransformation.py`    
	This is a script to batch process multiple raw probability files from the terminal shell. It runs 5 samples at a time in parallel, so we see an increase in speed. There are a few functions defined here that facilitate batch processing in parallel. This line `if __name__=='__main__':` determines whether the script acts like a python script or as a module. If you import `mpTransformation` `__name__ != '__main__'` so it will act like a module that allows you to call the functons defined in the first part of the file. If you run the script via `python mpTransformation` in the shell, `__name__ == '__main__'` and the script will execute the code below to batch process samples.

`new.py`  
	This module is a copy of `__init__.py` where I tried to modify the transformation functions to increase speed. It was generally disasterous, but I wanted to keep the file as a record.

`process_after_gui.py`  
	This script follows the same convention as `mpTransformation.py` by including a conditional statement to determine if it should behave as a module or a script. This code accompanied the scripts in the `gui` folder.  The gui allows rudimentar manual curation of commisure alignments and then saves the untransformed data to psi files. This script reads in the already aligned data from psi files and converts the coordinate system. For most purposes, this script will not be useful unless someone want to revive the gui. Generally, the alignment process implemented in jupyter notebooks replaces the need for a gui.

`test_transform.py`  
	I honestly can't remember why I wrote this code. It looks like I intended to run this script from the command line on a single file at a time. Maybe I was using it to benchmark speed...?

`utility.py`  
	The majority of this code was written to facilitate the manual alignment of commissures in jupyter notebooks. At some point, it could be appropriate to integrate this code into `__init__.py`, but I haven't done it because it will require going back and fixing the import statements in a bunch of notebooks.
