## Supplementary material for "ΔSCOPE: A new method to quantify 3D biological structures and identify differences in zebrafish forebrain development": An archived copy of the DeltaSCOPE code repository.: test_transform.py

```
import cranium

expname = 'test'
samplenum = '01'
directory = 'data'

#Create an embryo object that facilitates data processing
e = cranium.embryo(expname,samplenum,directory)

#For each channel in your sample, add a channel with a unique name, e.g. 'c1' or 'c2'
e.add_channel('data/C1/AT_01_Probabilities.h5','c1')
# e.add_channel('data/C2/ZRF1_01_Probabilities.h5','c2')

genthresh = 0.5
scale = [1,1,1]
microns = [0.16,0.16,0.21]

#Threshold each channel and scale points according to voxel dimension in microns
e.chnls['c1'].preprocess_data(genthresh,scale,microns)
# e.chnls['c2'].preprocess_data(genthresh,scale,microns)

medthresh = 0.25
radius = 20
microns = [0.16,0.16,0.21]

#Run PCA on the structural channel, in this case, c1
e.chnls['c1'].calculate_pca_median(e.chnls['c1'].raw_data,medthresh,radius,microns)

#Save the pca object that includes the transformation matrix
pca = e.chnls['c1'].pcamed

comporder = [0,2,1]
fitdim = ['x','z']

#Transform the structural channel using the saved pca object
e.chnls['c1'].pca_transform_3d(e.chnls['c1'].df_thresh,pca,comporder,fitdim,deg=2)

#Save the mathematical model and vertex (center point) of the structural channel
mm = e.chnls['c1'].mm
vertex = e.chnls['c1'].vertex

#Transform any additional channels using the pca object calculated based on the structural channel
# e.chnls['c2'].pca_transform_3d(e.chnls['c2'].df_thresh,pca,comporder,fitdim,deg=2,mm=mm,vertex=vertex)

#Transform each channel to cylindrical coordinates
e.chnls['c1'].transform_coordinates()
# e.chnls['c2'].transform_coordinates()

#Save processed data to .psi file
# e.save_psi()
```
