## Supplementary material for "ΔSCOPE: A new method to quantify 3D biological structures and identify differences in zebrafish forebrain development": An archived copy of the DeltaSCOPE code repository.: cranium.html

  


cranium — cranium 0.1.0 documentation


cranium

0.1.0

- Data Preparation
- Data Processing
- Checklist
- Parameter Reference
- API

cranium

- Docs »
- Module code »
- cranium

---

### Source code for cranium

```
import os
import numpy as np
import h5py
#import skimage.io as io
import matplotlib.pyplot as plt
import time
import itertools as it
import matplotlib as mpl
import multiprocessing as mp
from functools import partial
#import statsmodels.formula.api as smf
#from mpl_toolkits.mplot3d import Axes3D
import pandas as pd
#import plotly.plotly as py
#import plotly.graph_objs as go
from scipy.optimize import minimize
import scipy
from sklearn.decomposition import PCA
from skimage.filters import median
from skimage.morphology import disk
from sklearn.metrics import mean_squared_error
from scipy.integrate import simps
import scipy.stats as stats

[docs]class brain:
	''' Object to manage biological data and associated functions. '''

	def __init__(self):
		'''Initialize brain object'''

[docs]	def read_data(self,filepath):
		'''
		Reads 3D data from file and selects appropriate channel based on the assumption that the channel with the most zeros has zero as the value for no signal

		:param str filepath: Filepath to hdf5 probability file
		:return: Creates the variable :attr:`brain.raw_data`

		.. py:attribute:: brain.raw_data

			Array of shape [z,y,x] containing raw probability data
		'''

		#Read h5 file and extract probability data
		f = h5py.File(filepath,'r')

		#Select either channe0/1 or exported_data
		d = f.get('exported_data')
		if d != None:
			c1 = np.array(d[:,:,:,0])
			c2 = np.array(d[:,:,:,1])
		elif d == None:
			c1 = np.array(f.get('channel0'))
			c2 = np.array(f.get('channel1'))

		#Figure out which channels has more zeros and therefore is background
		if np.count_nonzero(c1<0.1) > np.count_nonzero(c1>0.9):
			#: Array of shape [z,y,x] containing raw probability data
			self.raw_data = c1
		else:
			#: Array of shape [z,y,x] containing raw probability data
			self.raw_data = c2


[docs]	def create_dataframe(self,data,scale):
		'''
		Creates a pandas dataframe containing the x,y,z and signal/probability value for each point in the :py:attr:`brain.raw_data` array

		:param array data: Raw probability data in 3D array
		:param array scale: Array of length three containing the micron values for [x,y,z]
		:return: Pandas DataFrame with xyz and probability value for each point
		'''

		#NB: scale variable actually contains microns dimensions

		dim = data.shape
		xyz = np.zeros((dim[0],dim[1],dim[2],4))

		#Generate array with xyz values for each point
		for x in range(dim[2]):

			xyz[:,:,x,2] = x
			xyz[:,:,x,3] = data[:,:,x]

			zindex = np.arange(0,dim[0],1)
			yindex = np.arange(0,dim[1],1)
			gy,gz = np.meshgrid(yindex,zindex)

			xyz[:,:,x,0] = gz
			xyz[:,:,x,1] = gy

		flat = np.reshape(xyz,(-1,4))

		#Create dataframe of points and scale to microns
		df = pd.DataFrame({'x':flat[:,2]*scale[0],'y':flat[:,1]*scale[1],'z':flat[:,0]*scale[2],'value':flat[:,3]})
		return(df)


[docs]	def plot_projections(self,df,subset):
		'''
		Plots the x, y, and z projections of the input dataframe in a matplotlib plot

		:param pd.DataFrame df: Dataframe with columns: 'x','y','z'
		:param float subset: Value between 0 and 1 indicating what percentage of the df to subsample
		:returns: Matplotlib figure with three labeled scatterplots
		'''

		df = df.sample(frac=subset)
    
    	#Create figure and subplots
		fig = plt.figure(figsize=(12,6))
		ax = fig.add_subplot(131)
		ay = fig.add_subplot(132)
		az = fig.add_subplot(133)

		#Create scatter plot for each projection
		ax.scatter(df.x,df.z)
		ay.scatter(df.x,df.y)
		az.scatter(df.z,df.y)

		#Plot model
		xvalues = np.arange(np.min(df.x),np.max(df.x))
		ax.plot(xvalues,self.mm.p(xvalues),c='y')

		#Add labels
		ax.set_title('Y projection')
		ay.set_title('Z projection')
		az.set_title('X projection')
		ax.set_xlabel('X')
		ax.set_ylabel('Z')
		ay.set_xlabel('X')
		ay.set_ylabel('Y')
		az.set_xlabel('Z')
		az.set_ylabel('Y')

		#Adjust spacing and show plot
		plt.subplots_adjust(wspace=0.4)
		
		return(fig)


[docs]	def preprocess_data(self,threshold,scale,microns):
		'''
		Thresholds and scales data prior to PCA

		Creates :py:attr:`brain.threshold`, :py:attr:`brain.df_thresh`, and :py:attr:`brain.df_scl`

		:param float threshold: Value between 0 and 1 to use as a cutoff for minimum pixel value
		:param array scale: Array with three values representing the constant by which to multiply x,y,z respectively
		:param array microns: Array with three values representing the x,y,z micron dimensions of the voxel

		.. py:attribute:: brain.threshold 
			
			Value used to threshold the data prior to calculating the model

		.. py:attribute:: brain.df_thresh

			Dataframe containing only points with values above the specified threshold

		.. py:attribute:: brain.df_scl

			Dataframe containing data from :py:attr:`brain.df_thresh` after a scaling value has been applied
		'''

		#: Dataframe with four columns: x,y,z,value with all points in :py:attr:`brain.raw_data`
		self.df = self.create_dataframe(self.raw_data,microns)

		#Create new dataframe with values above threshold
		self.threshold = threshold
		self.df_thresh = self.df[self.df.value > self.threshold]

		#Scale xyz by value in scale array to force PCA axis selection
		self.scale = scale
		self.df_scl = pd.DataFrame({
			'x':self.df_thresh.x * self.scale[0], 
			'y':self.df_thresh.y * self.scale[1],
			'z':self.df_thresh.z * self.scale[2]})


[docs]	def process_alignment_data(self,data,threshold,radius,microns):
		'''
		Applies a median filter twice to the data which is used for alignment

		Ensures than any noise in the structural data does not interfere with alignment

		:param array data: Raw data imported by the function :py:func:`brain.read_data`
		:param float threshold: Value between 0 and 1 to use as a cutoff for minimum pixel value
		:param int radius: Integer that determines the radius of the circle used for the median filter
		:param array microns: Array with three values representing the x,y,z micron dimensions of the voxel
		:returns: Dataframe containing data processed with the median filter and threshold
		'''

		#Iterate over each plane and apply median filter twice
		out = np.zeros(data.shape)
		for z in range(data.shape[0]):
			out[z] = median(median(data[z],disk(radius)),disk(radius))

		outdf = self.create_dataframe(out,microns)
		thresh = outdf[outdf.value > threshold]
		return(thresh)


[docs]	def calculate_pca_median(self,data,threshold,radius,microns):
		'''
		Calculate PCA transformation matrix, :py:attr:`brain.pcamed`, based on data (:py:attr:`brain.pcamed`) after applying median filter and threshold

		:param array data: 3D array containing raw probability data
		:param float threshold: Value between 0 and 1 indicating the lower cutoff for positive signal
		:param int radius: Radius of neighborhood that should be considered for the median filter
		:param array microns: Array with three values representing the x,y,z micron dimensions of the voxel
		
		.. py:attribute:: brain.median

			Pandas dataframe containing data that has been processed with a median filter twice and thresholded

		.. py:attribute:: brain.pcamed

			PCA object managing the transformation matrix and any resulting transformations

		'''

		self.median = self.process_alignment_data(data,threshold,radius,microns)

		self.pcamed = PCA()
		self.pcamed.fit(self.median[['x','y','z']])


[docs]	def calculate_pca_median_2d(self,data,threshold,radius,microns):
		'''
		Calculate PCA transformation matrix for 2 dimensions of data, :py:attr:`brain.pcamed`, based on data after applying median filter and threshold

		.. warning:: `fit_dim` is not used to determine which dimensions to fit. Defaults to x and z

		:param array data: 3D array containing raw probability data
		:param float threshold: Value between 0 and 1 indicating the lower cutoff for positive signal
		:param int radius: Radius of neighborhood that should be considered for the median filter
		:param array microns: Array with three values representing the x,y,z micron dimensions of the voxel
		'''

		self.median = self.process_alignment_data(data,threshold,radius,microns)

		self.pcamed = PCA()
		self.pcamed.fit(self.median[['y','z']])


[docs]	def pca_transform_2d(self,df,pca,comp_order,fit_dim,deg=2,mm=None,vertex=None,flip=None):
		'''
		Transforms `df` in 2D based on the PCA object, `pca`, whose transformation matrix has already been calculated

		Calling :py:func:`brain.align_data` creates :py:attr:`brain.df_align`

		.. warning:: `fit_dim` is not used to determine which dimensions to fit. Defaults to x and z

		:param pd.DataFrame df: Dataframe containing thresholded xyz data
		:param pca_object pca: A pca object containing a transformation object, e.g. :py:attr:`brain.pcamed`
		:param array comp_order: Array specifies the assignment of components to x,y,z. Form [x component index, y component index, z component index], e.g. [0,2,1]
		:param array fit_dim: Array of length two containing two strings describing the first and second axis for fitting the model, e.g. ['x','z']
		:param int deg: (or None) Degree of the function that should be fit to the model. deg=2 by default
		:param mm: (:py:class:`math_model` or None) Math model for primary channel
		:param array vertex: (or None) Array of type [vx,vy,vz] (:py:attr:`brain.vertex`) indicating the translation values
		:param Bool flip: (or None) Boolean value to determine if the data should be rotated by 180 degrees
		'''

		fit = pca.transform(df[['y','z']])
		df_fit = pd.DataFrame({
			'x':df.x,
			'y':fit[:,comp_order[1]-1],
			'z':fit[:,comp_order[2]-1]
			})

		self.align_data(df_fit,fit_dim,deg=2,mm=None,vertex=None,flip=None)


[docs]	def pca_transform_3d(self,df,pca,comp_order,fit_dim,deg=2,mm=None,vertex=None,flip=None):
		'''
		Transforms `df` in 3D based on the PCA object, `pca`, whose transformation matrix has already been calculated

		:param pd.DataFrame df: Dataframe containing thresholded xyz data
		:param pca_object pca: A pca object containing a transformation object, e.g. :py:attr:`brain.pcamed`
		:param array comp_order: Array specifies the assignment of components to x,y,z. Form [x component index, y component index, z component index], e.g. [0,2,1]
		:param array fit_dim: Array of length two containing two strings describing the first and second axis for fitting the model, e.g. ['x','z']
		:param int deg: (or None) Degree of the function that should be fit to the model. deg=2 by default
		:param mm: (:py:class:`math_model` or None) Math model for primary channel
		:param array vertex: (or None) Array of type [vx,vy,vz] (:py:attr:`brain.vertex`) indicating the translation values
		:param Bool flip: (or None) Boolean value to determine if the data should be rotated by 180 degrees
		'''

		fit = pca.transform(df[['x','y','z']])
		df_fit = pd.DataFrame({
			'x':fit[:,comp_order[0]],
			'y':fit[:,comp_order[1]],
			'z':fit[:,comp_order[2]]
			})

		self.align_data(df_fit,fit_dim,deg=2,mm=None,vertex=None,flip=None)


[docs]	def align_data(self,df_fit,fit_dim,deg=2,mm=None,vertex=None,flip=None):
		'''
		Apply PCA transformation matrix and align data so that the vertex is at the origin

		Creates :py:attr:`brain.df_align` and :py:attr:`brain.mm`

		:param pd.DataFrame df: dataframe containing thresholded xyz data
		:param array comp_order: Array specifies the assignment of components to x,y,z. Form [x component index, y component index, z component index], e.g. [0,2,1]
		:param array fit_dim: Array of length two containing two strings describing the first and second axis for fitting the model, e.g. ['x','z']
		:param int deg: (or None) Degree of the function that should be fit to the model. deg=2 by default
		:param mm: (:py:class:`math_model` or None) Math model for primary channel
		:param array vertex: (or None) Array of type [vx,vy,vz] (:py:attr:`brain.vertex`) indicating the translation values
		:param Bool flip: (or None) Boolean value to determine if the data should be rotated by 180 degrees

		.. py:attribute:: brain.df_align

			Dataframe containing point data aligned using PCA

		.. py:attribute:: brain.mm

			Math model object fit to data in brain object
		'''
		
		#If vertex for translation is not included
		if vertex == None and deg==2:
			#Calculate model
			model = np.polyfit(df_fit[fit_dim[0]],df_fit[fit_dim[1]],deg=deg)
			p = np.poly1d(model)

			#Find vertex
			a = -model[1]/(2*model[0])
			if fit_dim[0] == 'x':
				vx = a
				if fit_dim[1] == 'y':
					vy = p(vx)
					vz = df_fit.z.mean()
				else:
					vz = p(vx)
					vy = df_fit.y.mean()
			elif fit_dim[0] == 'y':
				vy = a
				if fit_dim[1] == 'x':
					vx = p(a)
					vz = df_fit.z.mean()
				else:
					vz = p(a)
					vx = df_fit.x.mean()
			elif fit_dim[0] == 'z':
				vz = a
				if fit_dim[1] == 'x':
					vx = p(a)
					vy = df_fit.y.mean()
				else:
					vy = p(a)
					vx = df_fit.x.mean()
			self.vertex = [vx,vy,vz]
		elif deg == 1:
			#Calculate model
			model = np.polyfit(df_fit[fit_dim[0]],df_fit[fit_dim[1]],deg=deg)
			p = np.poly1d(model)

			if fit_dim[0] == 'x':
				vx = df_fit.x.mean()
				if fit_dim[1] == 'y':
					vy = p(vx)
					vz = df_fit.z.mean()
				else:
					vz = p(vx)
					vy = df_fit.y.mean()
			elif fit_dim[0] == 'y':
				vy = df_fit.y.mean()
				if fit_dim[1] == 'x':
					vx = p(vy)
					vz = df_fit.z.mean()
				else:
					vz = p(vy)
					vx = df_fit.x.mean()
			elif fit_dim[0] == 'z':
				vz = df_fit.z.mean()
				if fit_dim[1] == 'x':
					vx = p(vz)
					vy = df_fit.y.mean()
				else:
					vy = p(vz)
					vx = df_fit.x.mean()
			self.vertex = [vx,vy,vz]
		else:
			self.vertex = vertex

		#Translate data so that the vertex is at the origin
		self.df_align = pd.DataFrame({
			'x': df_fit.x - self.vertex[0],
			'y': df_fit.y - self.vertex[1],
			'z': df_fit.z - self.vertex[2]
			})

		#Rotate data by 180 degrees if necessary
		if flip == None or flip == False:
			#Calculate model
			model = np.polyfit(df_fit[fit_dim[0]],df_fit[fit_dim[1]],deg=deg)
			p = np.poly1d(model)

			#If a is less than 0, rotate data
			if model[0] < 0:
				self.df_align = self.flip_data(self.df_align)
		elif flip == True:
			self.df_align = self.flip_data(self.df_align)

		#Calculate final math model
		if mm == None:
			self.mm = self.fit_model(self.df_align,deg,fit_dim)
		else:
			self.mm = mm


[docs]	def flip_data(self,df):
		'''
		Rotate data by 180 degrees

		:param dataframe df: Pandas dataframe containing x,y,z data
		:returns: Rotated dataframe
		'''

		r = np.array([[np.cos(np.pi),0,np.sin(np.pi)],
			[0,1,0],
			[-np.sin(np.pi),0,np.cos(np.pi)]])

		rot = np.dot(np.array(df),r)

		dfr = pd.DataFrame({'x':rot[:,0],'y':rot[:,1],'z':rot[:,2]})
		return(dfr)


[docs]	def fit_model(self,df,deg,fit_dim):
		'''Fit model to dataframe

		:param pd.DataFrame df: Dataframe containing at least x,y,z
		:param int deg: Degree of the function that should be fit to the model
		:param array fit_dim: Array of length two containing two strings describing the first and second axis for fitting the model, e.g. ['x','z']
		:returns: math model
		:rtype: :py:class:`math_model`
		'''

		mm = math_model(np.polyfit(df[fit_dim[0]], df[fit_dim[1]], deg=deg))
		return(mm)

###### Functions associated with alpha, r, theta coordinate system ######

[docs]	def find_distance(self,t,point):
		'''
		Find euclidean distance between math model(t) and data point in the xy plane

		:param float t: float value defining point on the line
		:param array point: array [x,y] defining data point
		:returns: distance between the two points
		:rtype: float
		'''

		x = float(t)
		z = self.mm.p(x)

		#Calculate distance between two points passed as array
		dist = np.linalg.norm(point - np.array([x,z]))

		return(dist)


[docs]	def find_min_distance(self,row):
		'''
		Find the point on the curve that produces the minimum distance between the point and the data point using scipy.optimize.minimize(:py:func:`brain.find_distance`)

		:param pd.Series row: row from dataframe in the form of a pandas Series
		:returns: point in the curve (xc, yc, zc) and r
		:rtype: floats
		'''

		dpoint = np.array([row.x,row.z])

		#Use scipy.optimize.minimize to find minimum solution of brain.find_distance, 
		#dpoint[0] is starting guess for x value
		result = minimize(self.find_distance, dpoint[0], args=(dpoint))

		x = result['x'][0]
		y = 0
		z = self.mm.p(x)
		#r = result['fun']

		return(x,y,z)

#return(pd.Series({'xc':x, 'yc':y, 'zc':z, 'r':r}))

[docs]	def integrand(self,x):
		'''
		Function to integrate to calculate arclength

		:param float x: integer value for x
		:returns: arclength value for integrating
		:rtype: float
		'''

		y_prime = self.mm.cf[0]*2*x + self.mm.cf[1]

		arclength = np.sqrt(1 + y_prime**2)
		return(arclength)


[docs]	def find_arclength(self,xc):
		'''
		Calculate arclength by integrating the derivative of the math model in xy plane

		.. math:: 

			\int_{vertex}^{point} \sqrt{1 + (2ax + b)^2}

		:param float row: Postion in the x axis along the curve
		:returns: Length of the arc along the curve between the row and the vertex
		:rtype: float
		'''

		ac,err = scipy.integrate.quad(self.integrand,xc,0)
		return(ac)


[docs]	def find_theta(self,row,zc,yc):
		'''
		Calculate theta for a row containing data point in relationship to the xz plane

		:param pd.Series row: row from dataframe in the form of a pandas Series
		:param float yc: Y position of the closest point in the curve to the data point
		:param float zc: Z position of the closest point in the curve to the data point
		:returns: theta, angle between point and the model plane
		:rtype: float
		'''

		theta = np.arctan2(row.y-yc,row.z-zc)
		return(theta)


[docs]	def find_r(self,row,zc,yc):
		'''
		Calculate r using the Pythagorean theorem

		:param pd.Series row: row from dataframe in the form of a pandas Series
		:param float yc: Y position of the closest point in the curve to the data point
		:param float zc: Z position of the closest point in the curve to the data point
		:returns: r, distance between the point and the model
		:rtype: float
		'''

		r = np.sqrt((row.z-zc)**2 + (row.y-yc)**2)
		return(r)


[docs]	def calc_coord(self,row):
		'''
		Calculate alpah, r, theta for a particular row

		:param pd.Series row: row from dataframe in the form of a pandas Series
		:returns: pd.Series populated with coordinate of closest point on the math model, r, theta, and ac (arclength)
		'''

		xc,yc,zc = self.find_min_distance(row)
		ac = self.find_arclength(xc)
		theta = self.find_theta(row,zc,yc)
		r = self.find_r(row,zc,yc)

		return(pd.Series({'x':row.x,'y':row.y,'z':row.z,'xc':xc, 'yc':yc, 'zc':zc,
					'r':r, 'ac':ac, 'theta':theta}))


[docs]	def transform_coordinates(self):
		'''
		Transform coordinate system so that each point is defined relative to math model by (alpha,theta,r) (only applied to :py:attr:`brain.df_align`)

		:returns: appends columns r, xc, yc, zc, ac, theta to :py:attr:`brain.df_align`
		'''

		#Calculate alpha, theta, r for each row in dataset
		self.df_align = self.df_align.merge(self.df_align.apply((lambda row: self.calc_coord(row)), axis=1))


[docs]	def subset_data(self,df,sample_frac=0.5):
		'''
		Takes a random sample of the data based on the value between 0 and 1 defined for sample_frac
		
		Creates the variable :py:attr:`brain.subset`

		:param pd.DataFrame: Dataframe which will be sampled
		:param float sample_frac: (or None) Value between 0 and 1 specifying proportion of the dataset that should be randomly sampled for plotting
		
		.. py:attribute:: brain.subset

			Random sample of the input dataframe
		'''

		self.subset = df.sample(frac=sample_frac)


[docs]	def add_thresh_df(self,df):
		'''
		Adds dataframe of thresholded and transformed data to :py:attr:`brain.df_thresh`

		:param pd.DataFrame df: dataframe of thesholded and transformed data
		:returns: :py:attr:`brain.df_thresh`
		'''

		self.df_thresh = df


[docs]	def add_aligned_df(self,df):
		'''
		Adds dataframe of aligned data

		.. warning:: Calculates model, but assumes that the dimensions of the fit are x and z

		:param pd.DataFrame df: Dataframe of aligned data
		:returns: :py:attr:`brain.df_align`
		'''

		self.df_align = df
		self.mm = self.fit_model(self.df_align,2,['x','z'])


[docs]class embryo:
	'''
	Class to managed multiple brain objects in a multichannel sample

	:param str name: Name of this sample set
	:param str number: Sample number corresponding to this embryo
	:param str outdir: Path to directory for output files

	.. py:attribute:: embryo.chnls

		Dictionary containing the :py:class:`brain` object for each channel

	.. py:attribute:: embryo.outdir

		Path to directory for output files

	.. py:attribute:: embryo.name

		Name of this sample set

	.. py:attribute:: embryo.number

		Sample number corresponding to this embryo
	'''

	def __init__(self,name,number,outdir):
		'''Initialize embryo object'''

		self.chnls = {}
		self.outdir = outdir
		self.name = name
		self.number = number

[docs]	def add_channel(self,filepath,key):
		'''
		Add channel to :py:attr:`embryo.chnls` dictionary

		:param str filepath: Complete filepath to image
		:param str key: Name of the channel
		'''

		s = brain()
		s.read_data(filepath)

		self.chnls[key] = s


[docs]	def process_channels(self,mthresh,gthresh,radius,scale,microns,deg,primary_key,comp_order,fit_dim):
		'''
		Process all channels through the production of the :py:attr:`brain.df_align` dataframe

		:param float mthresh: Value between 0 and 1 to use as a cutoff for minimum pixel value for median data
		:param float gthresh: Value between 0 and 1 to use as a cutoff for minimum pixel value for general data
		:param int radius: Size of the neighborhood area to examine with median filter
		:param array scale: Array with three values representing the constant by which to multiply x,y,z respectively
		:param array microns: Array with three values representing the x,y,z micron dimensions of the voxel
		:param int deg: Degree of the function that should be fit to the model
		:param str primary_key: Key for the primary structural channel which PCA and the model should be fit too
		:param array comp_order: Array specifies the assignment of components to x,y,z. Form [x component index, y component index, z component index], e.g. [0,2,1]
		:param array fit_dim: Array of length two containing two strings describing the first and second axis for fitting the model, e.g. ['x','z']
		'''

		#Process primary channel
		self.chnls[primary_key].preprocess_data(gthresh,scale,microns)

		self.chnls[primary_key].calculate_pca_median(self.chnls[primary_key].raw_data,
			mthresh,radius,microns)
		self.pca = self.chnls[primary_key].pcamed

		self.chnls[primary_key].align_data(self.chnls[primary_key].df_thresh,
			self.pca,comp_order,fit_dim,deg=deg)
		self.mm = self.chnls[primary_key].mm
		self.vertex = self.chnls[primary_key].vertex

		self.chnls[primary_key].transform_coordinates()

		print('Primary channel',primary_key,'processing complete')

		for ch in self.chnls.keys():
			if ch != primary_key:
				self.chnls[ch].preprocess_data(gthresh,scale,microns)
				
				self.chnls[ch].align_data(self.chnls[ch].df_thresh,
					self.pca,comp_order,fit_dim,deg=deg,
					mm = self.mm, vertex = self.vertex)

				self.chnls[ch].transform_coordinates()
				print(ch,'processed')


[docs]	def save_projections(self,subset):
		'''
		Save projections of both channels into png files in :py:attr:`embryo.outdir` following the naming scheme [:py:attr:`embryo.name`]_[:py:attr:`embryo.number`]_[`channel name`]_MIP.png

		:param float subset: Value between 0 and 1 to specify the fraction of the data to randomly sample for plotting
		'''

		for ch in self.chnls.keys():
			fig = self.chnls[ch].plot_projections(self.chnls[ch].df_align,subset)
			fig.savefig(os.path.join(self.outdir,
				self.name+'_'+self.number+'_'+ch+'_MIP.png'))

		print('Projections generated')


[docs]	def save_psi(self):
		'''
		Save all channels into psi files following the naming scheme [:py:attr:`embryo.name`]_[:py:attr:`embryo.number`]_[`channel name`].psi
		'''

		columns = ['x','y','z','ac','r','theta']

		for ch in self.chnls.keys():
			write_data(os.path.join(self.outdir,
				self.name+'_'+self.number+'_'+ch+'.psi'),
				self.chnls[ch].df_align[columns])

		print('PSIs generated')


[docs]	def add_psi_data(self,filepath,key):
		'''
		Read psi data into a channel dataframe

		:param str filepath: Complete filepath to data
		:param str key: Descriptive key for channel dataframe in dictionary
		'''

		self.chnls[key] = read_psi(filepath)


[docs]class math_model:
	'''
	Object to contain attributes associated with the math model of a sample

	:param array model: Array of coefficients calculated by np.polyfit

	.. py:attribute:: math_model.cf

		Array of coefficients for the math model

	.. py:attribute:: math_model.p

		Poly1d function for the math model to allow calculation and plotting of the model
	'''

	def __init__(self,model):

		self.cf = model
		self.p = np.poly1d(model)


[docs]class landmarks:
	'''
	Class to handle calculation of landmarks to describe structural data

	:param list percbins: (or None) Must be a list of integers between 0 and 100 
	:param int rnull: (or None) When the r value cannot be calculated it will be set to this value
	
	.. py:attribute:: brain.lm_wt_rf

		pd.DataFrame, which wildtype landmarks will be added to

	.. py:attribute:: brain.lm_mt_rf

		pd.DataFrame, which mutant landmarks will be added to

	.. py:attribute:: brain.rnull

		Integer specifying the value which null landmark calculations will be set to
	
	.. py:attribute:: brain.percbins

		Integer specifying the percentiles which will be used to calculate landmarks
	'''

	def __init__(self,percbins=[10,50,90],rnull=15):

		self.lm_wt_rf = pd.DataFrame()
		self.lm_mt_rf = pd.DataFrame()

		self.rnull = rnull
		self.percbins = percbins

[docs]	def calc_bins(self,Ldf,ac_num,tstep):
		'''
		Calculates alpha and theta bins based on ac_num and tstep

		Creates :py:attr:`landmarks.acbins` and :py:attr:`landmarks.tbins`

		.. warning:: `tstep` does not handle scenarios where 2pi is not evenly divisible by tstep

		:param list Ldf: List of dataframes that are being used for the analysis typically accessed by `dict.values()`
		:param int ac_num: Integer indicating the number of divisions that should be made along alpha
		:param float tstep: The size of each bin used for alpha

		.. py:attribute:: landmarks.acbins

			List containing the boundaries of each bin along alpha based on `ac_num`

		.. py:attribute:: landmarks.tbins

			List containing the boundaries of each bin along theta based on `tstep`
		'''

		#Find the minimum and maximum values of alpha in the dataset
		acmin,acmax = 0,0
		for df in Ldf:
			if df.ac.min() < acmin:
				acmin = df.ac.min()
			if df.ac.max() > acmax:
				acmax = df.ac.max()

		#Correct min and max values so that arclength is equal on each side
		if abs(acmin) > acmax:
			self.acbins = np.linspace(acmin,abs(acmin),ac_num)
		else:
			self.acbins = np.linspace(-acmax,acmax,ac_num)

		#Calculate tbins divisions based on tstep
		self.tbins = np.arange(-np.pi,np.pi+tstep,tstep)


[docs]	def calc_perc(self,df,snum,dtype,out):
		'''
		Calculate landmarks for a dataframe based on the bins and percentiles that have been previously defined 

		:param pd.DataFrame df: Dataframe containing columns x,y,z,alpha,r,theta
		:param str snum: String containing a sample identifier that can be converted to an integer
		:param str dtype: String describing the sample group to which the sample belongs, e.g. control or experimental
		:returns: pd.DataFrame with new landmarks appended
		'''

		D = {'stype':dtype}

		#Go through each arclength bins
		for a in range(len(self.acbins)):
			if a+1 < len(self.acbins):
				arange = [self.acbins[a],self.acbins[a+1]]

				#Go through each theta bin
				for t in range(len(self.tbins)):
					if t+1 < len(self.tbins):
						trange = [self.tbins[t],self.tbins[t+1]]

						#Go through each percentile
						for p in self.percbins:
							d = df[(df.ac > arange[0]) & (df.ac < arange[1])]
							d = d[(d.theta > trange[0]) & (d.theta < trange[1])]

							#Try to calculate percentile, but set to null if it fails
							try:
								r = np.percentile(d.r,p)
								pts = d[d.r < r].count()['i']
							except:
								r = self.rnull
								pts = 0

							#Create name of column based on bins
							L = []
							for s in [arange[0],arange[1],trange[0],trange[1]]:
								L.append(str(np.around(s,decimals=2)))
							name = '_'.join(L)

							D[name+'_'+str(p)+'_pts'] = pts
							D[name+'_'+str(p)+'_r'] = r

		out = out.append(pd.Series(D,name=int(snum)))
		return(out)


[docs]	def calc_wt_reformat(self,df,snum):
		'''
		.. warning:: Deprecated function, but includes code pertaining to calculating point based data
		'''

		D = {'stype':'wildtype'}

		for a in range(len(self.acbins)):
			if a+1 < len(self.acbins):
				arange = [self.acbins[a],self.acbins[a+1]]

				for t in range(len(self.tbins)):
					if t+1 < len(self.tbins):
						trange = [self.tbins[t],self.tbins[t+1]]

						for p in self.percbins:
							d = df[(df.ac > arange[0]) & (df.ac < arange[1])]
							d = d[(d.theta > trange[0]) & (d.theta < trange[1])]

							try:
								r = np.percentile(d.r,p)
								pts = d[d.r < r].count()['i']
							except:
								r = self.rnull
								pts = 0

							L = []
							for s in [arange[0],arange[1],trange[0],trange[1]]:
								L.append(str(np.around(s,decimals=2)))
							name = '_'.join(L)

							D[name+'_'+str(p)+'_pts-pts'] = pts
							D[name+'_'+str(p)+'_perc-pts'] = pts
							D[name+'_'+str(p)+'_pts-r'] = r
							D[name+'_'+str(p)+'_perc-r'] = r

		self.lm_wt_rf = self.lm_wt_rf.append(pd.Series(D,name=int(snum)))


[docs]	def calc_mt_landmarks(self,df,snum,wt):
		'''
		.. warning:: Deprecated function, but attempted to calculate mutant landmarks based on the number of points found in the wildtype standard
		'''

		D = {'stype':'mutant'}

		for c in wt.columns:
			if len(c.split('_')) == 6:
				amn,amx,tmn,tmx,p,dtype = c.split('_')
				p = int(p)

				# print(c)
				# print(df.count()['i'])
				d = df[(df.ac > float(amn))&(df.ac < float(amx))]
				# print(d.count()['i'])
				d = d[(d.theta > float(tmn))&(d.theta < float(tmx))]
				# print(d.count()['i'])

				if dtype.split('-')[0] == 'perc':
					try:
						perc_r = np.percentile(d.r,p)
						perc_pts = d[d.r < perc_r]
					except:
						perc_r = self.rnull
						perc_pts = 0

					if dtype.split('-')[1] == 'r':
						D[c] = perc_r
					else:
						D[c] = perc_pts

				else:
					if dtype.split('-')[1] == 'pts':
						#Calculate precent of points by dividing pts by total pts
						p = wt[c].mean()/d.count()['i']
						try:
							pt_r = np.percentile(d.r,p)
						except:
							pt_r = self.rnull
						D[c] = pt_r
					else:
						D[c] = np.nan

		self.lm_mt_rf = self.lm_mt_rf.append(pd.Series(D,name=int(snum)))


[docs]def reformat_to_cart(df):
	'''
	Take a dataframe in which columns contain the bin parameters and convert to a cartesian coordinate system

	:param pd.DataFrame df: Dataframe containing columns with string names that contain the bin parameter
	:returns: pd.DataFrame with each landmark as a row and columns: x,y,z,r,r_std,t,pts
	'''

	ndf = pd.DataFrame()
	for c in df.columns:
		if len(c.split('_')) == 6:
			amn,amx,tmn,tmx,p,dtype = c.split('_')
			x = np.mean([float(amn),float(amx)])
			t = np.mean([float(tmn),float(tmx)])

			if dtype == 'r':
				r = np.mean(df[c])
				r_std = stats.sem(df[c])
				y = np.sin(t)*r
				z = np.cos(t)*r

				pts = np.mean(df['_'.join([amn,amx,tmn,tmx,p,'pts'])])

				D = pd.Series({'x':x,'y':y,'z':z,'r':r,'r_sem':r_std,'t':t,'pts':pts})

				ndf = ndf.append(pd.Series(D),ignore_index=True)

	return(ndf)

def convert_to_arr(xarr,tarr,wt,mt):
	wtarr = np.zeros((len(xarr),len(tarr),wt.count(axis=0)['Unnamed: 0']))
	mtarr = np.zeros((len(xarr),len(tarr),mt.count(axis=0)['Unnamed: 0']))

	for c in mt.columns:
		if len(c.split('_')) == 6:
			amn,amx,tmn,tmx,p,dtype = c.split('_')
			x = np.mean([float(amn),float(amx)])
			t = np.mean([float(tmn),float(tmx)])

			if dtype=='r':
				wtarr[np.where(xarr==x)[0],np.where(tarr==t)[0]] = wt[c]
				mtarr[np.where(xarr==x)[0],np.where(tarr==t)[0]] = mt[c]

	return(wtarr,mtarr)

P = {
	'zln':2,'zpt':3,'zfb':1,
	'wtc':'b','mtc':'r',
	'alpha':0.3,
	'cmap':'Greys_r',
	'xarr':None,
	'tarr':None
}

[docs]def subplot_lmk(ax,p,avg,sem,parr,xarr,tarr,dtype,Pn=P):
	'''
	Plot a ribbon of average and standard error of the mean onto the subplot, `ax`

	:param plt.Subplot ax: Matplotlib subplot onto which the data should be plotted
	:param list p: List of two theta values that should be plotted
	:param np.array avg: Array of shape (xvalues,tvalues) containing the average values of the data
	:param np.array sem: Array of shape (xvalues,tvalues) containing the standard error of the mean values of the data
	:param np.array parr: Array of shape (xvalues,tvalues) containing the p values for the data
	:param str dtype: String describing sample type
	:param Pn: Dictionary containing the following values: 'zln':2,'zpt':3,'zfb':1,'wtc':'b','mtc':'r','alpha':0.3,'cmap':'Greys_r'
	:type: dict or None
	'''

	for k in Pn.keys():
		P[k] = Pn[k]

	if dtype=='wt':
		c = P['wtc']
	else:
		c = P['mtc']

	ti1 = np.where(tarr==p[0])[0][0]
	ti2 = np.where(tarr==p[1])[0][0]

	ax.fill_between(xarr,avg[:,ti1]+sem[:,ti1],avg[:,ti1]-sem[:,ti1],alpha=P['alpha'],color=c,zorder=P['zfb'])
	ax.fill_between(xarr,-avg[:,ti2]+sem[:,ti2],-avg[:,ti2]-sem[:,ti2],alpha=P['alpha'],color=c,zorder=P['zfb'])

	ax.plot(xarr,avg[:,ti1],c=c,zorder=P['zln'])
	ax.plot(xarr,-avg[:,ti2],c=c,zorder=P['zln'])

	if dtype=='mt':
		ax.scatter(xarr,avg[:,ti1],c=parr[:,ti1],cmap=P['cmap'],zorder=P['zpt'])
		ax.scatter(xarr,-avg[:,ti2],c=parr[:,ti2],cmap=P['cmap'],zorder=P['zpt'])

##### PSI file processing ############

[docs]def write_header(f):
	'''
	Writes header for PSI file with columns Id,x,y,z,ac,r,theta

	:param file f: file object created by 'open(filename,'w')`
	'''

	contents = [
		'PSI Format 1.0',
		'',
		'column[0] = "Id"',
		'column[1] = "x"',
		'column[2] = "y"',
		'column[3] = "z"',
		'column[4] = "ac"',
		'symbol[4] = "A"',
		'type[4] = float',
		'column[5] = "r"',
		'symbol[5] = "R"',
		'type[5] = float',
		'column[6] = "theta"',
		'symbol[6] = "T"',
		'type[6] = float'
	]

	for line in contents:
		f.write('# '+ line + '\n')


[docs]def write_data(filepath,df):
	'''
	Writes data in PSI format to file after writing header using :py:func:`write_header`. Closes file at the conclusion of writing data.

	:param str filepath: Complete filepath to output file
	:param pd.DataFrame df: dataframe containing columns x,y,z,ac,r,theta
	'''

	#Open new file at given filepath
	f = open(filepath,'w')

	#Write header contents to file
	write_header(f)

	n = df.count()['x']
    
	#Write line with sample number
	f.write(str(n)+' 0 0\n')

	#Write translation matrix
	f.write('1 0 0\n'+
			'0 1 0\n'+
			'0 0 1\n')

	#Write dataframe to file using pandas to_csv function to format
	try:
		f.write(df[['x','y','z','ac','theta','r']].to_csv(sep=' ', index=True, header=False))
	except:
		f.write(df[['x','y','z']].to_csv(sep=' ', index=True, header=False))

	f.close()

	print('Write to',filepath,'complete')


[docs]def read_psi(filepath):
	'''
	Reads psi file at the given filepath and returns data in a pandas DataFrame

	:param str filepath: Complete filepath to file
	:returns: pd.Dataframe containing data
	'''

	df = pd.read_csv(filepath,
		sep=' ',
		header=19)

	if len(df.columns) <= 4:
		df.columns = ['i','x','y','z']
	elif len(df.columns) >= 6:
		df.columns=['i','x','y','z','ac','theta','r']

	return(df)


[docs]def read_psi_to_dict(directory,dtype):
	'''
	Read psis from directory into dictionary of dfs with filtering based on dtype

	:param str directory: Directory to get psis from 
	:param str dtype: Usually 'AT' or 'ZRF1'
	:returns: Dictionary of pd.DataFrame
	'''

	dfs = {}
	for f in os.listdir(directory):
		if dtype in f:
			df = read_psi(os.path.join(directory,f))
			num = f.split('_')[-1].split('.')[0]
			print(num)
			dfs[num] = df

	return(dfs)

###### Stand alone functions

[docs]def process_sample(num,root,outdir,name,chs,prefixes,threshold,scale,deg,primary_key,comp_order,fit_dim,flip_dim):
	'''
	Process single sample through :py:class:`brain` class and saves df to csv

	.. warning:: Out of date and will probably fail

	:param str num: Sample number
	:param str root: Complete path to the root directory for this sample set
	:param str name: Name describing this sample set
	:param str outdir: Complete path to output directory
	:param array chs: Array containing strings specifying the directories for each channel
	:param array prefixes: Array containing strings specifying the file prefix for each channel
	:param float threshold: Value between 0 and 1 to use as a cutoff for minimum pixel value
	:param array scale: Array with three values representing the constant by which to multiply x,y,z respectively
	:param int deg: Degree of the function that should be fit to the model
	:param str primary_key: Key for the primary structural channel which PCA and the model should be fit too
	'''

	tic = time.time()

	print(num,'Starting',name)

	e = embryo(name,num,outdir)
	e.add_channel(os.path.join(root,chs[0],prefixes[0]+'_'+num+'_Probabilities.h5'),
		'at')
	e.add_channel(os.path.join(root,chs[1],prefixes[1]+'_'+num+'_Probabilities.h5'),'zrf1')
	e.process_channels(threshold,scale,deg,primary_key,[0,2,1],['x','z'],'z')
	
	print(num,'Data processing complete',name)

	e.save_projections(0.1)
	e.save_psi()

	toc = time.time()
	print(name,'complete',toc-tic)


[docs]def calculate_models(Ldf):
	'''
	Calculate model for each dataframe in list and add to new dataframe

	:param list Ldf: List of dataframes containing aligned data
	:returns: pd.Dataframe with a,b,c values for parabolic model
	'''

	modeldf = pd.DataFrame({'a':[],'b':[],'c':[]})

	for df in Ldf:
		s = cranium.brain()
		s.add_aligned_df(df)
		s.fit_model(s.df_align,2)

		modeldf = modeldf.append(pd.DataFrame({'a':[s.mm.cf[0]],'b':[s.mm.cf[1]],'c':[s.mm.cf[2]]}))

	return(modeldf)


[docs]def generate_kde(data,var,x,absv=False):
	'''
	Generate list of KDEs from either dictionary or list of data

	:param data: pd.DataFrames to convert
	:type: dict or list
	:param str var: Name of column to select from df
	:param array x: Array of datapoints to evaluate KDE on
	:param bool absv: (or None) Set to True to use absolute value of selected data for KDE calculation
	:returns: List of KDE arrays
	'''

	L = []

	#Dictionary workflow
	if type(data) ==  type({}):
		for key in data.keys():
			if absv == False:
				y = data[key][var].sample(frac=0.1)
			elif absv == True:
				y = np.abs(data[key][var].sample(frac=0.1))
			kde = scipy.stats.gaussian_kde(y).evaluate(x)
			L.append(kde)
	#List workflow
	elif type(data) == type([]):
		for df in data:
			if absv == False:
				y = df[var].sample(frac=0.1)
			elif absv == True:
				y = np.abs(df[var].sample(frac=0.1))
			kde = scipy.stats.gaussian_kde(y).evaluate(x)
			L.append(kde)

	return(L)


[docs]def calculate_area_error(pdf,Lkde,x):
	'''
	Calculate area between PDF and each kde in Lkde

	:param array pdf: Array of probability distribution function that is the same shape as kdes in Lkde
	:param list Lkde: List of arrays of Kdes 
	:param array x: Array of datapoints used to generate pdf and kdes
	:returns: List of error values for each kde in Lkde
	'''

	L = []

	for kde in Lkde:
		L.append(simps(np.abs(pdf-kde),x))

	return(L)


[docs]def rescale_variable(Ddfs,var,newvar):
	'''
	Rescale variable from -1 to 1 and save in newvar column on original dataframe

	:param dict Ddfs: Dictionary of pd.DataFrames
	:param str var: Name of column to select from dfs
	:param str newvar: Name to use for new data in appended column
	:returns: Dictionary of dataframes containing column of rescaled data
	'''

	Dout = {}

	for key in Ddfs.keys():
		df = Ddfs[key]
		y = df[var]

		#Find min and max values
		mi,ma = y.min(),y.max()

		#Divide dataframe
		dfmin = df[y < 0]
		dfmax = df[y >= 0]

		#Add rescaled variable to dataframe
		dfmin = dfmin.join(pd.DataFrame({newvar:y/np.abs(mi)}))
		dfmax = dfmax.join(pd.DataFrame({newvar:y/np.abs(ma)}))

		dfout = dfmin.append(dfmax)
		Dout[key] = dfout

	return(Dout)
```

---

© Copyright 2018, Morgan Schwartz.

Built with Sphinx using a theme provided by Read the Docs.
