## Supplementary material for "ΔSCOPE: A new method to quantify 3D biological structures and identify differences in zebrafish forebrain development": An archived copy of the DeltaSCOPE code repository.: index.rst.txt

.. cranium documentation master file, created by
   sphinx-quickstart on Thu Jan  4 13:33:50 2018.
   You can adapt this file completely to your liking, but it should at least
   contain the root `toctree` directive.

Welcome to cranium's documentation!
===================================

Contents:

.. toctree::
	:numbered:

	Data Preparation <data-prep>
    Data Processing <data-process>
    Checklist <checklist>
    Parameter Reference <param-ref>
    API

Is cranium right for you?
++++++++++++++++++++++++++

Can dos:
* Align single 3D microscopy collections to an orientation that is consistent between samples

Cannot:
* Align cryosections to create a 3D volume from 2D slices

Issues
+++++++


Indices and tables
==================

* :ref:`genindex`
* :ref:`modindex`
* :ref:`search`
