## Supplementary material for "ΔSCOPE: A new method to quantify 3D biological structures and identify differences in zebrafish forebrain development": An archived copy of the DeltaSCOPE code repository.: param-ref.rst.txt

.. _param ref:

Parameter Reference
====================

.. currentmodule:: cranium

.. envvar:: genthresh
		
	This parameter defines the cutoff point that will divide the :attr:`brain.raw_data` into a set of true signal points and background points based on the probability that each point is true signal. The :file:`_Probabilties.h5` dataset is generated after running the Ilastik pixel classification workflow described :ref:`here <Signal Normalization>`. Pixels with a value of 1 are likely to be background. Correspondingly, pixels with a value close to 0 are most likely to be true signal. We have found that a threshold of 0.5 is sufficient to divide true signal from background; however, if your data contains a lot of intermediate background values (0.4-0.7), you may benefit from a smaller threshold, e.g. 0.3.

	Recommended: ``0.5``

.. envvar:: microns

	This parameter is a list of the form :samp:`[{x},{y},{z}]` that specifies the dimensions of the voxel in microns. The data in :attr:`brain.raw_data` is scaled by :envvar:`microns` to control for datasets in which the ``z`` dimension is larger than the ``x`` and ``y`` dimensions.

	Example: ``[0.16,0.16,0.21]``

.. envvar:: scale

	This is a deprecated parameter that should always be set to ``[1,1,1]``.

	Required: ``[1,1,1]``

.. envvar:: medthresh

	This parameter serves the same purpose as :envvar:`genthresh`; however, it is used exclusively on data used for aligning samples using PCA. This threshold is typically more stringent than :envvar:`genthresh` to ensure that any noise in the data does not interfere with the alignment process.

	Recommended: ``0.25``

.. envvar:: radius

	The :envvar:`radius` of the median filter can  be tuned to eliminate noisy signal. The typical value for :envvar:`radius` is 20, which refers to the number of neighboring points that are considered in the median filter. A smaller value for :envvar:`radius` will preserve small variation in signal, while a larger value will cause even more blunting and smoothing of the data.

	Recommended: ``20``

.. envvar:: comporder

	This parameter controls how principle components are reassigned to the typical Cartesian coordinate system (XYZ) that most users are familiar with. It takes the form of an array of length 3 that specifies the index of the component that will be assigned to the X, Y, or Z axis: :samp:`[{x index},{y index},{z index}]`. Please note that the index that matches each principle component starts counting at 0, e.g. 1st PC = 0, 2nd PC = 1, and 3rd PC = 2.

	For example, if we want to assign the 1st PC to the x axis, the 2nd to the Z axis, and the 3rd to the y axis, the :envvar:`comporder` parameter would be :samp:`[0,2,1]`. 

	Example: ``[0,2,1]``

.. envvar:: fitdim

	This parameter determines which 2 axes will be used to fit the 2D model. It takes the form of a list of 2 of the 3 dimensions specified as a lowercase string, e.g. ``'x','y','z'``. 

	If we wanted to fit a model in the XZ plane, while holding the Y axis constant, the :envvar:`fitdim` parameter would be ``['x','z']``.

	Example: ``['x','z']``

.. envvar:: deg

	This parameter specifies the degree of the function that will be fit to the data. The default is ``2``, which specifies a parabolic function. A deg of ``1`` would fit a linear function.

	Default: ``2``

	.. warning:: The infrastructure to support degrees other than 2 is not currently in place. Check `here <https://github.com/msschwartz21/craniumPy/issues/23>`_ for updates.
