## Supplementary material for "ΔSCOPE: A new method to quantify 3D biological structures and identify differences in zebrafish forebrain development": An archived copy of the DeltaSCOPE code repository.: jquery-1.11.1.js

```
/*!
 * jQuery JavaScript Library v1.11.1
 * http://jquery.com/
 *
 * Includes Sizzle.js
 * http://sizzlejs.com/
 *
 * Copyright 2005, 2014 jQuery Foundation, Inc. and other contributors
 * Released under the MIT license
 * http://jquery.org/license
 *
 * Date: 2014-05-01T17:42Z
 */

(function( global, factory ) {

	if ( typeof module === "object" && typeof module.exports === "object" ) {
		// For CommonJS and CommonJS-like environments where a proper window is present,
		// execute the factory and get jQuery
		// For environments that do not inherently posses a window with a document
		// (such as Node.js), expose a jQuery-making factory as module.exports
		// This accentuates the need for the creation of a real window
		// e.g. var jQuery = require("jquery")(window);
		// See ticket #14549 for more info
		module.exports = global.document ?
			factory( global, true ) :
			function( w ) {
				if ( !w.document ) {
					throw new Error( "jQuery requires a window with a document" );
				}
				return factory( w );
			};
	} else {
		factory( global );
	}

// Pass this if window is not defined yet
}(typeof window !== "undefined" ? window : this, function( window, noGlobal ) {

// Can't do this because several apps including ASP.NET trace
// the stack via arguments.caller.callee and Firefox dies if
// you try to trace through "use strict" call chains. (#13335)
// Support: Firefox 18+
//

var deletedIds = [];

var slice = deletedIds.slice;

var concat = deletedIds.concat;

var push = deletedIds.push;

var indexOf = deletedIds.indexOf;

var class2type = {};

var toString = class2type.toString;

var hasOwn = class2type.hasOwnProperty;

var support = {};


var
	version = "1.11.1",

	// Define a local copy of jQuery
	jQuery = function( selector, context ) {
		// The jQuery object is actually just the init constructor 'enhanced'
		// Need init if jQuery is called (just allow error to be thrown if not included)
		return new jQuery.fn.init( selector, context );
	},

	// Support: Android<4.1, IE<9
	// Make sure we trim BOM and NBSP
	rtrim = /^[\s\uFEFF\xA0]+|[\s\uFEFF\xA0]+$/g,

	// Matches dashed string for camelizing
	rmsPrefix = /^-ms-/,
	rdashAlpha = /-([\da-z])/gi,

	// Used by jQuery.camelCase as callback to replace()
	fcamelCase = function( all, letter ) {
		return letter.toUpperCase();
	};

jQuery.fn = jQuery.prototype = {
	// The current version of jQuery being used
	jquery: version,

	constructor: jQuery,

	// Start with an empty selector
	selector: "",

	// The default length of a jQuery object is 0
	length: 0,

	toArray: function() {
		return slice.call( this );
	},

	// Get the Nth element in the matched element set OR
	// Get the whole matched element set as a clean array
	get: function( num ) {
		return num != null ?

			// Return just the one element from the set
			( num < 0 ? this[ num + this.length ] : this[ num ] ) :

			// Return all the elements in a clean array
			slice.call( this );
	},

	// Take an array of elements and push it onto the stack
	// (returning the new matched element set)
	pushStack: function( elems ) {

		// Build a new jQuery matched element set
		var ret = jQuery.merge( this.constructor(), elems );

		// Add the old object onto the stack (as a reference)
		ret.prevObject = this;
		ret.context = this.context;

		// Return the newly-formed element set
		return ret;
	},

	// Execute a callback for every element in the matched set.
	// (You can seed the arguments with an array of args, but this is
	// only used internally.)
	each: function( callback, args ) {
		return jQuery.each( this, callback, args );
	},

	map: function( callback ) {
		return this.pushStack( jQuery.map(this, function( elem, i ) {
			return callback.call( elem, i, elem );
		}));
	},

	slice: function() {
		return this.pushStack( slice.apply( this, arguments ) );
	},

	first: function() {
		return this.eq( 0 );
	},

	last: function() {
		return this.eq( -1 );
	},

	eq: function( i ) {
		var len = this.length,
			j = +i + ( i < 0 ? len : 0 );
		return this.pushStack( j >= 0 && j < len ? [ this[j] ] : [] );
	},

	end: function() {
		return this.prevObject || this.constructor(null);
	},

	// For internal use only.
	// Behaves like an Array's method, not like a jQuery method.
	push: push,
	sort: deletedIds.sort,
	splice: deletedIds.splice
};

jQuery.extend = jQuery.fn.extend = function() {
	var src, copyIsArray, copy, name, options, clone,
		target = arguments[0] || {},
		i = 1,
		length = arguments.length,
		deep = false;

	// Handle a deep copy situation
	if ( typeof target === "boolean" ) {
		deep = target;

		// skip the boolean and the target
		target = arguments[ i ] || {};
		i++;
	}

	// Handle case when target is a string or something (possible in deep copy)
	if ( typeof target !== "object" && !jQuery.isFunction(target) ) {
		target = {};
	}

	// extend jQuery itself if only one argument is passed
	if ( i === length ) {
		target = this;
		i--;
	}

	for ( ; i < length; i++ ) {
		// Only deal with non-null/undefined values
		if ( (options = arguments[ i ]) != null ) {
			// Extend the base object
			for ( name in options ) {
				src = target[ name ];
				copy = options[ name ];

				// Prevent never-ending loop
				if ( target === copy ) {
					continue;
				}

				// Recurse if we're merging plain objects or arrays
				if ( deep && copy && ( jQuery.isPlainObject(copy) || (copyIsArray = jQuery.isArray(copy)) ) ) {
					if ( copyIsArray ) {
						copyIsArray = false;
						clone = src && jQuery.isArray(src) ? src : [];

					} else {
						clone = src && jQuery.isPlainObject(src) ? src : {};
					}

					// Never move original objects, clone them
					target[ name ] = jQuery.extend( deep, clone, copy );

				// Don't bring in undefined values
				} else if ( copy !== undefined ) {
					target[ name ] = copy;
				}
			}
		}
	}

	// Return the modified object
	return target;
};

jQuery.extend({
	// Unique for each copy of jQuery on the page
	expando: "jQuery" + ( version + Math.random() ).replace( /\D/g, "" ),

	// Assume jQuery is ready without the ready module
	isReady: true,

	error: function( msg ) {
		throw new Error( msg );
	},

	noop: function() {},

	// See test/unit/core.js for details concerning isFunction.
	// Since version 1.3, DOM methods and functions like alert
	// aren't supported. They return false on IE (#2968).
	isFunction: function( obj ) {
		return jQuery.type(obj) === "function";
	},

	isArray: Array.isArray || function( obj ) {
		return jQuery.type(obj) === "array";
	},

	isWindow: function( obj ) {
		/* jshint eqeqeq: false */
		return obj != null && obj == obj.window;
	},

	isNumeric: function( obj ) {
		// parseFloat NaNs numeric-cast false positives (null|true|false|"")
		// ...but misinterprets leading-number strings, particularly hex literals ("0x...")
		// subtraction forces infinities to NaN
		return !jQuery.isArray( obj ) && obj - parseFloat( obj ) >= 0;
	},

	isEmptyObject: function( obj ) {
		var name;
		for ( name in obj ) {
			return false;
		}
		return true;
	},

	isPlainObject: function( obj ) {
		var key;

		// Must be an Object.
		// Because of IE, we also have to check the presence of the constructor property.
		// Make sure that DOM nodes and window objects don't pass through, as well
		if ( !obj || jQuery.type(obj) !== "object" || obj.nodeType || jQuery.isWindow( obj ) ) {
			return false;
		}

		try {
			// Not own constructor property must be Object
			if ( obj.constructor &&
				!hasOwn.call(obj, "constructor") &&
				!hasOwn.call(obj.constructor.prototype, "isPrototypeOf") ) {
				return false;
			}
		} catch ( e ) {
			// IE8,9 Will throw exceptions on certain host objects #9897
			return false;
		}

		// Support: IE<9
		// Handle iteration over inherited properties before own properties.
		if ( support.ownLast ) {
			for ( key in obj ) {
				return hasOwn.call( obj, key );
			}
		}

		// Own properties are enumerated firstly, so to speed up,
		// if last one is own, then all properties are own.
		for ( key in obj ) {}

		return key === undefined || hasOwn.call( obj, key );
	},

	type: function( obj ) {
		if ( obj == null ) {
			return obj + "";
		}
		return typeof obj === "object" || typeof obj === "function" ?
			class2type[ toString.call(obj) ] || "object" :
			typeof obj;
	},

	// Evaluates a script in a global context
	// Workarounds based on findings by Jim Driscoll
	// http://weblogs.java.net/blog/driscoll/archive/2009/09/08/eval-javascript-global-context
	globalEval: function( data ) {
		if ( data && jQuery.trim( data ) ) {
			// We use execScript on Internet Explorer
			// We use an anonymous function so that context is window
			// rather than jQuery in Firefox
			( window.execScript || function( data ) {
				window[ "eval" ].call( window, data );
			} )( data );
		}
	},

	// Convert dashed to camelCase; used by the css and data modules
	// Microsoft forgot to hump their vendor prefix (#9572)
	camelCase: function( string ) {
		return string.replace( rmsPrefix, "ms-" ).replace( rdashAlpha, fcamelCase );
	},

	nodeName: function( elem, name ) {
		return elem.nodeName && elem.nodeName.toLowerCase() === name.toLowerCase();
	},

	// args is for internal usage only
	each: function( obj, callback, args ) {
		var value,
			i = 0,
			length = obj.length,
			isArray = isArraylike( obj );

		if ( args ) {
			if ( isArray ) {
				for ( ; i < length; i++ ) {
					value = callback.apply( obj[ i ], args );

					if ( value === false ) {
						break;
					}
				}
			} else {
				for ( i in obj ) {
					value = callback.apply( obj[ i ], args );

					if ( value === false ) {
						break;
					}
				}
			}

		// A special, fast, case for the most common use of each
		} else {
			if ( isArray ) {
				for ( ; i < length; i++ ) {
					value = callback.call( obj[ i ], i, obj[ i ] );

					if ( value === false ) {
						break;
					}
				}
			} else {
				for ( i in obj ) {
					value = callback.call( obj[ i ], i, obj[ i ] );

					if ( value === false ) {
						break;
					}
				}
			}
		}

		return obj;
	},

	// Support: Android<4.1, IE<9
	trim: function( text ) {
		return text == null ?
			"" :
			( text + "" ).replace( rtrim, "" );
	},

	// results is for internal usage only
	makeArray: function( arr, results ) {
		var ret = results || [];

		if ( arr != null ) {
			if ( isArraylike( Object(arr) ) ) {
				jQuery.merge( ret,
					typeof arr === "string" ?
					[ arr ] : arr
				);
			} else {
				push.call( ret, arr );
			}
		}

		return ret;
	},

	inArray: function( elem, arr, i ) {
		var len;

		if ( arr ) {
			if ( indexOf ) {
				return indexOf.call( arr, elem, i );
			}

			len = arr.length;
			i = i ? i < 0 ? Math.max( 0, len + i ) : i : 0;

			for ( ; i < len; i++ ) {
				// Skip accessing in sparse arrays
				if ( i in arr && arr[ i ] === elem ) {
					return i;
				}
			}
		}

		return -1;
	},

	merge: function( first, second ) {
		var len = +second.length,
			j = 0,
			i = first.length;

		while ( j < len ) {
			first[ i++ ] = second[ j++ ];
		}

		// Support: IE<9
		// Workaround casting of .length to NaN on otherwise arraylike objects (e.g., NodeLists)
		if ( len !== len ) {
			while ( second[j] !== undefined ) {
				first[ i++ ] = second[ j++ ];
			}
		}

		first.length = i;

		return first;
	},

	grep: function( elems, callback, invert ) {
		var callbackInverse,
			matches = [],
			i = 0,
			length = elems.length,
			callbackExpect = !invert;

		// Go through the array, only saving the items
		// that pass the validator function
		for ( ; i < length; i++ ) {
			callbackInverse = !callback( elems[ i ], i );
			if ( callbackInverse !== callbackExpect ) {
				matches.push( elems[ i ] );
			}
		}

		return matches;
	},

	// arg is for internal usage only
	map: function( elems, callback, arg ) {
		var value,
			i = 0,
			length = elems.length,
			isArray = isArraylike( elems ),
			ret = [];

		// Go through the array, translating each of the items to their new values
		if ( isArray ) {
			for ( ; i < length; i++ ) {
				value = callback( elems[ i ], i, arg );

				if ( value != null ) {
					ret.push( value );
				}
			}

		// Go through every key on the object,
		} else {
			for ( i in elems ) {
				value = callback( elems[ i ], i, arg );

				if ( value != null ) {
					ret.push( value );
				}
			}
		}

		// Flatten any nested arrays
		return concat.apply( [], ret );
	},

	// A global GUID counter for objects
	guid: 1,

	// Bind a function to a context, optionally partially applying any
	// arguments.
	proxy: function( fn, context ) {
		var args, proxy, tmp;

		if ( typeof context === "string" ) {
			tmp = fn[ context ];
			context = fn;
			fn = tmp;
		}

		// Quick check to determine if target is callable, in the spec
		// this throws a TypeError, but we will just return undefined.
		if ( !jQuery.isFunction( fn ) ) {
			return undefined;
		}

		// Simulated bind
		args = slice.call( arguments, 2 );
		proxy = function() {
			return fn.apply( context || this, args.concat( slice.call( arguments ) ) );
		};

		// Set the guid of unique handler to the same of original handler, so it can be removed
		proxy.guid = fn.guid = fn.guid || jQuery.guid++;

		return proxy;
	},

	now: function() {
		return +( new Date() );
	},

	// jQuery.support is not used in Core but other projects attach their
	// properties to it so it needs to exist.
	support: support
});

// Populate the class2type map
jQuery.each("Boolean Number String Function Array Date RegExp Object Error".split(" "), function(i, name) {
	class2type[ "[object " + name + "]" ] = name.toLowerCase();
});

function isArraylike( obj ) {
	var length = obj.length,
		type = jQuery.type( obj );

	if ( type === "function" || jQuery.isWindow( obj ) ) {
		return false;
	}

	if ( obj.nodeType === 1 && length ) {
		return true;
	}

	return type === "array" || length === 0 ||
		typeof length === "number" && length > 0 && ( length - 1 ) in obj;
}
var Sizzle =
/*!
 * Sizzle CSS Selector Engine v1.10.19
 * http://sizzlejs.com/
 *
 * Copyright 2013 jQuery Foundation, Inc. and other contributors
 * Released under the MIT license
 * http://jquery.org/license
 *
 * Date: 2014-04-18
 */
(function( window ) {

var i,
	support,
	Expr,
	getText,
	isXML,
	tokenize,
	compile,
	select,
	outermostContext,
	sortInput,
	hasDuplicate,

	// Local document vars
	setDocument,
	document,
	docElem,
	documentIsHTML,
	rbuggyQSA,
	rbuggyMatches,
	matches,
	contains,

	// Instance-specific data
	expando = "sizzle" + -(new Date()),
	preferredDoc = window.document,
	dirruns = 0,
	done = 0,
	classCache = createCache(),
	tokenCache = createCache(),
	compilerCache = createCache(),
	sortOrder = function( a, b ) {
		if ( a === b ) {
			hasDuplicate = true;
		}
		return 0;
	},

	// General-purpose constants
	strundefined = typeof undefined,
	MAX_NEGATIVE = 1 << 31,

	// Instance methods
	hasOwn = ({}).hasOwnProperty,
	arr = [],
	pop = arr.pop,
	push_native = arr.push,
	push = arr.push,
	slice = arr.slice,
	// Use a stripped-down indexOf if we can't use a native one
	indexOf = arr.indexOf || function( elem ) {
		var i = 0,
			len = this.length;
		for ( ; i < len; i++ ) {
			if ( this[i] === elem ) {
				return i;
			}
		}
		return -1;
	},

	booleans = "checked|selected|async|autofocus|autoplay|controls|defer|disabled|hidden|ismap|loop|multiple|open|readonly|required|scoped",

	// Regular expressions

	// Whitespace characters http://www.w3.org/TR/css3-selectors/#whitespace
	whitespace = "[\\x20\\t\\r\\n\\f]",
	// http://www.w3.org/TR/css3-syntax/#characters
	characterEncoding = "(?:\\\\.|[\\w-]|[^\\x00-\\xa0])+",

	// Loosely modeled on CSS identifier characters
	// An unquoted value should be a CSS identifier http://www.w3.org/TR/css3-selectors/#attribute-selectors
	// Proper syntax: http://www.w3.org/TR/CSS21/syndata.html#value-def-identifier
	identifier = characterEncoding.replace( "w", "w#" ),

	// Attribute selectors: http://www.w3.org/TR/selectors/#attribute-selectors
	attributes = "\\[" + whitespace + "*(" + characterEncoding + ")(?:" + whitespace +
		// Operator (capture 2)
		"*([*^$|!~]?=)" + whitespace +
		// "Attribute values must be CSS identifiers [capture 5] or strings [capture 3 or capture 4]"
		"*(?:'((?:\\\\.|[^\\\\'])*)'|\"((?:\\\\.|[^\\\\\"])*)\"|(" + identifier + "))|)" + whitespace +
		"*\\]",

	pseudos = ":(" + characterEncoding + ")(?:\\((" +
		// To reduce the number of selectors needing tokenize in the preFilter, prefer arguments:
		// 1. quoted (capture 3; capture 4 or capture 5)
		"('((?:\\\\.|[^\\\\'])*)'|\"((?:\\\\.|[^\\\\\"])*)\")|" +
		// 2. simple (capture 6)
		"((?:\\\\.|[^\\\\()[\\]]|" + attributes + ")*)|" +
		// 3. anything else (capture 2)
		".*" +
		")\\)|)",

	// Leading and non-escaped trailing whitespace, capturing some non-whitespace characters preceding the latter
	rtrim = new RegExp( "^" + whitespace + "+|((?:^|[^\\\\])(?:\\\\.)*)" + whitespace + "+$", "g" ),

	rcomma = new RegExp( "^" + whitespace + "*," + whitespace + "*" ),
	rcombinators = new RegExp( "^" + whitespace + "*([>+~]|" + whitespace + ")" + whitespace + "*" ),

	rattributeQuotes = new RegExp( "=" + whitespace + "*([^\\]'\"]*?)" + whitespace + "*\\]", "g" ),

	rpseudo = new RegExp( pseudos ),
	ridentifier = new RegExp( "^" + identifier + "$" ),

	matchExpr = {
		"ID": new RegExp( "^#(" + characterEncoding + ")" ),
		"CLASS": new RegExp( "^\\.(" + characterEncoding + ")" ),
		"TAG": new RegExp( "^(" + characterEncoding.replace( "w", "w*" ) + ")" ),
		"ATTR": new RegExp( "^" + attributes ),
		"PSEUDO": new RegExp( "^" + pseudos ),
		"CHILD": new RegExp( "^:(only|first|last|nth|nth-last)-(child|of-type)(?:\\(" + whitespace +
			"*(even|odd|(([+-]|)(\\d*)n|)" + whitespace + "*(?:([+-]|)" + whitespace +
			"*(\\d+)|))" + whitespace + "*\\)|)", "i" ),
		"bool": new RegExp( "^(?:" + booleans + ")$", "i" ),
		// For use in libraries implementing .is()
		// We use this for POS matching in `select`
		"needsContext": new RegExp( "^" + whitespace + "*[>+~]|:(even|odd|eq|gt|lt|nth|first|last)(?:\\(" +
			whitespace + "*((?:-\\d)?\\d*)" + whitespace + "*\\)|)(?=[^-]|$)", "i" )
	},

	rinputs = /^(?:input|select|textarea|button)$/i,
	rheader = /^h\d$/i,

	rnative = /^[^{]+\{\s*\[native \w/,

	// Easily-parseable/retrievable ID or TAG or CLASS selectors
	rquickExpr = /^(?:#([\w-]+)|(\w+)|\.([\w-]+))$/,

	rsibling = /[+~]/,
	rescape = /'|\\/g,

	// CSS escapes http://www.w3.org/TR/CSS21/syndata.html#escaped-characters
	runescape = new RegExp( "\\\\([\\da-f]{1,6}" + whitespace + "?|(" + whitespace + ")|.)", "ig" ),
	funescape = function( _, escaped, escapedWhitespace ) {
		var high = "0x" + escaped - 0x10000;
		// NaN means non-codepoint
		// Support: Firefox<24
		// Workaround erroneous numeric interpretation of +"0x"
		return high !== high || escapedWhitespace ?
			escaped :
			high < 0 ?
				// BMP codepoint
				String.fromCharCode( high + 0x10000 ) :
				// Supplemental Plane codepoint (surrogate pair)
				String.fromCharCode( high >> 10 | 0xD800, high & 0x3FF | 0xDC00 );
	};

// Optimize for push.apply( _, NodeList )
try {
	push.apply(
		(arr = slice.call( preferredDoc.childNodes )),
		preferredDoc.childNodes
	);
	// Support: Android<4.0
	// Detect silently failing push.apply
	arr[ preferredDoc.childNodes.length ].nodeType;
} catch ( e ) {
	push = { apply: arr.length ?

		// Leverage slice if possible
		function( target, els ) {
			push_native.apply( target, slice.call(els) );
		} :

		// Support: IE<9
		// Otherwise append directly
		function( target, els ) {
			var j = target.length,
				i = 0;
			// Can't trust NodeList.length
			while ( (target[j++] = els[i++]) ) {}
			target.length = j - 1;
		}
	};
}

function Sizzle( selector, context, results, seed ) {
	var match, elem, m, nodeType,
		// QSA vars
		i, groups, old, nid, newContext, newSelector;

	if ( ( context ? context.ownerDocument || context : preferredDoc ) !== document ) {
		setDocument( context );
	}

	context = context || document;
	results = results || [];

	if ( !selector || typeof selector !== "string" ) {
		return results;
	}

	if ( (nodeType = context.nodeType) !== 1 && nodeType !== 9 ) {
		return [];
	}

	if ( documentIsHTML && !seed ) {

		// Shortcuts
		if ( (match = rquickExpr.exec( selector )) ) {
			// Speed-up: Sizzle("#ID")
			if ( (m = match[1]) ) {
				if ( nodeType === 9 ) {
					elem = context.getElementById( m );
					// Check parentNode to catch when Blackberry 4.6 returns
					// nodes that are no longer in the document (jQuery #6963)
					if ( elem && elem.parentNode ) {
						// Handle the case where IE, Opera, and Webkit return items
						// by name instead of ID
						if ( elem.id === m ) {
							results.push( elem );
							return results;
						}
					} else {
						return results;
					}
				} else {
					// Context is not a document
					if ( context.ownerDocument && (elem = context.ownerDocument.getElementById( m )) &&
						contains( context, elem ) && elem.id === m ) {
						results.push( elem );
						return results;
					}
				}

			// Speed-up: Sizzle("TAG")
			} else if ( match[2] ) {
				push.apply( results, context.getElementsByTagName( selector ) );
				return results;

			// Speed-up: Sizzle(".CLASS")
			} else if ( (m = match[3]) && support.getElementsByClassName && context.getElementsByClassName ) {
				push.apply( results, context.getElementsByClassName( m ) );
				return results;
			}
		}

		// QSA path
		if ( support.qsa && (!rbuggyQSA || !rbuggyQSA.test( selector )) ) {
			nid = old = expando;
			newContext = context;
			newSelector = nodeType === 9 && selector;

			// qSA works strangely on Element-rooted queries
			// We can work around this by specifying an extra ID on the root
			// and working up from there (Thanks to Andrew Dupont for the technique)
			// IE 8 doesn't work on object elements
			if ( nodeType === 1 && context.nodeName.toLowerCase() !== "object" ) {
				groups = tokenize( selector );

				if ( (old = context.getAttribute("id")) ) {
					nid = old.replace( rescape, "\\$&" );
				} else {
					context.setAttribute( "id", nid );
				}
				nid = "[id='" + nid + "'] ";

				i = groups.length;
				while ( i-- ) {
					groups[i] = nid + toSelector( groups[i] );
				}
				newContext = rsibling.test( selector ) && testContext( context.parentNode ) || context;
				newSelector = groups.join(",");
			}

			if ( newSelector ) {
				try {
					push.apply( results,
						newContext.querySelectorAll( newSelector )
					);
					return results;
				} catch(qsaError) {
				} finally {
					if ( !old ) {
						context.removeAttribute("id");
					}
				}
			}
		}
	}

	// All others
	return select( selector.replace( rtrim, "$1" ), context, results, seed );
}

/**
 * Create key-value caches of limited size
 * @returns {Function(string, Object)} Returns the Object data after storing it on itself with
 *	property name the (space-suffixed) string and (if the cache is larger than Expr.cacheLength)
 *	deleting the oldest entry
 */
function createCache() {
	var keys = [];

	function cache( key, value ) {
		// Use (key + " ") to avoid collision with native prototype properties (see Issue #157)
		if ( keys.push( key + " " ) > Expr.cacheLength ) {
			// Only keep the most recent entries
			delete cache[ keys.shift() ];
		}
		return (cache[ key + " " ] = value);
	}
	return cache;
}

/**
 * Mark a function for special use by Sizzle
 * @param {Function} fn The function to mark
 */
function markFunction( fn ) {
	fn[ expando ] = true;
	return fn;
}

/**
 * Support testing using an element
 * @param {Function} fn Passed the created div and expects a boolean result
 */
function assert( fn ) {
	var div = document.createElement("div");

	try {
		return !!fn( div );
	} catch (e) {
		return false;
	} finally {
		// Remove from its parent by default
		if ( div.parentNode ) {
			div.parentNode.removeChild( div );
		}
		// release memory in IE
		div = null;
	}
}

/**
 * Adds the same handler for all of the specified attrs
 * @param {String} attrs Pipe-separated list of attributes
 * @param {Function} handler The method that will be applied
 */
function addHandle( attrs, handler ) {
	var arr = attrs.split("|"),
		i = attrs.length;

	while ( i-- ) {
		Expr.attrHandle[ arr[i] ] = handler;
	}
}

/**
 * Checks document order of two siblings
 * @param {Element} a
 * @param {Element} b
 * @returns {Number} Returns less than 0 if a precedes b, greater than 0 if a follows b
 */
function siblingCheck( a, b ) {
	var cur = b && a,
		diff = cur && a.nodeType === 1 && b.nodeType === 1 &&
			( ~b.sourceIndex || MAX_NEGATIVE ) -
			( ~a.sourceIndex || MAX_NEGATIVE );

	// Use IE sourceIndex if available on both nodes
	if ( diff ) {
		return diff;
	}

	// Check if b follows a
	if ( cur ) {
		while ( (cur = cur.nextSibling) ) {
			if ( cur === b ) {
				return -1;
			}
		}
	}

	return a ? 1 : -1;
}

/**
 * Returns a function to use in pseudos for input types
 * @param {String} type
 */
function createInputPseudo( type ) {
	return function( elem ) {
		var name = elem.nodeName.toLowerCase();
		return name === "input" && elem.type === type;
	};
}

/**
 * Returns a function to use in pseudos for buttons
 * @param {String} type
 */
function createButtonPseudo( type ) {
	return function( elem ) {
		var name = elem.nodeName.toLowerCase();
		return (name === "input" || name === "button") && elem.type === type;
	};
}

/**
 * Returns a function to use in pseudos for positionals
 * @param {Function} fn
 */
function createPositionalPseudo( fn ) {
	return markFunction(function( argument ) {
		argument = +argument;
		return markFunction(function( seed, matches ) {
			var j,
				matchIndexes = fn( [], seed.length, argument ),
				i = matchIndexes.length;

			// Match elements found at the specified indexes
			while ( i-- ) {
				if ( seed[ (j = matchIndexes[i]) ] ) {
					seed[j] = !(matches[j] = seed[j]);
				}
			}
		});
	});
}

/**
 * Checks a node for validity as a Sizzle context
 * @param {Element|Object=} context
 * @returns {Element|Object|Boolean} The input node if acceptable, otherwise a falsy value
 */
function testContext( context ) {
	return context && typeof context.getElementsByTagName !== strundefined && context;
}

// Expose support vars for convenience
support = Sizzle.support = {};

/**
 * Detects XML nodes
 * @param {Element|Object} elem An element or a document
 * @returns {Boolean} True iff elem is a non-HTML XML node
 */
isXML = Sizzle.isXML = function( elem ) {
	// documentElement is verified for cases where it doesn't yet exist
	// (such as loading iframes in IE - #4833)
	var documentElement = elem && (elem.ownerDocument || elem).documentElement;
	return documentElement ? documentElement.nodeName !== "HTML" : false;
};

/**
 * Sets document-related variables once based on the current document
 * @param {Element|Object} [doc] An element or document object to use to set the document
 * @returns {Object} Returns the current document
 */
setDocument = Sizzle.setDocument = function( node ) {
	var hasCompare,
		doc = node ? node.ownerDocument || node : preferredDoc,
		parent = doc.defaultView;

	// If no document and documentElement is available, return
	if ( doc === document || doc.nodeType !== 9 || !doc.documentElement ) {
		return document;
	}

	// Set our document
	document = doc;
	docElem = doc.documentElement;

	// Support tests
	documentIsHTML = !isXML( doc );

	// Support: IE>8
	// If iframe document is assigned to "document" variable and if iframe has been reloaded,
	// IE will throw "permission denied" error when accessing "document" variable, see jQuery #13936
	// IE6-8 do not support the defaultView property so parent will be undefined
	if ( parent && parent !== parent.top ) {
		// IE11 does not have attachEvent, so all must suffer
		if ( parent.addEventListener ) {
			parent.addEventListener( "unload", function() {
				setDocument();
			}, false );
		} else if ( parent.attachEvent ) {
			parent.attachEvent( "onunload", function() {
				setDocument();
			});
		}
	}

	/* Attributes
	---------------------------------------------------------------------- */

	// Support: IE<8
	// Verify that getAttribute really returns attributes and not properties (excepting IE8 booleans)
	support.attributes = assert(function( div ) {
		div.className = "i";
		return !div.getAttribute("className");
	});

	/* getElement(s)By*
	---------------------------------------------------------------------- */

	// Check if getElementsByTagName("*") returns only elements
	support.getElementsByTagName = assert(function( div ) {
		div.appendChild( doc.createComment("") );
		return !div.getElementsByTagName("*").length;
	});

	// Check if getElementsByClassName can be trusted
	support.getElementsByClassName = rnative.test( doc.getElementsByClassName ) && assert(function( div ) {
		div.innerHTML = "<div class='a'></div><div class='a i'></div>";

		// Support: Safari<4
		// Catch class over-caching
		div.firstChild.className = "i";
		// Support: Opera<10
		// Catch gEBCN failure to find non-leading classes
		return div.getElementsByClassName("i").length === 2;
	});

	// Support: IE<10
	// Check if getElementById returns elements by name
	// The broken getElementById methods don't pick up programatically-set names,
	// so use a roundabout getElementsByName test
	support.getById = assert(function( div ) {
		docElem.appendChild( div ).id = expando;
		return !doc.getElementsByName || !doc.getElementsByName( expando ).length;
	});

	// ID find and filter
	if ( support.getById ) {
		Expr.find["ID"] = function( id, context ) {
			if ( typeof context.getElementById !== strundefined && documentIsHTML ) {
				var m = context.getElementById( id );
				// Check parentNode to catch when Blackberry 4.6 returns
				// nodes that are no longer in the document #6963
				return m && m.parentNode ? [ m ] : [];
			}
		};
		Expr.filter["ID"] = function( id ) {
			var attrId = id.replace( runescape, funescape );
			return function( elem ) {
				return elem.getAttribute("id") === attrId;
			};
		};
	} else {
		// Support: IE6/7
		// getElementById is not reliable as a find shortcut
		delete Expr.find["ID"];

		Expr.filter["ID"] =  function( id ) {
			var attrId = id.replace( runescape, funescape );
			return function( elem ) {
				var node = typeof elem.getAttributeNode !== strundefined && elem.getAttributeNode("id");
				return node && node.value === attrId;
			};
		};
	}

	// Tag
	Expr.find["TAG"] = support.getElementsByTagName ?
		function( tag, context ) {
			if ( typeof context.getElementsByTagName !== strundefined ) {
				return context.getElementsByTagName( tag );
			}
		} :
		function( tag, context ) {
			var elem,
				tmp = [],
				i = 0,
				results = context.getElementsByTagName( tag );

			// Filter out possible comments
			if ( tag === "*" ) {
				while ( (elem = results[i++]) ) {
					if ( elem.nodeType === 1 ) {
						tmp.push( elem );
					}
				}

				return tmp;
			}
			return results;
		};

	// Class
	Expr.find["CLASS"] = support.getElementsByClassName && function( className, context ) {
		if ( typeof context.getElementsByClassName !== strundefined && documentIsHTML ) {
			return context.getElementsByClassName( className );
		}
	};

	/* QSA/matchesSelector
	---------------------------------------------------------------------- */

	// QSA and matchesSelector support

	// matchesSelector(:active) reports false when true (IE9/Opera 11.5)
	rbuggyMatches = [];

	// qSa(:focus) reports false when true (Chrome 21)
	// We allow this because of a bug in IE8/9 that throws an error
	// whenever `document.activeElement` is accessed on an iframe
	// So, we allow :focus to pass through QSA all the time to avoid the IE error
	// See http://bugs.jquery.com/ticket/13378
	rbuggyQSA = [];

	if ( (support.qsa = rnative.test( doc.querySelectorAll )) ) {
		// Build QSA regex
		// Regex strategy adopted from Diego Perini
		assert(function( div ) {
			// Select is set to empty string on purpose
			// This is to test IE's treatment of not explicitly
			// setting a boolean content attribute,
			// since its presence should be enough
			// http://bugs.jquery.com/ticket/12359
			div.innerHTML = "<select msallowclip=''><option selected=''></option></select>";

			// Support: IE8, Opera 11-12.16
			// Nothing should be selected when empty strings follow ^= or $= or *=
			// The test attribute must be unknown in Opera but "safe" for WinRT
			// http://msdn.microsoft.com/en-us/library/ie/hh465388.aspx#attribute_section
			if ( div.querySelectorAll("[msallowclip^='']").length ) {
				rbuggyQSA.push( "[*^$]=" + whitespace + "*(?:''|\"\")" );
			}

			// Support: IE8
			// Boolean attributes and "value" are not treated correctly
			if ( !div.querySelectorAll("[selected]").length ) {
				rbuggyQSA.push( "\\[" + whitespace + "*(?:value|" + booleans + ")" );
			}

			// Webkit/Opera - :checked should return selected option elements
			// http://www.w3.org/TR/2011/REC-css3-selectors-20110929/#checked
			// IE8 throws error here and will not see later tests
			if ( !div.querySelectorAll(":checked").length ) {
				rbuggyQSA.push(":checked");
			}
		});

		assert(function( div ) {
			// Support: Windows 8 Native Apps
			// The type and name attributes are restricted during .innerHTML assignment
			var input = doc.createElement("input");
			input.setAttribute( "type", "hidden" );
			div.appendChild( input ).setAttribute( "name", "D" );

			// Support: IE8
			// Enforce case-sensitivity of name attribute
			if ( div.querySelectorAll("[name=d]").length ) {
				rbuggyQSA.push( "name" + whitespace + "*[*^$|!~]?=" );
			}

			// FF 3.5 - :enabled/:disabled and hidden elements (hidden elements are still enabled)
			// IE8 throws error here and will not see later tests
			if ( !div.querySelectorAll(":enabled").length ) {
				rbuggyQSA.push( ":enabled", ":disabled" );
			}

			// Opera 10-11 does not throw on post-comma invalid pseudos
			div.querySelectorAll("*,:x");
			rbuggyQSA.push(",.*:");
		});
	}

	if ( (support.matchesSelector = rnative.test( (matches = docElem.matches ||
		docElem.webkitMatchesSelector ||
		docElem.mozMatchesSelector ||
		docElem.oMatchesSelector ||
		docElem.msMatchesSelector) )) ) {

		assert(function( div ) {
			// Check to see if it's possible to do matchesSelector
			// on a disconnected node (IE 9)
			support.disconnectedMatch = matches.call( div, "div" );

			// This should fail with an exception
			// Gecko does not error, returns false instead
			matches.call( div, "[s!='']:x" );
			rbuggyMatches.push( "!=", pseudos );
		});
	}

	rbuggyQSA = rbuggyQSA.length && new RegExp( rbuggyQSA.join("|") );
	rbuggyMatches = rbuggyMatches.length && new RegExp( rbuggyMatches.join("|") );

	/* Contains
	---------------------------------------------------------------------- */
	hasCompare = rnative.test( docElem.compareDocumentPosition );

	// Element contains another
	// Purposefully does not implement inclusive descendent
	// As in, an element does not contain itself
	contains = hasCompare || rnative.test( docElem.contains ) ?
		function( a, b ) {
			var adown = a.nodeType === 9 ? a.documentElement : a,
				bup = b && b.parentNode;
			return a === bup || !!( bup && bup.nodeType === 1 && (
				adown.contains ?
					adown.contains( bup ) :
					a.compareDocumentPosition && a.compareDocumentPosition( bup ) & 16
			));
		} :
		function( a, b ) {
			if ( b ) {
				while ( (b = b.parentNode) ) {
					if ( b === a ) {
						return true;
					}
				}
			}
			return false;
		};

	/* Sorting
	---------------------------------------------------------------------- */

	// Document order sorting
	sortOrder = hasCompare ?
	function( a, b ) {

		// Flag for duplicate removal
		if ( a === b ) {
			hasDuplicate = true;
			return 0;
		}

		// Sort on method existence if only one input has compareDocumentPosition
		var compare = !a.compareDocumentPosition - !b.compareDocumentPosition;
		if ( compare ) {
			return compare;
		}

		// Calculate position if both inputs belong to the same document
		compare = ( a.ownerDocument || a ) === ( b.ownerDocument || b ) ?
			a.compareDocumentPosition( b ) :

			// Otherwise we know they are disconnected
			1;

		// Disconnected nodes
		if ( compare & 1 ||
			(!support.sortDetached && b.compareDocumentPosition( a ) === compare) ) {

			// Choose the first element that is related to our preferred document
			if ( a === doc || a.ownerDocument === preferredDoc && contains(preferredDoc, a) ) {
				return -1;
			}
			if ( b === doc || b.ownerDocument === preferredDoc && contains(preferredDoc, b) ) {
				return 1;
			}

			// Maintain original order
			return sortInput ?
				( indexOf.call( sortInput, a ) - indexOf.call( sortInput, b ) ) :
				0;
		}

		return compare & 4 ? -1 : 1;
	} :
	function( a, b ) {
		// Exit early if the nodes are identical
		if ( a === b ) {
			hasDuplicate = true;
			return 0;
		}

		var cur,
			i = 0,
			aup = a.parentNode,
			bup = b.parentNode,
			ap = [ a ],
			bp = [ b ];

		// Parentless nodes are either documents or disconnected
		if ( !aup || !bup ) {
			return a === doc ? -1 :
				b === doc ? 1 :
				aup ? -1 :
				bup ? 1 :
				sortInput ?
				( indexOf.call( sortInput, a ) - indexOf.call( sortInput, b ) ) :
				0;

		// If the nodes are siblings, we can do a quick check
		} else if ( aup === bup ) {
			return siblingCheck( a, b );
		}

		// Otherwise we need full lists of their ancestors for comparison
		cur = a;
		while ( (cur = cur.parentNode) ) {
			ap.unshift( cur );
		}
		cur = b;
		while ( (cur = cur.parentNode) ) {
			bp.unshift( cur );
		}

		// Walk down the tree looking for a discrepancy
		while ( ap[i] === bp[i] ) {
			i++;
		}

		return i ?
			// Do a sibling check if the nodes have a common ancestor
			siblingCheck( ap[i], bp[i] ) :

			// Otherwise nodes in our document sort first
			ap[i] === preferredDoc ? -1 :
			bp[i] === preferredDoc ? 1 :
			0;
	};

	return doc;
};

Sizzle.matches = function( expr, elements ) {
	return Sizzle( expr, null, null, elements );
};

Sizzle.matchesSelector = function( elem, expr ) {
	// Set document vars if needed
	if ( ( elem.ownerDocument || elem ) !== document ) {
		setDocument( elem );
	}

	// Make sure that attribute selectors are quoted
	expr = expr.replace( rattributeQuotes, "='$1']" );

	if ( support.matchesSelector && documentIsHTML &&
		( !rbuggyMatches || !rbuggyMatches.test( expr ) ) &&
		( !rbuggyQSA     || !rbuggyQSA.test( expr ) ) ) {

		try {
			var ret = matches.call( elem, expr );

			// IE 9's matchesSelector returns false on disconnected nodes
			if ( ret || support.disconnectedMatch ||
					// As well, disconnected nodes are said to be in a document
					// fragment in IE 9
					elem.document && elem.document.nodeType !== 11 ) {
				return ret;
			}
		} catch(e) {}
	}

	return Sizzle( expr, document, null, [ elem ] ).length > 0;
};

Sizzle.contains = function( context, elem ) {
	// Set document vars if needed
	if ( ( context.ownerDocument || context ) !== document ) {
		setDocument( context );
	}
	return contains( context, elem );
};

Sizzle.attr = function( elem, name ) {
	// Set document vars if needed
	if ( ( elem.ownerDocument || elem ) !== document ) {
		setDocument( elem );
	}

	var fn = Expr.attrHandle[ name.toLowerCase() ],
		// Don't get fooled by Object.prototype properties (jQuery #13807)
		val = fn && hasOwn.call( Expr.attrHandle, name.toLowerCase() ) ?
			fn( elem, name, !documentIsHTML ) :
			undefined;

	return val !== undefined ?
		val :
		support.attributes || !documentIsHTML ?
			elem.getAttribute( name ) :
			(val = elem.getAttributeNode(name)) && val.specified ?
				val.value :
				null;
};

Sizzle.error = function( msg ) {
	throw new Error( "Syntax error, unrecognized expression: " + msg );
};

/**
 * Document sorting and removing duplicates
 * @param {ArrayLike} results
 */
Sizzle.uniqueSort = function( results ) {
	var elem,
		duplicates = [],
		j = 0,
		i = 0;

	// Unless we *know* we can detect duplicates, assume their presence
	hasDuplicate = !support.detectDuplicates;
	sortInput = !support.sortStable && results.slice( 0 );
	results.sort( sortOrder );

	if ( hasDuplicate ) {
		while ( (elem = results[i++]) ) {
			if ( elem === results[ i ] ) {
				j = duplicates.push( i );
			}
		}
		while ( j-- ) {
			results.splice( duplicates[ j ], 1 );
		}
	}

	// Clear input after sorting to release objects
	// See https://github.com/jquery/sizzle/pull/225
	sortInput = null;

	return results;
};

/**
 * Utility function for retrieving the text value of an array of DOM nodes
 * @param {Array|Element} elem
 */
getText = Sizzle.getText = function( elem ) {
	var node,
		ret = "",
		i = 0,
		nodeType = elem.nodeType;

	if ( !nodeType ) {
		// If no nodeType, this is expected to be an array
		while ( (node = elem[i++]) ) {
			// Do not traverse comment nodes
			ret += getText( node );
		}
	} else if ( nodeType === 1 || nodeType === 9 || nodeType === 11 ) {
		// Use textContent for elements
		// innerText usage removed for consistency of new lines (jQuery #11153)
		if ( typeof elem.textContent === "string" ) {
			return elem.textContent;
		} else {
			// Traverse its children
			for ( elem = elem.firstChild; elem; elem = elem.nextSibling ) {
				ret += getText( elem );
			}
		}
	} else if ( nodeType === 3 || nodeType === 4 ) {
		return elem.nodeValue;
	}
	// Do not include comment or processing instruction nodes

	return ret;
};

Expr = Sizzle.selectors = {

	// Can be adjusted by the user
	cacheLength: 50,

	createPseudo: markFunction,

	match: matchExpr,

	attrHandle: {},

	find: {},

	relative: {
		">": { dir: "parentNode", first: true },
		" ": { dir: "parentNode" },
		"+": { dir: "previousSibling", first: true },
		"~": { dir: "previousSibling" }
	},

	preFilter: {
		"ATTR": function( match ) {
			match[1] = match[1].replace( runescape, funescape );

			// Move the given value to match[3] whether quoted or unquoted
			match[3] = ( match[3] || match[4] || match[5] || "" ).replace( runescape, funescape );

			if ( match[2] === "~=" ) {
				match[3] = " " + match[3] + " ";
			}

			return match.slice( 0, 4 );
		},

		"CHILD": function( match ) {
			/* matches from matchExpr["CHILD"]
				1 type (only|nth|...)
				2 what (child|of-type)
				3 argument (even|odd|\d*|\d*n([+-]\d+)?|...)
				4 xn-component of xn+y argument ([+-]?\d*n|)
				5 sign of xn-component
				6 x of xn-component
				7 sign of y-component
				8 y of y-component
			*/
			match[1] = match[1].toLowerCase();

			if ( match[1].slice( 0, 3 ) === "nth" ) {
				// nth-* requires argument
				if ( !match[3] ) {
					Sizzle.error( match[0] );
				}

				// numeric x and y parameters for Expr.filter.CHILD
				// remember that false/true cast respectively to 0/1
				match[4] = +( match[4] ? match[5] + (match[6] || 1) : 2 * ( match[3] === "even" || match[3] === "odd" ) );
				match[5] = +( ( match[7] + match[8] ) || match[3] === "odd" );

			// other types prohibit arguments
			} else if ( match[3] ) {
				Sizzle.error( match[0] );
			}

			return match;
		},

		"PSEUDO": function( match ) {
			var excess,
				unquoted = !match[6] && match[2];

			if ( matchExpr["CHILD"].test( match[0] ) ) {
				return null;
			}

			// Accept quoted arguments as-is
			if ( match[3] ) {
				match[2] = match[4] || match[5] || "";

			// Strip excess characters from unquoted arguments
			} else if ( unquoted && rpseudo.test( unquoted ) &&
				// Get excess from tokenize (recursively)
				(excess = tokenize( unquoted, true )) &&
				// advance to the next closing parenthesis
				(excess = unquoted.indexOf( ")", unquoted.length - excess ) - unquoted.length) ) {

				// excess is a negative index
				match[0] = match[0].slice( 0, excess );
				match[2] = unquoted.slice( 0, excess );
			}

			// Return only captures needed by the pseudo filter method (type and argument)
			return match.slice( 0, 3 );
		}
	},

	filter: {

		"TAG": function( nodeNameSelector ) {
			var nodeName = nodeNameSelector.replace( runescape, funescape ).toLowerCase();
			return nodeNameSelector === "*" ?
				function() { return true; } :
				function( elem ) {
					return elem.nodeName && elem.nodeName.toLowerCase() === nodeName;
				};
		},

		"CLASS": function( className ) {
			var pattern = classCache[ className + " " ];

			return pattern ||
				(pattern = new RegExp( "(^|" + whitespace + ")" + className + "(" + whitespace + "|$)" )) &&
				classCache( className, function( elem ) {
					return pattern.test( typeof elem.className === "string" && elem.className || typeof elem.getAttribute !== strundefined && elem.getAttribute("class") || "" );
				});
		},

		"ATTR": function( name, operator, check ) {
			return function( elem ) {
				var result = Sizzle.attr( elem, name );

				if ( result == null ) {
					return operator === "!=";
				}
				if ( !operator ) {
					return true;
				}

				result += "";

				return operator === "=" ? result === check :
					operator === "!=" ? result !== check :
					operator === "^=" ? check && result.indexOf( check ) === 0 :
					operator === "*=" ? check && result.indexOf( check ) > -1 :
					operator === "$=" ? check && result.slice( -check.length ) === check :
					operator === "~=" ? ( " " + result + " " ).indexOf( check ) > -1 :
					operator === "|=" ? result === check || result.slice( 0, check.length + 1 ) === check + "-" :
					false;
			};
		},

		"CHILD": function( type, what, argument, first, last ) {
			var simple = type.slice( 0, 3 ) !== "nth",
				forward = type.slice( -4 ) !== "last",
				ofType = what === "of-type";

			return first === 1 && last === 0 ?

				// Shortcut for :nth-*(n)
				function( elem ) {
					return !!elem.parentNode;
				} :

				function( elem, context, xml ) {
					var cache, outerCache, node, diff, nodeIndex, start,
						dir = simple !== forward ? "nextSibling" : "previousSibling",
						parent = elem.parentNode,
						name = ofType && elem.nodeName.toLowerCase(),
						useCache = !xml && !ofType;

					if ( parent ) {

						// :(first|last|only)-(child|of-type)
						if ( simple ) {
							while ( dir ) {
								node = elem;
								while ( (node = node[ dir ]) ) {
									if ( ofType ? node.nodeName.toLowerCase() === name : node.nodeType === 1 ) {
										return false;
									}
								}
								// Reverse direction for :only-* (if we haven't yet done so)
								start = dir = type === "only" && !start && "nextSibling";
							}
							return true;
						}

						start = [ forward ? parent.firstChild : parent.lastChild ];

						// non-xml :nth-child(...) stores cache data on `parent`
						if ( forward && useCache ) {
							// Seek `elem` from a previously-cached index
							outerCache = parent[ expando ] || (parent[ expando ] = {});
							cache = outerCache[ type ] || [];
							nodeIndex = cache[0] === dirruns && cache[1];
							diff = cache[0] === dirruns && cache[2];
							node = nodeIndex && parent.childNodes[ nodeIndex ];

							while ( (node = ++nodeIndex && node && node[ dir ] ||

								// Fallback to seeking `elem` from the start
								(diff = nodeIndex = 0) || start.pop()) ) {

								// When found, cache indexes on `parent` and break
								if ( node.nodeType === 1 && ++diff && node === elem ) {
									outerCache[ type ] = [ dirruns, nodeIndex, diff ];
									break;
								}
							}

						// Use previously-cached element index if available
						} else if ( useCache && (cache = (elem[ expando ] || (elem[ expando ] = {}))[ type ]) && cache[0] === dirruns ) {
							diff = cache[1];

						// xml :nth-child(...) or :nth-last-child(...) or :nth(-last)?-of-type(...)
						} else {
							// Use the same loop as above to seek `elem` from the start
							while ( (node = ++nodeIndex && node && node[ dir ] ||
								(diff = nodeIndex = 0) || start.pop()) ) {

								if ( ( ofType ? node.nodeName.toLowerCase() === name : node.nodeType === 1 ) && ++diff ) {
									// Cache the index of each encountered element
									if ( useCache ) {
										(node[ expando ] || (node[ expando ] = {}))[ type ] = [ dirruns, diff ];
									}

									if ( node === elem ) {
										break;
									}
								}
							}
						}

						// Incorporate the offset, then check against cycle size
						diff -= last;
						return diff === first || ( diff % first === 0 && diff / first >= 0 );
					}
				};
		},

		"PSEUDO": function( pseudo, argument ) {
			// pseudo-class names are case-insensitive
			// http://www.w3.org/TR/selectors/#pseudo-classes
			// Prioritize by case sensitivity in case custom pseudos are added with uppercase letters
			// Remember that setFilters inherits from pseudos
			var args,
				fn = Expr.pseudos[ pseudo ] || Expr.setFilters[ pseudo.toLowerCase() ] ||
					Sizzle.error( "unsupported pseudo: " + pseudo );

			// The user may use createPseudo to indicate that
			// arguments are needed to create the filter function
			// just as Sizzle does
			if ( fn[ expando ] ) {
				return fn( argument );
			}

			// But maintain support for old signatures
			if ( fn.length > 1 ) {
				args = [ pseudo, pseudo, "", argument ];
				return Expr.setFilters.hasOwnProperty( pseudo.toLowerCase() ) ?
					markFunction(function( seed, matches ) {
						var idx,
							matched = fn( seed, argument ),
							i = matched.length;
						while ( i-- ) {
							idx = indexOf.call( seed, matched[i] );
							seed[ idx ] = !( matches[ idx ] = matched[i] );
						}
					}) :
					function( elem ) {
						return fn( elem, 0, args );
					};
			}

			return fn;
		}
	},

	pseudos: {
		// Potentially complex pseudos
		"not": markFunction(function( selector ) {
			// Trim the selector passed to compile
			// to avoid treating leading and trailing
			// spaces as combinators
			var input = [],
				results = [],
				matcher = compile( selector.replace( rtrim, "$1" ) );

			return matcher[ expando ] ?
				markFunction(function( seed, matches, context, xml ) {
					var elem,
						unmatched = matcher( seed, null, xml, [] ),
						i = seed.length;

					// Match elements unmatched by `matcher`
					while ( i-- ) {
						if ( (elem = unmatched[i]) ) {
							seed[i] = !(matches[i] = elem);
						}
					}
				}) :
				function( elem, context, xml ) {
					input[0] = elem;
					matcher( input, null, xml, results );
					return !results.pop();
				};
		}),

		"has": markFunction(function( selector ) {
			return function( elem ) {
				return Sizzle( selector, elem ).length > 0;
			};
		}),

		"contains": markFunction(function( text ) {
			return function( elem ) {
				return ( elem.textContent || elem.innerText || getText( elem ) ).indexOf( text ) > -1;
			};
		}),

		// "Whether an element is represented by a :lang() selector
		// is based solely on the element's language value
		// being equal to the identifier C,
		// or beginning with the identifier C immediately followed by "-".
		// The matching of C against the element's language value is performed case-insensitively.
		// The identifier C does not have to be a valid language name."
		// http://www.w3.org/TR/selectors/#lang-pseudo
		"lang": markFunction( function( lang ) {
			// lang value must be a valid identifier
			if ( !ridentifier.test(lang || "") ) {
				Sizzle.error( "unsupported lang: " + lang );
			}
			lang = lang.replace( runescape, funescape ).toLowerCase();
			return function( elem ) {
				var elemLang;
				do {
					if ( (elemLang = documentIsHTML ?
						elem.lang :
						elem.getAttribute("xml:lang") || elem.getAttribute("lang")) ) {

						elemLang = elemLang.toLowerCase();
						return elemLang === lang || elemLang.indexOf( lang + "-" ) === 0;
					}
				} while ( (elem = elem.parentNode) && elem.nodeType === 1 );
				return false;
			};
		}),

		// Miscellaneous
		"target": function( elem ) {
			var hash = window.location && window.location.hash;
			return hash && hash.slice( 1 ) === elem.id;
		},

		"root": function( elem ) {
			return elem === docElem;
		},

		"focus": function( elem ) {
			return elem === document.activeElement && (!document.hasFocus || document.hasFocus()) && !!(elem.type || elem.href || ~elem.tabIndex);
		},

		// Boolean properties
		"enabled": function( elem ) {
			return elem.disabled === false;
		},

		"disabled": function( elem ) {
			return elem.disabled === true;
		},

		"checked": function( elem ) {
			// In CSS3, :checked should return both checked and selected elements
			// http://www.w3.org/TR/2011/REC-css3-selectors-20110929/#checked
			var nodeName = elem.nodeName.toLowerCase();
			return (nodeName === "input" && !!elem.checked) || (nodeName === "option" && !!elem.selected);
		},

		"selected": function( elem ) {
			// Accessing this property makes selected-by-default
			// options in Safari work properly
			if ( elem.parentNode ) {
				elem.parentNode.selectedIndex;
			}

			return elem.selected === true;
		},

		// Contents
		"empty": function( elem ) {
			// http://www.w3.org/TR/selectors/#empty-pseudo
			// :empty is negated by element (1) or content nodes (text: 3; cdata: 4; entity ref: 5),
			//   but not by others (comment: 8; processing instruction: 7; etc.)
			// nodeType < 6 works because attributes (2) do not appear as children
			for ( elem = elem.firstChild; elem; elem = elem.nextSibling ) {
				if ( elem.nodeType < 6 ) {
					return false;
				}
			}
			return true;
		},

		"parent": function( elem ) {
			return !Expr.pseudos["empty"]( elem );
		},

		// Element/input types
		"header": function( elem ) {
			return rheader.test( elem.nodeName );
		},

		"input": function( elem ) {
			return rinputs.test( elem.nodeName );
		},

		"button": function( elem ) {
			var name = elem.nodeName.toLowerCase();
			return name === "input" && elem.type === "button" || name === "button";
		},

		"text": function( elem ) {
			var attr;
			return elem.nodeName.toLowerCase() === "input" &&
				elem.type === "text" &&

				// Support: IE<8
				// New HTML5 attribute values (e.g., "search") appear with elem.type === "text"
				( (attr = elem.getAttribute("type")) == null || attr.toLowerCase() === "text" );
		},

		// Position-in-collection
		"first": createPositionalPseudo(function() {
			return [ 0 ];
		}),

		"last": createPositionalPseudo(function( matchIndexes, length ) {
			return [ length - 1 ];
		}),

		"eq": createPositionalPseudo(function( matchIndexes, length, argument ) {
			return [ argument < 0 ? argument + length : argument ];
		}),

		"even": createPositionalPseudo(function( matchIndexes, length ) {
			var i = 0;
			for ( ; i < length; i += 2 ) {
				matchIndexes.push( i );
			}
			return matchIndexes;
		}),

		"odd": createPositionalPseudo(function( matchIndexes, length ) {
			var i = 1;
			for ( ; i < length; i += 2 ) {
				matchIndexes.push( i );
			}
			return matchIndexes;
		}),

		"lt": createPositionalPseudo(function( matchIndexes, length, argument ) {
			var i = argument < 0 ? argument + length : argument;
			for ( ; --i >= 0; ) {
				matchIndexes.push( i );
			}
			return matchIndexes;
		}),

		"gt": createPositionalPseudo(function( matchIndexes, length, argument ) {
			var i = argument < 0 ? argument + length : argument;
			for ( ; ++i < length; ) {
				matchIndexes.push( i );
			}
			return matchIndexes;
		})
	}
};

Expr.pseudos["nth"] = Expr.pseudos["eq"];

// Add button/input type pseudos
for ( i in { radio: true, checkbox: true, file: true, password: true, image: true } ) {
	Expr.pseudos[ i ] = createInputPseudo( i );
}
for ( i in { submit: true, reset: true } ) {
	Expr.pseudos[ i ] = createButtonPseudo( i );
}

// Easy API for creating new setFilters
function setFilters() {}
setFilters.prototype = Expr.filters = Expr.pseudos;
Expr.setFilters = new setFilters();

tokenize = Sizzle.tokenize = function( selector, parseOnly ) {
	var matched, match, tokens, type,
		soFar, groups, preFilters,
		cached = tokenCache[ selector + " " ];

	if ( cached ) {
		return parseOnly ? 0 : cached.slice( 0 );
	}

	soFar = selector;
	groups = [];
	preFilters = Expr.preFilter;

	while ( soFar ) {

		// Comma and first run
		if ( !matched || (match = rcomma.exec( soFar )) ) {
			if ( match ) {
				// Don't consume trailing commas as valid
				soFar = soFar.slice( match[0].length ) || soFar;
			}
			groups.push( (tokens = []) );
		}

		matched = false;

		// Combinators
		if ( (match = rcombinators.exec( soFar )) ) {
			matched = match.shift();
			tokens.push({
				value: matched,
				// Cast descendant combinators to space
				type: match[0].replace( rtrim, " " )
			});
			soFar = soFar.slice( matched.length );
		}

		// Filters
		for ( type in Expr.filter ) {
			if ( (match = matchExpr[ type ].exec( soFar )) && (!preFilters[ type ] ||
				(match = preFilters[ type ]( match ))) ) {
				matched = match.shift();
				tokens.push({
					value: matched,
					type: type,
					matches: match
				});
				soFar = soFar.slice( matched.length );
			}
		}

		if ( !matched ) {
			break;
		}
	}

	// Return the length of the invalid excess
	// if we're just parsing
	// Otherwise, throw an error or return tokens
	return parseOnly ?
		soFar.length :
		soFar ?
			Sizzle.error( selector ) :
			// Cache the tokens
			tokenCache( selector, groups ).slice( 0 );
};

function toSelector( tokens ) {
	var i = 0,
		len = tokens.length,
		selector = "";
	for ( ; i < len; i++ ) {
		selector += tokens[i].value;
	}
	return selector;
}

function addCombinator( matcher, combinator, base ) {
	var dir = combinator.dir,
		checkNonElements = base && dir === "parentNode",
		doneName = done++;

	return combinator.first ?
		// Check against closest ancestor/preceding element
		function( elem, context, xml ) {
			while ( (elem = elem[ dir ]) ) {
				if ( elem.nodeType === 1 || checkNonElements ) {
					return matcher( elem, context, xml );
				}
			}
		} :

		// Check against all ancestor/preceding elements
		function( elem, context, xml ) {
			var oldCache, outerCache,
				newCache = [ dirruns, doneName ];

			// We can't set arbitrary data on XML nodes, so they don't benefit from dir caching
			if ( xml ) {
				while ( (elem = elem[ dir ]) ) {
					if ( elem.nodeType === 1 || checkNonElements ) {
						if ( matcher( elem, context, xml ) ) {
							return true;
						}
					}
				}
			} else {
				while ( (elem = elem[ dir ]) ) {
					if ( elem.nodeType === 1 || checkNonElements ) {
						outerCache = elem[ expando ] || (elem[ expando ] = {});
						if ( (oldCache = outerCache[ dir ]) &&
							oldCache[ 0 ] === dirruns && oldCache[ 1 ] === doneName ) {

							// Assign to newCache so results back-propagate to previous elements
							return (newCache[ 2 ] = oldCache[ 2 ]);
						} else {
							// Reuse newcache so results back-propagate to previous elements
							outerCache[ dir ] = newCache;

							// A match means we're done; a fail means we have to keep checking
							if ( (newCache[ 2 ] = matcher( elem, context, xml )) ) {
								return true;
							}
						}
					}
				}
			}
		};
}

function elementMatcher( matchers ) {
	return matchers.length > 1 ?
		function( elem, context, xml ) {
			var i = matchers.length;
			while ( i-- ) {
				if ( !matchers[i]( elem, context, xml ) ) {
					return false;
				}
			}
			return true;
		} :
		matchers[0];
}

function multipleContexts( selector, contexts, results ) {
	var i = 0,
		len = contexts.length;
	for ( ; i < len; i++ ) {
		Sizzle( selector, contexts[i], results );
	}
	return results;
}

function condense( unmatched, map, filter, context, xml ) {
	var elem,
		newUnmatched = [],
		i = 0,
		len = unmatched.length,
		mapped = map != null;

	for ( ; i < len; i++ ) {
		if ( (elem = unmatched[i]) ) {
			if ( !filter || filter( elem, context, xml ) ) {
				newUnmatched.push( elem );
				if ( mapped ) {
					map.push( i );
				}
			}
		}
	}

	return newUnmatched;
}

function setMatcher( preFilter, selector, matcher, postFilter, postFinder, postSelector ) {
	if ( postFilter && !postFilter[ expando ] ) {
		postFilter = setMatcher( postFilter );
	}
	if ( postFinder && !postFinder[ expando ] ) {
		postFinder = setMatcher( postFinder, postSelector );
	}
	return markFunction(function( seed, results, context, xml ) {
		var temp, i, elem,
			preMap = [],
			postMap = [],
			preexisting = results.length,

			// Get initial elements from seed or context
			elems = seed || multipleContexts( selector || "*", context.nodeType ? [ context ] : context, [] ),

			// Prefilter to get matcher input, preserving a map for seed-results synchronization
			matcherIn = preFilter && ( seed || !selector ) ?
				condense( elems, preMap, preFilter, context, xml ) :
				elems,

			matcherOut = matcher ?
				// If we have a postFinder, or filtered seed, or non-seed postFilter or preexisting results,
				postFinder || ( seed ? preFilter : preexisting || postFilter ) ?

					// ...intermediate processing is necessary
					[] :

					// ...otherwise use results directly
					results :
				matcherIn;

		// Find primary matches
		if ( matcher ) {
			matcher( matcherIn, matcherOut, context, xml );
		}

		// Apply postFilter
		if ( postFilter ) {
			temp = condense( matcherOut, postMap );
			postFilter( temp, [], context, xml );

			// Un-match failing elements by moving them back to matcherIn
			i = temp.length;
			while ( i-- ) {
				if ( (elem = temp[i]) ) {
					matcherOut[ postMap[i] ] = !(matcherIn[ postMap[i] ] = elem);
				}
			}
		}

		if ( seed ) {
			if ( postFinder || preFilter ) {
				if ( postFinder ) {
					// Get the final matcherOut by condensing this intermediate into postFinder contexts
					temp = [];
					i = matcherOut.length;
					while ( i-- ) {
						if ( (elem = matcherOut[i]) ) {
							// Restore matcherIn since elem is not yet a final match
							temp.push( (matcherIn[i] = elem) );
						}
					}
					postFinder( null, (matcherOut = []), temp, xml );
				}

				// Move matched elements from seed to results to keep them synchronized
				i = matcherOut.length;
				while ( i-- ) {
					if ( (elem = matcherOut[i]) &&
						(temp = postFinder ? indexOf.call( seed, elem ) : preMap[i]) > -1 ) {

						seed[temp] = !(results[temp] = elem);
					}
				}
			}

		// Add elements to results, through postFinder if defined
		} else {
			matcherOut = condense(
				matcherOut === results ?
					matcherOut.splice( preexisting, matcherOut.length ) :
					matcherOut
			);
			if ( postFinder ) {
				postFinder( null, results, matcherOut, xml );
			} else {
				push.apply( results, matcherOut );
			}
		}
	});
}

function matcherFromTokens( tokens ) {
	var checkContext, matcher, j,
		len = tokens.length,
		leadingRelative = Expr.relative[ tokens[0].type ],
		implicitRelative = leadingRelative || Expr.relative[" "],
		i = leadingRelative ? 1 : 0,

		// The foundational matcher ensures that elements are reachable from top-level context(s)
		matchContext = addCombinator( function( elem ) {
			return elem === checkContext;
		}, implicitRelative, true ),
		matchAnyContext = addCombinator( function( elem ) {
			return indexOf.call( checkContext, elem ) > -1;
		}, implicitRelative, true ),
		matchers = [ function( elem, context, xml ) {
			return ( !leadingRelative && ( xml || context !== outermostContext ) ) || (
				(checkContext = context).nodeType ?
					matchContext( elem, context, xml ) :
					matchAnyContext( elem, context, xml ) );
		} ];

	for ( ; i < len; i++ ) {
		if ( (matcher = Expr.relative[ tokens[i].type ]) ) {
			matchers = [ addCombinator(elementMatcher( matchers ), matcher) ];
		} else {
			matcher = Expr.filter[ tokens[i].type ].apply( null, tokens[i].matches );

			// Return special upon seeing a positional matcher
			if ( matcher[ expando ] ) {
				// Find the next relative operator (if any) for proper handling
				j = ++i;
				for ( ; j < len; j++ ) {
					if ( Expr.relative[ tokens[j].type ] ) {
						break;
					}
				}
				return setMatcher(
					i > 1 && elementMatcher( matchers ),
					i > 1 && toSelector(
						// If the preceding token was a descendant combinator, insert an implicit any-element `*`
						tokens.slice( 0, i - 1 ).concat({ value: tokens[ i - 2 ].type === " " ? "*" : "" })
					).replace( rtrim, "$1" ),
					matcher,
					i < j && matcherFromTokens( tokens.slice( i, j ) ),
					j < len && matcherFromTokens( (tokens = tokens.slice( j )) ),
					j < len && toSelector( tokens )
				);
			}
			matchers.push( matcher );
		}
	}

	return elementMatcher( matchers );
}

function matcherFromGroupMatchers( elementMatchers, setMatchers ) {
	var bySet = setMatchers.length > 0,
		byElement = elementMatchers.length > 0,
		superMatcher = function( seed, context, xml, results, outermost ) {
			var elem, j, matcher,
				matchedCount = 0,
				i = "0",
				unmatched = seed && [],
				setMatched = [],
				contextBackup = outermostContext,
				// We must always have either seed elements or outermost context
				elems = seed || byElement && Expr.find["TAG"]( "*", outermost ),
				// Use integer dirruns iff this is the outermost matcher
				dirrunsUnique = (dirruns += contextBackup == null ? 1 : Math.random() || 0.1),
				len = elems.length;

			if ( outermost ) {
				outermostContext = context !== document && context;
			}

			// Add elements passing elementMatchers directly to results
			// Keep `i` a string if there are no elements so `matchedCount` will be "00" below
			// Support: IE<9, Safari
			// Tolerate NodeList properties (IE: "length"; Safari: <number>) matching elements by id
			for ( ; i !== len && (elem = elems[i]) != null; i++ ) {
				if ( byElement && elem ) {
					j = 0;
					while ( (matcher = elementMatchers[j++]) ) {
						if ( matcher( elem, context, xml ) ) {
							results.push( elem );
							break;
						}
					}
					if ( outermost ) {
						dirruns = dirrunsUnique;
					}
				}

				// Track unmatched elements for set filters
				if ( bySet ) {
					// They will have gone through all possible matchers
					if ( (elem = !matcher && elem) ) {
						matchedCount--;
					}

					// Lengthen the array for every element, matched or not
					if ( seed ) {
						unmatched.push( elem );
					}
				}
			}

			// Apply set filters to unmatched elements
			matchedCount += i;
			if ( bySet && i !== matchedCount ) {
				j = 0;
				while ( (matcher = setMatchers[j++]) ) {
					matcher( unmatched, setMatched, context, xml );
				}

				if ( seed ) {
					// Reintegrate element matches to eliminate the need for sorting
					if ( matchedCount > 0 ) {
						while ( i-- ) {
							if ( !(unmatched[i] || setMatched[i]) ) {
								setMatched[i] = pop.call( results );
							}
						}
					}

					// Discard index placeholder values to get only actual matches
					setMatched = condense( setMatched );
				}

				// Add matches to results
				push.apply( results, setMatched );

				// Seedless set matches succeeding multiple successful matchers stipulate sorting
				if ( outermost && !seed && setMatched.length > 0 &&
					( matchedCount + setMatchers.length ) > 1 ) {

					Sizzle.uniqueSort( results );
				}
			}

			// Override manipulation of globals by nested matchers
			if ( outermost ) {
				dirruns = dirrunsUnique;
				outermostContext = contextBackup;
			}

			return unmatched;
		};

	return bySet ?
		markFunction( superMatcher ) :
		superMatcher;
}

compile = Sizzle.compile = function( selector, match /* Internal Use Only */ ) {
	var i,
		setMatchers = [],
		elementMatchers = [],
		cached = compilerCache[ selector + " " ];

	if ( !cached ) {
		// Generate a function of recursive functions that can be used to check each element
		if ( !match ) {
			match = tokenize( selector );
		}
		i = match.length;
		while ( i-- ) {
			cached = matcherFromTokens( match[i] );
			if ( cached[ expando ] ) {
				setMatchers.push( cached );
			} else {
				elementMatchers.push( cached );
			}
		}

		// Cache the compiled function
		cached = compilerCache( selector, matcherFromGroupMatchers( elementMatchers, setMatchers ) );

		// Save selector and tokenization
		cached.selector = selector;
	}
	return cached;
};

/**
 * A low-level selection function that works with Sizzle's compiled
 *  selector functions
 * @param {String|Function} selector A selector or a pre-compiled
 *  selector function built with Sizzle.compile
 * @param {Element} context
 * @param {Array} [results]
 * @param {Array} [seed] A set of elements to match against
 */
select = Sizzle.select = function( selector, context, results, seed ) {
	var i, tokens, token, type, find,
		compiled = typeof selector === "function" && selector,
		match = !seed && tokenize( (selector = compiled.selector || selector) );

	results = results || [];

	// Try to minimize operations if there is no seed and only one group
	if ( match.length === 1 ) {

		// Take a shortcut and set the context if the root selector is an ID
		tokens = match[0] = match[0].slice( 0 );
		if ( tokens.length > 2 && (token = tokens[0]).type === "ID" &&
				support.getById && context.nodeType === 9 && documentIsHTML &&
				Expr.relative[ tokens[1].type ] ) {

			context = ( Expr.find["ID"]( token.matches[0].replace(runescape, funescape), context ) || [] )[0];
			if ( !context ) {
				return results;

			// Precompiled matchers will still verify ancestry, so step up a level
			} else if ( compiled ) {
				context = context.parentNode;
			}

			selector = selector.slice( tokens.shift().value.length );
		}

		// Fetch a seed set for right-to-left matching
		i = matchExpr["needsContext"].test( selector ) ? 0 : tokens.length;
		while ( i-- ) {
			token = tokens[i];

			// Abort if we hit a combinator
			if ( Expr.relative[ (type = token.type) ] ) {
				break;
			}
			if ( (find = Expr.find[ type ]) ) {
				// Search, expanding context for leading sibling combinators
				if ( (seed = find(
					token.matches[0].replace( runescape, funescape ),
					rsibling.test( tokens[0].type ) && testContext( context.parentNode ) || context
				)) ) {

					// If seed is empty or no tokens remain, we can return early
					tokens.splice( i, 1 );
					selector = seed.length && toSelector( tokens );
					if ( !selector ) {
						push.apply( results, seed );
						return results;
					}

					break;
				}
			}
		}
	}

	// Compile and execute a filtering function if one is not provided
	// Provide `match` to avoid retokenization if we modified the selector above
	( compiled || compile( selector, match ) )(
		seed,
		context,
		!documentIsHTML,
		results,
		rsibling.test( selector ) && testContext( context.parentNode ) || context
	);
	return results;
};

// One-time assignments

// Sort stability
support.sortStable = expando.split("").sort( sortOrder ).join("") === expando;

// Support: Chrome<14
// Always assume duplicates if they aren't passed to the comparison function
support.detectDuplicates = !!hasDuplicate;

// Initialize against the default document
setDocument();

// Support: Webkit<537.32 - Safari 6.0.3/Chrome 25 (fixed in Chrome 27)
// Detached nodes confoundingly follow *each other*
support.sortDetached = assert(function( div1 ) {
	// Should return 1, but returns 4 (following)
	return div1.compareDocumentPosition( document.createElement("div") ) & 1;
});

// Support: IE<8
// Prevent attribute/property "interpolation"
// http://msdn.microsoft.com/en-us/library/ms536429%28VS.85%29.aspx
if ( !assert(function( div ) {
	div.innerHTML = "<a href='#'></a>";
	return div.firstChild.getAttribute("href") === "#" ;
}) ) {
	addHandle( "type|href|height|width", function( elem, name, isXML ) {
		if ( !isXML ) {
			return elem.getAttribute( name, name.toLowerCase() === "type" ? 1 : 2 );
		}
	});
}

// Support: IE<9
// Use defaultValue in place of getAttribute("value")
if ( !support.attributes || !assert(function( div ) {
	div.innerHTML = "<input/>";
	div.firstChild.setAttribute( "value", "" );
	return div.firstChild.getAttribute( "value" ) === "";
}) ) {
	addHandle( "value", function( elem, name, isXML ) {
		if ( !isXML && elem.nodeName.toLowerCase() === "input" ) {
			return elem.defaultValue;
		}
	});
}

// Support: IE<9
// Use getAttributeNode to fetch booleans when getAttribute lies
if ( !assert(function( div ) {
	return div.getAttribute("disabled") == null;
}) ) {
	addHandle( booleans, function( elem, name, isXML ) {
		var val;
		if ( !isXML ) {
			return elem[ name ] === true ? name.toLowerCase() :
					(val = elem.getAttributeNode( name )) && val.specified ?
					val.value :
				null;
		}
	});
}

return Sizzle;

})( window );


jQuery.find = Sizzle;
jQuery.expr = Sizzle.selectors;
jQuery.expr[":"] = jQuery.expr.pseudos;
jQuery.unique = Sizzle.uniqueSort;
jQuery.text = Sizzle.getText;
jQuery.isXMLDoc = Sizzle.isXML;
jQuery.contains = Sizzle.contains;


var rneedsContext = jQuery.expr.match.needsContext;

var rsingleTag = (/^<(\w+)\s*\/?>(?:<\/\1>|)$/);


var risSimple = /^.[^:#\[\.,]*$/;

// Implement the identical functionality for filter and not
function winnow( elements, qualifier, not ) {
	if ( jQuery.isFunction( qualifier ) ) {
		return jQuery.grep( elements, function( elem, i ) {
			/* jshint -W018 */
			return !!qualifier.call( elem, i, elem ) !== not;
		});

	}

	if ( qualifier.nodeType ) {
		return jQuery.grep( elements, function( elem ) {
			return ( elem === qualifier ) !== not;
		});

	}

	if ( typeof qualifier === "string" ) {
		if ( risSimple.test( qualifier ) ) {
			return jQuery.filter( qualifier, elements, not );
		}

		qualifier = jQuery.filter( qualifier, elements );
	}

	return jQuery.grep( elements, function( elem ) {
		return ( jQuery.inArray( elem, qualifier ) >= 0 ) !== not;
	});
}

jQuery.filter = function( expr, elems, not ) {
	var elem = elems[ 0 ];

	if ( not ) {
		expr = ":not(" + expr + ")";
	}

	return elems.length === 1 && elem.nodeType === 1 ?
		jQuery.find.matchesSelector( elem, expr ) ? [ elem ] : [] :
		jQuery.find.matches( expr, jQuery.grep( elems, function( elem ) {
			return elem.nodeType === 1;
		}));
};

jQuery.fn.extend({
	find: function( selector ) {
		var i,
			ret = [],
			self = this,
			len = self.length;

		if ( typeof selector !== "string" ) {
			return this.pushStack( jQuery( selector ).filter(function() {
				for ( i = 0; i < len; i++ ) {
					if ( jQuery.contains( self[ i ], this ) ) {
						return true;
					}
				}
			}) );
		}

		for ( i = 0; i < len; i++ ) {
			jQuery.find( selector, self[ i ], ret );
		}

		// Needed because $( selector, context ) becomes $( context ).find( selector )
		ret = this.pushStack( len > 1 ? jQuery.unique( ret ) : ret );
		ret.selector = this.selector ? this.selector + " " + selector : selector;
		return ret;
	},
	filter: function( selector ) {
		return this.pushStack( winnow(this, selector || [], false) );
	},
	not: function( selector ) {
		return this.pushStack( winnow(this, selector || [], true) );
	},
	is: function( selector ) {
		return !!winnow(
			this,

			// If this is a positional/relative selector, check membership in the returned set
			// so $("p:first").is("p:last") won't return true for a doc with two "p".
			typeof selector === "string" && rneedsContext.test( selector ) ?
				jQuery( selector ) :
				selector || [],
			false
		).length;
	}
});


// Initialize a jQuery object


// A central reference to the root jQuery(document)
var rootjQuery,

	// Use the correct document accordingly with window argument (sandbox)
	document = window.document,

	// A simple way to check for HTML strings
	// Prioritize #id over <tag> to avoid XSS via location.hash (#9521)
	// Strict HTML recognition (#11290: must start with <)
	rquickExpr = /^(?:\s*(<[\w\W]+>)[^>]*|#([\w-]*))$/,

	init = jQuery.fn.init = function( selector, context ) {
		var match, elem;

		// HANDLE: $(""), $(null), $(undefined), $(false)
		if ( !selector ) {
			return this;
		}

		// Handle HTML strings
		if ( typeof selector === "string" ) {
			if ( selector.charAt(0) === "<" && selector.charAt( selector.length - 1 ) === ">" && selector.length >= 3 ) {
				// Assume that strings that start and end with <> are HTML and skip the regex check
				match = [ null, selector, null ];

			} else {
				match = rquickExpr.exec( selector );
			}

			// Match html or make sure no context is specified for #id
			if ( match && (match[1] || !context) ) {

				// HANDLE: $(html) -> $(array)
				if ( match[1] ) {
					context = context instanceof jQuery ? context[0] : context;

					// scripts is true for back-compat
					// Intentionally let the error be thrown if parseHTML is not present
					jQuery.merge( this, jQuery.parseHTML(
						match[1],
						context && context.nodeType ? context.ownerDocument || context : document,
						true
					) );

					// HANDLE: $(html, props)
					if ( rsingleTag.test( match[1] ) && jQuery.isPlainObject( context ) ) {
						for ( match in context ) {
							// Properties of context are called as methods if possible
							if ( jQuery.isFunction( this[ match ] ) ) {
								this[ match ]( context[ match ] );

							// ...and otherwise set as attributes
							} else {
								this.attr( match, context[ match ] );
							}
						}
					}

					return this;

				// HANDLE: $(#id)
				} else {
					elem = document.getElementById( match[2] );

					// Check parentNode to catch when Blackberry 4.6 returns
					// nodes that are no longer in the document #6963
					if ( elem && elem.parentNode ) {
						// Handle the case where IE and Opera return items
						// by name instead of ID
						if ( elem.id !== match[2] ) {
							return rootjQuery.find( selector );
						}

						// Otherwise, we inject the element directly into the jQuery object
						this.length = 1;
						this[0] = elem;
					}

					this.context = document;
					this.selector = selector;
					return this;
				}

			// HANDLE: $(expr, $(...))
			} else if ( !context || context.jquery ) {
				return ( context || rootjQuery ).find( selector );

			// HANDLE: $(expr, context)
			// (which is just equivalent to: $(context).find(expr)
			} else {
				return this.constructor( context ).find( selector );
			}

		// HANDLE: $(DOMElement)
		} else if ( selector.nodeType ) {
			this.context = this[0] = selector;
			this.length = 1;
			return this;

		// HANDLE: $(function)
		// Shortcut for document ready
		} else if ( jQuery.isFunction( selector ) ) {
			return typeof rootjQuery.ready !== "undefined" ?
				rootjQuery.ready( selector ) :
				// Execute immediately if ready is not present
				selector( jQuery );
		}

		if ( selector.selector !== undefined ) {
			this.selector = selector.selector;
			this.context = selector.context;
		}

		return jQuery.makeArray( selector, this );
	};

// Give the init function the jQuery prototype for later instantiation
init.prototype = jQuery.fn;

// Initialize central reference
rootjQuery = jQuery( document );


var rparentsprev = /^(?:parents|prev(?:Until|All))/,
	// methods guaranteed to produce a unique set when starting from a unique set
	guaranteedUnique = {
		children: true,
		contents: true,
		next: true,
		prev: true
	};

jQuery.extend({
	dir: function( elem, dir, until ) {
		var matched = [],
			cur = elem[ dir ];

		while ( cur && cur.nodeType !== 9 && (until === undefined || cur.nodeType !== 1 || !jQuery( cur ).is( until )) ) {
			if ( cur.nodeType === 1 ) {
				matched.push( cur );
			}
			cur = cur[dir];
		}
		return matched;
	},

	sibling: function( n, elem ) {
		var r = [];

		for ( ; n; n = n.nextSibling ) {
			if ( n.nodeType === 1 && n !== elem ) {
				r.push( n );
			}
		}

		return r;
	}
});

jQuery.fn.extend({
	has: function( target ) {
		var i,
			targets = jQuery( target, this ),
			len = targets.length;

		return this.filter(function() {
			for ( i = 0; i < len; i++ ) {
				if ( jQuery.contains( this, targets[i] ) ) {
					return true;
				}
			}
		});
	},

	closest: function( selectors, context ) {
		var cur,
			i = 0,
			l = this.length,
			matched = [],
			pos = rneedsContext.test( selectors ) || typeof selectors !== "string" ?
				jQuery( selectors, context || this.context ) :
				0;

		for ( ; i < l; i++ ) {
			for ( cur = this[i]; cur && cur !== context; cur = cur.parentNode ) {
				// Always skip document fragments
				if ( cur.nodeType < 11 && (pos ?
					pos.index(cur) > -1 :

					// Don't pass non-elements to Sizzle
					cur.nodeType === 1 &&
						jQuery.find.matchesSelector(cur, selectors)) ) {

					matched.push( cur );
					break;
				}
			}
		}

		return this.pushStack( matched.length > 1 ? jQuery.unique( matched ) : matched );
	},

	// Determine the position of an element within
	// the matched set of elements
	index: function( elem ) {

		// No argument, return index in parent
		if ( !elem ) {
			return ( this[0] && this[0].parentNode ) ? this.first().prevAll().length : -1;
		}

		// index in selector
		if ( typeof elem === "string" ) {
			return jQuery.inArray( this[0], jQuery( elem ) );
		}

		// Locate the position of the desired element
		return jQuery.inArray(
			// If it receives a jQuery object, the first element is used
			elem.jquery ? elem[0] : elem, this );
	},

	add: function( selector, context ) {
		return this.pushStack(
			jQuery.unique(
				jQuery.merge( this.get(), jQuery( selector, context ) )
			)
		);
	},

	addBack: function( selector ) {
		return this.add( selector == null ?
			this.prevObject : this.prevObject.filter(selector)
		);
	}
});

function sibling( cur, dir ) {
	do {
		cur = cur[ dir ];
	} while ( cur && cur.nodeType !== 1 );

	return cur;
}

jQuery.each({
	parent: function( elem ) {
		var parent = elem.parentNode;
		return parent && parent.nodeType !== 11 ? parent : null;
	},
	parents: function( elem ) {
		return jQuery.dir( elem, "parentNode" );
	},
	parentsUntil: function( elem, i, until ) {
		return jQuery.dir( elem, "parentNode", until );
	},
	next: function( elem ) {
		return sibling( elem, "nextSibling" );
	},
	prev: function( elem ) {
		return sibling( elem, "previousSibling" );
	},
	nextAll: function( elem ) {
		return jQuery.dir( elem, "nextSibling" );
	},
	prevAll: function( elem ) {
		return jQuery.dir( elem, "previousSibling" );
	},
	nextUntil: function( elem, i, until ) {
		return jQuery.dir( elem, "nextSibling", until );
	},
	prevUntil: function( elem, i, until ) {
		return jQuery.dir( elem, "previousSibling", until );
	},
	siblings: function( elem ) {
		return jQuery.sibling( ( elem.parentNode || {} ).firstChild, elem );
	},
	children: function( elem ) {
		return jQuery.sibling( elem.firstChild );
	},
	contents: function( elem ) {
		return jQuery.nodeName( elem, "iframe" ) ?
			elem.contentDocument || elem.contentWindow.document :
			jQuery.merge( [], elem.childNodes );
	}
}, function( name, fn ) {
	jQuery.fn[ name ] = function( until, selector ) {
		var ret = jQuery.map( this, fn, until );

		if ( name.slice( -5 ) !== "Until" ) {
			selector = until;
		}

		if ( selector && typeof selector === "string" ) {
			ret = jQuery.filter( selector, ret );
		}

		if ( this.length > 1 ) {
			// Remove duplicates
			if ( !guaranteedUnique[ name ] ) {
				ret = jQuery.unique( ret );
			}

			// Reverse order for parents* and prev-derivatives
			if ( rparentsprev.test( name ) ) {
				ret = ret.reverse();
			}
		}

		return this.pushStack( ret );
	};
});
var rnotwhite = (/\S+/g);


// String to Object options format cache
var optionsCache = {};

// Convert String-formatted options into Object-formatted ones and store in cache
function createOptions( options ) {
	var object = optionsCache[ options ] = {};
	jQuery.each( options.match( rnotwhite ) || [], function( _, flag ) {
		object[ flag ] = true;
	});
	return object;
}

/*
 * Create a callback list using the following parameters:
 *
 *	options: an optional list of space-separated options that will change how
 *			the callback list behaves or a more traditional option object
 *
 * By default a callback list will act like an event callback list and can be
 * "fired" multiple times.
 *
 * Possible options:
 *
 *	once:			will ensure the callback list can only be fired once (like a Deferred)
 *
 *	memory:			will keep track of previous values and will call any callback added
 *					after the list has been fired right away with the latest "memorized"
 *					values (like a Deferred)
 *
 *	unique:			will ensure a callback can only be added once (no duplicate in the list)
 *
 *	stopOnFalse:	interrupt callings when a callback returns false
 *
 */
jQuery.Callbacks = function( options ) {

	// Convert options from String-formatted to Object-formatted if needed
	// (we check in cache first)
	options = typeof options === "string" ?
		( optionsCache[ options ] || createOptions( options ) ) :
		jQuery.extend( {}, options );

	var // Flag to know if list is currently firing
		firing,
		// Last fire value (for non-forgettable lists)
		memory,
		// Flag to know if list was already fired
		fired,
		// End of the loop when firing
		firingLength,
		// Index of currently firing callback (modified by remove if needed)
		firingIndex,
		// First callback to fire (used internally by add and fireWith)
		firingStart,
		// Actual callback list
		list = [],
		// Stack of fire calls for repeatable lists
		stack = !options.once && [],
		// Fire callbacks
		fire = function( data ) {
			memory = options.memory && data;
			fired = true;
			firingIndex = firingStart || 0;
			firingStart = 0;
			firingLength = list.length;
			firing = true;
			for ( ; list && firingIndex < firingLength; firingIndex++ ) {
				if ( list[ firingIndex ].apply( data[ 0 ], data[ 1 ] ) === false && options.stopOnFalse ) {
					memory = false; // To prevent further calls using add
					break;
				}
			}
			firing = false;
			if ( list ) {
				if ( stack ) {
					if ( stack.length ) {
						fire( stack.shift() );
					}
				} else if ( memory ) {
					list = [];
				} else {
					self.disable();
				}
			}
		},
		// Actual Callbacks object
		self = {
			// Add a callback or a collection of callbacks to the list
			add: function() {
				if ( list ) {
					// First, we save the current length
					var start = list.length;
					(function add( args ) {
						jQuery.each( args, function( _, arg ) {
							var type = jQuery.type( arg );
							if ( type === "function" ) {
								if ( !options.unique || !self.has( arg ) ) {
									list.push( arg );
								}
							} else if ( arg && arg.length && type !== "string" ) {
								// Inspect recursively
								add( arg );
							}
						});
					})( arguments );
					// Do we need to add the callbacks to the
					// current firing batch?
					if ( firing ) {
						firingLength = list.length;
					// With memory, if we're not firing then
					// we should call right away
					} else if ( memory ) {
						firingStart = start;
						fire( memory );
					}
				}
				return this;
			},
			// Remove a callback from the list
			remove: function() {
				if ( list ) {
					jQuery.each( arguments, function( _, arg ) {
						var index;
						while ( ( index = jQuery.inArray( arg, list, index ) ) > -1 ) {
							list.splice( index, 1 );
							// Handle firing indexes
							if ( firing ) {
								if ( index <= firingLength ) {
									firingLength--;
								}
								if ( index <= firingIndex ) {
									firingIndex--;
								}
							}
						}
					});
				}
				return this;
			},
			// Check if a given callback is in the list.
			// If no argument is given, return whether or not list has callbacks attached.
			has: function( fn ) {
				return fn ? jQuery.inArray( fn, list ) > -1 : !!( list && list.length );
			},
			// Remove all callbacks from the list
			empty: function() {
				list = [];
				firingLength = 0;
				return this;
			},
			// Have the list do nothing anymore
			disable: function() {
				list = stack = memory = undefined;
				return this;
			},
			// Is it disabled?
			disabled: function() {
				return !list;
			},
			// Lock the list in its current state
			lock: function() {
				stack = undefined;
				if ( !memory ) {
					self.disable();
				}
				return this;
			},
			// Is it locked?
			locked: function() {
				return !stack;
			},
			// Call all callbacks with the given context and arguments
			fireWith: function( context, args ) {
				if ( list && ( !fired || stack ) ) {
					args = args || [];
					args = [ context, args.slice ? args.slice() : args ];
					if ( firing ) {
						stack.push( args );
					} else {
						fire( args );
					}
				}
				return this;
			},
			// Call all the callbacks with the given arguments
			fire: function() {
				self.fireWith( this, arguments );
				return this;
			},
			// To know if the callbacks have already been called at least once
			fired: function() {
				return !!fired;
			}
		};

	return self;
};


jQuery.extend({

	Deferred: function( func ) {
		var tuples = [
				// action, add listener, listener list, final state
				[ "resolve", "done", jQuery.Callbacks("once memory"), "resolved" ],
				[ "reject", "fail", jQuery.Callbacks("once memory"), "rejected" ],
				[ "notify", "progress", jQuery.Callbacks("memory") ]
			],
			state = "pending",
			promise = {
				state: function() {
					return state;
				},
				always: function() {
					deferred.done( arguments ).fail( arguments );
					return this;
				},
				then: function( /* fnDone, fnFail, fnProgress */ ) {
					var fns = arguments;
					return jQuery.Deferred(function( newDefer ) {
						jQuery.each( tuples, function( i, tuple ) {
							var fn = jQuery.isFunction( fns[ i ] ) && fns[ i ];
							// deferred[ done | fail | progress ] for forwarding actions to newDefer
							deferred[ tuple[1] ](function() {
								var returned = fn && fn.apply( this, arguments );
								if ( returned && jQuery.isFunction( returned.promise ) ) {
									returned.promise()
										.done( newDefer.resolve )
										.fail( newDefer.reject )
										.progress( newDefer.notify );
								} else {
									newDefer[ tuple[ 0 ] + "With" ]( this === promise ? newDefer.promise() : this, fn ? [ returned ] : arguments );
								}
							});
						});
						fns = null;
					}).promise();
				},
				// Get a promise for this deferred
				// If obj is provided, the promise aspect is added to the object
				promise: function( obj ) {
					return obj != null ? jQuery.extend( obj, promise ) : promise;
				}
			},
			deferred = {};

		// Keep pipe for back-compat
		promise.pipe = promise.then;

		// Add list-specific methods
		jQuery.each( tuples, function( i, tuple ) {
			var list = tuple[ 2 ],
				stateString = tuple[ 3 ];

			// promise[ done | fail | progress ] = list.add
			promise[ tuple[1] ] = list.add;

			// Handle state
			if ( stateString ) {
				list.add(function() {
					// state = [ resolved | rejected ]
					state = stateString;

				// [ reject_list | resolve_list ].disable; progress_list.lock
				}, tuples[ i ^ 1 ][ 2 ].disable, tuples[ 2 ][ 2 ].lock );
			}

			// deferred[ resolve | reject | notify ]
			deferred[ tuple[0] ] = function() {
				deferred[ tuple[0] + "With" ]( this === deferred ? promise : this, arguments );
				return this;
			};
			deferred[ tuple[0] + "With" ] = list.fireWith;
		});

		// Make the deferred a promise
		promise.promise( deferred );

		// Call given func if any
		if ( func ) {
			func.call( deferred, deferred );
		}

		// All done!
		return deferred;
	},

	// Deferred helper
	when: function( subordinate /* , ..., subordinateN */ ) {
		var i = 0,
			resolveValues = slice.call( arguments ),
			length = resolveValues.length,

			// the count of uncompleted subordinates
			remaining = length !== 1 || ( subordinate && jQuery.isFunction( subordinate.promise ) ) ? length : 0,

			// the master Deferred. If resolveValues consist of only a single Deferred, just use that.
			deferred = remaining === 1 ? subordinate : jQuery.Deferred(),

			// Update function for both resolve and progress values
			updateFunc = function( i, contexts, values ) {
				return function( value ) {
					contexts[ i ] = this;
					values[ i ] = arguments.length > 1 ? slice.call( arguments ) : value;
					if ( values === progressValues ) {
						deferred.notifyWith( contexts, values );

					} else if ( !(--remaining) ) {
						deferred.resolveWith( contexts, values );
					}
				};
			},

			progressValues, progressContexts, resolveContexts;

		// add listeners to Deferred subordinates; treat others as resolved
		if ( length > 1 ) {
			progressValues = new Array( length );
			progressContexts = new Array( length );
			resolveContexts = new Array( length );
			for ( ; i < length; i++ ) {
				if ( resolveValues[ i ] && jQuery.isFunction( resolveValues[ i ].promise ) ) {
					resolveValues[ i ].promise()
						.done( updateFunc( i, resolveContexts, resolveValues ) )
						.fail( deferred.reject )
						.progress( updateFunc( i, progressContexts, progressValues ) );
				} else {
					--remaining;
				}
			}
		}

		// if we're not waiting on anything, resolve the master
		if ( !remaining ) {
			deferred.resolveWith( resolveContexts, resolveValues );
		}

		return deferred.promise();
	}
});


// The deferred used on DOM ready
var readyList;

jQuery.fn.ready = function( fn ) {
	// Add the callback
	jQuery.ready.promise().done( fn );

	return this;
};

jQuery.extend({
	// Is the DOM ready to be used? Set to true once it occurs.
	isReady: false,

	// A counter to track how many items to wait for before
	// the ready event fires. See #6781
	readyWait: 1,

	// Hold (or release) the ready event
	holdReady: function( hold ) {
		if ( hold ) {
			jQuery.readyWait++;
		} else {
			jQuery.ready( true );
		}
	},

	// Handle when the DOM is ready
	ready: function( wait ) {

		// Abort if there are pending holds or we're already ready
		if ( wait === true ? --jQuery.readyWait : jQuery.isReady ) {
			return;
		}

		// Make sure body exists, at least, in case IE gets a little overzealous (ticket #5443).
		if ( !document.body ) {
			return setTimeout( jQuery.ready );
		}

		// Remember that the DOM is ready
		jQuery.isReady = true;

		// If a normal DOM Ready event fired, decrement, and wait if need be
		if ( wait !== true && --jQuery.readyWait > 0 ) {
			return;
		}

		// If there are functions bound, to execute
		readyList.resolveWith( document, [ jQuery ] );

		// Trigger any bound ready events
		if ( jQuery.fn.triggerHandler ) {
			jQuery( document ).triggerHandler( "ready" );
			jQuery( document ).off( "ready" );
		}
	}
});

/**
 * Clean-up method for dom ready events
 */
function detach() {
	if ( document.addEventListener ) {
		document.removeEventListener( "DOMContentLoaded", completed, false );
		window.removeEventListener( "load", completed, false );

	} else {
		document.detachEvent( "onreadystatechange", completed );
		window.detachEvent( "onload", completed );
	}
}

/**
 * The ready event handler and self cleanup method
 */
function completed() {
	// readyState === "complete" is good enough for us to call the dom ready in oldIE
	if ( document.addEventListener || event.type === "load" || document.readyState === "complete" ) {
		detach();
		jQuery.ready();
	}
}

jQuery.ready.promise = function( obj ) {
	if ( !readyList ) {

		readyList = jQuery.Deferred();

		// Catch cases where $(document).ready() is called after the browser event has already occurred.
		// we once tried to use readyState "interactive" here, but it caused issues like the one
		// discovered by ChrisS here: http://bugs.jquery.com/ticket/12282#comment:15
		if ( document.readyState === "complete" ) {
			// Handle it asynchronously to allow scripts the opportunity to delay ready
			setTimeout( jQuery.ready );

		// Standards-based browsers support DOMContentLoaded
		} else if ( document.addEventListener ) {
			// Use the handy event callback
			document.addEventListener( "DOMContentLoaded", completed, false );

			// A fallback to window.onload, that will always work
			window.addEventListener( "load", completed, false );

		// If IE event model is used
		} else {
			// Ensure firing before onload, maybe late but safe also for iframes
			document.attachEvent( "onreadystatechange", completed );

			// A fallback to window.onload, that will always work
			window.attachEvent( "onload", completed );

			// If IE and not a frame
			// continually check to see if the document is ready
			var top = false;

			try {
				top = window.frameElement == null && document.documentElement;
			} catch(e) {}

			if ( top && top.doScroll ) {
				(function doScrollCheck() {
					if ( !jQuery.isReady ) {

						try {
							// Use the trick by Diego Perini
							// http://javascript.nwbox.com/IEContentLoaded/
							top.doScroll("left");
						} catch(e) {
							return setTimeout( doScrollCheck, 50 );
						}

						// detach all dom ready events
						detach();

						// and execute any waiting functions
						jQuery.ready();
					}
				})();
			}
		}
	}
	return readyList.promise( obj );
};


var strundefined = typeof undefined;


// Support: IE<9
// Iteration over object's inherited properties before its own
var i;
for ( i in jQuery( support ) ) {
	break;
}
support.ownLast = i !== "0";

// Note: most support tests are defined in their respective modules.
// false until the test is run
support.inlineBlockNeedsLayout = false;

// Execute ASAP in case we need to set body.style.zoom
jQuery(function() {
	// Minified: var a,b,c,d
	var val, div, body, container;

	body = document.getElementsByTagName( "body" )[ 0 ];
	if ( !body || !body.style ) {
		// Return for frameset docs that don't have a body
		return;
	}

	// Setup
	div = document.createElement( "div" );
	container = document.createElement( "div" );
	container.style.cssText = "position:absolute;border:0;width:0;height:0;top:0;left:-9999px";
	body.appendChild( container ).appendChild( div );

	if ( typeof div.style.zoom !== strundefined ) {
		// Support: IE<8
		// Check if natively block-level elements act like inline-block
		// elements when setting their display to 'inline' and giving
		// them layout
		div.style.cssText = "display:inline;margin:0;border:0;padding:1px;width:1px;zoom:1";

		support.inlineBlockNeedsLayout = val = div.offsetWidth === 3;
		if ( val ) {
			// Prevent IE 6 from affecting layout for positioned elements #11048
			// Prevent IE from shrinking the body in IE 7 mode #12869
			// Support: IE<8
			body.style.zoom = 1;
		}
	}

	body.removeChild( container );
});


(function() {
	var div = document.createElement( "div" );

	// Execute the test only if not already executed in another module.
	if (support.deleteExpando == null) {
		// Support: IE<9
		support.deleteExpando = true;
		try {
			delete div.test;
		} catch( e ) {
			support.deleteExpando = false;
		}
	}

	// Null elements to avoid leaks in IE.
	div = null;
})();


/**
 * Determines whether an object can have data
 */
jQuery.acceptData = function( elem ) {
	var noData = jQuery.noData[ (elem.nodeName + " ").toLowerCase() ],
		nodeType = +elem.nodeType || 1;

	// Do not set data on non-element DOM nodes because it will not be cleared (#8335).
	return nodeType !== 1 && nodeType !== 9 ?
		false :

		// Nodes accept data unless otherwise specified; rejection can be conditional
		!noData || noData !== true && elem.getAttribute("classid") === noData;
};


var rbrace = /^(?:\{[\w\W]*\}|\[[\w\W]*\])$/,
	rmultiDash = /([A-Z])/g;

function dataAttr( elem, key, data ) {
	// If nothing was found internally, try to fetch any
	// data from the HTML5 data-* attribute
	if ( data === undefined && elem.nodeType === 1 ) {

		var name = "data-" + key.replace( rmultiDash, "-$1" ).toLowerCase();

		data = elem.getAttribute( name );

		if ( typeof data === "string" ) {
			try {
				data = data === "true" ? true :
					data === "false" ? false :
					data === "null" ? null :
					// Only convert to a number if it doesn't change the string
					+data + "" === data ? +data :
					rbrace.test( data ) ? jQuery.parseJSON( data ) :
					data;
			} catch( e ) {}

			// Make sure we set the data so it isn't changed later
			jQuery.data( elem, key, data );

		} else {
			data = undefined;
		}
	}

	return data;
}

// checks a cache object for emptiness
function isEmptyDataObject( obj ) {
	var name;
	for ( name in obj ) {

		// if the public data object is empty, the private is still empty
		if ( name === "data" && jQuery.isEmptyObject( obj[name] ) ) {
			continue;
		}
		if ( name !== "toJSON" ) {
			return false;
		}
	}

	return true;
}

function internalData( elem, name, data, pvt /* Internal Use Only */ ) {
	if ( !jQuery.acceptData( elem ) ) {
		return;
	}

	var ret, thisCache,
		internalKey = jQuery.expando,

		// We have to handle DOM nodes and JS objects differently because IE6-7
		// can't GC object references properly across the DOM-JS boundary
		isNode = elem.nodeType,

		// Only DOM nodes need the global jQuery cache; JS object data is
		// attached directly to the object so GC can occur automatically
		cache = isNode ? jQuery.cache : elem,

		// Only defining an ID for JS objects if its cache already exists allows
		// the code to shortcut on the same path as a DOM node with no cache
		id = isNode ? elem[ internalKey ] : elem[ internalKey ] && internalKey;

	// Avoid doing any more work than we need to when trying to get data on an
	// object that has no data at all
	if ( (!id || !cache[id] || (!pvt && !cache[id].data)) && data === undefined && typeof name === "string" ) {
		return;
	}

	if ( !id ) {
		// Only DOM nodes need a new unique ID for each element since their data
		// ends up in the global cache
		if ( isNode ) {
			id = elem[ internalKey ] = deletedIds.pop() || jQuery.guid++;
		} else {
			id = internalKey;
		}
	}

	if ( !cache[ id ] ) {
		// Avoid exposing jQuery metadata on plain JS objects when the object
		// is serialized using JSON.stringify
		cache[ id ] = isNode ? {} : { toJSON: jQuery.noop };
	}

	// An object can be passed to jQuery.data instead of a key/value pair; this gets
	// shallow copied over onto the existing cache
	if ( typeof name === "object" || typeof name === "function" ) {
		if ( pvt ) {
			cache[ id ] = jQuery.extend( cache[ id ], name );
		} else {
			cache[ id ].data = jQuery.extend( cache[ id ].data, name );
		}
	}

	thisCache = cache[ id ];

	// jQuery data() is stored in a separate object inside the object's internal data
	// cache in order to avoid key collisions between internal data and user-defined
	// data.
	if ( !pvt ) {
		if ( !thisCache.data ) {
			thisCache.data = {};
		}

		thisCache = thisCache.data;
	}

	if ( data !== undefined ) {
		thisCache[ jQuery.camelCase( name ) ] = data;
	}

	// Check for both converted-to-camel and non-converted data property names
	// If a data property was specified
	if ( typeof name === "string" ) {

		// First Try to find as-is property data
		ret = thisCache[ name ];

		// Test for null|undefined property data
		if ( ret == null ) {

			// Try to find the camelCased property
			ret = thisCache[ jQuery.camelCase( name ) ];
		}
	} else {
		ret = thisCache;
	}

	return ret;
}

function internalRemoveData( elem, name, pvt ) {
	if ( !jQuery.acceptData( elem ) ) {
		return;
	}

	var thisCache, i,
		isNode = elem.nodeType,

		// See jQuery.data for more information
		cache = isNode ? jQuery.cache : elem,
		id = isNode ? elem[ jQuery.expando ] : jQuery.expando;

	// If there is already no cache entry for this object, there is no
	// purpose in continuing
	if ( !cache[ id ] ) {
		return;
	}

	if ( name ) {

		thisCache = pvt ? cache[ id ] : cache[ id ].data;

		if ( thisCache ) {

			// Support array or space separated string names for data keys
			if ( !jQuery.isArray( name ) ) {

				// try the string as a key before any manipulation
				if ( name in thisCache ) {
					name = [ name ];
				} else {

					// split the camel cased version by spaces unless a key with the spaces exists
					name = jQuery.camelCase( name );
					if ( name in thisCache ) {
						name = [ name ];
					} else {
						name = name.split(" ");
					}
				}
			} else {
				// If "name" is an array of keys...
				// When data is initially created, via ("key", "val") signature,
				// keys will be converted to camelCase.
				// Since there is no way to tell _how_ a key was added, remove
				// both plain key and camelCase key. #12786
				// This will only penalize the array argument path.
				name = name.concat( jQuery.map( name, jQuery.camelCase ) );
			}

			i = name.length;
			while ( i-- ) {
				delete thisCache[ name[i] ];
			}

			// If there is no data left in the cache, we want to continue
			// and let the cache object itself get destroyed
			if ( pvt ? !isEmptyDataObject(thisCache) : !jQuery.isEmptyObject(thisCache) ) {
				return;
			}
		}
	}

	// See jQuery.data for more information
	if ( !pvt ) {
		delete cache[ id ].data;

		// Don't destroy the parent cache unless the internal data object
		// had been the only thing left in it
		if ( !isEmptyDataObject( cache[ id ] ) ) {
			return;
		}
	}

	// Destroy the cache
	if ( isNode ) {
		jQuery.cleanData( [ elem ], true );

	// Use delete when supported for expandos or `cache` is not a window per isWindow (#10080)
	/* jshint eqeqeq: false */
	} else if ( support.deleteExpando || cache != cache.window ) {
		/* jshint eqeqeq: true */
		delete cache[ id ];

	// When all else fails, null
	} else {
		cache[ id ] = null;
	}
}

jQuery.extend({
	cache: {},

	// The following elements (space-suffixed to avoid Object.prototype collisions)
	// throw uncatchable exceptions if you attempt to set expando properties
	noData: {
		"applet ": true,
		"embed ": true,
		// ...but Flash objects (which have this classid) *can* handle expandos
		"object ": "clsid:D27CDB6E-AE6D-11cf-96B8-444553540000"
	},

	hasData: function( elem ) {
		elem = elem.nodeType ? jQuery.cache[ elem[jQuery.expando] ] : elem[ jQuery.expando ];
		return !!elem && !isEmptyDataObject( elem );
	},

	data: function( elem, name, data ) {
		return internalData( elem, name, data );
	},

	removeData: function( elem, name ) {
		return internalRemoveData( elem, name );
	},

	// For internal use only.
	_data: function( elem, name, data ) {
		return internalData( elem, name, data, true );
	},

	_removeData: function( elem, name ) {
		return internalRemoveData( elem, name, true );
	}
});

jQuery.fn.extend({
	data: function( key, value ) {
		var i, name, data,
			elem = this[0],
			attrs = elem && elem.attributes;

		// Special expections of .data basically thwart jQuery.access,
		// so implement the relevant behavior ourselves

		// Gets all values
		if ( key === undefined ) {
			if ( this.length ) {
				data = jQuery.data( elem );

				if ( elem.nodeType === 1 && !jQuery._data( elem, "parsedAttrs" ) ) {
					i = attrs.length;
					while ( i-- ) {

						// Support: IE11+
						// The attrs elements can be null (#14894)
						if ( attrs[ i ] ) {
							name = attrs[ i ].name;
							if ( name.indexOf( "data-" ) === 0 ) {
								name = jQuery.camelCase( name.slice(5) );
								dataAttr( elem, name, data[ name ] );
							}
						}
					}
					jQuery._data( elem, "parsedAttrs", true );
				}
			}

			return data;
		}

		// Sets multiple values
		if ( typeof key === "object" ) {
			return this.each(function() {
				jQuery.data( this, key );
			});
		}

		return arguments.length > 1 ?

			// Sets one value
			this.each(function() {
				jQuery.data( this, key, value );
			}) :

			// Gets one value
			// Try to fetch any internally stored data first
			elem ? dataAttr( elem, key, jQuery.data( elem, key ) ) : undefined;
	},

	removeData: function( key ) {
		return this.each(function() {
			jQuery.removeData( this, key );
		});
	}
});


jQuery.extend({
	queue: function( elem, type, data ) {
		var queue;

		if ( elem ) {
			type = ( type || "fx" ) + "queue";
			queue = jQuery._data( elem, type );

			// Speed up dequeue by getting out quickly if this is just a lookup
			if ( data ) {
				if ( !queue || jQuery.isArray(data) ) {
					queue = jQuery._data( elem, type, jQuery.makeArray(data) );
				} else {
					queue.push( data );
				}
			}
			return queue || [];
		}
	},

	dequeue: function( elem, type ) {
		type = type || "fx";

		var queue = jQuery.queue( elem, type ),
			startLength = queue.length,
			fn = queue.shift(),
			hooks = jQuery._queueHooks( elem, type ),
			next = function() {
				jQuery.dequeue( elem, type );
			};

		// If the fx queue is dequeued, always remove the progress sentinel
		if ( fn === "inprogress" ) {
			fn = queue.shift();
			startLength--;
		}

		if ( fn ) {

			// Add a progress sentinel to prevent the fx queue from being
			// automatically dequeued
			if ( type === "fx" ) {
				queue.unshift( "inprogress" );
			}

			// clear up the last queue stop function
			delete hooks.stop;
			fn.call( elem, next, hooks );
		}

		if ( !startLength && hooks ) {
			hooks.empty.fire();
		}
	},

	// not intended for public consumption - generates a queueHooks object, or returns the current one
	_queueHooks: function( elem, type ) {
		var key = type + "queueHooks";
		return jQuery._data( elem, key ) || jQuery._data( elem, key, {
			empty: jQuery.Callbacks("once memory").add(function() {
				jQuery._removeData( elem, type + "queue" );
				jQuery._removeData( elem, key );
			})
		});
	}
});

jQuery.fn.extend({
	queue: function( type, data ) {
		var setter = 2;

		if ( typeof type !== "string" ) {
			data = type;
			type = "fx";
			setter--;
		}

		if ( arguments.length < setter ) {
			return jQuery.queue( this[0], type );
		}

		return data === undefined ?
			this :
			this.each(function() {
				var queue = jQuery.queue( this, type, data );

				// ensure a hooks for this queue
				jQuery._queueHooks( this, type );

				if ( type === "fx" && queue[0] !== "inprogress" ) {
					jQuery.dequeue( this, type );
				}
			});
	},
	dequeue: function( type ) {
		return this.each(function() {
			jQuery.dequeue( this, type );
		});
	},
	clearQueue: function( type ) {
		return this.queue( type || "fx", [] );
	},
	// Get a promise resolved when queues of a certain type
	// are emptied (fx is the type by default)
	promise: function( type, obj ) {
		var tmp,
			count = 1,
			defer = jQuery.Deferred(),
			elements = this,
			i = this.length,
			resolve = function() {
				if ( !( --count ) ) {
					defer.resolveWith( elements, [ elements ] );
				}
			};

		if ( typeof type !== "string" ) {
			obj = type;
			type = undefined;
		}
		type = type || "fx";

		while ( i-- ) {
			tmp = jQuery._data( elements[ i ], type + "queueHooks" );
			if ( tmp && tmp.empty ) {
				count++;
				tmp.empty.add( resolve );
			}
		}
		resolve();
		return defer.promise( obj );
	}
});
var pnum = (/[+-]?(?:\d*\.|)\d+(?:[eE][+-]?\d+|)/).source;

var cssExpand = [ "Top", "Right", "Bottom", "Left" ];

var isHidden = function( elem, el ) {
		// isHidden might be called from jQuery#filter function;
		// in that case, element will be second argument
		elem = el || elem;
		return jQuery.css( elem, "display" ) === "none" || !jQuery.contains( elem.ownerDocument, elem );
	};


// Multifunctional method to get and set values of a collection
// The value/s can optionally be executed if it's a function
var access = jQuery.access = function( elems, fn, key, value, chainable, emptyGet, raw ) {
	var i = 0,
		length = elems.length,
		bulk = key == null;

	// Sets many values
	if ( jQuery.type( key ) === "object" ) {
		chainable = true;
		for ( i in key ) {
			jQuery.access( elems, fn, i, key[i], true, emptyGet, raw );
		}

	// Sets one value
	} else if ( value !== undefined ) {
		chainable = true;

		if ( !jQuery.isFunction( value ) ) {
			raw = true;
		}

		if ( bulk ) {
			// Bulk operations run against the entire set
			if ( raw ) {
				fn.call( elems, value );
				fn = null;

			// ...except when executing function values
			} else {
				bulk = fn;
				fn = function( elem, key, value ) {
					return bulk.call( jQuery( elem ), value );
				};
			}
		}

		if ( fn ) {
			for ( ; i < length; i++ ) {
				fn( elems[i], key, raw ? value : value.call( elems[i], i, fn( elems[i], key ) ) );
			}
		}
	}

	return chainable ?
		elems :

		// Gets
		bulk ?
			fn.call( elems ) :
			length ? fn( elems[0], key ) : emptyGet;
};
var rcheckableType = (/^(?:checkbox|radio)$/i);


(function() {
	// Minified: var a,b,c
	var input = document.createElement( "input" ),
		div = document.createElement( "div" ),
		fragment = document.createDocumentFragment();

	// Setup
	div.innerHTML = "  <link/><table></table><a href='/a'>a</a><input type='checkbox'/>";

	// IE strips leading whitespace when .innerHTML is used
	support.leadingWhitespace = div.firstChild.nodeType === 3;

	// Make sure that tbody elements aren't automatically inserted
	// IE will insert them into empty tables
	support.tbody = !div.getElementsByTagName( "tbody" ).length;

	// Make sure that link elements get serialized correctly by innerHTML
	// This requires a wrapper element in IE
	support.htmlSerialize = !!div.getElementsByTagName( "link" ).length;

	// Makes sure cloning an html5 element does not cause problems
	// Where outerHTML is undefined, this still works
	support.html5Clone =
		document.createElement( "nav" ).cloneNode( true ).outerHTML !== "<:nav></:nav>";

	// Check if a disconnected checkbox will retain its checked
	// value of true after appended to the DOM (IE6/7)
	input.type = "checkbox";
	input.checked = true;
	fragment.appendChild( input );
	support.appendChecked = input.checked;

	// Make sure textarea (and checkbox) defaultValue is properly cloned
	// Support: IE6-IE11+
	div.innerHTML = "<textarea>x</textarea>";
	support.noCloneChecked = !!div.cloneNode( true ).lastChild.defaultValue;

	// #11217 - WebKit loses check when the name is after the checked attribute
	fragment.appendChild( div );
	div.innerHTML = "<input type='radio' checked='checked' name='t'/>";

	// Support: Safari 5.1, iOS 5.1, Android 4.x, Android 2.3
	// old WebKit doesn't clone checked state correctly in fragments
	support.checkClone = div.cloneNode( true ).cloneNode( true ).lastChild.checked;

	// Support: IE<9
	// Opera does not clone events (and typeof div.attachEvent === undefined).
	// IE9-10 clones events bound via attachEvent, but they don't trigger with .click()
	support.noCloneEvent = true;
	if ( div.attachEvent ) {
		div.attachEvent( "onclick", function() {
			support.noCloneEvent = false;
		});

		div.cloneNode( true ).click();
	}

	// Execute the test only if not already executed in another module.
	if (support.deleteExpando == null) {
		// Support: IE<9
		support.deleteExpando = true;
		try {
			delete div.test;
		} catch( e ) {
			support.deleteExpando = false;
		}
	}
})();


(function() {
	var i, eventName,
		div = document.createElement( "div" );

	// Support: IE<9 (lack submit/change bubble), Firefox 23+ (lack focusin event)
	for ( i in { submit: true, change: true, focusin: true }) {
		eventName = "on" + i;

		if ( !(support[ i + "Bubbles" ] = eventName in window) ) {
			// Beware of CSP restrictions (https://developer.mozilla.org/en/Security/CSP)
			div.setAttribute( eventName, "t" );
			support[ i + "Bubbles" ] = div.attributes[ eventName ].expando === false;
		}
	}

	// Null elements to avoid leaks in IE.
	div = null;
})();


var rformElems = /^(?:input|select|textarea)$/i,
	rkeyEvent = /^key/,
	rmouseEvent = /^(?:mouse|pointer|contextmenu)|click/,
	rfocusMorph = /^(?:focusinfocus|focusoutblur)$/,
	rtypenamespace = /^([^.]*)(?:\.(.+)|)$/;

function returnTrue() {
	return true;
}

function returnFalse() {
	return false;
}

function safeActiveElement() {
	try {
		return document.activeElement;
	} catch ( err ) { }
}

/*
 * Helper functions for managing events -- not part of the public interface.
 * Props to Dean Edwards' addEvent library for many of the ideas.
 */
jQuery.event = {

	global: {},

	add: function( elem, types, handler, data, selector ) {
		var tmp, events, t, handleObjIn,
			special, eventHandle, handleObj,
			handlers, type, namespaces, origType,
			elemData = jQuery._data( elem );

		// Don't attach events to noData or text/comment nodes (but allow plain objects)
		if ( !elemData ) {
			return;
		}

		// Caller can pass in an object of custom data in lieu of the handler
		if ( handler.handler ) {
			handleObjIn = handler;
			handler = handleObjIn.handler;
			selector = handleObjIn.selector;
		}

		// Make sure that the handler has a unique ID, used to find/remove it later
		if ( !handler.guid ) {
			handler.guid = jQuery.guid++;
		}

		// Init the element's event structure and main handler, if this is the first
		if ( !(events = elemData.events) ) {
			events = elemData.events = {};
		}
		if ( !(eventHandle = elemData.handle) ) {
			eventHandle = elemData.handle = function( e ) {
				// Discard the second event of a jQuery.event.trigger() and
				// when an event is called after a page has unloaded
				return typeof jQuery !== strundefined && (!e || jQuery.event.triggered !== e.type) ?
					jQuery.event.dispatch.apply( eventHandle.elem, arguments ) :
					undefined;
			};
			// Add elem as a property of the handle fn to prevent a memory leak with IE non-native events
			eventHandle.elem = elem;
		}

		// Handle multiple events separated by a space
		types = ( types || "" ).match( rnotwhite ) || [ "" ];
		t = types.length;
		while ( t-- ) {
			tmp = rtypenamespace.exec( types[t] ) || [];
			type = origType = tmp[1];
			namespaces = ( tmp[2] || "" ).split( "." ).sort();

			// There *must* be a type, no attaching namespace-only handlers
			if ( !type ) {
				continue;
			}

			// If event changes its type, use the special event handlers for the changed type
			special = jQuery.event.special[ type ] || {};

			// If selector defined, determine special event api type, otherwise given type
			type = ( selector ? special.delegateType : special.bindType ) || type;

			// Update special based on newly reset type
			special = jQuery.event.special[ type ] || {};

			// handleObj is passed to all event handlers
			handleObj = jQuery.extend({
				type: type,
				origType: origType,
				data: data,
				handler: handler,
				guid: handler.guid,
				selector: selector,
				needsContext: selector && jQuery.expr.match.needsContext.test( selector ),
				namespace: namespaces.join(".")
			}, handleObjIn );

			// Init the event handler queue if we're the first
			if ( !(handlers = events[ type ]) ) {
				handlers = events[ type ] = [];
				handlers.delegateCount = 0;

				// Only use addEventListener/attachEvent if the special events handler returns false
				if ( !special.setup || special.setup.call( elem, data, namespaces, eventHandle ) === false ) {
					// Bind the global event handler to the element
					if ( elem.addEventListener ) {
						elem.addEventListener( type, eventHandle, false );

					} else if ( elem.attachEvent ) {
						elem.attachEvent( "on" + type, eventHandle );
					}
				}
			}

			if ( special.add ) {
				special.add.call( elem, handleObj );

				if ( !handleObj.handler.guid ) {
					handleObj.handler.guid = handler.guid;
				}
			}

			// Add to the element's handler list, delegates in front
			if ( selector ) {
				handlers.splice( handlers.delegateCount++, 0, handleObj );
			} else {
				handlers.push( handleObj );
			}

			// Keep track of which events have ever been used, for event optimization
			jQuery.event.global[ type ] = true;
		}

		// Nullify elem to prevent memory leaks in IE
		elem = null;
	},

	// Detach an event or set of events from an element
	remove: function( elem, types, handler, selector, mappedTypes ) {
		var j, handleObj, tmp,
			origCount, t, events,
			special, handlers, type,
			namespaces, origType,
			elemData = jQuery.hasData( elem ) && jQuery._data( elem );

		if ( !elemData || !(events = elemData.events) ) {
			return;
		}

		// Once for each type.namespace in types; type may be omitted
		types = ( types || "" ).match( rnotwhite ) || [ "" ];
		t = types.length;
		while ( t-- ) {
			tmp = rtypenamespace.exec( types[t] ) || [];
			type = origType = tmp[1];
			namespaces = ( tmp[2] || "" ).split( "." ).sort();

			// Unbind all events (on this namespace, if provided) for the element
			if ( !type ) {
				for ( type in events ) {
					jQuery.event.remove( elem, type + types[ t ], handler, selector, true );
				}
				continue;
			}

			special = jQuery.event.special[ type ] || {};
			type = ( selector ? special.delegateType : special.bindType ) || type;
			handlers = events[ type ] || [];
			tmp = tmp[2] && new RegExp( "(^|\\.)" + namespaces.join("\\.(?:.*\\.|)") + "(\\.|$)" );

			// Remove matching events
			origCount = j = handlers.length;
			while ( j-- ) {
				handleObj = handlers[ j ];

				if ( ( mappedTypes || origType === handleObj.origType ) &&
					( !handler || handler.guid === handleObj.guid ) &&
					( !tmp || tmp.test( handleObj.namespace ) ) &&
					( !selector || selector === handleObj.selector || selector === "**" && handleObj.selector ) ) {
					handlers.splice( j, 1 );

					if ( handleObj.selector ) {
						handlers.delegateCount--;
					}
					if ( special.remove ) {
						special.remove.call( elem, handleObj );
					}
				}
			}

			// Remove generic event handler if we removed something and no more handlers exist
			// (avoids potential for endless recursion during removal of special event handlers)
			if ( origCount && !handlers.length ) {
				if ( !special.teardown || special.teardown.call( elem, namespaces, elemData.handle ) === false ) {
					jQuery.removeEvent( elem, type, elemData.handle );
				}

				delete events[ type ];
			}
		}

		// Remove the expando if it's no longer used
		if ( jQuery.isEmptyObject( events ) ) {
			delete elemData.handle;

			// removeData also checks for emptiness and clears the expando if empty
			// so use it instead of delete
			jQuery._removeData( elem, "events" );
		}
	},

	trigger: function( event, data, elem, onlyHandlers ) {
		var handle, ontype, cur,
			bubbleType, special, tmp, i,
			eventPath = [ elem || document ],
			type = hasOwn.call( event, "type" ) ? event.type : event,
			namespaces = hasOwn.call( event, "namespace" ) ? event.namespace.split(".") : [];

		cur = tmp = elem = elem || document;

		// Don't do events on text and comment nodes
		if ( elem.nodeType === 3 || elem.nodeType === 8 ) {
			return;
		}

		// focus/blur morphs to focusin/out; ensure we're not firing them right now
		if ( rfocusMorph.test( type + jQuery.event.triggered ) ) {
			return;
		}

		if ( type.indexOf(".") >= 0 ) {
			// Namespaced trigger; create a regexp to match event type in handle()
			namespaces = type.split(".");
			type = namespaces.shift();
			namespaces.sort();
		}
		ontype = type.indexOf(":") < 0 && "on" + type;

		// Caller can pass in a jQuery.Event object, Object, or just an event type string
		event = event[ jQuery.expando ] ?
			event :
			new jQuery.Event( type, typeof event === "object" && event );

		// Trigger bitmask: & 1 for native handlers; & 2 for jQuery (always true)
		event.isTrigger = onlyHandlers ? 2 : 3;
		event.namespace = namespaces.join(".");
		event.namespace_re = event.namespace ?
			new RegExp( "(^|\\.)" + namespaces.join("\\.(?:.*\\.|)") + "(\\.|$)" ) :
			null;

		// Clean up the event in case it is being reused
		event.result = undefined;
		if ( !event.target ) {
			event.target = elem;
		}

		// Clone any incoming data and prepend the event, creating the handler arg list
		data = data == null ?
			[ event ] :
			jQuery.makeArray( data, [ event ] );

		// Allow special events to draw outside the lines
		special = jQuery.event.special[ type ] || {};
		if ( !onlyHandlers && special.trigger && special.trigger.apply( elem, data ) === false ) {
			return;
		}

		// Determine event propagation path in advance, per W3C events spec (#9951)
		// Bubble up to document, then to window; watch for a global ownerDocument var (#9724)
		if ( !onlyHandlers && !special.noBubble && !jQuery.isWindow( elem ) ) {

			bubbleType = special.delegateType || type;
			if ( !rfocusMorph.test( bubbleType + type ) ) {
				cur = cur.parentNode;
			}
			for ( ; cur; cur = cur.parentNode ) {
				eventPath.push( cur );
				tmp = cur;
			}

			// Only add window if we got to document (e.g., not plain obj or detached DOM)
			if ( tmp === (elem.ownerDocument || document) ) {
				eventPath.push( tmp.defaultView || tmp.parentWindow || window );
			}
		}

		// Fire handlers on the event path
		i = 0;
		while ( (cur = eventPath[i++]) && !event.isPropagationStopped() ) {

			event.type = i > 1 ?
				bubbleType :
				special.bindType || type;

			// jQuery handler
			handle = ( jQuery._data( cur, "events" ) || {} )[ event.type ] && jQuery._data( cur, "handle" );
			if ( handle ) {
				handle.apply( cur, data );
			}

			// Native handler
			handle = ontype && cur[ ontype ];
			if ( handle && handle.apply && jQuery.acceptData( cur ) ) {
				event.result = handle.apply( cur, data );
				if ( event.result === false ) {
					event.preventDefault();
				}
			}
		}
		event.type = type;

		// If nobody prevented the default action, do it now
		if ( !onlyHandlers && !event.isDefaultPrevented() ) {

			if ( (!special._default || special._default.apply( eventPath.pop(), data ) === false) &&
				jQuery.acceptData( elem ) ) {

				// Call a native DOM method on the target with the same name name as the event.
				// Can't use an .isFunction() check here because IE6/7 fails that test.
				// Don't do default actions on window, that's where global variables be (#6170)
				if ( ontype && elem[ type ] && !jQuery.isWindow( elem ) ) {

					// Don't re-trigger an onFOO event when we call its FOO() method
					tmp = elem[ ontype ];

					if ( tmp ) {
						elem[ ontype ] = null;
					}

					// Prevent re-triggering of the same event, since we already bubbled it above
					jQuery.event.triggered = type;
					try {
						elem[ type ]();
					} catch ( e ) {
						// IE<9 dies on focus/blur to hidden element (#1486,#12518)
						// only reproducible on winXP IE8 native, not IE9 in IE8 mode
					}
					jQuery.event.triggered = undefined;

					if ( tmp ) {
						elem[ ontype ] = tmp;
					}
				}
			}
		}

		return event.result;
	},

	dispatch: function( event ) {

		// Make a writable jQuery.Event from the native event object
		event = jQuery.event.fix( event );

		var i, ret, handleObj, matched, j,
			handlerQueue = [],
			args = slice.call( arguments ),
			handlers = ( jQuery._data( this, "events" ) || {} )[ event.type ] || [],
			special = jQuery.event.special[ event.type ] || {};

		// Use the fix-ed jQuery.Event rather than the (read-only) native event
		args[0] = event;
		event.delegateTarget = this;

		// Call the preDispatch hook for the mapped type, and let it bail if desired
		if ( special.preDispatch && special.preDispatch.call( this, event ) === false ) {
			return;
		}

		// Determine handlers
		handlerQueue = jQuery.event.handlers.call( this, event, handlers );

		// Run delegates first; they may want to stop propagation beneath us
		i = 0;
		while ( (matched = handlerQueue[ i++ ]) && !event.isPropagationStopped() ) {
			event.currentTarget = matched.elem;

			j = 0;
			while ( (handleObj = matched.handlers[ j++ ]) && !event.isImmediatePropagationStopped() ) {

				// Triggered event must either 1) have no namespace, or
				// 2) have namespace(s) a subset or equal to those in the bound event (both can have no namespace).
				if ( !event.namespace_re || event.namespace_re.test( handleObj.namespace ) ) {

					event.handleObj = handleObj;
					event.data = handleObj.data;

					ret = ( (jQuery.event.special[ handleObj.origType ] || {}).handle || handleObj.handler )
							.apply( matched.elem, args );

					if ( ret !== undefined ) {
						if ( (event.result = ret) === false ) {
							event.preventDefault();
							event.stopPropagation();
						}
					}
				}
			}
		}

		// Call the postDispatch hook for the mapped type
		if ( special.postDispatch ) {
			special.postDispatch.call( this, event );
		}

		return event.result;
	},

	handlers: function( event, handlers ) {
		var sel, handleObj, matches, i,
			handlerQueue = [],
			delegateCount = handlers.delegateCount,
			cur = event.target;

		// Find delegate handlers
		// Black-hole SVG <use> instance trees (#13180)
		// Avoid non-left-click bubbling in Firefox (#3861)
		if ( delegateCount && cur.nodeType && (!event.button || event.type !== "click") ) {

			/* jshint eqeqeq: false */
			for ( ; cur != this; cur = cur.parentNode || this ) {
				/* jshint eqeqeq: true */

				// Don't check non-elements (#13208)
				// Don't process clicks on disabled elements (#6911, #8165, #11382, #11764)
				if ( cur.nodeType === 1 && (cur.disabled !== true || event.type !== "click") ) {
					matches = [];
					for ( i = 0; i < delegateCount; i++ ) {
						handleObj = handlers[ i ];

						// Don't conflict with Object.prototype properties (#13203)
						sel = handleObj.selector + " ";

						if ( matches[ sel ] === undefined ) {
							matches[ sel ] = handleObj.needsContext ?
								jQuery( sel, this ).index( cur ) >= 0 :
								jQuery.find( sel, this, null, [ cur ] ).length;
						}
						if ( matches[ sel ] ) {
							matches.push( handleObj );
						}
					}
					if ( matches.length ) {
						handlerQueue.push({ elem: cur, handlers: matches });
					}
				}
			}
		}

		// Add the remaining (directly-bound) handlers
		if ( delegateCount < handlers.length ) {
			handlerQueue.push({ elem: this, handlers: handlers.slice( delegateCount ) });
		}

		return handlerQueue;
	},

	fix: function( event ) {
		if ( event[ jQuery.expando ] ) {
			return event;
		}

		// Create a writable copy of the event object and normalize some properties
		var i, prop, copy,
			type = event.type,
			originalEvent = event,
			fixHook = this.fixHooks[ type ];

		if ( !fixHook ) {
			this.fixHooks[ type ] = fixHook =
				rmouseEvent.test( type ) ? this.mouseHooks :
				rkeyEvent.test( type ) ? this.keyHooks :
				{};
		}
		copy = fixHook.props ? this.props.concat( fixHook.props ) : this.props;

		event = new jQuery.Event( originalEvent );

		i = copy.length;
		while ( i-- ) {
			prop = copy[ i ];
			event[ prop ] = originalEvent[ prop ];
		}

		// Support: IE<9
		// Fix target property (#1925)
		if ( !event.target ) {
			event.target = originalEvent.srcElement || document;
		}

		// Support: Chrome 23+, Safari?
		// Target should not be a text node (#504, #13143)
		if ( event.target.nodeType === 3 ) {
			event.target = event.target.parentNode;
		}

		// Support: IE<9
		// For mouse/key events, metaKey==false if it's undefined (#3368, #11328)
		event.metaKey = !!event.metaKey;

		return fixHook.filter ? fixHook.filter( event, originalEvent ) : event;
	},

	// Includes some event props shared by KeyEvent and MouseEvent
	props: "altKey bubbles cancelable ctrlKey currentTarget eventPhase metaKey relatedTarget shiftKey target timeStamp view which".split(" "),

	fixHooks: {},

	keyHooks: {
		props: "char charCode key keyCode".split(" "),
		filter: function( event, original ) {

			// Add which for key events
			if ( event.which == null ) {
				event.which = original.charCode != null ? original.charCode : original.keyCode;
			}

			return event;
		}
	},

	mouseHooks: {
		props: "button buttons clientX clientY fromElement offsetX offsetY pageX pageY screenX screenY toElement".split(" "),
		filter: function( event, original ) {
			var body, eventDoc, doc,
				button = original.button,
				fromElement = original.fromElement;

			// Calculate pageX/Y if missing and clientX/Y available
			if ( event.pageX == null && original.clientX != null ) {
				eventDoc = event.target.ownerDocument || document;
				doc = eventDoc.documentElement;
				body = eventDoc.body;

				event.pageX = original.clientX + ( doc && doc.scrollLeft || body && body.scrollLeft || 0 ) - ( doc && doc.clientLeft || body && body.clientLeft || 0 );
				event.pageY = original.clientY + ( doc && doc.scrollTop  || body && body.scrollTop  || 0 ) - ( doc && doc.clientTop  || body && body.clientTop  || 0 );
			}

			// Add relatedTarget, if necessary
			if ( !event.relatedTarget && fromElement ) {
				event.relatedTarget = fromElement === event.target ? original.toElement : fromElement;
			}

			// Add which for click: 1 === left; 2 === middle; 3 === right
			// Note: button is not normalized, so don't use it
			if ( !event.which && button !== undefined ) {
				event.which = ( button & 1 ? 1 : ( button & 2 ? 3 : ( button & 4 ? 2 : 0 ) ) );
			}

			return event;
		}
	},

	special: {
		load: {
			// Prevent triggered image.load events from bubbling to window.load
			noBubble: true
		},
		focus: {
			// Fire native event if possible so blur/focus sequence is correct
			trigger: function() {
				if ( this !== safeActiveElement() && this.focus ) {
					try {
						this.focus();
						return false;
					} catch ( e ) {
						// Support: IE<9
						// If we error on focus to hidden element (#1486, #12518),
						// let .trigger() run the handlers
					}
				}
			},
			delegateType: "focusin"
		},
		blur: {
			trigger: function() {
				if ( this === safeActiveElement() && this.blur ) {
					this.blur();
					return false;
				}
			},
			delegateType: "focusout"
		},
		click: {
			// For checkbox, fire native event so checked state will be right
			trigger: function() {
				if ( jQuery.nodeName( this, "input" ) && this.type === "checkbox" && this.click ) {
					this.click();
					return false;
				}
			},

			// For cross-browser consistency, don't fire native .click() on links
			_default: function( event ) {
				return jQuery.nodeName( event.target, "a" );
			}
		},

		beforeunload: {
			postDispatch: function( event ) {

				// Support: Firefox 20+
				// Firefox doesn't alert if the returnValue field is not set.
				if ( event.result !== undefined && event.originalEvent ) {
					event.originalEvent.returnValue = event.result;
				}
			}
		}
	},

	simulate: function( type, elem, event, bubble ) {
		// Piggyback on a donor event to simulate a different one.
		// Fake originalEvent to avoid donor's stopPropagation, but if the
		// simulated event prevents default then we do the same on the donor.
		var e = jQuery.extend(
			new jQuery.Event(),
			event,
			{
				type: type,
				isSimulated: true,
				originalEvent: {}
			}
		);
		if ( bubble ) {
			jQuery.event.trigger( e, null, elem );
		} else {
			jQuery.event.dispatch.call( elem, e );
		}
		if ( e.isDefaultPrevented() ) {
			event.preventDefault();
		}
	}
};

jQuery.removeEvent = document.removeEventListener ?
	function( elem, type, handle ) {
		if ( elem.removeEventListener ) {
			elem.removeEventListener( type, handle, false );
		}
	} :
	function( elem, type, handle ) {
		var name = "on" + type;

		if ( elem.detachEvent ) {

			// #8545, #7054, preventing memory leaks for custom events in IE6-8
			// detachEvent needed property on element, by name of that event, to properly expose it to GC
			if ( typeof elem[ name ] === strundefined ) {
				elem[ name ] = null;
			}

			elem.detachEvent( name, handle );
		}
	};

jQuery.Event = function( src, props ) {
	// Allow instantiation without the 'new' keyword
	if ( !(this instanceof jQuery.Event) ) {
		return new jQuery.Event( src, props );
	}

	// Event object
	if ( src && src.type ) {
		this.originalEvent = src;
		this.type = src.type;

		// Events bubbling up the document may have been marked as prevented
		// by a handler lower down the tree; reflect the correct value.
		this.isDefaultPrevented = src.defaultPrevented ||
				src.defaultPrevented === undefined &&
				// Support: IE < 9, Android < 4.0
				src.returnValue === false ?
			returnTrue :
			returnFalse;

	// Event type
	} else {
		this.type = src;
	}

	// Put explicitly provided properties onto the event object
	if ( props ) {
		jQuery.extend( this, props );
	}

	// Create a timestamp if incoming event doesn't have one
	this.timeStamp = src && src.timeStamp || jQuery.now();

	// Mark it as fixed
	this[ jQuery.expando ] = true;
};

// jQuery.Event is based on DOM3 Events as specified by the ECMAScript Language Binding
// http://www.w3.org/TR/2003/WD-DOM-Level-3-Events-20030331/ecma-script-binding.html
jQuery.Event.prototype = {
	isDefaultPrevented: returnFalse,
	isPropagationStopped: returnFalse,
	isImmediatePropagationStopped: returnFalse,

	preventDefault: function() {
		var e = this.originalEvent;

		this.isDefaultPrevented = returnTrue;
		if ( !e ) {
			return;
		}

		// If preventDefault exists, run it on the original event
		if ( e.preventDefault ) {
			e.preventDefault();

		// Support: IE
		// Otherwise set the returnValue property of the original event to false
		} else {
			e.returnValue = false;
		}
	},
	stopPropagation: function() {
		var e = this.originalEvent;

		this.isPropagationStopped = returnTrue;
		if ( !e ) {
			return;
		}
		// If stopPropagation exists, run it on the original event
		if ( e.stopPropagation ) {
			e.stopPropagation();
		}

		// Support: IE
		// Set the cancelBubble property of the original event to true
		e.cancelBubble = true;
	},
	stopImmediatePropagation: function() {
		var e = this.originalEvent;

		this.isImmediatePropagationStopped = returnTrue;

		if ( e && e.stopImmediatePropagation ) {
			e.stopImmediatePropagation();
		}

		this.stopPropagation();
	}
};

// Create mouseenter/leave events using mouseover/out and event-time checks
jQuery.each({
	mouseenter: "mouseover",
	mouseleave: "mouseout",
	pointerenter: "pointerover",
	pointerleave: "pointerout"
}, function( orig, fix ) {
	jQuery.event.special[ orig ] = {
		delegateType: fix,
		bindType: fix,

		handle: function( event ) {
			var ret,
				target = this,
				related = event.relatedTarget,
				handleObj = event.handleObj;

			// For mousenter/leave call the handler if related is outside the target.
			// NB: No relatedTarget if the mouse left/entered the browser window
			if ( !related || (related !== target && !jQuery.contains( target, related )) ) {
				event.type = handleObj.origType;
				ret = handleObj.handler.apply( this, arguments );
				event.type = fix;
			}
			return ret;
		}
	};
});

// IE submit delegation
if ( !support.submitBubbles ) {

	jQuery.event.special.submit = {
		setup: function() {
			// Only need this for delegated form submit events
			if ( jQuery.nodeName( this, "form" ) ) {
				return false;
			}

			// Lazy-add a submit handler when a descendant form may potentially be submitted
			jQuery.event.add( this, "click._submit keypress._submit", function( e ) {
				// Node name check avoids a VML-related crash in IE (#9807)
				var elem = e.target,
					form = jQuery.nodeName( elem, "input" ) || jQuery.nodeName( elem, "button" ) ? elem.form : undefined;
				if ( form && !jQuery._data( form, "submitBubbles" ) ) {
					jQuery.event.add( form, "submit._submit", function( event ) {
						event._submit_bubble = true;
					});
					jQuery._data( form, "submitBubbles", true );
				}
			});
			// return undefined since we don't need an event listener
		},

		postDispatch: function( event ) {
			// If form was submitted by the user, bubble the event up the tree
			if ( event._submit_bubble ) {
				delete event._submit_bubble;
				if ( this.parentNode && !event.isTrigger ) {
					jQuery.event.simulate( "submit", this.parentNode, event, true );
				}
			}
		},

		teardown: function() {
			// Only need this for delegated form submit events
			if ( jQuery.nodeName( this, "form" ) ) {
				return false;
			}

			// Remove delegated handlers; cleanData eventually reaps submit handlers attached above
			jQuery.event.remove( this, "._submit" );
		}
	};
}

// IE change delegation and checkbox/radio fix
if ( !support.changeBubbles ) {

	jQuery.event.special.change = {

		setup: function() {

			if ( rformElems.test( this.nodeName ) ) {
				// IE doesn't fire change on a check/radio until blur; trigger it on click
				// after a propertychange. Eat the blur-change in special.change.handle.
				// This still fires onchange a second time for check/radio after blur.
				if ( this.type === "checkbox" || this.type === "radio" ) {
					jQuery.event.add( this, "propertychange._change", function( event ) {
						if ( event.originalEvent.propertyName === "checked" ) {
							this._just_changed = true;
						}
					});
					jQuery.event.add( this, "click._change", function( event ) {
						if ( this._just_changed && !event.isTrigger ) {
							this._just_changed = false;
						}
						// Allow triggered, simulated change events (#11500)
						jQuery.event.simulate( "change", this, event, true );
					});
				}
				return false;
			}
			// Delegated event; lazy-add a change handler on descendant inputs
			jQuery.event.add( this, "beforeactivate._change", function( e ) {
				var elem = e.target;

				if ( rformElems.test( elem.nodeName ) && !jQuery._data( elem, "changeBubbles" ) ) {
					jQuery.event.add( elem, "change._change", function( event ) {
						if ( this.parentNode && !event.isSimulated && !event.isTrigger ) {
							jQuery.event.simulate( "change", this.parentNode, event, true );
						}
					});
					jQuery._data( elem, "changeBubbles", true );
				}
			});
		},

		handle: function( event ) {
			var elem = event.target;

			// Swallow native change events from checkbox/radio, we already triggered them above
			if ( this !== elem || event.isSimulated || event.isTrigger || (elem.type !== "radio" && elem.type !== "checkbox") ) {
				return event.handleObj.handler.apply( this, arguments );
			}
		},

		teardown: function() {
			jQuery.event.remove( this, "._change" );

			return !rformElems.test( this.nodeName );
		}
	};
}

// Create "bubbling" focus and blur events
if ( !support.focusinBubbles ) {
	jQuery.each({ focus: "focusin", blur: "focusout" }, function( orig, fix ) {

		// Attach a single capturing handler on the document while someone wants focusin/focusout
		var handler = function( event ) {
				jQuery.event.simulate( fix, event.target, jQuery.event.fix( event ), true );
			};

		jQuery.event.special[ fix ] = {
			setup: function() {
				var doc = this.ownerDocument || this,
					attaches = jQuery._data( doc, fix );

				if ( !attaches ) {
					doc.addEventListener( orig, handler, true );
				}
				jQuery._data( doc, fix, ( attaches || 0 ) + 1 );
			},
			teardown: function() {
				var doc = this.ownerDocument || this,
					attaches = jQuery._data( doc, fix ) - 1;

				if ( !attaches ) {
					doc.removeEventListener( orig, handler, true );
					jQuery._removeData( doc, fix );
				} else {
					jQuery._data( doc, fix, attaches );
				}
			}
		};
	});
}

jQuery.fn.extend({

	on: function( types, selector, data, fn, /*INTERNAL*/ one ) {
		var type, origFn;

		// Types can be a map of types/handlers
		if ( typeof types === "object" ) {
			// ( types-Object, selector, data )
			if ( typeof selector !== "string" ) {
				// ( types-Object, data )
				data = data || selector;
				selector = undefined;
			}
			for ( type in types ) {
				this.on( type, selector, data, types[ type ], one );
			}
			return this;
		}

		if ( data == null && fn == null ) {
			// ( types, fn )
			fn = selector;
			data = selector = undefined;
		} else if ( fn == null ) {
			if ( typeof selector === "string" ) {
				// ( types, selector, fn )
				fn = data;
				data = undefined;
			} else {
				// ( types, data, fn )
				fn = data;
				data = selector;
				selector = undefined;
			}
		}
		if ( fn === false ) {
			fn = returnFalse;
		} else if ( !fn ) {
			return this;
		}

		if ( one === 1 ) {
			origFn = fn;
			fn = function( event ) {
				// Can use an empty set, since event contains the info
				jQuery().off( event );
				return origFn.apply( this, arguments );
			};
			// Use same guid so caller can remove using origFn
			fn.guid = origFn.guid || ( origFn.guid = jQuery.guid++ );
		}
		return this.each( function() {
			jQuery.event.add( this, types, fn, data, selector );
		});
	},
	one: function( types, selector, data, fn ) {
		return this.on( types, selector, data, fn, 1 );
	},
	off: function( types, selector, fn ) {
		var handleObj, type;
		if ( types && types.preventDefault && types.handleObj ) {
			// ( event )  dispatched jQuery.Event
			handleObj = types.handleObj;
			jQuery( types.delegateTarget ).off(
				handleObj.namespace ? handleObj.origType + "." + handleObj.namespace : handleObj.origType,
				handleObj.selector,
				handleObj.handler
			);
			return this;
		}
		if ( typeof types === "object" ) {
			// ( types-object [, selector] )
			for ( type in types ) {
				this.off( type, selector, types[ type ] );
			}
			return this;
		}
		if ( selector === false || typeof selector === "function" ) {
			// ( types [, fn] )
			fn = selector;
			selector = undefined;
		}
		if ( fn === false ) {
			fn = returnFalse;
		}
		return this.each(function() {
			jQuery.event.remove( this, types, fn, selector );
		});
	},

	trigger: function( type, data ) {
		return this.each(function() {
			jQuery.event.trigger( type, data, this );
		});
	},
	triggerHandler: function( type, data ) {
		var elem = this[0];
		if ( elem ) {
			return jQuery.event.trigger( type, data, elem, true );
		}
	}
});


function createSafeFragment( document ) {
	var list = nodeNames.split( "|" ),
		safeFrag = document.createDocumentFragment();

	if ( safeFrag.createElement ) {
		while ( list.length ) {
			safeFrag.createElement(
				list.pop()
			);
		}
	}
	return safeFrag;
}

var nodeNames = "abbr|article|aside|audio|bdi|canvas|data|datalist|details|figcaption|figure|footer|" +
		"header|hgroup|mark|meter|nav|output|progress|section|summary|time|video",
	rinlinejQuery = / jQuery\d+="(?:null|\d+)"/g,
	rnoshimcache = new RegExp("<(?:" + nodeNames + ")[\\s/>]", "i"),
	rleadingWhitespace = /^\s+/,
	rxhtmlTag = /<(?!area|br|col|embed|hr|img|input|link|meta|param)(([\w:]+)[^>]*)\/>/gi,
	rtagName = /<([\w:]+)/,
	rtbody = /<tbody/i,
	rhtml = /<|&#?\w+;/,
	rnoInnerhtml = /<(?:script|style|link)/i,
	// checked="checked" or checked
	rchecked = /checked\s*(?:[^=]|=\s*.checked.)/i,
	rscriptType = /^$|\/(?:java|ecma)script/i,
	rscriptTypeMasked = /^true\/(.*)/,
	rcleanScript = /^\s*<!(?:\[CDATA\[|--)|(?:\]\]|--)>\s*$/g,

	// We have to close these tags to support XHTML (#13200)
	wrapMap = {
		option: [ 1, "<select multiple='multiple'>", "</select>" ],
		legend: [ 1, "<fieldset>", "</fieldset>" ],
		area: [ 1, "<map>", "</map>" ],
		param: [ 1, "<object>", "</object>" ],
		thead: [ 1, "<table>", "</table>" ],
		tr: [ 2, "<table><tbody>", "</tbody></table>" ],
		col: [ 2, "<table><tbody></tbody><colgroup>", "</colgroup></table>" ],
		td: [ 3, "<table><tbody><tr>", "</tr></tbody></table>" ],

		// IE6-8 can't serialize link, script, style, or any html5 (NoScope) tags,
		// unless wrapped in a div with non-breaking characters in front of it.
		_default: support.htmlSerialize ? [ 0, "", "" ] : [ 1, "X<div>", "</div>"  ]
	},
	safeFragment = createSafeFragment( document ),
	fragmentDiv = safeFragment.appendChild( document.createElement("div") );

wrapMap.optgroup = wrapMap.option;
wrapMap.tbody = wrapMap.tfoot = wrapMap.colgroup = wrapMap.caption = wrapMap.thead;
wrapMap.th = wrapMap.td;

function getAll( context, tag ) {
	var elems, elem,
		i = 0,
		found = typeof context.getElementsByTagName !== strundefined ? context.getElementsByTagName( tag || "*" ) :
			typeof context.querySelectorAll !== strundefined ? context.querySelectorAll( tag || "*" ) :
			undefined;

	if ( !found ) {
		for ( found = [], elems = context.childNodes || context; (elem = elems[i]) != null; i++ ) {
			if ( !tag || jQuery.nodeName( elem, tag ) ) {
				found.push( elem );
			} else {
				jQuery.merge( found, getAll( elem, tag ) );
			}
		}
	}

	return tag === undefined || tag && jQuery.nodeName( context, tag ) ?
		jQuery.merge( [ context ], found ) :
		found;
}

// Used in buildFragment, fixes the defaultChecked property
function fixDefaultChecked( elem ) {
	if ( rcheckableType.test( elem.type ) ) {
		elem.defaultChecked = elem.checked;
	}
}

// Support: IE<8
// Manipulating tables requires a tbody
function manipulationTarget( elem, content ) {
	return jQuery.nodeName( elem, "table" ) &&
		jQuery.nodeName( content.nodeType !== 11 ? content : content.firstChild, "tr" ) ?

		elem.getElementsByTagName("tbody")[0] ||
			elem.appendChild( elem.ownerDocument.createElement("tbody") ) :
		elem;
}

// Replace/restore the type attribute of script elements for safe DOM manipulation
function disableScript( elem ) {
	elem.type = (jQuery.find.attr( elem, "type" ) !== null) + "/" + elem.type;
	return elem;
}
function restoreScript( elem ) {
	var match = rscriptTypeMasked.exec( elem.type );
	if ( match ) {
		elem.type = match[1];
	} else {
		elem.removeAttribute("type");
	}
	return elem;
}

// Mark scripts as having already been evaluated
function setGlobalEval( elems, refElements ) {
	var elem,
		i = 0;
	for ( ; (elem = elems[i]) != null; i++ ) {
		jQuery._data( elem, "globalEval", !refElements || jQuery._data( refElements[i], "globalEval" ) );
	}
}

function cloneCopyEvent( src, dest ) {

	if ( dest.nodeType !== 1 || !jQuery.hasData( src ) ) {
		return;
	}

	var type, i, l,
		oldData = jQuery._data( src ),
		curData = jQuery._data( dest, oldData ),
		events = oldData.events;

	if ( events ) {
		delete curData.handle;
		curData.events = {};

		for ( type in events ) {
			for ( i = 0, l = events[ type ].length; i < l; i++ ) {
				jQuery.event.add( dest, type, events[ type ][ i ] );
			}
		}
	}

	// make the cloned public data object a copy from the original
	if ( curData.data ) {
		curData.data = jQuery.extend( {}, curData.data );
	}
}

function fixCloneNodeIssues( src, dest ) {
	var nodeName, e, data;

	// We do not need to do anything for non-Elements
	if ( dest.nodeType !== 1 ) {
		return;
	}

	nodeName = dest.nodeName.toLowerCase();

	// IE6-8 copies events bound via attachEvent when using cloneNode.
	if ( !support.noCloneEvent && dest[ jQuery.expando ] ) {
		data = jQuery._data( dest );

		for ( e in data.events ) {
			jQuery.removeEvent( dest, e, data.handle );
		}

		// Event data gets referenced instead of copied if the expando gets copied too
		dest.removeAttribute( jQuery.expando );
	}

	// IE blanks contents when cloning scripts, and tries to evaluate newly-set text
	if ( nodeName === "script" && dest.text !== src.text ) {
		disableScript( dest ).text = src.text;
		restoreScript( dest );

	// IE6-10 improperly clones children of object elements using classid.
	// IE10 throws NoModificationAllowedError if parent is null, #12132.
	} else if ( nodeName === "object" ) {
		if ( dest.parentNode ) {
			dest.outerHTML = src.outerHTML;
		}

		// This path appears unavoidable for IE9. When cloning an object
		// element in IE9, the outerHTML strategy above is not sufficient.
		// If the src has innerHTML and the destination does not,
		// copy the src.innerHTML into the dest.innerHTML. #10324
		if ( support.html5Clone && ( src.innerHTML && !jQuery.trim(dest.innerHTML) ) ) {
			dest.innerHTML = src.innerHTML;
		}

	} else if ( nodeName === "input" && rcheckableType.test( src.type ) ) {
		// IE6-8 fails to persist the checked state of a cloned checkbox
		// or radio button. Worse, IE6-7 fail to give the cloned element
		// a checked appearance if the defaultChecked value isn't also set

		dest.defaultChecked = dest.checked = src.checked;

		// IE6-7 get confused and end up setting the value of a cloned
		// checkbox/radio button to an empty string instead of "on"
		if ( dest.value !== src.value ) {
			dest.value = src.value;
		}

	// IE6-8 fails to return the selected option to the default selected
	// state when cloning options
	} else if ( nodeName === "option" ) {
		dest.defaultSelected = dest.selected = src.defaultSelected;

	// IE6-8 fails to set the defaultValue to the correct value when
	// cloning other types of input fields
	} else if ( nodeName === "input" || nodeName === "textarea" ) {
		dest.defaultValue = src.defaultValue;
	}
}

jQuery.extend({
	clone: function( elem, dataAndEvents, deepDataAndEvents ) {
		var destElements, node, clone, i, srcElements,
			inPage = jQuery.contains( elem.ownerDocument, elem );

		if ( support.html5Clone || jQuery.isXMLDoc(elem) || !rnoshimcache.test( "<" + elem.nodeName + ">" ) ) {
			clone = elem.cloneNode( true );

		// IE<=8 does not properly clone detached, unknown element nodes
		} else {
			fragmentDiv.innerHTML = elem.outerHTML;
			fragmentDiv.removeChild( clone = fragmentDiv.firstChild );
		}

		if ( (!support.noCloneEvent || !support.noCloneChecked) &&
				(elem.nodeType === 1 || elem.nodeType === 11) && !jQuery.isXMLDoc(elem) ) {

			// We eschew Sizzle here for performance reasons: http://jsperf.com/getall-vs-sizzle/2
			destElements = getAll( clone );
			srcElements = getAll( elem );

			// Fix all IE cloning issues
			for ( i = 0; (node = srcElements[i]) != null; ++i ) {
				// Ensure that the destination node is not null; Fixes #9587
				if ( destElements[i] ) {
					fixCloneNodeIssues( node, destElements[i] );
				}
			}
		}

		// Copy the events from the original to the clone
		if ( dataAndEvents ) {
			if ( deepDataAndEvents ) {
				srcElements = srcElements || getAll( elem );
				destElements = destElements || getAll( clone );

				for ( i = 0; (node = srcElements[i]) != null; i++ ) {
					cloneCopyEvent( node, destElements[i] );
				}
			} else {
				cloneCopyEvent( elem, clone );
			}
		}

		// Preserve script evaluation history
		destElements = getAll( clone, "script" );
		if ( destElements.length > 0 ) {
			setGlobalEval( destElements, !inPage && getAll( elem, "script" ) );
		}

		destElements = srcElements = node = null;

		// Return the cloned set
		return clone;
	},

	buildFragment: function( elems, context, scripts, selection ) {
		var j, elem, contains,
			tmp, tag, tbody, wrap,
			l = elems.length,

			// Ensure a safe fragment
			safe = createSafeFragment( context ),

			nodes = [],
			i = 0;

		for ( ; i < l; i++ ) {
			elem = elems[ i ];

			if ( elem || elem === 0 ) {

				// Add nodes directly
				if ( jQuery.type( elem ) === "object" ) {
					jQuery.merge( nodes, elem.nodeType ? [ elem ] : elem );

				// Convert non-html into a text node
				} else if ( !rhtml.test( elem ) ) {
					nodes.push( context.createTextNode( elem ) );

				// Convert html into DOM nodes
				} else {
					tmp = tmp || safe.appendChild( context.createElement("div") );

					// Deserialize a standard representation
					tag = (rtagName.exec( elem ) || [ "", "" ])[ 1 ].toLowerCase();
					wrap = wrapMap[ tag ] || wrapMap._default;

					tmp.innerHTML = wrap[1] + elem.replace( rxhtmlTag, "<$1></$2>" ) + wrap[2];

					// Descend through wrappers to the right content
					j = wrap[0];
					while ( j-- ) {
						tmp = tmp.lastChild;
					}

					// Manually add leading whitespace removed by IE
					if ( !support.leadingWhitespace && rleadingWhitespace.test( elem ) ) {
						nodes.push( context.createTextNode( rleadingWhitespace.exec( elem )[0] ) );
					}

					// Remove IE's autoinserted <tbody> from table fragments
					if ( !support.tbody ) {

						// String was a <table>, *may* have spurious <tbody>
						elem = tag === "table" && !rtbody.test( elem ) ?
							tmp.firstChild :

							// String was a bare <thead> or <tfoot>
							wrap[1] === "<table>" && !rtbody.test( elem ) ?
								tmp :
								0;

						j = elem && elem.childNodes.length;
						while ( j-- ) {
							if ( jQuery.nodeName( (tbody = elem.childNodes[j]), "tbody" ) && !tbody.childNodes.length ) {
								elem.removeChild( tbody );
							}
						}
					}

					jQuery.merge( nodes, tmp.childNodes );

					// Fix #12392 for WebKit and IE > 9
					tmp.textContent = "";

					// Fix #12392 for oldIE
					while ( tmp.firstChild ) {
						tmp.removeChild( tmp.firstChild );
					}

					// Remember the top-level container for proper cleanup
					tmp = safe.lastChild;
				}
			}
		}

		// Fix #11356: Clear elements from fragment
		if ( tmp ) {
			safe.removeChild( tmp );
		}

		// Reset defaultChecked for any radios and checkboxes
		// about to be appended to the DOM in IE 6/7 (#8060)
		if ( !support.appendChecked ) {
			jQuery.grep( getAll( nodes, "input" ), fixDefaultChecked );
		}

		i = 0;
		while ( (elem = nodes[ i++ ]) ) {

			// #4087 - If origin and destination elements are the same, and this is
			// that element, do not do anything
			if ( selection && jQuery.inArray( elem, selection ) !== -1 ) {
				continue;
			}

			contains = jQuery.contains( elem.ownerDocument, elem );

			// Append to fragment
			tmp = getAll( safe.appendChild( elem ), "script" );

			// Preserve script evaluation history
			if ( contains ) {
				setGlobalEval( tmp );
			}

			// Capture executables
			if ( scripts ) {
				j = 0;
				while ( (elem = tmp[ j++ ]) ) {
					if ( rscriptType.test( elem.type || "" ) ) {
						scripts.push( elem );
					}
				}
			}
		}

		tmp = null;

		return safe;
	},

	cleanData: function( elems, /* internal */ acceptData ) {
		var elem, type, id, data,
			i = 0,
			internalKey = jQuery.expando,
			cache = jQuery.cache,
			deleteExpando = support.deleteExpando,
			special = jQuery.event.special;

		for ( ; (elem = elems[i]) != null; i++ ) {
			if ( acceptData || jQuery.acceptData( elem ) ) {

				id = elem[ internalKey ];
				data = id && cache[ id ];

				if ( data ) {
					if ( data.events ) {
						for ( type in data.events ) {
							if ( special[ type ] ) {
								jQuery.event.remove( elem, type );

							// This is a shortcut to avoid jQuery.event.remove's overhead
							} else {
								jQuery.removeEvent( elem, type, data.handle );
							}
						}
					}

					// Remove cache only if it was not already removed by jQuery.event.remove
					if ( cache[ id ] ) {

						delete cache[ id ];

						// IE does not allow us to delete expando properties from nodes,
						// nor does it have a removeAttribute function on Document nodes;
						// we must handle all of these cases
						if ( deleteExpando ) {
							delete elem[ internalKey ];

						} else if ( typeof elem.removeAttribute !== strundefined ) {
							elem.removeAttribute( internalKey );

						} else {
							elem[ internalKey ] = null;
						}

						deletedIds.push( id );
					}
				}
			}
		}
	}
});

jQuery.fn.extend({
	text: function( value ) {
		return access( this, function( value ) {
			return value === undefined ?
				jQuery.text( this ) :
				this.empty().append( ( this[0] && this[0].ownerDocument || document ).createTextNode( value ) );
		}, null, value, arguments.length );
	},

	append: function() {
		return this.domManip( arguments, function( elem ) {
			if ( this.nodeType === 1 || this.nodeType === 11 || this.nodeType === 9 ) {
				var target = manipulationTarget( this, elem );
				target.appendChild( elem );
			}
		});
	},

	prepend: function() {
		return this.domManip( arguments, function( elem ) {
			if ( this.nodeType === 1 || this.nodeType === 11 || this.nodeType === 9 ) {
				var target = manipulationTarget( this, elem );
				target.insertBefore( elem, target.firstChild );
			}
		});
	},

	before: function() {
		return this.domManip( arguments, function( elem ) {
			if ( this.parentNode ) {
				this.parentNode.insertBefore( elem, this );
			}
		});
	},

	after: function() {
		return this.domManip( arguments, function( elem ) {
			if ( this.parentNode ) {
				this.parentNode.insertBefore( elem, this.nextSibling );
			}
		});
	},

	remove: function( selector, keepData /* Internal Use Only */ ) {
		var elem,
			elems = selector ? jQuery.filter( selector, this ) : this,
			i = 0;

		for ( ; (elem = elems[i]) != null; i++ ) {

			if ( !keepData && elem.nodeType === 1 ) {
				jQuery.cleanData( getAll( elem ) );
			}

			if ( elem.parentNode ) {
				if ( keepData && jQuery.contains( elem.ownerDocument, elem ) ) {
					setGlobalEval( getAll( elem, "script" ) );
				}
				elem.parentNode.removeChild( elem );
			}
		}

		return this;
	},

	empty: function() {
		var elem,
			i = 0;

		for ( ; (elem = this[i]) != null; i++ ) {
			// Remove element nodes and prevent memory leaks
			if ( elem.nodeType === 1 ) {
				jQuery.cleanData( getAll( elem, false ) );
			}

			// Remove any remaining nodes
			while ( elem.firstChild ) {
				elem.removeChild( elem.firstChild );
			}

			// If this is a select, ensure that it displays empty (#12336)
			// Support: IE<9
			if ( elem.options && jQuery.nodeName( elem, "select" ) ) {
				elem.options.length = 0;
			}
		}

		return this;
	},

	clone: function( dataAndEvents, deepDataAndEvents ) {
		dataAndEvents = dataAndEvents == null ? false : dataAndEvents;
		deepDataAndEvents = deepDataAndEvents == null ? dataAndEvents : deepDataAndEvents;

		return this.map(function() {
			return jQuery.clone( this, dataAndEvents, deepDataAndEvents );
		});
	},

	html: function( value ) {
		return access( this, function( value ) {
			var elem = this[ 0 ] || {},
				i = 0,
				l = this.length;

			if ( value === undefined ) {
				return elem.nodeType === 1 ?
					elem.innerHTML.replace( rinlinejQuery, "" ) :
					undefined;
			}

			// See if we can take a shortcut and just use innerHTML
			if ( typeof value === "string" && !rnoInnerhtml.test( value ) &&
				( support.htmlSerialize || !rnoshimcache.test( value )  ) &&
				( support.leadingWhitespace || !rleadingWhitespace.test( value ) ) &&
				!wrapMap[ (rtagName.exec( value ) || [ "", "" ])[ 1 ].toLowerCase() ] ) {

				value = value.replace( rxhtmlTag, "<$1></$2>" );

				try {
					for (; i < l; i++ ) {
						// Remove element nodes and prevent memory leaks
						elem = this[i] || {};
						if ( elem.nodeType === 1 ) {
							jQuery.cleanData( getAll( elem, false ) );
							elem.innerHTML = value;
						}
					}

					elem = 0;

				// If using innerHTML throws an exception, use the fallback method
				} catch(e) {}
			}

			if ( elem ) {
				this.empty().append( value );
			}
		}, null, value, arguments.length );
	},

	replaceWith: function() {
		var arg = arguments[ 0 ];

		// Make the changes, replacing each context element with the new content
		this.domManip( arguments, function( elem ) {
			arg = this.parentNode;

			jQuery.cleanData( getAll( this ) );

			if ( arg ) {
				arg.replaceChild( elem, this );
			}
		});

		// Force removal if there was no new content (e.g., from empty arguments)
		return arg && (arg.length || arg.nodeType) ? this : this.remove();
	},

	detach: function( selector ) {
		return this.remove( selector, true );
	},

	domManip: function( args, callback ) {

		// Flatten any nested arrays
		args = concat.apply( [], args );

		var first, node, hasScripts,
			scripts, doc, fragment,
			i = 0,
			l = this.length,
			set = this,
			iNoClone = l - 1,
			value = args[0],
			isFunction = jQuery.isFunction( value );

		// We can't cloneNode fragments that contain checked, in WebKit
		if ( isFunction ||
				( l > 1 && typeof value === "string" &&
					!support.checkClone && rchecked.test( value ) ) ) {
			return this.each(function( index ) {
				var self = set.eq( index );
				if ( isFunction ) {
					args[0] = value.call( this, index, self.html() );
				}
				self.domManip( args, callback );
			});
		}

		if ( l ) {
			fragment = jQuery.buildFragment( args, this[ 0 ].ownerDocument, false, this );
			first = fragment.firstChild;

			if ( fragment.childNodes.length === 1 ) {
				fragment = first;
			}

			if ( first ) {
				scripts = jQuery.map( getAll( fragment, "script" ), disableScript );
				hasScripts = scripts.length;

				// Use the original fragment for the last item instead of the first because it can end up
				// being emptied incorrectly in certain situations (#8070).
				for ( ; i < l; i++ ) {
					node = fragment;

					if ( i !== iNoClone ) {
						node = jQuery.clone( node, true, true );

						// Keep references to cloned scripts for later restoration
						if ( hasScripts ) {
							jQuery.merge( scripts, getAll( node, "script" ) );
						}
					}

					callback.call( this[i], node, i );
				}

				if ( hasScripts ) {
					doc = scripts[ scripts.length - 1 ].ownerDocument;

					// Reenable scripts
					jQuery.map( scripts, restoreScript );

					// Evaluate executable scripts on first document insertion
					for ( i = 0; i < hasScripts; i++ ) {
						node = scripts[ i ];
						if ( rscriptType.test( node.type || "" ) &&
							!jQuery._data( node, "globalEval" ) && jQuery.contains( doc, node ) ) {

							if ( node.src ) {
								// Optional AJAX dependency, but won't run scripts if not present
								if ( jQuery._evalUrl ) {
									jQuery._evalUrl( node.src );
								}
							} else {
								jQuery.globalEval( ( node.text || node.textContent || node.innerHTML || "" ).replace( rcleanScript, "" ) );
							}
						}
					}
				}

				// Fix #11809: Avoid leaking memory
				fragment = first = null;
			}
		}

		return this;
	}
});

jQuery.each({
	appendTo: "append",
	prependTo: "prepend",
	insertBefore: "before",
	insertAfter: "after",
	replaceAll: "replaceWith"
}, function( name, original ) {
	jQuery.fn[ name ] = function( selector ) {
		var elems,
			i = 0,
			ret = [],
			insert = jQuery( selector ),
			last = insert.length - 1;

		for ( ; i <= last; i++ ) {
			elems = i === last ? this : this.clone(true);
			jQuery( insert[i] )[ original ]( elems );

			// Modern browsers can apply jQuery collections as arrays, but oldIE needs a .get()
			push.apply( ret, elems.get() );
		}

		return this.pushStack( ret );
	};
});


var iframe,
	elemdisplay = {};

/**
 * Retrieve the actual display of a element
 * @param {String} name nodeName of the element
 * @param {Object} doc Document object
 */
// Called only from within defaultDisplay
function actualDisplay( name, doc ) {
	var style,
		elem = jQuery( doc.createElement( name ) ).appendTo( doc.body ),

		// getDefaultComputedStyle might be reliably used only on attached element
		display = window.getDefaultComputedStyle && ( style = window.getDefaultComputedStyle( elem[ 0 ] ) ) ?

			// Use of this method is a temporary fix (more like optmization) until something better comes along,
			// since it was removed from specification and supported only in FF
			style.display : jQuery.css( elem[ 0 ], "display" );

	// We don't have any data stored on the element,
	// so use "detach" method as fast way to get rid of the element
	elem.detach();

	return display;
}

/**
 * Try to determine the default display value of an element
 * @param {String} nodeName
 */
function defaultDisplay( nodeName ) {
	var doc = document,
		display = elemdisplay[ nodeName ];

	if ( !display ) {
		display = actualDisplay( nodeName, doc );

		// If the simple way fails, read from inside an iframe
		if ( display === "none" || !display ) {

			// Use the already-created iframe if possible
			iframe = (iframe || jQuery( "<iframe frameborder='0' width='0' height='0'/>" )).appendTo( doc.documentElement );

			// Always write a new HTML skeleton so Webkit and Firefox don't choke on reuse
			doc = ( iframe[ 0 ].contentWindow || iframe[ 0 ].contentDocument ).document;

			// Support: IE
			doc.write();
			doc.close();

			display = actualDisplay( nodeName, doc );
			iframe.detach();
		}

		// Store the correct default display
		elemdisplay[ nodeName ] = display;
	}

	return display;
}


(function() {
	var shrinkWrapBlocksVal;

	support.shrinkWrapBlocks = function() {
		if ( shrinkWrapBlocksVal != null ) {
			return shrinkWrapBlocksVal;
		}

		// Will be changed later if needed.
		shrinkWrapBlocksVal = false;

		// Minified: var b,c,d
		var div, body, container;

		body = document.getElementsByTagName( "body" )[ 0 ];
		if ( !body || !body.style ) {
			// Test fired too early or in an unsupported environment, exit.
			return;
		}

		// Setup
		div = document.createElement( "div" );
		container = document.createElement( "div" );
		container.style.cssText = "position:absolute;border:0;width:0;height:0;top:0;left:-9999px";
		body.appendChild( container ).appendChild( div );

		// Support: IE6
		// Check if elements with layout shrink-wrap their children
		if ( typeof div.style.zoom !== strundefined ) {
			// Reset CSS: box-sizing; display; margin; border
			div.style.cssText =
				// Support: Firefox<29, Android 2.3
				// Vendor-prefix box-sizing
				"-webkit-box-sizing:content-box;-moz-box-sizing:content-box;" +
				"box-sizing:content-box;display:block;margin:0;border:0;" +
				"padding:1px;width:1px;zoom:1";
			div.appendChild( document.createElement( "div" ) ).style.width = "5px";
			shrinkWrapBlocksVal = div.offsetWidth !== 3;
		}

		body.removeChild( container );

		return shrinkWrapBlocksVal;
	};

})();
var rmargin = (/^margin/);

var rnumnonpx = new RegExp( "^(" + pnum + ")(?!px)[a-z%]+$", "i" );


var getStyles, curCSS,
	rposition = /^(top|right|bottom|left)$/;

if ( window.getComputedStyle ) {
	getStyles = function( elem ) {
		return elem.ownerDocument.defaultView.getComputedStyle( elem, null );
	};

	curCSS = function( elem, name, computed ) {
		var width, minWidth, maxWidth, ret,
			style = elem.style;

		computed = computed || getStyles( elem );

		// getPropertyValue is only needed for .css('filter') in IE9, see #12537
		ret = computed ? computed.getPropertyValue( name ) || computed[ name ] : undefined;

		if ( computed ) {

			if ( ret === "" && !jQuery.contains( elem.ownerDocument, elem ) ) {
				ret = jQuery.style( elem, name );
			}

			// A tribute to the "awesome hack by Dean Edwards"
			// Chrome < 17 and Safari 5.0 uses "computed value" instead of "used value" for margin-right
			// Safari 5.1.7 (at least) returns percentage for a larger set of values, but width seems to be reliably pixels
			// this is against the CSSOM draft spec: http://dev.w3.org/csswg/cssom/#resolved-values
			if ( rnumnonpx.test( ret ) && rmargin.test( name ) ) {

				// Remember the original values
				width = style.width;
				minWidth = style.minWidth;
				maxWidth = style.maxWidth;

				// Put in the new values to get a computed value out
				style.minWidth = style.maxWidth = style.width = ret;
				ret = computed.width;

				// Revert the changed values
				style.width = width;
				style.minWidth = minWidth;
				style.maxWidth = maxWidth;
			}
		}

		// Support: IE
		// IE returns zIndex value as an integer.
		return ret === undefined ?
			ret :
			ret + "";
	};
} else if ( document.documentElement.currentStyle ) {
	getStyles = function( elem ) {
		return elem.currentStyle;
	};

	curCSS = function( elem, name, computed ) {
		var left, rs, rsLeft, ret,
			style = elem.style;

		computed = computed || getStyles( elem );
		ret = computed ? computed[ name ] : undefined;

		// Avoid setting ret to empty string here
		// so we don't default to auto
		if ( ret == null && style && style[ name ] ) {
			ret = style[ name ];
		}

		// From the awesome hack by Dean Edwards
		// http://erik.eae.net/archives/2007/07/27/18.54.15/#comment-102291

		// If we're not dealing with a regular pixel number
		// but a number that has a weird ending, we need to convert it to pixels
		// but not position css attributes, as those are proportional to the parent element instead
		// and we can't measure the parent instead because it might trigger a "stacking dolls" problem
		if ( rnumnonpx.test( ret ) && !rposition.test( name ) ) {

			// Remember the original values
			left = style.left;
			rs = elem.runtimeStyle;
			rsLeft = rs && rs.left;

			// Put in the new values to get a computed value out
			if ( rsLeft ) {
				rs.left = elem.currentStyle.left;
			}
			style.left = name === "fontSize" ? "1em" : ret;
			ret = style.pixelLeft + "px";

			// Revert the changed values
			style.left = left;
			if ( rsLeft ) {
				rs.left = rsLeft;
			}
		}

		// Support: IE
		// IE returns zIndex value as an integer.
		return ret === undefined ?
			ret :
			ret + "" || "auto";
	};
}


function addGetHookIf( conditionFn, hookFn ) {
	// Define the hook, we'll check on the first run if it's really needed.
	return {
		get: function() {
			var condition = conditionFn();

			if ( condition == null ) {
				// The test was not ready at this point; screw the hook this time
				// but check again when needed next time.
				return;
			}

			if ( condition ) {
				// Hook not needed (or it's not possible to use it due to missing dependency),
				// remove it.
				// Since there are no other hooks for marginRight, remove the whole object.
				delete this.get;
				return;
			}

			// Hook needed; redefine it so that the support test is not executed again.

			return (this.get = hookFn).apply( this, arguments );
		}
	};
}


(function() {
	// Minified: var b,c,d,e,f,g, h,i
	var div, style, a, pixelPositionVal, boxSizingReliableVal,
		reliableHiddenOffsetsVal, reliableMarginRightVal;

	// Setup
	div = document.createElement( "div" );
	div.innerHTML = "  <link/><table></table><a href='/a'>a</a><input type='checkbox'/>";
	a = div.getElementsByTagName( "a" )[ 0 ];
	style = a && a.style;

	// Finish early in limited (non-browser) environments
	if ( !style ) {
		return;
	}

	style.cssText = "float:left;opacity:.5";

	// Support: IE<9
	// Make sure that element opacity exists (as opposed to filter)
	support.opacity = style.opacity === "0.5";

	// Verify style float existence
	// (IE uses styleFloat instead of cssFloat)
	support.cssFloat = !!style.cssFloat;

	div.style.backgroundClip = "content-box";
	div.cloneNode( true ).style.backgroundClip = "";
	support.clearCloneStyle = div.style.backgroundClip === "content-box";

	// Support: Firefox<29, Android 2.3
	// Vendor-prefix box-sizing
	support.boxSizing = style.boxSizing === "" || style.MozBoxSizing === "" ||
		style.WebkitBoxSizing === "";

	jQuery.extend(support, {
		reliableHiddenOffsets: function() {
			if ( reliableHiddenOffsetsVal == null ) {
				computeStyleTests();
			}
			return reliableHiddenOffsetsVal;
		},

		boxSizingReliable: function() {
			if ( boxSizingReliableVal == null ) {
				computeStyleTests();
			}
			return boxSizingReliableVal;
		},

		pixelPosition: function() {
			if ( pixelPositionVal == null ) {
				computeStyleTests();
			}
			return pixelPositionVal;
		},

		// Support: Android 2.3
		reliableMarginRight: function() {
			if ( reliableMarginRightVal == null ) {
				computeStyleTests();
			}
			return reliableMarginRightVal;
		}
	});

	function computeStyleTests() {
		// Minified: var b,c,d,j
		var div, body, container, contents;

		body = document.getElementsByTagName( "body" )[ 0 ];
		if ( !body || !body.style ) {
			// Test fired too early or in an unsupported environment, exit.
			return;
		}

		// Setup
		div = document.createElement( "div" );
		container = document.createElement( "div" );
		container.style.cssText = "position:absolute;border:0;width:0;height:0;top:0;left:-9999px";
		body.appendChild( container ).appendChild( div );

		div.style.cssText =
			// Support: Firefox<29, Android 2.3
			// Vendor-prefix box-sizing
			"-webkit-box-sizing:border-box;-moz-box-sizing:border-box;" +
			"box-sizing:border-box;display:block;margin-top:1%;top:1%;" +
			"border:1px;padding:1px;width:4px;position:absolute";

		// Support: IE<9
		// Assume reasonable values in the absence of getComputedStyle
		pixelPositionVal = boxSizingReliableVal = false;
		reliableMarginRightVal = true;

		// Check for getComputedStyle so that this code is not run in IE<9.
		if ( window.getComputedStyle ) {
			pixelPositionVal = ( window.getComputedStyle( div, null ) || {} ).top !== "1%";
			boxSizingReliableVal =
				( window.getComputedStyle( div, null ) || { width: "4px" } ).width === "4px";

			// Support: Android 2.3
			// Div with explicit width and no margin-right incorrectly
			// gets computed margin-right based on width of container (#3333)
			// WebKit Bug 13343 - getComputedStyle returns wrong value for margin-right
			contents = div.appendChild( document.createElement( "div" ) );

			// Reset CSS: box-sizing; display; margin; border; padding
			contents.style.cssText = div.style.cssText =
				// Support: Firefox<29, Android 2.3
				// Vendor-prefix box-sizing
				"-webkit-box-sizing:content-box;-moz-box-sizing:content-box;" +
				"box-sizing:content-box;display:block;margin:0;border:0;padding:0";
			contents.style.marginRight = contents.style.width = "0";
			div.style.width = "1px";

			reliableMarginRightVal =
				!parseFloat( ( window.getComputedStyle( contents, null ) || {} ).marginRight );
		}

		// Support: IE8
		// Check if table cells still have offsetWidth/Height when they are set
		// to display:none and there are still other visible table cells in a
		// table row; if so, offsetWidth/Height are not reliable for use when
		// determining if an element has been hidden directly using
		// display:none (it is still safe to use offsets if a parent element is
		// hidden; don safety goggles and see bug #4512 for more information).
		div.innerHTML = "<table><tr><td></td><td>t</td></tr></table>";
		contents = div.getElementsByTagName( "td" );
		contents[ 0 ].style.cssText = "margin:0;border:0;padding:0;display:none";
		reliableHiddenOffsetsVal = contents[ 0 ].offsetHeight === 0;
		if ( reliableHiddenOffsetsVal ) {
			contents[ 0 ].style.display = "";
			contents[ 1 ].style.display = "none";
			reliableHiddenOffsetsVal = contents[ 0 ].offsetHeight === 0;
		}

		body.removeChild( container );
	}

})();


// A method for quickly swapping in/out CSS properties to get correct calculations.
jQuery.swap = function( elem, options, callback, args ) {
	var ret, name,
		old = {};

	// Remember the old values, and insert the new ones
	for ( name in options ) {
		old[ name ] = elem.style[ name ];
		elem.style[ name ] = options[ name ];
	}

	ret = callback.apply( elem, args || [] );

	// Revert the old values
	for ( name in options ) {
		elem.style[ name ] = old[ name ];
	}

	return ret;
};


var
		ralpha = /alpha\([^)]*\)/i,
	ropacity = /opacity\s*=\s*([^)]*)/,

	// swappable if display is none or starts with table except "table", "table-cell", or "table-caption"
	// see here for display values: https://developer.mozilla.org/en-US/docs/CSS/display
	rdisplayswap = /^(none|table(?!-c[ea]).+)/,
	rnumsplit = new RegExp( "^(" + pnum + ")(.*)$", "i" ),
	rrelNum = new RegExp( "^([+-])=(" + pnum + ")", "i" ),

	cssShow = { position: "absolute", visibility: "hidden", display: "block" },
	cssNormalTransform = {
		letterSpacing: "0",
		fontWeight: "400"
	},

	cssPrefixes = [ "Webkit", "O", "Moz", "ms" ];


// return a css property mapped to a potentially vendor prefixed property
function vendorPropName( style, name ) {

	// shortcut for names that are not vendor prefixed
	if ( name in style ) {
		return name;
	}

	// check for vendor prefixed names
	var capName = name.charAt(0).toUpperCase() + name.slice(1),
		origName = name,
		i = cssPrefixes.length;

	while ( i-- ) {
		name = cssPrefixes[ i ] + capName;
		if ( name in style ) {
			return name;
		}
	}

	return origName;
}

function showHide( elements, show ) {
	var display, elem, hidden,
		values = [],
		index = 0,
		length = elements.length;

	for ( ; index < length; index++ ) {
		elem = elements[ index ];
		if ( !elem.style ) {
			continue;
		}

		values[ index ] = jQuery._data( elem, "olddisplay" );
		display = elem.style.display;
		if ( show ) {
			// Reset the inline display of this element to learn if it is
			// being hidden by cascaded rules or not
			if ( !values[ index ] && display === "none" ) {
				elem.style.display = "";
			}

			// Set elements which have been overridden with display: none
			// in a stylesheet to whatever the default browser style is
			// for such an element
			if ( elem.style.display === "" && isHidden( elem ) ) {
				values[ index ] = jQuery._data( elem, "olddisplay", defaultDisplay(elem.nodeName) );
			}
		} else {
			hidden = isHidden( elem );

			if ( display && display !== "none" || !hidden ) {
				jQuery._data( elem, "olddisplay", hidden ? display : jQuery.css( elem, "display" ) );
			}
		}
	}

	// Set the display of most of the elements in a second loop
	// to avoid the constant reflow
	for ( index = 0; index < length; index++ ) {
		elem = elements[ index ];
		if ( !elem.style ) {
			continue;
		}
		if ( !show || elem.style.display === "none" || elem.style.display === "" ) {
			elem.style.display = show ? values[ index ] || "" : "none";
		}
	}

	return elements;
}

function setPositiveNumber( elem, value, subtract ) {
	var matches = rnumsplit.exec( value );
	return matches ?
		// Guard against undefined "subtract", e.g., when used as in cssHooks
		Math.max( 0, matches[ 1 ] - ( subtract || 0 ) ) + ( matches[ 2 ] || "px" ) :
		value;
}

function augmentWidthOrHeight( elem, name, extra, isBorderBox, styles ) {
	var i = extra === ( isBorderBox ? "border" : "content" ) ?
		// If we already have the right measurement, avoid augmentation
		4 :
		// Otherwise initialize for horizontal or vertical properties
		name === "width" ? 1 : 0,

		val = 0;

	for ( ; i < 4; i += 2 ) {
		// both box models exclude margin, so add it if we want it
		if ( extra === "margin" ) {
			val += jQuery.css( elem, extra + cssExpand[ i ], true, styles );
		}

		if ( isBorderBox ) {
			// border-box includes padding, so remove it if we want content
			if ( extra === "content" ) {
				val -= jQuery.css( elem, "padding" + cssExpand[ i ], true, styles );
			}

			// at this point, extra isn't border nor margin, so remove border
			if ( extra !== "margin" ) {
				val -= jQuery.css( elem, "border" + cssExpand[ i ] + "Width", true, styles );
			}
		} else {
			// at this point, extra isn't content, so add padding
			val += jQuery.css( elem, "padding" + cssExpand[ i ], true, styles );

			// at this point, extra isn't content nor padding, so add border
			if ( extra !== "padding" ) {
				val += jQuery.css( elem, "border" + cssExpand[ i ] + "Width", true, styles );
			}
		}
	}

	return val;
}

function getWidthOrHeight( elem, name, extra ) {

	// Start with offset property, which is equivalent to the border-box value
	var valueIsBorderBox = true,
		val = name === "width" ? elem.offsetWidth : elem.offsetHeight,
		styles = getStyles( elem ),
		isBorderBox = support.boxSizing && jQuery.css( elem, "boxSizing", false, styles ) === "border-box";

	// some non-html elements return undefined for offsetWidth, so check for null/undefined
	// svg - https://bugzilla.mozilla.org/show_bug.cgi?id=649285
	// MathML - https://bugzilla.mozilla.org/show_bug.cgi?id=491668
	if ( val <= 0 || val == null ) {
		// Fall back to computed then uncomputed css if necessary
		val = curCSS( elem, name, styles );
		if ( val < 0 || val == null ) {
			val = elem.style[ name ];
		}

		// Computed unit is not pixels. Stop here and return.
		if ( rnumnonpx.test(val) ) {
			return val;
		}

		// we need the check for style in case a browser which returns unreliable values
		// for getComputedStyle silently falls back to the reliable elem.style
		valueIsBorderBox = isBorderBox && ( support.boxSizingReliable() || val === elem.style[ name ] );

		// Normalize "", auto, and prepare for extra
		val = parseFloat( val ) || 0;
	}

	// use the active box-sizing model to add/subtract irrelevant styles
	return ( val +
		augmentWidthOrHeight(
			elem,
			name,
			extra || ( isBorderBox ? "border" : "content" ),
			valueIsBorderBox,
			styles
		)
	) + "px";
}

jQuery.extend({
	// Add in style property hooks for overriding the default
	// behavior of getting and setting a style property
	cssHooks: {
		opacity: {
			get: function( elem, computed ) {
				if ( computed ) {
					// We should always get a number back from opacity
					var ret = curCSS( elem, "opacity" );
					return ret === "" ? "1" : ret;
				}
			}
		}
	},

	// Don't automatically add "px" to these possibly-unitless properties
	cssNumber: {
		"columnCount": true,
		"fillOpacity": true,
		"flexGrow": true,
		"flexShrink": true,
		"fontWeight": true,
		"lineHeight": true,
		"opacity": true,
		"order": true,
		"orphans": true,
		"widows": true,
		"zIndex": true,
		"zoom": true
	},

	// Add in properties whose names you wish to fix before
	// setting or getting the value
	cssProps: {
		// normalize float css property
		"float": support.cssFloat ? "cssFloat" : "styleFloat"
	},

	// Get and set the style property on a DOM Node
	style: function( elem, name, value, extra ) {
		// Don't set styles on text and comment nodes
		if ( !elem || elem.nodeType === 3 || elem.nodeType === 8 || !elem.style ) {
			return;
		}

		// Make sure that we're working with the right name
		var ret, type, hooks,
			origName = jQuery.camelCase( name ),
			style = elem.style;

		name = jQuery.cssProps[ origName ] || ( jQuery.cssProps[ origName ] = vendorPropName( style, origName ) );

		// gets hook for the prefixed version
		// followed by the unprefixed version
		hooks = jQuery.cssHooks[ name ] || jQuery.cssHooks[ origName ];

		// Check if we're setting a value
		if ( value !== undefined ) {
			type = typeof value;

			// convert relative number strings (+= or -=) to relative numbers. #7345
			if ( type === "string" && (ret = rrelNum.exec( value )) ) {
				value = ( ret[1] + 1 ) * ret[2] + parseFloat( jQuery.css( elem, name ) );
				// Fixes bug #9237
				type = "number";
			}

			// Make sure that null and NaN values aren't set. See: #7116
			if ( value == null || value !== value ) {
				return;
			}

			// If a number was passed in, add 'px' to the (except for certain CSS properties)
			if ( type === "number" && !jQuery.cssNumber[ origName ] ) {
				value += "px";
			}

			// Fixes #8908, it can be done more correctly by specifing setters in cssHooks,
			// but it would mean to define eight (for every problematic property) identical functions
			if ( !support.clearCloneStyle && value === "" && name.indexOf("background") === 0 ) {
				style[ name ] = "inherit";
			}

			// If a hook was provided, use that value, otherwise just set the specified value
			if ( !hooks || !("set" in hooks) || (value = hooks.set( elem, value, extra )) !== undefined ) {

				// Support: IE
				// Swallow errors from 'invalid' CSS values (#5509)
				try {
					style[ name ] = value;
				} catch(e) {}
			}

		} else {
			// If a hook was provided get the non-computed value from there
			if ( hooks && "get" in hooks && (ret = hooks.get( elem, false, extra )) !== undefined ) {
				return ret;
			}

			// Otherwise just get the value from the style object
			return style[ name ];
		}
	},

	css: function( elem, name, extra, styles ) {
		var num, val, hooks,
			origName = jQuery.camelCase( name );

		// Make sure that we're working with the right name
		name = jQuery.cssProps[ origName ] || ( jQuery.cssProps[ origName ] = vendorPropName( elem.style, origName ) );

		// gets hook for the prefixed version
		// followed by the unprefixed version
		hooks = jQuery.cssHooks[ name ] || jQuery.cssHooks[ origName ];

		// If a hook was provided get the computed value from there
		if ( hooks && "get" in hooks ) {
			val = hooks.get( elem, true, extra );
		}

		// Otherwise, if a way to get the computed value exists, use that
		if ( val === undefined ) {
			val = curCSS( elem, name, styles );
		}

		//convert "normal" to computed value
		if ( val === "normal" && name in cssNormalTransform ) {
			val = cssNormalTransform[ name ];
		}

		// Return, converting to number if forced or a qualifier was provided and val looks numeric
		if ( extra === "" || extra ) {
			num = parseFloat( val );
			return extra === true || jQuery.isNumeric( num ) ? num || 0 : val;
		}
		return val;
	}
});

jQuery.each([ "height", "width" ], function( i, name ) {
	jQuery.cssHooks[ name ] = {
		get: function( elem, computed, extra ) {
			if ( computed ) {
				// certain elements can have dimension info if we invisibly show them
				// however, it must have a current display style that would benefit from this
				return rdisplayswap.test( jQuery.css( elem, "display" ) ) && elem.offsetWidth === 0 ?
					jQuery.swap( elem, cssShow, function() {
						return getWidthOrHeight( elem, name, extra );
					}) :
					getWidthOrHeight( elem, name, extra );
			}
		},

		set: function( elem, value, extra ) {
			var styles = extra && getStyles( elem );
			return setPositiveNumber( elem, value, extra ?
				augmentWidthOrHeight(
					elem,
					name,
					extra,
					support.boxSizing && jQuery.css( elem, "boxSizing", false, styles ) === "border-box",
					styles
				) : 0
			);
		}
	};
});

if ( !support.opacity ) {
	jQuery.cssHooks.opacity = {
		get: function( elem, computed ) {
			// IE uses filters for opacity
			return ropacity.test( (computed && elem.currentStyle ? elem.currentStyle.filter : elem.style.filter) || "" ) ?
				( 0.01 * parseFloat( RegExp.$1 ) ) + "" :
				computed ? "1" : "";
		},

		set: function( elem, value ) {
			var style = elem.style,
				currentStyle = elem.currentStyle,
				opacity = jQuery.isNumeric( value ) ? "alpha(opacity=" + value * 100 + ")" : "",
				filter = currentStyle && currentStyle.filter || style.filter || "";

			// IE has trouble with opacity if it does not have layout
			// Force it by setting the zoom level
			style.zoom = 1;

			// if setting opacity to 1, and no other filters exist - attempt to remove filter attribute #6652
			// if value === "", then remove inline opacity #12685
			if ( ( value >= 1 || value === "" ) &&
					jQuery.trim( filter.replace( ralpha, "" ) ) === "" &&
					style.removeAttribute ) {

				// Setting style.filter to null, "" & " " still leave "filter:" in the cssText
				// if "filter:" is present at all, clearType is disabled, we want to avoid this
				// style.removeAttribute is IE Only, but so apparently is this code path...
				style.removeAttribute( "filter" );

				// if there is no filter style applied in a css rule or unset inline opacity, we are done
				if ( value === "" || currentStyle && !currentStyle.filter ) {
					return;
				}
			}

			// otherwise, set new filter values
			style.filter = ralpha.test( filter ) ?
				filter.replace( ralpha, opacity ) :
				filter + " " + opacity;
		}
	};
}

jQuery.cssHooks.marginRight = addGetHookIf( support.reliableMarginRight,
	function( elem, computed ) {
		if ( computed ) {
			// WebKit Bug 13343 - getComputedStyle returns wrong value for margin-right
			// Work around by temporarily setting element display to inline-block
			return jQuery.swap( elem, { "display": "inline-block" },
				curCSS, [ elem, "marginRight" ] );
		}
	}
);

// These hooks are used by animate to expand properties
jQuery.each({
	margin: "",
	padding: "",
	border: "Width"
}, function( prefix, suffix ) {
	jQuery.cssHooks[ prefix + suffix ] = {
		expand: function( value ) {
			var i = 0,
				expanded = {},

				// assumes a single number if not a string
				parts = typeof value === "string" ? value.split(" ") : [ value ];

			for ( ; i < 4; i++ ) {
				expanded[ prefix + cssExpand[ i ] + suffix ] =
					parts[ i ] || parts[ i - 2 ] || parts[ 0 ];
			}

			return expanded;
		}
	};

	if ( !rmargin.test( prefix ) ) {
		jQuery.cssHooks[ prefix + suffix ].set = setPositiveNumber;
	}
});

jQuery.fn.extend({
	css: function( name, value ) {
		return access( this, function( elem, name, value ) {
			var styles, len,
				map = {},
				i = 0;

			if ( jQuery.isArray( name ) ) {
				styles = getStyles( elem );
				len = name.length;

				for ( ; i < len; i++ ) {
					map[ name[ i ] ] = jQuery.css( elem, name[ i ], false, styles );
				}

				return map;
			}

			return value !== undefined ?
				jQuery.style( elem, name, value ) :
				jQuery.css( elem, name );
		}, name, value, arguments.length > 1 );
	},
	show: function() {
		return showHide( this, true );
	},
	hide: function() {
		return showHide( this );
	},
	toggle: function( state ) {
		if ( typeof state === "boolean" ) {
			return state ? this.show() : this.hide();
		}

		return this.each(function() {
			if ( isHidden( this ) ) {
				jQuery( this ).show();
			} else {
				jQuery( this ).hide();
			}
		});
	}
});


function Tween( elem, options, prop, end, easing ) {
	return new Tween.prototype.init( elem, options, prop, end, easing );
}
jQuery.Tween = Tween;

Tween.prototype = {
	constructor: Tween,
	init: function( elem, options, prop, end, easing, unit ) {
		this.elem = elem;
		this.prop = prop;
		this.easing = easing || "swing";
		this.options = options;
		this.start = this.now = this.cur();
		this.end = end;
		this.unit = unit || ( jQuery.cssNumber[ prop ] ? "" : "px" );
	},
	cur: function() {
		var hooks = Tween.propHooks[ this.prop ];

		return hooks && hooks.get ?
			hooks.get( this ) :
			Tween.propHooks._default.get( this );
	},
	run: function( percent ) {
		var eased,
			hooks = Tween.propHooks[ this.prop ];

		if ( this.options.duration ) {
			this.pos = eased = jQuery.easing[ this.easing ](
				percent, this.options.duration * percent, 0, 1, this.options.duration
			);
		} else {
			this.pos = eased = percent;
		}
		this.now = ( this.end - this.start ) * eased + this.start;

		if ( this.options.step ) {
			this.options.step.call( this.elem, this.now, this );
		}

		if ( hooks && hooks.set ) {
			hooks.set( this );
		} else {
			Tween.propHooks._default.set( this );
		}
		return this;
	}
};

Tween.prototype.init.prototype = Tween.prototype;

Tween.propHooks = {
	_default: {
		get: function( tween ) {
			var result;

			if ( tween.elem[ tween.prop ] != null &&
				(!tween.elem.style || tween.elem.style[ tween.prop ] == null) ) {
				return tween.elem[ tween.prop ];
			}

			// passing an empty string as a 3rd parameter to .css will automatically
			// attempt a parseFloat and fallback to a string if the parse fails
			// so, simple values such as "10px" are parsed to Float.
			// complex values such as "rotate(1rad)" are returned as is.
			result = jQuery.css( tween.elem, tween.prop, "" );
			// Empty strings, null, undefined and "auto" are converted to 0.
			return !result || result === "auto" ? 0 : result;
		},
		set: function( tween ) {
			// use step hook for back compat - use cssHook if its there - use .style if its
			// available and use plain properties where available
			if ( jQuery.fx.step[ tween.prop ] ) {
				jQuery.fx.step[ tween.prop ]( tween );
			} else if ( tween.elem.style && ( tween.elem.style[ jQuery.cssProps[ tween.prop ] ] != null || jQuery.cssHooks[ tween.prop ] ) ) {
				jQuery.style( tween.elem, tween.prop, tween.now + tween.unit );
			} else {
				tween.elem[ tween.prop ] = tween.now;
			}
		}
	}
};

// Support: IE <=9
// Panic based approach to setting things on disconnected nodes

Tween.propHooks.scrollTop = Tween.propHooks.scrollLeft = {
	set: function( tween ) {
		if ( tween.elem.nodeType && tween.elem.parentNode ) {
			tween.elem[ tween.prop ] = tween.now;
		}
	}
};

jQuery.easing = {
	linear: function( p ) {
		return p;
	},
	swing: function( p ) {
		return 0.5 - Math.cos( p * Math.PI ) / 2;
	}
};

jQuery.fx = Tween.prototype.init;

// Back Compat <1.8 extension point
jQuery.fx.step = {};


var
	fxNow, timerId,
	rfxtypes = /^(?:toggle|show|hide)$/,
	rfxnum = new RegExp( "^(?:([+-])=|)(" + pnum + ")([a-z%]*)$", "i" ),
	rrun = /queueHooks$/,
	animationPrefilters = [ defaultPrefilter ],
	tweeners = {
		"*": [ function( prop, value ) {
			var tween = this.createTween( prop, value ),
				target = tween.cur(),
				parts = rfxnum.exec( value ),
				unit = parts && parts[ 3 ] || ( jQuery.cssNumber[ prop ] ? "" : "px" ),

				// Starting value computation is required for potential unit mismatches
				start = ( jQuery.cssNumber[ prop ] || unit !== "px" && +target ) &&
					rfxnum.exec( jQuery.css( tween.elem, prop ) ),
				scale = 1,
				maxIterations = 20;

			if ( start && start[ 3 ] !== unit ) {
				// Trust units reported by jQuery.css
				unit = unit || start[ 3 ];

				// Make sure we update the tween properties later on
				parts = parts || [];

				// Iteratively approximate from a nonzero starting point
				start = +target || 1;

				do {
					// If previous iteration zeroed out, double until we get *something*
					// Use a string for doubling factor so we don't accidentally see scale as unchanged below
					scale = scale || ".5";

					// Adjust and apply
					start = start / scale;
					jQuery.style( tween.elem, prop, start + unit );

				// Update scale, tolerating zero or NaN from tween.cur()
				// And breaking the loop if scale is unchanged or perfect, or if we've just had enough
				} while ( scale !== (scale = tween.cur() / target) && scale !== 1 && --maxIterations );
			}

			// Update tween properties
			if ( parts ) {
				start = tween.start = +start || +target || 0;
				tween.unit = unit;
				// If a +=/-= token was provided, we're doing a relative animation
				tween.end = parts[ 1 ] ?
					start + ( parts[ 1 ] + 1 ) * parts[ 2 ] :
					+parts[ 2 ];
			}

			return tween;
		} ]
	};

// Animations created synchronously will run synchronously
function createFxNow() {
	setTimeout(function() {
		fxNow = undefined;
	});
	return ( fxNow = jQuery.now() );
}

// Generate parameters to create a standard animation
function genFx( type, includeWidth ) {
	var which,
		attrs = { height: type },
		i = 0;

	// if we include width, step value is 1 to do all cssExpand values,
	// if we don't include width, step value is 2 to skip over Left and Right
	includeWidth = includeWidth ? 1 : 0;
	for ( ; i < 4 ; i += 2 - includeWidth ) {
		which = cssExpand[ i ];
		attrs[ "margin" + which ] = attrs[ "padding" + which ] = type;
	}

	if ( includeWidth ) {
		attrs.opacity = attrs.width = type;
	}

	return attrs;
}

function createTween( value, prop, animation ) {
	var tween,
		collection = ( tweeners[ prop ] || [] ).concat( tweeners[ "*" ] ),
		index = 0,
		length = collection.length;
	for ( ; index < length; index++ ) {
		if ( (tween = collection[ index ].call( animation, prop, value )) ) {

			// we're done with this property
			return tween;
		}
	}
}

function defaultPrefilter( elem, props, opts ) {
	/* jshint validthis: true */
	var prop, value, toggle, tween, hooks, oldfire, display, checkDisplay,
		anim = this,
		orig = {},
		style = elem.style,
		hidden = elem.nodeType && isHidden( elem ),
		dataShow = jQuery._data( elem, "fxshow" );

	// handle queue: false promises
	if ( !opts.queue ) {
		hooks = jQuery._queueHooks( elem, "fx" );
		if ( hooks.unqueued == null ) {
			hooks.unqueued = 0;
			oldfire = hooks.empty.fire;
			hooks.empty.fire = function() {
				if ( !hooks.unqueued ) {
					oldfire();
				}
			};
		}
		hooks.unqueued++;

		anim.always(function() {
			// doing this makes sure that the complete handler will be called
			// before this completes
			anim.always(function() {
				hooks.unqueued--;
				if ( !jQuery.queue( elem, "fx" ).length ) {
					hooks.empty.fire();
				}
			});
		});
	}

	// height/width overflow pass
	if ( elem.nodeType === 1 && ( "height" in props || "width" in props ) ) {
		// Make sure that nothing sneaks out
		// Record all 3 overflow attributes because IE does not
		// change the overflow attribute when overflowX and
		// overflowY are set to the same value
		opts.overflow = [ style.overflow, style.overflowX, style.overflowY ];

		// Set display property to inline-block for height/width
		// animations on inline elements that are having width/height animated
		display = jQuery.css( elem, "display" );

		// Test default display if display is currently "none"
		checkDisplay = display === "none" ?
			jQuery._data( elem, "olddisplay" ) || defaultDisplay( elem.nodeName ) : display;

		if ( checkDisplay === "inline" && jQuery.css( elem, "float" ) === "none" ) {

			// inline-level elements accept inline-block;
			// block-level elements need to be inline with layout
			if ( !support.inlineBlockNeedsLayout || defaultDisplay( elem.nodeName ) === "inline" ) {
				style.display = "inline-block";
			} else {
				style.zoom = 1;
			}
		}
	}

	if ( opts.overflow ) {
		style.overflow = "hidden";
		if ( !support.shrinkWrapBlocks() ) {
			anim.always(function() {
				style.overflow = opts.overflow[ 0 ];
				style.overflowX = opts.overflow[ 1 ];
				style.overflowY = opts.overflow[ 2 ];
			});
		}
	}

	// show/hide pass
	for ( prop in props ) {
		value = props[ prop ];
		if ( rfxtypes.exec( value ) ) {
			delete props[ prop ];
			toggle = toggle || value === "toggle";
			if ( value === ( hidden ? "hide" : "show" ) ) {

				// If there is dataShow left over from a stopped hide or show and we are going to proceed with show, we should pretend to be hidden
				if ( value === "show" && dataShow && dataShow[ prop ] !== undefined ) {
					hidden = true;
				} else {
					continue;
				}
			}
			orig[ prop ] = dataShow && dataShow[ prop ] || jQuery.style( elem, prop );

		// Any non-fx value stops us from restoring the original display value
		} else {
			display = undefined;
		}
	}

	if ( !jQuery.isEmptyObject( orig ) ) {
		if ( dataShow ) {
			if ( "hidden" in dataShow ) {
				hidden = dataShow.hidden;
			}
		} else {
			dataShow = jQuery._data( elem, "fxshow", {} );
		}

		// store state if its toggle - enables .stop().toggle() to "reverse"
		if ( toggle ) {
			dataShow.hidden = !hidden;
		}
		if ( hidden ) {
			jQuery( elem ).show();
		} else {
			anim.done(function() {
				jQuery( elem ).hide();
			});
		}
		anim.done(function() {
			var prop;
			jQuery._removeData( elem, "fxshow" );
			for ( prop in orig ) {
				jQuery.style( elem, prop, orig[ prop ] );
			}
		});
		for ( prop in orig ) {
			tween = createTween( hidden ? dataShow[ prop ] : 0, prop, anim );

			if ( !( prop in dataShow ) ) {
				dataShow[ prop ] = tween.start;
				if ( hidden ) {
					tween.end = tween.start;
					tween.start = prop === "width" || prop === "height" ? 1 : 0;
				}
			}
		}

	// If this is a noop like .hide().hide(), restore an overwritten display value
	} else if ( (display === "none" ? defaultDisplay( elem.nodeName ) : display) === "inline" ) {
		style.display = display;
	}
}

function propFilter( props, specialEasing ) {
	var index, name, easing, value, hooks;

	// camelCase, specialEasing and expand cssHook pass
	for ( index in props ) {
		name = jQuery.camelCase( index );
		easing = specialEasing[ name ];
		value = props[ index ];
		if ( jQuery.isArray( value ) ) {
			easing = value[ 1 ];
			value = props[ index ] = value[ 0 ];
		}

		if ( index !== name ) {
			props[ name ] = value;
			delete props[ index ];
		}

		hooks = jQuery.cssHooks[ name ];
		if ( hooks && "expand" in hooks ) {
			value = hooks.expand( value );
			delete props[ name ];

			// not quite $.extend, this wont overwrite keys already present.
			// also - reusing 'index' from above because we have the correct "name"
			for ( index in value ) {
				if ( !( index in props ) ) {
					props[ index ] = value[ index ];
					specialEasing[ index ] = easing;
				}
			}
		} else {
			specialEasing[ name ] = easing;
		}
	}
}

function Animation( elem, properties, options ) {
	var result,
		stopped,
		index = 0,
		length = animationPrefilters.length,
		deferred = jQuery.Deferred().always( function() {
			// don't match elem in the :animated selector
			delete tick.elem;
		}),
		tick = function() {
			if ( stopped ) {
				return false;
			}
			var currentTime = fxNow || createFxNow(),
				remaining = Math.max( 0, animation.startTime + animation.duration - currentTime ),
				// archaic crash bug won't allow us to use 1 - ( 0.5 || 0 ) (#12497)
				temp = remaining / animation.duration || 0,
				percent = 1 - temp,
				index = 0,
				length = animation.tweens.length;

			for ( ; index < length ; index++ ) {
				animation.tweens[ index ].run( percent );
			}

			deferred.notifyWith( elem, [ animation, percent, remaining ]);

			if ( percent < 1 && length ) {
				return remaining;
			} else {
				deferred.resolveWith( elem, [ animation ] );
				return false;
			}
		},
		animation = deferred.promise({
			elem: elem,
			props: jQuery.extend( {}, properties ),
			opts: jQuery.extend( true, { specialEasing: {} }, options ),
			originalProperties: properties,
			originalOptions: options,
			startTime: fxNow || createFxNow(),
			duration: options.duration,
			tweens: [],
			createTween: function( prop, end ) {
				var tween = jQuery.Tween( elem, animation.opts, prop, end,
						animation.opts.specialEasing[ prop ] || animation.opts.easing );
				animation.tweens.push( tween );
				return tween;
			},
			stop: function( gotoEnd ) {
				var index = 0,
					// if we are going to the end, we want to run all the tweens
					// otherwise we skip this part
					length = gotoEnd ? animation.tweens.length : 0;
				if ( stopped ) {
					return this;
				}
				stopped = true;
				for ( ; index < length ; index++ ) {
					animation.tweens[ index ].run( 1 );
				}

				// resolve when we played the last frame
				// otherwise, reject
				if ( gotoEnd ) {
					deferred.resolveWith( elem, [ animation, gotoEnd ] );
				} else {
					deferred.rejectWith( elem, [ animation, gotoEnd ] );
				}
				return this;
			}
		}),
		props = animation.props;

	propFilter( props, animation.opts.specialEasing );

	for ( ; index < length ; index++ ) {
		result = animationPrefilters[ index ].call( animation, elem, props, animation.opts );
		if ( result ) {
			return result;
		}
	}

	jQuery.map( props, createTween, animation );

	if ( jQuery.isFunction( animation.opts.start ) ) {
		animation.opts.start.call( elem, animation );
	}

	jQuery.fx.timer(
		jQuery.extend( tick, {
			elem: elem,
			anim: animation,
			queue: animation.opts.queue
		})
	);

	// attach callbacks from options
	return animation.progress( animation.opts.progress )
		.done( animation.opts.done, animation.opts.complete )
		.fail( animation.opts.fail )
		.always( animation.opts.always );
}

jQuery.Animation = jQuery.extend( Animation, {
	tweener: function( props, callback ) {
		if ( jQuery.isFunction( props ) ) {
			callback = props;
			props = [ "*" ];
		} else {
			props = props.split(" ");
		}

		var prop,
			index = 0,
			length = props.length;

		for ( ; index < length ; index++ ) {
			prop = props[ index ];
			tweeners[ prop ] = tweeners[ prop ] || [];
			tweeners[ prop ].unshift( callback );
		}
	},

	prefilter: function( callback, prepend ) {
		if ( prepend ) {
			animationPrefilters.unshift( callback );
		} else {
			animationPrefilters.push( callback );
		}
	}
});

jQuery.speed = function( speed, easing, fn ) {
	var opt = speed && typeof speed === "object" ? jQuery.extend( {}, speed ) : {
		complete: fn || !fn && easing ||
			jQuery.isFunction( speed ) && speed,
		duration: speed,
		easing: fn && easing || easing && !jQuery.isFunction( easing ) && easing
	};

	opt.duration = jQuery.fx.off ? 0 : typeof opt.duration === "number" ? opt.duration :
		opt.duration in jQuery.fx.speeds ? jQuery.fx.speeds[ opt.duration ] : jQuery.fx.speeds._default;

	// normalize opt.queue - true/undefined/null -> "fx"
	if ( opt.queue == null || opt.queue === true ) {
		opt.queue = "fx";
	}

	// Queueing
	opt.old = opt.complete;

	opt.complete = function() {
		if ( jQuery.isFunction( opt.old ) ) {
			opt.old.call( this );
		}

		if ( opt.queue ) {
			jQuery.dequeue( this, opt.queue );
		}
	};

	return opt;
};

jQuery.fn.extend({
	fadeTo: function( speed, to, easing, callback ) {

		// show any hidden elements after setting opacity to 0
		return this.filter( isHidden ).css( "opacity", 0 ).show()

			// animate to the value specified
			.end().animate({ opacity: to }, speed, easing, callback );
	},
	animate: function( prop, speed, easing, callback ) {
		var empty = jQuery.isEmptyObject( prop ),
			optall = jQuery.speed( speed, easing, callback ),
			doAnimation = function() {
				// Operate on a copy of prop so per-property easing won't be lost
				var anim = Animation( this, jQuery.extend( {}, prop ), optall );

				// Empty animations, or finishing resolves immediately
				if ( empty || jQuery._data( this, "finish" ) ) {
					anim.stop( true );
				}
			};
			doAnimation.finish = doAnimation;

		return empty || optall.queue === false ?
			this.each( doAnimation ) :
			this.queue( optall.queue, doAnimation );
	},
	stop: function( type, clearQueue, gotoEnd ) {
		var stopQueue = function( hooks ) {
			var stop = hooks.stop;
			delete hooks.stop;
			stop( gotoEnd );
		};

		if ( typeof type !== "string" ) {
			gotoEnd = clearQueue;
			clearQueue = type;
			type = undefined;
		}
		if ( clearQueue && type !== false ) {
			this.queue( type || "fx", [] );
		}

		return this.each(function() {
			var dequeue = true,
				index = type != null && type + "queueHooks",
				timers = jQuery.timers,
				data = jQuery._data( this );

			if ( index ) {
				if ( data[ index ] && data[ index ].stop ) {
					stopQueue( data[ index ] );
				}
			} else {
				for ( index in data ) {
					if ( data[ index ] && data[ index ].stop && rrun.test( index ) ) {
						stopQueue( data[ index ] );
					}
				}
			}

			for ( index = timers.length; index--; ) {
				if ( timers[ index ].elem === this && (type == null || timers[ index ].queue === type) ) {
					timers[ index ].anim.stop( gotoEnd );
					dequeue = false;
					timers.splice( index, 1 );
				}
			}

			// start the next in the queue if the last step wasn't forced
			// timers currently will call their complete callbacks, which will dequeue
			// but only if they were gotoEnd
			if ( dequeue || !gotoEnd ) {
				jQuery.dequeue( this, type );
			}
		});
	},
	finish: function( type ) {
		if ( type !== false ) {
			type = type || "fx";
		}
		return this.each(function() {
			var index,
				data = jQuery._data( this ),
				queue = data[ type + "queue" ],
				hooks = data[ type + "queueHooks" ],
				timers = jQuery.timers,
				length = queue ? queue.length : 0;

			// enable finishing flag on private data
			data.finish = true;

			// empty the queue first
			jQuery.queue( this, type, [] );

			if ( hooks && hooks.stop ) {
				hooks.stop.call( this, true );
			}

			// look for any active animations, and finish them
			for ( index = timers.length; index--; ) {
				if ( timers[ index ].elem === this && timers[ index ].queue === type ) {
					timers[ index ].anim.stop( true );
					timers.splice( index, 1 );
				}
			}

			// look for any animations in the old queue and finish them
			for ( index = 0; index < length; index++ ) {
				if ( queue[ index ] && queue[ index ].finish ) {
					queue[ index ].finish.call( this );
				}
			}

			// turn off finishing flag
			delete data.finish;
		});
	}
});

jQuery.each([ "toggle", "show", "hide" ], function( i, name ) {
	var cssFn = jQuery.fn[ name ];
	jQuery.fn[ name ] = function( speed, easing, callback ) {
		return speed == null || typeof speed === "boolean" ?
			cssFn.apply( this, arguments ) :
			this.animate( genFx( name, true ), speed, easing, callback );
	};
});

// Generate shortcuts for custom animations
jQuery.each({
	slideDown: genFx("show"),
	slideUp: genFx("hide"),
	slideToggle: genFx("toggle"),
	fadeIn: { opacity: "show" },
	fadeOut: { opacity: "hide" },
	fadeToggle: { opacity: "toggle" }
}, function( name, props ) {
	jQuery.fn[ name ] = function( speed, easing, callback ) {
		return this.animate( props, speed, easing, callback );
	};
});

jQuery.timers = [];
jQuery.fx.tick = function() {
	var timer,
		timers = jQuery.timers,
		i = 0;

	fxNow = jQuery.now();

	for ( ; i < timers.length; i++ ) {
		timer = timers[ i ];
		// Checks the timer has not already been removed
		if ( !timer() && timers[ i ] === timer ) {
			timers.splice( i--, 1 );
		}
	}

	if ( !timers.length ) {
		jQuery.fx.stop();
	}
	fxNow = undefined;
};

jQuery.fx.timer = function( timer ) {
	jQuery.timers.push( timer );
	if ( timer() ) {
		jQuery.fx.start();
	} else {
		jQuery.timers.pop();
	}
};

jQuery.fx.interval = 13;

jQuery.fx.start = function() {
	if ( !timerId ) {
		timerId = setInterval( jQuery.fx.tick, jQuery.fx.interval );
	}
};

jQuery.fx.stop = function() {
	clearInterval( timerId );
	timerId = null;
};

jQuery.fx.speeds = {
	slow: 600,
	fast: 200,
	// Default speed
	_default: 400
};


// Based off of the plugin by Clint Helfers, with permission.
// http://blindsignals.com/index.php/2009/07/jquery-delay/
jQuery.fn.delay = function( time, type ) {
	time = jQuery.fx ? jQuery.fx.speeds[ time ] || time : time;
	type = type || "fx";

	return this.queue( type, function( next, hooks ) {
		var timeout = setTimeout( next, time );
		hooks.stop = function() {
			clearTimeout( timeout );
		};
	});
};


(function() {
	// Minified: var a,b,c,d,e
	var input, div, select, a, opt;

	// Setup
	div = document.createElement( "div" );
	div.setAttribute( "className", "t" );
	div.innerHTML = "  <link/><table></table><a href='/a'>a</a><input type='checkbox'/>";
	a = div.getElementsByTagName("a")[ 0 ];

	// First batch of tests.
	select = document.createElement("select");
	opt = select.appendChild( document.createElement("option") );
	input = div.getElementsByTagName("input")[ 0 ];

	a.style.cssText = "top:1px";

	// Test setAttribute on camelCase class. If it works, we need attrFixes when doing get/setAttribute (ie6/7)
	support.getSetAttribute = div.className !== "t";

	// Get the style information from getAttribute
	// (IE uses .cssText instead)
	support.style = /top/.test( a.getAttribute("style") );

	// Make sure that URLs aren't manipulated
	// (IE normalizes it by default)
	support.hrefNormalized = a.getAttribute("href") === "/a";

	// Check the default checkbox/radio value ("" on WebKit; "on" elsewhere)
	support.checkOn = !!input.value;

	// Make sure that a selected-by-default option has a working selected property.
	// (WebKit defaults to false instead of true, IE too, if it's in an optgroup)
	support.optSelected = opt.selected;

	// Tests for enctype support on a form (#6743)
	support.enctype = !!document.createElement("form").enctype;

	// Make sure that the options inside disabled selects aren't marked as disabled
	// (WebKit marks them as disabled)
	select.disabled = true;
	support.optDisabled = !opt.disabled;

	// Support: IE8 only
	// Check if we can trust getAttribute("value")
	input = document.createElement( "input" );
	input.setAttribute( "value", "" );
	support.input = input.getAttribute( "value" ) === "";

	// Check if an input maintains its value after becoming a radio
	input.value = "t";
	input.setAttribute( "type", "radio" );
	support.radioValue = input.value === "t";
})();


var rreturn = /\r/g;

jQuery.fn.extend({
	val: function( value ) {
		var hooks, ret, isFunction,
			elem = this[0];

		if ( !arguments.length ) {
			if ( elem ) {
				hooks = jQuery.valHooks[ elem.type ] || jQuery.valHooks[ elem.nodeName.toLowerCase() ];

				if ( hooks && "get" in hooks && (ret = hooks.get( elem, "value" )) !== undefined ) {
					return ret;
				}

				ret = elem.value;

				return typeof ret === "string" ?
					// handle most common string cases
					ret.replace(rreturn, "") :
					// handle cases where value is null/undef or number
					ret == null ? "" : ret;
			}

			return;
		}

		isFunction = jQuery.isFunction( value );

		return this.each(function( i ) {
			var val;

			if ( this.nodeType !== 1 ) {
				return;
			}

			if ( isFunction ) {
				val = value.call( this, i, jQuery( this ).val() );
			} else {
				val = value;
			}

			// Treat null/undefined as ""; convert numbers to string
			if ( val == null ) {
				val = "";
			} else if ( typeof val === "number" ) {
				val += "";
			} else if ( jQuery.isArray( val ) ) {
				val = jQuery.map( val, function( value ) {
					return value == null ? "" : value + "";
				});
			}

			hooks = jQuery.valHooks[ this.type ] || jQuery.valHooks[ this.nodeName.toLowerCase() ];

			// If set returns undefined, fall back to normal setting
			if ( !hooks || !("set" in hooks) || hooks.set( this, val, "value" ) === undefined ) {
				this.value = val;
			}
		});
	}
});

jQuery.extend({
	valHooks: {
		option: {
			get: function( elem ) {
				var val = jQuery.find.attr( elem, "value" );
				return val != null ?
					val :
					// Support: IE10-11+
					// option.text throws exceptions (#14686, #14858)
					jQuery.trim( jQuery.text( elem ) );
			}
		},
		select: {
			get: function( elem ) {
				var value, option,
					options = elem.options,
					index = elem.selectedIndex,
					one = elem.type === "select-one" || index < 0,
					values = one ? null : [],
					max = one ? index + 1 : options.length,
					i = index < 0 ?
						max :
						one ? index : 0;

				// Loop through all the selected options
				for ( ; i < max; i++ ) {
					option = options[ i ];

					// oldIE doesn't update selected after form reset (#2551)
					if ( ( option.selected || i === index ) &&
							// Don't return options that are disabled or in a disabled optgroup
							( support.optDisabled ? !option.disabled : option.getAttribute("disabled") === null ) &&
							( !option.parentNode.disabled || !jQuery.nodeName( option.parentNode, "optgroup" ) ) ) {

						// Get the specific value for the option
						value = jQuery( option ).val();

						// We don't need an array for one selects
						if ( one ) {
							return value;
						}

						// Multi-Selects return an array
						values.push( value );
					}
				}

				return values;
			},

			set: function( elem, value ) {
				var optionSet, option,
					options = elem.options,
					values = jQuery.makeArray( value ),
					i = options.length;

				while ( i-- ) {
					option = options[ i ];

					if ( jQuery.inArray( jQuery.valHooks.option.get( option ), values ) >= 0 ) {

						// Support: IE6
						// When new option element is added to select box we need to
						// force reflow of newly added node in order to workaround delay
						// of initialization properties
						try {
							option.selected = optionSet = true;

						} catch ( _ ) {

							// Will be executed only in IE6
							option.scrollHeight;
						}

					} else {
						option.selected = false;
					}
				}

				// Force browsers to behave consistently when non-matching value is set
				if ( !optionSet ) {
					elem.selectedIndex = -1;
				}

				return options;
			}
		}
	}
});

// Radios and checkboxes getter/setter
jQuery.each([ "radio", "checkbox" ], function() {
	jQuery.valHooks[ this ] = {
		set: function( elem, value ) {
			if ( jQuery.isArray( value ) ) {
				return ( elem.checked = jQuery.inArray( jQuery(elem).val(), value ) >= 0 );
			}
		}
	};
	if ( !support.checkOn ) {
		jQuery.valHooks[ this ].get = function( elem ) {
			// Support: Webkit
			// "" is returned instead of "on" if a value isn't specified
			return elem.getAttribute("value") === null ? "on" : elem.value;
		};
	}
});


var nodeHook, boolHook,
	attrHandle = jQuery.expr.attrHandle,
	ruseDefault = /^(?:checked|selected)$/i,
	getSetAttribute = support.getSetAttribute,
	getSetInput = support.input;

jQuery.fn.extend({
	attr: function( name, value ) {
		return access( this, jQuery.attr, name, value, arguments.length > 1 );
	},

	removeAttr: function( name ) {
		return this.each(function() {
			jQuery.removeAttr( this, name );
		});
	}
});

jQuery.extend({
	attr: function( elem, name, value ) {
		var hooks, ret,
			nType = elem.nodeType;

		// don't get/set attributes on text, comment and attribute nodes
		if ( !elem || nType === 3 || nType === 8 || nType === 2 ) {
			return;
		}

		// Fallback to prop when attributes are not supported
		if ( typeof elem.getAttribute === strundefined ) {
			return jQuery.prop( elem, name, value );
		}

		// All attributes are lowercase
		// Grab necessary hook if one is defined
		if ( nType !== 1 || !jQuery.isXMLDoc( elem ) ) {
			name = name.toLowerCase();
			hooks = jQuery.attrHooks[ name ] ||
				( jQuery.expr.match.bool.test( name ) ? boolHook : nodeHook );
		}

		if ( value !== undefined ) {

			if ( value === null ) {
				jQuery.removeAttr( elem, name );

			} else if ( hooks && "set" in hooks && (ret = hooks.set( elem, value, name )) !== undefined ) {
				return ret;

			} else {
				elem.setAttribute( name, value + "" );
				return value;
			}

		} else if ( hooks && "get" in hooks && (ret = hooks.get( elem, name )) !== null ) {
			return ret;

		} else {
			ret = jQuery.find.attr( elem, name );

			// Non-existent attributes return null, we normalize to undefined
			return ret == null ?
				undefined :
				ret;
		}
	},

	removeAttr: function( elem, value ) {
		var name, propName,
			i = 0,
			attrNames = value && value.match( rnotwhite );

		if ( attrNames && elem.nodeType === 1 ) {
			while ( (name = attrNames[i++]) ) {
				propName = jQuery.propFix[ name ] || name;

				// Boolean attributes get special treatment (#10870)
				if ( jQuery.expr.match.bool.test( name ) ) {
					// Set corresponding property to false
					if ( getSetInput && getSetAttribute || !ruseDefault.test( name ) ) {
						elem[ propName ] = false;
					// Support: IE<9
					// Also clear defaultChecked/defaultSelected (if appropriate)
					} else {
						elem[ jQuery.camelCase( "default-" + name ) ] =
							elem[ propName ] = false;
					}

				// See #9699 for explanation of this approach (setting first, then removal)
				} else {
					jQuery.attr( elem, name, "" );
				}

				elem.removeAttribute( getSetAttribute ? name : propName );
			}
		}
	},

	attrHooks: {
		type: {
			set: function( elem, value ) {
				if ( !support.radioValue && value === "radio" && jQuery.nodeName(elem, "input") ) {
					// Setting the type on a radio button after the value resets the value in IE6-9
					// Reset value to default in case type is set after value during creation
					var val = elem.value;
					elem.setAttribute( "type", value );
					if ( val ) {
						elem.value = val;
					}
					return value;
				}
			}
		}
	}
});

// Hook for boolean attributes
boolHook = {
	set: function( elem, value, name ) {
		if ( value === false ) {
			// Remove boolean attributes when set to false
			jQuery.removeAttr( elem, name );
		} else if ( getSetInput && getSetAttribute || !ruseDefault.test( name ) ) {
			// IE<8 needs the *property* name
			elem.setAttribute( !getSetAttribute && jQuery.propFix[ name ] || name, name );

		// Use defaultChecked and defaultSelected for oldIE
		} else {
			elem[ jQuery.camelCase( "default-" + name ) ] = elem[ name ] = true;
		}

		return name;
	}
};

// Retrieve booleans specially
jQuery.each( jQuery.expr.match.bool.source.match( /\w+/g ), function( i, name ) {

	var getter = attrHandle[ name ] || jQuery.find.attr;

	attrHandle[ name ] = getSetInput && getSetAttribute || !ruseDefault.test( name ) ?
		function( elem, name, isXML ) {
			var ret, handle;
			if ( !isXML ) {
				// Avoid an infinite loop by temporarily removing this function from the getter
				handle = attrHandle[ name ];
				attrHandle[ name ] = ret;
				ret = getter( elem, name, isXML ) != null ?
					name.toLowerCase() :
					null;
				attrHandle[ name ] = handle;
			}
			return ret;
		} :
		function( elem, name, isXML ) {
			if ( !isXML ) {
				return elem[ jQuery.camelCase( "default-" + name ) ] ?
					name.toLowerCase() :
					null;
			}
		};
});

// fix oldIE attroperties
if ( !getSetInput || !getSetAttribute ) {
	jQuery.attrHooks.value = {
		set: function( elem, value, name ) {
			if ( jQuery.nodeName( elem, "input" ) ) {
				// Does not return so that setAttribute is also used
				elem.defaultValue = value;
			} else {
				// Use nodeHook if defined (#1954); otherwise setAttribute is fine
				return nodeHook && nodeHook.set( elem, value, name );
			}
		}
	};
}

// IE6/7 do not support getting/setting some attributes with get/setAttribute
if ( !getSetAttribute ) {

	// Use this for any attribute in IE6/7
	// This fixes almost every IE6/7 issue
	nodeHook = {
		set: function( elem, value, name ) {
			// Set the existing or create a new attribute node
			var ret = elem.getAttributeNode( name );
			if ( !ret ) {
				elem.setAttributeNode(
					(ret = elem.ownerDocument.createAttribute( name ))
				);
			}

			ret.value = value += "";

			// Break association with cloned elements by also using setAttribute (#9646)
			if ( name === "value" || value === elem.getAttribute( name ) ) {
				return value;
			}
		}
	};

	// Some attributes are constructed with empty-string values when not defined
	attrHandle.id = attrHandle.name = attrHandle.coords =
		function( elem, name, isXML ) {
			var ret;
			if ( !isXML ) {
				return (ret = elem.getAttributeNode( name )) && ret.value !== "" ?
					ret.value :
					null;
			}
		};

	// Fixing value retrieval on a button requires this module
	jQuery.valHooks.button = {
		get: function( elem, name ) {
			var ret = elem.getAttributeNode( name );
			if ( ret && ret.specified ) {
				return ret.value;
			}
		},
		set: nodeHook.set
	};

	// Set contenteditable to false on removals(#10429)
	// Setting to empty string throws an error as an invalid value
	jQuery.attrHooks.contenteditable = {
		set: function( elem, value, name ) {
			nodeHook.set( elem, value === "" ? false : value, name );
		}
	};

	// Set width and height to auto instead of 0 on empty string( Bug #8150 )
	// This is for removals
	jQuery.each([ "width", "height" ], function( i, name ) {
		jQuery.attrHooks[ name ] = {
			set: function( elem, value ) {
				if ( value === "" ) {
					elem.setAttribute( name, "auto" );
					return value;
				}
			}
		};
	});
}

if ( !support.style ) {
	jQuery.attrHooks.style = {
		get: function( elem ) {
			// Return undefined in the case of empty string
			// Note: IE uppercases css property names, but if we were to .toLowerCase()
			// .cssText, that would destroy case senstitivity in URL's, like in "background"
			return elem.style.cssText || undefined;
		},
		set: function( elem, value ) {
			return ( elem.style.cssText = value + "" );
		}
	};
}


var rfocusable = /^(?:input|select|textarea|button|object)$/i,
	rclickable = /^(?:a|area)$/i;

jQuery.fn.extend({
	prop: function( name, value ) {
		return access( this, jQuery.prop, name, value, arguments.length > 1 );
	},

	removeProp: function( name ) {
		name = jQuery.propFix[ name ] || name;
		return this.each(function() {
			// try/catch handles cases where IE balks (such as removing a property on window)
			try {
				this[ name ] = undefined;
				delete this[ name ];
			} catch( e ) {}
		});
	}
});

jQuery.extend({
	propFix: {
		"for": "htmlFor",
		"class": "className"
	},

	prop: function( elem, name, value ) {
		var ret, hooks, notxml,
			nType = elem.nodeType;

		// don't get/set properties on text, comment and attribute nodes
		if ( !elem || nType === 3 || nType === 8 || nType === 2 ) {
			return;
		}

		notxml = nType !== 1 || !jQuery.isXMLDoc( elem );

		if ( notxml ) {
			// Fix name and attach hooks
			name = jQuery.propFix[ name ] || name;
			hooks = jQuery.propHooks[ name ];
		}

		if ( value !== undefined ) {
			return hooks && "set" in hooks && (ret = hooks.set( elem, value, name )) !== undefined ?
				ret :
				( elem[ name ] = value );

		} else {
			return hooks && "get" in hooks && (ret = hooks.get( elem, name )) !== null ?
				ret :
				elem[ name ];
		}
	},

	propHooks: {
		tabIndex: {
			get: function( elem ) {
				// elem.tabIndex doesn't always return the correct value when it hasn't been explicitly set
				// http://fluidproject.org/blog/2008/01/09/getting-setting-and-removing-tabindex-values-with-javascript/
				// Use proper attribute retrieval(#12072)
				var tabindex = jQuery.find.attr( elem, "tabindex" );

				return tabindex ?
					parseInt( tabindex, 10 ) :
					rfocusable.test( elem.nodeName ) || rclickable.test( elem.nodeName ) && elem.href ?
						0 :
						-1;
			}
		}
	}
});

// Some attributes require a special call on IE
// http://msdn.microsoft.com/en-us/library/ms536429%28VS.85%29.aspx
if ( !support.hrefNormalized ) {
	// href/src property should get the full normalized URL (#10299/#12915)
	jQuery.each([ "href", "src" ], function( i, name ) {
		jQuery.propHooks[ name ] = {
			get: function( elem ) {
				return elem.getAttribute( name, 4 );
			}
		};
	});
}

// Support: Safari, IE9+
// mis-reports the default selected property of an option
// Accessing the parent's selectedIndex property fixes it
if ( !support.optSelected ) {
	jQuery.propHooks.selected = {
		get: function( elem ) {
			var parent = elem.parentNode;

			if ( parent ) {
				parent.selectedIndex;

				// Make sure that it also works with optgroups, see #5701
				if ( parent.parentNode ) {
					parent.parentNode.selectedIndex;
				}
			}
			return null;
		}
	};
}

jQuery.each([
	"tabIndex",
	"readOnly",
	"maxLength",
	"cellSpacing",
	"cellPadding",
	"rowSpan",
	"colSpan",
	"useMap",
	"frameBorder",
	"contentEditable"
], function() {
	jQuery.propFix[ this.toLowerCase() ] = this;
});

// IE6/7 call enctype encoding
if ( !support.enctype ) {
	jQuery.propFix.enctype = "encoding";
}


var rclass = /[\t\r\n\f]/g;

jQuery.fn.extend({
	addClass: function( value ) {
		var classes, elem, cur, clazz, j, finalValue,
			i = 0,
			len = this.length,
			proceed = typeof value === "string" && value;

		if ( jQuery.isFunction( value ) ) {
			return this.each(function( j ) {
				jQuery( this ).addClass( value.call( this, j, this.className ) );
			});
		}

		if ( proceed ) {
			// The disjunction here is for better compressibility (see removeClass)
			classes = ( value || "" ).match( rnotwhite ) || [];

			for ( ; i < len; i++ ) {
				elem = this[ i ];
				cur = elem.nodeType === 1 && ( elem.className ?
					( " " + elem.className + " " ).replace( rclass, " " ) :
					" "
				);

				if ( cur ) {
					j = 0;
					while ( (clazz = classes[j++]) ) {
						if ( cur.indexOf( " " + clazz + " " ) < 0 ) {
							cur += clazz + " ";
						}
					}

					// only assign if different to avoid unneeded rendering.
					finalValue = jQuery.trim( cur );
					if ( elem.className !== finalValue ) {
						elem.className = finalValue;
					}
				}
			}
		}

		return this;
	},

	removeClass: function( value ) {
		var classes, elem, cur, clazz, j, finalValue,
			i = 0,
			len = this.length,
			proceed = arguments.length === 0 || typeof value === "string" && value;

		if ( jQuery.isFunction( value ) ) {
			return this.each(function( j ) {
				jQuery( this ).removeClass( value.call( this, j, this.className ) );
			});
		}
		if ( proceed ) {
			classes = ( value || "" ).match( rnotwhite ) || [];

			for ( ; i < len; i++ ) {
				elem = this[ i ];
				// This expression is here for better compressibility (see addClass)
				cur = elem.nodeType === 1 && ( elem.className ?
					( " " + elem.className + " " ).replace( rclass, " " ) :
					""
				);

				if ( cur ) {
					j = 0;
					while ( (clazz = classes[j++]) ) {
						// Remove *all* instances
						while ( cur.indexOf( " " + clazz + " " ) >= 0 ) {
							cur = cur.replace( " " + clazz + " ", " " );
						}
					}

					// only assign if different to avoid unneeded rendering.
					finalValue = value ? jQuery.trim( cur ) : "";
					if ( elem.className !== finalValue ) {
						elem.className = finalValue;
					}
				}
			}
		}

		return this;
	},

	toggleClass: function( value, stateVal ) {
		var type = typeof value;

		if ( typeof stateVal === "boolean" && type === "string" ) {
			return stateVal ? this.addClass( value ) : this.removeClass( value );
		}

		if ( jQuery.isFunction( value ) ) {
			return this.each(function( i ) {
				jQuery( this ).toggleClass( value.call(this, i, this.className, stateVal), stateVal );
			});
		}

		return this.each(function() {
			if ( type === "string" ) {
				// toggle individual class names
				var className,
					i = 0,
					self = jQuery( this ),
					classNames = value.match( rnotwhite ) || [];

				while ( (className = classNames[ i++ ]) ) {
					// check each className given, space separated list
					if ( self.hasClass( className ) ) {
						self.removeClass( className );
					} else {
						self.addClass( className );
					}
				}

			// Toggle whole class name
			} else if ( type === strundefined || type === "boolean" ) {
				if ( this.className ) {
					// store className if set
					jQuery._data( this, "__className__", this.className );
				}

				// If the element has a class name or if we're passed "false",
				// then remove the whole classname (if there was one, the above saved it).
				// Otherwise bring back whatever was previously saved (if anything),
				// falling back to the empty string if nothing was stored.
				this.className = this.className || value === false ? "" : jQuery._data( this, "__className__" ) || "";
			}
		});
	},

	hasClass: function( selector ) {
		var className = " " + selector + " ",
			i = 0,
			l = this.length;
		for ( ; i < l; i++ ) {
			if ( this[i].nodeType === 1 && (" " + this[i].className + " ").replace(rclass, " ").indexOf( className ) >= 0 ) {
				return true;
			}
		}

		return false;
	}
});


// Return jQuery for attributes-only inclusion


jQuery.each( ("blur focus focusin focusout load resize scroll unload click dblclick " +
	"mousedown mouseup mousemove mouseover mouseout mouseenter mouseleave " +
	"change select submit keydown keypress keyup error contextmenu").split(" "), function( i, name ) {

	// Handle event binding
	jQuery.fn[ name ] = function( data, fn ) {
		return arguments.length > 0 ?
			this.on( name, null, data, fn ) :
			this.trigger( name );
	};
});

jQuery.fn.extend({
	hover: function( fnOver, fnOut ) {
		return this.mouseenter( fnOver ).mouseleave( fnOut || fnOver );
	},

	bind: function( types, data, fn ) {
		return this.on( types, null, data, fn );
	},
	unbind: function( types, fn ) {
		return this.off( types, null, fn );
	},

	delegate: function( selector, types, data, fn ) {
		return this.on( types, selector, data, fn );
	},
	undelegate: function( selector, types, fn ) {
		// ( namespace ) or ( selector, types [, fn] )
		return arguments.length === 1 ? this.off( selector, "**" ) : this.off( types, selector || "**", fn );
	}
});


var nonce = jQuery.now();

var rquery = (/\?/);


var rvalidtokens = /(,)|(\[|{)|(}|])|"(?:[^"\\\r\n]|\\["\\\/bfnrt]|\\u[\da-fA-F]{4})*"\s*:?|true|false|null|-?(?!0\d)\d+(?:\.\d+|)(?:[eE][+-]?\d+|)/g;

jQuery.parseJSON = function( data ) {
	// Attempt to parse using the native JSON parser first
	if ( window.JSON && window.JSON.parse ) {
		// Support: Android 2.3
		// Workaround failure to string-cast null input
		return window.JSON.parse( data + "" );
	}

	var requireNonComma,
		depth = null,
		str = jQuery.trim( data + "" );

	// Guard against invalid (and possibly dangerous) input by ensuring that nothing remains
	// after removing valid tokens
	return str && !jQuery.trim( str.replace( rvalidtokens, function( token, comma, open, close ) {

		// Force termination if we see a misplaced comma
		if ( requireNonComma && comma ) {
			depth = 0;
		}

		// Perform no more replacements after returning to outermost depth
		if ( depth === 0 ) {
			return token;
		}

		// Commas must not follow "[", "{", or ","
		requireNonComma = open || comma;

		// Determine new depth
		// array/object open ("[" or "{"): depth += true - false (increment)
		// array/object close ("]" or "}"): depth += false - true (decrement)
		// other cases ("," or primitive): depth += true - true (numeric cast)
		depth += !close - !open;

		// Remove this token
		return "";
	}) ) ?
		( Function( "return " + str ) )() :
		jQuery.error( "Invalid JSON: " + data );
};


// Cross-browser xml parsing
jQuery.parseXML = function( data ) {
	var xml, tmp;
	if ( !data || typeof data !== "string" ) {
		return null;
	}
	try {
		if ( window.DOMParser ) { // Standard
			tmp = new DOMParser();
			xml = tmp.parseFromString( data, "text/xml" );
		} else { // IE
			xml = new ActiveXObject( "Microsoft.XMLDOM" );
			xml.async = "false";
			xml.loadXML( data );
		}
	} catch( e ) {
		xml = undefined;
	}
	if ( !xml || !xml.documentElement || xml.getElementsByTagName( "parsererror" ).length ) {
		jQuery.error( "Invalid XML: " + data );
	}
	return xml;
};


var
	// Document location
	ajaxLocParts,
	ajaxLocation,

	rhash = /#.*$/,
	rts = /([?&])_=[^&]*/,
	rheaders = /^(.*?):[ \t]*([^\r\n]*)\r?$/mg, // IE leaves an \r character at EOL
	// #7653, #8125, #8152: local protocol detection
	rlocalProtocol = /^(?:about|app|app-storage|.+-extension|file|res|widget):$/,
	rnoContent = /^(?:GET|HEAD)$/,
	rprotocol = /^\/\//,
	rurl = /^([\w.+-]+:)(?:\/\/(?:[^\/?#]*@|)([^\/?#:]*)(?::(\d+)|)|)/,

	/* Prefilters
	 * 1) They are useful to introduce custom dataTypes (see ajax/jsonp.js for an example)
	 * 2) These are called:
	 *    - BEFORE asking for a transport
	 *    - AFTER param serialization (s.data is a string if s.processData is true)
	 * 3) key is the dataType
	 * 4) the catchall symbol "*" can be used
	 * 5) execution will start with transport dataType and THEN continue down to "*" if needed
	 */
	prefilters = {},

	/* Transports bindings
	 * 1) key is the dataType
	 * 2) the catchall symbol "*" can be used
	 * 3) selection will start with transport dataType and THEN go to "*" if needed
	 */
	transports = {},

	// Avoid comment-prolog char sequence (#10098); must appease lint and evade compression
	allTypes = "*/".concat("*");

// #8138, IE may throw an exception when accessing
// a field from window.location if document.domain has been set
try {
	ajaxLocation = location.href;
} catch( e ) {
	// Use the href attribute of an A element
	// since IE will modify it given document.location
	ajaxLocation = document.createElement( "a" );
	ajaxLocation.href = "";
	ajaxLocation = ajaxLocation.href;
}

// Segment location into parts
ajaxLocParts = rurl.exec( ajaxLocation.toLowerCase() ) || [];

// Base "constructor" for jQuery.ajaxPrefilter and jQuery.ajaxTransport
function addToPrefiltersOrTransports( structure ) {

	// dataTypeExpression is optional and defaults to "*"
	return function( dataTypeExpression, func ) {

		if ( typeof dataTypeExpression !== "string" ) {
			func = dataTypeExpression;
			dataTypeExpression = "*";
		}

		var dataType,
			i = 0,
			dataTypes = dataTypeExpression.toLowerCase().match( rnotwhite ) || [];

		if ( jQuery.isFunction( func ) ) {
			// For each dataType in the dataTypeExpression
			while ( (dataType = dataTypes[i++]) ) {
				// Prepend if requested
				if ( dataType.charAt( 0 ) === "+" ) {
					dataType = dataType.slice( 1 ) || "*";
					(structure[ dataType ] = structure[ dataType ] || []).unshift( func );

				// Otherwise append
				} else {
					(structure[ dataType ] = structure[ dataType ] || []).push( func );
				}
			}
		}
	};
}

// Base inspection function for prefilters and transports
function inspectPrefiltersOrTransports( structure, options, originalOptions, jqXHR ) {

	var inspected = {},
		seekingTransport = ( structure === transports );

	function inspect( dataType ) {
		var selected;
		inspected[ dataType ] = true;
		jQuery.each( structure[ dataType ] || [], function( _, prefilterOrFactory ) {
			var dataTypeOrTransport = prefilterOrFactory( options, originalOptions, jqXHR );
			if ( typeof dataTypeOrTransport === "string" && !seekingTransport && !inspected[ dataTypeOrTransport ] ) {
				options.dataTypes.unshift( dataTypeOrTransport );
				inspect( dataTypeOrTransport );
				return false;
			} else if ( seekingTransport ) {
				return !( selected = dataTypeOrTransport );
			}
		});
		return selected;
	}

	return inspect( options.dataTypes[ 0 ] ) || !inspected[ "*" ] && inspect( "*" );
}

// A special extend for ajax options
// that takes "flat" options (not to be deep extended)
// Fixes #9887
function ajaxExtend( target, src ) {
	var deep, key,
		flatOptions = jQuery.ajaxSettings.flatOptions || {};

	for ( key in src ) {
		if ( src[ key ] !== undefined ) {
			( flatOptions[ key ] ? target : ( deep || (deep = {}) ) )[ key ] = src[ key ];
		}
	}
	if ( deep ) {
		jQuery.extend( true, target, deep );
	}

	return target;
}

/* Handles responses to an ajax request:
 * - finds the right dataType (mediates between content-type and expected dataType)
 * - returns the corresponding response
 */
function ajaxHandleResponses( s, jqXHR, responses ) {
	var firstDataType, ct, finalDataType, type,
		contents = s.contents,
		dataTypes = s.dataTypes;

	// Remove auto dataType and get content-type in the process
	while ( dataTypes[ 0 ] === "*" ) {
		dataTypes.shift();
		if ( ct === undefined ) {
			ct = s.mimeType || jqXHR.getResponseHeader("Content-Type");
		}
	}

	// Check if we're dealing with a known content-type
	if ( ct ) {
		for ( type in contents ) {
			if ( contents[ type ] && contents[ type ].test( ct ) ) {
				dataTypes.unshift( type );
				break;
			}
		}
	}

	// Check to see if we have a response for the expected dataType
	if ( dataTypes[ 0 ] in responses ) {
		finalDataType = dataTypes[ 0 ];
	} else {
		// Try convertible dataTypes
		for ( type in responses ) {
			if ( !dataTypes[ 0 ] || s.converters[ type + " " + dataTypes[0] ] ) {
				finalDataType = type;
				break;
			}
			if ( !firstDataType ) {
				firstDataType = type;
			}
		}
		// Or just use first one
		finalDataType = finalDataType || firstDataType;
	}

	// If we found a dataType
	// We add the dataType to the list if needed
	// and return the corresponding response
	if ( finalDataType ) {
		if ( finalDataType !== dataTypes[ 0 ] ) {
			dataTypes.unshift( finalDataType );
		}
		return responses[ finalDataType ];
	}
}

/* Chain conversions given the request and the original response
 * Also sets the responseXXX fields on the jqXHR instance
 */
function ajaxConvert( s, response, jqXHR, isSuccess ) {
	var conv2, current, conv, tmp, prev,
		converters = {},
		// Work with a copy of dataTypes in case we need to modify it for conversion
		dataTypes = s.dataTypes.slice();

	// Create converters map with lowercased keys
	if ( dataTypes[ 1 ] ) {
		for ( conv in s.converters ) {
			converters[ conv.toLowerCase() ] = s.converters[ conv ];
		}
	}

	current = dataTypes.shift();

	// Convert to each sequential dataType
	while ( current ) {

		if ( s.responseFields[ current ] ) {
			jqXHR[ s.responseFields[ current ] ] = response;
		}

		// Apply the dataFilter if provided
		if ( !prev && isSuccess && s.dataFilter ) {
			response = s.dataFilter( response, s.dataType );
		}

		prev = current;
		current = dataTypes.shift();

		if ( current ) {

			// There's only work to do if current dataType is non-auto
			if ( current === "*" ) {

				current = prev;

			// Convert response if prev dataType is non-auto and differs from current
			} else if ( prev !== "*" && prev !== current ) {

				// Seek a direct converter
				conv = converters[ prev + " " + current ] || converters[ "* " + current ];

				// If none found, seek a pair
				if ( !conv ) {
					for ( conv2 in converters ) {

						// If conv2 outputs current
						tmp = conv2.split( " " );
						if ( tmp[ 1 ] === current ) {

							// If prev can be converted to accepted input
							conv = converters[ prev + " " + tmp[ 0 ] ] ||
								converters[ "* " + tmp[ 0 ] ];
							if ( conv ) {
								// Condense equivalence converters
								if ( conv === true ) {
									conv = converters[ conv2 ];

								// Otherwise, insert the intermediate dataType
								} else if ( converters[ conv2 ] !== true ) {
									current = tmp[ 0 ];
									dataTypes.unshift( tmp[ 1 ] );
								}
								break;
							}
						}
					}
				}

				// Apply converter (if not an equivalence)
				if ( conv !== true ) {

					// Unless errors are allowed to bubble, catch and return them
					if ( conv && s[ "throws" ] ) {
						response = conv( response );
					} else {
						try {
							response = conv( response );
						} catch ( e ) {
							return { state: "parsererror", error: conv ? e : "No conversion from " + prev + " to " + current };
						}
					}
				}
			}
		}
	}

	return { state: "success", data: response };
}

jQuery.extend({

	// Counter for holding the number of active queries
	active: 0,

	// Last-Modified header cache for next request
	lastModified: {},
	etag: {},

	ajaxSettings: {
		url: ajaxLocation,
		type: "GET",
		isLocal: rlocalProtocol.test( ajaxLocParts[ 1 ] ),
		global: true,
		processData: true,
		async: true,
		contentType: "application/x-www-form-urlencoded; charset=UTF-8",
		/*
		timeout: 0,
		data: null,
		dataType: null,
		username: null,
		password: null,
		cache: null,
		throws: false,
		traditional: false,
		headers: {},
		*/

		accepts: {
			"*": allTypes,
			text: "text/plain",
			html: "text/html",
			xml: "application/xml, text/xml",
			json: "application/json, text/javascript"
		},

		contents: {
			xml: /xml/,
			html: /html/,
			json: /json/
		},

		responseFields: {
			xml: "responseXML",
			text: "responseText",
			json: "responseJSON"
		},

		// Data converters
		// Keys separate source (or catchall "*") and destination types with a single space
		converters: {

			// Convert anything to text
			"* text": String,

			// Text to html (true = no transformation)
			"text html": true,

			// Evaluate text as a json expression
			"text json": jQuery.parseJSON,

			// Parse text as xml
			"text xml": jQuery.parseXML
		},

		// For options that shouldn't be deep extended:
		// you can add your own custom options here if
		// and when you create one that shouldn't be
		// deep extended (see ajaxExtend)
		flatOptions: {
			url: true,
			context: true
		}
	},

	// Creates a full fledged settings object into target
	// with both ajaxSettings and settings fields.
	// If target is omitted, writes into ajaxSettings.
	ajaxSetup: function( target, settings ) {
		return settings ?

			// Building a settings object
			ajaxExtend( ajaxExtend( target, jQuery.ajaxSettings ), settings ) :

			// Extending ajaxSettings
			ajaxExtend( jQuery.ajaxSettings, target );
	},

	ajaxPrefilter: addToPrefiltersOrTransports( prefilters ),
	ajaxTransport: addToPrefiltersOrTransports( transports ),

	// Main method
	ajax: function( url, options ) {

		// If url is an object, simulate pre-1.5 signature
		if ( typeof url === "object" ) {
			options = url;
			url = undefined;
		}

		// Force options to be an object
		options = options || {};

		var // Cross-domain detection vars
			parts,
			// Loop variable
			i,
			// URL without anti-cache param
			cacheURL,
			// Response headers as string
			responseHeadersString,
			// timeout handle
			timeoutTimer,

			// To know if global events are to be dispatched
			fireGlobals,

			transport,
			// Response headers
			responseHeaders,
			// Create the final options object
			s = jQuery.ajaxSetup( {}, options ),
			// Callbacks context
			callbackContext = s.context || s,
			// Context for global events is callbackContext if it is a DOM node or jQuery collection
			globalEventContext = s.context && ( callbackContext.nodeType || callbackContext.jquery ) ?
				jQuery( callbackContext ) :
				jQuery.event,
			// Deferreds
			deferred = jQuery.Deferred(),
			completeDeferred = jQuery.Callbacks("once memory"),
			// Status-dependent callbacks
			statusCode = s.statusCode || {},
			// Headers (they are sent all at once)
			requestHeaders = {},
			requestHeadersNames = {},
			// The jqXHR state
			state = 0,
			// Default abort message
			strAbort = "canceled",
			// Fake xhr
			jqXHR = {
				readyState: 0,

				// Builds headers hashtable if needed
				getResponseHeader: function( key ) {
					var match;
					if ( state === 2 ) {
						if ( !responseHeaders ) {
							responseHeaders = {};
							while ( (match = rheaders.exec( responseHeadersString )) ) {
								responseHeaders[ match[1].toLowerCase() ] = match[ 2 ];
							}
						}
						match = responseHeaders[ key.toLowerCase() ];
					}
					return match == null ? null : match;
				},

				// Raw string
				getAllResponseHeaders: function() {
					return state === 2 ? responseHeadersString : null;
				},

				// Caches the header
				setRequestHeader: function( name, value ) {
					var lname = name.toLowerCase();
					if ( !state ) {
						name = requestHeadersNames[ lname ] = requestHeadersNames[ lname ] || name;
						requestHeaders[ name ] = value;
					}
					return this;
				},

				// Overrides response content-type header
				overrideMimeType: function( type ) {
					if ( !state ) {
						s.mimeType = type;
					}
					return this;
				},

				// Status-dependent callbacks
				statusCode: function( map ) {
					var code;
					if ( map ) {
						if ( state < 2 ) {
							for ( code in map ) {
								// Lazy-add the new callback in a way that preserves old ones
								statusCode[ code ] = [ statusCode[ code ], map[ code ] ];
							}
						} else {
							// Execute the appropriate callbacks
							jqXHR.always( map[ jqXHR.status ] );
						}
					}
					return this;
				},

				// Cancel the request
				abort: function( statusText ) {
					var finalText = statusText || strAbort;
					if ( transport ) {
						transport.abort( finalText );
					}
					done( 0, finalText );
					return this;
				}
			};

		// Attach deferreds
		deferred.promise( jqXHR ).complete = completeDeferred.add;
		jqXHR.success = jqXHR.done;
		jqXHR.error = jqXHR.fail;

		// Remove hash character (#7531: and string promotion)
		// Add protocol if not provided (#5866: IE7 issue with protocol-less urls)
		// Handle falsy url in the settings object (#10093: consistency with old signature)
		// We also use the url parameter if available
		s.url = ( ( url || s.url || ajaxLocation ) + "" ).replace( rhash, "" ).replace( rprotocol, ajaxLocParts[ 1 ] + "//" );

		// Alias method option to type as per ticket #12004
		s.type = options.method || options.type || s.method || s.type;

		// Extract dataTypes list
		s.dataTypes = jQuery.trim( s.dataType || "*" ).toLowerCase().match( rnotwhite ) || [ "" ];

		// A cross-domain request is in order when we have a protocol:host:port mismatch
		if ( s.crossDomain == null ) {
			parts = rurl.exec( s.url.toLowerCase() );
			s.crossDomain = !!( parts &&
				( parts[ 1 ] !== ajaxLocParts[ 1 ] || parts[ 2 ] !== ajaxLocParts[ 2 ] ||
					( parts[ 3 ] || ( parts[ 1 ] === "http:" ? "80" : "443" ) ) !==
						( ajaxLocParts[ 3 ] || ( ajaxLocParts[ 1 ] === "http:" ? "80" : "443" ) ) )
			);
		}

		// Convert data if not already a string
		if ( s.data && s.processData && typeof s.data !== "string" ) {
			s.data = jQuery.param( s.data, s.traditional );
		}

		// Apply prefilters
		inspectPrefiltersOrTransports( prefilters, s, options, jqXHR );

		// If request was aborted inside a prefilter, stop there
		if ( state === 2 ) {
			return jqXHR;
		}

		// We can fire global events as of now if asked to
		fireGlobals = s.global;

		// Watch for a new set of requests
		if ( fireGlobals && jQuery.active++ === 0 ) {
			jQuery.event.trigger("ajaxStart");
		}

		// Uppercase the type
		s.type = s.type.toUpperCase();

		// Determine if request has content
		s.hasContent = !rnoContent.test( s.type );

		// Save the URL in case we're toying with the If-Modified-Since
		// and/or If-None-Match header later on
		cacheURL = s.url;

		// More options handling for requests with no content
		if ( !s.hasContent ) {

			// If data is available, append data to url
			if ( s.data ) {
				cacheURL = ( s.url += ( rquery.test( cacheURL ) ? "&" : "?" ) + s.data );
				// #9682: remove data so that it's not used in an eventual retry
				delete s.data;
			}

			// Add anti-cache in url if needed
			if ( s.cache === false ) {
				s.url = rts.test( cacheURL ) ?

					// If there is already a '_' parameter, set its value
					cacheURL.replace( rts, "$1_=" + nonce++ ) :

					// Otherwise add one to the end
					cacheURL + ( rquery.test( cacheURL ) ? "&" : "?" ) + "_=" + nonce++;
			}
		}

		// Set the If-Modified-Since and/or If-None-Match header, if in ifModified mode.
		if ( s.ifModified ) {
			if ( jQuery.lastModified[ cacheURL ] ) {
				jqXHR.setRequestHeader( "If-Modified-Since", jQuery.lastModified[ cacheURL ] );
			}
			if ( jQuery.etag[ cacheURL ] ) {
				jqXHR.setRequestHeader( "If-None-Match", jQuery.etag[ cacheURL ] );
			}
		}

		// Set the correct header, if data is being sent
		if ( s.data && s.hasContent && s.contentType !== false || options.contentType ) {
			jqXHR.setRequestHeader( "Content-Type", s.contentType );
		}

		// Set the Accepts header for the server, depending on the dataType
		jqXHR.setRequestHeader(
			"Accept",
			s.dataTypes[ 0 ] && s.accepts[ s.dataTypes[0] ] ?
				s.accepts[ s.dataTypes[0] ] + ( s.dataTypes[ 0 ] !== "*" ? ", " + allTypes + "; q=0.01" : "" ) :
				s.accepts[ "*" ]
		);

		// Check for headers option
		for ( i in s.headers ) {
			jqXHR.setRequestHeader( i, s.headers[ i ] );
		}

		// Allow custom headers/mimetypes and early abort
		if ( s.beforeSend && ( s.beforeSend.call( callbackContext, jqXHR, s ) === false || state === 2 ) ) {
			// Abort if not done already and return
			return jqXHR.abort();
		}

		// aborting is no longer a cancellation
		strAbort = "abort";

		// Install callbacks on deferreds
		for ( i in { success: 1, error: 1, complete: 1 } ) {
			jqXHR[ i ]( s[ i ] );
		}

		// Get transport
		transport = inspectPrefiltersOrTransports( transports, s, options, jqXHR );

		// If no transport, we auto-abort
		if ( !transport ) {
			done( -1, "No Transport" );
		} else {
			jqXHR.readyState = 1;

			// Send global event
			if ( fireGlobals ) {
				globalEventContext.trigger( "ajaxSend", [ jqXHR, s ] );
			}
			// Timeout
			if ( s.async && s.timeout > 0 ) {
				timeoutTimer = setTimeout(function() {
					jqXHR.abort("timeout");
				}, s.timeout );
			}

			try {
				state = 1;
				transport.send( requestHeaders, done );
			} catch ( e ) {
				// Propagate exception as error if not done
				if ( state < 2 ) {
					done( -1, e );
				// Simply rethrow otherwise
				} else {
					throw e;
				}
			}
		}

		// Callback for when everything is done
		function done( status, nativeStatusText, responses, headers ) {
			var isSuccess, success, error, response, modified,
				statusText = nativeStatusText;

			// Called once
			if ( state === 2 ) {
				return;
			}

			// State is "done" now
			state = 2;

			// Clear timeout if it exists
			if ( timeoutTimer ) {
				clearTimeout( timeoutTimer );
			}

			// Dereference transport for early garbage collection
			// (no matter how long the jqXHR object will be used)
			transport = undefined;

			// Cache response headers
			responseHeadersString = headers || "";

			// Set readyState
			jqXHR.readyState = status > 0 ? 4 : 0;

			// Determine if successful
			isSuccess = status >= 200 && status < 300 || status === 304;

			// Get response data
			if ( responses ) {
				response = ajaxHandleResponses( s, jqXHR, responses );
			}

			// Convert no matter what (that way responseXXX fields are always set)
			response = ajaxConvert( s, response, jqXHR, isSuccess );

			// If successful, handle type chaining
			if ( isSuccess ) {

				// Set the If-Modified-Since and/or If-None-Match header, if in ifModified mode.
				if ( s.ifModified ) {
					modified = jqXHR.getResponseHeader("Last-Modified");
					if ( modified ) {
						jQuery.lastModified[ cacheURL ] = modified;
					}
					modified = jqXHR.getResponseHeader("etag");
					if ( modified ) {
						jQuery.etag[ cacheURL ] = modified;
					}
				}

				// if no content
				if ( status === 204 || s.type === "HEAD" ) {
					statusText = "nocontent";

				// if not modified
				} else if ( status === 304 ) {
					statusText = "notmodified";

				// If we have data, let's convert it
				} else {
					statusText = response.state;
					success = response.data;
					error = response.error;
					isSuccess = !error;
				}
			} else {
				// We extract error from statusText
				// then normalize statusText and status for non-aborts
				error = statusText;
				if ( status || !statusText ) {
					statusText = "error";
					if ( status < 0 ) {
						status = 0;
					}
				}
			}

			// Set data for the fake xhr object
			jqXHR.status = status;
			jqXHR.statusText = ( nativeStatusText || statusText ) + "";

			// Success/Error
			if ( isSuccess ) {
				deferred.resolveWith( callbackContext, [ success, statusText, jqXHR ] );
			} else {
				deferred.rejectWith( callbackContext, [ jqXHR, statusText, error ] );
			}

			// Status-dependent callbacks
			jqXHR.statusCode( statusCode );
			statusCode = undefined;

			if ( fireGlobals ) {
				globalEventContext.trigger( isSuccess ? "ajaxSuccess" : "ajaxError",
					[ jqXHR, s, isSuccess ? success : error ] );
			}

			// Complete
			completeDeferred.fireWith( callbackContext, [ jqXHR, statusText ] );

			if ( fireGlobals ) {
				globalEventContext.trigger( "ajaxComplete", [ jqXHR, s ] );
				// Handle the global AJAX counter
				if ( !( --jQuery.active ) ) {
					jQuery.event.trigger("ajaxStop");
				}
			}
		}

		return jqXHR;
	},

	getJSON: function( url, data, callback ) {
		return jQuery.get( url, data, callback, "json" );
	},

	getScript: function( url, callback ) {
		return jQuery.get( url, undefined, callback, "script" );
	}
});

jQuery.each( [ "get", "post" ], function( i, method ) {
	jQuery[ method ] = function( url, data, callback, type ) {
		// shift arguments if data argument was omitted
		if ( jQuery.isFunction( data ) ) {
			type = type || callback;
			callback = data;
			data = undefined;
		}

		return jQuery.ajax({
			url: url,
			type: method,
			dataType: type,
			data: data,
			success: callback
		});
	};
});

// Attach a bunch of functions for handling common AJAX events
jQuery.each( [ "ajaxStart", "ajaxStop", "ajaxComplete", "ajaxError", "ajaxSuccess", "ajaxSend" ], function( i, type ) {
	jQuery.fn[ type ] = function( fn ) {
		return this.on( type, fn );
	};
});


jQuery._evalUrl = function( url ) {
	return jQuery.ajax({
		url: url,
		type: "GET",
		dataType: "script",
		async: false,
		global: false,
		"throws": true
	});
};


jQuery.fn.extend({
	wrapAll: function( html ) {
		if ( jQuery.isFunction( html ) ) {
			return this.each(function(i) {
				jQuery(this).wrapAll( html.call(this, i) );
			});
		}

		if ( this[0] ) {
			// The elements to wrap the target around
			var wrap = jQuery( html, this[0].ownerDocument ).eq(0).clone(true);

			if ( this[0].parentNode ) {
				wrap.insertBefore( this[0] );
			}

			wrap.map(function() {
				var elem = this;

				while ( elem.firstChild && elem.firstChild.nodeType === 1 ) {
					elem = elem.firstChild;
				}

				return elem;
			}).append( this );
		}

		return this;
	},

	wrapInner: function( html ) {
		if ( jQuery.isFunction( html ) ) {
			return this.each(function(i) {
				jQuery(this).wrapInner( html.call(this, i) );
			});
		}

		return this.each(function() {
			var self = jQuery( this ),
				contents = self.contents();

			if ( contents.length ) {
				contents.wrapAll( html );

			} else {
				self.append( html );
			}
		});
	},

	wrap: function( html ) {
		var isFunction = jQuery.isFunction( html );

		return this.each(function(i) {
			jQuery( this ).wrapAll( isFunction ? html.call(this, i) : html );
		});
	},

	unwrap: function() {
		return this.parent().each(function() {
			if ( !jQuery.nodeName( this, "body" ) ) {
				jQuery( this ).replaceWith( this.childNodes );
			}
		}).end();
	}
});


jQuery.expr.filters.hidden = function( elem ) {
	// Support: Opera <= 12.12
	// Opera reports offsetWidths and offsetHeights less than zero on some elements
	return elem.offsetWidth <= 0 && elem.offsetHeight <= 0 ||
		(!support.reliableHiddenOffsets() &&
			((elem.style && elem.style.display) || jQuery.css( elem, "display" )) === "none");
};

jQuery.expr.filters.visible = function( elem ) {
	return !jQuery.expr.filters.hidden( elem );
};


var r20 = /%20/g,
	rbracket = /\[\]$/,
	rCRLF = /\r?\n/g,
	rsubmitterTypes = /^(?:submit|button|image|reset|file)$/i,
	rsubmittable = /^(?:input|select|textarea|keygen)/i;

function buildParams( prefix, obj, traditional, add ) {
	var name;

	if ( jQuery.isArray( obj ) ) {
		// Serialize array item.
		jQuery.each( obj, function( i, v ) {
			if ( traditional || rbracket.test( prefix ) ) {
				// Treat each array item as a scalar.
				add( prefix, v );

			} else {
				// Item is non-scalar (array or object), encode its numeric index.
				buildParams( prefix + "[" + ( typeof v === "object" ? i : "" ) + "]", v, traditional, add );
			}
		});

	} else if ( !traditional && jQuery.type( obj ) === "object" ) {
		// Serialize object item.
		for ( name in obj ) {
			buildParams( prefix + "[" + name + "]", obj[ name ], traditional, add );
		}

	} else {
		// Serialize scalar item.
		add( prefix, obj );
	}
}

// Serialize an array of form elements or a set of
// key/values into a query string
jQuery.param = function( a, traditional ) {
	var prefix,
		s = [],
		add = function( key, value ) {
			// If value is a function, invoke it and return its value
			value = jQuery.isFunction( value ) ? value() : ( value == null ? "" : value );
			s[ s.length ] = encodeURIComponent( key ) + "=" + encodeURIComponent( value );
		};

	// Set traditional to true for jQuery <= 1.3.2 behavior.
	if ( traditional === undefined ) {
		traditional = jQuery.ajaxSettings && jQuery.ajaxSettings.traditional;
	}

	// If an array was passed in, assume that it is an array of form elements.
	if ( jQuery.isArray( a ) || ( a.jquery && !jQuery.isPlainObject( a ) ) ) {
		// Serialize the form elements
		jQuery.each( a, function() {
			add( this.name, this.value );
		});

	} else {
		// If traditional, encode the "old" way (the way 1.3.2 or older
		// did it), otherwise encode params recursively.
		for ( prefix in a ) {
			buildParams( prefix, a[ prefix ], traditional, add );
		}
	}

	// Return the resulting serialization
	return s.join( "&" ).replace( r20, "+" );
};

jQuery.fn.extend({
	serialize: function() {
		return jQuery.param( this.serializeArray() );
	},
	serializeArray: function() {
		return this.map(function() {
			// Can add propHook for "elements" to filter or add form elements
			var elements = jQuery.prop( this, "elements" );
			return elements ? jQuery.makeArray( elements ) : this;
		})
		.filter(function() {
			var type = this.type;
			// Use .is(":disabled") so that fieldset[disabled] works
			return this.name && !jQuery( this ).is( ":disabled" ) &&
				rsubmittable.test( this.nodeName ) && !rsubmitterTypes.test( type ) &&
				( this.checked || !rcheckableType.test( type ) );
		})
		.map(function( i, elem ) {
			var val = jQuery( this ).val();

			return val == null ?
				null :
				jQuery.isArray( val ) ?
					jQuery.map( val, function( val ) {
						return { name: elem.name, value: val.replace( rCRLF, "\r\n" ) };
					}) :
					{ name: elem.name, value: val.replace( rCRLF, "\r\n" ) };
		}).get();
	}
});


// Create the request object
// (This is still attached to ajaxSettings for backward compatibility)
jQuery.ajaxSettings.xhr = window.ActiveXObject !== undefined ?
	// Support: IE6+
	function() {

		// XHR cannot access local files, always use ActiveX for that case
		return !this.isLocal &&

			// Support: IE7-8
			// oldIE XHR does not support non-RFC2616 methods (#13240)
			// See http://msdn.microsoft.com/en-us/library/ie/ms536648(v=vs.85).aspx
			// and http://www.w3.org/Protocols/rfc2616/rfc2616-sec9.html#sec9
			// Although this check for six methods instead of eight
			// since IE also does not support "trace" and "connect"
			/^(get|post|head|put|delete|options)$/i.test( this.type ) &&

			createStandardXHR() || createActiveXHR();
	} :
	// For all other browsers, use the standard XMLHttpRequest object
	createStandardXHR;

var xhrId = 0,
	xhrCallbacks = {},
	xhrSupported = jQuery.ajaxSettings.xhr();

// Support: IE<10
// Open requests must be manually aborted on unload (#5280)
if ( window.ActiveXObject ) {
	jQuery( window ).on( "unload", function() {
		for ( var key in xhrCallbacks ) {
			xhrCallbacks[ key ]( undefined, true );
		}
	});
}

// Determine support properties
support.cors = !!xhrSupported && ( "withCredentials" in xhrSupported );
xhrSupported = support.ajax = !!xhrSupported;

// Create transport if the browser can provide an xhr
if ( xhrSupported ) {

	jQuery.ajaxTransport(function( options ) {
		// Cross domain only allowed if supported through XMLHttpRequest
		if ( !options.crossDomain || support.cors ) {

			var callback;

			return {
				send: function( headers, complete ) {
					var i,
						xhr = options.xhr(),
						id = ++xhrId;

					// Open the socket
					xhr.open( options.type, options.url, options.async, options.username, options.password );

					// Apply custom fields if provided
					if ( options.xhrFields ) {
						for ( i in options.xhrFields ) {
							xhr[ i ] = options.xhrFields[ i ];
						}
					}

					// Override mime type if needed
					if ( options.mimeType && xhr.overrideMimeType ) {
						xhr.overrideMimeType( options.mimeType );
					}

					// X-Requested-With header
					// For cross-domain requests, seeing as conditions for a preflight are
					// akin to a jigsaw puzzle, we simply never set it to be sure.
					// (it can always be set on a per-request basis or even using ajaxSetup)
					// For same-domain requests, won't change header if already provided.
					if ( !options.crossDomain && !headers["X-Requested-With"] ) {
						headers["X-Requested-With"] = "XMLHttpRequest";
					}

					// Set headers
					for ( i in headers ) {
						// Support: IE<9
						// IE's ActiveXObject throws a 'Type Mismatch' exception when setting
						// request header to a null-value.
						//
						// To keep consistent with other XHR implementations, cast the value
						// to string and ignore `undefined`.
						if ( headers[ i ] !== undefined ) {
							xhr.setRequestHeader( i, headers[ i ] + "" );
						}
					}

					// Do send the request
					// This may raise an exception which is actually
					// handled in jQuery.ajax (so no try/catch here)
					xhr.send( ( options.hasContent && options.data ) || null );

					// Listener
					callback = function( _, isAbort ) {
						var status, statusText, responses;

						// Was never called and is aborted or complete
						if ( callback && ( isAbort || xhr.readyState === 4 ) ) {
							// Clean up
							delete xhrCallbacks[ id ];
							callback = undefined;
							xhr.onreadystatechange = jQuery.noop;

							// Abort manually if needed
							if ( isAbort ) {
								if ( xhr.readyState !== 4 ) {
									xhr.abort();
								}
							} else {
								responses = {};
								status = xhr.status;

								// Support: IE<10
								// Accessing binary-data responseText throws an exception
								// (#11426)
								if ( typeof xhr.responseText === "string" ) {
									responses.text = xhr.responseText;
								}

								// Firefox throws an exception when accessing
								// statusText for faulty cross-domain requests
								try {
									statusText = xhr.statusText;
								} catch( e ) {
									// We normalize with Webkit giving an empty statusText
									statusText = "";
								}

								// Filter status for non standard behaviors

								// If the request is local and we have data: assume a success
								// (success with no data won't get notified, that's the best we
								// can do given current implementations)
								if ( !status && options.isLocal && !options.crossDomain ) {
									status = responses.text ? 200 : 404;
								// IE - #1450: sometimes returns 1223 when it should be 204
								} else if ( status === 1223 ) {
									status = 204;
								}
							}
						}

						// Call complete if needed
						if ( responses ) {
							complete( status, statusText, responses, xhr.getAllResponseHeaders() );
						}
					};

					if ( !options.async ) {
						// if we're in sync mode we fire the callback
						callback();
					} else if ( xhr.readyState === 4 ) {
						// (IE6 & IE7) if it's in cache and has been
						// retrieved directly we need to fire the callback
						setTimeout( callback );
					} else {
						// Add to the list of active xhr callbacks
						xhr.onreadystatechange = xhrCallbacks[ id ] = callback;
					}
				},

				abort: function() {
					if ( callback ) {
						callback( undefined, true );
					}
				}
			};
		}
	});
}

// Functions to create xhrs
function createStandardXHR() {
	try {
		return new window.XMLHttpRequest();
	} catch( e ) {}
}

function createActiveXHR() {
	try {
		return new window.ActiveXObject( "Microsoft.XMLHTTP" );
	} catch( e ) {}
}


// Install script dataType
jQuery.ajaxSetup({
	accepts: {
		script: "text/javascript, application/javascript, application/ecmascript, application/x-ecmascript"
	},
	contents: {
		script: /(?:java|ecma)script/
	},
	converters: {
		"text script": function( text ) {
			jQuery.globalEval( text );
			return text;
		}
	}
});

// Handle cache's special case and global
jQuery.ajaxPrefilter( "script", function( s ) {
	if ( s.cache === undefined ) {
		s.cache = false;
	}
	if ( s.crossDomain ) {
		s.type = "GET";
		s.global = false;
	}
});

// Bind script tag hack transport
jQuery.ajaxTransport( "script", function(s) {

	// This transport only deals with cross domain requests
	if ( s.crossDomain ) {

		var script,
			head = document.head || jQuery("head")[0] || document.documentElement;

		return {

			send: function( _, callback ) {

				script = document.createElement("script");

				script.async = true;

				if ( s.scriptCharset ) {
					script.charset = s.scriptCharset;
				}

				script.src = s.url;

				// Attach handlers for all browsers
				script.onload = script.onreadystatechange = function( _, isAbort ) {

					if ( isAbort || !script.readyState || /loaded|complete/.test( script.readyState ) ) {

						// Handle memory leak in IE
						script.onload = script.onreadystatechange = null;

						// Remove the script
						if ( script.parentNode ) {
							script.parentNode.removeChild( script );
						}

						// Dereference the script
						script = null;

						// Callback if not abort
						if ( !isAbort ) {
							callback( 200, "success" );
						}
					}
				};

				// Circumvent IE6 bugs with base elements (#2709 and #4378) by prepending
				// Use native DOM manipulation to avoid our domManip AJAX trickery
				head.insertBefore( script, head.firstChild );
			},

			abort: function() {
				if ( script ) {
					script.onload( undefined, true );
				}
			}
		};
	}
});


var oldCallbacks = [],
	rjsonp = /(=)\?(?=&|$)|\?\?/;

// Default jsonp settings
jQuery.ajaxSetup({
	jsonp: "callback",
	jsonpCallback: function() {
		var callback = oldCallbacks.pop() || ( jQuery.expando + "_" + ( nonce++ ) );
		this[ callback ] = true;
		return callback;
	}
});

// Detect, normalize options and install callbacks for jsonp requests
jQuery.ajaxPrefilter( "json jsonp", function( s, originalSettings, jqXHR ) {

	var callbackName, overwritten, responseContainer,
		jsonProp = s.jsonp !== false && ( rjsonp.test( s.url ) ?
			"url" :
			typeof s.data === "string" && !( s.contentType || "" ).indexOf("application/x-www-form-urlencoded") && rjsonp.test( s.data ) && "data"
		);

	// Handle iff the expected data type is "jsonp" or we have a parameter to set
	if ( jsonProp || s.dataTypes[ 0 ] === "jsonp" ) {

		// Get callback name, remembering preexisting value associated with it
		callbackName = s.jsonpCallback = jQuery.isFunction( s.jsonpCallback ) ?
			s.jsonpCallback() :
			s.jsonpCallback;

		// Insert callback into url or form data
		if ( jsonProp ) {
			s[ jsonProp ] = s[ jsonProp ].replace( rjsonp, "$1" + callbackName );
		} else if ( s.jsonp !== false ) {
			s.url += ( rquery.test( s.url ) ? "&" : "?" ) + s.jsonp + "=" + callbackName;
		}

		// Use data converter to retrieve json after script execution
		s.converters["script json"] = function() {
			if ( !responseContainer ) {
				jQuery.error( callbackName + " was not called" );
			}
			return responseContainer[ 0 ];
		};

		// force json dataType
		s.dataTypes[ 0 ] = "json";

		// Install callback
		overwritten = window[ callbackName ];
		window[ callbackName ] = function() {
			responseContainer = arguments;
		};

		// Clean-up function (fires after converters)
		jqXHR.always(function() {
			// Restore preexisting value
			window[ callbackName ] = overwritten;

			// Save back as free
			if ( s[ callbackName ] ) {
				// make sure that re-using the options doesn't screw things around
				s.jsonpCallback = originalSettings.jsonpCallback;

				// save the callback name for future use
				oldCallbacks.push( callbackName );
			}

			// Call if it was a function and we have a response
			if ( responseContainer && jQuery.isFunction( overwritten ) ) {
				overwritten( responseContainer[ 0 ] );
			}

			responseContainer = overwritten = undefined;
		});

		// Delegate to script
		return "script";
	}
});


// data: string of html
// context (optional): If specified, the fragment will be created in this context, defaults to document
// keepScripts (optional): If true, will include scripts passed in the html string
jQuery.parseHTML = function( data, context, keepScripts ) {
	if ( !data || typeof data !== "string" ) {
		return null;
	}
	if ( typeof context === "boolean" ) {
		keepScripts = context;
		context = false;
	}
	context = context || document;

	var parsed = rsingleTag.exec( data ),
		scripts = !keepScripts && [];

	// Single tag
	if ( parsed ) {
		return [ context.createElement( parsed[1] ) ];
	}

	parsed = jQuery.buildFragment( [ data ], context, scripts );

	if ( scripts && scripts.length ) {
		jQuery( scripts ).remove();
	}

	return jQuery.merge( [], parsed.childNodes );
};


// Keep a copy of the old load method
var _load = jQuery.fn.load;

/**
 * Load a url into a page
 */
jQuery.fn.load = function( url, params, callback ) {
	if ( typeof url !== "string" && _load ) {
		return _load.apply( this, arguments );
	}

	var selector, response, type,
		self = this,
		off = url.indexOf(" ");

	if ( off >= 0 ) {
		selector = jQuery.trim( url.slice( off, url.length ) );
		url = url.slice( 0, off );
	}

	// If it's a function
	if ( jQuery.isFunction( params ) ) {

		// We assume that it's the callback
		callback = params;
		params = undefined;

	// Otherwise, build a param string
	} else if ( params && typeof params === "object" ) {
		type = "POST";
	}

	// If we have elements to modify, make the request
	if ( self.length > 0 ) {
		jQuery.ajax({
			url: url,

			// if "type" variable is undefined, then "GET" method will be used
			type: type,
			dataType: "html",
			data: params
		}).done(function( responseText ) {

			// Save response for use in complete callback
			response = arguments;

			self.html( selector ?

				// If a selector was specified, locate the right elements in a dummy div
				// Exclude scripts to avoid IE 'Permission Denied' errors
				jQuery("<div>").append( jQuery.parseHTML( responseText ) ).find( selector ) :

				// Otherwise use the full result
				responseText );

		}).complete( callback && function( jqXHR, status ) {
			self.each( callback, response || [ jqXHR.responseText, status, jqXHR ] );
		});
	}

	return this;
};


jQuery.expr.filters.animated = function( elem ) {
	return jQuery.grep(jQuery.timers, function( fn ) {
		return elem === fn.elem;
	}).length;
};


var docElem = window.document.documentElement;

/**
 * Gets a window from an element
 */
function getWindow( elem ) {
	return jQuery.isWindow( elem ) ?
		elem :
		elem.nodeType === 9 ?
			elem.defaultView || elem.parentWindow :
			false;
}

jQuery.offset = {
	setOffset: function( elem, options, i ) {
		var curPosition, curLeft, curCSSTop, curTop, curOffset, curCSSLeft, calculatePosition,
			position = jQuery.css( elem, "position" ),
			curElem = jQuery( elem ),
			props = {};

		// set position first, in-case top/left are set even on static elem
		if ( position === "static" ) {
			elem.style.position = "relative";
		}

		curOffset = curElem.offset();
		curCSSTop = jQuery.css( elem, "top" );
		curCSSLeft = jQuery.css( elem, "left" );
		calculatePosition = ( position === "absolute" || position === "fixed" ) &&
			jQuery.inArray("auto", [ curCSSTop, curCSSLeft ] ) > -1;

		// need to be able to calculate position if either top or left is auto and position is either absolute or fixed
		if ( calculatePosition ) {
			curPosition = curElem.position();
			curTop = curPosition.top;
			curLeft = curPosition.left;
		} else {
			curTop = parseFloat( curCSSTop ) || 0;
			curLeft = parseFloat( curCSSLeft ) || 0;
		}

		if ( jQuery.isFunction( options ) ) {
			options = options.call( elem, i, curOffset );
		}

		if ( options.top != null ) {
			props.top = ( options.top - curOffset.top ) + curTop;
		}
		if ( options.left != null ) {
			props.left = ( options.left - curOffset.left ) + curLeft;
		}

		if ( "using" in options ) {
			options.using.call( elem, props );
		} else {
			curElem.css( props );
		}
	}
};

jQuery.fn.extend({
	offset: function( options ) {
		if ( arguments.length ) {
			return options === undefined ?
				this :
				this.each(function( i ) {
					jQuery.offset.setOffset( this, options, i );
				});
		}

		var docElem, win,
			box = { top: 0, left: 0 },
			elem = this[ 0 ],
			doc = elem && elem.ownerDocument;

		if ( !doc ) {
			return;
		}

		docElem = doc.documentElement;

		// Make sure it's not a disconnected DOM node
		if ( !jQuery.contains( docElem, elem ) ) {
			return box;
		}

		// If we don't have gBCR, just use 0,0 rather than error
		// BlackBerry 5, iOS 3 (original iPhone)
		if ( typeof elem.getBoundingClientRect !== strundefined ) {
			box = elem.getBoundingClientRect();
		}
		win = getWindow( doc );
		return {
			top: box.top  + ( win.pageYOffset || docElem.scrollTop )  - ( docElem.clientTop  || 0 ),
			left: box.left + ( win.pageXOffset || docElem.scrollLeft ) - ( docElem.clientLeft || 0 )
		};
	},

	position: function() {
		if ( !this[ 0 ] ) {
			return;
		}

		var offsetParent, offset,
			parentOffset = { top: 0, left: 0 },
			elem = this[ 0 ];

		// fixed elements are offset from window (parentOffset = {top:0, left: 0}, because it is its only offset parent
		if ( jQuery.css( elem, "position" ) === "fixed" ) {
			// we assume that getBoundingClientRect is available when computed position is fixed
			offset = elem.getBoundingClientRect();
		} else {
			// Get *real* offsetParent
			offsetParent = this.offsetParent();

			// Get correct offsets
			offset = this.offset();
			if ( !jQuery.nodeName( offsetParent[ 0 ], "html" ) ) {
				parentOffset = offsetParent.offset();
			}

			// Add offsetParent borders
			parentOffset.top  += jQuery.css( offsetParent[ 0 ], "borderTopWidth", true );
			parentOffset.left += jQuery.css( offsetParent[ 0 ], "borderLeftWidth", true );
		}

		// Subtract parent offsets and element margins
		// note: when an element has margin: auto the offsetLeft and marginLeft
		// are the same in Safari causing offset.left to incorrectly be 0
		return {
			top:  offset.top  - parentOffset.top - jQuery.css( elem, "marginTop", true ),
			left: offset.left - parentOffset.left - jQuery.css( elem, "marginLeft", true)
		};
	},

	offsetParent: function() {
		return this.map(function() {
			var offsetParent = this.offsetParent || docElem;

			while ( offsetParent && ( !jQuery.nodeName( offsetParent, "html" ) && jQuery.css( offsetParent, "position" ) === "static" ) ) {
				offsetParent = offsetParent.offsetParent;
			}
			return offsetParent || docElem;
		});
	}
});

// Create scrollLeft and scrollTop methods
jQuery.each( { scrollLeft: "pageXOffset", scrollTop: "pageYOffset" }, function( method, prop ) {
	var top = /Y/.test( prop );

	jQuery.fn[ method ] = function( val ) {
		return access( this, function( elem, method, val ) {
			var win = getWindow( elem );

			if ( val === undefined ) {
				return win ? (prop in win) ? win[ prop ] :
					win.document.documentElement[ method ] :
					elem[ method ];
			}

			if ( win ) {
				win.scrollTo(
					!top ? val : jQuery( win ).scrollLeft(),
					top ? val : jQuery( win ).scrollTop()
				);

			} else {
				elem[ method ] = val;
			}
		}, method, val, arguments.length, null );
	};
});

// Add the top/left cssHooks using jQuery.fn.position
// Webkit bug: https://bugs.webkit.org/show_bug.cgi?id=29084
// getComputedStyle returns percent when specified for top/left/bottom/right
// rather than make the css module depend on the offset module, we just check for it here
jQuery.each( [ "top", "left" ], function( i, prop ) {
	jQuery.cssHooks[ prop ] = addGetHookIf( support.pixelPosition,
		function( elem, computed ) {
			if ( computed ) {
				computed = curCSS( elem, prop );
				// if curCSS returns percentage, fallback to offset
				return rnumnonpx.test( computed ) ?
					jQuery( elem ).position()[ prop ] + "px" :
					computed;
			}
		}
	);
});


// Create innerHeight, innerWidth, height, width, outerHeight and outerWidth methods
jQuery.each( { Height: "height", Width: "width" }, function( name, type ) {
	jQuery.each( { padding: "inner" + name, content: type, "": "outer" + name }, function( defaultExtra, funcName ) {
		// margin is only for outerHeight, outerWidth
		jQuery.fn[ funcName ] = function( margin, value ) {
			var chainable = arguments.length && ( defaultExtra || typeof margin !== "boolean" ),
				extra = defaultExtra || ( margin === true || value === true ? "margin" : "border" );

			return access( this, function( elem, type, value ) {
				var doc;

				if ( jQuery.isWindow( elem ) ) {
					// As of 5/8/2012 this will yield incorrect results for Mobile Safari, but there
					// isn't a whole lot we can do. See pull request at this URL for discussion:
					// https://github.com/jquery/jquery/pull/764
					return elem.document.documentElement[ "client" + name ];
				}

				// Get document width or height
				if ( elem.nodeType === 9 ) {
					doc = elem.documentElement;

					// Either scroll[Width/Height] or offset[Width/Height] or client[Width/Height], whichever is greatest
					// unfortunately, this causes bug #3838 in IE6/8 only, but there is currently no good, small way to fix it.
					return Math.max(
						elem.body[ "scroll" + name ], doc[ "scroll" + name ],
						elem.body[ "offset" + name ], doc[ "offset" + name ],
						doc[ "client" + name ]
					);
				}

				return value === undefined ?
					// Get width or height on the element, requesting but not forcing parseFloat
					jQuery.css( elem, type, extra ) :

					// Set width or height on the element
					jQuery.style( elem, type, value, extra );
			}, type, chainable ? margin : undefined, chainable, null );
		};
	});
});


// The number of elements contained in the matched element set
jQuery.fn.size = function() {
	return this.length;
};

jQuery.fn.andSelf = jQuery.fn.addBack;


// Register as a named AMD module, since jQuery can be concatenated with other
// files that may use define, but not via a proper concatenation script that
// understands anonymous AMD modules. A named AMD is safest and most robust
// way to register. Lowercase jquery is used because AMD module names are
// derived from file names, and jQuery is normally delivered in a lowercase
// file name. Do this after creating the global so that if an AMD module wants
// to call noConflict to hide this version of jQuery, it will work.

// Note that for maximum portability, libraries that are not jQuery should
// declare themselves as anonymous modules, and avoid setting a global if an
// AMD loader is present. jQuery is a special case. For more information, see
// https://github.com/jrburke/requirejs/wiki/Updating-existing-libraries#wiki-anon

if ( typeof define === "function" && define.amd ) {
	define( "jquery", [], function() {
		return jQuery;
	});
}


var
	// Map over jQuery in case of overwrite
	_jQuery = window.jQuery,

	// Map over the $ in case of overwrite
	_$ = window.$;

jQuery.noConflict = function( deep ) {
	if ( window.$ === jQuery ) {
		window.$ = _$;
	}

	if ( deep && window.jQuery === jQuery ) {
		window.jQuery = _jQuery;
	}

	return jQuery;
};

// Expose jQuery and $ identifiers, even in
// AMD (#7102#comment:10, https://github.com/jquery/jquery/pull/557)
// and CommonJS for browser emulators (#13566)
if ( typeof noGlobal === strundefined ) {
	window.jQuery = window.$ = jQuery;
}


return jQuery;

}));
```
