## Supplementary material for "ΔSCOPE: A new method to quantify 3D biological structures and identify differences in zebrafish forebrain development": An archived copy of the DeltaSCOPE code repository.: jquery.js

```
/*! jQuery v3.1.0 | (c) jQuery Foundation | jquery.org/license */
!function(a,b){"use strict";"object"==typeof module&&"object"==typeof module.exports?module.exports=a.document?b(a,!0):function(a){if(!a.document)throw new Error("jQuery requires a window with a document");return b(a)}:b(a)}("undefined"!=typeof window?window:this,function(a,b){"use strict";var c=[],d=a.document,e=Object.getPrototypeOf,f=c.slice,g=c.concat,h=c.push,i=c.indexOf,j={},k=j.toString,l=j.hasOwnProperty,m=l.toString,n=m.call(Object),o={};function p(a,b){b=b||d;var c=b.createElement("script");c.text=a,b.head.appendChild(c).parentNode.removeChild(c)}var q="3.1.0",r=function(a,b){return new r.fn.init(a,b)},s=/^[\s\uFEFF\xA0]+|[\s\uFEFF\xA0]+$/g,t=/^-ms-/,u=/-([a-z])/g,v=function(a,b){return b.toUpperCase()};r.fn=r.prototype={jquery:q,constructor:r,length:0,toArray:function(){return f.call(this)},get:function(a){return null!=a?a<0?this[a+this.length]:this[a]:f.call(this)},pushStack:function(a){var b=r.merge(this.constructor(),a);return b.prevObject=this,b},each:function(a){return r.each(this,a)},map:function(a){return this.pushStack(r.map(this,function(b,c){return a.call(b,c,b)}))},slice:function(){return this.pushStack(f.apply(this,arguments))},first:function(){return this.eq(0)},last:function(){return this.eq(-1)},eq:function(a){var b=this.length,c=+a+(a<0?b:0);return this.pushStack(c>=0&&c<b?[this[c]]:[])},end:function(){return this.prevObject||this.constructor()},push:h,sort:c.sort,splice:c.splice},r.extend=r.fn.extend=function(){var a,b,c,d,e,f,g=arguments[0]||{},h=1,i=arguments.length,j=!1;for("boolean"==typeof g&&(j=g,g=arguments[h]||{},h++),"object"==typeof g||r.isFunction(g)||(g={}),h===i&&(g=this,h--);h<i;h++)if(null!=(a=arguments[h]))for(b in a)c=g[b],d=a[b],g!==d&&(j&&d&&(r.isPlainObject(d)||(e=r.isArray(d)))?(e?(e=!1,f=c&&r.isArray(c)?c:[]):f=c&&r.isPlainObject(c)?c:{},g[b]=r.extend(j,f,d)):void 0!==d&&(g[b]=d));return g},r.extend({expando:"jQuery"+(q+Math.random()).replace(/\D/g,""),isReady:!0,error:function(a){throw new Error(a)},noop:function(){},isFunction:function(a){return"function"===r.type(a)},isArray:Array.isArray,isWindow:function(a){return null!=a&&a===a.window},isNumeric:function(a){var b=r.type(a);return("number"===b||"string"===b)&&!isNaN(a-parseFloat(a))},isPlainObject:function(a){var b,c;return!(!a||"[object Object]"!==k.call(a))&&(!(b=e(a))||(c=l.call(b,"constructor")&&b.constructor,"function"==typeof c&&m.call(c)===n))},isEmptyObject:function(a){var b;for(b in a)return!1;return!0},type:function(a){return null==a?a+"":"object"==typeof a||"function"==typeof a?j[k.call(a)]||"object":typeof a},globalEval:function(a){p(a)},camelCase:function(a){return a.replace(t,"ms-").replace(u,v)},nodeName:function(a,b){return a.nodeName&&a.nodeName.toLowerCase()===b.toLowerCase()},each:function(a,b){var c,d=0;if(w(a)){for(c=a.length;d<c;d++)if(b.call(a[d],d,a[d])===!1)break}else for(d in a)if(b.call(a[d],d,a[d])===!1)break;return a},trim:function(a){return null==a?"":(a+"").replace(s,"")},makeArray:function(a,b){var c=b||[];return null!=a&&(w(Object(a))?r.merge(c,"string"==typeof a?[a]:a):h.call(c,a)),c},inArray:function(a,b,c){return null==b?-1:i.call(b,a,c)},merge:function(a,b){for(var c=+b.length,d=0,e=a.length;d<c;d++)a[e++]=b[d];return a.length=e,a},grep:function(a,b,c){for(var d,e=[],f=0,g=a.length,h=!c;f<g;f++)d=!b(a[f],f),d!==h&&e.push(a[f]);return e},map:function(a,b,c){var d,e,f=0,h=[];if(w(a))for(d=a.length;f<d;f++)e=b(a[f],f,c),null!=e&&h.push(e);else for(f in a)e=b(a[f],f,c),null!=e&&h.push(e);return g.apply([],h)},guid:1,proxy:function(a,b){var c,d,e;if("string"==typeof b&&(c=a[b],b=a,a=c),r.isFunction(a))return d=f.call(arguments,2),e=function(){return a.apply(b||this,d.concat(f.call(arguments)))},e.guid=a.guid=a.guid||r.guid++,e},now:Date.now,support:o}),"function"==typeof Symbol&&(r.fn[Symbol.iterator]=c[Symbol.iterator]),r.each("Boolean Number String Function Array Date RegExp Object Error Symbol".split(" "),function(a,b){j["[object "+b+"]"]=b.toLowerCase()});function w(a){var b=!!a&&"length"in a&&a.length,c=r.type(a);return"function"!==c&&!r.isWindow(a)&&("array"===c||0===b||"number"==typeof b&&b>0&&b-1 in a)}var x=function(a){var b,c,d,e,f,g,h,i,j,k,l,m,n,o,p,q,r,s,t,u="sizzle"+1*new Date,v=a.document,w=0,x=0,y=ha(),z=ha(),A=ha(),B=function(a,b){return a===b&&(l=!0),0},C={}.hasOwnProperty,D=[],E=D.pop,F=D.push,G=D.push,H=D.slice,I=function(a,b){for(var c=0,d=a.length;c<d;c++)if(a[c]===b)return c;return-1},J="checked|selected|async|autofocus|autoplay|controls|defer|disabled|hidden|ismap|loop|multiple|open|readonly|required|scoped",K="[\\x20\\t\\r\\n\\f]",L="(?:\\\\.|[\\w-]|[^\0-\\xa0])+",M="\\["+K+"*("+L+")(?:"+K+"*([*^$|!~]?=)"+K+"*(?:'((?:\\\\.|[^\\\\'])*)'|\"((?:\\\\.|[^\\\\\"])*)\"|("+L+"))|)"+K+"*\\]",N=":("+L+")(?:\\((('((?:\\\\.|[^\\\\'])*)'|\"((?:\\\\.|[^\\\\\"])*)\")|((?:\\\\.|[^\\\\()[\\]]|"+M+")*)|.*)\\)|)",O=new RegExp(K+"+","g"),P=new RegExp("^"+K+"+|((?:^|[^\\\\])(?:\\\\.)*)"+K+"+$","g"),Q=new RegExp("^"+K+"*,"+K+"*"),R=new RegExp("^"+K+"*([>+~]|"+K+")"+K+"*"),S=new RegExp("="+K+"*([^\\]'\"]*?)"+K+"*\\]","g"),T=new RegExp(N),U=new RegExp("^"+L+"$"),V={ID:new RegExp("^#("+L+")"),CLASS:new RegExp("^\\.("+L+")"),TAG:new RegExp("^("+L+"|[*])"),ATTR:new RegExp("^"+M),PSEUDO:new RegExp("^"+N),CHILD:new RegExp("^:(only|first|last|nth|nth-last)-(child|of-type)(?:\\("+K+"*(even|odd|(([+-]|)(\\d*)n|)"+K+"*(?:([+-]|)"+K+"*(\\d+)|))"+K+"*\\)|)","i"),bool:new RegExp("^(?:"+J+")$","i"),needsContext:new RegExp("^"+K+"*[>+~]|:(even|odd|eq|gt|lt|nth|first|last)(?:\\("+K+"*((?:-\\d)?\\d*)"+K+"*\\)|)(?=[^-]|$)","i")},W=/^(?:input|select|textarea|button)$/i,X=/^h\d$/i,Y=/^[^{]+\{\s*\[native \w/,Z=/^(?:#([\w-]+)|(\w+)|\.([\w-]+))$/,$=/[+~]/,_=new RegExp("\\\\([\\da-f]{1,6}"+K+"?|("+K+")|.)","ig"),aa=function(a,b,c){var d="0x"+b-65536;return d!==d||c?b:d<0?String.fromCharCode(d+65536):String.fromCharCode(d>>10|55296,1023&d|56320)},ba=/([\0-\x1f\x7f]|^-?\d)|^-$|[^\x80-\uFFFF\w-]/g,ca=function(a,b){return b?"\0"===a?"\ufffd":a.slice(0,-1)+"\\"+a.charCodeAt(a.length-1).toString(16)+" ":"\\"+a},da=function(){m()},ea=ta(function(a){return a.disabled===!0},{dir:"parentNode",next:"legend"});try{G.apply(D=H.call(v.childNodes),v.childNodes),D[v.childNodes.length].nodeType}catch(fa){G={apply:D.length?function(a,b){F.apply(a,H.call(b))}:function(a,b){var c=a.length,d=0;while(a[c++]=b[d++]);a.length=c-1}}}function ga(a,b,d,e){var f,h,j,k,l,o,r,s=b&&b.ownerDocument,w=b?b.nodeType:9;if(d=d||[],"string"!=typeof a||!a||1!==w&&9!==w&&11!==w)return d;if(!e&&((b?b.ownerDocument||b:v)!==n&&m(b),b=b||n,p)){if(11!==w&&(l=Z.exec(a)))if(f=l[1]){if(9===w){if(!(j=b.getElementById(f)))return d;if(j.id===f)return d.push(j),d}else if(s&&(j=s.getElementById(f))&&t(b,j)&&j.id===f)return d.push(j),d}else{if(l[2])return G.apply(d,b.getElementsByTagName(a)),d;if((f=l[3])&&c.getElementsByClassName&&b.getElementsByClassName)return G.apply(d,b.getElementsByClassName(f)),d}if(c.qsa&&!A[a+" "]&&(!q||!q.test(a))){if(1!==w)s=b,r=a;else if("object"!==b.nodeName.toLowerCase()){(k=b.getAttribute("id"))?k=k.replace(ba,ca):b.setAttribute("id",k=u),o=g(a),h=o.length;while(h--)o[h]="#"+k+" "+sa(o[h]);r=o.join(","),s=$.test(a)&&qa(b.parentNode)||b}if(r)try{return G.apply(d,s.querySelectorAll(r)),d}catch(x){}finally{k===u&&b.removeAttribute("id")}}}return i(a.replace(P,"$1"),b,d,e)}function ha(){var a=[];function b(c,e){return a.push(c+" ")>d.cacheLength&&delete b[a.shift()],b[c+" "]=e}return b}function ia(a){return a[u]=!0,a}function ja(a){var b=n.createElement("fieldset");try{return!!a(b)}catch(c){return!1}finally{b.parentNode&&b.parentNode.removeChild(b),b=null}}function ka(a,b){var c=a.split("|"),e=c.length;while(e--)d.attrHandle[c[e]]=b}function la(a,b){var c=b&&a,d=c&&1===a.nodeType&&1===b.nodeType&&a.sourceIndex-b.sourceIndex;if(d)return d;if(c)while(c=c.nextSibling)if(c===b)return-1;return a?1:-1}function ma(a){return function(b){var c=b.nodeName.toLowerCase();return"input"===c&&b.type===a}}function na(a){return function(b){var c=b.nodeName.toLowerCase();return("input"===c||"button"===c)&&b.type===a}}function oa(a){return function(b){return"label"in b&&b.disabled===a||"form"in b&&b.disabled===a||"form"in b&&b.disabled===!1&&(b.isDisabled===a||b.isDisabled!==!a&&("label"in b||!ea(b))!==a)}}function pa(a){return ia(function(b){return b=+b,ia(function(c,d){var e,f=a([],c.length,b),g=f.length;while(g--)c[e=f[g]]&&(c[e]=!(d[e]=c[e]))})})}function qa(a){return a&&"undefined"!=typeof a.getElementsByTagName&&a}c=ga.support={},f=ga.isXML=function(a){var b=a&&(a.ownerDocument||a).documentElement;return!!b&&"HTML"!==b.nodeName},m=ga.setDocument=function(a){var b,e,g=a?a.ownerDocument||a:v;return g!==n&&9===g.nodeType&&g.documentElement?(n=g,o=n.documentElement,p=!f(n),v!==n&&(e=n.defaultView)&&e.top!==e&&(e.addEventListener?e.addEventListener("unload",da,!1):e.attachEvent&&e.attachEvent("onunload",da)),c.attributes=ja(function(a){return a.className="i",!a.getAttribute("className")}),c.getElementsByTagName=ja(function(a){return a.appendChild(n.createComment("")),!a.getElementsByTagName("*").length}),c.getElementsByClassName=Y.test(n.getElementsByClassName),c.getById=ja(function(a){return o.appendChild(a).id=u,!n.getElementsByName||!n.getElementsByName(u).length}),c.getById?(d.find.ID=function(a,b){if("undefined"!=typeof b.getElementById&&p){var c=b.getElementById(a);return c?[c]:[]}},d.filter.ID=function(a){var b=a.replace(_,aa);return function(a){return a.getAttribute("id")===b}}):(delete d.find.ID,d.filter.ID=function(a){var b=a.replace(_,aa);return function(a){var c="undefined"!=typeof a.getAttributeNode&&a.getAttributeNode("id");return c&&c.value===b}}),d.find.TAG=c.getElementsByTagName?function(a,b){return"undefined"!=typeof b.getElementsByTagName?b.getElementsByTagName(a):c.qsa?b.querySelectorAll(a):void 0}:function(a,b){var c,d=[],e=0,f=b.getElementsByTagName(a);if("*"===a){while(c=f[e++])1===c.nodeType&&d.push(c);return d}return f},d.find.CLASS=c.getElementsByClassName&&function(a,b){if("undefined"!=typeof b.getElementsByClassName&&p)return b.getElementsByClassName(a)},r=[],q=[],(c.qsa=Y.test(n.querySelectorAll))&&(ja(function(a){o.appendChild(a).innerHTML="<a id='"+u+"'></a><select id='"+u+"-\r\\' msallowcapture=''><option selected=''></option></select>",a.querySelectorAll("[msallowcapture^='']").length&&q.push("[*^$]="+K+"*(?:''|\"\")"),a.querySelectorAll("[selected]").length||q.push("\\["+K+"*(?:value|"+J+")"),a.querySelectorAll("[id~="+u+"-]").length||q.push("~="),a.querySelectorAll(":checked").length||q.push(":checked"),a.querySelectorAll("a#"+u+"+*").length||q.push(".#.+[+~]")}),ja(function(a){a.innerHTML="<a href='' disabled='disabled'></a><select disabled='disabled'><option/></select>";var b=n.createElement("input");b.setAttribute("type","hidden"),a.appendChild(b).setAttribute("name","D"),a.querySelectorAll("[name=d]").length&&q.push("name"+K+"*[*^$|!~]?="),2!==a.querySelectorAll(":enabled").length&&q.push(":enabled",":disabled"),o.appendChild(a).disabled=!0,2!==a.querySelectorAll(":disabled").length&&q.push(":enabled",":disabled"),a.querySelectorAll("*,:x"),q.push(",.*:")})),(c.matchesSelector=Y.test(s=o.matches||o.webkitMatchesSelector||o.mozMatchesSelector||o.oMatchesSelector||o.msMatchesSelector))&&ja(function(a){c.disconnectedMatch=s.call(a,"*"),s.call(a,"[s!='']:x"),r.push("!=",N)}),q=q.length&&new RegExp(q.join("|")),r=r.length&&new RegExp(r.join("|")),b=Y.test(o.compareDocumentPosition),t=b||Y.test(o.contains)?function(a,b){var c=9===a.nodeType?a.documentElement:a,d=b&&b.parentNode;return a===d||!(!d||1!==d.nodeType||!(c.contains?c.contains(d):a.compareDocumentPosition&&16&a.compareDocumentPosition(d)))}:function(a,b){if(b)while(b=b.parentNode)if(b===a)return!0;return!1},B=b?function(a,b){if(a===b)return l=!0,0;var d=!a.compareDocumentPosition-!b.compareDocumentPosition;return d?d:(d=(a.ownerDocument||a)===(b.ownerDocument||b)?a.compareDocumentPosition(b):1,1&d||!c.sortDetached&&b.compareDocumentPosition(a)===d?a===n||a.ownerDocument===v&&t(v,a)?-1:b===n||b.ownerDocument===v&&t(v,b)?1:k?I(k,a)-I(k,b):0:4&d?-1:1)}:function(a,b){if(a===b)return l=!0,0;var c,d=0,e=a.parentNode,f=b.parentNode,g=[a],h=[b];if(!e||!f)return a===n?-1:b===n?1:e?-1:f?1:k?I(k,a)-I(k,b):0;if(e===f)return la(a,b);c=a;while(c=c.parentNode)g.unshift(c);c=b;while(c=c.parentNode)h.unshift(c);while(g[d]===h[d])d++;return d?la(g[d],h[d]):g[d]===v?-1:h[d]===v?1:0},n):n},ga.matches=function(a,b){return ga(a,null,null,b)},ga.matchesSelector=function(a,b){if((a.ownerDocument||a)!==n&&m(a),b=b.replace(S,"='$1']"),c.matchesSelector&&p&&!A[b+" "]&&(!r||!r.test(b))&&(!q||!q.test(b)))try{var d=s.call(a,b);if(d||c.disconnectedMatch||a.document&&11!==a.document.nodeType)return d}catch(e){}return ga(b,n,null,[a]).length>0},ga.contains=function(a,b){return(a.ownerDocument||a)!==n&&m(a),t(a,b)},ga.attr=function(a,b){(a.ownerDocument||a)!==n&&m(a);var e=d.attrHandle[b.toLowerCase()],f=e&&C.call(d.attrHandle,b.toLowerCase())?e(a,b,!p):void 0;return void 0!==f?f:c.attributes||!p?a.getAttribute(b):(f=a.getAttributeNode(b))&&f.specified?f.value:null},ga.escape=function(a){return(a+"").replace(ba,ca)},ga.error=function(a){throw new Error("Syntax error, unrecognized expression: "+a)},ga.uniqueSort=function(a){var b,d=[],e=0,f=0;if(l=!c.detectDuplicates,k=!c.sortStable&&a.slice(0),a.sort(B),l){while(b=a[f++])b===a[f]&&(e=d.push(f));while(e--)a.splice(d[e],1)}return k=null,a},e=ga.getText=function(a){var b,c="",d=0,f=a.nodeType;if(f){if(1===f||9===f||11===f){if("string"==typeof a.textContent)return a.textContent;for(a=a.firstChild;a;a=a.nextSibling)c+=e(a)}else if(3===f||4===f)return a.nodeValue}else while(b=a[d++])c+=e(b);return c},d=ga.selectors={cacheLength:50,createPseudo:ia,match:V,attrHandle:{},find:{},relative:{">":{dir:"parentNode",first:!0}," ":{dir:"parentNode"},"+":{dir:"previousSibling",first:!0},"~":{dir:"previousSibling"}},preFilter:{ATTR:function(a){return a[1]=a[1].replace(_,aa),a[3]=(a[3]||a[4]||a[5]||"").replace(_,aa),"~="===a[2]&&(a[3]=" "+a[3]+" "),a.slice(0,4)},CHILD:function(a){return a[1]=a[1].toLowerCase(),"nth"===a[1].slice(0,3)?(a[3]||ga.error(a[0]),a[4]=+(a[4]?a[5]+(a[6]||1):2*("even"===a[3]||"odd"===a[3])),a[5]=+(a[7]+a[8]||"odd"===a[3])):a[3]&&ga.error(a[0]),a},PSEUDO:function(a){var b,c=!a[6]&&a[2];return V.CHILD.test(a[0])?null:(a[3]?a[2]=a[4]||a[5]||"":c&&T.test(c)&&(b=g(c,!0))&&(b=c.indexOf(")",c.length-b)-c.length)&&(a[0]=a[0].slice(0,b),a[2]=c.slice(0,b)),a.slice(0,3))}},filter:{TAG:function(a){var b=a.replace(_,aa).toLowerCase();return"*"===a?function(){return!0}:function(a){return a.nodeName&&a.nodeName.toLowerCase()===b}},CLASS:function(a){var b=y[a+" "];return b||(b=new RegExp("(^|"+K+")"+a+"("+K+"|$)"))&&y(a,function(a){return b.test("string"==typeof a.className&&a.className||"undefined"!=typeof a.getAttribute&&a.getAttribute("class")||"")})},ATTR:function(a,b,c){return function(d){var e=ga.attr(d,a);return null==e?"!="===b:!b||(e+="","="===b?e===c:"!="===b?e!==c:"^="===b?c&&0===e.indexOf(c):"*="===b?c&&e.indexOf(c)>-1:"$="===b?c&&e.slice(-c.length)===c:"~="===b?(" "+e.replace(O," ")+" ").indexOf(c)>-1:"|="===b&&(e===c||e.slice(0,c.length+1)===c+"-"))}},CHILD:function(a,b,c,d,e){var f="nth"!==a.slice(0,3),g="last"!==a.slice(-4),h="of-type"===b;return 1===d&&0===e?function(a){return!!a.parentNode}:function(b,c,i){var j,k,l,m,n,o,p=f!==g?"nextSibling":"previousSibling",q=b.parentNode,r=h&&b.nodeName.toLowerCase(),s=!i&&!h,t=!1;if(q){if(f){while(p){m=b;while(m=m[p])if(h?m.nodeName.toLowerCase()===r:1===m.nodeType)return!1;o=p="only"===a&&!o&&"nextSibling"}return!0}if(o=[g?q.firstChild:q.lastChild],g&&s){m=q,l=m[u]||(m[u]={}),k=l[m.uniqueID]||(l[m.uniqueID]={}),j=k[a]||[],n=j[0]===w&&j[1],t=n&&j[2],m=n&&q.childNodes[n];while(m=++n&&m&&m[p]||(t=n=0)||o.pop())if(1===m.nodeType&&++t&&m===b){k[a]=[w,n,t];break}}else if(s&&(m=b,l=m[u]||(m[u]={}),k=l[m.uniqueID]||(l[m.uniqueID]={}),j=k[a]||[],n=j[0]===w&&j[1],t=n),t===!1)while(m=++n&&m&&m[p]||(t=n=0)||o.pop())if((h?m.nodeName.toLowerCase()===r:1===m.nodeType)&&++t&&(s&&(l=m[u]||(m[u]={}),k=l[m.uniqueID]||(l[m.uniqueID]={}),k[a]=[w,t]),m===b))break;return t-=e,t===d||t%d===0&&t/d>=0}}},PSEUDO:function(a,b){var c,e=d.pseudos[a]||d.setFilters[a.toLowerCase()]||ga.error("unsupported pseudo: "+a);return e[u]?e(b):e.length>1?(c=[a,a,"",b],d.setFilters.hasOwnProperty(a.toLowerCase())?ia(function(a,c){var d,f=e(a,b),g=f.length;while(g--)d=I(a,f[g]),a[d]=!(c[d]=f[g])}):function(a){return e(a,0,c)}):e}},pseudos:{not:ia(function(a){var b=[],c=[],d=h(a.replace(P,"$1"));return d[u]?ia(function(a,b,c,e){var f,g=d(a,null,e,[]),h=a.length;while(h--)(f=g[h])&&(a[h]=!(b[h]=f))}):function(a,e,f){return b[0]=a,d(b,null,f,c),b[0]=null,!c.pop()}}),has:ia(function(a){return function(b){return ga(a,b).length>0}}),contains:ia(function(a){return a=a.replace(_,aa),function(b){return(b.textContent||b.innerText||e(b)).indexOf(a)>-1}}),lang:ia(function(a){return U.test(a||"")||ga.error("unsupported lang: "+a),a=a.replace(_,aa).toLowerCase(),function(b){var c;do if(c=p?b.lang:b.getAttribute("xml:lang")||b.getAttribute("lang"))return c=c.toLowerCase(),c===a||0===c.indexOf(a+"-");while((b=b.parentNode)&&1===b.nodeType);return!1}}),target:function(b){var c=a.location&&a.location.hash;return c&&c.slice(1)===b.id},root:function(a){return a===o},focus:function(a){return a===n.activeElement&&(!n.hasFocus||n.hasFocus())&&!!(a.type||a.href||~a.tabIndex)},enabled:oa(!1),disabled:oa(!0),checked:function(a){var b=a.nodeName.toLowerCase();return"input"===b&&!!a.checked||"option"===b&&!!a.selected},selected:function(a){return a.parentNode&&a.parentNode.selectedIndex,a.selected===!0},empty:function(a){for(a=a.firstChild;a;a=a.nextSibling)if(a.nodeType<6)return!1;return!0},parent:function(a){return!d.pseudos.empty(a)},header:function(a){return X.test(a.nodeName)},input:function(a){return W.test(a.nodeName)},button:function(a){var b=a.nodeName.toLowerCase();return"input"===b&&"button"===a.type||"button"===b},text:function(a){var b;return"input"===a.nodeName.toLowerCase()&&"text"===a.type&&(null==(b=a.getAttribute("type"))||"text"===b.toLowerCase())},first:pa(function(){return[0]}),last:pa(function(a,b){return[b-1]}),eq:pa(function(a,b,c){return[c<0?c+b:c]}),even:pa(function(a,b){for(var c=0;c<b;c+=2)a.push(c);return a}),odd:pa(function(a,b){for(var c=1;c<b;c+=2)a.push(c);return a}),lt:pa(function(a,b,c){for(var d=c<0?c+b:c;--d>=0;)a.push(d);return a}),gt:pa(function(a,b,c){for(var d=c<0?c+b:c;++d<b;)a.push(d);return a})}},d.pseudos.nth=d.pseudos.eq;for(b in{radio:!0,checkbox:!0,file:!0,password:!0,image:!0})d.pseudos[b]=ma(b);for(b in{submit:!0,reset:!0})d.pseudos[b]=na(b);function ra(){}ra.prototype=d.filters=d.pseudos,d.setFilters=new ra,g=ga.tokenize=function(a,b){var c,e,f,g,h,i,j,k=z[a+" "];if(k)return b?0:k.slice(0);h=a,i=[],j=d.preFilter;while(h){c&&!(e=Q.exec(h))||(e&&(h=h.slice(e[0].length)||h),i.push(f=[])),c=!1,(e=R.exec(h))&&(c=e.shift(),f.push({value:c,type:e[0].replace(P," ")}),h=h.slice(c.length));for(g in d.filter)!(e=V[g].exec(h))||j[g]&&!(e=j[g](e))||(c=e.shift(),f.push({value:c,type:g,matches:e}),h=h.slice(c.length));if(!c)break}return b?h.length:h?ga.error(a):z(a,i).slice(0)};function sa(a){for(var b=0,c=a.length,d="";b<c;b++)d+=a[b].value;return d}function ta(a,b,c){var d=b.dir,e=b.next,f=e||d,g=c&&"parentNode"===f,h=x++;return b.first?function(b,c,e){while(b=b[d])if(1===b.nodeType||g)return a(b,c,e)}:function(b,c,i){var j,k,l,m=[w,h];if(i){while(b=b[d])if((1===b.nodeType||g)&&a(b,c,i))return!0}else while(b=b[d])if(1===b.nodeType||g)if(l=b[u]||(b[u]={}),k=l[b.uniqueID]||(l[b.uniqueID]={}),e&&e===b.nodeName.toLowerCase())b=b[d]||b;else{if((j=k[f])&&j[0]===w&&j[1]===h)return m[2]=j[2];if(k[f]=m,m[2]=a(b,c,i))return!0}}}function ua(a){return a.length>1?function(b,c,d){var e=a.length;while(e--)if(!a[e](b,c,d))return!1;return!0}:a[0]}function va(a,b,c){for(var d=0,e=b.length;d<e;d++)ga(a,b[d],c);return c}function wa(a,b,c,d,e){for(var f,g=[],h=0,i=a.length,j=null!=b;h<i;h++)(f=a[h])&&(c&&!c(f,d,e)||(g.push(f),j&&b.push(h)));return g}function xa(a,b,c,d,e,f){return d&&!d[u]&&(d=xa(d)),e&&!e[u]&&(e=xa(e,f)),ia(function(f,g,h,i){var j,k,l,m=[],n=[],o=g.length,p=f||va(b||"*",h.nodeType?[h]:h,[]),q=!a||!f&&b?p:wa(p,m,a,h,i),r=c?e||(f?a:o||d)?[]:g:q;if(c&&c(q,r,h,i),d){j=wa(r,n),d(j,[],h,i),k=j.length;while(k--)(l=j[k])&&(r[n[k]]=!(q[n[k]]=l))}if(f){if(e||a){if(e){j=[],k=r.length;while(k--)(l=r[k])&&j.push(q[k]=l);e(null,r=[],j,i)}k=r.length;while(k--)(l=r[k])&&(j=e?I(f,l):m[k])>-1&&(f[j]=!(g[j]=l))}}else r=wa(r===g?r.splice(o,r.length):r),e?e(null,g,r,i):G.apply(g,r)})}function ya(a){for(var b,c,e,f=a.length,g=d.relative[a[0].type],h=g||d.relative[" "],i=g?1:0,k=ta(function(a){return a===b},h,!0),l=ta(function(a){return I(b,a)>-1},h,!0),m=[function(a,c,d){var e=!g&&(d||c!==j)||((b=c).nodeType?k(a,c,d):l(a,c,d));return b=null,e}];i<f;i++)if(c=d.relative[a[i].type])m=[ta(ua(m),c)];else{if(c=d.filter[a[i].type].apply(null,a[i].matches),c[u]){for(e=++i;e<f;e++)if(d.relative[a[e].type])break;return xa(i>1&&ua(m),i>1&&sa(a.slice(0,i-1).concat({value:" "===a[i-2].type?"*":""})).replace(P,"$1"),c,i<e&&ya(a.slice(i,e)),e<f&&ya(a=a.slice(e)),e<f&&sa(a))}m.push(c)}return ua(m)}function za(a,b){var c=b.length>0,e=a.length>0,f=function(f,g,h,i,k){var l,o,q,r=0,s="0",t=f&&[],u=[],v=j,x=f||e&&d.find.TAG("*",k),y=w+=null==v?1:Math.random()||.1,z=x.length;for(k&&(j=g===n||g||k);s!==z&&null!=(l=x[s]);s++){if(e&&l){o=0,g||l.ownerDocument===n||(m(l),h=!p);while(q=a[o++])if(q(l,g||n,h)){i.push(l);break}k&&(w=y)}c&&((l=!q&&l)&&r--,f&&t.push(l))}if(r+=s,c&&s!==r){o=0;while(q=b[o++])q(t,u,g,h);if(f){if(r>0)while(s--)t[s]||u[s]||(u[s]=E.call(i));u=wa(u)}G.apply(i,u),k&&!f&&u.length>0&&r+b.length>1&&ga.uniqueSort(i)}return k&&(w=y,j=v),t};return c?ia(f):f}return h=ga.compile=function(a,b){var c,d=[],e=[],f=A[a+" "];if(!f){b||(b=g(a)),c=b.length;while(c--)f=ya(b[c]),f[u]?d.push(f):e.push(f);f=A(a,za(e,d)),f.selector=a}return f},i=ga.select=function(a,b,e,f){var i,j,k,l,m,n="function"==typeof a&&a,o=!f&&g(a=n.selector||a);if(e=e||[],1===o.length){if(j=o[0]=o[0].slice(0),j.length>2&&"ID"===(k=j[0]).type&&c.getById&&9===b.nodeType&&p&&d.relative[j[1].type]){if(b=(d.find.ID(k.matches[0].replace(_,aa),b)||[])[0],!b)return e;n&&(b=b.parentNode),a=a.slice(j.shift().value.length)}i=V.needsContext.test(a)?0:j.length;while(i--){if(k=j[i],d.relative[l=k.type])break;if((m=d.find[l])&&(f=m(k.matches[0].replace(_,aa),$.test(j[0].type)&&qa(b.parentNode)||b))){if(j.splice(i,1),a=f.length&&sa(j),!a)return G.apply(e,f),e;break}}}return(n||h(a,o))(f,b,!p,e,!b||$.test(a)&&qa(b.parentNode)||b),e},c.sortStable=u.split("").sort(B).join("")===u,c.detectDuplicates=!!l,m(),c.sortDetached=ja(function(a){return 1&a.compareDocumentPosition(n.createElement("fieldset"))}),ja(function(a){return a.innerHTML="<a href='#'></a>","#"===a.firstChild.getAttribute("href")})||ka("type|href|height|width",function(a,b,c){if(!c)return a.getAttribute(b,"type"===b.toLowerCase()?1:2)}),c.attributes&&ja(function(a){return a.innerHTML="<input/>",a.firstChild.setAttribute("value",""),""===a.firstChild.getAttribute("value")})||ka("value",function(a,b,c){if(!c&&"input"===a.nodeName.toLowerCase())return a.defaultValue}),ja(function(a){return null==a.getAttribute("disabled")})||ka(J,function(a,b,c){var d;if(!c)return a[b]===!0?b.toLowerCase():(d=a.getAttributeNode(b))&&d.specified?d.value:null}),ga}(a);r.find=x,r.expr=x.selectors,r.expr[":"]=r.expr.pseudos,r.uniqueSort=r.unique=x.uniqueSort,r.text=x.getText,r.isXMLDoc=x.isXML,r.contains=x.contains,r.escapeSelector=x.escape;var y=function(a,b,c){var d=[],e=void 0!==c;while((a=a[b])&&9!==a.nodeType)if(1===a.nodeType){if(e&&r(a).is(c))break;d.push(a)}return d},z=function(a,b){for(var c=[];a;a=a.nextSibling)1===a.nodeType&&a!==b&&c.push(a);return c},A=r.expr.match.needsContext,B=/^<([a-z][^\/\0>:\x20\t\r\n\f]*)[\x20\t\r\n\f]*\/?>(?:<\/\1>|)$/i,C=/^.[^:#\[\.,]*$/;function D(a,b,c){if(r.isFunction(b))return r.grep(a,function(a,d){return!!b.call(a,d,a)!==c});if(b.nodeType)return r.grep(a,function(a){return a===b!==c});if("string"==typeof b){if(C.test(b))return r.filter(b,a,c);b=r.filter(b,a)}return r.grep(a,function(a){return i.call(b,a)>-1!==c&&1===a.nodeType})}r.filter=function(a,b,c){var d=b[0];return c&&(a=":not("+a+")"),1===b.length&&1===d.nodeType?r.find.matchesSelector(d,a)?[d]:[]:r.find.matches(a,r.grep(b,function(a){return 1===a.nodeType}))},r.fn.extend({find:function(a){var b,c,d=this.length,e=this;if("string"!=typeof a)return this.pushStack(r(a).filter(function(){for(b=0;b<d;b++)if(r.contains(e[b],this))return!0}));for(c=this.pushStack([]),b=0;b<d;b++)r.find(a,e[b],c);return d>1?r.uniqueSort(c):c},filter:function(a){return this.pushStack(D(this,a||[],!1))},not:function(a){return this.pushStack(D(this,a||[],!0))},is:function(a){return!!D(this,"string"==typeof a&&A.test(a)?r(a):a||[],!1).length}});var E,F=/^(?:\s*(<[\w\W]+>)[^>]*|#([\w-]+))$/,G=r.fn.init=function(a,b,c){var e,f;if(!a)return this;if(c=c||E,"string"==typeof a){if(e="<"===a[0]&&">"===a[a.length-1]&&a.length>=3?[null,a,null]:F.exec(a),!e||!e[1]&&b)return!b||b.jquery?(b||c).find(a):this.constructor(b).find(a);if(e[1]){if(b=b instanceof r?b[0]:b,r.merge(this,r.parseHTML(e[1],b&&b.nodeType?b.ownerDocument||b:d,!0)),B.test(e[1])&&r.isPlainObject(b))for(e in b)r.isFunction(this[e])?this[e](b[e]):this.attr(e,b[e]);return this}return f=d.getElementById(e[2]),f&&(this[0]=f,this.length=1),this}return a.nodeType?(this[0]=a,this.length=1,this):r.isFunction(a)?void 0!==c.ready?c.ready(a):a(r):r.makeArray(a,this)};G.prototype=r.fn,E=r(d);var H=/^(?:parents|prev(?:Until|All))/,I={children:!0,contents:!0,next:!0,prev:!0};r.fn.extend({has:function(a){var b=r(a,this),c=b.length;return this.filter(function(){for(var a=0;a<c;a++)if(r.contains(this,b[a]))return!0})},closest:function(a,b){var c,d=0,e=this.length,f=[],g="string"!=typeof a&&r(a);if(!A.test(a))for(;d<e;d++)for(c=this[d];c&&c!==b;c=c.parentNode)if(c.nodeType<11&&(g?g.index(c)>-1:1===c.nodeType&&r.find.matchesSelector(c,a))){f.push(c);break}return this.pushStack(f.length>1?r.uniqueSort(f):f)},index:function(a){return a?"string"==typeof a?i.call(r(a),this[0]):i.call(this,a.jquery?a[0]:a):this[0]&&this[0].parentNode?this.first().prevAll().length:-1},add:function(a,b){return this.pushStack(r.uniqueSort(r.merge(this.get(),r(a,b))))},addBack:function(a){return this.add(null==a?this.prevObject:this.prevObject.filter(a))}});function J(a,b){while((a=a[b])&&1!==a.nodeType);return a}r.each({parent:function(a){var b=a.parentNode;return b&&11!==b.nodeType?b:null},parents:function(a){return y(a,"parentNode")},parentsUntil:function(a,b,c){return y(a,"parentNode",c)},next:function(a){return J(a,"nextSibling")},prev:function(a){return J(a,"previousSibling")},nextAll:function(a){return y(a,"nextSibling")},prevAll:function(a){return y(a,"previousSibling")},nextUntil:function(a,b,c){return y(a,"nextSibling",c)},prevUntil:function(a,b,c){return y(a,"previousSibling",c)},siblings:function(a){return z((a.parentNode||{}).firstChild,a)},children:function(a){return z(a.firstChild)},contents:function(a){return a.contentDocument||r.merge([],a.childNodes)}},function(a,b){r.fn[a]=function(c,d){var e=r.map(this,b,c);return"Until"!==a.slice(-5)&&(d=c),d&&"string"==typeof d&&(e=r.filter(d,e)),this.length>1&&(I[a]||r.uniqueSort(e),H.test(a)&&e.reverse()),this.pushStack(e)}});var K=/\S+/g;function L(a){var b={};return r.each(a.match(K)||[],function(a,c){b[c]=!0}),b}r.Callbacks=function(a){a="string"==typeof a?L(a):r.extend({},a);var b,c,d,e,f=[],g=[],h=-1,i=function(){for(e=a.once,d=b=!0;g.length;h=-1){c=g.shift();while(++h<f.length)f[h].apply(c[0],c[1])===!1&&a.stopOnFalse&&(h=f.length,c=!1)}a.memory||(c=!1),b=!1,e&&(f=c?[]:"")},j={add:function(){return f&&(c&&!b&&(h=f.length-1,g.push(c)),function d(b){r.each(b,function(b,c){r.isFunction(c)?a.unique&&j.has(c)||f.push(c):c&&c.length&&"string"!==r.type(c)&&d(c)})}(arguments),c&&!b&&i()),this},remove:function(){return r.each(arguments,function(a,b){var c;while((c=r.inArray(b,f,c))>-1)f.splice(c,1),c<=h&&h--}),this},has:function(a){return a?r.inArray(a,f)>-1:f.length>0},empty:function(){return f&&(f=[]),this},disable:function(){return e=g=[],f=c="",this},disabled:function(){return!f},lock:function(){return e=g=[],c||b||(f=c=""),this},locked:function(){return!!e},fireWith:function(a,c){return e||(c=c||[],c=[a,c.slice?c.slice():c],g.push(c),b||i()),this},fire:function(){return j.fireWith(this,arguments),this},fired:function(){return!!d}};return j};function M(a){return a}function N(a){throw a}function O(a,b,c){var d;try{a&&r.isFunction(d=a.promise)?d.call(a).done(b).fail(c):a&&r.isFunction(d=a.then)?d.call(a,b,c):b.call(void 0,a)}catch(a){c.call(void 0,a)}}r.extend({Deferred:function(b){var c=[["notify","progress",r.Callbacks("memory"),r.Callbacks("memory"),2],["resolve","done",r.Callbacks("once memory"),r.Callbacks("once memory"),0,"resolved"],["reject","fail",r.Callbacks("once memory"),r.Callbacks("once memory"),1,"rejected"]],d="pending",e={state:function(){return d},always:function(){return f.done(arguments).fail(arguments),this},"catch":function(a){return e.then(null,a)},pipe:function(){var a=arguments;return r.Deferred(function(b){r.each(c,function(c,d){var e=r.isFunction(a[d[4]])&&a[d[4]];f[d[1]](function(){var a=e&&e.apply(this,arguments);a&&r.isFunction(a.promise)?a.promise().progress(b.notify).done(b.resolve).fail(b.reject):b[d[0]+"With"](this,e?[a]:arguments)})}),a=null}).promise()},then:function(b,d,e){var f=0;function g(b,c,d,e){return function(){var h=this,i=arguments,j=function(){var a,j;if(!(b<f)){if(a=d.apply(h,i),a===c.promise())throw new TypeError("Thenable self-resolution");j=a&&("object"==typeof a||"function"==typeof a)&&a.then,r.isFunction(j)?e?j.call(a,g(f,c,M,e),g(f,c,N,e)):(f++,j.call(a,g(f,c,M,e),g(f,c,N,e),g(f,c,M,c.notifyWith))):(d!==M&&(h=void 0,i=[a]),(e||c.resolveWith)(h,i))}},k=e?j:function(){try{j()}catch(a){r.Deferred.exceptionHook&&r.Deferred.exceptionHook(a,k.stackTrace),b+1>=f&&(d!==N&&(h=void 0,i=[a]),c.rejectWith(h,i))}};b?k():(r.Deferred.getStackHook&&(k.stackTrace=r.Deferred.getStackHook()),a.setTimeout(k))}}return r.Deferred(function(a){c[0][3].add(g(0,a,r.isFunction(e)?e:M,a.notifyWith)),c[1][3].add(g(0,a,r.isFunction(b)?b:M)),c[2][3].add(g(0,a,r.isFunction(d)?d:N))}).promise()},promise:function(a){return null!=a?r.extend(a,e):e}},f={};return r.each(c,function(a,b){var g=b[2],h=b[5];e[b[1]]=g.add,h&&g.add(function(){d=h},c[3-a][2].disable,c[0][2].lock),g.add(b[3].fire),f[b[0]]=function(){return f[b[0]+"With"](this===f?void 0:this,arguments),this},f[b[0]+"With"]=g.fireWith}),e.promise(f),b&&b.call(f,f),f},when:function(a){var b=arguments.length,c=b,d=Array(c),e=f.call(arguments),g=r.Deferred(),h=function(a){return function(c){d[a]=this,e[a]=arguments.length>1?f.call(arguments):c,--b||g.resolveWith(d,e)}};if(b<=1&&(O(a,g.done(h(c)).resolve,g.reject),"pending"===g.state()||r.isFunction(e[c]&&e[c].then)))return g.then();while(c--)O(e[c],h(c),g.reject);return g.promise()}});var P=/^(Eval|Internal|Range|Reference|Syntax|Type|URI)Error$/;r.Deferred.exceptionHook=function(b,c){a.console&&a.console.warn&&b&&P.test(b.name)&&a.console.warn("jQuery.Deferred exception: "+b.message,b.stack,c)},r.readyException=function(b){a.setTimeout(function(){throw b})};var Q=r.Deferred();r.fn.ready=function(a){return Q.then(a)["catch"](function(a){r.readyException(a)}),this},r.extend({isReady:!1,readyWait:1,holdReady:function(a){a?r.readyWait++:r.ready(!0)},ready:function(a){(a===!0?--r.readyWait:r.isReady)||(r.isReady=!0,a!==!0&&--r.readyWait>0||Q.resolveWith(d,[r]))}}),r.ready.then=Q.then;function R(){d.removeEventListener("DOMContentLoaded",R),a.removeEventListener("load",R),r.ready()}"complete"===d.readyState||"loading"!==d.readyState&&!d.documentElement.doScroll?a.setTimeout(r.ready):(d.addEventListener("DOMContentLoaded",R),a.addEventListener("load",R));var S=function(a,b,c,d,e,f,g){var h=0,i=a.length,j=null==c;if("object"===r.type(c)){e=!0;for(h in c)S(a,b,h,c[h],!0,f,g)}else if(void 0!==d&&(e=!0,
r.isFunction(d)||(g=!0),j&&(g?(b.call(a,d),b=null):(j=b,b=function(a,b,c){return j.call(r(a),c)})),b))for(;h<i;h++)b(a[h],c,g?d:d.call(a[h],h,b(a[h],c)));return e?a:j?b.call(a):i?b(a[0],c):f},T=function(a){return 1===a.nodeType||9===a.nodeType||!+a.nodeType};function U(){this.expando=r.expando+U.uid++}U.uid=1,U.prototype={cache:function(a){var b=a[this.expando];return b||(b={},T(a)&&(a.nodeType?a[this.expando]=b:Object.defineProperty(a,this.expando,{value:b,configurable:!0}))),b},set:function(a,b,c){var d,e=this.cache(a);if("string"==typeof b)e[r.camelCase(b)]=c;else for(d in b)e[r.camelCase(d)]=b[d];return e},get:function(a,b){return void 0===b?this.cache(a):a[this.expando]&&a[this.expando][r.camelCase(b)]},access:function(a,b,c){return void 0===b||b&&"string"==typeof b&&void 0===c?this.get(a,b):(this.set(a,b,c),void 0!==c?c:b)},remove:function(a,b){var c,d=a[this.expando];if(void 0!==d){if(void 0!==b){r.isArray(b)?b=b.map(r.camelCase):(b=r.camelCase(b),b=b in d?[b]:b.match(K)||[]),c=b.length;while(c--)delete d[b[c]]}(void 0===b||r.isEmptyObject(d))&&(a.nodeType?a[this.expando]=void 0:delete a[this.expando])}},hasData:function(a){var b=a[this.expando];return void 0!==b&&!r.isEmptyObject(b)}};var V=new U,W=new U,X=/^(?:\{[\w\W]*\}|\[[\w\W]*\])$/,Y=/[A-Z]/g;function Z(a,b,c){var d;if(void 0===c&&1===a.nodeType)if(d="data-"+b.replace(Y,"-$&").toLowerCase(),c=a.getAttribute(d),"string"==typeof c){try{c="true"===c||"false"!==c&&("null"===c?null:+c+""===c?+c:X.test(c)?JSON.parse(c):c)}catch(e){}W.set(a,b,c)}else c=void 0;return c}r.extend({hasData:function(a){return W.hasData(a)||V.hasData(a)},data:function(a,b,c){return W.access(a,b,c)},removeData:function(a,b){W.remove(a,b)},_data:function(a,b,c){return V.access(a,b,c)},_removeData:function(a,b){V.remove(a,b)}}),r.fn.extend({data:function(a,b){var c,d,e,f=this[0],g=f&&f.attributes;if(void 0===a){if(this.length&&(e=W.get(f),1===f.nodeType&&!V.get(f,"hasDataAttrs"))){c=g.length;while(c--)g[c]&&(d=g[c].name,0===d.indexOf("data-")&&(d=r.camelCase(d.slice(5)),Z(f,d,e[d])));V.set(f,"hasDataAttrs",!0)}return e}return"object"==typeof a?this.each(function(){W.set(this,a)}):S(this,function(b){var c;if(f&&void 0===b){if(c=W.get(f,a),void 0!==c)return c;if(c=Z(f,a),void 0!==c)return c}else this.each(function(){W.set(this,a,b)})},null,b,arguments.length>1,null,!0)},removeData:function(a){return this.each(function(){W.remove(this,a)})}}),r.extend({queue:function(a,b,c){var d;if(a)return b=(b||"fx")+"queue",d=V.get(a,b),c&&(!d||r.isArray(c)?d=V.access(a,b,r.makeArray(c)):d.push(c)),d||[]},dequeue:function(a,b){b=b||"fx";var c=r.queue(a,b),d=c.length,e=c.shift(),f=r._queueHooks(a,b),g=function(){r.dequeue(a,b)};"inprogress"===e&&(e=c.shift(),d--),e&&("fx"===b&&c.unshift("inprogress"),delete f.stop,e.call(a,g,f)),!d&&f&&f.empty.fire()},_queueHooks:function(a,b){var c=b+"queueHooks";return V.get(a,c)||V.access(a,c,{empty:r.Callbacks("once memory").add(function(){V.remove(a,[b+"queue",c])})})}}),r.fn.extend({queue:function(a,b){var c=2;return"string"!=typeof a&&(b=a,a="fx",c--),arguments.length<c?r.queue(this[0],a):void 0===b?this:this.each(function(){var c=r.queue(this,a,b);r._queueHooks(this,a),"fx"===a&&"inprogress"!==c[0]&&r.dequeue(this,a)})},dequeue:function(a){return this.each(function(){r.dequeue(this,a)})},clearQueue:function(a){return this.queue(a||"fx",[])},promise:function(a,b){var c,d=1,e=r.Deferred(),f=this,g=this.length,h=function(){--d||e.resolveWith(f,[f])};"string"!=typeof a&&(b=a,a=void 0),a=a||"fx";while(g--)c=V.get(f[g],a+"queueHooks"),c&&c.empty&&(d++,c.empty.add(h));return h(),e.promise(b)}});var $=/[+-]?(?:\d*\.|)\d+(?:[eE][+-]?\d+|)/.source,_=new RegExp("^(?:([+-])=|)("+$+")([a-z%]*)$","i"),aa=["Top","Right","Bottom","Left"],ba=function(a,b){return a=b||a,"none"===a.style.display||""===a.style.display&&r.contains(a.ownerDocument,a)&&"none"===r.css(a,"display")},ca=function(a,b,c,d){var e,f,g={};for(f in b)g[f]=a.style[f],a.style[f]=b[f];e=c.apply(a,d||[]);for(f in b)a.style[f]=g[f];return e};function da(a,b,c,d){var e,f=1,g=20,h=d?function(){return d.cur()}:function(){return r.css(a,b,"")},i=h(),j=c&&c[3]||(r.cssNumber[b]?"":"px"),k=(r.cssNumber[b]||"px"!==j&&+i)&&_.exec(r.css(a,b));if(k&&k[3]!==j){j=j||k[3],c=c||[],k=+i||1;do f=f||".5",k/=f,r.style(a,b,k+j);while(f!==(f=h()/i)&&1!==f&&--g)}return c&&(k=+k||+i||0,e=c[1]?k+(c[1]+1)*c[2]:+c[2],d&&(d.unit=j,d.start=k,d.end=e)),e}var ea={};function fa(a){var b,c=a.ownerDocument,d=a.nodeName,e=ea[d];return e?e:(b=c.body.appendChild(c.createElement(d)),e=r.css(b,"display"),b.parentNode.removeChild(b),"none"===e&&(e="block"),ea[d]=e,e)}function ga(a,b){for(var c,d,e=[],f=0,g=a.length;f<g;f++)d=a[f],d.style&&(c=d.style.display,b?("none"===c&&(e[f]=V.get(d,"display")||null,e[f]||(d.style.display="")),""===d.style.display&&ba(d)&&(e[f]=fa(d))):"none"!==c&&(e[f]="none",V.set(d,"display",c)));for(f=0;f<g;f++)null!=e[f]&&(a[f].style.display=e[f]);return a}r.fn.extend({show:function(){return ga(this,!0)},hide:function(){return ga(this)},toggle:function(a){return"boolean"==typeof a?a?this.show():this.hide():this.each(function(){ba(this)?r(this).show():r(this).hide()})}});var ha=/^(?:checkbox|radio)$/i,ia=/<([a-z][^\/\0>\x20\t\r\n\f]+)/i,ja=/^$|\/(?:java|ecma)script/i,ka={option:[1,"<select multiple='multiple'>","</select>"],thead:[1,"<table>","</table>"],col:[2,"<table><colgroup>","</colgroup></table>"],tr:[2,"<table><tbody>","</tbody></table>"],td:[3,"<table><tbody><tr>","</tr></tbody></table>"],_default:[0,"",""]};ka.optgroup=ka.option,ka.tbody=ka.tfoot=ka.colgroup=ka.caption=ka.thead,ka.th=ka.td;function la(a,b){var c="undefined"!=typeof a.getElementsByTagName?a.getElementsByTagName(b||"*"):"undefined"!=typeof a.querySelectorAll?a.querySelectorAll(b||"*"):[];return void 0===b||b&&r.nodeName(a,b)?r.merge([a],c):c}function ma(a,b){for(var c=0,d=a.length;c<d;c++)V.set(a[c],"globalEval",!b||V.get(b[c],"globalEval"))}var na=/<|&#?\w+;/;function oa(a,b,c,d,e){for(var f,g,h,i,j,k,l=b.createDocumentFragment(),m=[],n=0,o=a.length;n<o;n++)if(f=a[n],f||0===f)if("object"===r.type(f))r.merge(m,f.nodeType?[f]:f);else if(na.test(f)){g=g||l.appendChild(b.createElement("div")),h=(ia.exec(f)||["",""])[1].toLowerCase(),i=ka[h]||ka._default,g.innerHTML=i[1]+r.htmlPrefilter(f)+i[2],k=i[0];while(k--)g=g.lastChild;r.merge(m,g.childNodes),g=l.firstChild,g.textContent=""}else m.push(b.createTextNode(f));l.textContent="",n=0;while(f=m[n++])if(d&&r.inArray(f,d)>-1)e&&e.push(f);else if(j=r.contains(f.ownerDocument,f),g=la(l.appendChild(f),"script"),j&&ma(g),c){k=0;while(f=g[k++])ja.test(f.type||"")&&c.push(f)}return l}!function(){var a=d.createDocumentFragment(),b=a.appendChild(d.createElement("div")),c=d.createElement("input");c.setAttribute("type","radio"),c.setAttribute("checked","checked"),c.setAttribute("name","t"),b.appendChild(c),o.checkClone=b.cloneNode(!0).cloneNode(!0).lastChild.checked,b.innerHTML="<textarea>x</textarea>",o.noCloneChecked=!!b.cloneNode(!0).lastChild.defaultValue}();var pa=d.documentElement,qa=/^key/,ra=/^(?:mouse|pointer|contextmenu|drag|drop)|click/,sa=/^([^.]*)(?:\.(.+)|)/;function ta(){return!0}function ua(){return!1}function va(){try{return d.activeElement}catch(a){}}function wa(a,b,c,d,e,f){var g,h;if("object"==typeof b){"string"!=typeof c&&(d=d||c,c=void 0);for(h in b)wa(a,h,c,d,b[h],f);return a}if(null==d&&null==e?(e=c,d=c=void 0):null==e&&("string"==typeof c?(e=d,d=void 0):(e=d,d=c,c=void 0)),e===!1)e=ua;else if(!e)return a;return 1===f&&(g=e,e=function(a){return r().off(a),g.apply(this,arguments)},e.guid=g.guid||(g.guid=r.guid++)),a.each(function(){r.event.add(this,b,e,d,c)})}r.event={global:{},add:function(a,b,c,d,e){var f,g,h,i,j,k,l,m,n,o,p,q=V.get(a);if(q){c.handler&&(f=c,c=f.handler,e=f.selector),e&&r.find.matchesSelector(pa,e),c.guid||(c.guid=r.guid++),(i=q.events)||(i=q.events={}),(g=q.handle)||(g=q.handle=function(b){return"undefined"!=typeof r&&r.event.triggered!==b.type?r.event.dispatch.apply(a,arguments):void 0}),b=(b||"").match(K)||[""],j=b.length;while(j--)h=sa.exec(b[j])||[],n=p=h[1],o=(h[2]||"").split(".").sort(),n&&(l=r.event.special[n]||{},n=(e?l.delegateType:l.bindType)||n,l=r.event.special[n]||{},k=r.extend({type:n,origType:p,data:d,handler:c,guid:c.guid,selector:e,needsContext:e&&r.expr.match.needsContext.test(e),namespace:o.join(".")},f),(m=i[n])||(m=i[n]=[],m.delegateCount=0,l.setup&&l.setup.call(a,d,o,g)!==!1||a.addEventListener&&a.addEventListener(n,g)),l.add&&(l.add.call(a,k),k.handler.guid||(k.handler.guid=c.guid)),e?m.splice(m.delegateCount++,0,k):m.push(k),r.event.global[n]=!0)}},remove:function(a,b,c,d,e){var f,g,h,i,j,k,l,m,n,o,p,q=V.hasData(a)&&V.get(a);if(q&&(i=q.events)){b=(b||"").match(K)||[""],j=b.length;while(j--)if(h=sa.exec(b[j])||[],n=p=h[1],o=(h[2]||"").split(".").sort(),n){l=r.event.special[n]||{},n=(d?l.delegateType:l.bindType)||n,m=i[n]||[],h=h[2]&&new RegExp("(^|\\.)"+o.join("\\.(?:.*\\.|)")+"(\\.|$)"),g=f=m.length;while(f--)k=m[f],!e&&p!==k.origType||c&&c.guid!==k.guid||h&&!h.test(k.namespace)||d&&d!==k.selector&&("**"!==d||!k.selector)||(m.splice(f,1),k.selector&&m.delegateCount--,l.remove&&l.remove.call(a,k));g&&!m.length&&(l.teardown&&l.teardown.call(a,o,q.handle)!==!1||r.removeEvent(a,n,q.handle),delete i[n])}else for(n in i)r.event.remove(a,n+b[j],c,d,!0);r.isEmptyObject(i)&&V.remove(a,"handle events")}},dispatch:function(a){var b=r.event.fix(a),c,d,e,f,g,h,i=new Array(arguments.length),j=(V.get(this,"events")||{})[b.type]||[],k=r.event.special[b.type]||{};for(i[0]=b,c=1;c<arguments.length;c++)i[c]=arguments[c];if(b.delegateTarget=this,!k.preDispatch||k.preDispatch.call(this,b)!==!1){h=r.event.handlers.call(this,b,j),c=0;while((f=h[c++])&&!b.isPropagationStopped()){b.currentTarget=f.elem,d=0;while((g=f.handlers[d++])&&!b.isImmediatePropagationStopped())b.rnamespace&&!b.rnamespace.test(g.namespace)||(b.handleObj=g,b.data=g.data,e=((r.event.special[g.origType]||{}).handle||g.handler).apply(f.elem,i),void 0!==e&&(b.result=e)===!1&&(b.preventDefault(),b.stopPropagation()))}return k.postDispatch&&k.postDispatch.call(this,b),b.result}},handlers:function(a,b){var c,d,e,f,g=[],h=b.delegateCount,i=a.target;if(h&&i.nodeType&&("click"!==a.type||isNaN(a.button)||a.button<1))for(;i!==this;i=i.parentNode||this)if(1===i.nodeType&&(i.disabled!==!0||"click"!==a.type)){for(d=[],c=0;c<h;c++)f=b[c],e=f.selector+" ",void 0===d[e]&&(d[e]=f.needsContext?r(e,this).index(i)>-1:r.find(e,this,null,[i]).length),d[e]&&d.push(f);d.length&&g.push({elem:i,handlers:d})}return h<b.length&&g.push({elem:this,handlers:b.slice(h)}),g},addProp:function(a,b){Object.defineProperty(r.Event.prototype,a,{enumerable:!0,configurable:!0,get:r.isFunction(b)?function(){if(this.originalEvent)return b(this.originalEvent)}:function(){if(this.originalEvent)return this.originalEvent[a]},set:function(b){Object.defineProperty(this,a,{enumerable:!0,configurable:!0,writable:!0,value:b})}})},fix:function(a){return a[r.expando]?a:new r.Event(a)},special:{load:{noBubble:!0},focus:{trigger:function(){if(this!==va()&&this.focus)return this.focus(),!1},delegateType:"focusin"},blur:{trigger:function(){if(this===va()&&this.blur)return this.blur(),!1},delegateType:"focusout"},click:{trigger:function(){if("checkbox"===this.type&&this.click&&r.nodeName(this,"input"))return this.click(),!1},_default:function(a){return r.nodeName(a.target,"a")}},beforeunload:{postDispatch:function(a){void 0!==a.result&&a.originalEvent&&(a.originalEvent.returnValue=a.result)}}}},r.removeEvent=function(a,b,c){a.removeEventListener&&a.removeEventListener(b,c)},r.Event=function(a,b){return this instanceof r.Event?(a&&a.type?(this.originalEvent=a,this.type=a.type,this.isDefaultPrevented=a.defaultPrevented||void 0===a.defaultPrevented&&a.returnValue===!1?ta:ua,this.target=a.target&&3===a.target.nodeType?a.target.parentNode:a.target,this.currentTarget=a.currentTarget,this.relatedTarget=a.relatedTarget):this.type=a,b&&r.extend(this,b),this.timeStamp=a&&a.timeStamp||r.now(),void(this[r.expando]=!0)):new r.Event(a,b)},r.Event.prototype={constructor:r.Event,isDefaultPrevented:ua,isPropagationStopped:ua,isImmediatePropagationStopped:ua,isSimulated:!1,preventDefault:function(){var a=this.originalEvent;this.isDefaultPrevented=ta,a&&!this.isSimulated&&a.preventDefault()},stopPropagation:function(){var a=this.originalEvent;this.isPropagationStopped=ta,a&&!this.isSimulated&&a.stopPropagation()},stopImmediatePropagation:function(){var a=this.originalEvent;this.isImmediatePropagationStopped=ta,a&&!this.isSimulated&&a.stopImmediatePropagation(),this.stopPropagation()}},r.each({altKey:!0,bubbles:!0,cancelable:!0,changedTouches:!0,ctrlKey:!0,detail:!0,eventPhase:!0,metaKey:!0,pageX:!0,pageY:!0,shiftKey:!0,view:!0,"char":!0,charCode:!0,key:!0,keyCode:!0,button:!0,buttons:!0,clientX:!0,clientY:!0,offsetX:!0,offsetY:!0,pointerId:!0,pointerType:!0,screenX:!0,screenY:!0,targetTouches:!0,toElement:!0,touches:!0,which:function(a){var b=a.button;return null==a.which&&qa.test(a.type)?null!=a.charCode?a.charCode:a.keyCode:!a.which&&void 0!==b&&ra.test(a.type)?1&b?1:2&b?3:4&b?2:0:a.which}},r.event.addProp),r.each({mouseenter:"mouseover",mouseleave:"mouseout",pointerenter:"pointerover",pointerleave:"pointerout"},function(a,b){r.event.special[a]={delegateType:b,bindType:b,handle:function(a){var c,d=this,e=a.relatedTarget,f=a.handleObj;return e&&(e===d||r.contains(d,e))||(a.type=f.origType,c=f.handler.apply(this,arguments),a.type=b),c}}}),r.fn.extend({on:function(a,b,c,d){return wa(this,a,b,c,d)},one:function(a,b,c,d){return wa(this,a,b,c,d,1)},off:function(a,b,c){var d,e;if(a&&a.preventDefault&&a.handleObj)return d=a.handleObj,r(a.delegateTarget).off(d.namespace?d.origType+"."+d.namespace:d.origType,d.selector,d.handler),this;if("object"==typeof a){for(e in a)this.off(e,b,a[e]);return this}return b!==!1&&"function"!=typeof b||(c=b,b=void 0),c===!1&&(c=ua),this.each(function(){r.event.remove(this,a,c,b)})}});var xa=/<(?!area|br|col|embed|hr|img|input|link|meta|param)(([a-z][^\/\0>\x20\t\r\n\f]*)[^>]*)\/>/gi,ya=/<script|<style|<link/i,za=/checked\s*(?:[^=]|=\s*.checked.)/i,Aa=/^true\/(.*)/,Ba=/^\s*<!(?:\[CDATA\[|--)|(?:\]\]|--)>\s*$/g;function Ca(a,b){return r.nodeName(a,"table")&&r.nodeName(11!==b.nodeType?b:b.firstChild,"tr")?a.getElementsByTagName("tbody")[0]||a:a}function Da(a){return a.type=(null!==a.getAttribute("type"))+"/"+a.type,a}function Ea(a){var b=Aa.exec(a.type);return b?a.type=b[1]:a.removeAttribute("type"),a}function Fa(a,b){var c,d,e,f,g,h,i,j;if(1===b.nodeType){if(V.hasData(a)&&(f=V.access(a),g=V.set(b,f),j=f.events)){delete g.handle,g.events={};for(e in j)for(c=0,d=j[e].length;c<d;c++)r.event.add(b,e,j[e][c])}W.hasData(a)&&(h=W.access(a),i=r.extend({},h),W.set(b,i))}}function Ga(a,b){var c=b.nodeName.toLowerCase();"input"===c&&ha.test(a.type)?b.checked=a.checked:"input"!==c&&"textarea"!==c||(b.defaultValue=a.defaultValue)}function Ha(a,b,c,d){b=g.apply([],b);var e,f,h,i,j,k,l=0,m=a.length,n=m-1,q=b[0],s=r.isFunction(q);if(s||m>1&&"string"==typeof q&&!o.checkClone&&za.test(q))return a.each(function(e){var f=a.eq(e);s&&(b[0]=q.call(this,e,f.html())),Ha(f,b,c,d)});if(m&&(e=oa(b,a[0].ownerDocument,!1,a,d),f=e.firstChild,1===e.childNodes.length&&(e=f),f||d)){for(h=r.map(la(e,"script"),Da),i=h.length;l<m;l++)j=e,l!==n&&(j=r.clone(j,!0,!0),i&&r.merge(h,la(j,"script"))),c.call(a[l],j,l);if(i)for(k=h[h.length-1].ownerDocument,r.map(h,Ea),l=0;l<i;l++)j=h[l],ja.test(j.type||"")&&!V.access(j,"globalEval")&&r.contains(k,j)&&(j.src?r._evalUrl&&r._evalUrl(j.src):p(j.textContent.replace(Ba,""),k))}return a}function Ia(a,b,c){for(var d,e=b?r.filter(b,a):a,f=0;null!=(d=e[f]);f++)c||1!==d.nodeType||r.cleanData(la(d)),d.parentNode&&(c&&r.contains(d.ownerDocument,d)&&ma(la(d,"script")),d.parentNode.removeChild(d));return a}r.extend({htmlPrefilter:function(a){return a.replace(xa,"<$1></$2>")},clone:function(a,b,c){var d,e,f,g,h=a.cloneNode(!0),i=r.contains(a.ownerDocument,a);if(!(o.noCloneChecked||1!==a.nodeType&&11!==a.nodeType||r.isXMLDoc(a)))for(g=la(h),f=la(a),d=0,e=f.length;d<e;d++)Ga(f[d],g[d]);if(b)if(c)for(f=f||la(a),g=g||la(h),d=0,e=f.length;d<e;d++)Fa(f[d],g[d]);else Fa(a,h);return g=la(h,"script"),g.length>0&&ma(g,!i&&la(a,"script")),h},cleanData:function(a){for(var b,c,d,e=r.event.special,f=0;void 0!==(c=a[f]);f++)if(T(c)){if(b=c[V.expando]){if(b.events)for(d in b.events)e[d]?r.event.remove(c,d):r.removeEvent(c,d,b.handle);c[V.expando]=void 0}c[W.expando]&&(c[W.expando]=void 0)}}}),r.fn.extend({detach:function(a){return Ia(this,a,!0)},remove:function(a){return Ia(this,a)},text:function(a){return S(this,function(a){return void 0===a?r.text(this):this.empty().each(function(){1!==this.nodeType&&11!==this.nodeType&&9!==this.nodeType||(this.textContent=a)})},null,a,arguments.length)},append:function(){return Ha(this,arguments,function(a){if(1===this.nodeType||11===this.nodeType||9===this.nodeType){var b=Ca(this,a);b.appendChild(a)}})},prepend:function(){return Ha(this,arguments,function(a){if(1===this.nodeType||11===this.nodeType||9===this.nodeType){var b=Ca(this,a);b.insertBefore(a,b.firstChild)}})},before:function(){return Ha(this,arguments,function(a){this.parentNode&&this.parentNode.insertBefore(a,this)})},after:function(){return Ha(this,arguments,function(a){this.parentNode&&this.parentNode.insertBefore(a,this.nextSibling)})},empty:function(){for(var a,b=0;null!=(a=this[b]);b++)1===a.nodeType&&(r.cleanData(la(a,!1)),a.textContent="");return this},clone:function(a,b){return a=null!=a&&a,b=null==b?a:b,this.map(function(){return r.clone(this,a,b)})},html:function(a){return S(this,function(a){var b=this[0]||{},c=0,d=this.length;if(void 0===a&&1===b.nodeType)return b.innerHTML;if("string"==typeof a&&!ya.test(a)&&!ka[(ia.exec(a)||["",""])[1].toLowerCase()]){a=r.htmlPrefilter(a);try{for(;c<d;c++)b=this[c]||{},1===b.nodeType&&(r.cleanData(la(b,!1)),b.innerHTML=a);b=0}catch(e){}}b&&this.empty().append(a)},null,a,arguments.length)},replaceWith:function(){var a=[];return Ha(this,arguments,function(b){var c=this.parentNode;r.inArray(this,a)<0&&(r.cleanData(la(this)),c&&c.replaceChild(b,this))},a)}}),r.each({appendTo:"append",prependTo:"prepend",insertBefore:"before",insertAfter:"after",replaceAll:"replaceWith"},function(a,b){r.fn[a]=function(a){for(var c,d=[],e=r(a),f=e.length-1,g=0;g<=f;g++)c=g===f?this:this.clone(!0),r(e[g])[b](c),h.apply(d,c.get());return this.pushStack(d)}});var Ja=/^margin/,Ka=new RegExp("^("+$+")(?!px)[a-z%]+$","i"),La=function(b){var c=b.ownerDocument.defaultView;return c&&c.opener||(c=a),c.getComputedStyle(b)};!function(){function b(){if(i){i.style.cssText="box-sizing:border-box;position:relative;display:block;margin:auto;border:1px;padding:1px;top:1%;width:50%",i.innerHTML="",pa.appendChild(h);var b=a.getComputedStyle(i);c="1%"!==b.top,g="2px"===b.marginLeft,e="4px"===b.width,i.style.marginRight="50%",f="4px"===b.marginRight,pa.removeChild(h),i=null}}var c,e,f,g,h=d.createElement("div"),i=d.createElement("div");i.style&&(i.style.backgroundClip="content-box",i.cloneNode(!0).style.backgroundClip="",o.clearCloneStyle="content-box"===i.style.backgroundClip,h.style.cssText="border:0;width:8px;height:0;top:0;left:-9999px;padding:0;margin-top:1px;position:absolute",h.appendChild(i),r.extend(o,{pixelPosition:function(){return b(),c},boxSizingReliable:function(){return b(),e},pixelMarginRight:function(){return b(),f},reliableMarginLeft:function(){return b(),g}}))}();function Ma(a,b,c){var d,e,f,g,h=a.style;return c=c||La(a),c&&(g=c.getPropertyValue(b)||c[b],""!==g||r.contains(a.ownerDocument,a)||(g=r.style(a,b)),!o.pixelMarginRight()&&Ka.test(g)&&Ja.test(b)&&(d=h.width,e=h.minWidth,f=h.maxWidth,h.minWidth=h.maxWidth=h.width=g,g=c.width,h.width=d,h.minWidth=e,h.maxWidth=f)),void 0!==g?g+"":g}function Na(a,b){return{get:function(){return a()?void delete this.get:(this.get=b).apply(this,arguments)}}}var Oa=/^(none|table(?!-c[ea]).+)/,Pa={position:"absolute",visibility:"hidden",display:"block"},Qa={letterSpacing:"0",fontWeight:"400"},Ra=["Webkit","Moz","ms"],Sa=d.createElement("div").style;function Ta(a){if(a in Sa)return a;var b=a[0].toUpperCase()+a.slice(1),c=Ra.length;while(c--)if(a=Ra[c]+b,a in Sa)return a}function Ua(a,b,c){var d=_.exec(b);return d?Math.max(0,d[2]-(c||0))+(d[3]||"px"):b}function Va(a,b,c,d,e){for(var f=c===(d?"border":"content")?4:"width"===b?1:0,g=0;f<4;f+=2)"margin"===c&&(g+=r.css(a,c+aa[f],!0,e)),d?("content"===c&&(g-=r.css(a,"padding"+aa[f],!0,e)),"margin"!==c&&(g-=r.css(a,"border"+aa[f]+"Width",!0,e))):(g+=r.css(a,"padding"+aa[f],!0,e),"padding"!==c&&(g+=r.css(a,"border"+aa[f]+"Width",!0,e)));return g}function Wa(a,b,c){var d,e=!0,f=La(a),g="border-box"===r.css(a,"boxSizing",!1,f);if(a.getClientRects().length&&(d=a.getBoundingClientRect()[b]),d<=0||null==d){if(d=Ma(a,b,f),(d<0||null==d)&&(d=a.style[b]),Ka.test(d))return d;e=g&&(o.boxSizingReliable()||d===a.style[b]),d=parseFloat(d)||0}return d+Va(a,b,c||(g?"border":"content"),e,f)+"px"}r.extend({cssHooks:{opacity:{get:function(a,b){if(b){var c=Ma(a,"opacity");return""===c?"1":c}}}},cssNumber:{animationIterationCount:!0,columnCount:!0,fillOpacity:!0,flexGrow:!0,flexShrink:!0,fontWeight:!0,lineHeight:!0,opacity:!0,order:!0,orphans:!0,widows:!0,zIndex:!0,zoom:!0},cssProps:{"float":"cssFloat"},style:function(a,b,c,d){if(a&&3!==a.nodeType&&8!==a.nodeType&&a.style){var e,f,g,h=r.camelCase(b),i=a.style;return b=r.cssProps[h]||(r.cssProps[h]=Ta(h)||h),g=r.cssHooks[b]||r.cssHooks[h],void 0===c?g&&"get"in g&&void 0!==(e=g.get(a,!1,d))?e:i[b]:(f=typeof c,"string"===f&&(e=_.exec(c))&&e[1]&&(c=da(a,b,e),f="number"),null!=c&&c===c&&("number"===f&&(c+=e&&e[3]||(r.cssNumber[h]?"":"px")),o.clearCloneStyle||""!==c||0!==b.indexOf("background")||(i[b]="inherit"),g&&"set"in g&&void 0===(c=g.set(a,c,d))||(i[b]=c)),void 0)}},css:function(a,b,c,d){var e,f,g,h=r.camelCase(b);return b=r.cssProps[h]||(r.cssProps[h]=Ta(h)||h),g=r.cssHooks[b]||r.cssHooks[h],g&&"get"in g&&(e=g.get(a,!0,c)),void 0===e&&(e=Ma(a,b,d)),"normal"===e&&b in Qa&&(e=Qa[b]),""===c||c?(f=parseFloat(e),c===!0||isFinite(f)?f||0:e):e}}),r.each(["height","width"],function(a,b){r.cssHooks[b]={get:function(a,c,d){if(c)return!Oa.test(r.css(a,"display"))||a.getClientRects().length&&a.getBoundingClientRect().width?Wa(a,b,d):ca(a,Pa,function(){return Wa(a,b,d)})},set:function(a,c,d){var e,f=d&&La(a),g=d&&Va(a,b,d,"border-box"===r.css(a,"boxSizing",!1,f),f);return g&&(e=_.exec(c))&&"px"!==(e[3]||"px")&&(a.style[b]=c,c=r.css(a,b)),Ua(a,c,g)}}}),r.cssHooks.marginLeft=Na(o.reliableMarginLeft,function(a,b){if(b)return(parseFloat(Ma(a,"marginLeft"))||a.getBoundingClientRect().left-ca(a,{marginLeft:0},function(){return a.getBoundingClientRect().left}))+"px"}),r.each({margin:"",padding:"",border:"Width"},function(a,b){r.cssHooks[a+b]={expand:function(c){for(var d=0,e={},f="string"==typeof c?c.split(" "):[c];d<4;d++)e[a+aa[d]+b]=f[d]||f[d-2]||f[0];return e}},Ja.test(a)||(r.cssHooks[a+b].set=Ua)}),r.fn.extend({css:function(a,b){return S(this,function(a,b,c){var d,e,f={},g=0;if(r.isArray(b)){for(d=La(a),e=b.length;g<e;g++)f[b[g]]=r.css(a,b[g],!1,d);return f}return void 0!==c?r.style(a,b,c):r.css(a,b)},a,b,arguments.length>1)}});function Xa(a,b,c,d,e){return new Xa.prototype.init(a,b,c,d,e)}r.Tween=Xa,Xa.prototype={constructor:Xa,init:function(a,b,c,d,e,f){this.elem=a,this.prop=c,this.easing=e||r.easing._default,this.options=b,this.start=this.now=this.cur(),this.end=d,this.unit=f||(r.cssNumber[c]?"":"px")},cur:function(){var a=Xa.propHooks[this.prop];return a&&a.get?a.get(this):Xa.propHooks._default.get(this)},run:function(a){var b,c=Xa.propHooks[this.prop];return this.options.duration?this.pos=b=r.easing[this.easing](a,this.options.duration*a,0,1,this.options.duration):this.pos=b=a,this.now=(this.end-this.start)*b+this.start,this.options.step&&this.options.step.call(this.elem,this.now,this),c&&c.set?c.set(this):Xa.propHooks._default.set(this),this}},Xa.prototype.init.prototype=Xa.prototype,Xa.propHooks={_default:{get:function(a){var b;return 1!==a.elem.nodeType||null!=a.elem[a.prop]&&null==a.elem.style[a.prop]?a.elem[a.prop]:(b=r.css(a.elem,a.prop,""),b&&"auto"!==b?b:0)},set:function(a){r.fx.step[a.prop]?r.fx.step[a.prop](a):1!==a.elem.nodeType||null==a.elem.style[r.cssProps[a.prop]]&&!r.cssHooks[a.prop]?a.elem[a.prop]=a.now:r.style(a.elem,a.prop,a.now+a.unit)}}},Xa.propHooks.scrollTop=Xa.propHooks.scrollLeft={set:function(a){a.elem.nodeType&&a.elem.parentNode&&(a.elem[a.prop]=a.now)}},r.easing={linear:function(a){return a},swing:function(a){return.5-Math.cos(a*Math.PI)/2},_default:"swing"},r.fx=Xa.prototype.init,r.fx.step={};var Ya,Za,$a=/^(?:toggle|show|hide)$/,_a=/queueHooks$/;function ab(){Za&&(a.requestAnimationFrame(ab),r.fx.tick())}function bb(){return a.setTimeout(function(){Ya=void 0}),Ya=r.now()}function cb(a,b){var c,d=0,e={height:a};for(b=b?1:0;d<4;d+=2-b)c=aa[d],e["margin"+c]=e["padding"+c]=a;return b&&(e.opacity=e.width=a),e}function db(a,b,c){for(var d,e=(gb.tweeners[b]||[]).concat(gb.tweeners["*"]),f=0,g=e.length;f<g;f++)if(d=e[f].call(c,b,a))return d}function eb(a,b,c){var d,e,f,g,h,i,j,k,l="width"in b||"height"in b,m=this,n={},o=a.style,p=a.nodeType&&ba(a),q=V.get(a,"fxshow");c.queue||(g=r._queueHooks(a,"fx"),null==g.unqueued&&(g.unqueued=0,h=g.empty.fire,g.empty.fire=function(){g.unqueued||h()}),g.unqueued++,m.always(function(){m.always(function(){g.unqueued--,r.queue(a,"fx").length||g.empty.fire()})}));for(d in b)if(e=b[d],$a.test(e)){if(delete b[d],f=f||"toggle"===e,e===(p?"hide":"show")){if("show"!==e||!q||void 0===q[d])continue;p=!0}n[d]=q&&q[d]||r.style(a,d)}if(i=!r.isEmptyObject(b),i||!r.isEmptyObject(n)){l&&1===a.nodeType&&(c.overflow=[o.overflow,o.overflowX,o.overflowY],j=q&&q.display,null==j&&(j=V.get(a,"display")),k=r.css(a,"display"),"none"===k&&(j?k=j:(ga([a],!0),j=a.style.display||j,k=r.css(a,"display"),ga([a]))),("inline"===k||"inline-block"===k&&null!=j)&&"none"===r.css(a,"float")&&(i||(m.done(function(){o.display=j}),null==j&&(k=o.display,j="none"===k?"":k)),o.display="inline-block")),c.overflow&&(o.overflow="hidden",m.always(function(){o.overflow=c.overflow[0],o.overflowX=c.overflow[1],o.overflowY=c.overflow[2]})),i=!1;for(d in n)i||(q?"hidden"in q&&(p=q.hidden):q=V.access(a,"fxshow",{display:j}),f&&(q.hidden=!p),p&&ga([a],!0),m.done(function(){p||ga([a]),V.remove(a,"fxshow");for(d in n)r.style(a,d,n[d])})),i=db(p?q[d]:0,d,m),d in q||(q[d]=i.start,p&&(i.end=i.start,i.start=0))}}function fb(a,b){var c,d,e,f,g;for(c in a)if(d=r.camelCase(c),e=b[d],f=a[c],r.isArray(f)&&(e=f[1],f=a[c]=f[0]),c!==d&&(a[d]=f,delete a[c]),g=r.cssHooks[d],g&&"expand"in g){f=g.expand(f),delete a[d];for(c in f)c in a||(a[c]=f[c],b[c]=e)}else b[d]=e}function gb(a,b,c){var d,e,f=0,g=gb.prefilters.length,h=r.Deferred().always(function(){delete i.elem}),i=function(){if(e)return!1;for(var b=Ya||bb(),c=Math.max(0,j.startTime+j.duration-b),d=c/j.duration||0,f=1-d,g=0,i=j.tweens.length;g<i;g++)j.tweens[g].run(f);return h.notifyWith(a,[j,f,c]),f<1&&i?c:(h.resolveWith(a,[j]),!1)},j=h.promise({elem:a,props:r.extend({},b),opts:r.extend(!0,{specialEasing:{},easing:r.easing._default},c),originalProperties:b,originalOptions:c,startTime:Ya||bb(),duration:c.duration,tweens:[],createTween:function(b,c){var d=r.Tween(a,j.opts,b,c,j.opts.specialEasing[b]||j.opts.easing);return j.tweens.push(d),d},stop:function(b){var c=0,d=b?j.tweens.length:0;if(e)return this;for(e=!0;c<d;c++)j.tweens[c].run(1);return b?(h.notifyWith(a,[j,1,0]),h.resolveWith(a,[j,b])):h.rejectWith(a,[j,b]),this}}),k=j.props;for(fb(k,j.opts.specialEasing);f<g;f++)if(d=gb.prefilters[f].call(j,a,k,j.opts))return r.isFunction(d.stop)&&(r._queueHooks(j.elem,j.opts.queue).stop=r.proxy(d.stop,d)),d;return r.map(k,db,j),r.isFunction(j.opts.start)&&j.opts.start.call(a,j),r.fx.timer(r.extend(i,{elem:a,anim:j,queue:j.opts.queue})),j.progress(j.opts.progress).done(j.opts.done,j.opts.complete).fail(j.opts.fail).always(j.opts.always)}r.Animation=r.extend(gb,{tweeners:{"*":[function(a,b){var c=this.createTween(a,b);return da(c.elem,a,_.exec(b),c),c}]},tweener:function(a,b){r.isFunction(a)?(b=a,a=["*"]):a=a.match(K);for(var c,d=0,e=a.length;d<e;d++)c=a[d],gb.tweeners[c]=gb.tweeners[c]||[],gb.tweeners[c].unshift(b)},prefilters:[eb],prefilter:function(a,b){b?gb.prefilters.unshift(a):gb.prefilters.push(a)}}),r.speed=function(a,b,c){var e=a&&"object"==typeof a?r.extend({},a):{complete:c||!c&&b||r.isFunction(a)&&a,duration:a,easing:c&&b||b&&!r.isFunction(b)&&b};return r.fx.off||d.hidden?e.duration=0:e.duration="number"==typeof e.duration?e.duration:e.duration in r.fx.speeds?r.fx.speeds[e.duration]:r.fx.speeds._default,null!=e.queue&&e.queue!==!0||(e.queue="fx"),e.old=e.complete,e.complete=function(){r.isFunction(e.old)&&e.old.call(this),e.queue&&r.dequeue(this,e.queue)},e},r.fn.extend({fadeTo:function(a,b,c,d){return this.filter(ba).css("opacity",0).show().end().animate({opacity:b},a,c,d)},animate:function(a,b,c,d){var e=r.isEmptyObject(a),f=r.speed(b,c,d),g=function(){var b=gb(this,r.extend({},a),f);(e||V.get(this,"finish"))&&b.stop(!0)};return g.finish=g,e||f.queue===!1?this.each(g):this.queue(f.queue,g)},stop:function(a,b,c){var d=function(a){var b=a.stop;delete a.stop,b(c)};return"string"!=typeof a&&(c=b,b=a,a=void 0),b&&a!==!1&&this.queue(a||"fx",[]),this.each(function(){var b=!0,e=null!=a&&a+"queueHooks",f=r.timers,g=V.get(this);if(e)g[e]&&g[e].stop&&d(g[e]);else for(e in g)g[e]&&g[e].stop&&_a.test(e)&&d(g[e]);for(e=f.length;e--;)f[e].elem!==this||null!=a&&f[e].queue!==a||(f[e].anim.stop(c),b=!1,f.splice(e,1));!b&&c||r.dequeue(this,a)})},finish:function(a){return a!==!1&&(a=a||"fx"),this.each(function(){var b,c=V.get(this),d=c[a+"queue"],e=c[a+"queueHooks"],f=r.timers,g=d?d.length:0;for(c.finish=!0,r.queue(this,a,[]),e&&e.stop&&e.stop.call(this,!0),b=f.length;b--;)f[b].elem===this&&f[b].queue===a&&(f[b].anim.stop(!0),f.splice(b,1));for(b=0;b<g;b++)d[b]&&d[b].finish&&d[b].finish.call(this);delete c.finish})}}),r.each(["toggle","show","hide"],function(a,b){var c=r.fn[b];r.fn[b]=function(a,d,e){return null==a||"boolean"==typeof a?c.apply(this,arguments):this.animate(cb(b,!0),a,d,e)}}),r.each({slideDown:cb("show"),slideUp:cb("hide"),slideToggle:cb("toggle"),fadeIn:{opacity:"show"},fadeOut:{opacity:"hide"},fadeToggle:{opacity:"toggle"}},function(a,b){r.fn[a]=function(a,c,d){return this.animate(b,a,c,d)}}),r.timers=[],r.fx.tick=function(){var a,b=0,c=r.timers;for(Ya=r.now();b<c.length;b++)a=c[b],a()||c[b]!==a||c.splice(b--,1);c.length||r.fx.stop(),Ya=void 0},r.fx.timer=function(a){r.timers.push(a),a()?r.fx.start():r.timers.pop()},r.fx.interval=13,r.fx.start=function(){Za||(Za=a.requestAnimationFrame?a.requestAnimationFrame(ab):a.setInterval(r.fx.tick,r.fx.interval))},r.fx.stop=function(){a.cancelAnimationFrame?a.cancelAnimationFrame(Za):a.clearInterval(Za),Za=null},r.fx.speeds={slow:600,fast:200,_default:400},r.fn.delay=function(b,c){return b=r.fx?r.fx.speeds[b]||b:b,c=c||"fx",this.queue(c,function(c,d){var e=a.setTimeout(c,b);d.stop=function(){a.clearTimeout(e)}})},function(){var a=d.createElement("input"),b=d.createElement("select"),c=b.appendChild(d.createElement("option"));a.type="checkbox",o.checkOn=""!==a.value,o.optSelected=c.selected,a=d.createElement("input"),a.value="t",a.type="radio",o.radioValue="t"===a.value}();var hb,ib=r.expr.attrHandle;r.fn.extend({attr:function(a,b){return S(this,r.attr,a,b,arguments.length>1)},removeAttr:function(a){return this.each(function(){r.removeAttr(this,a)})}}),r.extend({attr:function(a,b,c){var d,e,f=a.nodeType;if(3!==f&&8!==f&&2!==f)return"undefined"==typeof a.getAttribute?r.prop(a,b,c):(1===f&&r.isXMLDoc(a)||(e=r.attrHooks[b.toLowerCase()]||(r.expr.match.bool.test(b)?hb:void 0)),void 0!==c?null===c?void r.removeAttr(a,b):e&&"set"in e&&void 0!==(d=e.set(a,c,b))?d:(a.setAttribute(b,c+""),c):e&&"get"in e&&null!==(d=e.get(a,b))?d:(d=r.find.attr(a,b),null==d?void 0:d))},attrHooks:{type:{set:function(a,b){if(!o.radioValue&&"radio"===b&&r.nodeName(a,"input")){var c=a.value;return a.setAttribute("type",b),c&&(a.value=c),b}}}},removeAttr:function(a,b){var c,d=0,e=b&&b.match(K);
if(e&&1===a.nodeType)while(c=e[d++])a.removeAttribute(c)}}),hb={set:function(a,b,c){return b===!1?r.removeAttr(a,c):a.setAttribute(c,c),c}},r.each(r.expr.match.bool.source.match(/\w+/g),function(a,b){var c=ib[b]||r.find.attr;ib[b]=function(a,b,d){var e,f,g=b.toLowerCase();return d||(f=ib[g],ib[g]=e,e=null!=c(a,b,d)?g:null,ib[g]=f),e}});var jb=/^(?:input|select|textarea|button)$/i,kb=/^(?:a|area)$/i;r.fn.extend({prop:function(a,b){return S(this,r.prop,a,b,arguments.length>1)},removeProp:function(a){return this.each(function(){delete this[r.propFix[a]||a]})}}),r.extend({prop:function(a,b,c){var d,e,f=a.nodeType;if(3!==f&&8!==f&&2!==f)return 1===f&&r.isXMLDoc(a)||(b=r.propFix[b]||b,e=r.propHooks[b]),void 0!==c?e&&"set"in e&&void 0!==(d=e.set(a,c,b))?d:a[b]=c:e&&"get"in e&&null!==(d=e.get(a,b))?d:a[b]},propHooks:{tabIndex:{get:function(a){var b=r.find.attr(a,"tabindex");return b?parseInt(b,10):jb.test(a.nodeName)||kb.test(a.nodeName)&&a.href?0:-1}}},propFix:{"for":"htmlFor","class":"className"}}),o.optSelected||(r.propHooks.selected={get:function(a){var b=a.parentNode;return b&&b.parentNode&&b.parentNode.selectedIndex,null},set:function(a){var b=a.parentNode;b&&(b.selectedIndex,b.parentNode&&b.parentNode.selectedIndex)}}),r.each(["tabIndex","readOnly","maxLength","cellSpacing","cellPadding","rowSpan","colSpan","useMap","frameBorder","contentEditable"],function(){r.propFix[this.toLowerCase()]=this});var lb=/[\t\r\n\f]/g;function mb(a){return a.getAttribute&&a.getAttribute("class")||""}r.fn.extend({addClass:function(a){var b,c,d,e,f,g,h,i=0;if(r.isFunction(a))return this.each(function(b){r(this).addClass(a.call(this,b,mb(this)))});if("string"==typeof a&&a){b=a.match(K)||[];while(c=this[i++])if(e=mb(c),d=1===c.nodeType&&(" "+e+" ").replace(lb," ")){g=0;while(f=b[g++])d.indexOf(" "+f+" ")<0&&(d+=f+" ");h=r.trim(d),e!==h&&c.setAttribute("class",h)}}return this},removeClass:function(a){var b,c,d,e,f,g,h,i=0;if(r.isFunction(a))return this.each(function(b){r(this).removeClass(a.call(this,b,mb(this)))});if(!arguments.length)return this.attr("class","");if("string"==typeof a&&a){b=a.match(K)||[];while(c=this[i++])if(e=mb(c),d=1===c.nodeType&&(" "+e+" ").replace(lb," ")){g=0;while(f=b[g++])while(d.indexOf(" "+f+" ")>-1)d=d.replace(" "+f+" "," ");h=r.trim(d),e!==h&&c.setAttribute("class",h)}}return this},toggleClass:function(a,b){var c=typeof a;return"boolean"==typeof b&&"string"===c?b?this.addClass(a):this.removeClass(a):r.isFunction(a)?this.each(function(c){r(this).toggleClass(a.call(this,c,mb(this),b),b)}):this.each(function(){var b,d,e,f;if("string"===c){d=0,e=r(this),f=a.match(K)||[];while(b=f[d++])e.hasClass(b)?e.removeClass(b):e.addClass(b)}else void 0!==a&&"boolean"!==c||(b=mb(this),b&&V.set(this,"__className__",b),this.setAttribute&&this.setAttribute("class",b||a===!1?"":V.get(this,"__className__")||""))})},hasClass:function(a){var b,c,d=0;b=" "+a+" ";while(c=this[d++])if(1===c.nodeType&&(" "+mb(c)+" ").replace(lb," ").indexOf(b)>-1)return!0;return!1}});var nb=/\r/g,ob=/[\x20\t\r\n\f]+/g;r.fn.extend({val:function(a){var b,c,d,e=this[0];{if(arguments.length)return d=r.isFunction(a),this.each(function(c){var e;1===this.nodeType&&(e=d?a.call(this,c,r(this).val()):a,null==e?e="":"number"==typeof e?e+="":r.isArray(e)&&(e=r.map(e,function(a){return null==a?"":a+""})),b=r.valHooks[this.type]||r.valHooks[this.nodeName.toLowerCase()],b&&"set"in b&&void 0!==b.set(this,e,"value")||(this.value=e))});if(e)return b=r.valHooks[e.type]||r.valHooks[e.nodeName.toLowerCase()],b&&"get"in b&&void 0!==(c=b.get(e,"value"))?c:(c=e.value,"string"==typeof c?c.replace(nb,""):null==c?"":c)}}}),r.extend({valHooks:{option:{get:function(a){var b=r.find.attr(a,"value");return null!=b?b:r.trim(r.text(a)).replace(ob," ")}},select:{get:function(a){for(var b,c,d=a.options,e=a.selectedIndex,f="select-one"===a.type,g=f?null:[],h=f?e+1:d.length,i=e<0?h:f?e:0;i<h;i++)if(c=d[i],(c.selected||i===e)&&!c.disabled&&(!c.parentNode.disabled||!r.nodeName(c.parentNode,"optgroup"))){if(b=r(c).val(),f)return b;g.push(b)}return g},set:function(a,b){var c,d,e=a.options,f=r.makeArray(b),g=e.length;while(g--)d=e[g],(d.selected=r.inArray(r.valHooks.option.get(d),f)>-1)&&(c=!0);return c||(a.selectedIndex=-1),f}}}}),r.each(["radio","checkbox"],function(){r.valHooks[this]={set:function(a,b){if(r.isArray(b))return a.checked=r.inArray(r(a).val(),b)>-1}},o.checkOn||(r.valHooks[this].get=function(a){return null===a.getAttribute("value")?"on":a.value})});var pb=/^(?:focusinfocus|focusoutblur)$/;r.extend(r.event,{trigger:function(b,c,e,f){var g,h,i,j,k,m,n,o=[e||d],p=l.call(b,"type")?b.type:b,q=l.call(b,"namespace")?b.namespace.split("."):[];if(h=i=e=e||d,3!==e.nodeType&&8!==e.nodeType&&!pb.test(p+r.event.triggered)&&(p.indexOf(".")>-1&&(q=p.split("."),p=q.shift(),q.sort()),k=p.indexOf(":")<0&&"on"+p,b=b[r.expando]?b:new r.Event(p,"object"==typeof b&&b),b.isTrigger=f?2:3,b.namespace=q.join("."),b.rnamespace=b.namespace?new RegExp("(^|\\.)"+q.join("\\.(?:.*\\.|)")+"(\\.|$)"):null,b.result=void 0,b.target||(b.target=e),c=null==c?[b]:r.makeArray(c,[b]),n=r.event.special[p]||{},f||!n.trigger||n.trigger.apply(e,c)!==!1)){if(!f&&!n.noBubble&&!r.isWindow(e)){for(j=n.delegateType||p,pb.test(j+p)||(h=h.parentNode);h;h=h.parentNode)o.push(h),i=h;i===(e.ownerDocument||d)&&o.push(i.defaultView||i.parentWindow||a)}g=0;while((h=o[g++])&&!b.isPropagationStopped())b.type=g>1?j:n.bindType||p,m=(V.get(h,"events")||{})[b.type]&&V.get(h,"handle"),m&&m.apply(h,c),m=k&&h[k],m&&m.apply&&T(h)&&(b.result=m.apply(h,c),b.result===!1&&b.preventDefault());return b.type=p,f||b.isDefaultPrevented()||n._default&&n._default.apply(o.pop(),c)!==!1||!T(e)||k&&r.isFunction(e[p])&&!r.isWindow(e)&&(i=e[k],i&&(e[k]=null),r.event.triggered=p,e[p](),r.event.triggered=void 0,i&&(e[k]=i)),b.result}},simulate:function(a,b,c){var d=r.extend(new r.Event,c,{type:a,isSimulated:!0});r.event.trigger(d,null,b)}}),r.fn.extend({trigger:function(a,b){return this.each(function(){r.event.trigger(a,b,this)})},triggerHandler:function(a,b){var c=this[0];if(c)return r.event.trigger(a,b,c,!0)}}),r.each("blur focus focusin focusout resize scroll click dblclick mousedown mouseup mousemove mouseover mouseout mouseenter mouseleave change select submit keydown keypress keyup contextmenu".split(" "),function(a,b){r.fn[b]=function(a,c){return arguments.length>0?this.on(b,null,a,c):this.trigger(b)}}),r.fn.extend({hover:function(a,b){return this.mouseenter(a).mouseleave(b||a)}}),o.focusin="onfocusin"in a,o.focusin||r.each({focus:"focusin",blur:"focusout"},function(a,b){var c=function(a){r.event.simulate(b,a.target,r.event.fix(a))};r.event.special[b]={setup:function(){var d=this.ownerDocument||this,e=V.access(d,b);e||d.addEventListener(a,c,!0),V.access(d,b,(e||0)+1)},teardown:function(){var d=this.ownerDocument||this,e=V.access(d,b)-1;e?V.access(d,b,e):(d.removeEventListener(a,c,!0),V.remove(d,b))}}});var qb=a.location,rb=r.now(),sb=/\?/;r.parseXML=function(b){var c;if(!b||"string"!=typeof b)return null;try{c=(new a.DOMParser).parseFromString(b,"text/xml")}catch(d){c=void 0}return c&&!c.getElementsByTagName("parsererror").length||r.error("Invalid XML: "+b),c};var tb=/\[\]$/,ub=/\r?\n/g,vb=/^(?:submit|button|image|reset|file)$/i,wb=/^(?:input|select|textarea|keygen)/i;function xb(a,b,c,d){var e;if(r.isArray(b))r.each(b,function(b,e){c||tb.test(a)?d(a,e):xb(a+"["+("object"==typeof e&&null!=e?b:"")+"]",e,c,d)});else if(c||"object"!==r.type(b))d(a,b);else for(e in b)xb(a+"["+e+"]",b[e],c,d)}r.param=function(a,b){var c,d=[],e=function(a,b){var c=r.isFunction(b)?b():b;d[d.length]=encodeURIComponent(a)+"="+encodeURIComponent(null==c?"":c)};if(r.isArray(a)||a.jquery&&!r.isPlainObject(a))r.each(a,function(){e(this.name,this.value)});else for(c in a)xb(c,a[c],b,e);return d.join("&")},r.fn.extend({serialize:function(){return r.param(this.serializeArray())},serializeArray:function(){return this.map(function(){var a=r.prop(this,"elements");return a?r.makeArray(a):this}).filter(function(){var a=this.type;return this.name&&!r(this).is(":disabled")&&wb.test(this.nodeName)&&!vb.test(a)&&(this.checked||!ha.test(a))}).map(function(a,b){var c=r(this).val();return null==c?null:r.isArray(c)?r.map(c,function(a){return{name:b.name,value:a.replace(ub,"\r\n")}}):{name:b.name,value:c.replace(ub,"\r\n")}}).get()}});var yb=/%20/g,zb=/#.*$/,Ab=/([?&])_=[^&]*/,Bb=/^(.*?):[ \t]*([^\r\n]*)$/gm,Cb=/^(?:about|app|app-storage|.+-extension|file|res|widget):$/,Db=/^(?:GET|HEAD)$/,Eb=/^\/\//,Fb={},Gb={},Hb="*/".concat("*"),Ib=d.createElement("a");Ib.href=qb.href;function Jb(a){return function(b,c){"string"!=typeof b&&(c=b,b="*");var d,e=0,f=b.toLowerCase().match(K)||[];if(r.isFunction(c))while(d=f[e++])"+"===d[0]?(d=d.slice(1)||"*",(a[d]=a[d]||[]).unshift(c)):(a[d]=a[d]||[]).push(c)}}function Kb(a,b,c,d){var e={},f=a===Gb;function g(h){var i;return e[h]=!0,r.each(a[h]||[],function(a,h){var j=h(b,c,d);return"string"!=typeof j||f||e[j]?f?!(i=j):void 0:(b.dataTypes.unshift(j),g(j),!1)}),i}return g(b.dataTypes[0])||!e["*"]&&g("*")}function Lb(a,b){var c,d,e=r.ajaxSettings.flatOptions||{};for(c in b)void 0!==b[c]&&((e[c]?a:d||(d={}))[c]=b[c]);return d&&r.extend(!0,a,d),a}function Mb(a,b,c){var d,e,f,g,h=a.contents,i=a.dataTypes;while("*"===i[0])i.shift(),void 0===d&&(d=a.mimeType||b.getResponseHeader("Content-Type"));if(d)for(e in h)if(h[e]&&h[e].test(d)){i.unshift(e);break}if(i[0]in c)f=i[0];else{for(e in c){if(!i[0]||a.converters[e+" "+i[0]]){f=e;break}g||(g=e)}f=f||g}if(f)return f!==i[0]&&i.unshift(f),c[f]}function Nb(a,b,c,d){var e,f,g,h,i,j={},k=a.dataTypes.slice();if(k[1])for(g in a.converters)j[g.toLowerCase()]=a.converters[g];f=k.shift();while(f)if(a.responseFields[f]&&(c[a.responseFields[f]]=b),!i&&d&&a.dataFilter&&(b=a.dataFilter(b,a.dataType)),i=f,f=k.shift())if("*"===f)f=i;else if("*"!==i&&i!==f){if(g=j[i+" "+f]||j["* "+f],!g)for(e in j)if(h=e.split(" "),h[1]===f&&(g=j[i+" "+h[0]]||j["* "+h[0]])){g===!0?g=j[e]:j[e]!==!0&&(f=h[0],k.unshift(h[1]));break}if(g!==!0)if(g&&a["throws"])b=g(b);else try{b=g(b)}catch(l){return{state:"parsererror",error:g?l:"No conversion from "+i+" to "+f}}}return{state:"success",data:b}}r.extend({active:0,lastModified:{},etag:{},ajaxSettings:{url:qb.href,type:"GET",isLocal:Cb.test(qb.protocol),global:!0,processData:!0,async:!0,contentType:"application/x-www-form-urlencoded; charset=UTF-8",accepts:{"*":Hb,text:"text/plain",html:"text/html",xml:"application/xml, text/xml",json:"application/json, text/javascript"},contents:{xml:/\bxml\b/,html:/\bhtml/,json:/\bjson\b/},responseFields:{xml:"responseXML",text:"responseText",json:"responseJSON"},converters:{"* text":String,"text html":!0,"text json":JSON.parse,"text xml":r.parseXML},flatOptions:{url:!0,context:!0}},ajaxSetup:function(a,b){return b?Lb(Lb(a,r.ajaxSettings),b):Lb(r.ajaxSettings,a)},ajaxPrefilter:Jb(Fb),ajaxTransport:Jb(Gb),ajax:function(b,c){"object"==typeof b&&(c=b,b=void 0),c=c||{};var e,f,g,h,i,j,k,l,m,n,o=r.ajaxSetup({},c),p=o.context||o,q=o.context&&(p.nodeType||p.jquery)?r(p):r.event,s=r.Deferred(),t=r.Callbacks("once memory"),u=o.statusCode||{},v={},w={},x="canceled",y={readyState:0,getResponseHeader:function(a){var b;if(k){if(!h){h={};while(b=Bb.exec(g))h[b[1].toLowerCase()]=b[2]}b=h[a.toLowerCase()]}return null==b?null:b},getAllResponseHeaders:function(){return k?g:null},setRequestHeader:function(a,b){return null==k&&(a=w[a.toLowerCase()]=w[a.toLowerCase()]||a,v[a]=b),this},overrideMimeType:function(a){return null==k&&(o.mimeType=a),this},statusCode:function(a){var b;if(a)if(k)y.always(a[y.status]);else for(b in a)u[b]=[u[b],a[b]];return this},abort:function(a){var b=a||x;return e&&e.abort(b),A(0,b),this}};if(s.promise(y),o.url=((b||o.url||qb.href)+"").replace(Eb,qb.protocol+"//"),o.type=c.method||c.type||o.method||o.type,o.dataTypes=(o.dataType||"*").toLowerCase().match(K)||[""],null==o.crossDomain){j=d.createElement("a");try{j.href=o.url,j.href=j.href,o.crossDomain=Ib.protocol+"//"+Ib.host!=j.protocol+"//"+j.host}catch(z){o.crossDomain=!0}}if(o.data&&o.processData&&"string"!=typeof o.data&&(o.data=r.param(o.data,o.traditional)),Kb(Fb,o,c,y),k)return y;l=r.event&&o.global,l&&0===r.active++&&r.event.trigger("ajaxStart"),o.type=o.type.toUpperCase(),o.hasContent=!Db.test(o.type),f=o.url.replace(zb,""),o.hasContent?o.data&&o.processData&&0===(o.contentType||"").indexOf("application/x-www-form-urlencoded")&&(o.data=o.data.replace(yb,"+")):(n=o.url.slice(f.length),o.data&&(f+=(sb.test(f)?"&":"?")+o.data,delete o.data),o.cache===!1&&(f=f.replace(Ab,""),n=(sb.test(f)?"&":"?")+"_="+rb++ +n),o.url=f+n),o.ifModified&&(r.lastModified[f]&&y.setRequestHeader("If-Modified-Since",r.lastModified[f]),r.etag[f]&&y.setRequestHeader("If-None-Match",r.etag[f])),(o.data&&o.hasContent&&o.contentType!==!1||c.contentType)&&y.setRequestHeader("Content-Type",o.contentType),y.setRequestHeader("Accept",o.dataTypes[0]&&o.accepts[o.dataTypes[0]]?o.accepts[o.dataTypes[0]]+("*"!==o.dataTypes[0]?", "+Hb+"; q=0.01":""):o.accepts["*"]);for(m in o.headers)y.setRequestHeader(m,o.headers[m]);if(o.beforeSend&&(o.beforeSend.call(p,y,o)===!1||k))return y.abort();if(x="abort",t.add(o.complete),y.done(o.success),y.fail(o.error),e=Kb(Gb,o,c,y)){if(y.readyState=1,l&&q.trigger("ajaxSend",[y,o]),k)return y;o.async&&o.timeout>0&&(i=a.setTimeout(function(){y.abort("timeout")},o.timeout));try{k=!1,e.send(v,A)}catch(z){if(k)throw z;A(-1,z)}}else A(-1,"No Transport");function A(b,c,d,h){var j,m,n,v,w,x=c;k||(k=!0,i&&a.clearTimeout(i),e=void 0,g=h||"",y.readyState=b>0?4:0,j=b>=200&&b<300||304===b,d&&(v=Mb(o,y,d)),v=Nb(o,v,y,j),j?(o.ifModified&&(w=y.getResponseHeader("Last-Modified"),w&&(r.lastModified[f]=w),w=y.getResponseHeader("etag"),w&&(r.etag[f]=w)),204===b||"HEAD"===o.type?x="nocontent":304===b?x="notmodified":(x=v.state,m=v.data,n=v.error,j=!n)):(n=x,!b&&x||(x="error",b<0&&(b=0))),y.status=b,y.statusText=(c||x)+"",j?s.resolveWith(p,[m,x,y]):s.rejectWith(p,[y,x,n]),y.statusCode(u),u=void 0,l&&q.trigger(j?"ajaxSuccess":"ajaxError",[y,o,j?m:n]),t.fireWith(p,[y,x]),l&&(q.trigger("ajaxComplete",[y,o]),--r.active||r.event.trigger("ajaxStop")))}return y},getJSON:function(a,b,c){return r.get(a,b,c,"json")},getScript:function(a,b){return r.get(a,void 0,b,"script")}}),r.each(["get","post"],function(a,b){r[b]=function(a,c,d,e){return r.isFunction(c)&&(e=e||d,d=c,c=void 0),r.ajax(r.extend({url:a,type:b,dataType:e,data:c,success:d},r.isPlainObject(a)&&a))}}),r._evalUrl=function(a){return r.ajax({url:a,type:"GET",dataType:"script",cache:!0,async:!1,global:!1,"throws":!0})},r.fn.extend({wrapAll:function(a){var b;return this[0]&&(r.isFunction(a)&&(a=a.call(this[0])),b=r(a,this[0].ownerDocument).eq(0).clone(!0),this[0].parentNode&&b.insertBefore(this[0]),b.map(function(){var a=this;while(a.firstElementChild)a=a.firstElementChild;return a}).append(this)),this},wrapInner:function(a){return r.isFunction(a)?this.each(function(b){r(this).wrapInner(a.call(this,b))}):this.each(function(){var b=r(this),c=b.contents();c.length?c.wrapAll(a):b.append(a)})},wrap:function(a){var b=r.isFunction(a);return this.each(function(c){r(this).wrapAll(b?a.call(this,c):a)})},unwrap:function(a){return this.parent(a).not("body").each(function(){r(this).replaceWith(this.childNodes)}),this}}),r.expr.pseudos.hidden=function(a){return!r.expr.pseudos.visible(a)},r.expr.pseudos.visible=function(a){return!!(a.offsetWidth||a.offsetHeight||a.getClientRects().length)},r.ajaxSettings.xhr=function(){try{return new a.XMLHttpRequest}catch(b){}};var Ob={0:200,1223:204},Pb=r.ajaxSettings.xhr();o.cors=!!Pb&&"withCredentials"in Pb,o.ajax=Pb=!!Pb,r.ajaxTransport(function(b){var c,d;if(o.cors||Pb&&!b.crossDomain)return{send:function(e,f){var g,h=b.xhr();if(h.open(b.type,b.url,b.async,b.username,b.password),b.xhrFields)for(g in b.xhrFields)h[g]=b.xhrFields[g];b.mimeType&&h.overrideMimeType&&h.overrideMimeType(b.mimeType),b.crossDomain||e["X-Requested-With"]||(e["X-Requested-With"]="XMLHttpRequest");for(g in e)h.setRequestHeader(g,e[g]);c=function(a){return function(){c&&(c=d=h.onload=h.onerror=h.onabort=h.onreadystatechange=null,"abort"===a?h.abort():"error"===a?"number"!=typeof h.status?f(0,"error"):f(h.status,h.statusText):f(Ob[h.status]||h.status,h.statusText,"text"!==(h.responseType||"text")||"string"!=typeof h.responseText?{binary:h.response}:{text:h.responseText},h.getAllResponseHeaders()))}},h.onload=c(),d=h.onerror=c("error"),void 0!==h.onabort?h.onabort=d:h.onreadystatechange=function(){4===h.readyState&&a.setTimeout(function(){c&&d()})},c=c("abort");try{h.send(b.hasContent&&b.data||null)}catch(i){if(c)throw i}},abort:function(){c&&c()}}}),r.ajaxPrefilter(function(a){a.crossDomain&&(a.contents.script=!1)}),r.ajaxSetup({accepts:{script:"text/javascript, application/javascript, application/ecmascript, application/x-ecmascript"},contents:{script:/\b(?:java|ecma)script\b/},converters:{"text script":function(a){return r.globalEval(a),a}}}),r.ajaxPrefilter("script",function(a){void 0===a.cache&&(a.cache=!1),a.crossDomain&&(a.type="GET")}),r.ajaxTransport("script",function(a){if(a.crossDomain){var b,c;return{send:function(e,f){b=r("<script>").prop({charset:a.scriptCharset,src:a.url}).on("load error",c=function(a){b.remove(),c=null,a&&f("error"===a.type?404:200,a.type)}),d.head.appendChild(b[0])},abort:function(){c&&c()}}}});var Qb=[],Rb=/(=)\?(?=&|$)|\?\?/;r.ajaxSetup({jsonp:"callback",jsonpCallback:function(){var a=Qb.pop()||r.expando+"_"+rb++;return this[a]=!0,a}}),r.ajaxPrefilter("json jsonp",function(b,c,d){var e,f,g,h=b.jsonp!==!1&&(Rb.test(b.url)?"url":"string"==typeof b.data&&0===(b.contentType||"").indexOf("application/x-www-form-urlencoded")&&Rb.test(b.data)&&"data");if(h||"jsonp"===b.dataTypes[0])return e=b.jsonpCallback=r.isFunction(b.jsonpCallback)?b.jsonpCallback():b.jsonpCallback,h?b[h]=b[h].replace(Rb,"$1"+e):b.jsonp!==!1&&(b.url+=(sb.test(b.url)?"&":"?")+b.jsonp+"="+e),b.converters["script json"]=function(){return g||r.error(e+" was not called"),g[0]},b.dataTypes[0]="json",f=a[e],a[e]=function(){g=arguments},d.always(function(){void 0===f?r(a).removeProp(e):a[e]=f,b[e]&&(b.jsonpCallback=c.jsonpCallback,Qb.push(e)),g&&r.isFunction(f)&&f(g[0]),g=f=void 0}),"script"}),o.createHTMLDocument=function(){var a=d.implementation.createHTMLDocument("").body;return a.innerHTML="<form></form><form></form>",2===a.childNodes.length}(),r.parseHTML=function(a,b,c){if("string"!=typeof a)return[];"boolean"==typeof b&&(c=b,b=!1);var e,f,g;return b||(o.createHTMLDocument?(b=d.implementation.createHTMLDocument(""),e=b.createElement("base"),e.href=d.location.href,b.head.appendChild(e)):b=d),f=B.exec(a),g=!c&&[],f?[b.createElement(f[1])]:(f=oa([a],b,g),g&&g.length&&r(g).remove(),r.merge([],f.childNodes))},r.fn.load=function(a,b,c){var d,e,f,g=this,h=a.indexOf(" ");return h>-1&&(d=r.trim(a.slice(h)),a=a.slice(0,h)),r.isFunction(b)?(c=b,b=void 0):b&&"object"==typeof b&&(e="POST"),g.length>0&&r.ajax({url:a,type:e||"GET",dataType:"html",data:b}).done(function(a){f=arguments,g.html(d?r("<div>").append(r.parseHTML(a)).find(d):a)}).always(c&&function(a,b){g.each(function(){c.apply(this,f||[a.responseText,b,a])})}),this},r.each(["ajaxStart","ajaxStop","ajaxComplete","ajaxError","ajaxSuccess","ajaxSend"],function(a,b){r.fn[b]=function(a){return this.on(b,a)}}),r.expr.pseudos.animated=function(a){return r.grep(r.timers,function(b){return a===b.elem}).length};function Sb(a){return r.isWindow(a)?a:9===a.nodeType&&a.defaultView}r.offset={setOffset:function(a,b,c){var d,e,f,g,h,i,j,k=r.css(a,"position"),l=r(a),m={};"static"===k&&(a.style.position="relative"),h=l.offset(),f=r.css(a,"top"),i=r.css(a,"left"),j=("absolute"===k||"fixed"===k)&&(f+i).indexOf("auto")>-1,j?(d=l.position(),g=d.top,e=d.left):(g=parseFloat(f)||0,e=parseFloat(i)||0),r.isFunction(b)&&(b=b.call(a,c,r.extend({},h))),null!=b.top&&(m.top=b.top-h.top+g),null!=b.left&&(m.left=b.left-h.left+e),"using"in b?b.using.call(a,m):l.css(m)}},r.fn.extend({offset:function(a){if(arguments.length)return void 0===a?this:this.each(function(b){r.offset.setOffset(this,a,b)});var b,c,d,e,f=this[0];if(f)return f.getClientRects().length?(d=f.getBoundingClientRect(),d.width||d.height?(e=f.ownerDocument,c=Sb(e),b=e.documentElement,{top:d.top+c.pageYOffset-b.clientTop,left:d.left+c.pageXOffset-b.clientLeft}):d):{top:0,left:0}},position:function(){if(this[0]){var a,b,c=this[0],d={top:0,left:0};return"fixed"===r.css(c,"position")?b=c.getBoundingClientRect():(a=this.offsetParent(),b=this.offset(),r.nodeName(a[0],"html")||(d=a.offset()),d={top:d.top+r.css(a[0],"borderTopWidth",!0),left:d.left+r.css(a[0],"borderLeftWidth",!0)}),{top:b.top-d.top-r.css(c,"marginTop",!0),left:b.left-d.left-r.css(c,"marginLeft",!0)}}},offsetParent:function(){return this.map(function(){var a=this.offsetParent;while(a&&"static"===r.css(a,"position"))a=a.offsetParent;return a||pa})}}),r.each({scrollLeft:"pageXOffset",scrollTop:"pageYOffset"},function(a,b){var c="pageYOffset"===b;r.fn[a]=function(d){return S(this,function(a,d,e){var f=Sb(a);return void 0===e?f?f[b]:a[d]:void(f?f.scrollTo(c?f.pageXOffset:e,c?e:f.pageYOffset):a[d]=e)},a,d,arguments.length)}}),r.each(["top","left"],function(a,b){r.cssHooks[b]=Na(o.pixelPosition,function(a,c){if(c)return c=Ma(a,b),Ka.test(c)?r(a).position()[b]+"px":c})}),r.each({Height:"height",Width:"width"},function(a,b){r.each({padding:"inner"+a,content:b,"":"outer"+a},function(c,d){r.fn[d]=function(e,f){var g=arguments.length&&(c||"boolean"!=typeof e),h=c||(e===!0||f===!0?"margin":"border");return S(this,function(b,c,e){var f;return r.isWindow(b)?0===d.indexOf("outer")?b["inner"+a]:b.document.documentElement["client"+a]:9===b.nodeType?(f=b.documentElement,Math.max(b.body["scroll"+a],f["scroll"+a],b.body["offset"+a],f["offset"+a],f["client"+a])):void 0===e?r.css(b,c,h):r.style(b,c,e,h)},b,g?e:void 0,g)}})}),r.fn.extend({bind:function(a,b,c){return this.on(a,null,b,c)},unbind:function(a,b){return this.off(a,null,b)},delegate:function(a,b,c,d){return this.on(b,a,c,d)},undelegate:function(a,b,c){return 1===arguments.length?this.off(a,"**"):this.off(b,a||"**",c)}}),r.parseJSON=JSON.parse,"function"==typeof define&&define.amd&&define("jquery",[],function(){return r});var Tb=a.jQuery,Ub=a.$;return r.noConflict=function(b){return a.$===r&&(a.$=Ub),b&&a.jQuery===r&&(a.jQuery=Tb),r},b||(a.jQuery=a.$=r),r});
```
