## Supplementary material for "ΔSCOPE: A new method to quantify 3D biological structures and identify differences in zebrafish forebrain development": An archived copy of the DeltaSCOPE code repository.: modernizr.min.js

```
/* Modernizr 2.6.2 (Custom Build) | MIT & BSD
 * Build: http://modernizr.com/download/#-fontface-backgroundsize-borderimage-borderradius-boxshadow-flexbox-hsla-multiplebgs-opacity-rgba-textshadow-cssanimations-csscolumns-generatedcontent-cssgradients-cssreflections-csstransforms-csstransforms3d-csstransitions-applicationcache-canvas-canvastext-draganddrop-hashchange-history-audio-video-indexeddb-input-inputtypes-localstorage-postmessage-sessionstorage-websockets-websqldatabase-webworkers-geolocation-inlinesvg-smil-svg-svgclippaths-touch-webgl-shiv-mq-cssclasses-addtest-prefixed-teststyles-testprop-testallprops-hasevent-prefixes-domprefixes-load
 */
;window.Modernizr=function(a,b,c){function D(a){j.cssText=a}function E(a,b){return D(n.join(a+";")+(b||""))}function F(a,b){return typeof a===b}function G(a,b){return!!~(""+a).indexOf(b)}function H(a,b){for(var d in a){var e=a[d];if(!G(e,"-")&&j[e]!==c)return b=="pfx"?e:!0}return!1}function I(a,b,d){for(var e in a){var f=b[a[e]];if(f!==c)return d===!1?a[e]:F(f,"function")?f.bind(d||b):f}return!1}function J(a,b,c){var d=a.charAt(0).toUpperCase()+a.slice(1),e=(a+" "+p.join(d+" ")+d).split(" ");return F(b,"string")||F(b,"undefined")?H(e,b):(e=(a+" "+q.join(d+" ")+d).split(" "),I(e,b,c))}function K(){e.input=function(c){for(var d=0,e=c.length;d<e;d++)u[c[d]]=c[d]in k;return u.list&&(u.list=!!b.createElement("datalist")&&!!a.HTMLDataListElement),u}("autocomplete autofocus list placeholder max min multiple pattern required step".split(" ")),e.inputtypes=function(a){for(var d=0,e,f,h,i=a.length;d<i;d++)k.setAttribute("type",f=a[d]),e=k.type!=="text",e&&(k.value=l,k.style.cssText="position:absolute;visibility:hidden;",/^range$/.test(f)&&k.style.WebkitAppearance!==c?(g.appendChild(k),h=b.defaultView,e=h.getComputedStyle&&h.getComputedStyle(k,null).WebkitAppearance!=="textfield"&&k.offsetHeight!==0,g.removeChild(k)):/^(search|tel)$/.test(f)||(/^(url|email)$/.test(f)?e=k.checkValidity&&k.checkValidity()===!1:e=k.value!=l)),t[a[d]]=!!e;return t}("search tel url email datetime date month week time datetime-local number range color".split(" "))}var d="2.6.2",e={},f=!0,g=b.documentElement,h="modernizr",i=b.createElement(h),j=i.style,k=b.createElement("input"),l=":)",m={}.toString,n=" -webkit- -moz- -o- -ms- ".split(" "),o="Webkit Moz O ms",p=o.split(" "),q=o.toLowerCase().split(" "),r={svg:"http://www.w3.org/2000/svg"},s={},t={},u={},v=[],w=v.slice,x,y=function(a,c,d,e){var f,i,j,k,l=b.createElement("div"),m=b.body,n=m||b.createElement("body");if(parseInt(d,10))while(d--)j=b.createElement("div"),j.id=e?e[d]:h+(d+1),l.appendChild(j);return f=["&#173;",'<style id="s',h,'">',a,"</style>"].join(""),l.id=h,(m?l:n).innerHTML+=f,n.appendChild(l),m||(n.style.background="",n.style.overflow="hidden",k=g.style.overflow,g.style.overflow="hidden",g.appendChild(n)),i=c(l,a),m?l.parentNode.removeChild(l):(n.parentNode.removeChild(n),g.style.overflow=k),!!i},z=function(b){var c=a.matchMedia||a.msMatchMedia;if(c)return c(b).matches;var d;return y("@media "+b+" { #"+h+" { position: absolute; } }",function(b){d=(a.getComputedStyle?getComputedStyle(b,null):b.currentStyle)["position"]=="absolute"}),d},A=function(){function d(d,e){e=e||b.createElement(a[d]||"div"),d="on"+d;var f=d in e;return f||(e.setAttribute||(e=b.createElement("div")),e.setAttribute&&e.removeAttribute&&(e.setAttribute(d,""),f=F(e[d],"function"),F(e[d],"undefined")||(e[d]=c),e.removeAttribute(d))),e=null,f}var a={select:"input",change:"input",submit:"form",reset:"form",error:"img",load:"img",abort:"img"};return d}(),B={}.hasOwnProperty,C;!F(B,"undefined")&&!F(B.call,"undefined")?C=function(a,b){return B.call(a,b)}:C=function(a,b){return b in a&&F(a.constructor.prototype[b],"undefined")},Function.prototype.bind||(Function.prototype.bind=function(b){var c=this;if(typeof c!="function")throw new TypeError;var d=w.call(arguments,1),e=function(){if(this instanceof e){var a=function(){};a.prototype=c.prototype;var f=new a,g=c.apply(f,d.concat(w.call(arguments)));return Object(g)===g?g:f}return c.apply(b,d.concat(w.call(arguments)))};return e}),s.flexbox=function(){return J("flexWrap")},s.canvas=function(){var a=b.createElement("canvas");return!!a.getContext&&!!a.getContext("2d")},s.canvastext=function(){return!!e.canvas&&!!F(b.createElement("canvas").getContext("2d").fillText,"function")},s.webgl=function(){return!!a.WebGLRenderingContext},s.touch=function(){var c;return"ontouchstart"in a||a.DocumentTouch&&b instanceof DocumentTouch?c=!0:y(["@media (",n.join("touch-enabled),("),h,")","{#modernizr{top:9px;position:absolute}}"].join(""),function(a){c=a.offsetTop===9}),c},s.geolocation=function(){return"geolocation"in navigator},s.postmessage=function(){return!!a.postMessage},s.websqldatabase=function(){return!!a.openDatabase},s.indexedDB=function(){return!!J("indexedDB",a)},s.hashchange=function(){return A("hashchange",a)&&(b.documentMode===c||b.documentMode>7)},s.history=function(){return!!a.history&&!!history.pushState},s.draganddrop=function(){var a=b.createElement("div");return"draggable"in a||"ondragstart"in a&&"ondrop"in a},s.websockets=function(){return"WebSocket"in a||"MozWebSocket"in a},s.rgba=function(){return D("background-color:rgba(150,255,150,.5)"),G(j.backgroundColor,"rgba")},s.hsla=function(){return D("background-color:hsla(120,40%,100%,.5)"),G(j.backgroundColor,"rgba")||G(j.backgroundColor,"hsla")},s.multiplebgs=function(){return D("background:url(https://),url(https://),red url(https://)"),/(url\s*\(.*?){3}/.test(j.background)},s.backgroundsize=function(){return J("backgroundSize")},s.borderimage=function(){return J("borderImage")},s.borderradius=function(){return J("borderRadius")},s.boxshadow=function(){return J("boxShadow")},s.textshadow=function(){return b.createElement("div").style.textShadow===""},s.opacity=function(){return E("opacity:.55"),/^0.55$/.test(j.opacity)},s.cssanimations=function(){return J("animationName")},s.csscolumns=function(){return J("columnCount")},s.cssgradients=function(){var a="background-image:",b="gradient(linear,left top,right bottom,from(#9f9),to(white));",c="linear-gradient(left top,#9f9, white);";return D((a+"-webkit- ".split(" ").join(b+a)+n.join(c+a)).slice(0,-a.length)),G(j.backgroundImage,"gradient")},s.cssreflections=function(){return J("boxReflect")},s.csstransforms=function(){return!!J("transform")},s.csstransforms3d=function(){var a=!!J("perspective");return a&&"webkitPerspective"in g.style&&y("@media (transform-3d),(-webkit-transform-3d){#modernizr{left:9px;position:absolute;height:3px;}}",function(b,c){a=b.offsetLeft===9&&b.offsetHeight===3}),a},s.csstransitions=function(){return J("transition")},s.fontface=function(){var a;return y('@font-face {font-family:"font";src:url("https://")}',function(c,d){var e=b.getElementById("smodernizr"),f=e.sheet||e.styleSheet,g=f?f.cssRules&&f.cssRules[0]?f.cssRules[0].cssText:f.cssText||"":"";a=/src/i.test(g)&&g.indexOf(d.split(" ")[0])===0}),a},s.generatedcontent=function(){var a;return y(["#",h,"{font:0/0 a}#",h,':after{content:"',l,'";visibility:hidden;font:3px/1 a}'].join(""),function(b){a=b.offsetHeight>=3}),a},s.video=function(){var a=b.createElement("video"),c=!1;try{if(c=!!a.canPlayType)c=new Boolean(c),c.ogg=a.canPlayType('video/ogg; codecs="theora"').replace(/^no$/,""),c.h264=a.canPlayType('video/mp4; codecs="avc1.42E01E"').replace(/^no$/,""),c.webm=a.canPlayType('video/webm; codecs="vp8, vorbis"').replace(/^no$/,"")}catch(d){}return c},s.audio=function(){var a=b.createElement("audio"),c=!1;try{if(c=!!a.canPlayType)c=new Boolean(c),c.ogg=a.canPlayType('audio/ogg; codecs="vorbis"').replace(/^no$/,""),c.mp3=a.canPlayType("audio/mpeg;").replace(/^no$/,""),c.wav=a.canPlayType('audio/wav; codecs="1"').replace(/^no$/,""),c.m4a=(a.canPlayType("audio/x-m4a;")||a.canPlayType("audio/aac;")).replace(/^no$/,"")}catch(d){}return c},s.localstorage=function(){try{return localStorage.setItem(h,h),localStorage.removeItem(h),!0}catch(a){return!1}},s.sessionstorage=function(){try{return sessionStorage.setItem(h,h),sessionStorage.removeItem(h),!0}catch(a){return!1}},s.webworkers=function(){return!!a.Worker},s.applicationcache=function(){return!!a.applicationCache},s.svg=function(){return!!b.createElementNS&&!!b.createElementNS(r.svg,"svg").createSVGRect},s.inlinesvg=function(){var a=b.createElement("div");return a.innerHTML="<svg/>",(a.firstChild&&a.firstChild.namespaceURI)==r.svg},s.smil=function(){return!!b.createElementNS&&/SVGAnimate/.test(m.call(b.createElementNS(r.svg,"animate")))},s.svgclippaths=function(){return!!b.createElementNS&&/SVGClipPath/.test(m.call(b.createElementNS(r.svg,"clipPath")))};for(var L in s)C(s,L)&&(x=L.toLowerCase(),e[x]=s[L](),v.push((e[x]?"":"no-")+x));return e.input||K(),e.addTest=function(a,b){if(typeof a=="object")for(var d in a)C(a,d)&&e.addTest(d,a[d]);else{a=a.toLowerCase();if(e[a]!==c)return e;b=typeof b=="function"?b():b,typeof f!="undefined"&&f&&(g.className+=" "+(b?"":"no-")+a),e[a]=b}return e},D(""),i=k=null,function(a,b){function k(a,b){var c=a.createElement("p"),d=a.getElementsByTagName("head")[0]||a.documentElement;return c.innerHTML="x<style>"+b+"</style>",d.insertBefore(c.lastChild,d.firstChild)}function l(){var a=r.elements;return typeof a=="string"?a.split(" "):a}function m(a){var b=i[a[g]];return b||(b={},h++,a[g]=h,i[h]=b),b}function n(a,c,f){c||(c=b);if(j)return c.createElement(a);f||(f=m(c));var g;return f.cache[a]?g=f.cache[a].cloneNode():e.test(a)?g=(f.cache[a]=f.createElem(a)).cloneNode():g=f.createElem(a),g.canHaveChildren&&!d.test(a)?f.frag.appendChild(g):g}function o(a,c){a||(a=b);if(j)return a.createDocumentFragment();c=c||m(a);var d=c.frag.cloneNode(),e=0,f=l(),g=f.length;for(;e<g;e++)d.createElement(f[e]);return d}function p(a,b){b.cache||(b.cache={},b.createElem=a.createElement,b.createFrag=a.createDocumentFragment,b.frag=b.createFrag()),a.createElement=function(c){return r.shivMethods?n(c,a,b):b.createElem(c)},a.createDocumentFragment=Function("h,f","return function(){var n=f.cloneNode(),c=n.createElement;h.shivMethods&&("+l().join().replace(/\w+/g,function(a){return b.createElem(a),b.frag.createElement(a),'c("'+a+'")'})+");return n}")(r,b.frag)}function q(a){a||(a=b);var c=m(a);return r.shivCSS&&!f&&!c.hasCSS&&(c.hasCSS=!!k(a,"article,aside,figcaption,figure,footer,header,hgroup,nav,section{display:block}mark{background:#FF0;color:#000}")),j||p(a,c),a}var c=a.html5||{},d=/^<|^(?:button|map|select|textarea|object|iframe|option|optgroup)$/i,e=/^(?:a|b|code|div|fieldset|h1|h2|h3|h4|h5|h6|i|label|li|ol|p|q|span|strong|style|table|tbody|td|th|tr|ul)$/i,f,g="_html5shiv",h=0,i={},j;(function(){try{var a=b.createElement("a");a.innerHTML="<xyz></xyz>",f="hidden"in a,j=a.childNodes.length==1||function(){b.createElement("a");var a=b.createDocumentFragment();return typeof a.cloneNode=="undefined"||typeof a.createDocumentFragment=="undefined"||typeof a.createElement=="undefined"}()}catch(c){f=!0,j=!0}})();var r={elements:c.elements||"abbr article aside audio bdi canvas data datalist details figcaption figure footer header hgroup mark meter nav output progress section summary time video",shivCSS:c.shivCSS!==!1,supportsUnknownElements:j,shivMethods:c.shivMethods!==!1,type:"default",shivDocument:q,createElement:n,createDocumentFragment:o};a.html5=r,q(b)}(this,b),e._version=d,e._prefixes=n,e._domPrefixes=q,e._cssomPrefixes=p,e.mq=z,e.hasEvent=A,e.testProp=function(a){return H([a])},e.testAllProps=J,e.testStyles=y,e.prefixed=function(a,b,c){return b?J(a,b,c):J(a,"pfx")},g.className=g.className.replace(/(^|\s)no-js(\s|$)/,"$1$2")+(f?" js "+v.join(" "):""),e}(this,this.document),function(a,b,c){function d(a){return"[object Function]"==o.call(a)}function e(a){return"string"==typeof a}function f(){}function g(a){return!a||"loaded"==a||"complete"==a||"uninitialized"==a}function h(){var a=p.shift();q=1,a?a.t?m(function(){("c"==a.t?B.injectCss:B.injectJs)(a.s,0,a.a,a.x,a.e,1)},0):(a(),h()):q=0}function i(a,c,d,e,f,i,j){function k(b){if(!o&&g(l.readyState)&&(u.r=o=1,!q&&h(),l.onload=l.onreadystatechange=null,b)){"img"!=a&&m(function(){t.removeChild(l)},50);for(var d in y[c])y[c].hasOwnProperty(d)&&y[c][d].onload()}}var j=j||B.errorTimeout,l=b.createElement(a),o=0,r=0,u={t:d,s:c,e:f,a:i,x:j};1===y[c]&&(r=1,y[c]=[]),"object"==a?l.data=c:(l.src=c,l.type=a),l.width=l.height="0",l.onerror=l.onload=l.onreadystatechange=function(){k.call(this,r)},p.splice(e,0,u),"img"!=a&&(r||2===y[c]?(t.insertBefore(l,s?null:n),m(k,j)):y[c].push(l))}function j(a,b,c,d,f){return q=0,b=b||"j",e(a)?i("c"==b?v:u,a,b,this.i++,c,d,f):(p.splice(this.i++,0,a),1==p.length&&h()),this}function k(){var a=B;return a.loader={load:j,i:0},a}var l=b.documentElement,m=a.setTimeout,n=b.getElementsByTagName("script")[0],o={}.toString,p=[],q=0,r="MozAppearance"in l.style,s=r&&!!b.createRange().compareNode,t=s?l:n.parentNode,l=a.opera&&"[object Opera]"==o.call(a.opera),l=!!b.attachEvent&&!l,u=r?"object":l?"script":"img",v=l?"script":u,w=Array.isArray||function(a){return"[object Array]"==o.call(a)},x=[],y={},z={timeout:function(a,b){return b.length&&(a.timeout=b[0]),a}},A,B;B=function(a){function b(a){var a=a.split("!"),b=x.length,c=a.pop(),d=a.length,c={url:c,origUrl:c,prefixes:a},e,f,g;for(f=0;f<d;f++)g=a[f].split("="),(e=z[g.shift()])&&(c=e(c,g));for(f=0;f<b;f++)c=x[f](c);return c}function g(a,e,f,g,h){var i=b(a),j=i.autoCallback;i.url.split(".").pop().split("?").shift(),i.bypass||(e&&(e=d(e)?e:e[a]||e[g]||e[a.split("/").pop().split("?")[0]]),i.instead?i.instead(a,e,f,g,h):(y[i.url]?i.noexec=!0:y[i.url]=1,f.load(i.url,i.forceCSS||!i.forceJS&&"css"==i.url.split(".").pop().split("?").shift()?"c":c,i.noexec,i.attrs,i.timeout),(d(e)||d(j))&&f.load(function(){k(),e&&e(i.origUrl,h,g),j&&j(i.origUrl,h,g),y[i.url]=2})))}function h(a,b){function c(a,c){if(a){if(e(a))c||(j=function(){var a=[].slice.call(arguments);k.apply(this,a),l()}),g(a,j,b,0,h);else if(Object(a)===a)for(n in m=function(){var b=0,c;for(c in a)a.hasOwnProperty(c)&&b++;return b}(),a)a.hasOwnProperty(n)&&(!c&&!--m&&(d(j)?j=function(){var a=[].slice.call(arguments);k.apply(this,a),l()}:j[n]=function(a){return function(){var b=[].slice.call(arguments);a&&a.apply(this,b),l()}}(k[n])),g(a[n],j,b,n,h))}else!c&&l()}var h=!!a.test,i=a.load||a.both,j=a.callback||f,k=j,l=a.complete||f,m,n;c(h?a.yep:a.nope,!!i),i&&c(i)}var i,j,l=this.yepnope.loader;if(e(a))g(a,0,l,0);else if(w(a))for(i=0;i<a.length;i++)j=a[i],e(j)?g(j,0,l,0):w(j)?B(j):Object(j)===j&&h(j,l);else Object(a)===a&&h(a,l)},B.addPrefix=function(a,b){z[a]=b},B.addFilter=function(a){x.push(a)},B.errorTimeout=1e4,null==b.readyState&&b.addEventListener&&(b.readyState="loading",b.addEventListener("DOMContentLoaded",A=function(){b.removeEventListener("DOMContentLoaded",A,0),b.readyState="complete"},0)),a.yepnope=k(),a.yepnope.executeStack=h,a.yepnope.injectJs=function(a,c,d,e,i,j){var k=b.createElement("script"),l,o,e=e||B.errorTimeout;k.src=a;for(o in d)k.setAttribute(o,d[o]);c=j?h:c||f,k.onreadystatechange=k.onload=function(){!l&&g(k.readyState)&&(l=1,c(),k.onload=k.onreadystatechange=null)},m(function(){l||(l=1,c(1))},e),i?k.onload():n.parentNode.insertBefore(k,n)},a.yepnope.injectCss=function(a,c,d,e,g,i){var e=b.createElement("link"),j,c=i?h:c||f;e.href=a,e.rel="stylesheet",e.type="text/css";for(j in d)e.setAttribute(j,d[j]);g||(n.parentNode.insertBefore(e,n),m(c,0))}}(this,document),Modernizr.load=function(){yepnope.apply(window,[].slice.call(arguments,0))};
```
