## Supplementary material for "ΔSCOPE: A new method to quantify 3D biological structures and identify differences in zebrafish forebrain development": An archived copy of the DeltaSCOPE code repository.: theme.js

```
require=(function e(t,n,r){function s(o,u){if(!n[o]){if(!t[o]){var a=typeof require=="function"&&require;if(!u&&a)return a(o,!0);if(i)return i(o,!0);var f=new Error("Cannot find module '"+o+"'");throw f.code="MODULE_NOT_FOUND",f}var l=n[o]={exports:{}};t[o][0].call(l.exports,function(e){var n=t[o][1][e];return s(n?n:e)},l,l.exports,e,t,n,r)}return n[o].exports}var i=typeof require=="function"&&require;for(var o=0;o<r.length;o++)s(r[o]);return s})({"sphinx-rtd-theme":[function(require,module,exports){
var jQuery = (typeof(window) != 'undefined') ? window.jQuery : require('jquery');

// Sphinx theme nav state
function ThemeNav () {

    var nav = {
        navBar: null,
        win: null,
        winScroll: false,
        winResize: false,
        linkScroll: false,
        winPosition: 0,
        winHeight: null,
        docHeight: null,
        isRunning: null
    };

    nav.enable = function () {
        var self = this;

        jQuery(function ($) {
            self.init($);

            self.reset();
            self.win.on('hashchange', self.reset);

            // Set scroll monitor
            self.win.on('scroll', function () {
                if (!self.linkScroll) {
                    self.winScroll = true;
                }
            });
            setInterval(function () { if (self.winScroll) self.onScroll(); }, 25);

            // Set resize monitor
            self.win.on('resize', function () {
                self.winResize = true;
            });
            setInterval(function () { if (self.winResize) self.onResize(); }, 25);
            self.onResize();
        });
    };

    nav.init = function ($) {
        var doc = $(document),
            self = this;

        this.navBar = $('div.wy-side-scroll:first');
        this.win = $(window);

        // Set up javascript UX bits
        $(document)
            // Shift nav in mobile when clicking the menu.
            .on('click', "[data-toggle='wy-nav-top']", function() {
                $("[data-toggle='wy-nav-shift']").toggleClass("shift");
                $("[data-toggle='rst-versions']").toggleClass("shift");
            })

            // Nav menu link click operations
            .on('click', ".wy-menu-vertical .current ul li a", function() {
                var target = $(this);
                // Close menu when you click a link.
                $("[data-toggle='wy-nav-shift']").removeClass("shift");
                $("[data-toggle='rst-versions']").toggleClass("shift");
                // Handle dynamic display of l3 and l4 nav lists
                self.toggleCurrent(target);
                self.hashChange();
            })
            .on('click', "[data-toggle='rst-current-version']", function() {
                $("[data-toggle='rst-versions']").toggleClass("shift-up");
            })

        // Make tables responsive
        $("table.docutils:not(.field-list)")
            .wrap("<div class='wy-table-responsive'></div>");

        // Add expand links to all parents of nested ul
        $('.wy-menu-vertical ul').not('.simple').siblings('a').each(function () {
            var link = $(this);
                expand = $('<span class="toctree-expand"></span>');
            expand.on('click', function (ev) {
                self.toggleCurrent(link);
                ev.stopPropagation();
                return false;
            });
            link.prepend(expand);
        });
    };

    nav.reset = function () {
        // Get anchor from URL and open up nested nav
        var anchor = encodeURI(window.location.hash);
        if (anchor) {
            try {
                var link = $('.wy-menu-vertical')
                    .find('[href="' + anchor + '"]');
                $('.wy-menu-vertical li.toctree-l1 li.current')
                    .removeClass('current');
                link.closest('li.toctree-l2').addClass('current');
                link.closest('li.toctree-l3').addClass('current');
                link.closest('li.toctree-l4').addClass('current');
            }
            catch (err) {
                console.log("Error expanding nav for anchor", err);
            }
        }
    };

    nav.onScroll = function () {
        this.winScroll = false;
        var newWinPosition = this.win.scrollTop(),
            winBottom = newWinPosition + this.winHeight,
            navPosition = this.navBar.scrollTop(),
            newNavPosition = navPosition + (newWinPosition - this.winPosition);
        if (newWinPosition < 0 || winBottom > this.docHeight) {
            return;
        }
        this.navBar.scrollTop(newNavPosition);
        this.winPosition = newWinPosition;
    };

    nav.onResize = function () {
        this.winResize = false;
        this.winHeight = this.win.height();
        this.docHeight = $(document).height();
    };

    nav.hashChange = function () {
        this.linkScroll = true;
        this.win.one('hashchange', function () {
            this.linkScroll = false;
        });
    };

    nav.toggleCurrent = function (elem) {
        var parent_li = elem.closest('li');
        parent_li.siblings('li.current').removeClass('current');
        parent_li.siblings().find('li.current').removeClass('current');
        parent_li.find('> ul li.current').removeClass('current');
        parent_li.toggleClass('current');
    }

    return nav;
};

module.exports.ThemeNav = ThemeNav();

if (typeof(window) != 'undefined') {
    window.SphinxRtdTheme = { StickyNav: module.exports.ThemeNav };
}

},{"jquery":"jquery"}]},{},["sphinx-rtd-theme"]);
```
