## Supplementary material for "ΔSCOPE: A new method to quantify 3D biological structures and identify differences in zebrafish forebrain development": An archived copy of the DeltaSCOPE code repository.: searchtools.js

```
/*
 * searchtools.js_t
 * ~~~~~~~~~~~~~~~~
 *
 * Sphinx JavaScript utilities for the full-text search.
 *
 * :copyright: Copyright 2007-2018 by the Sphinx team, see AUTHORS.
 * :license: BSD, see LICENSE for details.
 *
 */


/* Non-minified version JS is _stemmer.js if file is provided */ 
/**
 * Porter Stemmer
 */
var Stemmer = function() {

  var step2list = {
    ational: 'ate',
    tional: 'tion',
    enci: 'ence',
    anci: 'ance',
    izer: 'ize',
    bli: 'ble',
    alli: 'al',
    entli: 'ent',
    eli: 'e',
    ousli: 'ous',
    ization: 'ize',
    ation: 'ate',
    ator: 'ate',
    alism: 'al',
    iveness: 'ive',
    fulness: 'ful',
    ousness: 'ous',
    aliti: 'al',
    iviti: 'ive',
    biliti: 'ble',
    logi: 'log'
  };

  var step3list = {
    icate: 'ic',
    ative: '',
    alize: 'al',
    iciti: 'ic',
    ical: 'ic',
    ful: '',
    ness: ''
  };

  var c = "[^aeiou]";          // consonant
  var v = "[aeiouy]";          // vowel
  var C = c + "[^aeiouy]*";    // consonant sequence
  var V = v + "[aeiou]*";      // vowel sequence

  var mgr0 = "^(" + C + ")?" + V + C;                      // [C]VC... is m>0
  var meq1 = "^(" + C + ")?" + V + C + "(" + V + ")?$";    // [C]VC[V] is m=1
  var mgr1 = "^(" + C + ")?" + V + C + V + C;              // [C]VCVC... is m>1
  var s_v   = "^(" + C + ")?" + v;                         // vowel in stem

  this.stemWord = function (w) {
    var stem;
    var suffix;
    var firstch;
    var origword = w;

    if (w.length < 3)
      return w;

    var re;
    var re2;
    var re3;
    var re4;

    firstch = w.substr(0,1);
    if (firstch == "y")
      w = firstch.toUpperCase() + w.substr(1);

    // Step 1a
    re = /^(.+?)(ss|i)es$/;
    re2 = /^(.+?)([^s])s$/;

    if (re.test(w))
      w = w.replace(re,"$1$2");
    else if (re2.test(w))
      w = w.replace(re2,"$1$2");

    // Step 1b
    re = /^(.+?)eed$/;
    re2 = /^(.+?)(ed|ing)$/;
    if (re.test(w)) {
      var fp = re.exec(w);
      re = new RegExp(mgr0);
      if (re.test(fp[1])) {
        re = /.$/;
        w = w.replace(re,"");
      }
    }
    else if (re2.test(w)) {
      var fp = re2.exec(w);
      stem = fp[1];
      re2 = new RegExp(s_v);
      if (re2.test(stem)) {
        w = stem;
        re2 = /(at|bl|iz)$/;
        re3 = new RegExp("([^aeiouylsz])\\1$");
        re4 = new RegExp("^" + C + v + "[^aeiouwxy]$");
        if (re2.test(w))
          w = w + "e";
        else if (re3.test(w)) {
          re = /.$/;
          w = w.replace(re,"");
        }
        else if (re4.test(w))
          w = w + "e";
      }
    }

    // Step 1c
    re = /^(.+?)y$/;
    if (re.test(w)) {
      var fp = re.exec(w);
      stem = fp[1];
      re = new RegExp(s_v);
      if (re.test(stem))
        w = stem + "i";
    }

    // Step 2
    re = /^(.+?)(ational|tional|enci|anci|izer|bli|alli|entli|eli|ousli|ization|ation|ator|alism|iveness|fulness|ousness|aliti|iviti|biliti|logi)$/;
    if (re.test(w)) {
      var fp = re.exec(w);
      stem = fp[1];
      suffix = fp[2];
      re = new RegExp(mgr0);
      if (re.test(stem))
        w = stem + step2list[suffix];
    }

    // Step 3
    re = /^(.+?)(icate|ative|alize|iciti|ical|ful|ness)$/;
    if (re.test(w)) {
      var fp = re.exec(w);
      stem = fp[1];
      suffix = fp[2];
      re = new RegExp(mgr0);
      if (re.test(stem))
        w = stem + step3list[suffix];
    }

    // Step 4
    re = /^(.+?)(al|ance|ence|er|ic|able|ible|ant|ement|ment|ent|ou|ism|ate|iti|ous|ive|ize)$/;
    re2 = /^(.+?)(s|t)(ion)$/;
    if (re.test(w)) {
      var fp = re.exec(w);
      stem = fp[1];
      re = new RegExp(mgr1);
      if (re.test(stem))
        w = stem;
    }
    else if (re2.test(w)) {
      var fp = re2.exec(w);
      stem = fp[1] + fp[2];
      re2 = new RegExp(mgr1);
      if (re2.test(stem))
        w = stem;
    }

    // Step 5
    re = /^(.+?)e$/;
    if (re.test(w)) {
      var fp = re.exec(w);
      stem = fp[1];
      re = new RegExp(mgr1);
      re2 = new RegExp(meq1);
      re3 = new RegExp("^" + C + v + "[^aeiouwxy]$");
      if (re.test(stem) || (re2.test(stem) && !(re3.test(stem))))
        w = stem;
    }
    re = /ll$/;
    re2 = new RegExp(mgr1);
    if (re.test(w) && re2.test(w)) {
      re = /.$/;
      w = w.replace(re,"");
    }

    // and turn initial Y back to y
    if (firstch == "y")
      w = firstch.toLowerCase() + w.substr(1);
    return w;
  }
}


/**
 * Simple result scoring code.
 */
var Scorer = {
  // Implement the following function to further tweak the score for each result
  // The function takes a result array [filename, title, anchor, descr, score]
  // and returns the new score.
  /*
  score: function(result) {
    return result[4];
  },
  */

  // query matches the full name of an object
  objNameMatch: 11,
  // or matches in the last dotted part of the object name
  objPartialMatch: 6,
  // Additive scores depending on the priority of the object
  objPrio: {0:  15,   // used to be importantResults
            1:  5,   // used to be objectResults
            2: -5},  // used to be unimportantResults
  //  Used when the priority is not in the mapping.
  objPrioDefault: 0,

  // query found in title
  title: 15,
  // query found in terms
  term: 5
};


var splitChars = (function() {
    var result = {};
    var singles = [96, 180, 187, 191, 215, 247, 749, 885, 903, 907, 909, 930, 1014, 1648,
         1748, 1809, 2416, 2473, 2481, 2526, 2601, 2609, 2612, 2615, 2653, 2702,
         2706, 2729, 2737, 2740, 2857, 2865, 2868, 2910, 2928, 2948, 2961, 2971,
         2973, 3085, 3089, 3113, 3124, 3213, 3217, 3241, 3252, 3295, 3341, 3345,
         3369, 3506, 3516, 3633, 3715, 3721, 3736, 3744, 3748, 3750, 3756, 3761,
         3781, 3912, 4239, 4347, 4681, 4695, 4697, 4745, 4785, 4799, 4801, 4823,
         4881, 5760, 5901, 5997, 6313, 7405, 8024, 8026, 8028, 8030, 8117, 8125,
         8133, 8181, 8468, 8485, 8487, 8489, 8494, 8527, 11311, 11359, 11687, 11695,
         11703, 11711, 11719, 11727, 11735, 12448, 12539, 43010, 43014, 43019, 43587,
         43696, 43713, 64286, 64297, 64311, 64317, 64319, 64322, 64325, 65141];
    var i, j, start, end;
    for (i = 0; i < singles.length; i++) {
        result[singles[i]] = true;
    }
    var ranges = [[0, 47], [58, 64], [91, 94], [123, 169], [171, 177], [182, 184], [706, 709],
         [722, 735], [741, 747], [751, 879], [888, 889], [894, 901], [1154, 1161],
         [1318, 1328], [1367, 1368], [1370, 1376], [1416, 1487], [1515, 1519], [1523, 1568],
         [1611, 1631], [1642, 1645], [1750, 1764], [1767, 1773], [1789, 1790], [1792, 1807],
         [1840, 1868], [1958, 1968], [1970, 1983], [2027, 2035], [2038, 2041], [2043, 2047],
         [2070, 2073], [2075, 2083], [2085, 2087], [2089, 2307], [2362, 2364], [2366, 2383],
         [2385, 2391], [2402, 2405], [2419, 2424], [2432, 2436], [2445, 2446], [2449, 2450],
         [2483, 2485], [2490, 2492], [2494, 2509], [2511, 2523], [2530, 2533], [2546, 2547],
         [2554, 2564], [2571, 2574], [2577, 2578], [2618, 2648], [2655, 2661], [2672, 2673],
         [2677, 2692], [2746, 2748], [2750, 2767], [2769, 2783], [2786, 2789], [2800, 2820],
         [2829, 2830], [2833, 2834], [2874, 2876], [2878, 2907], [2914, 2917], [2930, 2946],
         [2955, 2957], [2966, 2968], [2976, 2978], [2981, 2983], [2987, 2989], [3002, 3023],
         [3025, 3045], [3059, 3076], [3130, 3132], [3134, 3159], [3162, 3167], [3170, 3173],
         [3184, 3191], [3199, 3204], [3258, 3260], [3262, 3293], [3298, 3301], [3312, 3332],
         [3386, 3388], [3390, 3423], [3426, 3429], [3446, 3449], [3456, 3460], [3479, 3481],
         [3518, 3519], [3527, 3584], [3636, 3647], [3655, 3663], [3674, 3712], [3717, 3718],
         [3723, 3724], [3726, 3731], [3752, 3753], [3764, 3772], [3774, 3775], [3783, 3791],
         [3802, 3803], [3806, 3839], [3841, 3871], [3892, 3903], [3949, 3975], [3980, 4095],
         [4139, 4158], [4170, 4175], [4182, 4185], [4190, 4192], [4194, 4196], [4199, 4205],
         [4209, 4212], [4226, 4237], [4250, 4255], [4294, 4303], [4349, 4351], [4686, 4687],
         [4702, 4703], [4750, 4751], [4790, 4791], [4806, 4807], [4886, 4887], [4955, 4968],
         [4989, 4991], [5008, 5023], [5109, 5120], [5741, 5742], [5787, 5791], [5867, 5869],
         [5873, 5887], [5906, 5919], [5938, 5951], [5970, 5983], [6001, 6015], [6068, 6102],
         [6104, 6107], [6109, 6111], [6122, 6127], [6138, 6159], [6170, 6175], [6264, 6271],
         [6315, 6319], [6390, 6399], [6429, 6469], [6510, 6511], [6517, 6527], [6572, 6592],
         [6600, 6607], [6619, 6655], [6679, 6687], [6741, 6783], [6794, 6799], [6810, 6822],
         [6824, 6916], [6964, 6980], [6988, 6991], [7002, 7042], [7073, 7085], [7098, 7167],
         [7204, 7231], [7242, 7244], [7294, 7400], [7410, 7423], [7616, 7679], [7958, 7959],
         [7966, 7967], [8006, 8007], [8014, 8015], [8062, 8063], [8127, 8129], [8141, 8143],
         [8148, 8149], [8156, 8159], [8173, 8177], [8189, 8303], [8306, 8307], [8314, 8318],
         [8330, 8335], [8341, 8449], [8451, 8454], [8456, 8457], [8470, 8472], [8478, 8483],
         [8506, 8507], [8512, 8516], [8522, 8525], [8586, 9311], [9372, 9449], [9472, 10101],
         [10132, 11263], [11493, 11498], [11503, 11516], [11518, 11519], [11558, 11567],
         [11622, 11630], [11632, 11647], [11671, 11679], [11743, 11822], [11824, 12292],
         [12296, 12320], [12330, 12336], [12342, 12343], [12349, 12352], [12439, 12444],
         [12544, 12548], [12590, 12592], [12687, 12689], [12694, 12703], [12728, 12783],
         [12800, 12831], [12842, 12880], [12896, 12927], [12938, 12976], [12992, 13311],
         [19894, 19967], [40908, 40959], [42125, 42191], [42238, 42239], [42509, 42511],
         [42540, 42559], [42592, 42593], [42607, 42622], [42648, 42655], [42736, 42774],
         [42784, 42785], [42889, 42890], [42893, 43002], [43043, 43055], [43062, 43071],
         [43124, 43137], [43188, 43215], [43226, 43249], [43256, 43258], [43260, 43263],
         [43302, 43311], [43335, 43359], [43389, 43395], [43443, 43470], [43482, 43519],
         [43561, 43583], [43596, 43599], [43610, 43615], [43639, 43641], [43643, 43647],
         [43698, 43700], [43703, 43704], [43710, 43711], [43715, 43738], [43742, 43967],
         [44003, 44015], [44026, 44031], [55204, 55215], [55239, 55242], [55292, 55295],
         [57344, 63743], [64046, 64047], [64110, 64111], [64218, 64255], [64263, 64274],
         [64280, 64284], [64434, 64466], [64830, 64847], [64912, 64913], [64968, 65007],
         [65020, 65135], [65277, 65295], [65306, 65312], [65339, 65344], [65371, 65381],
         [65471, 65473], [65480, 65481], [65488, 65489], [65496, 65497]];
    for (i = 0; i < ranges.length; i++) {
        start = ranges[i][0];
        end = ranges[i][1];
        for (j = start; j <= end; j++) {
            result[j] = true;
        }
    }
    return result;
})();

function splitQuery(query) {
    var result = [];
    var start = -1;
    for (var i = 0; i < query.length; i++) {
        if (splitChars[query.charCodeAt(i)]) {
            if (start !== -1) {
                result.push(query.slice(start, i));
                start = -1;
            }
        } else if (start === -1) {
            start = i;
        }
    }
    if (start !== -1) {
        result.push(query.slice(start));
    }
    return result;
}


/**
 * Search Module
 */
var Search = {

  _index : null,
  _queued_query : null,
  _pulse_status : -1,

  init : function() {
      var params = $.getQueryParameters();
      if (params.q) {
          var query = params.q[0];
          $('input[name="q"]')[0].value = query;
          this.performSearch(query);
      }
  },

  loadIndex : function(url) {
    $.ajax({type: "GET", url: url, data: null,
            dataType: "script", cache: true,
            complete: function(jqxhr, textstatus) {
              if (textstatus != "success") {
                document.getElementById("searchindexloader").src = url;
              }
            }});
  },

  setIndex : function(index) {
    var q;
    this._index = index;
    if ((q = this._queued_query) !== null) {
      this._queued_query = null;
      Search.query(q);
    }
  },

  hasIndex : function() {
      return this._index !== null;
  },

  deferQuery : function(query) {
      this._queued_query = query;
  },

  stopPulse : function() {
      this._pulse_status = 0;
  },

  startPulse : function() {
    if (this._pulse_status >= 0)
        return;
    function pulse() {
      var i;
      Search._pulse_status = (Search._pulse_status + 1) % 4;
      var dotString = '';
      for (i = 0; i < Search._pulse_status; i++)
        dotString += '.';
      Search.dots.text(dotString);
      if (Search._pulse_status > -1)
        window.setTimeout(pulse, 500);
    }
    pulse();
  },

  /**
   * perform a search for something (or wait until index is loaded)
   */
  performSearch : function(query) {
    // create the required interface elements
    this.out = $('#search-results');
    this.title = $('<h2>' + _('Searching') + '</h2>').appendTo(this.out);
    this.dots = $('<span></span>').appendTo(this.title);
    this.status = $('<p style="display: none"></p>').appendTo(this.out);
    this.output = $('<ul class="search"/>').appendTo(this.out);

    $('#search-progress').text(_('Preparing search...'));
    this.startPulse();

    // index already loaded, the browser was quick!
    if (this.hasIndex())
      this.query(query);
    else
      this.deferQuery(query);
  },

  /**
   * execute search (requires search index to be loaded)
   */
  query : function(query) {
    var i;
    var stopwords = ["a","and","are","as","at","be","but","by","for","if","in","into","is","it","near","no","not","of","on","or","such","that","the","their","then","there","these","they","this","to","was","will","with"];

    // stem the searchterms and add them to the correct list
    var stemmer = new Stemmer();
    var searchterms = [];
    var excluded = [];
    var hlterms = [];
    var tmp = splitQuery(query);
    var objectterms = [];
    for (i = 0; i < tmp.length; i++) {
      if (tmp[i] !== "") {
          objectterms.push(tmp[i].toLowerCase());
      }

      if ($u.indexOf(stopwords, tmp[i].toLowerCase()) != -1 || tmp[i].match(/^\d+$/) ||
          tmp[i] === "") {
        // skip this "word"
        continue;
      }
      // stem the word
      var word = stemmer.stemWord(tmp[i].toLowerCase());
      // prevent stemmer from cutting word smaller than two chars
      if(word.length < 3 && tmp[i].length >= 3) {
        word = tmp[i];
      }
      var toAppend;
      // select the correct list
      if (word[0] == '-') {
        toAppend = excluded;
        word = word.substr(1);
      }
      else {
        toAppend = searchterms;
        hlterms.push(tmp[i].toLowerCase());
      }
      // only add if not already in the list
      if (!$u.contains(toAppend, word))
        toAppend.push(word);
    }
    var highlightstring = '?highlight=' + $.urlencode(hlterms.join(" "));

    // console.debug('SEARCH: searching for:');
    // console.info('required: ', searchterms);
    // console.info('excluded: ', excluded);

    // prepare search
    var terms = this._index.terms;
    var titleterms = this._index.titleterms;

    // array of [filename, title, anchor, descr, score]
    var results = [];
    $('#search-progress').empty();

    // lookup as object
    for (i = 0; i < objectterms.length; i++) {
      var others = [].concat(objectterms.slice(0, i),
                             objectterms.slice(i+1, objectterms.length));
      results = results.concat(this.performObjectSearch(objectterms[i], others));
    }

    // lookup as search terms in fulltext
    results = results.concat(this.performTermsSearch(searchterms, excluded, terms, titleterms));

    // let the scorer override scores with a custom scoring function
    if (Scorer.score) {
      for (i = 0; i < results.length; i++)
        results[i][4] = Scorer.score(results[i]);
    }

    // now sort the results by score (in opposite order of appearance, since the
    // display function below uses pop() to retrieve items) and then
    // alphabetically
    results.sort(function(a, b) {
      var left = a[4];
      var right = b[4];
      if (left > right) {
        return 1;
      } else if (left < right) {
        return -1;
      } else {
        // same score: sort alphabetically
        left = a[1].toLowerCase();
        right = b[1].toLowerCase();
        return (left > right) ? -1 : ((left < right) ? 1 : 0);
      }
    });

    // for debugging
    //Search.lastresults = results.slice();  // a copy
    //console.info('search results:', Search.lastresults);

    // print the results
    var resultCount = results.length;
    function displayNextItem() {
      // results left, load the summary and display it
      if (results.length) {
        var item = results.pop();
        var listItem = $('<li style="display:none"></li>');
        if (DOCUMENTATION_OPTIONS.FILE_SUFFIX === '') {
          // dirhtml builder
          var dirname = item[0] + '/';
          if (dirname.match(/\/index\/$/)) {
            dirname = dirname.substring(0, dirname.length-6);
          } else if (dirname == 'index/') {
            dirname = '';
          }
          listItem.append($('<a/>').attr('href',
            DOCUMENTATION_OPTIONS.URL_ROOT + dirname +
            highlightstring + item[2]).html(item[1]));
        } else {
          // normal html builders
          listItem.append($('<a/>').attr('href',
            item[0] + DOCUMENTATION_OPTIONS.FILE_SUFFIX +
            highlightstring + item[2]).html(item[1]));
        }
        if (item[3]) {
          listItem.append($('<span> (' + item[3] + ')</span>'));
          Search.output.append(listItem);
          listItem.slideDown(5, function() {
            displayNextItem();
          });
        } else if (DOCUMENTATION_OPTIONS.HAS_SOURCE) {
          var suffix = DOCUMENTATION_OPTIONS.SOURCELINK_SUFFIX;
          if (suffix === undefined) {
            suffix = '.txt';
          }
          $.ajax({url: DOCUMENTATION_OPTIONS.URL_ROOT + '_sources/' + item[5] + (item[5].slice(-suffix.length) === suffix ? '' : suffix),
                  dataType: "text",
                  complete: function(jqxhr, textstatus) {
                    var data = jqxhr.responseText;
                    if (data !== '' && data !== undefined) {
                      listItem.append(Search.makeSearchSummary(data, searchterms, hlterms));
                    }
                    Search.output.append(listItem);
                    listItem.slideDown(5, function() {
                      displayNextItem();
                    });
                  }});
        } else {
          // no source available, just display title
          Search.output.append(listItem);
          listItem.slideDown(5, function() {
            displayNextItem();
          });
        }
      }
      // search finished, update title and status message
      else {
        Search.stopPulse();
        Search.title.text(_('Search Results'));
        if (!resultCount)
          Search.status.text(_('Your search did not match any documents. Please make sure that all words are spelled correctly and that you\'ve selected enough categories.'));
        else
            Search.status.text(_('Search finished, found %s page(s) matching the search query.').replace('%s', resultCount));
        Search.status.fadeIn(500);
      }
    }
    displayNextItem();
  },

  /**
   * search for object names
   */
  performObjectSearch : function(object, otherterms) {
    var filenames = this._index.filenames;
    var docnames = this._index.docnames;
    var objects = this._index.objects;
    var objnames = this._index.objnames;
    var titles = this._index.titles;

    var i;
    var results = [];

    for (var prefix in objects) {
      for (var name in objects[prefix]) {
        var fullname = (prefix ? prefix + '.' : '') + name;
        if (fullname.toLowerCase().indexOf(object) > -1) {
          var score = 0;
          var parts = fullname.split('.');
          // check for different match types: exact matches of full name or
          // "last name" (i.e. last dotted part)
          if (fullname == object || parts[parts.length - 1] == object) {
            score += Scorer.objNameMatch;
          // matches in last name
          } else if (parts[parts.length - 1].indexOf(object) > -1) {
            score += Scorer.objPartialMatch;
          }
          var match = objects[prefix][name];
          var objname = objnames[match[1]][2];
          var title = titles[match[0]];
          // If more than one term searched for, we require other words to be
          // found in the name/title/description
          if (otherterms.length > 0) {
            var haystack = (prefix + ' ' + name + ' ' +
                            objname + ' ' + title).toLowerCase();
            var allfound = true;
            for (i = 0; i < otherterms.length; i++) {
              if (haystack.indexOf(otherterms[i]) == -1) {
                allfound = false;
                break;
              }
            }
            if (!allfound) {
              continue;
            }
          }
          var descr = objname + _(', in ') + title;

          var anchor = match[3];
          if (anchor === '')
            anchor = fullname;
          else if (anchor == '-')
            anchor = objnames[match[1]][1] + '-' + fullname;
          // add custom score for some objects according to scorer
          if (Scorer.objPrio.hasOwnProperty(match[2])) {
            score += Scorer.objPrio[match[2]];
          } else {
            score += Scorer.objPrioDefault;
          }
          results.push([docnames[match[0]], fullname, '#'+anchor, descr, score, filenames[match[0]]]);
        }
      }
    }

    return results;
  },

  /**
   * search for full-text terms in the index
   */
  performTermsSearch : function(searchterms, excluded, terms, titleterms) {
    var docnames = this._index.docnames;
    var filenames = this._index.filenames;
    var titles = this._index.titles;

    var i, j, file;
    var fileMap = {};
    var scoreMap = {};
    var results = [];

    // perform the search on the required terms
    for (i = 0; i < searchterms.length; i++) {
      var word = searchterms[i];
      var files = [];
      var _o = [
        {files: terms[word], score: Scorer.term},
        {files: titleterms[word], score: Scorer.title}
      ];

      // no match but word was a required one
      if ($u.every(_o, function(o){return o.files === undefined;})) {
        break;
      }
      // found search word in contents
      $u.each(_o, function(o) {
        var _files = o.files;
        if (_files === undefined)
          return

        if (_files.length === undefined)
          _files = [_files];
        files = files.concat(_files);

        // set score for the word in each file to Scorer.term
        for (j = 0; j < _files.length; j++) {
          file = _files[j];
          if (!(file in scoreMap))
            scoreMap[file] = {}
          scoreMap[file][word] = o.score;
        }
      });

      // create the mapping
      for (j = 0; j < files.length; j++) {
        file = files[j];
        if (file in fileMap)
          fileMap[file].push(word);
        else
          fileMap[file] = [word];
      }
    }

    // now check if the files don't contain excluded terms
    for (file in fileMap) {
      var valid = true;

      // check if all requirements are matched
      if (fileMap[file].length != searchterms.length)
          continue;

      // ensure that none of the excluded terms is in the search result
      for (i = 0; i < excluded.length; i++) {
        if (terms[excluded[i]] == file ||
            titleterms[excluded[i]] == file ||
            $u.contains(terms[excluded[i]] || [], file) ||
            $u.contains(titleterms[excluded[i]] || [], file)) {
          valid = false;
          break;
        }
      }

      // if we have still a valid result we can add it to the result list
      if (valid) {
        // select one (max) score for the file.
        // for better ranking, we should calculate ranking by using words statistics like basic tf-idf...
        var score = $u.max($u.map(fileMap[file], function(w){return scoreMap[file][w]}));
        results.push([docnames[file], titles[file], '', null, score, filenames[file]]);
      }
    }
    return results;
  },

  /**
   * helper function to return a node containing the
   * search summary for a given text. keywords is a list
   * of stemmed words, hlwords is the list of normal, unstemmed
   * words. the first one is used to find the occurrence, the
   * latter for highlighting it.
   */
  makeSearchSummary : function(text, keywords, hlwords) {
    var textLower = text.toLowerCase();
    var start = 0;
    $.each(keywords, function() {
      var i = textLower.indexOf(this.toLowerCase());
      if (i > -1)
        start = i;
    });
    start = Math.max(start - 120, 0);
    var excerpt = ((start > 0) ? '...' : '') +
      $.trim(text.substr(start, 240)) +
      ((start + 240 - text.length) ? '...' : '');
    var rv = $('<div class="context"></div>').text(excerpt);
    $.each(hlwords, function() {
      rv = rv.highlightText(this, 'highlighted');
    });
    return rv;
  }
};

$(document).ready(function() {
  Search.init();
});
```
