## Supplementary material for "ΔSCOPE: A new method to quantify 3D biological structures and identify differences in zebrafish forebrain development": An archived copy of the DeltaSCOPE code repository.: underscore-1.3.1.js

```
//     Underscore.js 1.3.1
//     (c) 2009-2012 Jeremy Ashkenas, DocumentCloud Inc.
//     Underscore is freely distributable under the MIT license.
//     Portions of Underscore are inspired or borrowed from Prototype,
//     Oliver Steele's Functional, and John Resig's Micro-Templating.
//     For all details and documentation:
//     http://documentcloud.github.com/underscore

(function() {

  // Baseline setup
  // --------------

  // Establish the root object, `window` in the browser, or `global` on the server.
  var root = this;

  // Save the previous value of the `_` variable.
  var previousUnderscore = root._;

  // Establish the object that gets returned to break out of a loop iteration.
  var breaker = {};

  // Save bytes in the minified (but not gzipped) version:
  var ArrayProto = Array.prototype, ObjProto = Object.prototype, FuncProto = Function.prototype;

  // Create quick reference variables for speed access to core prototypes.
  var slice            = ArrayProto.slice,
      unshift          = ArrayProto.unshift,
      toString         = ObjProto.toString,
      hasOwnProperty   = ObjProto.hasOwnProperty;

  // All **ECMAScript 5** native function implementations that we hope to use
  // are declared here.
  var
    nativeForEach      = ArrayProto.forEach,
    nativeMap          = ArrayProto.map,
    nativeReduce       = ArrayProto.reduce,
    nativeReduceRight  = ArrayProto.reduceRight,
    nativeFilter       = ArrayProto.filter,
    nativeEvery        = ArrayProto.every,
    nativeSome         = ArrayProto.some,
    nativeIndexOf      = ArrayProto.indexOf,
    nativeLastIndexOf  = ArrayProto.lastIndexOf,
    nativeIsArray      = Array.isArray,
    nativeKeys         = Object.keys,
    nativeBind         = FuncProto.bind;

  // Create a safe reference to the Underscore object for use below.
  var _ = function(obj) { return new wrapper(obj); };

  // Export the Underscore object for **Node.js**, with
  // backwards-compatibility for the old `require()` API. If we're in
  // the browser, add `_` as a global object via a string identifier,
  // for Closure Compiler "advanced" mode.
  if (typeof exports !== 'undefined') {
    if (typeof module !== 'undefined' && module.exports) {
      exports = module.exports = _;
    }
    exports._ = _;
  } else {
    root['_'] = _;
  }

  // Current version.
  _.VERSION = '1.3.1';

  // Collection Functions
  // --------------------

  // The cornerstone, an `each` implementation, aka `forEach`.
  // Handles objects with the built-in `forEach`, arrays, and raw objects.
  // Delegates to **ECMAScript 5**'s native `forEach` if available.
  var each = _.each = _.forEach = function(obj, iterator, context) {
    if (obj == null) return;
    if (nativeForEach && obj.forEach === nativeForEach) {
      obj.forEach(iterator, context);
    } else if (obj.length === +obj.length) {
      for (var i = 0, l = obj.length; i < l; i++) {
        if (i in obj && iterator.call(context, obj[i], i, obj) === breaker) return;
      }
    } else {
      for (var key in obj) {
        if (_.has(obj, key)) {
          if (iterator.call(context, obj[key], key, obj) === breaker) return;
        }
      }
    }
  };

  // Return the results of applying the iterator to each element.
  // Delegates to **ECMAScript 5**'s native `map` if available.
  _.map = _.collect = function(obj, iterator, context) {
    var results = [];
    if (obj == null) return results;
    if (nativeMap && obj.map === nativeMap) return obj.map(iterator, context);
    each(obj, function(value, index, list) {
      results[results.length] = iterator.call(context, value, index, list);
    });
    if (obj.length === +obj.length) results.length = obj.length;
    return results;
  };

  // **Reduce** builds up a single result from a list of values, aka `inject`,
  // or `foldl`. Delegates to **ECMAScript 5**'s native `reduce` if available.
  _.reduce = _.foldl = _.inject = function(obj, iterator, memo, context) {
    var initial = arguments.length > 2;
    if (obj == null) obj = [];
    if (nativeReduce && obj.reduce === nativeReduce) {
      if (context) iterator = _.bind(iterator, context);
      return initial ? obj.reduce(iterator, memo) : obj.reduce(iterator);
    }
    each(obj, function(value, index, list) {
      if (!initial) {
        memo = value;
        initial = true;
      } else {
        memo = iterator.call(context, memo, value, index, list);
      }
    });
    if (!initial) throw new TypeError('Reduce of empty array with no initial value');
    return memo;
  };

  // The right-associative version of reduce, also known as `foldr`.
  // Delegates to **ECMAScript 5**'s native `reduceRight` if available.
  _.reduceRight = _.foldr = function(obj, iterator, memo, context) {
    var initial = arguments.length > 2;
    if (obj == null) obj = [];
    if (nativeReduceRight && obj.reduceRight === nativeReduceRight) {
      if (context) iterator = _.bind(iterator, context);
      return initial ? obj.reduceRight(iterator, memo) : obj.reduceRight(iterator);
    }
    var reversed = _.toArray(obj).reverse();
    if (context && !initial) iterator = _.bind(iterator, context);
    return initial ? _.reduce(reversed, iterator, memo, context) : _.reduce(reversed, iterator);
  };

  // Return the first value which passes a truth test. Aliased as `detect`.
  _.find = _.detect = function(obj, iterator, context) {
    var result;
    any(obj, function(value, index, list) {
      if (iterator.call(context, value, index, list)) {
        result = value;
        return true;
      }
    });
    return result;
  };

  // Return all the elements that pass a truth test.
  // Delegates to **ECMAScript 5**'s native `filter` if available.
  // Aliased as `select`.
  _.filter = _.select = function(obj, iterator, context) {
    var results = [];
    if (obj == null) return results;
    if (nativeFilter && obj.filter === nativeFilter) return obj.filter(iterator, context);
    each(obj, function(value, index, list) {
      if (iterator.call(context, value, index, list)) results[results.length] = value;
    });
    return results;
  };

  // Return all the elements for which a truth test fails.
  _.reject = function(obj, iterator, context) {
    var results = [];
    if (obj == null) return results;
    each(obj, function(value, index, list) {
      if (!iterator.call(context, value, index, list)) results[results.length] = value;
    });
    return results;
  };

  // Determine whether all of the elements match a truth test.
  // Delegates to **ECMAScript 5**'s native `every` if available.
  // Aliased as `all`.
  _.every = _.all = function(obj, iterator, context) {
    var result = true;
    if (obj == null) return result;
    if (nativeEvery && obj.every === nativeEvery) return obj.every(iterator, context);
    each(obj, function(value, index, list) {
      if (!(result = result && iterator.call(context, value, index, list))) return breaker;
    });
    return result;
  };

  // Determine if at least one element in the object matches a truth test.
  // Delegates to **ECMAScript 5**'s native `some` if available.
  // Aliased as `any`.
  var any = _.some = _.any = function(obj, iterator, context) {
    iterator || (iterator = _.identity);
    var result = false;
    if (obj == null) return result;
    if (nativeSome && obj.some === nativeSome) return obj.some(iterator, context);
    each(obj, function(value, index, list) {
      if (result || (result = iterator.call(context, value, index, list))) return breaker;
    });
    return !!result;
  };

  // Determine if a given value is included in the array or object using `===`.
  // Aliased as `contains`.
  _.include = _.contains = function(obj, target) {
    var found = false;
    if (obj == null) return found;
    if (nativeIndexOf && obj.indexOf === nativeIndexOf) return obj.indexOf(target) != -1;
    found = any(obj, function(value) {
      return value === target;
    });
    return found;
  };

  // Invoke a method (with arguments) on every item in a collection.
  _.invoke = function(obj, method) {
    var args = slice.call(arguments, 2);
    return _.map(obj, function(value) {
      return (_.isFunction(method) ? method || value : value[method]).apply(value, args);
    });
  };

  // Convenience version of a common use case of `map`: fetching a property.
  _.pluck = function(obj, key) {
    return _.map(obj, function(value){ return value[key]; });
  };

  // Return the maximum element or (element-based computation).
  _.max = function(obj, iterator, context) {
    if (!iterator && _.isArray(obj)) return Math.max.apply(Math, obj);
    if (!iterator && _.isEmpty(obj)) return -Infinity;
    var result = {computed : -Infinity};
    each(obj, function(value, index, list) {
      var computed = iterator ? iterator.call(context, value, index, list) : value;
      computed >= result.computed && (result = {value : value, computed : computed});
    });
    return result.value;
  };

  // Return the minimum element (or element-based computation).
  _.min = function(obj, iterator, context) {
    if (!iterator && _.isArray(obj)) return Math.min.apply(Math, obj);
    if (!iterator && _.isEmpty(obj)) return Infinity;
    var result = {computed : Infinity};
    each(obj, function(value, index, list) {
      var computed = iterator ? iterator.call(context, value, index, list) : value;
      computed < result.computed && (result = {value : value, computed : computed});
    });
    return result.value;
  };

  // Shuffle an array.
  _.shuffle = function(obj) {
    var shuffled = [], rand;
    each(obj, function(value, index, list) {
      if (index == 0) {
        shuffled[0] = value;
      } else {
        rand = Math.floor(Math.random() * (index + 1));
        shuffled[index] = shuffled[rand];
        shuffled[rand] = value;
      }
    });
    return shuffled;
  };

  // Sort the object's values by a criterion produced by an iterator.
  _.sortBy = function(obj, iterator, context) {
    return _.pluck(_.map(obj, function(value, index, list) {
      return {
        value : value,
        criteria : iterator.call(context, value, index, list)
      };
    }).sort(function(left, right) {
      var a = left.criteria, b = right.criteria;
      return a < b ? -1 : a > b ? 1 : 0;
    }), 'value');
  };

  // Groups the object's values by a criterion. Pass either a string attribute
  // to group by, or a function that returns the criterion.
  _.groupBy = function(obj, val) {
    var result = {};
    var iterator = _.isFunction(val) ? val : function(obj) { return obj[val]; };
    each(obj, function(value, index) {
      var key = iterator(value, index);
      (result[key] || (result[key] = [])).push(value);
    });
    return result;
  };

  // Use a comparator function to figure out at what index an object should
  // be inserted so as to maintain order. Uses binary search.
  _.sortedIndex = function(array, obj, iterator) {
    iterator || (iterator = _.identity);
    var low = 0, high = array.length;
    while (low < high) {
      var mid = (low + high) >> 1;
      iterator(array[mid]) < iterator(obj) ? low = mid + 1 : high = mid;
    }
    return low;
  };

  // Safely convert anything iterable into a real, live array.
  _.toArray = function(iterable) {
    if (!iterable)                return [];
    if (iterable.toArray)         return iterable.toArray();
    if (_.isArray(iterable))      return slice.call(iterable);
    if (_.isArguments(iterable))  return slice.call(iterable);
    return _.values(iterable);
  };

  // Return the number of elements in an object.
  _.size = function(obj) {
    return _.toArray(obj).length;
  };

  // Array Functions
  // ---------------

  // Get the first element of an array. Passing **n** will return the first N
  // values in the array. Aliased as `head`. The **guard** check allows it to work
  // with `_.map`.
  _.first = _.head = function(array, n, guard) {
    return (n != null) && !guard ? slice.call(array, 0, n) : array[0];
  };

  // Returns everything but the last entry of the array. Especcialy useful on
  // the arguments object. Passing **n** will return all the values in
  // the array, excluding the last N. The **guard** check allows it to work with
  // `_.map`.
  _.initial = function(array, n, guard) {
    return slice.call(array, 0, array.length - ((n == null) || guard ? 1 : n));
  };

  // Get the last element of an array. Passing **n** will return the last N
  // values in the array. The **guard** check allows it to work with `_.map`.
  _.last = function(array, n, guard) {
    if ((n != null) && !guard) {
      return slice.call(array, Math.max(array.length - n, 0));
    } else {
      return array[array.length - 1];
    }
  };

  // Returns everything but the first entry of the array. Aliased as `tail`.
  // Especially useful on the arguments object. Passing an **index** will return
  // the rest of the values in the array from that index onward. The **guard**
  // check allows it to work with `_.map`.
  _.rest = _.tail = function(array, index, guard) {
    return slice.call(array, (index == null) || guard ? 1 : index);
  };

  // Trim out all falsy values from an array.
  _.compact = function(array) {
    return _.filter(array, function(value){ return !!value; });
  };

  // Return a completely flattened version of an array.
  _.flatten = function(array, shallow) {
    return _.reduce(array, function(memo, value) {
      if (_.isArray(value)) return memo.concat(shallow ? value : _.flatten(value));
      memo[memo.length] = value;
      return memo;
    }, []);
  };

  // Return a version of the array that does not contain the specified value(s).
  _.without = function(array) {
    return _.difference(array, slice.call(arguments, 1));
  };

  // Produce a duplicate-free version of the array. If the array has already
  // been sorted, you have the option of using a faster algorithm.
  // Aliased as `unique`.
  _.uniq = _.unique = function(array, isSorted, iterator) {
    var initial = iterator ? _.map(array, iterator) : array;
    var result = [];
    _.reduce(initial, function(memo, el, i) {
      if (0 == i || (isSorted === true ? _.last(memo) != el : !_.include(memo, el))) {
        memo[memo.length] = el;
        result[result.length] = array[i];
      }
      return memo;
    }, []);
    return result;
  };

  // Produce an array that contains the union: each distinct element from all of
  // the passed-in arrays.
  _.union = function() {
    return _.uniq(_.flatten(arguments, true));
  };

  // Produce an array that contains every item shared between all the
  // passed-in arrays. (Aliased as "intersect" for back-compat.)
  _.intersection = _.intersect = function(array) {
    var rest = slice.call(arguments, 1);
    return _.filter(_.uniq(array), function(item) {
      return _.every(rest, function(other) {
        return _.indexOf(other, item) >= 0;
      });
    });
  };

  // Take the difference between one array and a number of other arrays.
  // Only the elements present in just the first array will remain.
  _.difference = function(array) {
    var rest = _.flatten(slice.call(arguments, 1));
    return _.filter(array, function(value){ return !_.include(rest, value); });
  };

  // Zip together multiple lists into a single array -- elements that share
  // an index go together.
  _.zip = function() {
    var args = slice.call(arguments);
    var length = _.max(_.pluck(args, 'length'));
    var results = new Array(length);
    for (var i = 0; i < length; i++) results[i] = _.pluck(args, "" + i);
    return results;
  };

  // If the browser doesn't supply us with indexOf (I'm looking at you, **MSIE**),
  // we need this function. Return the position of the first occurrence of an
  // item in an array, or -1 if the item is not included in the array.
  // Delegates to **ECMAScript 5**'s native `indexOf` if available.
  // If the array is large and already in sort order, pass `true`
  // for **isSorted** to use binary search.
  _.indexOf = function(array, item, isSorted) {
    if (array == null) return -1;
    var i, l;
    if (isSorted) {
      i = _.sortedIndex(array, item);
      return array[i] === item ? i : -1;
    }
    if (nativeIndexOf && array.indexOf === nativeIndexOf) return array.indexOf(item);
    for (i = 0, l = array.length; i < l; i++) if (i in array && array[i] === item) return i;
    return -1;
  };

  // Delegates to **ECMAScript 5**'s native `lastIndexOf` if available.
  _.lastIndexOf = function(array, item) {
    if (array == null) return -1;
    if (nativeLastIndexOf && array.lastIndexOf === nativeLastIndexOf) return array.lastIndexOf(item);
    var i = array.length;
    while (i--) if (i in array && array[i] === item) return i;
    return -1;
  };

  // Generate an integer Array containing an arithmetic progression. A port of
  // the native Python `range()` function. See
  // [the Python documentation](http://docs.python.org/library/functions.html#range).
  _.range = function(start, stop, step) {
    if (arguments.length <= 1) {
      stop = start || 0;
      start = 0;
    }
    step = arguments[2] || 1;

    var len = Math.max(Math.ceil((stop - start) / step), 0);
    var idx = 0;
    var range = new Array(len);

    while(idx < len) {
      range[idx++] = start;
      start += step;
    }

    return range;
  };

  // Function (ahem) Functions
  // ------------------

  // Reusable constructor function for prototype setting.
  var ctor = function(){};

  // Create a function bound to a given object (assigning `this`, and arguments,
  // optionally). Binding with arguments is also known as `curry`.
  // Delegates to **ECMAScript 5**'s native `Function.bind` if available.
  // We check for `func.bind` first, to fail fast when `func` is undefined.
  _.bind = function bind(func, context) {
    var bound, args;
    if (func.bind === nativeBind && nativeBind) return nativeBind.apply(func, slice.call(arguments, 1));
    if (!_.isFunction(func)) throw new TypeError;
    args = slice.call(arguments, 2);
    return bound = function() {
      if (!(this instanceof bound)) return func.apply(context, args.concat(slice.call(arguments)));
      ctor.prototype = func.prototype;
      var self = new ctor;
      var result = func.apply(self, args.concat(slice.call(arguments)));
      if (Object(result) === result) return result;
      return self;
    };
  };

  // Bind all of an object's methods to that object. Useful for ensuring that
  // all callbacks defined on an object belong to it.
  _.bindAll = function(obj) {
    var funcs = slice.call(arguments, 1);
    if (funcs.length == 0) funcs = _.functions(obj);
    each(funcs, function(f) { obj[f] = _.bind(obj[f], obj); });
    return obj;
  };

  // Memoize an expensive function by storing its results.
  _.memoize = function(func, hasher) {
    var memo = {};
    hasher || (hasher = _.identity);
    return function() {
      var key = hasher.apply(this, arguments);
      return _.has(memo, key) ? memo[key] : (memo[key] = func.apply(this, arguments));
    };
  };

  // Delays a function for the given number of milliseconds, and then calls
  // it with the arguments supplied.
  _.delay = function(func, wait) {
    var args = slice.call(arguments, 2);
    return setTimeout(function(){ return func.apply(func, args); }, wait);
  };

  // Defers a function, scheduling it to run after the current call stack has
  // cleared.
  _.defer = function(func) {
    return _.delay.apply(_, [func, 1].concat(slice.call(arguments, 1)));
  };

  // Returns a function, that, when invoked, will only be triggered at most once
  // during a given window of time.
  _.throttle = function(func, wait) {
    var context, args, timeout, throttling, more;
    var whenDone = _.debounce(function(){ more = throttling = false; }, wait);
    return function() {
      context = this; args = arguments;
      var later = function() {
        timeout = null;
        if (more) func.apply(context, args);
        whenDone();
      };
      if (!timeout) timeout = setTimeout(later, wait);
      if (throttling) {
        more = true;
      } else {
        func.apply(context, args);
      }
      whenDone();
      throttling = true;
    };
  };

  // Returns a function, that, as long as it continues to be invoked, will not
  // be triggered. The function will be called after it stops being called for
  // N milliseconds.
  _.debounce = function(func, wait) {
    var timeout;
    return function() {
      var context = this, args = arguments;
      var later = function() {
        timeout = null;
        func.apply(context, args);
      };
      clearTimeout(timeout);
      timeout = setTimeout(later, wait);
    };
  };

  // Returns a function that will be executed at most one time, no matter how
  // often you call it. Useful for lazy initialization.
  _.once = function(func) {
    var ran = false, memo;
    return function() {
      if (ran) return memo;
      ran = true;
      return memo = func.apply(this, arguments);
    };
  };

  // Returns the first function passed as an argument to the second,
  // allowing you to adjust arguments, run code before and after, and
  // conditionally execute the original function.
  _.wrap = function(func, wrapper) {
    return function() {
      var args = [func].concat(slice.call(arguments, 0));
      return wrapper.apply(this, args);
    };
  };

  // Returns a function that is the composition of a list of functions, each
  // consuming the return value of the function that follows.
  _.compose = function() {
    var funcs = arguments;
    return function() {
      var args = arguments;
      for (var i = funcs.length - 1; i >= 0; i--) {
        args = [funcs[i].apply(this, args)];
      }
      return args[0];
    };
  };

  // Returns a function that will only be executed after being called N times.
  _.after = function(times, func) {
    if (times <= 0) return func();
    return function() {
      if (--times < 1) { return func.apply(this, arguments); }
    };
  };

  // Object Functions
  // ----------------

  // Retrieve the names of an object's properties.
  // Delegates to **ECMAScript 5**'s native `Object.keys`
  _.keys = nativeKeys || function(obj) {
    if (obj !== Object(obj)) throw new TypeError('Invalid object');
    var keys = [];
    for (var key in obj) if (_.has(obj, key)) keys[keys.length] = key;
    return keys;
  };

  // Retrieve the values of an object's properties.
  _.values = function(obj) {
    return _.map(obj, _.identity);
  };

  // Return a sorted list of the function names available on the object.
  // Aliased as `methods`
  _.functions = _.methods = function(obj) {
    var names = [];
    for (var key in obj) {
      if (_.isFunction(obj[key])) names.push(key);
    }
    return names.sort();
  };

  // Extend a given object with all the properties in passed-in object(s).
  _.extend = function(obj) {
    each(slice.call(arguments, 1), function(source) {
      for (var prop in source) {
        obj[prop] = source[prop];
      }
    });
    return obj;
  };

  // Fill in a given object with default properties.
  _.defaults = function(obj) {
    each(slice.call(arguments, 1), function(source) {
      for (var prop in source) {
        if (obj[prop] == null) obj[prop] = source[prop];
      }
    });
    return obj;
  };

  // Create a (shallow-cloned) duplicate of an object.
  _.clone = function(obj) {
    if (!_.isObject(obj)) return obj;
    return _.isArray(obj) ? obj.slice() : _.extend({}, obj);
  };

  // Invokes interceptor with the obj, and then returns obj.
  // The primary purpose of this method is to "tap into" a method chain, in
  // order to perform operations on intermediate results within the chain.
  _.tap = function(obj, interceptor) {
    interceptor(obj);
    return obj;
  };

  // Internal recursive comparison function.
  function eq(a, b, stack) {
    // Identical objects are equal. `0 === -0`, but they aren't identical.
    // See the Harmony `egal` proposal: http://wiki.ecmascript.org/doku.php?id=harmony:egal.
    if (a === b) return a !== 0 || 1 / a == 1 / b;
    // A strict comparison is necessary because `null == undefined`.
    if (a == null || b == null) return a === b;
    // Unwrap any wrapped objects.
    if (a._chain) a = a._wrapped;
    if (b._chain) b = b._wrapped;
    // Invoke a custom `isEqual` method if one is provided.
    if (a.isEqual && _.isFunction(a.isEqual)) return a.isEqual(b);
    if (b.isEqual && _.isFunction(b.isEqual)) return b.isEqual(a);
    // Compare `[[Class]]` names.
    var className = toString.call(a);
    if (className != toString.call(b)) return false;
    switch (className) {
      // Strings, numbers, dates, and booleans are compared by value.
      case '[object String]':
        // Primitives and their corresponding object wrappers are equivalent; thus, `"5"` is
        // equivalent to `new String("5")`.
        return a == String(b);
      case '[object Number]':
        // `NaN`s are equivalent, but non-reflexive. An `egal` comparison is performed for
        // other numeric values.
        return a != +a ? b != +b : (a == 0 ? 1 / a == 1 / b : a == +b);
      case '[object Date]':
      case '[object Boolean]':
        // Coerce dates and booleans to numeric primitive values. Dates are compared by their
        // millisecond representations. Note that invalid dates with millisecond representations
        // of `NaN` are not equivalent.
        return +a == +b;
      // RegExps are compared by their source patterns and flags.
      case '[object RegExp]':
        return a.source == b.source &&
               a.global == b.global &&
               a.multiline == b.multiline &&
               a.ignoreCase == b.ignoreCase;
    }
    if (typeof a != 'object' || typeof b != 'object') return false;
    // Assume equality for cyclic structures. The algorithm for detecting cyclic
    // structures is adapted from ES 5.1 section 15.12.3, abstract operation `JO`.
    var length = stack.length;
    while (length--) {
      // Linear search. Performance is inversely proportional to the number of
      // unique nested structures.
      if (stack[length] == a) return true;
    }
    // Add the first object to the stack of traversed objects.
    stack.push(a);
    var size = 0, result = true;
    // Recursively compare objects and arrays.
    if (className == '[object Array]') {
      // Compare array lengths to determine if a deep comparison is necessary.
      size = a.length;
      result = size == b.length;
      if (result) {
        // Deep compare the contents, ignoring non-numeric properties.
        while (size--) {
          // Ensure commutative equality for sparse arrays.
          if (!(result = size in a == size in b && eq(a[size], b[size], stack))) break;
        }
      }
    } else {
      // Objects with different constructors are not equivalent.
      if ('constructor' in a != 'constructor' in b || a.constructor != b.constructor) return false;
      // Deep compare objects.
      for (var key in a) {
        if (_.has(a, key)) {
          // Count the expected number of properties.
          size++;
          // Deep compare each member.
          if (!(result = _.has(b, key) && eq(a[key], b[key], stack))) break;
        }
      }
      // Ensure that both objects contain the same number of properties.
      if (result) {
        for (key in b) {
          if (_.has(b, key) && !(size--)) break;
        }
        result = !size;
      }
    }
    // Remove the first object from the stack of traversed objects.
    stack.pop();
    return result;
  }

  // Perform a deep comparison to check if two objects are equal.
  _.isEqual = function(a, b) {
    return eq(a, b, []);
  };

  // Is a given array, string, or object empty?
  // An "empty" object has no enumerable own-properties.
  _.isEmpty = function(obj) {
    if (_.isArray(obj) || _.isString(obj)) return obj.length === 0;
    for (var key in obj) if (_.has(obj, key)) return false;
    return true;
  };

  // Is a given value a DOM element?
  _.isElement = function(obj) {
    return !!(obj && obj.nodeType == 1);
  };

  // Is a given value an array?
  // Delegates to ECMA5's native Array.isArray
  _.isArray = nativeIsArray || function(obj) {
    return toString.call(obj) == '[object Array]';
  };

  // Is a given variable an object?
  _.isObject = function(obj) {
    return obj === Object(obj);
  };

  // Is a given variable an arguments object?
  _.isArguments = function(obj) {
    return toString.call(obj) == '[object Arguments]';
  };
  if (!_.isArguments(arguments)) {
    _.isArguments = function(obj) {
      return !!(obj && _.has(obj, 'callee'));
    };
  }

  // Is a given value a function?
  _.isFunction = function(obj) {
    return toString.call(obj) == '[object Function]';
  };

  // Is a given value a string?
  _.isString = function(obj) {
    return toString.call(obj) == '[object String]';
  };

  // Is a given value a number?
  _.isNumber = function(obj) {
    return toString.call(obj) == '[object Number]';
  };

  // Is the given value `NaN`?
  _.isNaN = function(obj) {
    // `NaN` is the only value for which `===` is not reflexive.
    return obj !== obj;
  };

  // Is a given value a boolean?
  _.isBoolean = function(obj) {
    return obj === true || obj === false || toString.call(obj) == '[object Boolean]';
  };

  // Is a given value a date?
  _.isDate = function(obj) {
    return toString.call(obj) == '[object Date]';
  };

  // Is the given value a regular expression?
  _.isRegExp = function(obj) {
    return toString.call(obj) == '[object RegExp]';
  };

  // Is a given value equal to null?
  _.isNull = function(obj) {
    return obj === null;
  };

  // Is a given variable undefined?
  _.isUndefined = function(obj) {
    return obj === void 0;
  };

  // Has own property?
  _.has = function(obj, key) {
    return hasOwnProperty.call(obj, key);
  };

  // Utility Functions
  // -----------------

  // Run Underscore.js in *noConflict* mode, returning the `_` variable to its
  // previous owner. Returns a reference to the Underscore object.
  _.noConflict = function() {
    root._ = previousUnderscore;
    return this;
  };

  // Keep the identity function around for default iterators.
  _.identity = function(value) {
    return value;
  };

  // Run a function **n** times.
  _.times = function (n, iterator, context) {
    for (var i = 0; i < n; i++) iterator.call(context, i);
  };

  // Escape a string for HTML interpolation.
  _.escape = function(string) {
    return (''+string).replace(/&/g, '&amp;').replace(/</g, '&lt;').replace(/>/g, '&gt;').replace(/"/g, '&quot;').replace(/'/g, '&#x27;').replace(/\//g,'&#x2F;');
  };

  // Add your own custom functions to the Underscore object, ensuring that
  // they're correctly added to the OOP wrapper as well.
  _.mixin = function(obj) {
    each(_.functions(obj), function(name){
      addToWrapper(name, _[name] = obj[name]);
    });
  };

  // Generate a unique integer id (unique within the entire client session).
  // Useful for temporary DOM ids.
  var idCounter = 0;
  _.uniqueId = function(prefix) {
    var id = idCounter++;
    return prefix ? prefix + id : id;
  };

  // By default, Underscore uses ERB-style template delimiters, change the
  // following template settings to use alternative delimiters.
  _.templateSettings = {
    evaluate    : /<%([\s\S]+?)%>/g,
    interpolate : /<%=([\s\S]+?)%>/g,
    escape      : /<%-([\s\S]+?)%>/g
  };

  // When customizing `templateSettings`, if you don't want to define an
  // interpolation, evaluation or escaping regex, we need one that is
  // guaranteed not to match.
  var noMatch = /.^/;

  // Within an interpolation, evaluation, or escaping, remove HTML escaping
  // that had been previously added.
  var unescape = function(code) {
    return code.replace(/\\\\/g, '\\').replace(/\\'/g, "'");
  };

  // JavaScript micro-templating, similar to John Resig's implementation.
  // Underscore templating handles arbitrary delimiters, preserves whitespace,
  // and correctly escapes quotes within interpolated code.
  _.template = function(str, data) {
    var c  = _.templateSettings;
    var tmpl = 'var __p=[],print=function(){__p.push.apply(__p,arguments);};' +
      'with(obj||{}){__p.push(\'' +
      str.replace(/\\/g, '\\\\')
         .replace(/'/g, "\\'")
         .replace(c.escape || noMatch, function(match, code) {
           return "',_.escape(" + unescape(code) + "),'";
         })
         .replace(c.interpolate || noMatch, function(match, code) {
           return "'," + unescape(code) + ",'";
         })
         .replace(c.evaluate || noMatch, function(match, code) {
           return "');" + unescape(code).replace(/[\r\n\t]/g, ' ') + ";__p.push('";
         })
         .replace(/\r/g, '\\r')
         .replace(/\n/g, '\\n')
         .replace(/\t/g, '\\t')
         + "');}return __p.join('');";
    var func = new Function('obj', '_', tmpl);
    if (data) return func(data, _);
    return function(data) {
      return func.call(this, data, _);
    };
  };

  // Add a "chain" function, which will delegate to the wrapper.
  _.chain = function(obj) {
    return _(obj).chain();
  };

  // The OOP Wrapper
  // ---------------

  // If Underscore is called as a function, it returns a wrapped object that
  // can be used OO-style. This wrapper holds altered versions of all the
  // underscore functions. Wrapped objects may be chained.
  var wrapper = function(obj) { this._wrapped = obj; };

  // Expose `wrapper.prototype` as `_.prototype`
  _.prototype = wrapper.prototype;

  // Helper function to continue chaining intermediate results.
  var result = function(obj, chain) {
    return chain ? _(obj).chain() : obj;
  };

  // A method to easily add functions to the OOP wrapper.
  var addToWrapper = function(name, func) {
    wrapper.prototype[name] = function() {
      var args = slice.call(arguments);
      unshift.call(args, this._wrapped);
      return result(func.apply(_, args), this._chain);
    };
  };

  // Add all of the Underscore functions to the wrapper object.
  _.mixin(_);

  // Add all mutator Array functions to the wrapper.
  each(['pop', 'push', 'reverse', 'shift', 'sort', 'splice', 'unshift'], function(name) {
    var method = ArrayProto[name];
    wrapper.prototype[name] = function() {
      var wrapped = this._wrapped;
      method.apply(wrapped, arguments);
      var length = wrapped.length;
      if ((name == 'shift' || name == 'splice') && length === 0) delete wrapped[0];
      return result(wrapped, this._chain);
    };
  });

  // Add all accessor Array functions to the wrapper.
  each(['concat', 'join', 'slice'], function(name) {
    var method = ArrayProto[name];
    wrapper.prototype[name] = function() {
      return result(method.apply(this._wrapped, arguments), this._chain);
    };
  });

  // Start chaining a wrapped Underscore object.
  wrapper.prototype.chain = function() {
    this._chain = true;
    return this;
  };

  // Extracts the result from a wrapped and chained object.
  wrapper.prototype.value = function() {
    return this._wrapped;
  };

}).call(this);
```
