## Supplementary material for "ΔSCOPE: A new method to quantify 3D biological structures and identify differences in zebrafish forebrain development": An archived copy of the DeltaSCOPE code repository.: underscore.js

```
// Underscore.js 1.3.1
// (c) 2009-2012 Jeremy Ashkenas, DocumentCloud Inc.
// Underscore is freely distributable under the MIT license.
// Portions of Underscore are inspired or borrowed from Prototype,
// Oliver Steele's Functional, and John Resig's Micro-Templating.
// For all details and documentation:
// http://documentcloud.github.com/underscore
(function(){function q(a,c,d){if(a===c)return a!==0||1/a==1/c;if(a==null||c==null)return a===c;if(a._chain)a=a._wrapped;if(c._chain)c=c._wrapped;if(a.isEqual&&b.isFunction(a.isEqual))return a.isEqual(c);if(c.isEqual&&b.isFunction(c.isEqual))return c.isEqual(a);var e=l.call(a);if(e!=l.call(c))return false;switch(e){case "[object String]":return a==String(c);case "[object Number]":return a!=+a?c!=+c:a==0?1/a==1/c:a==+c;case "[object Date]":case "[object Boolean]":return+a==+c;case "[object RegExp]":return a.source==
c.source&&a.global==c.global&&a.multiline==c.multiline&&a.ignoreCase==c.ignoreCase}if(typeof a!="object"||typeof c!="object")return false;for(var f=d.length;f--;)if(d[f]==a)return true;d.push(a);var f=0,g=true;if(e=="[object Array]"){if(f=a.length,g=f==c.length)for(;f--;)if(!(g=f in a==f in c&&q(a[f],c[f],d)))break}else{if("constructor"in a!="constructor"in c||a.constructor!=c.constructor)return false;for(var h in a)if(b.has(a,h)&&(f++,!(g=b.has(c,h)&&q(a[h],c[h],d))))break;if(g){for(h in c)if(b.has(c,
h)&&!f--)break;g=!f}}d.pop();return g}var r=this,G=r._,n={},k=Array.prototype,o=Object.prototype,i=k.slice,H=k.unshift,l=o.toString,I=o.hasOwnProperty,w=k.forEach,x=k.map,y=k.reduce,z=k.reduceRight,A=k.filter,B=k.every,C=k.some,p=k.indexOf,D=k.lastIndexOf,o=Array.isArray,J=Object.keys,s=Function.prototype.bind,b=function(a){return new m(a)};if(typeof exports!=="undefined"){if(typeof module!=="undefined"&&module.exports)exports=module.exports=b;exports._=b}else r._=b;b.VERSION="1.3.1";var j=b.each=
b.forEach=function(a,c,d){if(a!=null)if(w&&a.forEach===w)a.forEach(c,d);else if(a.length===+a.length)for(var e=0,f=a.length;e<f;e++){if(e in a&&c.call(d,a[e],e,a)===n)break}else for(e in a)if(b.has(a,e)&&c.call(d,a[e],e,a)===n)break};b.map=b.collect=function(a,c,b){var e=[];if(a==null)return e;if(x&&a.map===x)return a.map(c,b);j(a,function(a,g,h){e[e.length]=c.call(b,a,g,h)});if(a.length===+a.length)e.length=a.length;return e};b.reduce=b.foldl=b.inject=function(a,c,d,e){var f=arguments.length>2;a==
null&&(a=[]);if(y&&a.reduce===y)return e&&(c=b.bind(c,e)),f?a.reduce(c,d):a.reduce(c);j(a,function(a,b,i){f?d=c.call(e,d,a,b,i):(d=a,f=true)});if(!f)throw new TypeError("Reduce of empty array with no initial value");return d};b.reduceRight=b.foldr=function(a,c,d,e){var f=arguments.length>2;a==null&&(a=[]);if(z&&a.reduceRight===z)return e&&(c=b.bind(c,e)),f?a.reduceRight(c,d):a.reduceRight(c);var g=b.toArray(a).reverse();e&&!f&&(c=b.bind(c,e));return f?b.reduce(g,c,d,e):b.reduce(g,c)};b.find=b.detect=
function(a,c,b){var e;E(a,function(a,g,h){if(c.call(b,a,g,h))return e=a,true});return e};b.filter=b.select=function(a,c,b){var e=[];if(a==null)return e;if(A&&a.filter===A)return a.filter(c,b);j(a,function(a,g,h){c.call(b,a,g,h)&&(e[e.length]=a)});return e};b.reject=function(a,c,b){var e=[];if(a==null)return e;j(a,function(a,g,h){c.call(b,a,g,h)||(e[e.length]=a)});return e};b.every=b.all=function(a,c,b){var e=true;if(a==null)return e;if(B&&a.every===B)return a.every(c,b);j(a,function(a,g,h){if(!(e=
e&&c.call(b,a,g,h)))return n});return e};var E=b.some=b.any=function(a,c,d){c||(c=b.identity);var e=false;if(a==null)return e;if(C&&a.some===C)return a.some(c,d);j(a,function(a,b,h){if(e||(e=c.call(d,a,b,h)))return n});return!!e};b.include=b.contains=function(a,c){var b=false;if(a==null)return b;return p&&a.indexOf===p?a.indexOf(c)!=-1:b=E(a,function(a){return a===c})};b.invoke=function(a,c){var d=i.call(arguments,2);return b.map(a,function(a){return(b.isFunction(c)?c||a:a[c]).apply(a,d)})};b.pluck=
function(a,c){return b.map(a,function(a){return a[c]})};b.max=function(a,c,d){if(!c&&b.isArray(a))return Math.max.apply(Math,a);if(!c&&b.isEmpty(a))return-Infinity;var e={computed:-Infinity};j(a,function(a,b,h){b=c?c.call(d,a,b,h):a;b>=e.computed&&(e={value:a,computed:b})});return e.value};b.min=function(a,c,d){if(!c&&b.isArray(a))return Math.min.apply(Math,a);if(!c&&b.isEmpty(a))return Infinity;var e={computed:Infinity};j(a,function(a,b,h){b=c?c.call(d,a,b,h):a;b<e.computed&&(e={value:a,computed:b})});
return e.value};b.shuffle=function(a){var b=[],d;j(a,function(a,f){f==0?b[0]=a:(d=Math.floor(Math.random()*(f+1)),b[f]=b[d],b[d]=a)});return b};b.sortBy=function(a,c,d){return b.pluck(b.map(a,function(a,b,g){return{value:a,criteria:c.call(d,a,b,g)}}).sort(function(a,b){var c=a.criteria,d=b.criteria;return c<d?-1:c>d?1:0}),"value")};b.groupBy=function(a,c){var d={},e=b.isFunction(c)?c:function(a){return a[c]};j(a,function(a,b){var c=e(a,b);(d[c]||(d[c]=[])).push(a)});return d};b.sortedIndex=function(a,
c,d){d||(d=b.identity);for(var e=0,f=a.length;e<f;){var g=e+f>>1;d(a[g])<d(c)?e=g+1:f=g}return e};b.toArray=function(a){return!a?[]:a.toArray?a.toArray():b.isArray(a)?i.call(a):b.isArguments(a)?i.call(a):b.values(a)};b.size=function(a){return b.toArray(a).length};b.first=b.head=function(a,b,d){return b!=null&&!d?i.call(a,0,b):a[0]};b.initial=function(a,b,d){return i.call(a,0,a.length-(b==null||d?1:b))};b.last=function(a,b,d){return b!=null&&!d?i.call(a,Math.max(a.length-b,0)):a[a.length-1]};b.rest=
b.tail=function(a,b,d){return i.call(a,b==null||d?1:b)};b.compact=function(a){return b.filter(a,function(a){return!!a})};b.flatten=function(a,c){return b.reduce(a,function(a,e){if(b.isArray(e))return a.concat(c?e:b.flatten(e));a[a.length]=e;return a},[])};b.without=function(a){return b.difference(a,i.call(arguments,1))};b.uniq=b.unique=function(a,c,d){var d=d?b.map(a,d):a,e=[];b.reduce(d,function(d,g,h){if(0==h||(c===true?b.last(d)!=g:!b.include(d,g)))d[d.length]=g,e[e.length]=a[h];return d},[]);
return e};b.union=function(){return b.uniq(b.flatten(arguments,true))};b.intersection=b.intersect=function(a){var c=i.call(arguments,1);return b.filter(b.uniq(a),function(a){return b.every(c,function(c){return b.indexOf(c,a)>=0})})};b.difference=function(a){var c=b.flatten(i.call(arguments,1));return b.filter(a,function(a){return!b.include(c,a)})};b.zip=function(){for(var a=i.call(arguments),c=b.max(b.pluck(a,"length")),d=Array(c),e=0;e<c;e++)d[e]=b.pluck(a,""+e);return d};b.indexOf=function(a,c,
d){if(a==null)return-1;var e;if(d)return d=b.sortedIndex(a,c),a[d]===c?d:-1;if(p&&a.indexOf===p)return a.indexOf(c);for(d=0,e=a.length;d<e;d++)if(d in a&&a[d]===c)return d;return-1};b.lastIndexOf=function(a,b){if(a==null)return-1;if(D&&a.lastIndexOf===D)return a.lastIndexOf(b);for(var d=a.length;d--;)if(d in a&&a[d]===b)return d;return-1};b.range=function(a,b,d){arguments.length<=1&&(b=a||0,a=0);for(var d=arguments[2]||1,e=Math.max(Math.ceil((b-a)/d),0),f=0,g=Array(e);f<e;)g[f++]=a,a+=d;return g};
var F=function(){};b.bind=function(a,c){var d,e;if(a.bind===s&&s)return s.apply(a,i.call(arguments,1));if(!b.isFunction(a))throw new TypeError;e=i.call(arguments,2);return d=function(){if(!(this instanceof d))return a.apply(c,e.concat(i.call(arguments)));F.prototype=a.prototype;var b=new F,g=a.apply(b,e.concat(i.call(arguments)));return Object(g)===g?g:b}};b.bindAll=function(a){var c=i.call(arguments,1);c.length==0&&(c=b.functions(a));j(c,function(c){a[c]=b.bind(a[c],a)});return a};b.memoize=function(a,
c){var d={};c||(c=b.identity);return function(){var e=c.apply(this,arguments);return b.has(d,e)?d[e]:d[e]=a.apply(this,arguments)}};b.delay=function(a,b){var d=i.call(arguments,2);return setTimeout(function(){return a.apply(a,d)},b)};b.defer=function(a){return b.delay.apply(b,[a,1].concat(i.call(arguments,1)))};b.throttle=function(a,c){var d,e,f,g,h,i=b.debounce(function(){h=g=false},c);return function(){d=this;e=arguments;var b;f||(f=setTimeout(function(){f=null;h&&a.apply(d,e);i()},c));g?h=true:
a.apply(d,e);i();g=true}};b.debounce=function(a,b){var d;return function(){var e=this,f=arguments;clearTimeout(d);d=setTimeout(function(){d=null;a.apply(e,f)},b)}};b.once=function(a){var b=false,d;return function(){if(b)return d;b=true;return d=a.apply(this,arguments)}};b.wrap=function(a,b){return function(){var d=[a].concat(i.call(arguments,0));return b.apply(this,d)}};b.compose=function(){var a=arguments;return function(){for(var b=arguments,d=a.length-1;d>=0;d--)b=[a[d].apply(this,b)];return b[0]}};
b.after=function(a,b){return a<=0?b():function(){if(--a<1)return b.apply(this,arguments)}};b.keys=J||function(a){if(a!==Object(a))throw new TypeError("Invalid object");var c=[],d;for(d in a)b.has(a,d)&&(c[c.length]=d);return c};b.values=function(a){return b.map(a,b.identity)};b.functions=b.methods=function(a){var c=[],d;for(d in a)b.isFunction(a[d])&&c.push(d);return c.sort()};b.extend=function(a){j(i.call(arguments,1),function(b){for(var d in b)a[d]=b[d]});return a};b.defaults=function(a){j(i.call(arguments,
1),function(b){for(var d in b)a[d]==null&&(a[d]=b[d])});return a};b.clone=function(a){return!b.isObject(a)?a:b.isArray(a)?a.slice():b.extend({},a)};b.tap=function(a,b){b(a);return a};b.isEqual=function(a,b){return q(a,b,[])};b.isEmpty=function(a){if(b.isArray(a)||b.isString(a))return a.length===0;for(var c in a)if(b.has(a,c))return false;return true};b.isElement=function(a){return!!(a&&a.nodeType==1)};b.isArray=o||function(a){return l.call(a)=="[object Array]"};b.isObject=function(a){return a===Object(a)};
b.isArguments=function(a){return l.call(a)=="[object Arguments]"};if(!b.isArguments(arguments))b.isArguments=function(a){return!(!a||!b.has(a,"callee"))};b.isFunction=function(a){return l.call(a)=="[object Function]"};b.isString=function(a){return l.call(a)=="[object String]"};b.isNumber=function(a){return l.call(a)=="[object Number]"};b.isNaN=function(a){return a!==a};b.isBoolean=function(a){return a===true||a===false||l.call(a)=="[object Boolean]"};b.isDate=function(a){return l.call(a)=="[object Date]"};
b.isRegExp=function(a){return l.call(a)=="[object RegExp]"};b.isNull=function(a){return a===null};b.isUndefined=function(a){return a===void 0};b.has=function(a,b){return I.call(a,b)};b.noConflict=function(){r._=G;return this};b.identity=function(a){return a};b.times=function(a,b,d){for(var e=0;e<a;e++)b.call(d,e)};b.escape=function(a){return(""+a).replace(/&/g,"&amp;").replace(/</g,"&lt;").replace(/>/g,"&gt;").replace(/"/g,"&quot;").replace(/'/g,"&#x27;").replace(/\//g,"&#x2F;")};b.mixin=function(a){j(b.functions(a),
function(c){K(c,b[c]=a[c])})};var L=0;b.uniqueId=function(a){var b=L++;return a?a+b:b};b.templateSettings={evaluate:/<%([\s\S]+?)%>/g,interpolate:/<%=([\s\S]+?)%>/g,escape:/<%-([\s\S]+?)%>/g};var t=/.^/,u=function(a){return a.replace(/\\\\/g,"\\").replace(/\\'/g,"'")};b.template=function(a,c){var d=b.templateSettings,d="var __p=[],print=function(){__p.push.apply(__p,arguments);};with(obj||{}){__p.push('"+a.replace(/\\/g,"\\\\").replace(/'/g,"\\'").replace(d.escape||t,function(a,b){return"',_.escape("+
u(b)+"),'"}).replace(d.interpolate||t,function(a,b){return"',"+u(b)+",'"}).replace(d.evaluate||t,function(a,b){return"');"+u(b).replace(/[\r\n\t]/g," ")+";__p.push('"}).replace(/\r/g,"\\r").replace(/\n/g,"\\n").replace(/\t/g,"\\t")+"');}return __p.join('');",e=new Function("obj","_",d);return c?e(c,b):function(a){return e.call(this,a,b)}};b.chain=function(a){return b(a).chain()};var m=function(a){this._wrapped=a};b.prototype=m.prototype;var v=function(a,c){return c?b(a).chain():a},K=function(a,c){m.prototype[a]=
function(){var a=i.call(arguments);H.call(a,this._wrapped);return v(c.apply(b,a),this._chain)}};b.mixin(b);j("pop,push,reverse,shift,sort,splice,unshift".split(","),function(a){var b=k[a];m.prototype[a]=function(){var d=this._wrapped;b.apply(d,arguments);var e=d.length;(a=="shift"||a=="splice")&&e===0&&delete d[0];return v(d,this._chain)}});j(["concat","join","slice"],function(a){var b=k[a];m.prototype[a]=function(){return v(b.apply(this._wrapped,arguments),this._chain)}});m.prototype.chain=function(){this._chain=
true;return this};m.prototype.value=function(){return this._wrapped}}).call(this);
```
