## Supplementary material for "ΔSCOPE: A new method to quantify 3D biological structures and identify differences in zebrafish forebrain development": An archived copy of the DeltaSCOPE code repository.: websupport.js

```
/*
 * websupport.js
 * ~~~~~~~~~~~~~
 *
 * sphinx.websupport utilities for all documentation.
 *
 * :copyright: Copyright 2007-2018 by the Sphinx team, see AUTHORS.
 * :license: BSD, see LICENSE for details.
 *
 */

(function($) {
  $.fn.autogrow = function() {
    return this.each(function() {
    var textarea = this;

    $.fn.autogrow.resize(textarea);

    $(textarea)
      .focus(function() {
        textarea.interval = setInterval(function() {
          $.fn.autogrow.resize(textarea);
        }, 500);
      })
      .blur(function() {
        clearInterval(textarea.interval);
      });
    });
  };

  $.fn.autogrow.resize = function(textarea) {
    var lineHeight = parseInt($(textarea).css('line-height'), 10);
    var lines = textarea.value.split('\n');
    var columns = textarea.cols;
    var lineCount = 0;
    $.each(lines, function() {
      lineCount += Math.ceil(this.length / columns) || 1;
    });
    var height = lineHeight * (lineCount + 1);
    $(textarea).css('height', height);
  };
})(jQuery);

(function($) {
  var comp, by;

  function init() {
    initEvents();
    initComparator();
  }

  function initEvents() {
    $(document).on("click", 'a.comment-close', function(event) {
      event.preventDefault();
      hide($(this).attr('id').substring(2));
    });
    $(document).on("click", 'a.vote', function(event) {
      event.preventDefault();
      handleVote($(this));
    });
    $(document).on("click", 'a.reply', function(event) {
      event.preventDefault();
      openReply($(this).attr('id').substring(2));
    });
    $(document).on("click", 'a.close-reply', function(event) {
      event.preventDefault();
      closeReply($(this).attr('id').substring(2));
    });
    $(document).on("click", 'a.sort-option', function(event) {
      event.preventDefault();
      handleReSort($(this));
    });
    $(document).on("click", 'a.show-proposal', function(event) {
      event.preventDefault();
      showProposal($(this).attr('id').substring(2));
    });
    $(document).on("click", 'a.hide-proposal', function(event) {
      event.preventDefault();
      hideProposal($(this).attr('id').substring(2));
    });
    $(document).on("click", 'a.show-propose-change', function(event) {
      event.preventDefault();
      showProposeChange($(this).attr('id').substring(2));
    });
    $(document).on("click", 'a.hide-propose-change', function(event) {
      event.preventDefault();
      hideProposeChange($(this).attr('id').substring(2));
    });
    $(document).on("click", 'a.accept-comment', function(event) {
      event.preventDefault();
      acceptComment($(this).attr('id').substring(2));
    });
    $(document).on("click", 'a.delete-comment', function(event) {
      event.preventDefault();
      deleteComment($(this).attr('id').substring(2));
    });
    $(document).on("click", 'a.comment-markup', function(event) {
      event.preventDefault();
      toggleCommentMarkupBox($(this).attr('id').substring(2));
    });
  }

  /**
   * Set comp, which is a comparator function used for sorting and
   * inserting comments into the list.
   */
  function setComparator() {
    // If the first three letters are "asc", sort in ascending order
    // and remove the prefix.
    if (by.substring(0,3) == 'asc') {
      var i = by.substring(3);
      comp = function(a, b) { return a[i] - b[i]; };
    } else {
      // Otherwise sort in descending order.
      comp = function(a, b) { return b[by] - a[by]; };
    }

    // Reset link styles and format the selected sort option.
    $('a.sel').attr('href', '#').removeClass('sel');
    $('a.by' + by).removeAttr('href').addClass('sel');
  }

  /**
   * Create a comp function. If the user has preferences stored in
   * the sortBy cookie, use those, otherwise use the default.
   */
  function initComparator() {
    by = 'rating'; // Default to sort by rating.
    // If the sortBy cookie is set, use that instead.
    if (document.cookie.length > 0) {
      var start = document.cookie.indexOf('sortBy=');
      if (start != -1) {
        start = start + 7;
        var end = document.cookie.indexOf(";", start);
        if (end == -1) {
          end = document.cookie.length;
          by = unescape(document.cookie.substring(start, end));
        }
      }
    }
    setComparator();
  }

  /**
   * Show a comment div.
   */
  function show(id) {
    $('#ao' + id).hide();
    $('#ah' + id).show();
    var context = $.extend({id: id}, opts);
    var popup = $(renderTemplate(popupTemplate, context)).hide();
    popup.find('textarea[name="proposal"]').hide();
    popup.find('a.by' + by).addClass('sel');
    var form = popup.find('#cf' + id);
    form.submit(function(event) {
      event.preventDefault();
      addComment(form);
    });
    $('#s' + id).after(popup);
    popup.slideDown('fast', function() {
      getComments(id);
    });
  }

  /**
   * Hide a comment div.
   */
  function hide(id) {
    $('#ah' + id).hide();
    $('#ao' + id).show();
    var div = $('#sc' + id);
    div.slideUp('fast', function() {
      div.remove();
    });
  }

  /**
   * Perform an ajax request to get comments for a node
   * and insert the comments into the comments tree.
   */
  function getComments(id) {
    $.ajax({
     type: 'GET',
     url: opts.getCommentsURL,
     data: {node: id},
     success: function(data, textStatus, request) {
       var ul = $('#cl' + id);
       var speed = 100;
       $('#cf' + id)
         .find('textarea[name="proposal"]')
         .data('source', data.source);

       if (data.comments.length === 0) {
         ul.html('<li>No comments yet.</li>');
         ul.data('empty', true);
       } else {
         // If there are comments, sort them and put them in the list.
         var comments = sortComments(data.comments);
         speed = data.comments.length * 100;
         appendComments(comments, ul);
         ul.data('empty', false);
       }
       $('#cn' + id).slideUp(speed + 200);
       ul.slideDown(speed);
     },
     error: function(request, textStatus, error) {
       showError('Oops, there was a problem retrieving the comments.');
     },
     dataType: 'json'
    });
  }

  /**
   * Add a comment via ajax and insert the comment into the comment tree.
   */
  function addComment(form) {
    var node_id = form.find('input[name="node"]').val();
    var parent_id = form.find('input[name="parent"]').val();
    var text = form.find('textarea[name="comment"]').val();
    var proposal = form.find('textarea[name="proposal"]').val();

    if (text == '') {
      showError('Please enter a comment.');
      return;
    }

    // Disable the form that is being submitted.
    form.find('textarea,input').attr('disabled', 'disabled');

    // Send the comment to the server.
    $.ajax({
      type: "POST",
      url: opts.addCommentURL,
      dataType: 'json',
      data: {
        node: node_id,
        parent: parent_id,
        text: text,
        proposal: proposal
      },
      success: function(data, textStatus, error) {
        // Reset the form.
        if (node_id) {
          hideProposeChange(node_id);
        }
        form.find('textarea')
          .val('')
          .add(form.find('input'))
          .removeAttr('disabled');
	var ul = $('#cl' + (node_id || parent_id));
        if (ul.data('empty')) {
          $(ul).empty();
          ul.data('empty', false);
        }
        insertComment(data.comment);
        var ao = $('#ao' + node_id);
        ao.find('img').attr({'src': opts.commentBrightImage});
        if (node_id) {
          // if this was a "root" comment, remove the commenting box
          // (the user can get it back by reopening the comment popup)
          $('#ca' + node_id).slideUp();
        }
      },
      error: function(request, textStatus, error) {
        form.find('textarea,input').removeAttr('disabled');
        showError('Oops, there was a problem adding the comment.');
      }
    });
  }

  /**
   * Recursively append comments to the main comment list and children
   * lists, creating the comment tree.
   */
  function appendComments(comments, ul) {
    $.each(comments, function() {
      var div = createCommentDiv(this);
      ul.append($(document.createElement('li')).html(div));
      appendComments(this.children, div.find('ul.comment-children'));
      // To avoid stagnating data, don't store the comments children in data.
      this.children = null;
      div.data('comment', this);
    });
  }

  /**
   * After adding a new comment, it must be inserted in the correct
   * location in the comment tree.
   */
  function insertComment(comment) {
    var div = createCommentDiv(comment);

    // To avoid stagnating data, don't store the comments children in data.
    comment.children = null;
    div.data('comment', comment);

    var ul = $('#cl' + (comment.node || comment.parent));
    var siblings = getChildren(ul);

    var li = $(document.createElement('li'));
    li.hide();

    // Determine where in the parents children list to insert this comment.
    for(i=0; i < siblings.length; i++) {
      if (comp(comment, siblings[i]) <= 0) {
        $('#cd' + siblings[i].id)
          .parent()
          .before(li.html(div));
        li.slideDown('fast');
        return;
      }
    }

    // If we get here, this comment rates lower than all the others,
    // or it is the only comment in the list.
    ul.append(li.html(div));
    li.slideDown('fast');
  }

  function acceptComment(id) {
    $.ajax({
      type: 'POST',
      url: opts.acceptCommentURL,
      data: {id: id},
      success: function(data, textStatus, request) {
        $('#cm' + id).fadeOut('fast');
        $('#cd' + id).removeClass('moderate');
      },
      error: function(request, textStatus, error) {
        showError('Oops, there was a problem accepting the comment.');
      }
    });
  }

  function deleteComment(id) {
    $.ajax({
      type: 'POST',
      url: opts.deleteCommentURL,
      data: {id: id},
      success: function(data, textStatus, request) {
        var div = $('#cd' + id);
        if (data == 'delete') {
          // Moderator mode: remove the comment and all children immediately
          div.slideUp('fast', function() {
            div.remove();
          });
          return;
        }
        // User mode: only mark the comment as deleted
        div
          .find('span.user-id:first')
          .text('[deleted]').end()
          .find('div.comment-text:first')
          .text('[deleted]').end()
          .find('#cm' + id + ', #dc' + id + ', #ac' + id + ', #rc' + id +
                ', #sp' + id + ', #hp' + id + ', #cr' + id + ', #rl' + id)
          .remove();
        var comment = div.data('comment');
        comment.username = '[deleted]';
        comment.text = '[deleted]';
        div.data('comment', comment);
      },
      error: function(request, textStatus, error) {
        showError('Oops, there was a problem deleting the comment.');
      }
    });
  }

  function showProposal(id) {
    $('#sp' + id).hide();
    $('#hp' + id).show();
    $('#pr' + id).slideDown('fast');
  }

  function hideProposal(id) {
    $('#hp' + id).hide();
    $('#sp' + id).show();
    $('#pr' + id).slideUp('fast');
  }

  function showProposeChange(id) {
    $('#pc' + id).hide();
    $('#hc' + id).show();
    var textarea = $('#pt' + id);
    textarea.val(textarea.data('source'));
    $.fn.autogrow.resize(textarea[0]);
    textarea.slideDown('fast');
  }

  function hideProposeChange(id) {
    $('#hc' + id).hide();
    $('#pc' + id).show();
    var textarea = $('#pt' + id);
    textarea.val('').removeAttr('disabled');
    textarea.slideUp('fast');
  }

  function toggleCommentMarkupBox(id) {
    $('#mb' + id).toggle();
  }

  /** Handle when the user clicks on a sort by link. */
  function handleReSort(link) {
    var classes = link.attr('class').split(/\s+/);
    for (var i=0; i<classes.length; i++) {
      if (classes[i] != 'sort-option') {
	by = classes[i].substring(2);
      }
    }
    setComparator();
    // Save/update the sortBy cookie.
    var expiration = new Date();
    expiration.setDate(expiration.getDate() + 365);
    document.cookie= 'sortBy=' + escape(by) +
                     ';expires=' + expiration.toUTCString();
    $('ul.comment-ul').each(function(index, ul) {
      var comments = getChildren($(ul), true);
      comments = sortComments(comments);
      appendComments(comments, $(ul).empty());
    });
  }

  /**
   * Function to process a vote when a user clicks an arrow.
   */
  function handleVote(link) {
    if (!opts.voting) {
      showError("You'll need to login to vote.");
      return;
    }

    var id = link.attr('id');
    if (!id) {
      // Didn't click on one of the voting arrows.
      return;
    }
    // If it is an unvote, the new vote value is 0,
    // Otherwise it's 1 for an upvote, or -1 for a downvote.
    var value = 0;
    if (id.charAt(1) != 'u') {
      value = id.charAt(0) == 'u' ? 1 : -1;
    }
    // The data to be sent to the server.
    var d = {
      comment_id: id.substring(2),
      value: value
    };

    // Swap the vote and unvote links.
    link.hide();
    $('#' + id.charAt(0) + (id.charAt(1) == 'u' ? 'v' : 'u') + d.comment_id)
      .show();

    // The div the comment is displayed in.
    var div = $('div#cd' + d.comment_id);
    var data = div.data('comment');

    // If this is not an unvote, and the other vote arrow has
    // already been pressed, unpress it.
    if ((d.value !== 0) && (data.vote === d.value * -1)) {
      $('#' + (d.value == 1 ? 'd' : 'u') + 'u' + d.comment_id).hide();
      $('#' + (d.value == 1 ? 'd' : 'u') + 'v' + d.comment_id).show();
    }

    // Update the comments rating in the local data.
    data.rating += (data.vote === 0) ? d.value : (d.value - data.vote);
    data.vote = d.value;
    div.data('comment', data);

    // Change the rating text.
    div.find('.rating:first')
      .text(data.rating + ' point' + (data.rating == 1 ? '' : 's'));

    // Send the vote information to the server.
    $.ajax({
      type: "POST",
      url: opts.processVoteURL,
      data: d,
      error: function(request, textStatus, error) {
        showError('Oops, there was a problem casting that vote.');
      }
    });
  }

  /**
   * Open a reply form used to reply to an existing comment.
   */
  function openReply(id) {
    // Swap out the reply link for the hide link
    $('#rl' + id).hide();
    $('#cr' + id).show();

    // Add the reply li to the children ul.
    var div = $(renderTemplate(replyTemplate, {id: id})).hide();
    $('#cl' + id)
      .prepend(div)
      // Setup the submit handler for the reply form.
      .find('#rf' + id)
      .submit(function(event) {
        event.preventDefault();
        addComment($('#rf' + id));
        closeReply(id);
      })
      .find('input[type=button]')
      .click(function() {
        closeReply(id);
      });
    div.slideDown('fast', function() {
      $('#rf' + id).find('textarea').focus();
    });
  }

  /**
   * Close the reply form opened with openReply.
   */
  function closeReply(id) {
    // Remove the reply div from the DOM.
    $('#rd' + id).slideUp('fast', function() {
      $(this).remove();
    });

    // Swap out the hide link for the reply link
    $('#cr' + id).hide();
    $('#rl' + id).show();
  }

  /**
   * Recursively sort a tree of comments using the comp comparator.
   */
  function sortComments(comments) {
    comments.sort(comp);
    $.each(comments, function() {
      this.children = sortComments(this.children);
    });
    return comments;
  }

  /**
   * Get the children comments from a ul. If recursive is true,
   * recursively include childrens' children.
   */
  function getChildren(ul, recursive) {
    var children = [];
    ul.children().children("[id^='cd']")
      .each(function() {
        var comment = $(this).data('comment');
        if (recursive)
          comment.children = getChildren($(this).find('#cl' + comment.id), true);
        children.push(comment);
      });
    return children;
  }

  /** Create a div to display a comment in. */
  function createCommentDiv(comment) {
    if (!comment.displayed && !opts.moderator) {
      return $('<div class="moderate">Thank you!  Your comment will show up '
               + 'once it is has been approved by a moderator.</div>');
    }
    // Prettify the comment rating.
    comment.pretty_rating = comment.rating + ' point' +
      (comment.rating == 1 ? '' : 's');
    // Make a class (for displaying not yet moderated comments differently)
    comment.css_class = comment.displayed ? '' : ' moderate';
    // Create a div for this comment.
    var context = $.extend({}, opts, comment);
    var div = $(renderTemplate(commentTemplate, context));

    // If the user has voted on this comment, highlight the correct arrow.
    if (comment.vote) {
      var direction = (comment.vote == 1) ? 'u' : 'd';
      div.find('#' + direction + 'v' + comment.id).hide();
      div.find('#' + direction + 'u' + comment.id).show();
    }

    if (opts.moderator || comment.text != '[deleted]') {
      div.find('a.reply').show();
      if (comment.proposal_diff)
        div.find('#sp' + comment.id).show();
      if (opts.moderator && !comment.displayed)
        div.find('#cm' + comment.id).show();
      if (opts.moderator || (opts.username == comment.username))
        div.find('#dc' + comment.id).show();
    }
    return div;
  }

  /**
   * A simple template renderer. Placeholders such as <%id%> are replaced
   * by context['id'] with items being escaped. Placeholders such as <#id#>
   * are not escaped.
   */
  function renderTemplate(template, context) {
    var esc = $(document.createElement('div'));

    function handle(ph, escape) {
      var cur = context;
      $.each(ph.split('.'), function() {
        cur = cur[this];
      });
      return escape ? esc.text(cur || "").html() : cur;
    }

    return template.replace(/<([%#])([\w\.]*)\1>/g, function() {
      return handle(arguments[2], arguments[1] == '%' ? true : false);
    });
  }

  /** Flash an error message briefly. */
  function showError(message) {
    $(document.createElement('div')).attr({'class': 'popup-error'})
      .append($(document.createElement('div'))
               .attr({'class': 'error-message'}).text(message))
      .appendTo('body')
      .fadeIn("slow")
      .delay(2000)
      .fadeOut("slow");
  }

  /** Add a link the user uses to open the comments popup. */
  $.fn.comment = function() {
    return this.each(function() {
      var id = $(this).attr('id').substring(1);
      var count = COMMENT_METADATA[id];
      var title = count + ' comment' + (count == 1 ? '' : 's');
      var image = count > 0 ? opts.commentBrightImage : opts.commentImage;
      var addcls = count == 0 ? ' nocomment' : '';
      $(this)
        .append(
          $(document.createElement('a')).attr({
            href: '#',
            'class': 'sphinx-comment-open' + addcls,
            id: 'ao' + id
          })
            .append($(document.createElement('img')).attr({
              src: image,
              alt: 'comment',
              title: title
            }))
            .click(function(event) {
              event.preventDefault();
              show($(this).attr('id').substring(2));
            })
        )
        .append(
          $(document.createElement('a')).attr({
            href: '#',
            'class': 'sphinx-comment-close hidden',
            id: 'ah' + id
          })
            .append($(document.createElement('img')).attr({
              src: opts.closeCommentImage,
              alt: 'close',
              title: 'close'
            }))
            .click(function(event) {
              event.preventDefault();
              hide($(this).attr('id').substring(2));
            })
        );
    });
  };

  var opts = {
    processVoteURL: '/_process_vote',
    addCommentURL: '/_add_comment',
    getCommentsURL: '/_get_comments',
    acceptCommentURL: '/_accept_comment',
    deleteCommentURL: '/_delete_comment',
    commentImage: '/static/_static/comment.png',
    closeCommentImage: '/static/_static/comment-close.png',
    loadingImage: '/static/_static/ajax-loader.gif',
    commentBrightImage: '/static/_static/comment-bright.png',
    upArrow: '/static/_static/up.png',
    downArrow: '/static/_static/down.png',
    upArrowPressed: '/static/_static/up-pressed.png',
    downArrowPressed: '/static/_static/down-pressed.png',
    voting: false,
    moderator: false
  };

  if (typeof COMMENT_OPTIONS != "undefined") {
    opts = jQuery.extend(opts, COMMENT_OPTIONS);
  }

  var popupTemplate = '\
    <div class="sphinx-comments" id="sc<%id%>">\
      <p class="sort-options">\
        Sort by:\
        <a href="#" class="sort-option byrating">best rated</a>\
        <a href="#" class="sort-option byascage">newest</a>\
        <a href="#" class="sort-option byage">oldest</a>\
      </p>\
      <div class="comment-header">Comments</div>\
      <div class="comment-loading" id="cn<%id%>">\
        loading comments... <img src="<%loadingImage%>" alt="" /></div>\
      <ul id="cl<%id%>" class="comment-ul"></ul>\
      <div id="ca<%id%>">\
      <p class="add-a-comment">Add a comment\
        (<a href="#" class="comment-markup" id="ab<%id%>">markup</a>):</p>\
      <div class="comment-markup-box" id="mb<%id%>">\
        reStructured text markup: <i>*emph*</i>, <b>**strong**</b>, \
        <code>``code``</code>, \
        code blocks: <code>::</code> and an indented block after blank line</div>\
      <form method="post" id="cf<%id%>" class="comment-form" action="">\
        <textarea name="comment" cols="80"></textarea>\
        <p class="propose-button">\
          <a href="#" id="pc<%id%>" class="show-propose-change">\
            Propose a change &#9657;\
          </a>\
          <a href="#" id="hc<%id%>" class="hide-propose-change">\
            Propose a change &#9663;\
          </a>\
        </p>\
        <textarea name="proposal" id="pt<%id%>" cols="80"\
                  spellcheck="false"></textarea>\
        <input type="submit" value="Add comment" />\
        <input type="hidden" name="node" value="<%id%>" />\
        <input type="hidden" name="parent" value="" />\
      </form>\
      </div>\
    </div>';

  var commentTemplate = '\
    <div id="cd<%id%>" class="sphinx-comment<%css_class%>">\
      <div class="vote">\
        <div class="arrow">\
          <a href="#" id="uv<%id%>" class="vote" title="vote up">\
            <img src="<%upArrow%>" />\
          </a>\
          <a href="#" id="uu<%id%>" class="un vote" title="vote up">\
            <img src="<%upArrowPressed%>" />\
          </a>\
        </div>\
        <div class="arrow">\
          <a href="#" id="dv<%id%>" class="vote" title="vote down">\
            <img src="<%downArrow%>" id="da<%id%>" />\
          </a>\
          <a href="#" id="du<%id%>" class="un vote" title="vote down">\
            <img src="<%downArrowPressed%>" />\
          </a>\
        </div>\
      </div>\
      <div class="comment-content">\
        <p class="tagline comment">\
          <span class="user-id"><%username%></span>\
          <span class="rating"><%pretty_rating%></span>\
          <span class="delta"><%time.delta%></span>\
        </p>\
        <div class="comment-text comment"><#text#></div>\
        <p class="comment-opts comment">\
          <a href="#" class="reply hidden" id="rl<%id%>">reply &#9657;</a>\
          <a href="#" class="close-reply" id="cr<%id%>">reply &#9663;</a>\
          <a href="#" id="sp<%id%>" class="show-proposal">proposal &#9657;</a>\
          <a href="#" id="hp<%id%>" class="hide-proposal">proposal &#9663;</a>\
          <a href="#" id="dc<%id%>" class="delete-comment hidden">delete</a>\
          <span id="cm<%id%>" class="moderation hidden">\
            <a href="#" id="ac<%id%>" class="accept-comment">accept</a>\
          </span>\
        </p>\
        <pre class="proposal" id="pr<%id%>">\
<#proposal_diff#>\
        </pre>\
          <ul class="comment-children" id="cl<%id%>"></ul>\
        </div>\
        <div class="clearleft"></div>\
      </div>\
    </div>';

  var replyTemplate = '\
    <li>\
      <div class="reply-div" id="rd<%id%>">\
        <form id="rf<%id%>">\
          <textarea name="comment" cols="80"></textarea>\
          <input type="submit" value="Add reply" />\
          <input type="button" value="Cancel" />\
          <input type="hidden" name="parent" value="<%id%>" />\
          <input type="hidden" name="node" value="" />\
        </form>\
      </div>\
    </li>';

  $(document).ready(function() {
    init();
  });
})(jQuery);

$(document).ready(function() {
  // add comment anchors for all paragraphs that are commentable
  $('.sphinx-has-comment').comment();

  // highlight search words in search results
  $("div.context").each(function() {
    var params = $.getQueryParameters();
    var terms = (params.q) ? params.q[0].split(/\s+/) : [];
    var result = $(this);
    $.each(terms, function() {
      result.highlightText(this.toLowerCase(), 'highlighted');
    });
  });

  // directly open comment window if requested
  var anchor = document.location.hash;
  if (anchor.substring(0, 9) == '#comment-') {
    $('#ao' + anchor.substring(9)).click();
    document.location.hash = '#s' + anchor.substring(9);
  }
});
```
