## Supplementary material for "ΔSCOPE: A new method to quantify 3D biological structures and identify differences in zebrafish forebrain development": An archived copy of the DeltaSCOPE code repository.: API.html

  


API — cranium 0.1.0 documentation


cranium

0.1.0

- Data Preparation
- Data Processing
- Checklist
- Parameter Reference
- API

cranium

- Docs »
- API
- View page source

---

### API¶

*class* `cranium.``brain`[source]¶
:   Object to manage biological data and associated functions.

    `read_data`(*filepath*)[source]¶
    :   Reads 3D data from file and selects appropriate channel based on the assumption that the channel with the most zeros has zero as the value for no signal

        |  |  |
        | --- | --- |
        | Parameters: | **filepath** (*str*) – Filepath to hdf5 probability file |
        | Returns: | Creates the variable `brain.raw_data` |

        `raw_data`¶
        :   Array of shape [z,y,x] containing raw probability data

    `create_dataframe`(*data*, *scale*)[source]¶
    :   Creates a pandas dataframe containing the x,y,z and signal/probability value for each point in the `brain.raw_data` array

        |  |  |
        | --- | --- |
        | Parameters: | - **data** (*array*) – Raw probability data in 3D array - **scale** (*array*) – Array of length three containing the micron values for [x,y,z] |
        | Returns: | Pandas DataFrame with xyz and probability value for each point |

    `plot_projections`(*df*, *subset*)[source]¶
    :   Plots the x, y, and z projections of the input dataframe in a matplotlib plot

        |  |  |
        | --- | --- |
        | Parameters: | - **df** (*pd.DataFrame*) – Dataframe with columns: ‘x’,’y’,’z’ - **subset** (*float*) – Value between 0 and 1 indicating what percentage of the df to subsample |
        | Returns: | Matplotlib figure with three labeled scatterplots |

    `preprocess_data`(*threshold*, *scale*, *microns*)[source]¶
    :   Thresholds and scales data prior to PCA

        Creates `brain.threshold`, `brain.df_thresh`, and `brain.df_scl`

        |  |  |
        | --- | --- |
        | Parameters: | - **threshold** (*float*) – Value between 0 and 1 to use as a cutoff for minimum pixel value - **scale** (*array*) – Array with three values representing the constant by which to multiply x,y,z respectively - **microns** (*array*) – Array with three values representing the x,y,z micron dimensions of the voxel |

        `threshold`¶
        :   Value used to threshold the data prior to calculating the model

        `df_thresh`¶
        :   Dataframe containing only points with values above the specified threshold

        `df_scl`¶
        :   Dataframe containing data from `brain.df_thresh` after a scaling value has been applied

    `process_alignment_data`(*data*, *threshold*, *radius*, *microns*)[source]¶
    :   Applies a median filter twice to the data which is used for alignment

        Ensures than any noise in the structural data does not interfere with alignment

        |  |  |
        | --- | --- |
        | Parameters: | - **data** (*array*) – Raw data imported by the function `brain.read_data()` - **threshold** (*float*) – Value between 0 and 1 to use as a cutoff for minimum pixel value - **radius** (*int*) – Integer that determines the radius of the circle used for the median filter - **microns** (*array*) – Array with three values representing the x,y,z micron dimensions of the voxel |
        | Returns: | Dataframe containing data processed with the median filter and threshold |

    `calculate_pca_median`(*data*, *threshold*, *radius*, *microns*)[source]¶
    :   Calculate PCA transformation matrix, `brain.pcamed`, based on data (`brain.pcamed`) after applying median filter and threshold

        |  |  |
        | --- | --- |
        | Parameters: | - **data** (*array*) – 3D array containing raw probability data - **threshold** (*float*) – Value between 0 and 1 indicating the lower cutoff for positive signal - **radius** (*int*) – Radius of neighborhood that should be considered for the median filter - **microns** (*array*) – Array with three values representing the x,y,z micron dimensions of the voxel |

        `median`¶
        :   Pandas dataframe containing data that has been processed with a median filter twice and thresholded

        `pcamed`¶
        :   PCA object managing the transformation matrix and any resulting transformations

    `calculate_pca_median_2d`(*data*, *threshold*, *radius*, *microns*)[source]¶
    :   Calculate PCA transformation matrix for 2 dimensions of data, `brain.pcamed`, based on data after applying median filter and threshold

        Warning

        fit\_dim is not used to determine which dimensions to fit. Defaults to x and z

        |  |  |
        | --- | --- |
        | Parameters: | - **data** (*array*) – 3D array containing raw probability data - **threshold** (*float*) – Value between 0 and 1 indicating the lower cutoff for positive signal - **radius** (*int*) – Radius of neighborhood that should be considered for the median filter - **microns** (*array*) – Array with three values representing the x,y,z micron dimensions of the voxel |

    `pca_transform_2d`(*df*, *pca*, *comp\_order*, *fit\_dim*, *deg=2*, *mm=None*, *vertex=None*, *flip=None*)[source]¶
    :   Transforms df in 2D based on the PCA object, pca, whose transformation matrix has already been calculated

        Calling `brain.align_data()` creates `brain.df_align`

        Warning

        fit\_dim is not used to determine which dimensions to fit. Defaults to x and z

        |  |  |
        | --- | --- |
        | Parameters: | - **df** (*pd.DataFrame*) – Dataframe containing thresholded xyz data - **pca** (*pca\_object*) – A pca object containing a transformation object, e.g. `brain.pcamed` - **comp\_order** (*array*) – Array specifies the assignment of components to x,y,z. Form [x component index, y component index, z component index], e.g. [0,2,1] - **fit\_dim** (*array*) – Array of length two containing two strings describing the first and second axis for fitting the model, e.g. [‘x’,’z’] - **deg** (*int*) – (or None) Degree of the function that should be fit to the model. deg=2 by default - **mm** – (`math_model` or None) Math model for primary channel - **vertex** (*array*) – (or None) Array of type [vx,vy,vz] (`brain.vertex`) indicating the translation values - **flip** (*Bool*) – (or None) Boolean value to determine if the data should be rotated by 180 degrees |

    `pca_transform_3d`(*df*, *pca*, *comp\_order*, *fit\_dim*, *deg=2*, *mm=None*, *vertex=None*, *flip=None*)[source]¶
    :   Transforms df in 3D based on the PCA object, pca, whose transformation matrix has already been calculated

        |  |  |
        | --- | --- |
        | Parameters: | - **df** (*pd.DataFrame*) – Dataframe containing thresholded xyz data - **pca** (*pca\_object*) – A pca object containing a transformation object, e.g. `brain.pcamed` - **comp\_order** (*array*) – Array specifies the assignment of components to x,y,z. Form [x component index, y component index, z component index], e.g. [0,2,1] - **fit\_dim** (*array*) – Array of length two containing two strings describing the first and second axis for fitting the model, e.g. [‘x’,’z’] - **deg** (*int*) – (or None) Degree of the function that should be fit to the model. deg=2 by default - **mm** – (`math_model` or None) Math model for primary channel - **vertex** (*array*) – (or None) Array of type [vx,vy,vz] (`brain.vertex`) indicating the translation values - **flip** (*Bool*) – (or None) Boolean value to determine if the data should be rotated by 180 degrees |

    `align_data`(*df\_fit*, *fit\_dim*, *deg=2*, *mm=None*, *vertex=None*, *flip=None*)[source]¶
    :   Apply PCA transformation matrix and align data so that the vertex is at the origin

        Creates `brain.df_align` and `brain.mm`

        |  |  |
        | --- | --- |
        | Parameters: | - **df** (*pd.DataFrame*) – dataframe containing thresholded xyz data - **comp\_order** (*array*) – Array specifies the assignment of components to x,y,z. Form [x component index, y component index, z component index], e.g. [0,2,1] - **fit\_dim** (*array*) – Array of length two containing two strings describing the first and second axis for fitting the model, e.g. [‘x’,’z’] - **deg** (*int*) – (or None) Degree of the function that should be fit to the model. deg=2 by default - **mm** – (`math_model` or None) Math model for primary channel - **vertex** (*array*) – (or None) Array of type [vx,vy,vz] (`brain.vertex`) indicating the translation values - **flip** (*Bool*) – (or None) Boolean value to determine if the data should be rotated by 180 degrees |

        `df_align`¶
        :   Dataframe containing point data aligned using PCA

        `mm`¶
        :   Math model object fit to data in brain object

    `flip_data`(*df*)[source]¶
    :   Rotate data by 180 degrees

        |  |  |
        | --- | --- |
        | Parameters: | **df** (*dataframe*) – Pandas dataframe containing x,y,z data |
        | Returns: | Rotated dataframe |

    `fit_model`(*df*, *deg*, *fit\_dim*)[source]¶
    :   Fit model to dataframe

        |  |  |
        | --- | --- |
        | Parameters: | - **df** (*pd.DataFrame*) – Dataframe containing at least x,y,z - **deg** (*int*) – Degree of the function that should be fit to the model - **fit\_dim** (*array*) – Array of length two containing two strings describing the first and second axis for fitting the model, e.g. [‘x’,’z’] |
        | Returns: | math model |
        | Return type: | `math_model` |

    `find_distance`(*t*, *point*)[source]¶
    :   Find euclidean distance between math model(t) and data point in the xy plane

        |  |  |
        | --- | --- |
        | Parameters: | - **t** (*float*) – float value defining point on the line - **point** (*array*) – array [x,y] defining data point |
        | Returns: | distance between the two points |
        | Return type: | float |

    `find_min_distance`(*row*)[source]¶
    :   Find the point on the curve that produces the minimum distance between the point and the data point using scipy.optimize.minimize(`brain.find_distance()`)

        |  |  |
        | --- | --- |
        | Parameters: | **row** (*pd.Series*) – row from dataframe in the form of a pandas Series |
        | Returns: | point in the curve (xc, yc, zc) and r |
        | Return type: | floats |

    `integrand`(*x*)[source]¶
    :   Function to integrate to calculate arclength

        |  |  |
        | --- | --- |
        | Parameters: | **x** (*float*) – integer value for x |
        | Returns: | arclength value for integrating |
        | Return type: | float |

    `find_arclength`(*xc*)[source]¶
    :   Calculate arclength by integrating the derivative of the math model in xy plane

        System Message: WARNING/2 (\int\_{vertex}^{point} \sqrt{1 + (2ax + b)^2})

        latex exited with error
        [stderr]
        Unfortunately, the package anyfontsize could not be installed.Please check the log file:
        C:/Users/zfishlab/AppData/Local/MiKTeX/2.9/miktex/log/latex.log
        [stdout]
        This is pdfTeX, Version 3.14159265-2.6-1.40.17 (MiKTeX 2.9 64-bit)
        entering extended mode
        (math.tex
        LaTeX2e <2016/03/31> patch level 1
        Babel <3.9r> and hyphenation patterns for 75 language(s) loaded.
        ("C:\Users\zfishlab\AppData\Local\Programs\MiKTeX 2.9\tex\latex\base\article.cl
        s"
        Document Class: article 2014/09/29 v1.4h Standard LaTeX document class
        ("C:\Users\zfishlab\AppData\Local\Programs\MiKTeX 2.9\tex\latex\base\size12.clo
        "))
        ("C:\Users\zfishlab\AppData\Local\Programs\MiKTeX 2.9\tex\latex\base\inputenc.s
        ty"
        ("C:\Users\zfishlab\AppData\Local\Programs\MiKTeX 2.9\tex\latex\ucs\utf8x.def")
        ) ("C:\Users\zfishlab\AppData\Local\Programs\MiKTeX 2.9\tex\latex\ucs\ucs.sty"
        ("C:\Users\zfishlab\AppData\Local\Programs\MiKTeX 2.9\tex\latex\ucs\uni-global.
        def"))
        ("C:\Users\zfishlab\AppData\Local\Programs\MiKTeX 2.9\tex\latex\amsmath\amsmath
        .sty"
        For additional information on amsmath, use the `?' option.
        ("C:\Users\zfishlab\AppData\Local\Programs\MiKTeX 2.9\tex\latex\amsmath\amstext
        .sty"
        ("C:\Users\zfishlab\AppData\Local\Programs\MiKTeX 2.9\tex\latex\amsmath\amsgen.
        sty"))
        ("C:\Users\zfishlab\AppData\Local\Programs\MiKTeX 2.9\tex\latex\amsmath\amsbsy.
        sty")
        ("C:\Users\zfishlab\AppData\Local\Programs\MiKTeX 2.9\tex\latex\amsmath\amsopn.
        sty"))
        ("C:\Users\zfishlab\AppData\Local\Programs\MiKTeX 2.9\tex\latex\amscls\amsthm.s
        ty")
        ("C:\Users\zfishlab\AppData\Local\Programs\MiKTeX 2.9\tex\latex\amsfonts\amssym
        b.sty"
        ("C:\Users\zfishlab\AppData\Local\Programs\MiKTeX 2.9\tex\latex\amsfonts\amsfon
        ts.sty"))
        ======================================================================
        ======================================================================
        ! LaTeX Error: File `anyfontsize.sty' not found.
        Type X to quit or <RETURN> to proceed,
        or enter new name. (Default extension: sty)
        Enter file name:
        ! Emergency stop.
        <read \*>
        l.9 \usepackage
        {bm}
        No pages of output.
        Transcript written on math.log.

        |  |  |
        | --- | --- |
        | Parameters: | **row** (*float*) – Postion in the x axis along the curve |
        | Returns: | Length of the arc along the curve between the row and the vertex |
        | Return type: | float |

    `find_theta`(*row*, *zc*, *yc*)[source]¶
    :   Calculate theta for a row containing data point in relationship to the xz plane

        |  |  |
        | --- | --- |
        | Parameters: | - **row** (*pd.Series*) – row from dataframe in the form of a pandas Series - **yc** (*float*) – Y position of the closest point in the curve to the data point - **zc** (*float*) – Z position of the closest point in the curve to the data point |
        | Returns: | theta, angle between point and the model plane |
        | Return type: | float |

    `find_r`(*row*, *zc*, *yc*)[source]¶
    :   Calculate r using the Pythagorean theorem

        |  |  |
        | --- | --- |
        | Parameters: | - **row** (*pd.Series*) – row from dataframe in the form of a pandas Series - **yc** (*float*) – Y position of the closest point in the curve to the data point - **zc** (*float*) – Z position of the closest point in the curve to the data point |
        | Returns: | r, distance between the point and the model |
        | Return type: | float |

    `calc_coord`(*row*)[source]¶
    :   Calculate alpah, r, theta for a particular row

        |  |  |
        | --- | --- |
        | Parameters: | **row** (*pd.Series*) – row from dataframe in the form of a pandas Series |
        | Returns: | pd.Series populated with coordinate of closest point on the math model, r, theta, and ac (arclength) |

    `transform_coordinates`()[source]¶
    :   Transform coordinate system so that each point is defined relative to math model by (alpha,theta,r) (only applied to `brain.df_align`)

        |  |  |
        | --- | --- |
        | Returns: | appends columns r, xc, yc, zc, ac, theta to `brain.df_align` |

    `subset_data`(*df*, *sample\_frac=0.5*)[source]¶
    :   Takes a random sample of the data based on the value between 0 and 1 defined for sample\_frac

        Creates the variable `brain.subset`

        |  |  |
        | --- | --- |
        | Parameters: | - **pd.DataFrame** – Dataframe which will be sampled - **sample\_frac** (*float*) – (or None) Value between 0 and 1 specifying proportion of the dataset that should be randomly sampled for plotting |

        `subset`¶
        :   Random sample of the input dataframe

    `add_thresh_df`(*df*)[source]¶
    :   Adds dataframe of thresholded and transformed data to `brain.df_thresh`

        |  |  |
        | --- | --- |
        | Parameters: | **df** (*pd.DataFrame*) – dataframe of thesholded and transformed data |
        | Returns: | `brain.df_thresh` |

    `add_aligned_df`(*df*)[source]¶
    :   Adds dataframe of aligned data

        Warning

        Calculates model, but assumes that the dimensions of the fit are x and z

        |  |  |
        | --- | --- |
        | Parameters: | **df** (*pd.DataFrame*) – Dataframe of aligned data |
        | Returns: | `brain.df_align` |

*class* `cranium.``embryo`(*name*, *number*, *outdir*)[source]¶
:   Class to managed multiple brain objects in a multichannel sample

    |  |  |
    | --- | --- |
    | Parameters: | - **name** (*str*) – Name of this sample set - **number** (*str*) – Sample number corresponding to this embryo - **outdir** (*str*) – Path to directory for output files |

    `chnls`¶
    :   Dictionary containing the `brain` object for each channel

    `outdir`¶
    :   Path to directory for output files

    `name`¶
    :   Name of this sample set

    `number`¶
    :   Sample number corresponding to this embryo

    `add_channel`(*filepath*, *key*)[source]¶
    :   Add channel to `embryo.chnls` dictionary

        |  |  |
        | --- | --- |
        | Parameters: | - **filepath** (*str*) – Complete filepath to image - **key** (*str*) – Name of the channel |

    `process_channels`(*mthresh*, *gthresh*, *radius*, *scale*, *microns*, *deg*, *primary\_key*, *comp\_order*, *fit\_dim*)[source]¶
    :   Process all channels through the production of the `brain.df_align` dataframe

        |  |  |
        | --- | --- |
        | Parameters: | - **mthresh** (*float*) – Value between 0 and 1 to use as a cutoff for minimum pixel value for median data - **gthresh** (*float*) – Value between 0 and 1 to use as a cutoff for minimum pixel value for general data - **radius** (*int*) – Size of the neighborhood area to examine with median filter - **scale** (*array*) – Array with three values representing the constant by which to multiply x,y,z respectively - **microns** (*array*) – Array with three values representing the x,y,z micron dimensions of the voxel - **deg** (*int*) – Degree of the function that should be fit to the model - **primary\_key** (*str*) – Key for the primary structural channel which PCA and the model should be fit too - **comp\_order** (*array*) – Array specifies the assignment of components to x,y,z. Form [x component index, y component index, z component index], e.g. [0,2,1] - **fit\_dim** (*array*) – Array of length two containing two strings describing the first and second axis for fitting the model, e.g. [‘x’,’z’] |

    `save_projections`(*subset*)[source]¶
    :   Save projections of both channels into png files in `embryo.outdir` following the naming scheme [`embryo.name`]\_[`embryo.number`]\_[channel name]\_MIP.png

        |  |  |
        | --- | --- |
        | Parameters: | **subset** (*float*) – Value between 0 and 1 to specify the fraction of the data to randomly sample for plotting |

    `save_psi`()[source]¶
    :   Save all channels into psi files following the naming scheme [`embryo.name`]\_[`embryo.number`]\_[channel name].psi

    `add_psi_data`(*filepath*, *key*)[source]¶
    :   Read psi data into a channel dataframe

        |  |  |
        | --- | --- |
        | Parameters: | - **filepath** (*str*) – Complete filepath to data - **key** (*str*) – Descriptive key for channel dataframe in dictionary |

*class* `cranium.``math_model`(*model*)[source]¶
:   Object to contain attributes associated with the math model of a sample

    |  |  |
    | --- | --- |
    | Parameters: | **model** (*array*) – Array of coefficients calculated by np.polyfit |

    `cf`¶
    :   Array of coefficients for the math model

    `p`¶
    :   Poly1d function for the math model to allow calculation and plotting of the model

*class* `cranium.``landmarks`(*percbins=[10, 50, 90], rnull=15*)[source]¶
:   Class to handle calculation of landmarks to describe structural data

    |  |  |
    | --- | --- |
    | Parameters: | - **percbins** (*list*) – (or None) Must be a list of integers between 0 and 100 - **rnull** (*int*) – (or None) When the r value cannot be calculated it will be set to this value |

    `brain.``lm_wt_rf`¶
    :   pd.DataFrame, which wildtype landmarks will be added to

    `brain.``lm_mt_rf`¶
    :   pd.DataFrame, which mutant landmarks will be added to

    `brain.``rnull`¶
    :   Integer specifying the value which null landmark calculations will be set to

    `brain.``percbins`¶
    :   Integer specifying the percentiles which will be used to calculate landmarks

    `calc_bins`(*Ldf*, *ac\_num*, *tstep*)[source]¶
    :   Calculates alpha and theta bins based on ac\_num and tstep

        Creates `landmarks.acbins` and `landmarks.tbins`

        Warning

        tstep does not handle scenarios where 2pi is not evenly divisible by tstep

        |  |  |
        | --- | --- |
        | Parameters: | - **Ldf** (*list*) – List of dataframes that are being used for the analysis typically accessed by dict.values() - **ac\_num** (*int*) – Integer indicating the number of divisions that should be made along alpha - **tstep** (*float*) – The size of each bin used for alpha |

        `acbins`¶
        :   List containing the boundaries of each bin along alpha based on ac\_num

        `tbins`¶
        :   List containing the boundaries of each bin along theta based on tstep

    `calc_perc`(*df*, *snum*, *dtype*, *out*)[source]¶
    :   Calculate landmarks for a dataframe based on the bins and percentiles that have been previously defined

        |  |  |
        | --- | --- |
        | Parameters: | - **df** (*pd.DataFrame*) – Dataframe containing columns x,y,z,alpha,r,theta - **snum** (*str*) – String containing a sample identifier that can be converted to an integer - **dtype** (*str*) – String describing the sample group to which the sample belongs, e.g. control or experimental |
        | Returns: | pd.DataFrame with new landmarks appended |

    `calc_wt_reformat`(*df*, *snum*)[source]¶
    :   Warning

        Deprecated function, but includes code pertaining to calculating point based data

    `calc_mt_landmarks`(*df*, *snum*, *wt*)[source]¶
    :   Warning

        Deprecated function, but attempted to calculate mutant landmarks based on the number of points found in the wildtype standard

`cranium.``reformat_to_cart`(*df*)[source]¶
:   Take a dataframe in which columns contain the bin parameters and convert to a cartesian coordinate system

    |  |  |
    | --- | --- |
    | Parameters: | **df** (*pd.DataFrame*) – Dataframe containing columns with string names that contain the bin parameter |
    | Returns: | pd.DataFrame with each landmark as a row and columns: x,y,z,r,r\_std,t,pts |

`cranium.``subplot_lmk`(*ax*, *p*, *avg*, *sem*, *parr*, *xarr*, *tarr*, *dtype*, *Pn={'alpha': 0.3*, *'cmap': 'Greys\_r'*, *'mtc': 'r'*, *'tarr': None*, *'wtc': 'b'*, *'xarr': None*, *'zfb': 1*, *'zln': 2*, *'zpt': 3}*)[source]¶
:   Plot a ribbon of average and standard error of the mean onto the subplot, ax

    |  |  |
    | --- | --- |
    | Parameters: | - **ax** (*plt.Subplot*) – Matplotlib subplot onto which the data should be plotted - **p** (*list*) – List of two theta values that should be plotted - **avg** (*np.array*) – Array of shape (xvalues,tvalues) containing the average values of the data - **sem** (*np.array*) – Array of shape (xvalues,tvalues) containing the standard error of the mean values of the data - **parr** (*np.array*) – Array of shape (xvalues,tvalues) containing the p values for the data - **dtype** (*str*) – String describing sample type - **Pn** – Dictionary containing the following values: ‘zln’:2,’zpt’:3,’zfb’:1,’wtc’:’b’,’mtc’:’r’,’alpha’:0.3,’cmap’:’Greys\_r’ |
    | Type: | dict or None |

`cranium.``write_header`(*f*)[source]¶
:   Writes header for PSI file with columns Id,x,y,z,ac,r,theta

    |  |  |
    | --- | --- |
    | Parameters: | **f** (*file*) – file object created by ‘open(filename,’w’)` |

`cranium.``write_data`(*filepath*, *df*)[source]¶
:   Writes data in PSI format to file after writing header using `write_header()`. Closes file at the conclusion of writing data.

    |  |  |
    | --- | --- |
    | Parameters: | - **filepath** (*str*) – Complete filepath to output file - **df** (*pd.DataFrame*) – dataframe containing columns x,y,z,ac,r,theta |

`cranium.``read_psi`(*filepath*)[source]¶
:   Reads psi file at the given filepath and returns data in a pandas DataFrame

    |  |  |
    | --- | --- |
    | Parameters: | **filepath** (*str*) – Complete filepath to file |
    | Returns: | pd.Dataframe containing data |

`cranium.``read_psi_to_dict`(*directory*, *dtype*)[source]¶
:   Read psis from directory into dictionary of dfs with filtering based on dtype

    |  |  |
    | --- | --- |
    | Parameters: | - **directory** (*str*) – Directory to get psis from - **dtype** (*str*) – Usually ‘AT’ or ‘ZRF1’ |
    | Returns: | Dictionary of pd.DataFrame |

`cranium.``process_sample`(*num*, *root*, *outdir*, *name*, *chs*, *prefixes*, *threshold*, *scale*, *deg*, *primary\_key*, *comp\_order*, *fit\_dim*, *flip\_dim*)[source]¶
:   Process single sample through `brain` class and saves df to csv

    Warning

    Out of date and will probably fail

    |  |  |
    | --- | --- |
    | Parameters: | - **num** (*str*) – Sample number - **root** (*str*) – Complete path to the root directory for this sample set - **name** (*str*) – Name describing this sample set - **outdir** (*str*) – Complete path to output directory - **chs** (*array*) – Array containing strings specifying the directories for each channel - **prefixes** (*array*) – Array containing strings specifying the file prefix for each channel - **threshold** (*float*) – Value between 0 and 1 to use as a cutoff for minimum pixel value - **scale** (*array*) – Array with three values representing the constant by which to multiply x,y,z respectively - **deg** (*int*) – Degree of the function that should be fit to the model - **primary\_key** (*str*) – Key for the primary structural channel which PCA and the model should be fit too |

`cranium.``calculate_models`(*Ldf*)[source]¶
:   Calculate model for each dataframe in list and add to new dataframe

    |  |  |
    | --- | --- |
    | Parameters: | **Ldf** (*list*) – List of dataframes containing aligned data |
    | Returns: | pd.Dataframe with a,b,c values for parabolic model |

`cranium.``generate_kde`(*data*, *var*, *x*, *absv=False*)[source]¶
:   Generate list of KDEs from either dictionary or list of data

    |  |  |
    | --- | --- |
    | Parameters: | - **data** – pd.DataFrames to convert - **var** (*str*) – Name of column to select from df - **x** (*array*) – Array of datapoints to evaluate KDE on - **absv** (*bool*) – (or None) Set to True to use absolute value of selected data for KDE calculation |
    | Type: | dict or list |
    | Returns: | List of KDE arrays |

`cranium.``calculate_area_error`(*pdf*, *Lkde*, *x*)[source]¶
:   Calculate area between PDF and each kde in Lkde

    |  |  |
    | --- | --- |
    | Parameters: | - **pdf** (*array*) – Array of probability distribution function that is the same shape as kdes in Lkde - **Lkde** (*list*) – List of arrays of Kdes - **x** (*array*) – Array of datapoints used to generate pdf and kdes |
    | Returns: | List of error values for each kde in Lkde |

`cranium.``rescale_variable`(*Ddfs*, *var*, *newvar*)[source]¶
:   Rescale variable from -1 to 1 and save in newvar column on original dataframe

    |  |  |
    | --- | --- |
    | Parameters: | - **Ddfs** (*dict*) – Dictionary of pd.DataFrames - **var** (*str*) – Name of column to select from dfs - **newvar** (*str*) – Name to use for new data in appended column |
    | Returns: | Dictionary of dataframes containing column of rescaled data |

Previous

---

© Copyright 2018, Morgan Schwartz.

Built with Sphinx using a theme provided by Read the Docs.
