## Supplementary material for "ΔSCOPE: A new method to quantify 3D biological structures and identify differences in zebrafish forebrain development": An archived copy of the DeltaSCOPE code repository.: checklist.html

  


Checklist — cranium 0.1.0 documentation


cranium

0.1.0

- Data Preparation
- Data Processing
- Checklist
- Parameter Reference
- API

cranium

- Docs »
- Checklist
- View page source

---

### Checklist¶

1. Convert data to HDF5 files containing a single sample/channel per file
2. Train Ilastik on each channel and process all files to create ‘\_Probability.h5’ files

Next 
 Previous

---

© Copyright 2018, Morgan Schwartz.

Built with Sphinx using a theme provided by Read the Docs.
