## Supplementary material for "ΔSCOPE: A new method to quantify 3D biological structures and identify differences in zebrafish forebrain development": An archived copy of the DeltaSCOPE code repository.: data-prep.html

  


Data Preparation — cranium 0.1.0 documentation


cranium

0.1.0

- Data Preparation
  - Biological Questions
  - File Format
  - Signal Normalization
- Data Processing
- Checklist
- Parameter Reference
- API

cranium

- Docs »
- Data Preparation
- View page source

---

### Data Preparation¶

#### Biological Questions¶

Cranium is designed to compare biological structures in three dimensions in order to cultivate sensitivity that is typically lost in maximum intensity projections (MIPs). In order to apply Cranium to a new biological structure, certain conditions must be satisfied:

1. The gross morphology of the structure must be roughly consistent between samples.
2. The x, y, and z dimensions cannot have the same size/extent. For example, in the zebrafish spinal cord, the anterior-posterior axis is the longest and the medial-lateral axis is the smallest with the dorsal-ventral axis falling between these two dimensions. The different proportional sizes of these axes enables us to consistently align the structure in 3D space regardless of the sample’s orientation during image collection.
3. The gross morphology of the structure can be described with a simple polynomial equation. For example, the spinal cord can be described by a line that falls at the midline of the medial-lateral axis: y = mx + b

#### File Format¶

Most microscopes save data in their own data format: for example, Zeiss, `.czi`; Leica, `.lif`. In order to ensure that image data is legible to all components of the workflow, files need to be converted to the HDF5 (`.h5`) format. This conversion can be easily executed in Fiji using the BioFormats and HDF5 plugin. At this point, individual channels need to be split and saved in their own files.

#### Signal Normalization¶

Biological fluorescence microscopy data contains variation in signal intensity due to both biological and technical error. For example, the top of the sample is frequently brighter than the bottom because it is closer to the objective and has not been bleached by collecting previous optical sections. If we were to try to select the set of points that represent ‘true signal’ by applying a single intensity threshold, points that represent background at the top of the stack may have the same intensity as points of true signal at the bottom of the stack.

In order to implement an adaptive thresholding protocol that avoids this challenges, we are using the open-source software, Ilastik. This software uses machine learning principles in order to predict the likelihood that a particular pixel contains true signal. The probability is calculated based on user annotation of images, in which regions of true signal and background are labeled. This protocol allows the user to apply their domain knowledge of the sample in order to best distinguish signal from background. Tutorials describing how to install and implement Ilastik’s pixel classification workflows are available on Ilastik’s website.

Next 
 Previous

---

© Copyright 2018, Morgan Schwartz.

Built with Sphinx using a theme provided by Read the Docs.
