## Supplementary material for "ΔSCOPE: A new method to quantify 3D biological structures and identify differences in zebrafish forebrain development": An archived copy of the DeltaSCOPE code repository.: data-process.html

  


Cranium Data Processing — cranium 0.1.0 documentation


cranium

0.1.0

- Data Preparation
- Data Processing
  - Intensity Thresholding
    - Goal
    - Setting Parameters
    - Code Instructions
  - Sample alignment using principle component analysis (PCA)
    - Goal
      - Principle Component Analysis
      - Structural Channel Processing
      - Model Fitting
    - Setting Parameters
    - Code Instructions
  - Cylindrical Coordinates
    - Goal
    - Code Instructions
- Checklist
- Parameter Reference
- API

cranium

- Docs »
- Cranium Data Processing
- View page source

---

### Cranium Data Processing¶

#### Intensity Thresholding¶

##### Goal¶

At this point, each sample/channel should have been processed by Ilastik to create a new `c1_10_Probabilities.h5` file. If you open this file in Fiji, it should contain three dimensions and two channels (signal and background, but for our purposes the two are interchangeable). Each pixel should have a value ranging between 0 and 1. If we are inspecting the signal channel, pixels with a value close to 0 are highly likely to be true signal. Correspondingly, pixels with a value close to 1 are likely to be background. The `c1_10_Probabilities.h5` file by Ilastik contains two channels (signal and background), which contain pixel values that are direct opposites. For a given pixel, the background intensity value is 1 minus the signal intensity value. In order to simplify our data, we will apply a threshold that will divide the data into two sets of pixels: signal and background. In the steps that follow, we will only use the set of pixels that correspond to true signal. This set of pixels may be also referred to as a set of points or a point cloud. In order to avoid keeping track of which channel is the signal channel, we assume that after applying a threshold there will be more points will fall in the background group than in the signal group.

Warning

If your data contains more points of signal than background, cranium will select your background channel as the signal channel. In order to correct this assumption, the inequality in `brain.read_data()` will need to be changed to equate a large number of points with signal instead of background.

##### Setting Parameters¶

In order to divide the image into two sets of pixels (signal and background), we use a parameter, `genthresh`, to determine which group the pixels fall into. We have found that a value of 0.5 is sufficient to divide the data and have not found that the results vary greatly if it is changed; however, if your data contains a lot of intermediate background values (0.4-0.7), you may benefit from a smaller threshold, e.g. 0.3.

The following code requires a parameter `micron`, which specifies the dimensions of the image voxel in microns. It is a list of the form `[x,y,z]`. This information can typically be found in the metadata of a microscope collection file.

This section of the code also includes a deprecated parameter `scale`, which must be set to `[1,1,1]`.

##### Code Instructions¶

The following instructions apply to processing a single sample. Details regarding function parameters can be found under `embryo` and `brain.preprocess_data()`.

```
#Create an embryo object that facilitates data processing
e = cranium.embryo(experiment-name,sample-number,directory)

#For each channel in your sample, add a channel with a unique name, e.g. 'c1' or 'c2'
e.add_channel(c1-filepath,c1-name)
e.add_channel(c2-filepath,c2-name)

#Threshold each channel and scale points according to voxel dimension in microns
e.chnls[c1-name].preprocess_data(genthresh,scale,microns)
e.chnls[c2-name].preprocess_data(genthresh,scale,microns)
```

#### Sample alignment using principle component analysis (PCA)¶

##### Goal¶

###### Principle Component Analysis¶

During typical collections of biological samples, each sample will be oriented slightly differently in relation to the microscope due to variations in the shape and size of the sample as well as human error during the mounting process. As a result of this variation, we cannot directly compare samples in 3D space without realigning them. We have implemented principle component analysis in order to automate the process of alignment without a need for human supervision. Fig. 1 shown below illustrates how PCA can be used to align a set of points in 2D. Cranium uses the same process in 3D to identify biologically meaningful axes present in biological structures.

Fig. 1 Principle component analysis (PCA) can be used to identify and align samples along consistent axes. (1) In 2D, PCA first selects the axis that captures the most variability in the data (1st PC). The 2nd PC is selected in a position orthogonal to the 1st PC that captures the remaining variation in the data. (2) The data are then rotated so that the 1st and 2nd PCs correspond with x and y axes respectively. (3) Finally, we have added a step of identifying the center of the data and shifting it to the origin.

###### Structural Channel Processing¶

We designate one channel, the `structural channel`, which we will use for PCA to align samples. Since we are interested in the gross morphology of this channel, we apply two data preprocessing steps to reduce the data down to only essential points. First, we return to `brain.raw_data` and apply a new threshold, `medthresh`, which is typically more stringent than `genthresh`. This step ensures we are only considering points of signal with extremely high likelihood of being real. Second, we apply a median filter to the data twice, which smooths out the structure and eliminates small points of variation that may interfere with the alignment process of the gross structure.

PCA outputs three new dimensions, the 1st, 2nd, and 3rd PCs. These components will be reassigned to the X, Y, and Z axes to help the user maintain orientation in regards to their data. In the case of the zebrafish post-optic commissure shown below (Fig. 2), the 1st PC is reassigned to the X axis and the 2nd and 3rd PCs are assigned to Z and Y respectively. These assignments honor the user’s expectation of the sample’s alignment in 3D space. The assignment of components to axes can be modified using the parameter `comporder`.

Warning

In order for PCA to consistently align your samples in the same orientation, we are assuming that the three dimensions of your data are of different relative sizes. Since PCA looks for the axes that capture the most variation in your data, a sample that has axes of the same relative size will not have any distinguishing characteristics that PCA can use to identify and separate different axes.

Fig. 2 This example illustrates the efficacy of PCA at changing the orientation of the zebrafish post optic commissure. In this case, the 1st PC is significantly longer than the 2nd and 3rd. While these two remaining components are similar in size, the typically longer depth of the 2nd PC distinguishes it from the 3rd PC.

###### Model Fitting¶

In addition to rotating the orientation of the data in 3D space, we also want to align the center of all samples at the origin. In order to determine the center of the data, we fit a simple mathematical model to the data that will also be used later in the analysis. The zebrafish post optic commissure shown above forms a parabolic structure, which can be described by y = ax^2 + bx + c. For simplicity, we fit the model in two dimensions, while holding one dimension constant. In the case of the POC, the parabolic structure lies flat in the XZ plane, which means that the structure can be described using exclusively the X and Z dimensions. The dimensions, which will be used to fit the 2D model, are specified in the parameter `fitdim`. Additionally the `deg` parameter specifies the degree of the function that fits to the data.

##### Setting Parameters¶

`medthresh` is typically set to 0.25, in comparison to a value of 0.5 for `genthresh`. If your data contains aberrant signal that does not contribute to the gross morphology of the structure, an even lower `medthresh` may help limit the negative influence of noisy signal. Additionally, the `radius` of the median filter can also be tuned to eliminate noisy signal. The typical value for `radius` is 20, which refers to the number of neighboring points that are considered in the median filter. A smaller value for `radius` will preserve small variation in signal, while a larger value will cause even more blunting and smoothing of the data.

The `comporder` parameter controls how principle components are reassigned to the typical Cartesian coordinate system (XYZ) that most users are familiar with. It takes the form of an array of length 3 that specifies the index of the component that will be assigned to the X, Y, or Z axis: `[x index,y index,z index]`. Please note that the index that matches each principle component starts counting at 0, e.g. 1st PC = 0, 2nd PC = 1, and 3rd PC = 2. For example, if we want to assign the 1st PC to the x axis, the 2nd to the Z axis, and the 3rd to the y axis, the `comporder` parameter would be `[0,2,1]`.

Finally, the remaining two parameters determines how the model will be fit to the data. `fitdim` determines which 2 axes will be used to fit the 2D model. It takes the form of a list of 2 of the 3 dimensions specified as a lowercase string, e.g. `'x','y','z'`. If we wanted to fit a model in the XZ plane, while holding the Y axis constant, the `fitdim` parameter would be `['x','z']`. `deg` specifies the degree of the function that will be fit to the data. The default is `2`, which specifies a parabolic function. A deg of `1` would fit a linear function.

Warning

The ability to specify degrees other than 2 is still being developed. Check here for updates.

##### Code Instructions¶

```
#Run PCA on the structural channel, in this case, c1
e.chnls['c1'].calculate_pca_median(e.chnls['c1'].raw_data,medthresh,radius,microns)

#Save the pca object that includes the transformation matrix
pca = e.chnls['c1'].pcamed

#Transform the structural channel using the saved pca object
e.chnls['c1'].pca_transform_3d(e.chnls['c1'].df_thresh,pca,comporder,fitdim,deg=2)

#Save the mathematical model and vertex (center point) of the structural channel
mm = e.chnls['AT'].mm
vertex = e.chnls['AT'].vertex

#Transform any additional channels using the pca object calculated based on the structural channel
e.chnls['c2'].pca_transform_3d(e.chnls['c2'].df_thresh,pca,comporder,fitdim,deg=2,mm=mm,vertex=vertex)
```

#### Cylindrical Coordinates¶

##### Goal¶

The ultimate goal with this workflow of data processing is to enable us to study small differences in biological structures when comparing a set of control samples to experimental samples. While our samples are now aligned to all fall in the same region of 3D space, our points are still defined by the xyz coordinates defined by the microscope. In order to detect changes in our structure, we will define the position of points relative to the structure using a cylindrical coordinate system. We will rely on the previously defined mathematical model, `brain.mm`, to represent the underlying shape of our structure. From there we will define the position of each point relative to the mathematical model (Fig. 3). The first dimension, R, is defined as the shortest distance between the point and the model. Second, alpha is defined as the distance from the point’s intersection with the model to the midline or center of the structure. Third, the position of the point around the model is defined in theta. Following the completion of the transformation, the final dataset is saved to a `.psi` file.

Fig. 3 To enable analysis of data point relative to a biological structure, points are transformed from a Cartesian coordinate system (x,y,z) into a cylindrical coordinate system ($alpha$,$theta$,R) defined relative to the structure.

##### Code Instructions¶

This transformation does not require defining any parameters; however, it assumes the data has already been thresholded and aligned using PCA.

```
#Transform each channel to cylindrical coordinates
e.chnls['c1'].transform_coordinates()
e.chnls['c2'].transform_coordinates()

#Save processed data to .psi file
e.save_psi()
```

Warning

This processing step is time consuming. We recommend running multiple samples in parallel in order to reduce the total amount of computational time required.

Todo

Add multiprocessing instructions

Next 
 Previous

---

© Copyright 2018, Morgan Schwartz.

Built with Sphinx using a theme provided by Read the Docs.
