## Supplementary material for "ΔSCOPE: A new method to quantify 3D biological structures and identify differences in zebrafish forebrain development": An archived copy of the DeltaSCOPE code repository.: genindex.html

  


Index — cranium 0.1.0 documentation


cranium

0.1.0

- Data Preparation
- Data Processing
- Checklist
- Parameter Reference
- API

cranium

- Docs »
- Index

---

### Index

**A**
| **B**
| **C**
| **D**
| **E**
| **F**
| **G**
| **I**
| **L**
| **M**
| **N**
| **O**
| **P**
| **R**
| **S**
| **T**
| **W**

## A

|  |  |
| --- | --- |
| - acbins (cranium.landmarks attribute) - add\_aligned\_df() (cranium.brain method) - add\_channel() (cranium.embryo method) | - add\_psi\_data() (cranium.embryo method) - add\_thresh\_df() (cranium.brain method) - align\_data() (cranium.brain method) |

## B

- brain (class in cranium)

## C

|  |  |
| --- | --- |
| - calc\_bins() (cranium.landmarks method) - calc\_coord() (cranium.brain method) - calc\_mt\_landmarks() (cranium.landmarks method) - calc\_perc() (cranium.landmarks method) - calc\_wt\_reformat() (cranium.landmarks method) - calculate\_area\_error() (in module cranium) - calculate\_models() (in module cranium) | - calculate\_pca\_median() (cranium.brain method) - calculate\_pca\_median\_2d() (cranium.brain method) - cf (cranium.math\_model attribute) - chnls (cranium.embryo attribute) - comporder, [1], [2], [3] - cranium (module) - create\_dataframe() (cranium.brain method) |

## D

|  |  |
| --- | --- |
| - deg, [1] - df\_align (cranium.brain attribute) | - df\_scl (cranium.brain attribute) - df\_thresh (cranium.brain attribute) |

## E

- embryo (class in cranium)
- environment variable
  - comporder, [1], [2], [3], [4]
  - deg, [1], [2]
  - fitdim, [1], [2], [3], [4]
  - genthresh, [1], [2], [3], [4], [5]
  - medthresh, [1], [2], [3]
  - micron
  - microns, [1]
  - radius, [1], [2], [3], [4], [5], [6]
  - scale, [1]

## F

|  |  |
| --- | --- |
| - find\_arclength() (cranium.brain method) - find\_distance() (cranium.brain method) - find\_min\_distance() (cranium.brain method) - find\_r() (cranium.brain method) | - find\_theta() (cranium.brain method) - fit\_model() (cranium.brain method) - fitdim, [1], [2], [3] - flip\_data() (cranium.brain method) |

## G

|  |  |
| --- | --- |
| - generate\_kde() (in module cranium) | - genthresh, [1], [2], [3], [4] |

## I

- integrand() (cranium.brain method)

## L

|  |  |
| --- | --- |
| - landmarks (class in cranium) | - lm\_mt\_rf (cranium.landmarks.brain attribute) - lm\_wt\_rf (cranium.landmarks.brain attribute) |

## M

|  |  |
| --- | --- |
| - math\_model (class in cranium) - median (cranium.brain attribute) - medthresh, [1], [2] | - micron - microns - mm (cranium.brain attribute) |

## N

|  |  |
| --- | --- |
| - name (cranium.embryo attribute) | - number (cranium.embryo attribute) |

## O

- outdir (cranium.embryo attribute)

## P

|  |  |
| --- | --- |
| - p (cranium.math\_model attribute) - pca\_transform\_2d() (cranium.brain method) - pca\_transform\_3d() (cranium.brain method) - pcamed (cranium.brain attribute) - percbins (cranium.landmarks.brain attribute) | - plot\_projections() (cranium.brain method) - preprocess\_data() (cranium.brain method) - process\_alignment\_data() (cranium.brain method) - process\_channels() (cranium.embryo method) - process\_sample() (in module cranium) |

## R

|  |  |
| --- | --- |
| - radius, [1], [2], [3], [4], [5] - raw\_data (cranium.brain attribute) - read\_data() (cranium.brain method) - read\_psi() (in module cranium) | - read\_psi\_to\_dict() (in module cranium) - reformat\_to\_cart() (in module cranium) - rescale\_variable() (in module cranium) - rnull (cranium.landmarks.brain attribute) |

## S

|  |  |
| --- | --- |
| - save\_projections() (cranium.embryo method) - save\_psi() (cranium.embryo method) - scale | - subplot\_lmk() (in module cranium) - subset (cranium.brain attribute) - subset\_data() (cranium.brain method) |

## T

|  |  |
| --- | --- |
| - tbins (cranium.landmarks attribute) | - threshold (cranium.brain attribute) - transform\_coordinates() (cranium.brain method) |

## W

|  |  |
| --- | --- |
| - write\_data() (in module cranium) | - write\_header() (in module cranium) |

---

© Copyright 2018, Morgan Schwartz.

Built with Sphinx using a theme provided by Read the Docs.
