## Supplementary material for "ΔSCOPE: A new method to quantify 3D biological structures and identify differences in zebrafish forebrain development": An archived copy of the DeltaSCOPE code repository.: index.html

  


Welcome to cranium’s documentation! — cranium 0.1.0 documentation


cranium

0.1.0

- Data Preparation
- Data Processing
- Checklist
- Parameter Reference
- API

cranium

- Docs »
- Welcome to cranium’s documentation!
- View page source

---

### Welcome to cranium’s documentation!¶

Contents:

- Data Preparation
  - Biological Questions
  - File Format
  - Signal Normalization
- Data Processing
  - Intensity Thresholding
    - Goal
    - Setting Parameters
    - Code Instructions
  - Sample alignment using principle component analysis (PCA)
    - Goal
      - Principle Component Analysis
      - Structural Channel Processing
      - Model Fitting
    - Setting Parameters
    - Code Instructions
  - Cylindrical Coordinates
    - Goal
    - Code Instructions
- Checklist
- Parameter Reference
- API

#### Is cranium right for you?¶

Can dos:
\* Align single 3D microscopy collections to an orientation that is consistent between samples

Cannot:
\* Align cryosections to create a 3D volume from 2D slices

#### Issues¶

### Indices and tables¶

- Index
- Module Index
- Search Page

Next

---

© Copyright 2018, Morgan Schwartz.

Built with Sphinx using a theme provided by Read the Docs.
