## Supplementary material for "ΔSCOPE: A new method to quantify 3D biological structures and identify differences in zebrafish forebrain development": An archived copy of the DeltaSCOPE code repository.: param-ref.html

  


Parameter Reference — cranium 0.1.0 documentation


cranium

0.1.0

- Data Preparation
- Data Processing
- Checklist
- Parameter Reference
- API

cranium

- Docs »
- Parameter Reference
- View page source

---

### Parameter Reference¶

`genthresh`¶
:   This parameter defines the cutoff point that will divide the `brain.raw_data` into a set of true signal points and background points based on the probability that each point is true signal. The `_Probabilties.h5` dataset is generated after running the Ilastik pixel classification workflow described here. Pixels with a value of 1 are likely to be background. Correspondingly, pixels with a value close to 0 are most likely to be true signal. We have found that a threshold of 0.5 is sufficient to divide true signal from background; however, if your data contains a lot of intermediate background values (0.4-0.7), you may benefit from a smaller threshold, e.g. 0.3.

    Recommended: `0.5`

`microns`¶
:   This parameter is a list of the form `[x,y,z]` that specifies the dimensions of the voxel in microns. The data in `brain.raw_data` is scaled by `microns` to control for datasets in which the `z` dimension is larger than the `x` and `y` dimensions.

    Example: `[0.16,0.16,0.21]`

`scale`¶
:   This is a deprecated parameter that should always be set to `[1,1,1]`.

    Required: `[1,1,1]`

`medthresh`¶
:   This parameter serves the same purpose as `genthresh`; however, it is used exclusively on data used for aligning samples using PCA. This threshold is typically more stringent than `genthresh` to ensure that any noise in the data does not interfere with the alignment process.

    Recommended: `0.25`

`radius`¶
:   The `radius` of the median filter can be tuned to eliminate noisy signal. The typical value for `radius` is 20, which refers to the number of neighboring points that are considered in the median filter. A smaller value for `radius` will preserve small variation in signal, while a larger value will cause even more blunting and smoothing of the data.

    Recommended: `20`

`comporder`¶
:   This parameter controls how principle components are reassigned to the typical Cartesian coordinate system (XYZ) that most users are familiar with. It takes the form of an array of length 3 that specifies the index of the component that will be assigned to the X, Y, or Z axis: `[x index,y index,z index]`. Please note that the index that matches each principle component starts counting at 0, e.g. 1st PC = 0, 2nd PC = 1, and 3rd PC = 2.

    For example, if we want to assign the 1st PC to the x axis, the 2nd to the Z axis, and the 3rd to the y axis, the `comporder` parameter would be `[0,2,1]`.

    Example: `[0,2,1]`

`fitdim`¶
:   This parameter determines which 2 axes will be used to fit the 2D model. It takes the form of a list of 2 of the 3 dimensions specified as a lowercase string, e.g. `'x','y','z'`.

    If we wanted to fit a model in the XZ plane, while holding the Y axis constant, the `fitdim` parameter would be `['x','z']`.

    Example: `['x','z']`

`deg`¶
:   This parameter specifies the degree of the function that will be fit to the data. The default is `2`, which specifies a parabolic function. A deg of `1` would fit a linear function.

    Default: `2`

    Warning

    The infrastructure to support degrees other than 2 is not currently in place. Check here for updates.

Next 
 Previous

---

© Copyright 2018, Morgan Schwartz.

Built with Sphinx using a theme provided by Read the Docs.
