## Supplementary material for "ΔSCOPE: A new method to quantify 3D biological structures and identify differences in zebrafish forebrain development": An archived copy of the DeltaSCOPE code repository.: py-modindex.html

  

Python Module Index — cranium 0.1.0 documentation

cranium

0.1.0

- Data Preparation
- Data Processing
- Checklist
- Parameter Reference
- API

cranium

- Docs »
- Python Module Index

---

### Python Module Index

**c**

|  |  |
| --- | --- |
|  | **c** |
|  | `cranium` |

---

© Copyright 2018, Morgan Schwartz.

Built with Sphinx using a theme provided by Read the Docs.
