## Supplementary material for "ΔSCOPE: A new method to quantify 3D biological structures and identify differences in zebrafish forebrain development": An archived copy of the DeltaSCOPE code repository.: search.html

  


Search — cranium 0.1.0 documentation


cranium

0.1.0

- Data Preparation
- Data Processing
- Checklist
- Parameter Reference
- API

cranium

- Docs »
- Search

---

Please activate JavaScript to enable the search
functionality.

---

© Copyright 2018, Morgan Schwartz.

Built with Sphinx using a theme provided by Read the Docs.
