## Supplementary material for "ΔSCOPE: A new method to quantify 3D biological structures and identify differences in zebrafish forebrain development": An archived copy of the DeltaSCOPE code repository.: searchindex.js

```
Search.setIndex({docnames:["API","checklist","data-prep","data-process","index","param-ref","start-here"],envversion:53,filenames:["API.rst","checklist.rst","data-prep.rst","data-process.rst","index.rst","param-ref.rst","start-here.rst"],objects:{"":{comporder:[5,5,1,"-"],cranium:[0,0,0,"-"],deg:[5,5,1,"-"],fitdim:[5,5,1,"-"],genthresh:[5,5,1,"-"],medthresh:[5,5,1,"-"],microns:[5,5,1,"-"],radius:[5,5,1,"-"],scale:[5,5,1,"-"]},"cranium.brain":{add_aligned_df:[0,2,1,""],add_thresh_df:[0,2,1,""],align_data:[0,2,1,""],calc_coord:[0,2,1,""],calculate_pca_median:[0,2,1,""],calculate_pca_median_2d:[0,2,1,""],create_dataframe:[0,2,1,""],df_align:[0,3,1,""],df_scl:[0,3,1,""],df_thresh:[0,3,1,""],find_arclength:[0,2,1,""],find_distance:[0,2,1,""],find_min_distance:[0,2,1,""],find_r:[0,2,1,""],find_theta:[0,2,1,""],fit_model:[0,2,1,""],flip_data:[0,2,1,""],integrand:[0,2,1,""],median:[0,3,1,""],mm:[0,3,1,""],pca_transform_2d:[0,2,1,""],pca_transform_3d:[0,2,1,""],pcamed:[0,3,1,""],plot_projections:[0,2,1,""],preprocess_data:[0,2,1,""],process_alignment_data:[0,2,1,""],raw_data:[0,3,1,""],read_data:[0,2,1,""],subset:[0,3,1,""],subset_data:[0,2,1,""],threshold:[0,3,1,""],transform_coordinates:[0,2,1,""]},"cranium.embryo":{add_channel:[0,2,1,""],add_psi_data:[0,2,1,""],chnls:[0,3,1,""],name:[0,3,1,""],number:[0,3,1,""],outdir:[0,3,1,""],process_channels:[0,2,1,""],save_projections:[0,2,1,""],save_psi:[0,2,1,""]},"cranium.landmarks":{acbins:[0,3,1,""],calc_bins:[0,2,1,""],calc_mt_landmarks:[0,2,1,""],calc_perc:[0,2,1,""],calc_wt_reformat:[0,2,1,""],tbins:[0,3,1,""]},"cranium.landmarks.brain":{lm_mt_rf:[0,3,1,""],lm_wt_rf:[0,3,1,""],percbins:[0,3,1,""],rnull:[0,3,1,""]},"cranium.math_model":{cf:[0,3,1,""],p:[0,3,1,""]},cranium:{brain:[0,1,1,""],calculate_area_error:[0,4,1,""],calculate_models:[0,4,1,""],embryo:[0,1,1,""],generate_kde:[0,4,1,""],landmarks:[0,1,1,""],math_model:[0,1,1,""],process_sample:[0,4,1,""],read_psi:[0,4,1,""],read_psi_to_dict:[0,4,1,""],reformat_to_cart:[0,4,1,""],rescale_variable:[0,4,1,""],subplot_lmk:[0,4,1,""],write_data:[0,4,1,""],write_header:[0,4,1,""]}},objnames:{"0":["py","module","Python module"],"1":["py","class","Python class"],"2":["py","method","Python method"],"3":["py","attribute","Python attribute"],"4":["py","function","Python function"],"5":["std","envvar","environment variable"]},objtypes:{"0":"py:module","1":"py:class","2":"py:method","3":"py:attribute","4":"py:function","5":"std:envvar"},terms:{"1st":[3,5],"2nd":[3,5],"2pi":0,"3rd":[3,5],"boolean":0,"case":3,"class":0,"default":[0,3,5],"final":3,"float":0,"function":[0,3,5],"import":0,"int":0,"new":[0,2,3],"null":0,"return":[0,3],"true":[0,2,3,5],"try":2,"var":0,"while":[3,5],For:[2,3,5],PCs:3,The:[0,2,3,5],These:3,_mip:0,_probabilti:5,_probabl:[1,3],aberr:3,abil:3,abov:[0,3],absolut:0,absv:0,ac_num:0,acbin:0,access:0,accord:3,adapt:2,add:[0,3],add_aligned_df:0,add_channel:[0,3],add_psi_data:0,add_thresh_df:0,added:[0,3],addit:3,addition:3,after:[0,3,5],align:[0,2,4,5],align_data:0,all:[0,1,2,3],allow:[0,2],along:[0,3],alpah:0,alpha:[0,3],alreadi:[0,3],also:3,alwai:5,amount:3,analysi:[0,4],angl:0,ani:[0,3,5],annot:2,anterior:2,api:4,append:0,appli:[0,2,3],appropri:0,arc:0,arclength:0,area:0,around:3,arrai:[0,3,5],assign:[0,3,5],associ:0,assum:[0,3],assumpt:[0,3],attempt:0,attribut:0,autom:3,avail:2,averag:0,avg:0,avoid:[2,3],axes:[2,3,5],axi:[0,2,3,5],background:[2,3,5],base:[0,2,3,5],becaus:2,been:[0,2,3],being:[0,3],belong:0,below:3,benefit:[3,5],best:2,between:[0,2,3,4],bin:0,bioformat:2,biolog:[0,3,4],bleach:2,blunt:[3,5],bool:0,both:[0,2],bottom:2,boundari:0,brain:[0,3,5],brighter:2,c1_10:3,calc_bin:0,calc_coord:0,calc_mt_landmark:0,calc_perc:0,calc_wt_reformat:0,calcul:[0,2,3],calculate_area_error:0,calculate_model:0,calculate_pca_median:[0,3],calculate_pca_median_2d:0,call:0,can:[0,2,3,4,5],cannot:[0,2,3,4],captur:3,cartesian:[0,3,5],caus:[3,5],center:3,certain:2,challeng:2,chang:3,channel:[0,1,2,4],characterist:3,check:[3,5],checklist:4,chnl:[0,3],chs:0,circl:0,classif:[2,5],close:[0,3,5],closer:2,closest:0,cloud:3,cmap:0,code:[0,4],coeffici:0,collect:[2,3,4],column:0,commissur:3,comp_ord:0,compar:[2,3],comparison:3,complet:[0,3],compon:[0,2,4,5],compord:[3,5],comput:3,conclus:0,condit:2,consid:[0,3,5],consist:[2,3,4],constant:[0,3,5],consum:3,contain:[0,1,2,3,5],content:4,contribut:3,control:[0,3,5],convers:2,convert:[0,1,2],coordin:[0,4,5],cord:2,correct:3,correspond:[0,3],correspondingli:[3,5],count:[3,5],cranium:[0,2],creat:[0,1,3,4],create_datafram:0,cryosect:4,csv:0,cultiv:2,current:5,curv:0,cutoff:[0,5],cylindr:4,czi:2,data:[0,1,4,5],datafram:0,datapoint:0,dataset:[0,3,5],date:0,ddf:0,defin:[0,3,5],deg:[0,3,5],degre:[0,3,5],deprec:[0,3,5],depth:3,deriv:0,describ:[0,2,3,5],descript:0,design:[2,3],detail:3,detect:3,determin:[0,3,5],develop:3,df_align:0,df_fit:0,df_scl:0,df_thresh:[0,3],dfs:0,dict:0,dictionari:0,differ:[2,3],dimens:[0,2,3,5],direct:3,directli:3,directori:[0,3],distanc:[0,3],distinguish:[2,3],distribut:0,divid:[3,5],divis:0,doe:[0,3,5],domain:2,dorsal:2,dos:4,down:3,dtype:0,due:[2,3],dure:[2,3],each:[0,1,3,5],easili:2,efficaci:3,either:0,elimin:[3,5],embryo:[0,3],enabl:[2,3],ensur:[0,2,3,5],equat:[2,3],error:[0,2,3],essenti:3,euclidean:0,evalu:0,even:[3,5],evenli:0,examin:0,exampl:[2,3,5],exclus:[3,5],execut:2,expect:3,experi:3,experiment:[0,3],extent:2,extrem:3,facilit:3,fail:0,fall:[2,3],fals:0,familiar:[3,5],fig:3,figur:0,fiji:[2,3],file:[0,1,3,4],filenam:0,filepath:[0,3],filter:[0,3,5],find:0,find_arclength:0,find_dist:0,find_min_dist:0,find_r:0,find_theta:0,first:[0,3],fit:[0,4,5],fit_dim:0,fit_model:0,fitdim:[3,5],flat:3,flip:0,flip_data:0,flip_dim:0,fluoresc:2,follow:[0,3],form:[0,3,5],format:[0,4],found:[0,3,5],fraction:0,frequent:2,from:[0,2,3,4,5],gener:[0,5],generate_kd:0,genthresh:[3,5],get:0,given:[0,3],goal:4,greatli:3,greys_r:0,gross:[2,3],group:[0,3],gthresh:0,handl:0,has:[0,2,3],have:[0,2,3,5],hdf5:[0,1,2],header:0,help:3,here:[3,5],high:3,highli:3,hold:[3,5],honor:3,how:[2,3,5],howev:[3,5],human:3,identifi:[0,3],ilastik:[1,2,3,5],illustr:3,imag:[0,2,3],implement:[2,3],includ:[0,3],index:[0,3,4,5],indic:0,individu:2,inequ:3,influenc:3,inform:3,infrastructur:5,input:0,inspect:3,instal:2,instead:3,instruct:4,integ:0,integr:0,integrand:0,intens:[2,4],interchang:3,interest:3,interfer:[0,3,5],intermedi:[3,5],intersect:3,kde:0,keep:3,kei:0,knowledg:2,label:[0,2],landmark:0,larg:3,larger:[3,5],later:[2,3],ldf:0,learn:2,least:0,legibl:2,leica:2,length:[0,3,5],lies:3,lif:2,like:[3,5],likelihood:[2,3],limit:3,line:[0,2],linear:[3,5],list:[0,3,5],lkde:0,lm_mt_rf:0,lm_wt_rf:0,longer:3,longest:2,look:3,lost:2,lot:[3,5],lower:[0,3],lowercas:[3,5],machin:2,made:0,mai:[2,3,5],maintain:3,manag:0,match:[3,5],math:0,math_model:0,mathemat:3,matplotlib:0,matrix:[0,3],maximum:2,mean:[0,3],meaning:3,medial:2,median:[0,3,5],medthresh:[3,5],metadata:3,micron:[0,3,5],microscop:[2,3],microscopi:[2,4],midlin:[2,3],minim:0,minimum:0,minu:3,mip:2,model:[0,4,5],modifi:3,modul:4,more:[3,5],morpholog:[2,3],most:[0,2,3,5],mount:3,mtc:0,mthresh:0,multichannel:0,multipl:[0,3],multipli:0,multiprocess:3,must:[0,2,3],mutant:0,name:[0,3],need:[2,3],neg:3,neighbor:[3,5],neighborhood:0,newvar:0,nois:[0,5],noisi:[3,5],none:0,normal:4,note:[3,5],now:3,num:0,number:[0,3,5],object:[0,2,3],one:3,onli:[0,3],onto:0,open:[0,2,3],opposit:3,optic:[2,3],optim:0,order:[2,3],orient:[2,3,4],origin:[0,3],orthogon:3,other:[3,5],our:3,out:[0,3],outdir:0,output:[0,3],own:2,page:4,panda:0,parabol:[0,3,5],parallel:3,paramet:[0,4],parr:0,particular:[0,2],path:0,pca:[0,4,5],pca_object:0,pca_transform_2d:0,pca_transform_3d:[0,3],pcame:[0,3],pdf:0,per:1,percbin:0,percentag:0,percentil:0,pertain:0,pixel:[0,2,3,5],place:5,plane:[0,3,5],pleas:[3,5],plot:0,plot_project:0,plt:0,plugin:2,png:0,poc:3,point:[0,2,3,5],poly1d:0,polyfit:0,polynomi:2,popul:0,posit:[0,3],post:3,posterior:2,postion:0,predict:2,prefix:0,prepar:4,preprocess:3,preprocess_data:[0,3],present:3,preserv:[3,5],previou:2,previous:[0,3],primari:0,primary_kei:0,principl:[2,4,5],prior:0,probabl:[0,2,5],process:[0,1,4,5],process_alignment_data:0,process_channel:0,process_sampl:0,produc:0,product:0,project:[0,2],proport:[0,2],protocol:2,psi:[0,3],pts:0,purpos:[3,5],pythagorean:0,question:4,r_std:0,radiu:[0,3,5],random:0,randomli:0,rang:3,raw:0,raw_data:[0,3,5],read:0,read_data:[0,3],read_psi:0,read_psi_to_dict:0,real:3,realign:3,reassign:[3,5],recommend:[3,5],reduc:3,refer:[3,4],reformat_to_cart:0,regard:3,regardless:2,region:[2,3],rel:[0,3],relat:3,relationship:0,reli:3,remain:3,repres:[0,2,3],requir:[3,5],rescal:0,rescale_vari:0,respect:[0,3],result:[0,3],ribbon:0,rnull:0,root:0,rotat:[0,3],roughli:2,row:0,run:[3,5],same:[0,2,3,5],sampl:[0,1,2,4,5],sample_frac:0,satisfi:2,save:[0,2,3],save_project:0,save_psi:[0,3],scale:[0,3,5],scatterplot:0,scenario:0,scheme:0,scipi:0,search:4,second:[0,3],section:[2,3],select:[0,2,3],sem:0,sensit:2,separ:3,seri:0,serv:5,set:[0,2,4,5],shape:[0,3],shift:3,shortest:3,should:[0,3,5],shown:3,signal:[0,3,4,5],significantli:3,similar:3,simpl:[2,3],simplic:3,simplifi:3,sinc:3,singl:[0,1,2,3,4],size:[0,2,3],slice:4,slightli:3,small:[3,5],smaller:[3,5],smallest:2,smooth:[3,5],snum:0,softwar:2,sourc:[0,2],space:[2,3],specifi:[0,3,5],spinal:2,split:2,stack:2,standard:0,start:[3,5],step:3,still:3,str:0,string:[0,3,5],stringent:[3,5],structur:[0,2,4],studi:3,subplot:0,subplot_lmk:0,subsampl:0,subset:0,subset_data:0,suffici:[3,5],supervis:3,support:5,system:[0,3,5],take:[0,3,5],tarr:0,tbin:0,technic:2,than:[0,2,3,5],them:3,theorem:0,theshold:0,theta:[0,3],thi:[0,2,3,5],third:3,three:[0,2,3],threshold:[0,2,4,5],through:0,time:3,too:0,top:2,total:3,track:3,train:1,transform:[0,3],transform_coordin:[0,3],translat:0,tstep:0,tune:[3,5],tutori:2,tvalu:0,twice:[0,3],two:[0,2,3],type:0,typic:[0,2,3,5],ultim:3,under:3,underli:3,uniqu:3,updat:[3,5],use:[0,3],used:[0,3,5],user:[2,3,5],uses:[2,3],using:[0,2,4,5],usual:0,valu:[0,3,5],vari:3,variabl:[0,3],variat:[2,3,5],ventral:2,vertex:[0,3],volum:4,voxel:[0,3,5],want:[3,5],websit:2,well:3,were:2,what:0,when:[0,3],where:0,which:[0,2,3,5],whose:0,wildtyp:0,without:3,workflow:[2,3,5],would:[3,5],write:0,write_data:0,write_head:0,wtc:0,xarr:0,xvalu:0,xyz:[0,3,5],you:[3,5],your:[3,5],zebrafish:[2,3],zeiss:2,zero:0,zfb:0,zln:0,zpt:0,zrf1:0},titles:["API","Checklist","Data Preparation","Cranium Data Processing","Welcome to cranium\u2019s documentation!","Parameter Reference","Start Here"],titleterms:{align:3,analysi:3,api:0,biolog:2,channel:3,checklist:1,code:3,compon:3,coordin:3,cranium:[3,4],cylindr:3,data:[2,3],document:4,file:2,fit:3,format:2,goal:3,here:6,indic:4,instal:6,instruct:3,intens:3,issu:4,model:3,normal:2,paramet:[3,5],pca:3,prepar:2,principl:3,process:3,question:2,refer:5,right:4,sampl:3,set:3,signal:2,start:6,structur:3,tabl:4,threshold:3,todo:3,using:3,welcom:4,you:4}})
```
