## Supplementary material for "ΔSCOPE: A new method to quantify 3D biological structures and identify differences in zebrafish forebrain development": An archived copy of the DeltaSCOPE code repository.: Makefile

```
# Makefile for Sphinx documentation
#

# You can set these variables from the command line.
SPHINXOPTS    =
SPHINXBUILD   = sphinx-build
PAPER         =
BUILDDIR      = build

# User-friendly check for sphinx-build
ifeq ($(shell which $(SPHINXBUILD) >/dev/null 2>&1; echo $$?), 1)
$(error The '$(SPHINXBUILD)' command was not found. Make sure you have Sphinx installed, then set the SPHINXBUILD environment variable to point to the full path of the '$(SPHINXBUILD)' executable. Alternatively you can add the directory with the executable to your PATH. If you don't have Sphinx installed, grab it from http://sphinx-doc.org/)
endif

# Internal variables.
PAPEROPT_a4     = -D latex_paper_size=a4
PAPEROPT_letter = -D latex_paper_size=letter
ALLSPHINXOPTS   = -d $(BUILDDIR)/doctrees $(PAPEROPT_$(PAPER)) $(SPHINXOPTS) source
# the i18n builder cannot share the environment and doctrees with the others
I18NSPHINXOPTS  = $(PAPEROPT_$(PAPER)) $(SPHINXOPTS) source

.PHONY: help clean html dirhtml singlehtml pickle json htmlhelp qthelp devhelp epub latex latexpdf text man changes linkcheck doctest coverage gettext

help:
	@echo "Please use \`make <target>' where <target> is one of"
	@echo "  html       to make standalone HTML files"
	@echo "  dirhtml    to make HTML files named index.html in directories"
	@echo "  singlehtml to make a single large HTML file"
	@echo "  pickle     to make pickle files"
	@echo "  json       to make JSON files"
	@echo "  htmlhelp   to make HTML files and a HTML help project"
	@echo "  qthelp     to make HTML files and a qthelp project"
	@echo "  applehelp  to make an Apple Help Book"
	@echo "  devhelp    to make HTML files and a Devhelp project"
	@echo "  epub       to make an epub"
	@echo "  latex      to make LaTeX files, you can set PAPER=a4 or PAPER=letter"
	@echo "  latexpdf   to make LaTeX files and run them through pdflatex"
	@echo "  latexpdfja to make LaTeX files and run them through platex/dvipdfmx"
	@echo "  text       to make text files"
	@echo "  man        to make manual pages"
	@echo "  texinfo    to make Texinfo files"
	@echo "  info       to make Texinfo files and run them through makeinfo"
	@echo "  gettext    to make PO message catalogs"
	@echo "  changes    to make an overview of all changed/added/deprecated items"
	@echo "  xml        to make Docutils-native XML files"
	@echo "  pseudoxml  to make pseudoxml-XML files for display purposes"
	@echo "  linkcheck  to check all external links for integrity"
	@echo "  doctest    to run all doctests embedded in the documentation (if enabled)"
	@echo "  coverage   to run coverage check of the documentation (if enabled)"

clean:
	rm -rf $(BUILDDIR)/*

html:
	$(SPHINXBUILD) -b html $(ALLSPHINXOPTS) $(BUILDDIR)/html
	@echo
	@echo "Build finished. The HTML pages are in $(BUILDDIR)/html."

dirhtml:
	$(SPHINXBUILD) -b dirhtml $(ALLSPHINXOPTS) $(BUILDDIR)/dirhtml
	@echo
	@echo "Build finished. The HTML pages are in $(BUILDDIR)/dirhtml."

singlehtml:
	$(SPHINXBUILD) -b singlehtml $(ALLSPHINXOPTS) $(BUILDDIR)/singlehtml
	@echo
	@echo "Build finished. The HTML page is in $(BUILDDIR)/singlehtml."

pickle:
	$(SPHINXBUILD) -b pickle $(ALLSPHINXOPTS) $(BUILDDIR)/pickle
	@echo
	@echo "Build finished; now you can process the pickle files."

json:
	$(SPHINXBUILD) -b json $(ALLSPHINXOPTS) $(BUILDDIR)/json
	@echo
	@echo "Build finished; now you can process the JSON files."

htmlhelp:
	$(SPHINXBUILD) -b htmlhelp $(ALLSPHINXOPTS) $(BUILDDIR)/htmlhelp
	@echo
	@echo "Build finished; now you can run HTML Help Workshop with the" \
	      ".hhp project file in $(BUILDDIR)/htmlhelp."

qthelp:
	$(SPHINXBUILD) -b qthelp $(ALLSPHINXOPTS) $(BUILDDIR)/qthelp
	@echo
	@echo "Build finished; now you can run "qcollectiongenerator" with the" \
	      ".qhcp project file in $(BUILDDIR)/qthelp, like this:"
	@echo "# qcollectiongenerator $(BUILDDIR)/qthelp/cranium.qhcp"
	@echo "To view the help file:"
	@echo "# assistant -collectionFile $(BUILDDIR)/qthelp/cranium.qhc"

applehelp:
	$(SPHINXBUILD) -b applehelp $(ALLSPHINXOPTS) $(BUILDDIR)/applehelp
	@echo
	@echo "Build finished. The help book is in $(BUILDDIR)/applehelp."
	@echo "N.B. You won't be able to view it unless you put it in" \
	      "~/Library/Documentation/Help or install it in your application" \
	      "bundle."

devhelp:
	$(SPHINXBUILD) -b devhelp $(ALLSPHINXOPTS) $(BUILDDIR)/devhelp
	@echo
	@echo "Build finished."
	@echo "To view the help file:"
	@echo "# mkdir -p $$HOME/.local/share/devhelp/cranium"
	@echo "# ln -s $(BUILDDIR)/devhelp $$HOME/.local/share/devhelp/cranium"
	@echo "# devhelp"

epub:
	$(SPHINXBUILD) -b epub $(ALLSPHINXOPTS) $(BUILDDIR)/epub
	@echo
	@echo "Build finished. The epub file is in $(BUILDDIR)/epub."

latex:
	$(SPHINXBUILD) -b latex $(ALLSPHINXOPTS) $(BUILDDIR)/latex
	@echo
	@echo "Build finished; the LaTeX files are in $(BUILDDIR)/latex."
	@echo "Run \`make' in that directory to run these through (pdf)latex" \
	      "(use \`make latexpdf' here to do that automatically)."

latexpdf:
	$(SPHINXBUILD) -b latex $(ALLSPHINXOPTS) $(BUILDDIR)/latex
	@echo "Running LaTeX files through pdflatex..."
	$(MAKE) -C $(BUILDDIR)/latex all-pdf
	@echo "pdflatex finished; the PDF files are in $(BUILDDIR)/latex."

latexpdfja:
	$(SPHINXBUILD) -b latex $(ALLSPHINXOPTS) $(BUILDDIR)/latex
	@echo "Running LaTeX files through platex and dvipdfmx..."
	$(MAKE) -C $(BUILDDIR)/latex all-pdf-ja
	@echo "pdflatex finished; the PDF files are in $(BUILDDIR)/latex."

text:
	$(SPHINXBUILD) -b text $(ALLSPHINXOPTS) $(BUILDDIR)/text
	@echo
	@echo "Build finished. The text files are in $(BUILDDIR)/text."

man:
	$(SPHINXBUILD) -b man $(ALLSPHINXOPTS) $(BUILDDIR)/man
	@echo
	@echo "Build finished. The manual pages are in $(BUILDDIR)/man."

texinfo:
	$(SPHINXBUILD) -b texinfo $(ALLSPHINXOPTS) $(BUILDDIR)/texinfo
	@echo
	@echo "Build finished. The Texinfo files are in $(BUILDDIR)/texinfo."
	@echo "Run \`make' in that directory to run these through makeinfo" \
	      "(use \`make info' here to do that automatically)."

info:
	$(SPHINXBUILD) -b texinfo $(ALLSPHINXOPTS) $(BUILDDIR)/texinfo
	@echo "Running Texinfo files through makeinfo..."
	make -C $(BUILDDIR)/texinfo info
	@echo "makeinfo finished; the Info files are in $(BUILDDIR)/texinfo."

gettext:
	$(SPHINXBUILD) -b gettext $(I18NSPHINXOPTS) $(BUILDDIR)/locale
	@echo
	@echo "Build finished. The message catalogs are in $(BUILDDIR)/locale."

changes:
	$(SPHINXBUILD) -b changes $(ALLSPHINXOPTS) $(BUILDDIR)/changes
	@echo
	@echo "The overview file is in $(BUILDDIR)/changes."

linkcheck:
	$(SPHINXBUILD) -b linkcheck $(ALLSPHINXOPTS) $(BUILDDIR)/linkcheck
	@echo
	@echo "Link check complete; look for any errors in the above output " \
	      "or in $(BUILDDIR)/linkcheck/output.txt."

doctest:
	$(SPHINXBUILD) -b doctest $(ALLSPHINXOPTS) $(BUILDDIR)/doctest
	@echo "Testing of doctests in the sources finished, look at the " \
	      "results in $(BUILDDIR)/doctest/output.txt."

coverage:
	$(SPHINXBUILD) -b coverage $(ALLSPHINXOPTS) $(BUILDDIR)/coverage
	@echo "Testing of coverage in the sources finished, look at the " \
	      "results in $(BUILDDIR)/coverage/python.txt."

xml:
	$(SPHINXBUILD) -b xml $(ALLSPHINXOPTS) $(BUILDDIR)/xml
	@echo
	@echo "Build finished. The XML files are in $(BUILDDIR)/xml."

pseudoxml:
	$(SPHINXBUILD) -b pseudoxml $(ALLSPHINXOPTS) $(BUILDDIR)/pseudoxml
	@echo
	@echo "Build finished. The pseudo-XML files are in $(BUILDDIR)/pseudoxml."
```
