## Supplementary material for "ΔSCOPE: A new method to quantify 3D biological structures and identify differences in zebrafish forebrain development": An archived copy of the DeltaSCOPE code repository.: conf.py

```
#!/usr/bin/env python3
# -*- coding: utf-8 -*-
#
# deltascope documentation build configuration file, created by
# sphinx-quickstart on Thu Jan  4 13:33:50 2018.
#
# This file is execfile()d with the current directory set to its
# containing dir.
#
# Note that not all possible configuration values are present in this
# autogenerated file.
#
# All configuration values have a default; values that are commented out
# serve to show the default.

import deltascope
from mock import MagicMock
import sphinx_rtd_theme
import sys
import os
import shlex

sys.path.insert(0, os.path.abspath('../..'))
# sys.path.append('C:\Users\zfishlab\Code\deltascope')
# sys.path.append(os.path.join(os.path.dirname(__name__), '..'))
# sys.path.insert(0,'C:\\Users\\zfishlab\\Code\\deltascope\\deltascope')
# sys.path.append(os.path.abspath('../../deltascope'))


class Mock(MagicMock):
    @classmethod
    def __getattr__(cls, name):
        return MagicMock()


MOCK_MODULES = ['scipy', 'scipy.optimize', 'scipy.integrate', 'scipy.stats',
                'numpy', 'pandas', 'h5py',
                'matplotlib', 'matplotlib.pyplot',
                'sklearn', 'sklearn.preprocessing', 'sklearn.decomposition', 'sklearn.metrics', 'sklearn.ensemble',
                'skimage', 'skimage.filters', 'skimage.morphology']
sys.modules.update((mod_name, Mock()) for mod_name in MOCK_MODULES)


# If extensions (or modules to document with autodoc) are in another directory,
# add these directories to sys.path here. If the directory is relative to the
# documentation root, use os.path.abspath to make it absolute, like shown here.
#sys.path.insert(0, os.path.abspath('.'))

# -- General configuration ------------------------------------------------

# If your documentation needs a minimal Sphinx version, state it here.
#needs_sphinx = '1.0'

# Add any Sphinx extension module names here, as strings. They can be
# extensions coming with Sphinx (named 'sphinx.ext.*') or your custom
# ones.
extensions = [
    'sphinx.ext.autodoc',
    'sphinx.ext.todo',
    'sphinx.ext.coverage',
    'sphinx.ext.ifconfig',
    'sphinx.ext.viewcode',
    'sphinx.ext.imgmath',
    'sphinx.ext.intersphinx'
]

numfig = True

# Add any paths that contain templates here, relative to this directory.
templates_path = ['_templates']

# The suffix(es) of source filenames.
# You can specify multiple suffix as a list of string:
# source_suffix = ['.rst', '.md']
source_suffix = '.rst'

# The encoding of source files.
#source_encoding = 'utf-8-sig'

# The master toctree document.
master_doc = 'index'

# General information about the project.
project = 'deltascope'
copyright = '2018, Morgan Schwartz'
author = 'Morgan Schwartz'

# The version info for the project you're documenting, acts as replacement for
# |version| and |release|, also used in various other places throughout the
# built documents.
#
# The short X.Y version.
version = '1.0'
# The full version, including alpha/beta/rc tags.
release = '1.0.0'

# The language for content autogenerated by Sphinx. Refer to documentation
# for a list of supported languages.
#
# This is also used if you do content translation via gettext catalogs.
# Usually you set "language" from the command line for these cases.
language = None

# There are two options for replacing |today|: either, you set today to some
# non-false value, then it is used:
#today = ''
# Else, today_fmt is used as the format for a strftime call.
#today_fmt = '%B %d, %Y'

# List of patterns, relative to source directory, that match files and
# directories to ignore when looking for source files.
exclude_patterns = []

# The reST default role (used for this markup: `text`) to use for all
# documents.
#default_role = None

# If true, '()' will be appended to :func: etc. cross-reference text.
#add_function_parentheses = True

# If true, the current module name will be prepended to all description
# unit titles (such as .. function::).
#add_module_names = True

# If true, sectionauthor and moduleauthor directives will be shown in the
# output. They are ignored by default.
#show_authors = False

# The name of the Pygments (syntax highlighting) style to use.
pygments_style = 'sphinx'

# A list of ignored prefixes for module index sorting.
#modindex_common_prefix = []

# If true, keep warnings as "system message" paragraphs in the built documents.
#keep_warnings = False

# If true, `todo` and `todoList` produce output, else they produce nothing.
todo_include_todos = True

autodoc_member_order = 'bysource'


# -- Options for HTML output ----------------------------------------------

# The theme to use for HTML and HTML Help pages.  See the documentation for
# a list of builtin themes.
html_theme = 'sphinx_rtd_theme'
html_theme_path = [sphinx_rtd_theme.get_html_theme_path()]

# Theme options are theme-specific and customize the look and feel of a theme
# further.  For a list of options available for each theme, see the
# documentation.
#html_theme_options = {}

# Add any paths that contain custom themes here, relative to this directory.
#html_theme_path = []

# The name for this set of Sphinx documents.  If None, it defaults to
# "<project> v<release> documentation".
#html_title = None

# A shorter title for the navigation bar.  Default is the same as html_title.
#html_short_title = None

# The name of an image file (relative to this directory) to place at the top
# of the sidebar.
#html_logo = None

# The name of an image file (within the static path) to use as favicon of the
# docs.  This file should be a Windows icon file (.ico) being 16x16 or 32x32
# pixels large.
#html_favicon = None

# Add any paths that contain custom static files (such as style sheets) here,
# relative to this directory. They are copied after the builtin static files,
# so a file named "default.css" will overwrite the builtin "default.css".
html_static_path = ['_static']

# Add any extra paths that contain custom files (such as robots.txt or
# .htaccess) here, relative to this directory. These files are copied
# directly to the root of the documentation.
#html_extra_path = []

# If not '', a 'Last updated on:' timestamp is inserted at every page bottom,
# using the given strftime format.
#html_last_updated_fmt = '%b %d, %Y'

# If true, SmartyPants will be used to convert quotes and dashes to
# typographically correct entities.
#html_use_smartypants = True

# Custom sidebar templates, maps document names to template names.
#html_sidebars = {}

# Additional templates that should be rendered to pages, maps page names to
# template names.
#html_additional_pages = {}

# If false, no module index is generated.
#html_domain_indices = True

# If false, no index is generated.
#html_use_index = True

# If true, the index is split into individual pages for each letter.
#html_split_index = False

# If true, links to the reST sources are added to the pages.
#html_show_sourcelink = True

# If true, "Created using Sphinx" is shown in the HTML footer. Default is True.
#html_show_sphinx = True

# If true, "(C) Copyright ..." is shown in the HTML footer. Default is True.
#html_show_copyright = True

# If true, an OpenSearch description file will be output, and all pages will
# contain a <link> tag referring to it.  The value of this option must be the
# base URL from which the finished HTML is served.
#html_use_opensearch = ''

# This is the file name suffix for HTML files (e.g. ".xhtml").
#html_file_suffix = None

# Language to be used for generating the HTML full-text search index.
# Sphinx supports the following languages:
#   'da', 'de', 'en', 'es', 'fi', 'fr', 'h', 'it', 'ja'
#   'nl', 'no', 'pt', 'ro', 'r', 'sv', 'tr'
#html_search_language = 'en'

# A dictionary with options for the search language support, empty by default.
# Now only 'ja' uses this config value
#html_search_options = {'type': 'default'}

# The name of a javascript file (relative to the configuration directory) that
# implements a search results scorer. If empty, the default will be used.
#html_search_scorer = 'scorer.js'

# Output file base name for HTML help builder.
htmlhelp_basename = 'deltascopedoc'

# -- Options for LaTeX output ---------------------------------------------

latex_elements = {
    # The paper size ('letterpaper' or 'a4paper').
    # 'papersize': 'letterpaper',

    # The font size ('10pt', '11pt' or '12pt').
    # 'pointsize': '10pt',

    # Additional stuff for the LaTeX preamble.
    # 'preamble': '',

    # Latex figure (float) alignment
    # 'figure_align': 'htbp',
}

# Grouping the document tree into LaTeX files. List of tuples
# (source start file, target name, title,
#  author, documentclass [howto, manual, or own class]).
latex_documents = [
    (master_doc, 'deltascope.tex', 'deltascope Documentation',
     'Morgan Schwartz', 'manual'),
]

# The name of an image file (relative to this directory) to place at the top of
# the title page.
#latex_logo = None

# For "manual" documents, if this is true, then toplevel headings are parts,
# not chapters.
#latex_use_parts = False

# If true, show page references after internal links.
#latex_show_pagerefs = False

# If true, show URL addresses after external links.
#latex_show_urls = False

# Documents to append as an appendix to all manuals.
#latex_appendices = []

# If false, no module index is generated.
#latex_domain_indices = True


# -- Options for manual page output ---------------------------------------

# One entry per manual page. List of tuples
# (source start file, name, description, authors, manual section).
man_pages = [
    (master_doc, 'deltascope', 'deltascope Documentation',
     [author], 1)
]

# If true, show URL addresses after external links.
#man_show_urls = False


# -- Options for Texinfo output -------------------------------------------

# Grouping the document tree into Texinfo files. List of tuples
# (source start file, target name, title, author,
#  dir menu entry, description, category)
texinfo_documents = [
    (master_doc, 'deltascope', 'deltascope Documentation',
     author, 'deltascope', 'One line description of project.',
     'Miscellaneous'),
]

# Documents to append as an appendix to all manuals.
#texinfo_appendices = []

# If false, no module index is generated.
#texinfo_domain_indices = True

# How to display URL addresses: 'footnote', 'no', or 'inline'.
#texinfo_show_urls = 'footnote'

# If true, do not generate a @detailmenu in the "Top" node's menu.
#texinfo_no_detailmenu = False
```
