## Supplementary material for "ΔSCOPE: A new method to quantify 3D biological structures and identify differences in zebrafish forebrain development": An archived copy of the DeltaSCOPE code repository.: alignment-22hpf.html


In [1]:

```
import numpy as np
import pandas as pd
import matplotlib.pyplot as plt

import os
import re
from imp import reload
import h5py
import sys
```

In [2]:

```
import deltascope as cranium
import deltascope.alignment as ut
```

In [3]:

```
at = ".\\data\\22hpf\\AT\\Prob"
zrf = ".\\data\\22hpf\\Gfap\\Prob"
```

In [4]:

```
outdir = ".\\data\\22hpf\\Output-02-14-2019"
```

In [5]:

```
os.mkdir(outdir)
```

```
---------------------------------------------------------------------------
FileExistsError                           Traceback (most recent call last)
<ipython-input-5-97d3b4674911> in <module>
----> 1 os.mkdir(outdir)

FileExistsError: [WinError 183] Cannot create a file when that file already exists: '.\\data\\22hpf\\Output-09-15'
```

In [6]:

```
Dat = {}
for f in os.listdir(at):
    if 'h5' in f:
        num  = re.findall(r'\d+',f.split('.')[0])[-1]
        Dat[num] = os.path.join(at,f)
```

In [7]:

```
Dzrf = {}
for f in os.listdir(zrf):
    if 'h5' in f:
        num  = re.findall(r'\d+',f.split('.')[0])[-1]
        Dzrf[num] = os.path.join(zrf,f)
```

In [8]:

```
Dbat = {}
```

In [9]:

```
Dbzrf = {}
```

### Data Preprocessing¶

In [10]:

```
klist = Dat.keys()
```

In [11]:

```
param = {
    'gthresh':0.5,
    'scale':[1,1,1],
    'microns':[0.16,0.16,0.21],
    'mthresh':0.8,
    'radius':10,
    'comp_order':[0,2,1],
    'fit_dim':['x','z'],
    'deg':2
}
```

In [12]:

```
for k in klist:
    try:
        Dbat[k] = ut.preprocess(Dat[k],param)
        Dbzrf[k] = ut.preprocess(Dzrf[k],param,pca=Dbat[k].pcamed,mm=Dbat[k].mm,vertex=Dbat[k].vertex)
        print(k)
    except:
        print(k,'failed')
        e = sys.exc_info()
        print(e)
```

```
C:\Users\zfishlab\AppData\Local\Continuum\anaconda3\envs\test\lib\site-packages\skimage\util\dtype.py:141: UserWarning: Possible precision loss when converting from float32 to uint8
  .format(dtypeobj_in, dtypeobj_out))
```

```
10
```

```
C:\Users\zfishlab\AppData\Local\Continuum\anaconda3\envs\test\lib\site-packages\skimage\util\dtype.py:141: UserWarning: Possible precision loss when converting from float32 to uint8
  .format(dtypeobj_in, dtypeobj_out))
```

```
12
```

```
C:\Users\zfishlab\AppData\Local\Continuum\anaconda3\envs\test\lib\site-packages\skimage\util\dtype.py:141: UserWarning: Possible precision loss when converting from float32 to uint8
  .format(dtypeobj_in, dtypeobj_out))
```

```
13
```

```
C:\Users\zfishlab\AppData\Local\Continuum\anaconda3\envs\test\lib\site-packages\skimage\util\dtype.py:141: UserWarning: Possible precision loss when converting from float32 to uint8
  .format(dtypeobj_in, dtypeobj_out))
```

```
14
```

```
C:\Users\zfishlab\AppData\Local\Continuum\anaconda3\envs\test\lib\site-packages\skimage\util\dtype.py:141: UserWarning: Possible precision loss when converting from float32 to uint8
  .format(dtypeobj_in, dtypeobj_out))
```

```
1
```

```
C:\Users\zfishlab\AppData\Local\Continuum\anaconda3\envs\test\lib\site-packages\skimage\util\dtype.py:141: UserWarning: Possible precision loss when converting from float32 to uint8
  .format(dtypeobj_in, dtypeobj_out))
```

```
2
```

```
C:\Users\zfishlab\AppData\Local\Continuum\anaconda3\envs\test\lib\site-packages\skimage\util\dtype.py:141: UserWarning: Possible precision loss when converting from float32 to uint8
  .format(dtypeobj_in, dtypeobj_out))
```

```
3
```

```
C:\Users\zfishlab\AppData\Local\Continuum\anaconda3\envs\test\lib\site-packages\skimage\util\dtype.py:141: UserWarning: Possible precision loss when converting from float32 to uint8
  .format(dtypeobj_in, dtypeobj_out))
```

```
4
```

```
C:\Users\zfishlab\AppData\Local\Continuum\anaconda3\envs\test\lib\site-packages\skimage\util\dtype.py:141: UserWarning: Possible precision loss when converting from float32 to uint8
  .format(dtypeobj_in, dtypeobj_out))
```

```
5
```

```
C:\Users\zfishlab\AppData\Local\Continuum\anaconda3\envs\test\lib\site-packages\skimage\util\dtype.py:141: UserWarning: Possible precision loss when converting from float32 to uint8
  .format(dtypeobj_in, dtypeobj_out))
```

```
6
```

```
C:\Users\zfishlab\AppData\Local\Continuum\anaconda3\envs\test\lib\site-packages\skimage\util\dtype.py:141: UserWarning: Possible precision loss when converting from float32 to uint8
  .format(dtypeobj_in, dtypeobj_out))
```

```
7
```

```
C:\Users\zfishlab\AppData\Local\Continuum\anaconda3\envs\test\lib\site-packages\skimage\util\dtype.py:141: UserWarning: Possible precision loss when converting from float32 to uint8
  .format(dtypeobj_in, dtypeobj_out))
```

```
8
```

```
C:\Users\zfishlab\AppData\Local\Continuum\anaconda3\envs\test\lib\site-packages\skimage\util\dtype.py:141: UserWarning: Possible precision loss when converting from float32 to uint8
  .format(dtypeobj_in, dtypeobj_out))
```

```
9
```

### Functions¶

In [13]:

```
def start(k):
    return(ut.start(k,Dbat,[Dbzrf],im=True))
def save_at(k,df):
    ut.save_at(k,df,outdir,'22hpf')
def save_both(k,dfa,dfb):
    ut.save_both(k,dfa,dfb,outdir,'22hpf')
```

In [14]:

```
def fit_model(axi,df,mm=None):
    if mm == None:
        mm = np.polyfit(df.x,df.z,2)
    p = np.poly1d(mm)
    xrange = np.arange(np.min(df.x),np.max(df.x))
    axi.plot(xrange,p(xrange),c='m')
    return(mm)
```

In [15]:

```
model = pd.DataFrame({'a':[],'b':[],'c':[]})
def save_model(k,mm,model):
    row = pd.Series({'a':mm[0],'b':mm[1],'c':mm[2]},name=k)
    model = model.append(row)
    return(model)
```

In [16]:

```
def pick_pts(x1,z1,vx,vz,x2,z2):
    pts = pd.DataFrame({'x':[x1,vx,x2],'z':[z1,vz,z2]})
    return(pts)
```

In [17]:

```
klist
```

Out[17]:

```
dict_keys(['10', '12', '13', '14', '1', '2', '3', '4', '5', '6', '7', '8', '9'])
```

In [18]:

```
Dbzrf.keys()
```

Out[18]:

```
dict_keys(['10', '12', '13', '14', '1', '2', '3', '4', '5', '6', '7', '8', '9'])
```

# 14¶

In [19]:

```
Dbat['14']
```

Out[19]:

```
<deltascope.brain at 0x244f0a9fa90>
```

In [20]:

```
k,df,Ldf,ax = start('14')
```

In [21]:

```
df1,Ldf1,pts,ax = ut.check_pts(df,Ldf,'z')
```

In [22]:

```
pts.iloc[0].x = -40
pts.iloc[0].z = 26
pts.iloc[1].x = 45
pts.iloc[1].z = 22
ax[0,1].scatter(pts.x,pts.z,c='y')
pts
```

Out[22]:

|  | x | z |
| --- | --- | --- |
| 0 | -40.0 | 26.0 |
| 1 | 45.0 | 22.0 |

In [23]:

```
df2,Ldf2,ax = ut.revise_pts(df,Ldf,'z',pts=pts)
```

In [24]:

```
mm = fit_model(ax[1,1],df2)
```

In [25]:

```
pts = pick_pts(-40,26,0,-5,45,22)
ax[1,1].scatter(pts.x,pts.z,c='m')
```

Out[25]:

```
<matplotlib.collections.PathCollection at 0x2451de638d0>
```

In [26]:

```
pts = pick_pts(-43,26,0,-5,45,22)
```

In [27]:

```
df3,Ldf3,mm,ax = ut.ch_vertex(df2,Ldf2,pts=pts)
```

In [28]:

```
model = save_model(k,mm,model)
```

In [29]:

```
save_both(k,df3,Ldf3[0])
```

```
Write to .\data\22hpf\Output-09-15\AT_14_22hpf.psi complete
Write to .\data\22hpf\Output-09-15\ZRF_14_22hpf.psi complete
```

# 8¶

In [30]:

```
k,df,Ldf,ax = start('8')
```

In [31]:

```
mm = fit_model(ax[1,1],df)
```

In [32]:

```
model = save_model(k,mm,model)
```

In [33]:

```
save_both(k,df,Ldf[0])
```

```
Write to .\data\22hpf\Output-09-15\AT_8_22hpf.psi complete
Write to .\data\22hpf\Output-09-15\ZRF_8_22hpf.psi complete
```

# 6¶

In [34]:

```
k,df,Ldf,ax = start('6')
```

In [35]:

```
df1,Ldf1 = ut.zyswitch(df,Ldf)
ax = ut.make_graph([df1]+Ldf1)
```

In [36]:

```
df2,Ldf2 = ut.flip(df1,Ldf1)
ax = ut.make_graph([df2]+Ldf2)
```

In [37]:

```
mm = fit_model(ax[0,1],df2)
```

In [38]:

```
pts = pick_pts(-42,4,0,-15,40,5)
ax[0,1].scatter(pts.x,pts.z,c='m',s=50)
```

Out[38]:

```
<matplotlib.collections.PathCollection at 0x2451b5da400>
```

In [39]:

```
df3,Ldf3,mm,ax = ut.ch_vertex(df2,Ldf2,pts=pts)
```

In [40]:

```
model = save_model(k,mm,model)
save_both(k,df3,Ldf3[0])
```

```
Write to .\data\22hpf\Output-09-15\AT_6_22hpf.psi complete
Write to .\data\22hpf\Output-09-15\ZRF_6_22hpf.psi complete
```

# 10¶

In [41]:

```
k,df,Ldf,ax = start('10')
```

In [42]:

```
pts = pick_pts(-32,18,0,-12,40,22)
ax[0,1].scatter(pts.x,pts.z,c='m',s=50)
```

Out[42]:

```
<matplotlib.collections.PathCollection at 0x2451ded0198>
```

In [43]:

```
df1,Ldf1,mm,ax = ut.ch_vertex(df,Ldf,pts=pts)
```

In [44]:

```
model = save_model(k,mm,model)
save_both(k,df1,Ldf1[0])
```

```
Write to .\data\22hpf\Output-09-15\AT_10_22hpf.psi complete
Write to .\data\22hpf\Output-09-15\ZRF_10_22hpf.psi complete
```

# 7¶

In [45]:

```
k,df,Ldf,ax = start('7')
```

In [46]:

```
mm = fit_model(ax[0,1],df)
```

In [47]:

```
model = save_model(k,mm,model)
```

In [48]:

```
save_both(k,df,Ldf[0])
```

```
Write to .\data\22hpf\Output-09-15\AT_7_22hpf.psi complete
Write to .\data\22hpf\Output-09-15\ZRF_7_22hpf.psi complete
```

# 1¶

In [49]:

```
k,df,Ldf,ax = start('1')
```

In [50]:

```
mm = fit_model(ax[0,1],df)
```

In [51]:

```
pts = pick_pts(-48,15,0,-3,50,15)
ax[0,1].scatter(pts.x,pts.z,c='m',s=50)
```

Out[51]:

```
<matplotlib.collections.PathCollection at 0x2451ed3e748>
```

In [52]:

```
df1,Ldf1,mm,ax = ut.ch_vertex(df,Ldf,pts=pts)
```

In [53]:

```
model = save_model(k,mm,model)
```

In [54]:

```
save_both(k,df1,Ldf1[0])
```

```
Write to .\data\22hpf\Output-09-15\AT_1_22hpf.psi complete
Write to .\data\22hpf\Output-09-15\ZRF_1_22hpf.psi complete
```

# 13¶

In [55]:

```
k,df,Ldf,ax = start('13')
```

In [56]:

```
df1,Ldf1,pts,ax = ut.check_pts(df,Ldf,'z')
```

In [57]:

```
pts.iloc[0].x = -45
pts.iloc[0].z = 17
pts.iloc[1].x = 42
pts.iloc[1].z = 14
ax[0,1].scatter(pts.x,pts.z,c='y')
pts
```

Out[57]:

|  | x | z |
| --- | --- | --- |
| 0 | -45.0 | 17.0 |
| 1 | 42.0 | 14.0 |

In [58]:

```
df2,Ldf2,ax = ut.revise_pts(df,Ldf,'z',pts=pts)
```

In [59]:

```
df3,Ldf3,mm,ax = ut.ch_vertex(df2,Ldf2)
```

In [60]:

```
pts = pick_pts(-39,16,0,-6,47,16)
ax[0,1].scatter(pts.x,pts.z,c='m',s=50)
```

Out[60]:

```
<matplotlib.collections.PathCollection at 0x2451f730748>
```

In [61]:

```
df4,Ldf4,mm,ax = ut.ch_vertex(df3,Ldf3,pts=pts)
```

In [62]:

```
model = save_model(k,mm,model)
```

In [63]:

```
save_both(k,df4,Ldf4[0])
```

```
Write to .\data\22hpf\Output-09-15\AT_13_22hpf.psi complete
Write to .\data\22hpf\Output-09-15\ZRF_13_22hpf.psi complete
```

# 2¶

In [64]:

```
k,df,Ldf,ax = start('2')
```

In [65]:

```
mm = fit_model(ax[0,1],df)
```

In [66]:

```
model = save_model(k,mm,model)
save_both(k,df,Ldf[0])
```

```
Write to .\data\22hpf\Output-09-15\AT_2_22hpf.psi complete
Write to .\data\22hpf\Output-09-15\ZRF_2_22hpf.psi complete
```

# 12¶

In [67]:

```
k,df,Ldf,ax = start('12')
```

In [68]:

```
mm = fit_model(ax[0,1],df)
```

In [69]:

```
pts = pick_pts(-36,14,0,-11,43,16)
ax[0,1].scatter(pts.x,pts.z,c='m',s=50)
```

Out[69]:

```
<matplotlib.collections.PathCollection at 0x2451fb0a898>
```

In [70]:

```
df1,Ldf1,mm,ax = ut.ch_vertex(df,Ldf,pts=pts)
```

In [71]:

```
model = save_model(k,mm,model)
save_both(k,df1,Ldf1[0])
```

```
Write to .\data\22hpf\Output-09-15\AT_12_22hpf.psi complete
Write to .\data\22hpf\Output-09-15\ZRF_12_22hpf.psi complete
```

# 4¶

In [72]:

```
k,df,Ldf,ax = start('4')
```

In [73]:

```
pts = pick_pts(-38,28,0,-13,40,30)
ax[0,1].scatter(pts.x,pts.z,c='m',s=50)
```

Out[73]:

```
<matplotlib.collections.PathCollection at 0x2451ee190f0>
```

In [74]:

```
df1,Ldf1,mm,ax = ut.ch_vertex(df,Ldf,pts=pts)
```

In [75]:

```
model = save_model(k,mm,model)
save_both(k,df1,Ldf1[0])
```

```
Write to .\data\22hpf\Output-09-15\AT_4_22hpf.psi complete
Write to .\data\22hpf\Output-09-15\ZRF_4_22hpf.psi complete
```

# 9¶

In [76]:

```
k,df,Ldf,ax = start('9')
```

In [77]:

```
mm = fit_model(ax[0,1],df)
```

In [78]:

```
model = save_model(k,mm,model)
```

In [79]:

```
save_both(k,df,Ldf[0])
```

```
Write to .\data\22hpf\Output-09-15\AT_9_22hpf.psi complete
Write to .\data\22hpf\Output-09-15\ZRF_9_22hpf.psi complete
```

# 5¶

In [80]:

```
k,df,Ldf,ax = start('5')
```

In [81]:

```
mm = fit_model(ax[0,1],df)
```

In [82]:

```
model = save_model(k,mm,model)
save_both(k,df,Ldf[0])
```

```
Write to .\data\22hpf\Output-09-15\AT_5_22hpf.psi complete
Write to .\data\22hpf\Output-09-15\ZRF_5_22hpf.psi complete
```

# 3¶

In [83]:

```
k,df,Ldf,ax = start('3')
```

In [84]:

```
mm = fit_model(ax[0,1],df)
```

In [85]:

```
pts = pick_pts(-36,22,0,-5,38,20)
ax[0,1].scatter(pts.x,pts.z,c='m',s=50)
```

Out[85]:

```
<matplotlib.collections.PathCollection at 0x2457e9b7e48>
```

In [86]:

```
df1,Ldf1,mm,ax = ut.ch_vertex(df,Ldf,pts=pts)
```

In [87]:

```
model = save_model(k,mm,model)
save_both(k,df1,Ldf1[0])
```

```
Write to .\data\22hpf\Output-09-15\AT_3_22hpf.psi complete
Write to .\data\22hpf\Output-09-15\ZRF_3_22hpf.psi complete
```

In [88]:

```
model
```

Out[88]:

|  | a | b | c |
| --- | --- | --- | --- |
| 14 | 0.015011 | 7.442408e-17 | -5.864644e-15 |
| 8 | 0.005242 | -1.504477e-16 | 2.311078e-14 |
| 6 | 0.011614 | -8.318195e-17 | -3.975859e-15 |
| 10 | 0.024826 | 1.920145e-17 | -1.037776e-14 |
| 7 | 0.005284 | -1.208995e-16 | 4.298055e-15 |
| 1 | 0.007500 | 9.491636e-17 | 2.051180e-15 |
| 13 | 0.012002 | 1.511626e-16 | 3.079346e-15 |
| 2 | 0.007366 | -7.175444e-17 | 7.848694e-16 |
| 12 | 0.016739 | 2.391703e-17 | -1.238302e-14 |
| 4 | 0.027615 | -6.906357e-17 | -8.116310e-15 |
| 9 | 0.009719 | -7.102912e-16 | -1.980932e-15 |
| 5 | 0.007165 | 9.423889e-16 | 4.467195e-14 |
| 3 | 0.019026 | 4.753098e-17 | 1.243870e-14 |

In [89]:

```
model.to_csv(os.path.join(outdir,'model.csv'))
```

In [90]:

```
outdir
```

Out[90]:

```
'.\\data\\22hpf\\Output-09-15'
```
