## Supplementary material for "ΔSCOPE: A new method to quantify 3D biological structures and identify differences in zebrafish forebrain development": An archived copy of the DeltaSCOPE code repository.: alignment-24hpf.html


In [2]:

```
import numpy as np
import pandas as pd
import matplotlib.pyplot as plt

import os
import re
from imp import reload
import h5py
import sys
```

In [3]:

```
import deltascope as cranium
import deltascope.alignment as ut
```

In [4]:

```
at = ".\\data\\24hpf\\AT\\Prob"
gfap = ".\\data\\24hpf\\Gfap\\Prob"
```

In [5]:

```
outdir = ".\\data\\24hpf\\Output-02-15-2019"
```

In [6]:

```
os.mkdir(outdir)
```

In [7]:

```
Dat = {}
for f in os.listdir(at):
    if 'h5' in f:
        num  = re.findall(r'\d+',f.split('.')[0])[-1]
        Dat[num] = os.path.join(at,f)
```

In [8]:

```
Dzrf = {}
for f in os.listdir(gfap):
    if 'h5' in f:
        num  = re.findall(r'\d+',f.split('.')[0])[-1]
        Dzrf[num] = os.path.join(gfap,f)
```

In [9]:

```
Dat.keys()
```

Out[9]:

```
dict_keys(['10', '11', '12', '17', '19', '1', '21', '23', '24', '25', '26', '27', '28', '3', '4', '5', '6', '7', '8', '9'])
```

In [10]:

```
Dbat = {}
Dbzrf={}
```

### Data Preprocessing¶

In [11]:

```
klist = Dat.keys()
```

In [12]:

```
param = {
    'gthresh':0.5,
    'scale':[1,1,1],
    'microns':[0.16,0.16,0.21],
    'mthresh':0.8,
    'radius':10,
    'comp_order':[0,2,1],
    'fit_dim':['x','z'],
    'deg':2
}
```

In [13]:

```
for k in klist:
    try:
        Dbat[k] = ut.preprocess(Dat[k],param)
        Dbzrf[k] = ut.preprocess(Dzrf[k],param,pca=Dbat[k].pcamed,mm=Dbat[k].mm,vertex=Dbat[k].vertex)
        print(k)
    except:
        print(k,'failed')
        e = sys.exc_info()
        print(e)
```

```
C:\Users\zfishlab\AppData\Local\Continuum\anaconda3\envs\test\lib\site-packages\skimage\util\dtype.py:141: UserWarning: Possible precision loss when converting from float32 to uint8
  .format(dtypeobj_in, dtypeobj_out))
```

```
10
```

```
C:\Users\zfishlab\AppData\Local\Continuum\anaconda3\envs\test\lib\site-packages\skimage\util\dtype.py:141: UserWarning: Possible precision loss when converting from float32 to uint8
  .format(dtypeobj_in, dtypeobj_out))
```

```
11
```

```
C:\Users\zfishlab\AppData\Local\Continuum\anaconda3\envs\test\lib\site-packages\skimage\util\dtype.py:141: UserWarning: Possible precision loss when converting from float32 to uint8
  .format(dtypeobj_in, dtypeobj_out))
```

```
12
```

```
C:\Users\zfishlab\AppData\Local\Continuum\anaconda3\envs\test\lib\site-packages\skimage\util\dtype.py:141: UserWarning: Possible precision loss when converting from float32 to uint8
  .format(dtypeobj_in, dtypeobj_out))
```

```
17 failed
(<class 'ValueError'>, ValueError('Found array with 0 sample(s) (shape=(0, 3)) while a minimum of 1 is required.'), <traceback object at 0x000002072ED89048>)
```

```
C:\Users\zfishlab\AppData\Local\Continuum\anaconda3\envs\test\lib\site-packages\skimage\util\dtype.py:141: UserWarning: Possible precision loss when converting from float32 to uint8
  .format(dtypeobj_in, dtypeobj_out))
```

```
19
```

```
C:\Users\zfishlab\AppData\Local\Continuum\anaconda3\envs\test\lib\site-packages\skimage\util\dtype.py:141: UserWarning: Possible precision loss when converting from float32 to uint8
  .format(dtypeobj_in, dtypeobj_out))
```

```
1
```

```
C:\Users\zfishlab\AppData\Local\Continuum\anaconda3\envs\test\lib\site-packages\skimage\util\dtype.py:141: UserWarning: Possible precision loss when converting from float32 to uint8
  .format(dtypeobj_in, dtypeobj_out))
```

```
21
```

```
C:\Users\zfishlab\AppData\Local\Continuum\anaconda3\envs\test\lib\site-packages\skimage\util\dtype.py:141: UserWarning: Possible precision loss when converting from float32 to uint8
  .format(dtypeobj_in, dtypeobj_out))
```

```
23
```

```
C:\Users\zfishlab\AppData\Local\Continuum\anaconda3\envs\test\lib\site-packages\skimage\util\dtype.py:141: UserWarning: Possible precision loss when converting from float32 to uint8
  .format(dtypeobj_in, dtypeobj_out))
```

```
24 failed
(<class 'ValueError'>, ValueError('Found array with 0 sample(s) (shape=(0, 3)) while a minimum of 1 is required.'), <traceback object at 0x000002072EDB2B08>)
```

```
C:\Users\zfishlab\AppData\Local\Continuum\anaconda3\envs\test\lib\site-packages\skimage\util\dtype.py:141: UserWarning: Possible precision loss when converting from float32 to uint8
  .format(dtypeobj_in, dtypeobj_out))
```

```
25
```

```
C:\Users\zfishlab\AppData\Local\Continuum\anaconda3\envs\test\lib\site-packages\skimage\util\dtype.py:141: UserWarning: Possible precision loss when converting from float32 to uint8
  .format(dtypeobj_in, dtypeobj_out))
```

```
26
```

```
C:\Users\zfishlab\AppData\Local\Continuum\anaconda3\envs\test\lib\site-packages\skimage\util\dtype.py:141: UserWarning: Possible precision loss when converting from float32 to uint8
  .format(dtypeobj_in, dtypeobj_out))
```

```
27
```

```
C:\Users\zfishlab\AppData\Local\Continuum\anaconda3\envs\test\lib\site-packages\skimage\util\dtype.py:141: UserWarning: Possible precision loss when converting from float32 to uint8
  .format(dtypeobj_in, dtypeobj_out))
```

```
28
```

```
C:\Users\zfishlab\AppData\Local\Continuum\anaconda3\envs\test\lib\site-packages\skimage\util\dtype.py:141: UserWarning: Possible precision loss when converting from float32 to uint8
  .format(dtypeobj_in, dtypeobj_out))
```

```
3
```

```
C:\Users\zfishlab\AppData\Local\Continuum\anaconda3\envs\test\lib\site-packages\skimage\util\dtype.py:141: UserWarning: Possible precision loss when converting from float32 to uint8
  .format(dtypeobj_in, dtypeobj_out))
```

```
4
```

```
C:\Users\zfishlab\AppData\Local\Continuum\anaconda3\envs\test\lib\site-packages\skimage\util\dtype.py:141: UserWarning: Possible precision loss when converting from float32 to uint8
  .format(dtypeobj_in, dtypeobj_out))
```

```
5
```

```
C:\Users\zfishlab\AppData\Local\Continuum\anaconda3\envs\test\lib\site-packages\skimage\util\dtype.py:141: UserWarning: Possible precision loss when converting from float32 to uint8
  .format(dtypeobj_in, dtypeobj_out))
```

```
6
```

```
C:\Users\zfishlab\AppData\Local\Continuum\anaconda3\envs\test\lib\site-packages\skimage\util\dtype.py:141: UserWarning: Possible precision loss when converting from float32 to uint8
  .format(dtypeobj_in, dtypeobj_out))
```

```
7
```

```
C:\Users\zfishlab\AppData\Local\Continuum\anaconda3\envs\test\lib\site-packages\skimage\util\dtype.py:141: UserWarning: Possible precision loss when converting from float32 to uint8
  .format(dtypeobj_in, dtypeobj_out))
```

```
8
```

```
C:\Users\zfishlab\AppData\Local\Continuum\anaconda3\envs\test\lib\site-packages\skimage\util\dtype.py:141: UserWarning: Possible precision loss when converting from float32 to uint8
  .format(dtypeobj_in, dtypeobj_out))
```

```
9 failed
(<class 'ValueError'>, ValueError('Found array with 0 sample(s) (shape=(0, 3)) while a minimum of 1 is required.'), <traceback object at 0x000002072F1CA448>)
```

In [14]:

```
def start(k):
    return(ut.start(k,Dbat,[Dbzrf],im=False))
def save_at(k,df):
    ut.save_at(k,df,outdir,'24hpf')
def save_both(k,dfa,dfb):
    ut.save_both(k,dfa,dfb,outdir,'24hpf')
```

In [15]:

```
def fit_model(axi,df,mm=None):
    if mm == None:
        mm = np.polyfit(df.x,df.z,2)
    p = np.poly1d(mm)
    xrange = np.arange(np.min(df.x),np.max(df.x))
    axi.plot(xrange,p(xrange),c='m')
    return(mm)
```

In [24]:

```
model = pd.DataFrame({'a':[],'b':[],'c':[]})
def save_model(k,mm,model):
    row = pd.Series({'a':mm[0],'b':mm[1],'c':mm[2]},name=k)
    model = model.append(row)
    return(model)
```

In [35]:

```
def pick_pts(x1,z1,vx,vz,x2,z2):
    pts = pd.DataFrame({'x':[x1,vx,x2],'z':[z1,vz,z2]})
    return(pts)
```

In [25]:

```
klist
```

Out[25]:

```
dict_keys(['10', '11', '12', '17', '19', '1', '21', '23', '24', '25', '26', '27', '28', '3', '4', '5', '6', '7', '8', '9'])
```

# 7¶

In [26]:

```
k,df,Ldf,ax = start('7')
```

In [27]:

```
df1,Ldf1,pts,ax = ut.check_pts(df,Ldf,'z')
```

In [28]:

```
pts.iloc[1].x = 36
pts.iloc[1].z = 22
ax[0,1].scatter(pts.x,pts.z,c='y')
```

Out[28]:

```
<matplotlib.collections.PathCollection at 0x20758c1dcc0>
```

In [29]:

```
df2,Ldf2,ax = ut.revise_pts(df,Ldf,'z',pts=pts)
```

In [30]:

```
df3,Ldf3,p,ax = ut.check_yz(df2,Ldf2)
```

In [31]:

```
df4,Ldf4,mm,ax = ut.ch_vertex(df3,Ldf3)
```

In [36]:

```
pts = pick_pts(-42,27,0,-4,40,27)
ax = ut.make_graph([df4]+Ldf4)
ax[0,1].scatter(pts.x,pts.z,c='m',s=40)
```

Out[36]:

```
<matplotlib.collections.PathCollection at 0x20758a5cf60>
```

In [37]:

```
df5,Ldf5,mm,ax = ut.ch_vertex(df4,Ldf4,pts=pts)
```

In [38]:

```
save_both(k,df5,Ldf5[0])
model = save_model(k,mm,model)
```

```
Write to .\data\24hpf\Output-02-15-2019\AT_7_24hpf.psi complete
Write to .\data\24hpf\Output-02-15-2019\ZRF_7_24hpf.psi complete
```

# 26¶

In [39]:

```
k,df,Ldf,ax = start('26')
```

In [40]:

```
df1,Ldf1,pts,ax = ut.check_pts(df,Ldf,'z')
```

In [41]:

```
pts.iloc[1].x = 18
pts.iloc[1].z = 2
ax[0,1].scatter(pts.x,pts.z,c='y')
pts
```

Out[41]:

|  | x | z |
| --- | --- | --- |
| 0 | -52.087549 | 41.197433 |
| 1 | 18.000000 | 2.000000 |

In [42]:

```
df2,Ldf2,ax = ut.revise_pts(df,Ldf,'z',pts=pts)
```

In [43]:

```
df3,Ldf3,pts,ax = ut.check_pts(df2,Ldf2,'y')
```

In [44]:

```
pts.iloc[1].x = 16
pts.iloc[1].y = -1
pts.iloc[0].y = 18
ax[0,0].scatter(pts.x,pts.y,c='y')
pts
```

Out[44]:

|  | x | y |
| --- | --- | --- |
| 0 | -65.223682 | 18.0 |
| 1 | 16.000000 | -1.0 |

In [45]:

```
df4,Ldf4,ax = ut.revise_pts(df2,Ldf2,'y',pts=pts)
```

In [46]:

```
df5,Ldf5,p,ax = ut.check_yz(df4,Ldf4)
```

In [47]:

```
p2 = np.poly1d([2/5,0])
xrange = np.arange(-20,20)
p[0,2].plot(xrange,p2(xrange),c='c')
```

Out[47]:

```
[<matplotlib.lines.Line2D at 0x20758522668>]
```

In [48]:

```
df6,Ldf6,ax,p = ut.check_yz(df4,Ldf4,mm=[2/5,0])
```

In [49]:

```
df7,Ldf7,mm,ax = ut.ch_vertex(df6,Ldf6)
```

In [50]:

```
pts = pick_pts(-40,20,0,-8,40,20)
ax[0,1].scatter(pts.x,pts.z,c='m',s=40)
```

Out[50]:

```
<matplotlib.collections.PathCollection at 0x20806cba940>
```

In [51]:

```
df8,Ldf8,mm,ax = ut.ch_vertex(df7,Ldf7,pts=pts)
```

In [52]:

```
save_both(k,df8,Ldf8[0])
model = save_model(k,mm,model)
```

```
Write to .\data\24hpf\Output-02-15-2019\AT_26_24hpf.psi complete
Write to .\data\24hpf\Output-02-15-2019\ZRF_26_24hpf.psi complete
```

In [53]:

```
model = model.iloc[-2:]
```

In [54]:

```
model
```

Out[54]:

|  | a | b | c |
| --- | --- | --- | --- |
| 7 | 0.018452 | 1.914462e-17 | -5.692721e-21 |
| 26 | 0.017500 | 2.761276e-17 | 0.000000e+00 |

# 9¶

In [56]:

```
Dbat['9'] = ut.preprocess(os.path.join(at,'AT_24hpf_9_Probabilities.h5'),param,stop='df_thresh')
Dbzrf['9'] = ut.preprocess(os.path.join(gfap,'Gfap_24hpf_9_Probabilities.h5'),param,stop='df_thresh')
```

In [57]:

```
k = '9'
df = Dbat[k].df_thresh
Ldf = [Dbzrf[k].df_thresh]
```

In [58]:

```
ax = ut.make_graph([df]+Ldf)
```

In [59]:

```
df1,Ldf1,mm,ax = ut.ch_vertex(df,Ldf)
```

In [60]:

```
save_both(k,df1,Ldf1[0])
model = save_model(k,mm,model)
model
```

```
Write to .\data\24hpf\Output-02-15-2019\AT_9_24hpf.psi complete
Write to .\data\24hpf\Output-02-15-2019\ZRF_9_24hpf.psi complete
```

Out[60]:

|  | a | b | c |
| --- | --- | --- | --- |
| 7 | 0.018452 | 1.914462e-17 | -5.692721e-21 |
| 26 | 0.017500 | 2.761276e-17 | 0.000000e+00 |
| 9 | 0.004727 | 2.061185e-16 | 6.484843e-15 |

# 5¶

In [61]:

```
k,df,Ldf,ax = start('5')
```

In [62]:

```
df1,Ldf1,pts,ax = ut.check_pts(df,Ldf,'z')
```

In [63]:

```
df2,Ldf2,mm,ax = ut.ch_vertex(df1,Ldf1)
```

In [64]:

```
pts = pick_pts(-55,22,0,-8,58,20)
ax[0,1].scatter(pts.x,pts.z,s=50,c='m')
```

Out[64]:

```
<matplotlib.collections.PathCollection at 0x20806e3c4a8>
```

In [65]:

```
df3,Ldf3,mm,ax = ut.ch_vertex(df2,Ldf2,pts=pts)
```

In [66]:

```
save_both(k,df3,Ldf3[0])
model = save_model(k,mm,model)
model
```

```
Write to .\data\24hpf\Output-02-15-2019\AT_5_24hpf.psi complete
Write to .\data\24hpf\Output-02-15-2019\ZRF_5_24hpf.psi complete
```

Out[66]:

|  | a | b | c |
| --- | --- | --- | --- |
| 7 | 0.018452 | 1.914462e-17 | -5.692721e-21 |
| 26 | 0.017500 | 2.761276e-17 | 0.000000e+00 |
| 9 | 0.004727 | 2.061185e-16 | 6.484843e-15 |
| 5 | 0.009099 | 5.287413e-17 | -1.423254e-14 |

# 1¶

In [67]:

```
k = '1'
df = Dbat[k].df_thresh
Ldf = [Dbzrf[k].df_thresh]
```

In [68]:

```
ax = ut.make_graph([df]+Ldf)
```

In [69]:

```
df1,Ldf1,ax,p = ut.check_yz(df,Ldf)
```

In [70]:

```
df2,Ldf2,mm,ax = ut.ch_vertex(df1,Ldf1)
```

In [71]:

```
pts = pick_pts(-55,20,-6,-20,35,20)
ax[0,1].scatter(pts.x,pts.z,c='m',s=50)
```

Out[71]:

```
<matplotlib.collections.PathCollection at 0x2072fcb6e10>
```

In [72]:

```
df3,Ldf3,mm,ax = ut.ch_vertex(df2,Ldf2,pts=pts)
```

In [73]:

```
save_both(k,df3,Ldf3[0])
model = save_model(k,mm,model)
model
```

```
Write to .\data\24hpf\Output-02-15-2019\AT_1_24hpf.psi complete
Write to .\data\24hpf\Output-02-15-2019\ZRF_1_24hpf.psi complete
```

Out[73]:

|  | a | b | c |
| --- | --- | --- | --- |
| 7 | 0.018452 | 1.914462e-17 | -5.692721e-21 |
| 26 | 0.017500 | 2.761276e-17 | 0.000000e+00 |
| 9 | 0.004727 | 2.061185e-16 | 6.484843e-15 |
| 5 | 0.009099 | 5.287413e-17 | -1.423254e-14 |
| 1 | 0.019910 | -4.560282e-16 | 1.231414e-14 |

# 21¶

In [74]:

```
k,df,Ldf,ax = start('21')
```

In [75]:

```
df1,Ldf1,ax,p = ut.check_yz(df,Ldf)
```

In [76]:

```
mm = [3/5,0]
p = np.poly1d(mm)
xrange = np.arange(-10,30)
ax[0,2].plot(xrange,p(xrange),c='y')
```

Out[76]:

```
[<matplotlib.lines.Line2D at 0x208084a22e8>]
```

In [77]:

```
df2,Ldf2,ax,p = ut.check_yz(df,Ldf,mm=mm)
```

In [78]:

```
df3,Ldf3,mm,ax = ut.ch_vertex(df2,Ldf2)
```

In [79]:

```
pts = pick_pts(-48,24,0,-6,45,20)
ax[0,1].scatter(pts.x,pts.z,c='m',s=50)
```

Out[79]:

```
<matplotlib.collections.PathCollection at 0x20758d0c940>
```

In [80]:

```
df4,Ldf4,mm,ax = ut.ch_vertex(df3,Ldf3,pts=pts)
```

In [81]:

```
save_both(k,df4,Ldf4[0])
model = save_model(k,mm,model)
model
```

```
Write to .\data\24hpf\Output-02-15-2019\AT_21_24hpf.psi complete
Write to .\data\24hpf\Output-02-15-2019\ZRF_21_24hpf.psi complete
```

Out[81]:

|  | a | b | c |
| --- | --- | --- | --- |
| 7 | 0.018452 | 1.914462e-17 | -5.692721e-21 |
| 26 | 0.017500 | 2.761276e-17 | 0.000000e+00 |
| 9 | 0.004727 | 2.061185e-16 | 6.484843e-15 |
| 5 | 0.009099 | 5.287413e-17 | -1.423254e-14 |
| 1 | 0.019910 | -4.560282e-16 | 1.231414e-14 |
| 21 | 0.012933 | -1.773224e-16 | -8.838063e-16 |

# 12¶

In [82]:

```
k,df,Ldf,ax = start('12')
```

In [83]:

```
df1,Ldf1,pts,ax = ut.check_pts(df,Ldf,'z')
```

In [84]:

```
df2,Ldf2,mm,ax = ut.ch_vertex(df1,Ldf1)
```

In [85]:

```
pts = pick_pts(-30,10,5,-8,35,10)
ax[0,1].scatter(pts.x,pts.z,c='m',s=50)
```

Out[85]:

```
<matplotlib.collections.PathCollection at 0x20809078f98>
```

In [86]:

```
df3,Ldf3,mm,ax = ut.ch_vertex(df2,Ldf2,pts=pts)
```

In [87]:

```
save_both(k,df3,Ldf3[0])
model = save_model(k,mm,model)
```

```
Write to .\data\24hpf\Output-02-15-2019\AT_12_24hpf.psi complete
Write to .\data\24hpf\Output-02-15-2019\ZRF_12_24hpf.psi complete
```

# 24¶

In [89]:

```
Dbat['24'] = ut.preprocess(os.path.join(at,'AT_24hpf_24_Probabilities.h5'),param,stop='df_thresh')
Dbzrf['24'] = ut.preprocess(os.path.join(gfap,'Gfap_24hpf_24_Probabilities.h5'),param,stop='df_thresh')
```

In [90]:

```
k = '24'
df = Dbat[k].df_thresh
Ldf = [Dbzrf[k].df_thresh]
ax = ut.make_graph([df]+Ldf)
```

In [91]:

```
df1,Ldf1,p,ax = ut.check_yz(df,Ldf)
```

In [92]:

```
df2,Ldf2,mm,ax = ut.ch_vertex(df1,Ldf1)
```

In [93]:

```
save_both(k,df2,Ldf2[0])
model = save_model(k,mm,model)
```

```
Write to .\data\24hpf\Output-02-15-2019\AT_24_24hpf.psi complete
Write to .\data\24hpf\Output-02-15-2019\ZRF_24_24hpf.psi complete
```

# 28¶

In [94]:

```
k = '28'
df = Dbat[k].df_thresh
Ldf = [Dbzrf[k].df_thresh]
ax = ut.make_graph([df]+Ldf)
```

In [95]:

```
df1,Ldf1,pts,ax = ut.check_pts(df,Ldf,'y')
```

In [96]:

```
pts.iloc[0].x = 60
pts.iloc[0].y = 17
ax[0,0].scatter(pts.x,pts.y,c='c')
pts
```

Out[96]:

|  | x | y |
| --- | --- | --- |
| 0 | 60.00 | 17.00 |
| 1 | 6.72 | 8.64 |

In [97]:

```
df2,Ldf2,ax = ut.revise_pts(df,Ldf,'y',pts=pts)
```

In [98]:

```
df3,Ldf3,mm,ax = ut.ch_vertex(df2,Ldf2)
```

In [99]:

```
pts = pick_pts(-28,9,0,-3,23,9)
ax[0,1].scatter(pts.x,pts.z,c='m',s=50)
```

Out[99]:

```
<matplotlib.collections.PathCollection at 0x208636768d0>
```

In [100]:

```
df4,Ldf4,mm,ax = ut.ch_vertex(df3,Ldf3,pts=pts)
```

In [101]:

```
save_both(k,df4,Ldf4[0])
model = save_model(k,mm,model)
```

```
Write to .\data\24hpf\Output-02-15-2019\AT_28_24hpf.psi complete
Write to .\data\24hpf\Output-02-15-2019\ZRF_28_24hpf.psi complete
```

# 3¶

In [102]:

```
k,df,Ldf,ax = start('3')
```

In [103]:

```
df1,Ldf1,pts,ax = ut.check_pts(df,Ldf,'z')
```

In [104]:

```
pts.iloc[1].z = 7
ax[0,1].scatter(pts.x,pts.z,c='y')
pts
```

Out[104]:

|  | x | z |
| --- | --- | --- |
| 0 | 58.142891 | 42.86977 |
| 1 | -34.095611 | 7.00000 |

In [105]:

```
df2,Ldf2,ax = ut.revise_pts(df,Ldf,'z',pts=pts)
```

In [106]:

```
df3,Ldf3,mm,ax = ut.ch_vertex(df2,Ldf2)
```

In [107]:

```
pts = pick_pts(-50,22,0,-10,47,20)
ax[0,1].scatter(pts.x,pts.z,c='m',s=50)
```

Out[107]:

```
<matplotlib.collections.PathCollection at 0x20897388dd8>
```

In [108]:

```
df4,Ldf4,mm,ax = ut.ch_vertex(df3,Ldf3,pts=pts)
```

In [109]:

```
save_both(k,df4,Ldf4[0])
model = save_model(k,mm,model)
```

```
Write to .\data\24hpf\Output-02-15-2019\AT_3_24hpf.psi complete
Write to .\data\24hpf\Output-02-15-2019\ZRF_3_24hpf.psi complete
```

# 10¶

In [110]:

```
k,df,Ldf,ax = start('10')
```

In [111]:

```
df1,Ldf1,pts,ax = ut.check_pts(df,Ldf,'z')
```

In [112]:

```
pts.iloc[1].x = -28
pts.iloc[1].z = 10
ax[0,1].scatter(pts.x,pts.z,c='y')
pts
```

Out[112]:

|  | x | z |
| --- | --- | --- |
| 0 | 46.552267 | 55.507086 |
| 1 | -28.000000 | 10.000000 |

In [113]:

```
df2,Ldf2,mm,ax = ut.ch_vertex(df1,Ldf1)
```

In [114]:

```
pts = pick_pts(-58,25,0,-2,45,25)
ax[0,1].scatter(pts.x,pts.z,c='m',s=50)
```

Out[114]:

```
<matplotlib.collections.PathCollection at 0x208632bf160>
```

In [115]:

```
df3,Ldf3,mm,ax = ut.ch_vertex(df2,Ldf2,pts=pts)
```

In [116]:

```
save_both(k,df3,Ldf3[0])
model = save_model(k,mm,model)
```

```
Write to .\data\24hpf\Output-02-15-2019\AT_10_24hpf.psi complete
Write to .\data\24hpf\Output-02-15-2019\ZRF_10_24hpf.psi complete
```

# 27¶

In [117]:

```
k,df,Ldf,ax = start('27')
```

In [118]:

```
df1,Ldf1 = ut.zyswitch(df,Ldf)
```

In [119]:

```
ax = ut.make_graph([df1]+Ldf1)
```

In [120]:

```
df2,Ldf2,pts,ax = ut.check_pts(df1,Ldf1,'y')
```

In [121]:

```
pts
pts.iloc[0].x = -55
pts.iloc[0].y = 10
pts.iloc[1].x = 30
pts.iloc[1].y = 5
ax[0,0].scatter(pts.x,pts.y,c='c')
```

Out[121]:

```
<matplotlib.collections.PathCollection at 0x20806ebb780>
```

In [122]:

```
df3,Ldf3,ax = ut.revise_pts(df1,Ldf1,'y',pts=pts)
```

In [123]:

```
df4,Ldf4,mm,ax = ut.ch_vertex(df3,Ldf3)
```

In [124]:

```
pts = pick_pts(-40,9,0,-7,40,11)
ax[0,1].scatter(pts.x,pts.z,c='m',s=50)
```

Out[124]:

```
<matplotlib.collections.PathCollection at 0x20897665e48>
```

In [125]:

```
df5,Ldf5,mm,ax = ut.ch_vertex(df4,Ldf4,pts=pts)
```

In [126]:

```
save_both(k,df5,Ldf5[0])
model = save_model(k,mm,model)
```

```
Write to .\data\24hpf\Output-02-15-2019\AT_27_24hpf.psi complete
Write to .\data\24hpf\Output-02-15-2019\ZRF_27_24hpf.psi complete
```

# 17¶

In [128]:

```
k = '17'
Dbat[k] = ut.preprocess(os.path.join(at,'AT_24hpf_17_Probabilities.h5'),param,stop='df_thresh')
Dbzrf[k] = ut.preprocess(os.path.join(gfap,'Gfap_24hpf_17_Probabilities.h5'),param,stop='df_thresh')
```

In [129]:

```
df = Dbat[k].df_thresh
Ldf = [Dbzrf[k].df_thresh]
ax = ut.make_graph([df]+Ldf)
```

In [130]:

```
df1,Ldf1,p,ax = ut.check_yz(df,Ldf)
```

In [131]:

```
df2,Ldf2,mm,ax = ut.ch_vertex(df1,Ldf1)
```

In [132]:

```
pts = pick_pts(-38,16,0,-8,45,16)
ax[0,1].scatter(pts.x,pts.z,c='m',s=50)
```

Out[132]:

```
<matplotlib.collections.PathCollection at 0x208974277b8>
```

In [133]:

```
df3,Ldf3,mm,ax = ut.ch_vertex(df2,Ldf2,pts=pts)
```

In [134]:

```
save_both(k,df3,Ldf3[0])
model = save_model(k,mm,model)
```

```
Write to .\data\24hpf\Output-02-15-2019\AT_17_24hpf.psi complete
Write to .\data\24hpf\Output-02-15-2019\ZRF_17_24hpf.psi complete
```

# 4¶

In [135]:

```
k,df,Ldf,ax = start('4')
```

In [136]:

```
ut.make_graph([df,Dbat[k].df_thresh])
```

Out[136]:

```
array([[<matplotlib.axes._subplots.AxesSubplot object at 0x000002089818B438>,
        <matplotlib.axes._subplots.AxesSubplot object at 0x0000020898447128>,
        <matplotlib.axes._subplots.AxesSubplot object at 0x000002089846E550>],
       [<matplotlib.axes._subplots.AxesSubplot object at 0x0000020898495AC8>,
        <matplotlib.axes._subplots.AxesSubplot object at 0x00000208984C8080>,
        <matplotlib.axes._subplots.AxesSubplot object at 0x00000208984ED5C0>]],
      dtype=object)
```

In [137]:

```
df1,Ldf1,mm,ax = ut.ch_vertex(Dbat[k].df_thresh,[Dbzrf[k].df_thresh])
```

In [138]:

```
pts = pick_pts(-55,6,0,-5,52,6)
ax[0,1].scatter(pts.x,pts.z,c='m',s=50)
```

Out[138]:

```
<matplotlib.collections.PathCollection at 0x208980d6a20>
```

In [139]:

```
df2,Ldf2,mm,ax = ut.ch_vertex(df1,Ldf1,pts=pts)
```

In [140]:

```
save_both(k,df2,Ldf2[0])
model = save_model(k,mm,model)
```

```
Write to .\data\24hpf\Output-02-15-2019\AT_4_24hpf.psi complete
Write to .\data\24hpf\Output-02-15-2019\ZRF_4_24hpf.psi complete
```

# 11¶

In [141]:

```
k,df,Ldf,ax = start('11')
```

In [142]:

```
df1,Ldf1,mm,ax = ut.ch_vertex(df,Ldf)
```

In [143]:

```
pts = pick_pts(-48,11,-5,-8,36,8)
ax[0,1].scatter(pts.x,pts.z,c='m',s=50)
```

Out[143]:

```
<matplotlib.collections.PathCollection at 0x20863127c18>
```

In [144]:

```
df2,Ldf2,mm,ax = ut.ch_vertex(df1,Ldf1,pts=pts)
```

In [145]:

```
save_both(k,df2,Ldf2[0])
model = save_model(k,mm,model)
```

```
Write to .\data\24hpf\Output-02-15-2019\AT_11_24hpf.psi complete
Write to .\data\24hpf\Output-02-15-2019\ZRF_11_24hpf.psi complete
```

# 6¶

In [146]:

```
k,df,Ldf,ax = start('6')
```

In [147]:

```
df1,Ldf1,pts,ax = ut.check_pts(df,Ldf,'z')
```

In [148]:

```
pts.iloc[1].x = -56
pts.iloc[1].z = 24
ax[0,1].scatter(pts.x,pts.z,c='y')
pts
```

Out[148]:

|  | x | z |
| --- | --- | --- |
| 0 | 44.665085 | 36.879888 |
| 1 | -56.000000 | 24.000000 |

In [149]:

```
df2,Ldf2,ax = ut.revise_pts(df,Ldf,'z',pts=pts)
```

In [150]:

```
df3,Ldf3,mm,ax = ut.ch_vertex(df2,Ldf2)
```

In [151]:

```
save_both(k,df3,Ldf3[0])
model = save_model(k,mm,model)
```

```
Write to .\data\24hpf\Output-02-15-2019\AT_6_24hpf.psi complete
Write to .\data\24hpf\Output-02-15-2019\ZRF_6_24hpf.psi complete
```

# 23¶

In [153]:

```
k = '23'
Dbat[k] = ut.preprocess(os.path.join(at,'AT_24hpf_23_Probabilities.h5'),param,stop='df_thresh')
Dbzrf[k] = ut.preprocess(os.path.join(gfap,'Gfap_24hpf_23_Probabilities.h5'),param,stop='df_thresh')
```

In [154]:

```
df = Dbat[k].df_thresh
Ldf = [Dbzrf[k].df_thresh]
ax = ut.make_graph([df]+Ldf)
```

In [155]:

```
df1,Ldf1,p,ax = ut.check_yz(df,Ldf)
```

In [156]:

```
df2,Ldf2,mm,ax = ut.ch_vertex(df1,Ldf1)
```

In [157]:

```
save_both(k,df2,Ldf2[0])
model = save_model(k,mm,model)
```

```
Write to .\data\24hpf\Output-02-15-2019\AT_23_24hpf.psi complete
Write to .\data\24hpf\Output-02-15-2019\ZRF_23_24hpf.psi complete
```

# 19¶

In [158]:

```
k,df,Ldf,ax = start('19')
```

In [159]:

```
pts = pick_pts(-65,20,0,-10,55,17)
ax[0,1].scatter(pts.x,pts.z,c='m',s=50)
```

Out[159]:

```
<matplotlib.collections.PathCollection at 0x208d11e05f8>
```

In [160]:

```
df1,Ldf1,mm,ax = ut.ch_vertex(df,Ldf,pts=pts)
```

In [161]:

```
save_both(k,df1,Ldf1[0])
model = save_model(k,mm,model)
```

```
Write to .\data\24hpf\Output-02-15-2019\AT_19_24hpf.psi complete
Write to .\data\24hpf\Output-02-15-2019\ZRF_19_24hpf.psi complete
```

# 25¶

In [162]:

```
k,df,Ldf,ax = start('25')
```

In [163]:

```
df1,Ldf1,ax,p = ut.check_yz(df,Ldf)
```

In [164]:

```
p
```

Out[164]:

```
poly1d([ 0.40313491, -2.74690804])
```

In [165]:

```
df2,Ldf2,ax,p = ut.check_yz(df,Ldf,mm=[1,-2.7])
```

In [166]:

```
df3,Ldf3,mm,ax = ut.ch_vertex(df2,Ldf2)
```

In [167]:

```
save_both(k,df3,Ldf3[0])
model = save_model(k,mm,model)
```

```
Write to .\data\24hpf\Output-02-15-2019\AT_25_24hpf.psi complete
Write to .\data\24hpf\Output-02-15-2019\ZRF_25_24hpf.psi complete
```

# 8¶

In [168]:

```
k,df,Ldf,ax = start('8')
```

In [169]:

```
df1,Ldf1,ax,p = ut.check_yz(df,Ldf)
```

In [170]:

```
df2,Ldf2,pts,ax = ut.check_pts(df1,Ldf1,'z')
```

In [171]:

```
pts.iloc[1].x = 43
pts.iloc[1].z = 25
ax[0,1].scatter(pts.x,pts.z,c='y')
pts
```

Out[171]:

|  | x | z |
| --- | --- | --- |
| 0 | -54.177133 | 31.661079 |
| 1 | 43.000000 | 25.000000 |

In [172]:

```
df3,Ldf3,ax = ut.revise_pts(df1,Ldf1,'z',pts=pts)
```

In [173]:

```
df4,Ldf4,mm,ax = ut.ch_vertex(df3,Ldf3)
```

In [174]:

```
save_both(k,df4,Ldf4[0])
model = save_model(k,mm,model)
```

```
Write to .\data\24hpf\Output-02-15-2019\AT_8_24hpf.psi complete
Write to .\data\24hpf\Output-02-15-2019\ZRF_8_24hpf.psi complete
```

In [175]:

```
model
```

Out[175]:

|  | a | b | c |
| --- | --- | --- | --- |
| 7 | 0.018452 | 1.914462e-17 | -5.692721e-21 |
| 26 | 0.017500 | 2.761276e-17 | 0.000000e+00 |
| 9 | 0.004727 | 2.061185e-16 | 6.484843e-15 |
| 5 | 0.009099 | 5.287413e-17 | -1.423254e-14 |
| 1 | 0.019910 | -4.560282e-16 | 1.231414e-14 |
| 21 | 0.012933 | -1.773224e-16 | -8.838063e-16 |
| 12 | 0.017143 | 2.387306e-16 | -5.129661e-15 |
| 24 | 0.011212 | 1.409065e-16 | 3.431518e-15 |
| 28 | 0.018634 | -2.287230e-16 | 3.079476e-15 |
| 3 | 0.013178 | -9.440544e-17 | 5.942779e-15 |
| 10 | 0.010345 | 2.094698e-16 | -2.061921e-15 |
| 27 | 0.010625 | -1.097348e-16 | 1.418498e-15 |
| 17 | 0.014035 | -6.426921e-17 | 6.152684e-15 |
| 4 | 0.003846 | 3.740995e-17 | -2.563972e-15 |
| 11 | 0.009906 | 7.740906e-17 | 2.494437e-15 |
| 6 | 0.008516 | 1.994954e-16 | -3.326923e-15 |
| 23 | 0.010611 | -4.715987e-16 | -2.876130e-14 |
| 19 | 0.007937 | -1.523041e-16 | -6.897951e-15 |
| 25 | 0.003600 | 4.781800e-18 | 3.397189e-15 |
| 8 | 0.007573 | -8.906381e-18 | 3.158467e-15 |

In [176]:

```
model.to_csv(os.path.join(outdir,'model.csv'))
```
