## Supplementary material for "ΔSCOPE: A new method to quantify 3D biological structures and identify differences in zebrafish forebrain development": An archived copy of the DeltaSCOPE code repository.: alignment-24hpf.ipynb

```
# %% [code]
import numpy as np
import pandas as pd
import matplotlib.pyplot as plt

import os
import re
from imp import reload
import h5py
import sys

# %% [code]
import deltascope as cranium
import deltascope.alignment as ut

# %% [code]
at = ".\\data\\24hpf\\AT\\Prob"
gfap = ".\\data\\24hpf\\Gfap\\Prob"

# %% [code]
outdir = ".\\data\\24hpf\\Output-02-15-2019"

# %% [code]
os.mkdir(outdir)

# %% [code]
Dat = {}
for f in os.listdir(at):
    if 'h5' in f:
        num  = re.findall(r'\d+',f.split('.')[0])[-1]
        Dat[num] = os.path.join(at,f)

# %% [code]
Dzrf = {}
for f in os.listdir(gfap):
    if 'h5' in f:
        num  = re.findall(r'\d+',f.split('.')[0])[-1]
        Dzrf[num] = os.path.join(gfap,f)

# %% [code]
Dat.keys()

# %% [code]
Dbat = {}
Dbzrf={}

# %% [markdown]
# Data Preprocessing

# %% [code]
klist = Dat.keys()

# %% [code]
param = {
    'gthresh':0.5,
    'scale':[1,1,1],
    'microns':[0.16,0.16,0.21],
    'mthresh':0.8,
    'radius':10,
    'comp_order':[0,2,1],
    'fit_dim':['x','z'],
    'deg':2
}

# %% [code]
for k in klist:
    try:
        Dbat[k] = ut.preprocess(Dat[k],param)
        Dbzrf[k] = ut.preprocess(Dzrf[k],param,pca=Dbat[k].pcamed,mm=Dbat[k].mm,vertex=Dbat[k].vertex)
        print(k)
    except:
        print(k,'failed')
        e = sys.exc_info()
        print(e)

# %% [code]
def start(k):
    return(ut.start(k,Dbat,[Dbzrf],im=False))
def save_at(k,df):
    ut.save_at(k,df,outdir,'24hpf')
def save_both(k,dfa,dfb):
    ut.save_both(k,dfa,dfb,outdir,'24hpf')

# %% [code]
def fit_model(axi,df,mm=None):
    if mm == None:
        mm = np.polyfit(df.x,df.z,2)
    p = np.poly1d(mm)
    xrange = np.arange(np.min(df.x),np.max(df.x))
    axi.plot(xrange,p(xrange),c='m')
    return(mm)

# %% [code]
model = pd.DataFrame({'a':[],'b':[],'c':[]})
def save_model(k,mm,model):
    row = pd.Series({'a':mm[0],'b':mm[1],'c':mm[2]},name=k)
    model = model.append(row)
    return(model)

# %% [code]
def pick_pts(x1,z1,vx,vz,x2,z2):
    pts = pd.DataFrame({'x':[x1,vx,x2],'z':[z1,vz,z2]})
    return(pts)

# %% [code]
klist

# %% [markdown]
# 7

# %% [code]
k,df,Ldf,ax = start('7')

# %% [code]
df1,Ldf1,pts,ax = ut.check_pts(df,Ldf,'z')

# %% [code]
pts.iloc[1].x = 36
pts.iloc[1].z = 22
ax[0,1].scatter(pts.x,pts.z,c='y')

# %% [code]
df2,Ldf2,ax = ut.revise_pts(df,Ldf,'z',pts=pts)

# %% [code]
df3,Ldf3,p,ax = ut.check_yz(df2,Ldf2)

# %% [code]
df4,Ldf4,mm,ax = ut.ch_vertex(df3,Ldf3)

# %% [code]
pts = pick_pts(-42,27,0,-4,40,27)
ax = ut.make_graph([df4]+Ldf4)
ax[0,1].scatter(pts.x,pts.z,c='m',s=40)

# %% [code]
df5,Ldf5,mm,ax = ut.ch_vertex(df4,Ldf4,pts=pts)

# %% [code]
save_both(k,df5,Ldf5[0])
model = save_model(k,mm,model)

# %% [markdown]
# 26

# %% [code]
k,df,Ldf,ax = start('26')

# %% [code]
df1,Ldf1,pts,ax = ut.check_pts(df,Ldf,'z')

# %% [code]
pts.iloc[1].x = 18
pts.iloc[1].z = 2
ax[0,1].scatter(pts.x,pts.z,c='y')
pts

# %% [code]
df2,Ldf2,ax = ut.revise_pts(df,Ldf,'z',pts=pts)

# %% [code]
df3,Ldf3,pts,ax = ut.check_pts(df2,Ldf2,'y')

# %% [code]
pts.iloc[1].x = 16
pts.iloc[1].y = -1
pts.iloc[0].y = 18
ax[0,0].scatter(pts.x,pts.y,c='y')
pts

# %% [code]
df4,Ldf4,ax = ut.revise_pts(df2,Ldf2,'y',pts=pts)

# %% [code]
df5,Ldf5,p,ax = ut.check_yz(df4,Ldf4)

# %% [code]
p2 = np.poly1d([2/5,0])
xrange = np.arange(-20,20)
p[0,2].plot(xrange,p2(xrange),c='c')

# %% [code]
df6,Ldf6,ax,p = ut.check_yz(df4,Ldf4,mm=[2/5,0])

# %% [code]
df7,Ldf7,mm,ax = ut.ch_vertex(df6,Ldf6)

# %% [code]
pts = pick_pts(-40,20,0,-8,40,20)
ax[0,1].scatter(pts.x,pts.z,c='m',s=40)

# %% [code]
df8,Ldf8,mm,ax = ut.ch_vertex(df7,Ldf7,pts=pts)

# %% [code]
save_both(k,df8,Ldf8[0])
model = save_model(k,mm,model)

# %% [code]
model = model.iloc[-2:]

# %% [code]
model

# %% [markdown]
# 9

# %% [code]
Dbat['9'] = ut.preprocess(os.path.join(at,'AT_24hpf_9_Probabilities.h5'),param,stop='df_thresh')
Dbzrf['9'] = ut.preprocess(os.path.join(gfap,'Gfap_24hpf_9_Probabilities.h5'),param,stop='df_thresh')

# %% [code]
k = '9'
df = Dbat[k].df_thresh
Ldf = [Dbzrf[k].df_thresh]

# %% [code]
ax = ut.make_graph([df]+Ldf)

# %% [code]
df1,Ldf1,mm,ax = ut.ch_vertex(df,Ldf)

# %% [code]
save_both(k,df1,Ldf1[0])
model = save_model(k,mm,model)
model

# %% [markdown]
# 5

# %% [code]
k,df,Ldf,ax = start('5')

# %% [code]
df1,Ldf1,pts,ax = ut.check_pts(df,Ldf,'z')

# %% [code]
df2,Ldf2,mm,ax = ut.ch_vertex(df1,Ldf1)

# %% [code]
pts = pick_pts(-55,22,0,-8,58,20)
ax[0,1].scatter(pts.x,pts.z,s=50,c='m')

# %% [code]
df3,Ldf3,mm,ax = ut.ch_vertex(df2,Ldf2,pts=pts)

# %% [code]
save_both(k,df3,Ldf3[0])
model = save_model(k,mm,model)
model

# %% [markdown]
# 1

# %% [code]
k = '1'
df = Dbat[k].df_thresh
Ldf = [Dbzrf[k].df_thresh]

# %% [code]
ax = ut.make_graph([df]+Ldf)

# %% [code]
df1,Ldf1,ax,p = ut.check_yz(df,Ldf)

# %% [code]
df2,Ldf2,mm,ax = ut.ch_vertex(df1,Ldf1)

# %% [code]
pts = pick_pts(-55,20,-6,-20,35,20)
ax[0,1].scatter(pts.x,pts.z,c='m',s=50)

# %% [code]
df3,Ldf3,mm,ax = ut.ch_vertex(df2,Ldf2,pts=pts)

# %% [code]
save_both(k,df3,Ldf3[0])
model = save_model(k,mm,model)
model

# %% [markdown]
# 21

# %% [code]
k,df,Ldf,ax = start('21')

# %% [code]
df1,Ldf1,ax,p = ut.check_yz(df,Ldf)

# %% [code]
mm = [3/5,0]
p = np.poly1d(mm)
xrange = np.arange(-10,30)
ax[0,2].plot(xrange,p(xrange),c='y')

# %% [code]
df2,Ldf2,ax,p = ut.check_yz(df,Ldf,mm=mm)

# %% [code]
df3,Ldf3,mm,ax = ut.ch_vertex(df2,Ldf2)

# %% [code]
pts = pick_pts(-48,24,0,-6,45,20)
ax[0,1].scatter(pts.x,pts.z,c='m',s=50)

# %% [code]
df4,Ldf4,mm,ax = ut.ch_vertex(df3,Ldf3,pts=pts)

# %% [code]
save_both(k,df4,Ldf4[0])
model = save_model(k,mm,model)
model

# %% [markdown]
# 12

# %% [code]
k,df,Ldf,ax = start('12')

# %% [code]
df1,Ldf1,pts,ax = ut.check_pts(df,Ldf,'z')

# %% [code]
df2,Ldf2,mm,ax = ut.ch_vertex(df1,Ldf1)

# %% [code]
pts = pick_pts(-30,10,5,-8,35,10)
ax[0,1].scatter(pts.x,pts.z,c='m',s=50)

# %% [code]
df3,Ldf3,mm,ax = ut.ch_vertex(df2,Ldf2,pts=pts)

# %% [code]
save_both(k,df3,Ldf3[0])
model = save_model(k,mm,model)

# %% [markdown]
# 24

# %% [code]
Dbat['24'] = ut.preprocess(os.path.join(at,'AT_24hpf_24_Probabilities.h5'),param,stop='df_thresh')
Dbzrf['24'] = ut.preprocess(os.path.join(gfap,'Gfap_24hpf_24_Probabilities.h5'),param,stop='df_thresh')

# %% [code]
k = '24'
df = Dbat[k].df_thresh
Ldf = [Dbzrf[k].df_thresh]
ax = ut.make_graph([df]+Ldf)

# %% [code]
df1,Ldf1,p,ax = ut.check_yz(df,Ldf)

# %% [code]
df2,Ldf2,mm,ax = ut.ch_vertex(df1,Ldf1)

# %% [code]
save_both(k,df2,Ldf2[0])
model = save_model(k,mm,model)

# %% [markdown]
# 28

# %% [code]
k = '28'
df = Dbat[k].df_thresh
Ldf = [Dbzrf[k].df_thresh]
ax = ut.make_graph([df]+Ldf)

# %% [code]
df1,Ldf1,pts,ax = ut.check_pts(df,Ldf,'y')

# %% [code]
pts.iloc[0].x = 60
pts.iloc[0].y = 17
ax[0,0].scatter(pts.x,pts.y,c='c')
pts

# %% [code]
df2,Ldf2,ax = ut.revise_pts(df,Ldf,'y',pts=pts)

# %% [code]
df3,Ldf3,mm,ax = ut.ch_vertex(df2,Ldf2)

# %% [code]
pts = pick_pts(-28,9,0,-3,23,9)
ax[0,1].scatter(pts.x,pts.z,c='m',s=50)

# %% [code]
df4,Ldf4,mm,ax = ut.ch_vertex(df3,Ldf3,pts=pts)

# %% [code]
save_both(k,df4,Ldf4[0])
model = save_model(k,mm,model)

# %% [markdown]
# 3

# %% [code]
k,df,Ldf,ax = start('3')

# %% [code]
df1,Ldf1,pts,ax = ut.check_pts(df,Ldf,'z')

# %% [code]
pts.iloc[1].z = 7
ax[0,1].scatter(pts.x,pts.z,c='y')
pts

# %% [code]
df2,Ldf2,ax = ut.revise_pts(df,Ldf,'z',pts=pts)

# %% [code]
df3,Ldf3,mm,ax = ut.ch_vertex(df2,Ldf2)

# %% [code]
pts = pick_pts(-50,22,0,-10,47,20)
ax[0,1].scatter(pts.x,pts.z,c='m',s=50)

# %% [code]
df4,Ldf4,mm,ax = ut.ch_vertex(df3,Ldf3,pts=pts)

# %% [code]
save_both(k,df4,Ldf4[0])
model = save_model(k,mm,model)

# %% [markdown]
# 10

# %% [code]
k,df,Ldf,ax = start('10')

# %% [code]
df1,Ldf1,pts,ax = ut.check_pts(df,Ldf,'z')

# %% [code]
pts.iloc[1].x = -28
pts.iloc[1].z = 10
ax[0,1].scatter(pts.x,pts.z,c='y')
pts

# %% [code]
df2,Ldf2,mm,ax = ut.ch_vertex(df1,Ldf1)

# %% [code]
pts = pick_pts(-58,25,0,-2,45,25)
ax[0,1].scatter(pts.x,pts.z,c='m',s=50)

# %% [code]
df3,Ldf3,mm,ax = ut.ch_vertex(df2,Ldf2,pts=pts)

# %% [code]
save_both(k,df3,Ldf3[0])
model = save_model(k,mm,model)

# %% [markdown]
# 27

# %% [code]
k,df,Ldf,ax = start('27')

# %% [code]
df1,Ldf1 = ut.zyswitch(df,Ldf)

# %% [code]
ax = ut.make_graph([df1]+Ldf1)

# %% [code]
df2,Ldf2,pts,ax = ut.check_pts(df1,Ldf1,'y')

# %% [code]
pts
pts.iloc[0].x = -55
pts.iloc[0].y = 10
pts.iloc[1].x = 30
pts.iloc[1].y = 5
ax[0,0].scatter(pts.x,pts.y,c='c')

# %% [code]
df3,Ldf3,ax = ut.revise_pts(df1,Ldf1,'y',pts=pts)

# %% [code]
df4,Ldf4,mm,ax = ut.ch_vertex(df3,Ldf3)

# %% [code]
pts = pick_pts(-40,9,0,-7,40,11)
ax[0,1].scatter(pts.x,pts.z,c='m',s=50)

# %% [code]
df5,Ldf5,mm,ax = ut.ch_vertex(df4,Ldf4,pts=pts)

# %% [code]
save_both(k,df5,Ldf5[0])
model = save_model(k,mm,model)

# %% [markdown]
# 17

# %% [code]
k = '17'
Dbat[k] = ut.preprocess(os.path.join(at,'AT_24hpf_17_Probabilities.h5'),param,stop='df_thresh')
Dbzrf[k] = ut.preprocess(os.path.join(gfap,'Gfap_24hpf_17_Probabilities.h5'),param,stop='df_thresh')

# %% [code]
df = Dbat[k].df_thresh
Ldf = [Dbzrf[k].df_thresh]
ax = ut.make_graph([df]+Ldf)

# %% [code]
df1,Ldf1,p,ax = ut.check_yz(df,Ldf)

# %% [code]
df2,Ldf2,mm,ax = ut.ch_vertex(df1,Ldf1)

# %% [code]
pts = pick_pts(-38,16,0,-8,45,16)
ax[0,1].scatter(pts.x,pts.z,c='m',s=50)

# %% [code]
df3,Ldf3,mm,ax = ut.ch_vertex(df2,Ldf2,pts=pts)

# %% [code]
save_both(k,df3,Ldf3[0])
model = save_model(k,mm,model)

# %% [markdown]
# 4

# %% [code]
k,df,Ldf,ax = start('4')

# %% [code]
ut.make_graph([df,Dbat[k].df_thresh])

# %% [code]
df1,Ldf1,mm,ax = ut.ch_vertex(Dbat[k].df_thresh,[Dbzrf[k].df_thresh])

# %% [code]
pts = pick_pts(-55,6,0,-5,52,6)
ax[0,1].scatter(pts.x,pts.z,c='m',s=50)

# %% [code]
df2,Ldf2,mm,ax = ut.ch_vertex(df1,Ldf1,pts=pts)

# %% [code]
save_both(k,df2,Ldf2[0])
model = save_model(k,mm,model)

# %% [markdown]
# 11

# %% [code]
k,df,Ldf,ax = start('11')

# %% [code]
df1,Ldf1,mm,ax = ut.ch_vertex(df,Ldf)

# %% [code]
pts = pick_pts(-48,11,-5,-8,36,8)
ax[0,1].scatter(pts.x,pts.z,c='m',s=50)

# %% [code]
df2,Ldf2,mm,ax = ut.ch_vertex(df1,Ldf1,pts=pts)

# %% [code]
save_both(k,df2,Ldf2[0])
model = save_model(k,mm,model)

# %% [markdown]
# 6

# %% [code]
k,df,Ldf,ax = start('6')

# %% [code]
df1,Ldf1,pts,ax = ut.check_pts(df,Ldf,'z')

# %% [code]
pts.iloc[1].x = -56
pts.iloc[1].z = 24
ax[0,1].scatter(pts.x,pts.z,c='y')
pts

# %% [code]
df2,Ldf2,ax = ut.revise_pts(df,Ldf,'z',pts=pts)

# %% [code]
df3,Ldf3,mm,ax = ut.ch_vertex(df2,Ldf2)

# %% [code]
save_both(k,df3,Ldf3[0])
model = save_model(k,mm,model)

# %% [markdown]
# 23

# %% [code]
k = '23'
Dbat[k] = ut.preprocess(os.path.join(at,'AT_24hpf_23_Probabilities.h5'),param,stop='df_thresh')
Dbzrf[k] = ut.preprocess(os.path.join(gfap,'Gfap_24hpf_23_Probabilities.h5'),param,stop='df_thresh')

# %% [code]
df = Dbat[k].df_thresh
Ldf = [Dbzrf[k].df_thresh]
ax = ut.make_graph([df]+Ldf)

# %% [code]
df1,Ldf1,p,ax = ut.check_yz(df,Ldf)

# %% [code]
df2,Ldf2,mm,ax = ut.ch_vertex(df1,Ldf1)

# %% [code]
save_both(k,df2,Ldf2[0])
model = save_model(k,mm,model)

# %% [markdown]
# 19

# %% [code]
k,df,Ldf,ax = start('19')

# %% [code]
pts = pick_pts(-65,20,0,-10,55,17)
ax[0,1].scatter(pts.x,pts.z,c='m',s=50)

# %% [code]
df1,Ldf1,mm,ax = ut.ch_vertex(df,Ldf,pts=pts)

# %% [code]
save_both(k,df1,Ldf1[0])
model = save_model(k,mm,model)

# %% [markdown]
# 25

# %% [code]
k,df,Ldf,ax = start('25')

# %% [code]
df1,Ldf1,ax,p = ut.check_yz(df,Ldf)

# %% [code]
p

# %% [code]
df2,Ldf2,ax,p = ut.check_yz(df,Ldf,mm=[1,-2.7])

# %% [code]
df3,Ldf3,mm,ax = ut.ch_vertex(df2,Ldf2)

# %% [code]
save_both(k,df3,Ldf3[0])
model = save_model(k,mm,model)

# %% [markdown]
# 8

# %% [code]
k,df,Ldf,ax = start('8')

# %% [code]
df1,Ldf1,ax,p = ut.check_yz(df,Ldf)

# %% [code]
df2,Ldf2,pts,ax = ut.check_pts(df1,Ldf1,'z')

# %% [code]
pts.iloc[1].x = 43
pts.iloc[1].z = 25
ax[0,1].scatter(pts.x,pts.z,c='y')
pts

# %% [code]
df3,Ldf3,ax = ut.revise_pts(df1,Ldf1,'z',pts=pts)

# %% [code]
df4,Ldf4,mm,ax = ut.ch_vertex(df3,Ldf3)

# %% [code]
save_both(k,df4,Ldf4[0])
model = save_model(k,mm,model)

# %% [code]
model

# %% [code]
model.to_csv(os.path.join(outdir,'model.csv'))
```
