## Supplementary material for "ΔSCOPE: A new method to quantify 3D biological structures and identify differences in zebrafish forebrain development": An archived copy of the DeltaSCOPE code repository.: alignment-26hpf.html


In [1]:

```
import numpy as np
import pandas as pd
import matplotlib.pyplot as plt

import os
import re
from imp import reload
import h5py
```

In [2]:

```
import deltascope as cranium
import deltascope.alignment as ut
```

In [3]:

```
at = ".\\data\\26hpf\\AT\\Prob"
gfap = ".\\data\\26hpf\\Gfap\\Prob"
```

In [4]:

```
outdir = ".\\data\\26hpf\\Output-02-14-2019"
```

In [5]:

```
os.mkdir(outdir)
```

In [6]:

```
Dat = {}
for f in os.listdir(at):
    if 'h5' in f:
        num  = re.findall(r'\d+',f.split('.')[0])[-1]
        Dat[num] = os.path.join(at,f)
```

In [7]:

```
Dzrf = {}
for f in os.listdir(gfap):
    if 'h5' in f:
        num  = re.findall(r'\d+',f.split('.')[0])[-1]
        Dzrf[num] = os.path.join(gfap,f)
```

In [8]:

```
Dbat = {}
Dbzrf = {}
```

### Data Preprocessing¶

In [9]:

```
klist = Dat.keys()
```

In [10]:

```
param = {
    'gthresh':0.5,
    'scale':[1,1,1],
    'microns':[0.16,0.16,0.21],
    'mthresh':0.5,
    'radius':10,
    'comp_order':[0,2,1],
    'fit_dim':['x','z'],
    'deg':2
}
```

In [11]:

```
%%time
for k in klist:
    Dbat[k] = ut.preprocess(Dat[k],param)
    Dbzrf[k] = ut.preprocess(Dzrf[k],param,pca=Dbat[k].pcamed,mm=Dbat[k].mm,vertex=Dbat[k].vertex)
    print(k)
```

```
C:\Users\zfishlab\AppData\Local\Continuum\anaconda3\envs\test\lib\site-packages\skimage\util\dtype.py:141: UserWarning: Possible precision loss when converting from float32 to uint8
  .format(dtypeobj_in, dtypeobj_out))
```

```
10
```

```
C:\Users\zfishlab\AppData\Local\Continuum\anaconda3\envs\test\lib\site-packages\skimage\util\dtype.py:141: UserWarning: Possible precision loss when converting from float32 to uint8
  .format(dtypeobj_in, dtypeobj_out))
```

```
11
```

```
C:\Users\zfishlab\AppData\Local\Continuum\anaconda3\envs\test\lib\site-packages\skimage\util\dtype.py:141: UserWarning: Possible precision loss when converting from float32 to uint8
  .format(dtypeobj_in, dtypeobj_out))
```

```
12
```

```
C:\Users\zfishlab\AppData\Local\Continuum\anaconda3\envs\test\lib\site-packages\skimage\util\dtype.py:141: UserWarning: Possible precision loss when converting from float32 to uint8
  .format(dtypeobj_in, dtypeobj_out))
```

```
13
```

```
C:\Users\zfishlab\AppData\Local\Continuum\anaconda3\envs\test\lib\site-packages\skimage\util\dtype.py:141: UserWarning: Possible precision loss when converting from float32 to uint8
  .format(dtypeobj_in, dtypeobj_out))
```

```
14
```

```
C:\Users\zfishlab\AppData\Local\Continuum\anaconda3\envs\test\lib\site-packages\skimage\util\dtype.py:141: UserWarning: Possible precision loss when converting from float32 to uint8
  .format(dtypeobj_in, dtypeobj_out))
```

```
15
```

```
C:\Users\zfishlab\AppData\Local\Continuum\anaconda3\envs\test\lib\site-packages\skimage\util\dtype.py:141: UserWarning: Possible precision loss when converting from float32 to uint8
  .format(dtypeobj_in, dtypeobj_out))
```

```
1
```

```
C:\Users\zfishlab\AppData\Local\Continuum\anaconda3\envs\test\lib\site-packages\skimage\util\dtype.py:141: UserWarning: Possible precision loss when converting from float32 to uint8
  .format(dtypeobj_in, dtypeobj_out))
```

```
2
```

```
C:\Users\zfishlab\AppData\Local\Continuum\anaconda3\envs\test\lib\site-packages\skimage\util\dtype.py:141: UserWarning: Possible precision loss when converting from float32 to uint8
  .format(dtypeobj_in, dtypeobj_out))
```

```
3
```

```
C:\Users\zfishlab\AppData\Local\Continuum\anaconda3\envs\test\lib\site-packages\skimage\util\dtype.py:141: UserWarning: Possible precision loss when converting from float32 to uint8
  .format(dtypeobj_in, dtypeobj_out))
```

```
4
```

```
C:\Users\zfishlab\AppData\Local\Continuum\anaconda3\envs\test\lib\site-packages\skimage\util\dtype.py:141: UserWarning: Possible precision loss when converting from float32 to uint8
  .format(dtypeobj_in, dtypeobj_out))
```

```
5
```

```
C:\Users\zfishlab\AppData\Local\Continuum\anaconda3\envs\test\lib\site-packages\skimage\util\dtype.py:141: UserWarning: Possible precision loss when converting from float32 to uint8
  .format(dtypeobj_in, dtypeobj_out))
```

```
64
```

```
C:\Users\zfishlab\AppData\Local\Continuum\anaconda3\envs\test\lib\site-packages\skimage\util\dtype.py:141: UserWarning: Possible precision loss when converting from float32 to uint8
  .format(dtypeobj_in, dtypeobj_out))
```

```
66
```

```
C:\Users\zfishlab\AppData\Local\Continuum\anaconda3\envs\test\lib\site-packages\skimage\util\dtype.py:141: UserWarning: Possible precision loss when converting from float32 to uint8
  .format(dtypeobj_in, dtypeobj_out))
```

```
68
```

```
C:\Users\zfishlab\AppData\Local\Continuum\anaconda3\envs\test\lib\site-packages\skimage\util\dtype.py:141: UserWarning: Possible precision loss when converting from float32 to uint8
  .format(dtypeobj_in, dtypeobj_out))
```

```
6
```

```
C:\Users\zfishlab\AppData\Local\Continuum\anaconda3\envs\test\lib\site-packages\skimage\util\dtype.py:141: UserWarning: Possible precision loss when converting from float32 to uint8
  .format(dtypeobj_in, dtypeobj_out))
```

```
72
```

```
C:\Users\zfishlab\AppData\Local\Continuum\anaconda3\envs\test\lib\site-packages\skimage\util\dtype.py:141: UserWarning: Possible precision loss when converting from float32 to uint8
  .format(dtypeobj_in, dtypeobj_out))
```

```
7
```

```
C:\Users\zfishlab\AppData\Local\Continuum\anaconda3\envs\test\lib\site-packages\skimage\util\dtype.py:141: UserWarning: Possible precision loss when converting from float32 to uint8
  .format(dtypeobj_in, dtypeobj_out))
```

```
8
```

```
C:\Users\zfishlab\AppData\Local\Continuum\anaconda3\envs\test\lib\site-packages\skimage\util\dtype.py:141: UserWarning: Possible precision loss when converting from float32 to uint8
  .format(dtypeobj_in, dtypeobj_out))
```

```
9
Wall time: 10min 42s
```

In [12]:

```
def start(k):
    return(ut.start(k,Dbat,[Dbzrf],im=False))
def save_both(k,dfa,dfb):
    ut.save_both(k,dfa,dfb,outdir,'26hpf')
```

In [13]:

```
model = pd.DataFrame({'a':[],'b':[],'c':[]})
def save_model(k,mm,model):
    row = pd.Series({'a':mm[0],'b':mm[1],'c':mm[2]},name=k)
    model = model.append(row)
    return(model)
```

In [14]:

```
def fit_model(axi,df,mm=None):
    if mm == None:
        mm = np.polyfit(df.x,df.z,2)
    p = np.poly1d(mm)
    xrange = np.arange(np.min(df.x),np.max(df.x))
    axi.plot(xrange,p(xrange),c='m')
    return(mm)
```

In [15]:

```
def pick_pts(x1,z1,vx,vz,x2,z2):
    pts = pd.DataFrame({'x':[x1,vx,x2],'z':[z1,vz,z2]})
    return(pts)
```

# 72¶

In [16]:

```
k,df,Ldf,ax = start('72')
```

In [17]:

```
mm = fit_model(ax[0,1],df)
```

In [18]:

```
model = save_model(k,mm,model)
```

In [19]:

```
save_both(k,df,Ldf[0])
```

```
Write to .\data\26hpf\Output-02-14-2019\AT_72_26hpf.psi complete
Write to .\data\26hpf\Output-02-14-2019\ZRF_72_26hpf.psi complete
```

# 68¶

Really minimal signal; Discard

In [20]:

```
k,df,Ldf,ax = start('68')
```

# 3¶

In [21]:

```
k,df,Ldf,ax = start('3')
```

In [22]:

```
mm = fit_model(ax[0,1],df)
```

In [23]:

```
save_both(k,df,Ldf[0])
```

```
Write to .\data\26hpf\Output-02-14-2019\AT_3_26hpf.psi complete
Write to .\data\26hpf\Output-02-14-2019\ZRF_3_26hpf.psi complete
```

In [24]:

```
model = save_model(k,mm,model)
```

# 13¶

In [25]:

```
k,df,Ldf,ax = start('13')
```

In [26]:

```
save_both(k,df,Ldf[0])
```

```
Write to .\data\26hpf\Output-02-14-2019\AT_13_26hpf.psi complete
Write to .\data\26hpf\Output-02-14-2019\ZRF_13_26hpf.psi complete
```

In [27]:

```
mm = fit_model(ax[0,1],df)
```

In [28]:

```
model = save_model(k,mm,model)
```

# 12¶

In [29]:

```
k,df,Ldf,ax = start('12')
```

In [30]:

```
mm = fit_model(ax[0,1],df)
```

In [31]:

```
save_both(k,df,Ldf[0])
```

```
Write to .\data\26hpf\Output-02-14-2019\AT_12_26hpf.psi complete
Write to .\data\26hpf\Output-02-14-2019\ZRF_12_26hpf.psi complete
```

In [32]:

```
model = save_model(k,mm,model)
```

# 7¶

In [33]:

```
k,df,Ldf,ax = start('7')
```

In [34]:

```
df1,Ldf1,pts,ax = ut.check_pts(df,Ldf,'z')
```

In [35]:

```
pts.iloc[0].x = 28
pts.iloc[0].z = 15
ax[0,1].scatter(pts.x,pts.z,c='y')
pts
```

Out[35]:

|  | x | z |
| --- | --- | --- |
| 0 | 28.0000 | 15.000000 |
| 1 | -28.5364 | 11.898168 |

In [36]:

```
df2,Ldf2,ax = ut.revise_pts(df,Ldf,'z',pts=pts)
```

In [37]:

```
df3,Ldf3,mm,ax = ut.ch_vertex(df2,Ldf2)
```

In [38]:

```
save_both(k,df3,Ldf3[0])
```

```
Write to .\data\26hpf\Output-02-14-2019\AT_7_26hpf.psi complete
Write to .\data\26hpf\Output-02-14-2019\ZRF_7_26hpf.psi complete
```

In [39]:

```
model = save_model(k,mm,model)
```

# 4¶

In [40]:

```
k,df,Ldf,ax = start('4')
```

In [41]:

```
mm = fit_model(ax[0,1],df)
```

In [42]:

```
save_both(k,df,Ldf[0])
```

```
Write to .\data\26hpf\Output-02-14-2019\AT_4_26hpf.psi complete
Write to .\data\26hpf\Output-02-14-2019\ZRF_4_26hpf.psi complete
```

In [43]:

```
model = save_model(k,mm,model)
```

# 1¶

In [44]:

```
k,df,Ldf,ax = start('1')
```

In [45]:

```
mm = fit_model(ax[0,1],df)
```

In [46]:

```
save_both(k,df,Ldf[0])
```

```
Write to .\data\26hpf\Output-02-14-2019\AT_1_26hpf.psi complete
Write to .\data\26hpf\Output-02-14-2019\ZRF_1_26hpf.psi complete
```

In [47]:

```
model = save_model(k,mm,model)
```

# 10¶

In [48]:

```
k,df,Ldf,ax = start('10')
```

In [49]:

```
mm = fit_model(ax[0,1],df)
```

In [50]:

```
save_both(k,df,Ldf[0])
```

```
Write to .\data\26hpf\Output-02-14-2019\AT_10_26hpf.psi complete
Write to .\data\26hpf\Output-02-14-2019\ZRF_10_26hpf.psi complete
```

In [51]:

```
model = save_model(k,mm,model)
```

# 64¶

In [52]:

```
k,df,Ldf,ax = start('64')
```

In [53]:

```
mm = fit_model(ax[0,1],df)
```

In [54]:

```
save_both(k,df,Ldf[0])
```

```
Write to .\data\26hpf\Output-02-14-2019\AT_64_26hpf.psi complete
Write to .\data\26hpf\Output-02-14-2019\ZRF_64_26hpf.psi complete
```

In [55]:

```
model = save_model(k,mm,model)
```

# 14¶

In [56]:

```
k,df,Ldf,ax = start('14')
```

In [57]:

```
mm = fit_model(ax[0,1],df)
```

In [58]:

```
save_both(k,df,Ldf[0])
```

```
Write to .\data\26hpf\Output-02-14-2019\AT_14_26hpf.psi complete
Write to .\data\26hpf\Output-02-14-2019\ZRF_14_26hpf.psi complete
```

In [59]:

```
model = save_model(k,mm,model)
```

# 9¶

In [60]:

```
k,df,Ldf,ax = start('9')
```

In [61]:

```
df1,Ldf1,pts,ax = ut.check_pts(df,Ldf,'z')
```

In [62]:

```
df2,Ldf2,mm,ax = ut.ch_vertex(df1,Ldf1)
```

In [63]:

```
save_both(k,df2,Ldf2[0])
```

```
Write to .\data\26hpf\Output-02-14-2019\AT_9_26hpf.psi complete
Write to .\data\26hpf\Output-02-14-2019\ZRF_9_26hpf.psi complete
```

In [64]:

```
model = save_model(k,mm,model)
```

# 2¶

In [65]:

```
k,df,Ldf,ax = start('2')
```

In [66]:

```
mm = fit_model(ax[0,1],df)
```

In [67]:

```
save_both(k,df,Ldf[0])
```

```
Write to .\data\26hpf\Output-02-14-2019\AT_2_26hpf.psi complete
Write to .\data\26hpf\Output-02-14-2019\ZRF_2_26hpf.psi complete
```

In [68]:

```
model = save_model(k,mm,model)
```

# 5¶

In [69]:

```
k,df,Ldf,ax = start('5')
```

In [70]:

```
mm = fit_model(ax[0,1],df)
```

In [71]:

```
save_both(k,df,Ldf[0])
```

```
Write to .\data\26hpf\Output-02-14-2019\AT_5_26hpf.psi complete
Write to .\data\26hpf\Output-02-14-2019\ZRF_5_26hpf.psi complete
```

In [72]:

```
model = save_model(k,mm,model)
```

# 15¶

In [73]:

```
k,df,Ldf,ax = start('15')
```

In [74]:

```
pts = pick_pts(-25,13,0,-2,28,15)
ax[0,1].scatter(pts.x,pts.z,c='m',s=50)
```

Out[74]:

```
<matplotlib.collections.PathCollection at 0x1bf102ee780>
```

In [75]:

```
df1,Ldf1,mm,ax = ut.ch_vertex(df,Ldf,pts=pts)
```

In [76]:

```
save_both(k,df1,Ldf1[0])
```

```
Write to .\data\26hpf\Output-02-14-2019\AT_15_26hpf.psi complete
Write to .\data\26hpf\Output-02-14-2019\ZRF_15_26hpf.psi complete
```

In [77]:

```
model = save_model(k,mm,model)
```

# 8¶

In [78]:

```
k,df,Ldf,ax = start('8')
```

In [79]:

```
df1,Ldf1,pts,ax = ut.check_pts(df,Ldf,'z')
```

In [80]:

```
df2,Ldf2,mm,ax = ut.ch_vertex(df1,Ldf1)
```

In [81]:

```
save_both(k,df2,Ldf2[0])
```

```
Write to .\data\26hpf\Output-02-14-2019\AT_8_26hpf.psi complete
Write to .\data\26hpf\Output-02-14-2019\ZRF_8_26hpf.psi complete
```

In [82]:

```
model = save_model(k,mm,model)
```

# 6¶

In [83]:

```
k,df,Ldf,ax = start('6')
```

In [84]:

```
mm = fit_model(ax[0,1],df)
```

In [85]:

```
pts = pick_pts(-28,16,0,-1,30,13)
ax[0,1].scatter(pts.x,pts.z,c='m',s=50)
```

Out[85]:

```
<matplotlib.collections.PathCollection at 0x1bf37d88d30>
```

In [86]:

```
df1,Ldf1,mm,ax = ut.ch_vertex(df,Ldf,pts=pts)
```

In [87]:

```
save_both(k,df1,Ldf1[0])
```

```
Write to .\data\26hpf\Output-02-14-2019\AT_6_26hpf.psi complete
Write to .\data\26hpf\Output-02-14-2019\ZRF_6_26hpf.psi complete
```

In [88]:

```
model = save_model(k,mm,model)
```

# 11¶

In [89]:

```
k,df,Ldf,ax = start('11')
```

In [90]:

```
df1,Ldf1,pts,ax = ut.check_pts(df,Ldf,'z')
```

In [91]:

```
pts.iloc[1].x = 24
pts.iloc[1].z = 14
ax[0,1].scatter(pts.x,pts.z,c='y',s=50)
pts
```

Out[91]:

|  | x | z |
| --- | --- | --- |
| 0 | -25.476737 | 17.941593 |
| 1 | 24.000000 | 14.000000 |

In [92]:

```
df2,Ldf2,ax = ut.revise_pts(df,Ldf,'z',pts=pts)
```

In [93]:

```
df3,Ldf3,mm,ax = ut.ch_vertex(df2,Ldf2)
```

In [94]:

```
pts = pick_pts(-24,15,0,-1,26,16)
ax[0,1].scatter(pts.x,pts.z,c='m',s=50)
```

Out[94]:

```
<matplotlib.collections.PathCollection at 0x1c00412a668>
```

In [95]:

```
df4,Ldf4,mm,ax = ut.ch_vertex(df3,Ldf2,pts=pts)
```

In [96]:

```
save_both(k,df4,Ldf4[0])
model = save_model(k,mm,model)
```

```
Write to .\data\26hpf\Output-02-14-2019\AT_11_26hpf.psi complete
Write to .\data\26hpf\Output-02-14-2019\ZRF_11_26hpf.psi complete
```

# 66¶

In [97]:

```
k,df,Ldf,ax = start('66')
```

In [98]:

```
df1,Ldf1,pts,ax = ut.check_pts(df,Ldf,'z')
```

In [99]:

```
pts.iloc[0].z = 22
pts.iloc[1].x = -32
ax[0,1].scatter(pts.x,pts.z,c='y')
pts
```

Out[99]:

|  | x | z |
| --- | --- | --- |
| 0 | 32.42549 | 22.000000 |
| 1 | -32.00000 | 20.119464 |

In [100]:

```
df2,Ldf2,ax = ut.revise_pts(df,Ldf,'z',pts=pts)
```

In [101]:

```
df3,Ldf3,mm,ax = ut.ch_vertex(df2,Ldf2)
```

In [102]:

```
save_both(k,df3,Ldf3[0])
model = save_model(k,mm,model)
```

```
Write to .\data\26hpf\Output-02-14-2019\AT_66_26hpf.psi complete
Write to .\data\26hpf\Output-02-14-2019\ZRF_66_26hpf.psi complete
```

### Save model¶

In [103]:

```
model
```

Out[103]:

|  | a | b | c |
| --- | --- | --- | --- |
| 72 | 0.010760 | 2.129977e-17 | 1.604895e-15 |
| 3 | 0.015594 | 1.290557e-16 | -2.831030e-15 |
| 13 | 0.009664 | -9.035462e-17 | -6.184060e-15 |
| 12 | 0.015505 | -6.485233e-17 | -1.201199e-14 |
| 7 | 0.016419 | -1.036174e-17 | 2.291216e-15 |
| 4 | 0.013391 | 6.121736e-19 | -1.964892e-15 |
| 1 | 0.013496 | 6.928385e-17 | -4.674151e-15 |
| 10 | 0.008401 | 3.526229e-17 | -8.332603e-16 |
| 64 | 0.008415 | 2.266621e-17 | 3.172082e-15 |
| 14 | 0.013801 | 3.301717e-17 | 5.034357e-15 |
| 9 | 0.010984 | 5.337687e-17 | -4.015839e-15 |
| 2 | 0.015657 | -5.224502e-17 | -1.213049e-14 |
| 5 | 0.010840 | 5.833752e-17 | 2.663782e-14 |
| 15 | 0.022776 | -1.811466e-16 | -3.679489e-15 |
| 8 | 0.011622 | -8.047862e-18 | -2.442401e-15 |
| 6 | 0.018514 | 1.280987e-16 | 1.576550e-15 |
| 11 | 0.026410 | -3.338110e-16 | -3.729712e-15 |
| 66 | 0.017254 | 8.384075e-17 | 3.545331e-15 |

In [104]:

```
model.to_csv(os.path.join(outdir,'model.csv'))
```

In [105]:

```
outdir
```

Out[105]:

```
'.\\data\\26hpf\\Output-02-14-2019'
```
