## Supplementary material for "ΔSCOPE: A new method to quantify 3D biological structures and identify differences in zebrafish forebrain development": An archived copy of the DeltaSCOPE code repository.: alignment-28hpf.html


In [1]:

```
import numpy as np
import pandas as pd
import matplotlib.pyplot as plt

import os
import re
from imp import reload
import h5py
```

In [2]:

```
import deltascope as cranium
import deltascope.alignment as ut
```

In [3]:

```
at = ".\\data\\28hpf\\AT\\Prob"
gfap = ".\\data\\28hpf\\Gfap\\Prob-yot-zrf-ilastik"
```

In [4]:

```
outdir = ".\\data\\28hpf\\Output-02-14-2019-yot-ilastik"
```

In [5]:

```
os.mkdir(outdir)
```

In [6]:

```
Dat = {}
for f in os.listdir(at):
    if 'h5' in f:
        num  = re.findall(r'\d+',f.split('.')[0])[-1]
        Dat[num] = os.path.join(at,f)
```

In [7]:

```
Dzrf = {}
for f in os.listdir(gfap):
    if 'h5' in f:
        num  = re.findall(r'\d+',f.split('.')[0])[-1]
        Dzrf[num] = os.path.join(gfap,f)
```

In [8]:

```
Dbat = {}
Dbzrf = {}
```

### Data Preprocessing¶

In [9]:

```
klist = Dat.keys()
```

In [10]:

```
param = {
    'gthresh':0.5,
    'scale':[1,1,1],
    'microns':[0.16,0.16,0.21],
    'mthresh':0.5,
    'radius':10,
    'comp_order':[0,2,1],
    'fit_dim':['x','z'],
    'deg':2
}
```

In [11]:

```
%%time
for k in klist:
    if k not in list(Dbat.keys()):
        Dbat[k] = ut.preprocess(Dat[k],param)
        Dbzrf[k] = ut.preprocess(Dzrf[k],param,pca=Dbat[k].pcamed,mm=Dbat[k].mm,vertex=Dbat[k].vertex)
        print(k)
    else:
        print(k,'already processed')
```

```
C:\Users\zfishlab\AppData\Local\Continuum\anaconda3\envs\test\lib\site-packages\skimage\util\dtype.py:141: UserWarning: Possible precision loss when converting from float32 to uint8
  .format(dtypeobj_in, dtypeobj_out))
```

```
104
```

```
C:\Users\zfishlab\AppData\Local\Continuum\anaconda3\envs\test\lib\site-packages\skimage\util\dtype.py:141: UserWarning: Possible precision loss when converting from float32 to uint8
  .format(dtypeobj_in, dtypeobj_out))
```

```
10
```

```
C:\Users\zfishlab\AppData\Local\Continuum\anaconda3\envs\test\lib\site-packages\skimage\util\dtype.py:141: UserWarning: Possible precision loss when converting from float32 to uint8
  .format(dtypeobj_in, dtypeobj_out))
```

```
112
```

```
C:\Users\zfishlab\AppData\Local\Continuum\anaconda3\envs\test\lib\site-packages\skimage\util\dtype.py:141: UserWarning: Possible precision loss when converting from float32 to uint8
  .format(dtypeobj_in, dtypeobj_out))
```

```
115
```

```
C:\Users\zfishlab\AppData\Local\Continuum\anaconda3\envs\test\lib\site-packages\skimage\util\dtype.py:141: UserWarning: Possible precision loss when converting from float32 to uint8
  .format(dtypeobj_in, dtypeobj_out))
```

```
116
```

```
C:\Users\zfishlab\AppData\Local\Continuum\anaconda3\envs\test\lib\site-packages\skimage\util\dtype.py:141: UserWarning: Possible precision loss when converting from float32 to uint8
  .format(dtypeobj_in, dtypeobj_out))
```

```
117
```

```
C:\Users\zfishlab\AppData\Local\Continuum\anaconda3\envs\test\lib\site-packages\skimage\util\dtype.py:141: UserWarning: Possible precision loss when converting from float32 to uint8
  .format(dtypeobj_in, dtypeobj_out))
```

```
118
```

```
C:\Users\zfishlab\AppData\Local\Continuum\anaconda3\envs\test\lib\site-packages\skimage\util\dtype.py:141: UserWarning: Possible precision loss when converting from float32 to uint8
  .format(dtypeobj_in, dtypeobj_out))
```

```
119
```

```
C:\Users\zfishlab\AppData\Local\Continuum\anaconda3\envs\test\lib\site-packages\skimage\util\dtype.py:141: UserWarning: Possible precision loss when converting from float32 to uint8
  .format(dtypeobj_in, dtypeobj_out))
```

```
120
```

```
C:\Users\zfishlab\AppData\Local\Continuum\anaconda3\envs\test\lib\site-packages\skimage\util\dtype.py:141: UserWarning: Possible precision loss when converting from float32 to uint8
  .format(dtypeobj_in, dtypeobj_out))
```

```
121
```

```
C:\Users\zfishlab\AppData\Local\Continuum\anaconda3\envs\test\lib\site-packages\skimage\util\dtype.py:141: UserWarning: Possible precision loss when converting from float32 to uint8
  .format(dtypeobj_in, dtypeobj_out))
```

```
12
```

```
C:\Users\zfishlab\AppData\Local\Continuum\anaconda3\envs\test\lib\site-packages\skimage\util\dtype.py:141: UserWarning: Possible precision loss when converting from float32 to uint8
  .format(dtypeobj_in, dtypeobj_out))
```

```
14
```

```
C:\Users\zfishlab\AppData\Local\Continuum\anaconda3\envs\test\lib\site-packages\skimage\util\dtype.py:141: UserWarning: Possible precision loss when converting from float32 to uint8
  .format(dtypeobj_in, dtypeobj_out))
```

```
16
```

```
C:\Users\zfishlab\AppData\Local\Continuum\anaconda3\envs\test\lib\site-packages\skimage\util\dtype.py:141: UserWarning: Possible precision loss when converting from float32 to uint8
  .format(dtypeobj_in, dtypeobj_out))
```

```
2
```

```
C:\Users\zfishlab\AppData\Local\Continuum\anaconda3\envs\test\lib\site-packages\skimage\util\dtype.py:141: UserWarning: Possible precision loss when converting from float32 to uint8
  .format(dtypeobj_in, dtypeobj_out))
```

```
4
```

```
C:\Users\zfishlab\AppData\Local\Continuum\anaconda3\envs\test\lib\site-packages\skimage\util\dtype.py:141: UserWarning: Possible precision loss when converting from float32 to uint8
  .format(dtypeobj_in, dtypeobj_out))
```

```
5
```

```
C:\Users\zfishlab\AppData\Local\Continuum\anaconda3\envs\test\lib\site-packages\skimage\util\dtype.py:141: UserWarning: Possible precision loss when converting from float32 to uint8
  .format(dtypeobj_in, dtypeobj_out))
```

```
6
```

```
C:\Users\zfishlab\AppData\Local\Continuum\anaconda3\envs\test\lib\site-packages\skimage\util\dtype.py:141: UserWarning: Possible precision loss when converting from float32 to uint8
  .format(dtypeobj_in, dtypeobj_out))
```

```
8
Wall time: 10min 33s
```

In [12]:

```
def start(k):
    return(ut.start(k,Dbat,[Dbzrf],im=True))
def save_both(k,dfa,dfb):
    ut.save_both(k,dfa,dfb,outdir,'28hpf')
```

In [13]:

```
model = pd.DataFrame({'a':[],'b':[],'c':[]})
def save_model(k,mm,model):
    row = pd.Series({'a':mm[0],'b':mm[1],'c':mm[2]},name=k)
    model = model.append(row)
    return(model)
```

In [14]:

```
def fit_model(axi,df,mm=None):
    if mm == None:
        mm = np.polyfit(df.x,df.z,2)
    p = np.poly1d(mm)
    xrange = np.arange(np.min(df.x),np.max(df.x))
    axi.plot(xrange,p(xrange),c='m')
    return(mm)
```

In [15]:

```
def pick_pts(x1,z1,vx,vz,x2,z2):
    pts = pd.DataFrame({'x':[x1,vx,x2],'z':[z1,vz,z2]})
    return(pts)
```

# 104¶

In [16]:

```
k,df,Ldf,ax = start('104')
```

In [17]:

```
mm = fit_model(ax[0,1],df)
```

In [18]:

```
model = save_model(k,mm,model)
```

# 115¶

In [19]:

```
k,df,Ldf,ax = start('115')
```

In [20]:

```
df1,Ldf1 = ut.zyswitch(df,Ldf)
ax = ut.make_graph([df1]+Ldf1)
```

In [21]:

```
df2,Ldf2,mm,ax = ut.ch_vertex(df1,Ldf1)
```

In [22]:

```
save_both(k,df2,Ldf2[0])
```

```
Write to .\data\28hpf\Output-02-14-2019-yot-ilastik\AT_115_28hpf.psi complete
Write to .\data\28hpf\Output-02-14-2019-yot-ilastik\ZRF_115_28hpf.psi complete
```

In [23]:

```
model = save_model(k,mm,model)
```

# 10¶

In [24]:

```
k,df,Ldf,ax = start('10')
```

In [25]:

```
df1,Ldf1,pts,ax = ut.check_pts(df,Ldf,'z')
```

In [26]:

```
pts.iloc[1].x = -46
pts.iloc[1].z = 30
pts.iloc[0].z = 38
ax[0,1].scatter(pts.x,pts.z,c='y')
pts
```

Out[26]:

|  | x | z |
| --- | --- | --- |
| 0 | 50.036549 | 38.0 |
| 1 | -46.000000 | 30.0 |

In [27]:

```
df2,Ldf2,ax = ut.revise_pts(df,Ldf,'z',pts=pts)
```

In [28]:

```
df3,Ldf3,mm,ax = ut.ch_vertex(df2,Ldf2)
```

In [29]:

```
save_both(k,df3,Ldf3[0])
```

```
Write to .\data\28hpf\Output-02-14-2019-yot-ilastik\AT_10_28hpf.psi complete
Write to .\data\28hpf\Output-02-14-2019-yot-ilastik\ZRF_10_28hpf.psi complete
```

In [30]:

```
model = save_model(k,mm,model)
```

# 116¶

In [31]:

```
k,df,Ldf,ax = start('116')
```

In [32]:

```
df1,Ldf1,ax,p = ut.check_yz(df,Ldf)
```

In [33]:

```
mm = [-1,0]
xrange = np.arange(-10,10)
ax[0,2].plot(xrange,np.poly1d(mm)(xrange),c='g')
```

Out[33]:

```
[<matplotlib.lines.Line2D at 0x2096a9ff6d8>]
```

In [34]:

```
df2,Ldf2,ax,p = ut.check_yz(df,Ldf,mm=mm)
```

In [35]:

```
df3,Ldf3,mm,ax = ut.ch_vertex(df2,Ldf2)
```

In [36]:

```
save_both(k,df3,Ldf3[0])
```

```
Write to .\data\28hpf\Output-02-14-2019-yot-ilastik\AT_116_28hpf.psi complete
Write to .\data\28hpf\Output-02-14-2019-yot-ilastik\ZRF_116_28hpf.psi complete
```

In [37]:

```
model = save_model(k,mm,model)
```

# 16¶

In [38]:

```
k,df,Ldf,ax = start('16')
```

In [39]:

```
df1,Ldf1,pts,ax = ut.check_pts(df,Ldf,'z')
```

In [40]:

```
df2,Ldf2,mm,ax = ut.ch_vertex(df1,Ldf1)
```

In [41]:

```
save_both(k,df2,Ldf2[0])
```

```
Write to .\data\28hpf\Output-02-14-2019-yot-ilastik\AT_16_28hpf.psi complete
Write to .\data\28hpf\Output-02-14-2019-yot-ilastik\ZRF_16_28hpf.psi complete
```

In [42]:

```
model = save_model(k,mm,model)
```

# 6¶

In [43]:

```
k,df,Ldf,ax = start('6')
```

In [44]:

```
mm = fit_model(ax[0,1],df)
```

In [45]:

```
save_both(k,df,Ldf[0])
```

```
Write to .\data\28hpf\Output-02-14-2019-yot-ilastik\AT_6_28hpf.psi complete
Write to .\data\28hpf\Output-02-14-2019-yot-ilastik\ZRF_6_28hpf.psi complete
```

In [46]:

```
model = save_model(k,mm,model)
```

# 119¶

In [47]:

```
k,df,Ldf,ax = start('119')
```

In [48]:

```
df1,Ldf1,ax,p = ut.check_yz(df,Ldf)
```

In [49]:

```
mm = [1.2,0]
xrange = np.arange(-5,5)
ax[0,2].plot(xrange,np.poly1d(mm)(xrange),c='y')
```

Out[49]:

```
[<matplotlib.lines.Line2D at 0x2096a9f7208>]
```

In [50]:

```
df2,Ldf2,ax,p = ut.check_yz(df,Ldf,mm=mm)
```

In [51]:

```
df3,Ldf3,mm,ax = ut.ch_vertex(df2,Ldf2)
```

In [52]:

```
save_both(k,df3,Ldf3[0])
```

```
Write to .\data\28hpf\Output-02-14-2019-yot-ilastik\AT_119_28hpf.psi complete
Write to .\data\28hpf\Output-02-14-2019-yot-ilastik\ZRF_119_28hpf.psi complete
```

In [53]:

```
model = save_model(k,mm,model)
```

# 120¶

In [54]:

```
k,df,Ldf,ax = start('120')
```

In [55]:

```
mm = [-1/2,0]
xrange = np.arange(-5,5)
ax[0,2].plot(xrange,np.poly1d(mm)(xrange),c='y')
```

Out[55]:

```
[<matplotlib.lines.Line2D at 0x2089a783860>]
```

In [56]:

```
df1,Ldf1,ax,p = ut.check_yz(df,Ldf,mm=mm)
```

In [57]:

```
df2,Ldf2,mm,ax = ut.ch_vertex(df1,Ldf1)
```

In [58]:

```
pts = pick_pts(-35,15,0,-2,40,16)
ax[0,1].scatter(pts.x,pts.z,c='m',s=50)
```

Out[58]:

```
<matplotlib.collections.PathCollection at 0x20a32aba208>
```

In [59]:

```
df3,Ldf3,mm,ax = ut.ch_vertex(df1,Ldf1,pts=pts)
```

In [60]:

```
save_both(k,df3,Ldf3[0])
```

```
Write to .\data\28hpf\Output-02-14-2019-yot-ilastik\AT_120_28hpf.psi complete
Write to .\data\28hpf\Output-02-14-2019-yot-ilastik\ZRF_120_28hpf.psi complete
```

In [61]:

```
model = save_model(k,mm,model)
```

# 14¶

In [62]:

```
k,df,Ldf,ax = start('14')
```

In [63]:

```
df1,Ldf1,pts,ax = ut.check_pts(df,Ldf,'z')
```

In [64]:

```
df2,Ldf2,mm,ax = ut.ch_vertex(df1,Ldf1)
```

In [65]:

```
pts = pick_pts(-20,15,0,0,22,15)
ax[0,1].scatter(pts.x,pts.z,c='m',s=50)
```

Out[65]:

```
<matplotlib.collections.PathCollection at 0x2070e010898>
```

In [66]:

```
df3,Ldf3,mm,ax = ut.ch_vertex(df2,Ldf2,pts=pts)
```

In [67]:

```
save_both(k,df3,Ldf3[0])
```

```
Write to .\data\28hpf\Output-02-14-2019-yot-ilastik\AT_14_28hpf.psi complete
Write to .\data\28hpf\Output-02-14-2019-yot-ilastik\ZRF_14_28hpf.psi complete
```

In [68]:

```
model = save_model(k,mm,model)
```

# 8¶

In [69]:

```
k,df,Ldf,ax = start('8')
```

In [70]:

```
mm = [-2/5,0]
xrange = np.arange(-5,15)
ax[0,2].plot(xrange,np.poly1d(mm)(xrange),c='y')
```

Out[70]:

```
[<matplotlib.lines.Line2D at 0x2091d3636d8>]
```

In [71]:

```
df1,Ldf1,ax,p = ut.check_yz(df,Ldf,mm=mm)
```

In [72]:

```
df2,Ldf2,mm,ax = ut.ch_vertex(df1,Ldf1)
```

In [73]:

```
pts = pick_pts(-48,20,0,-2,48,20)
ax[0,1].scatter(pts.x,pts.z,c='m',s=50)
```

Out[73]:

```
<matplotlib.collections.PathCollection at 0x20a35762278>
```

In [74]:

```
df3,Ldf3,mm,ax = ut.ch_vertex(df2,Ldf2,pts=pts)
```

In [75]:

```
save_both(k,df3,Ldf3[0])
```

```
Write to .\data\28hpf\Output-02-14-2019-yot-ilastik\AT_8_28hpf.psi complete
Write to .\data\28hpf\Output-02-14-2019-yot-ilastik\ZRF_8_28hpf.psi complete
```

In [76]:

```
model = save_model(k,mm,model)
```

# 121¶

In [77]:

```
k,df,Ldf,ax = start('121')
```

In [78]:

```
df1,Ldf1 = ut.zyswitch(df,Ldf)
ax = ut.make_graph([df1]+Ldf1)
```

In [79]:

```
df2,Ldf2,mm,ax = ut.ch_vertex(df1,Ldf1)
```

In [80]:

```
save_both(k,df2,Ldf2[0])
```

```
Write to .\data\28hpf\Output-02-14-2019-yot-ilastik\AT_121_28hpf.psi complete
Write to .\data\28hpf\Output-02-14-2019-yot-ilastik\ZRF_121_28hpf.psi complete
```

In [81]:

```
model = save_model(k,mm,model)
```

# 12¶

In [82]:

```
k,df,Ldf,ax = start('12')
```

In [83]:

```
save_both(k,df,Ldf[0])
```

```
Write to .\data\28hpf\Output-02-14-2019-yot-ilastik\AT_12_28hpf.psi complete
Write to .\data\28hpf\Output-02-14-2019-yot-ilastik\ZRF_12_28hpf.psi complete
```

In [84]:

```
mm = fit_model(ax[0,1],df)
```

In [85]:

```
model = save_model(k,mm,model)
```

# 117¶

In [86]:

```
k,df,Ldf,ax = start('117')
```

In [87]:

```
df1,Ldf1 = ut.zyswitch(df,Ldf)
ax = ut.make_graph([df1]+Ldf1)
```

In [88]:

```
df2,Ldf2,mm,ax = ut.ch_vertex(df1,Ldf1)
```

In [89]:

```
save_both(k,df2,Ldf2[0])
```

```
Write to .\data\28hpf\Output-02-14-2019-yot-ilastik\AT_117_28hpf.psi complete
Write to .\data\28hpf\Output-02-14-2019-yot-ilastik\ZRF_117_28hpf.psi complete
```

In [90]:

```
model = save_model(k,mm,model)
```

# 5¶

In [91]:

```
k,df,Ldf,ax = start('5')
```

In [92]:

```
df1,Ldf1 = ut.zyswitch(df,Ldf)
ax = ut.make_graph([df1]+Ldf1)
```

In [93]:

```
pts = pick_pts(-35,10,-2,-6,30,10)
ax[0,1].scatter(pts.x,pts.z,c='m',s=50)
```

Out[93]:

```
<matplotlib.collections.PathCollection at 0x20aa9b90d68>
```

In [94]:

```
df2,Ldf2,mm,ax = ut.ch_vertex(df1,Ldf1,pts=pts)
```

In [95]:

```
save_both(k,df2,Ldf2[0])
```

```
Write to .\data\28hpf\Output-02-14-2019-yot-ilastik\AT_5_28hpf.psi complete
Write to .\data\28hpf\Output-02-14-2019-yot-ilastik\ZRF_5_28hpf.psi complete
```

In [96]:

```
model = save_model(k,mm,model)
```

# 4¶

In [97]:

```
k,df,Ldf,ax = start('4')
```

In [98]:

```
mm = fit_model(ax[0,1],df)
```

In [99]:

```
save_both(k,df,Ldf[0])
```

```
Write to .\data\28hpf\Output-02-14-2019-yot-ilastik\AT_4_28hpf.psi complete
Write to .\data\28hpf\Output-02-14-2019-yot-ilastik\ZRF_4_28hpf.psi complete
```

In [100]:

```
model = save_model(k,mm,model)
```

# 112¶

In [101]:

```
k,df,Ldf,ax = start('112')
```

In [102]:

```
mm = fit_model(ax[0,1],df)
```

In [103]:

```
save_both(k,df,Ldf[0])
```

```
Write to .\data\28hpf\Output-02-14-2019-yot-ilastik\AT_112_28hpf.psi complete
Write to .\data\28hpf\Output-02-14-2019-yot-ilastik\ZRF_112_28hpf.psi complete
```

In [104]:

```
model = save_model(k,mm,model)
```

# 118¶

In [105]:

```
k,df,Ldf,ax = start('118')
```

In [106]:

```
mm = [-2/5,0]
xrange = np.arange(-5,10)
ax[0,2].plot(xrange,np.poly1d(mm)(xrange),c='y')
```

Out[106]:

```
[<matplotlib.lines.Line2D at 0x20aa97d34a8>]
```

In [107]:

```
df1,Ldf1,ax,p = ut.check_yz(df,Ldf,mm=mm)
```

In [108]:

```
df2,Ldf2,mm,ax = ut.ch_vertex(df1,Ldf1)
```

In [109]:

```
save_both(k,df2,Ldf2[0])
```

```
Write to .\data\28hpf\Output-02-14-2019-yot-ilastik\AT_118_28hpf.psi complete
Write to .\data\28hpf\Output-02-14-2019-yot-ilastik\ZRF_118_28hpf.psi complete
```

In [110]:

```
model = save_model(k,mm,model)
```

# 2¶

In [111]:

```
k,df,Ldf,ax = start('2')
```

In [112]:

```
mm = fit_model(ax[0,1],df)
```

In [113]:

```
save_both(k,df,Ldf[0])
```

```
Write to .\data\28hpf\Output-02-14-2019-yot-ilastik\AT_2_28hpf.psi complete
Write to .\data\28hpf\Output-02-14-2019-yot-ilastik\ZRF_2_28hpf.psi complete
```

In [114]:

```
model = save_model(k,mm,model)
```

### Model¶

In [115]:

```
model
```

Out[115]:

|  | a | b | c |
| --- | --- | --- | --- |
| 104 | 0.010879 | 2.779127e-17 | 3.700548e-15 |
| 115 | 0.004570 | -2.487537e-18 | 2.795294e-15 |
| 10 | 0.012008 | -1.843857e-16 | -3.456267e-15 |
| 116 | 0.008391 | -7.337595e-18 | -2.478101e-17 |
| 16 | 0.016922 | 1.489012e-16 | -6.835019e-15 |
| 6 | 0.018744 | 1.090573e-16 | 2.577010e-15 |
| 119 | 0.011514 | 3.235386e-17 | -3.617943e-15 |
| 120 | 0.012476 | 3.148644e-17 | -5.183311e-15 |
| 14 | 0.034091 | 2.970435e-16 | -4.101984e-15 |
| 8 | 0.009549 | 3.889675e-17 | -3.076740e-15 |
| 121 | 0.008379 | 3.764361e-17 | 6.024008e-15 |
| 12 | 0.014052 | -5.706871e-17 | 2.742478e-15 |
| 117 | 0.008438 | -5.317300e-17 | 4.287112e-16 |
| 5 | 0.015152 | -1.086983e-17 | -7.179060e-15 |
| 4 | 0.018001 | 1.627579e-16 | -3.179114e-15 |
| 112 | 0.013030 | -4.923055e-17 | 6.889791e-15 |
| 118 | 0.006414 | -1.021713e-18 | 2.909860e-16 |
| 2 | 0.010410 | -3.945576e-17 | -6.589945e-15 |

In [116]:

```
model.to_csv(os.path.join(outdir,'model.csv'))
```

In [117]:

```
outdir
```

Out[117]:

```
'.\\data\\28hpf\\Output-02-14-2019-yot-ilastik'
```
