## Supplementary material for "ΔSCOPE: A new method to quantify 3D biological structures and identify differences in zebrafish forebrain development": An archived copy of the DeltaSCOPE code repository.: alignment-28hpf.ipynb

```
# %% [code]
import numpy as np
import pandas as pd
import matplotlib.pyplot as plt

import os
import re
from imp import reload
import h5py

# %% [code]
import deltascope as cranium
import deltascope.alignment as ut

# %% [code]
at = ".\\data\\28hpf\\AT\\Prob"
gfap = ".\\data\\28hpf\\Gfap\\Prob-yot-zrf-ilastik"

# %% [code]
outdir = ".\\data\\28hpf\\Output-02-14-2019-yot-ilastik"

# %% [code]
os.mkdir(outdir)

# %% [code]
Dat = {}
for f in os.listdir(at):
    if 'h5' in f:
        num  = re.findall(r'\d+',f.split('.')[0])[-1]
        Dat[num] = os.path.join(at,f)

# %% [code]
Dzrf = {}
for f in os.listdir(gfap):
    if 'h5' in f:
        num  = re.findall(r'\d+',f.split('.')[0])[-1]
        Dzrf[num] = os.path.join(gfap,f)

# %% [code]
Dbat = {}
Dbzrf = {}

# %% [markdown]
# Data Preprocessing

# %% [code]
klist = Dat.keys()

# %% [code]
param = {
    'gthresh':0.5,
    'scale':[1,1,1],
    'microns':[0.16,0.16,0.21],
    'mthresh':0.5,
    'radius':10,
    'comp_order':[0,2,1],
    'fit_dim':['x','z'],
    'deg':2
}

# %% [code]
%%time
for k in klist:
    if k not in list(Dbat.keys()):
        Dbat[k] = ut.preprocess(Dat[k],param)
        Dbzrf[k] = ut.preprocess(Dzrf[k],param,pca=Dbat[k].pcamed,mm=Dbat[k].mm,vertex=Dbat[k].vertex)
        print(k)
    else:
        print(k,'already processed')

# %% [code]
def start(k):
    return(ut.start(k,Dbat,[Dbzrf],im=True))
def save_both(k,dfa,dfb):
    ut.save_both(k,dfa,dfb,outdir,'28hpf')

# %% [code]
model = pd.DataFrame({'a':[],'b':[],'c':[]})
def save_model(k,mm,model):
    row = pd.Series({'a':mm[0],'b':mm[1],'c':mm[2]},name=k)
    model = model.append(row)
    return(model)

# %% [code]
def fit_model(axi,df,mm=None):
    if mm == None:
        mm = np.polyfit(df.x,df.z,2)
    p = np.poly1d(mm)
    xrange = np.arange(np.min(df.x),np.max(df.x))
    axi.plot(xrange,p(xrange),c='m')
    return(mm)

# %% [code]
def pick_pts(x1,z1,vx,vz,x2,z2):
    pts = pd.DataFrame({'x':[x1,vx,x2],'z':[z1,vz,z2]})
    return(pts)

# %% [markdown]
# 104

# %% [code]
k,df,Ldf,ax = start('104')

# %% [code]
mm = fit_model(ax[0,1],df)

# %% [code]
model = save_model(k,mm,model)

# %% [markdown]
# 115

# %% [code]
k,df,Ldf,ax = start('115')

# %% [code]
df1,Ldf1 = ut.zyswitch(df,Ldf)
ax = ut.make_graph([df1]+Ldf1)

# %% [code]
df2,Ldf2,mm,ax = ut.ch_vertex(df1,Ldf1)

# %% [code]
save_both(k,df2,Ldf2[0])

# %% [code]
model = save_model(k,mm,model)

# %% [markdown]
# 10

# %% [code]
k,df,Ldf,ax = start('10')

# %% [code]
df1,Ldf1,pts,ax = ut.check_pts(df,Ldf,'z')

# %% [code]
pts.iloc[1].x = -46
pts.iloc[1].z = 30
pts.iloc[0].z = 38
ax[0,1].scatter(pts.x,pts.z,c='y')
pts

# %% [code]
df2,Ldf2,ax = ut.revise_pts(df,Ldf,'z',pts=pts)

# %% [code]
df3,Ldf3,mm,ax = ut.ch_vertex(df2,Ldf2)

# %% [code]
save_both(k,df3,Ldf3[0])

# %% [code]
model = save_model(k,mm,model)

# %% [markdown]
# 116

# %% [code]
k,df,Ldf,ax = start('116')

# %% [code]
df1,Ldf1,ax,p = ut.check_yz(df,Ldf)

# %% [code]
mm = [-1,0]
xrange = np.arange(-10,10)
ax[0,2].plot(xrange,np.poly1d(mm)(xrange),c='g')

# %% [code]
df2,Ldf2,ax,p = ut.check_yz(df,Ldf,mm=mm)

# %% [code]
df3,Ldf3,mm,ax = ut.ch_vertex(df2,Ldf2)

# %% [code]
save_both(k,df3,Ldf3[0])

# %% [code]
model = save_model(k,mm,model)

# %% [markdown]
# 16

# %% [code]
k,df,Ldf,ax = start('16')

# %% [code]
df1,Ldf1,pts,ax = ut.check_pts(df,Ldf,'z')

# %% [code]
df2,Ldf2,mm,ax = ut.ch_vertex(df1,Ldf1)

# %% [code]
save_both(k,df2,Ldf2[0])

# %% [code]
model = save_model(k,mm,model)

# %% [markdown]
# 6

# %% [code]
k,df,Ldf,ax = start('6')

# %% [code]
mm = fit_model(ax[0,1],df)

# %% [code]
save_both(k,df,Ldf[0])

# %% [code]
model = save_model(k,mm,model)

# %% [markdown]
# 119

# %% [code]
k,df,Ldf,ax = start('119')

# %% [code]
df1,Ldf1,ax,p = ut.check_yz(df,Ldf)

# %% [code]
mm = [1.2,0]
xrange = np.arange(-5,5)
ax[0,2].plot(xrange,np.poly1d(mm)(xrange),c='y')

# %% [code]
df2,Ldf2,ax,p = ut.check_yz(df,Ldf,mm=mm)

# %% [code]
df3,Ldf3,mm,ax = ut.ch_vertex(df2,Ldf2)

# %% [code]
save_both(k,df3,Ldf3[0])

# %% [code]
model = save_model(k,mm,model)

# %% [markdown]
# 120

# %% [code]
k,df,Ldf,ax = start('120')

# %% [code]
mm = [-1/2,0]
xrange = np.arange(-5,5)
ax[0,2].plot(xrange,np.poly1d(mm)(xrange),c='y')

# %% [code]
df1,Ldf1,ax,p = ut.check_yz(df,Ldf,mm=mm)

# %% [code]
df2,Ldf2,mm,ax = ut.ch_vertex(df1,Ldf1)

# %% [code]
pts = pick_pts(-35,15,0,-2,40,16)
ax[0,1].scatter(pts.x,pts.z,c='m',s=50)

# %% [code]
df3,Ldf3,mm,ax = ut.ch_vertex(df1,Ldf1,pts=pts)

# %% [code]
save_both(k,df3,Ldf3[0])

# %% [code]
model = save_model(k,mm,model)

# %% [markdown]
# 14

# %% [code]
k,df,Ldf,ax = start('14')

# %% [code]
df1,Ldf1,pts,ax = ut.check_pts(df,Ldf,'z')

# %% [code]
df2,Ldf2,mm,ax = ut.ch_vertex(df1,Ldf1)

# %% [code]
pts = pick_pts(-20,15,0,0,22,15)
ax[0,1].scatter(pts.x,pts.z,c='m',s=50)

# %% [code]
df3,Ldf3,mm,ax = ut.ch_vertex(df2,Ldf2,pts=pts)

# %% [code]
save_both(k,df3,Ldf3[0])

# %% [code]
model = save_model(k,mm,model)

# %% [markdown]
# 8

# %% [code]
k,df,Ldf,ax = start('8')

# %% [code]
mm = [-2/5,0]
xrange = np.arange(-5,15)
ax[0,2].plot(xrange,np.poly1d(mm)(xrange),c='y')

# %% [code]
df1,Ldf1,ax,p = ut.check_yz(df,Ldf,mm=mm)

# %% [code]
df2,Ldf2,mm,ax = ut.ch_vertex(df1,Ldf1)

# %% [code]
pts = pick_pts(-48,20,0,-2,48,20)
ax[0,1].scatter(pts.x,pts.z,c='m',s=50)

# %% [code]
df3,Ldf3,mm,ax = ut.ch_vertex(df2,Ldf2,pts=pts)

# %% [code]
save_both(k,df3,Ldf3[0])

# %% [code]
model = save_model(k,mm,model)

# %% [markdown]
# 121

# %% [code]
k,df,Ldf,ax = start('121')

# %% [code]
df1,Ldf1 = ut.zyswitch(df,Ldf)
ax = ut.make_graph([df1]+Ldf1)

# %% [code]
df2,Ldf2,mm,ax = ut.ch_vertex(df1,Ldf1)

# %% [code]
save_both(k,df2,Ldf2[0])

# %% [code]
model = save_model(k,mm,model)

# %% [markdown]
# 12

# %% [code]
k,df,Ldf,ax = start('12')

# %% [code]
save_both(k,df,Ldf[0])

# %% [code]
mm = fit_model(ax[0,1],df)

# %% [code]
model = save_model(k,mm,model)

# %% [markdown]
# 117

# %% [code]
k,df,Ldf,ax = start('117')

# %% [code]
df1,Ldf1 = ut.zyswitch(df,Ldf)
ax = ut.make_graph([df1]+Ldf1)

# %% [code]
df2,Ldf2,mm,ax = ut.ch_vertex(df1,Ldf1)

# %% [code]
save_both(k,df2,Ldf2[0])

# %% [code]
model = save_model(k,mm,model)

# %% [markdown]
# 5

# %% [code]
k,df,Ldf,ax = start('5')

# %% [code]
df1,Ldf1 = ut.zyswitch(df,Ldf)
ax = ut.make_graph([df1]+Ldf1)

# %% [code]
pts = pick_pts(-35,10,-2,-6,30,10)
ax[0,1].scatter(pts.x,pts.z,c='m',s=50)

# %% [code]
df2,Ldf2,mm,ax = ut.ch_vertex(df1,Ldf1,pts=pts)

# %% [code]
save_both(k,df2,Ldf2[0])

# %% [code]
model = save_model(k,mm,model)

# %% [markdown]
# 4

# %% [code]
k,df,Ldf,ax = start('4')

# %% [code]
mm = fit_model(ax[0,1],df)

# %% [code]
save_both(k,df,Ldf[0])

# %% [code]
model = save_model(k,mm,model)

# %% [markdown]
# 112

# %% [code]
k,df,Ldf,ax = start('112')

# %% [code]
mm = fit_model(ax[0,1],df)

# %% [code]
save_both(k,df,Ldf[0])

# %% [code]
model = save_model(k,mm,model)

# %% [markdown]
# 118

# %% [code]
k,df,Ldf,ax = start('118')

# %% [code]
mm = [-2/5,0]
xrange = np.arange(-5,10)
ax[0,2].plot(xrange,np.poly1d(mm)(xrange),c='y')

# %% [code]
df1,Ldf1,ax,p = ut.check_yz(df,Ldf,mm=mm)

# %% [code]
df2,Ldf2,mm,ax = ut.ch_vertex(df1,Ldf1)

# %% [code]
save_both(k,df2,Ldf2[0])

# %% [code]
model = save_model(k,mm,model)

# %% [markdown]
# 2

# %% [code]
k,df,Ldf,ax = start('2')

# %% [code]
mm = fit_model(ax[0,1],df)

# %% [code]
save_both(k,df,Ldf[0])

# %% [code]
model = save_model(k,mm,model)

# %% [markdown]
# Model

# %% [code]
model

# %% [code]
model.to_csv(os.path.join(outdir,'model.csv'))

# %% [code]
outdir
```
