## Supplementary material for "ΔSCOPE: A new method to quantify 3D biological structures and identify differences in zebrafish forebrain development": An archived copy of the DeltaSCOPE code repository.: alignment-30hpf.html


In [1]:

```
import numpy as np
import pandas as pd
import matplotlib.pyplot as plt

import os
import re
from imp import reload
import h5py
```

In [2]:

```
import deltascope as cranium
import deltascope.alignment as ut
```

In [3]:

```
at = ".\\Data\\30hpf\\AT\\Prob"
gfap = ".\\Data\\30hpf\\Gfap\\Prob"
```

In [4]:

```
outdir = ".\\Data\\30hpf\\Output-02-14-2019"
```

In [5]:

```
os.mkdir(outdir)
```

In [6]:

```
Dat = {}
for f in os.listdir(at):
    if 'h5' in f:
        num  = re.findall(r'\d+',f.split('.')[0])[-1]
        Dat[num] = os.path.join(at,f)
```

In [7]:

```
Dzrf = {}
for f in os.listdir(gfap):
    if 'h5' in f:
        num  = re.findall(r'\d+',f.split('.')[0])[-1]
        Dzrf[num] = os.path.join(gfap,f)
```

In [8]:

```
Dbat = {}
Dbzrf = {}
```

### Data Preprocessing¶

In [9]:

```
klist = Dat.keys()
```

In [10]:

```
param = {
    'gthresh':0.5,
    'scale':[1,1,1],
    'microns':[0.16,0.16,0.21],
    'mthresh':0.5,
    'radius':10,
    'comp_order':[0,2,1],
    'fit_dim':['x','z'],
    'deg':2
}
```

In [11]:

```
%%time
for k in klist:
    if k not in list(Dbat.keys()):
        Dbat[k] = ut.preprocess(Dat[k],param)
        Dbzrf[k] = ut.preprocess(Dzrf[k],param,pca=Dbat[k].pcamed,mm=Dbat[k].mm,vertex=Dbat[k].vertex)
        print(k)
    else:
        print(k,'already processed')
```

```
C:\Users\zfishlab\AppData\Local\Continuum\anaconda3\envs\test\lib\site-packages\skimage\util\dtype.py:141: UserWarning: Possible precision loss when converting from float32 to uint8
  .format(dtypeobj_in, dtypeobj_out))
```

```
102
```

```
C:\Users\zfishlab\AppData\Local\Continuum\anaconda3\envs\test\lib\site-packages\skimage\util\dtype.py:141: UserWarning: Possible precision loss when converting from float32 to uint8
  .format(dtypeobj_in, dtypeobj_out))
```

```
103
```

```
C:\Users\zfishlab\AppData\Local\Continuum\anaconda3\envs\test\lib\site-packages\skimage\util\dtype.py:141: UserWarning: Possible precision loss when converting from float32 to uint8
  .format(dtypeobj_in, dtypeobj_out))
```

```
104
```

```
C:\Users\zfishlab\AppData\Local\Continuum\anaconda3\envs\test\lib\site-packages\skimage\util\dtype.py:141: UserWarning: Possible precision loss when converting from float32 to uint8
  .format(dtypeobj_in, dtypeobj_out))
```

```
105
```

```
C:\Users\zfishlab\AppData\Local\Continuum\anaconda3\envs\test\lib\site-packages\skimage\util\dtype.py:141: UserWarning: Possible precision loss when converting from float32 to uint8
  .format(dtypeobj_in, dtypeobj_out))
```

```
106
```

```
C:\Users\zfishlab\AppData\Local\Continuum\anaconda3\envs\test\lib\site-packages\skimage\util\dtype.py:141: UserWarning: Possible precision loss when converting from float32 to uint8
  .format(dtypeobj_in, dtypeobj_out))
```

```
107
```

```
C:\Users\zfishlab\AppData\Local\Continuum\anaconda3\envs\test\lib\site-packages\skimage\util\dtype.py:141: UserWarning: Possible precision loss when converting from float32 to uint8
  .format(dtypeobj_in, dtypeobj_out))
```

```
108
```

```
C:\Users\zfishlab\AppData\Local\Continuum\anaconda3\envs\test\lib\site-packages\skimage\util\dtype.py:141: UserWarning: Possible precision loss when converting from float32 to uint8
  .format(dtypeobj_in, dtypeobj_out))
```

```
110
```

```
C:\Users\zfishlab\AppData\Local\Continuum\anaconda3\envs\test\lib\site-packages\skimage\util\dtype.py:141: UserWarning: Possible precision loss when converting from float32 to uint8
  .format(dtypeobj_in, dtypeobj_out))
```

```
1
```

```
C:\Users\zfishlab\AppData\Local\Continuum\anaconda3\envs\test\lib\site-packages\skimage\util\dtype.py:141: UserWarning: Possible precision loss when converting from float32 to uint8
  .format(dtypeobj_in, dtypeobj_out))
```

```
2
```

```
C:\Users\zfishlab\AppData\Local\Continuum\anaconda3\envs\test\lib\site-packages\skimage\util\dtype.py:141: UserWarning: Possible precision loss when converting from float32 to uint8
  .format(dtypeobj_in, dtypeobj_out))
```

```
36
```

```
C:\Users\zfishlab\AppData\Local\Continuum\anaconda3\envs\test\lib\site-packages\skimage\util\dtype.py:141: UserWarning: Possible precision loss when converting from float32 to uint8
  .format(dtypeobj_in, dtypeobj_out))
```

```
3
```

```
C:\Users\zfishlab\AppData\Local\Continuum\anaconda3\envs\test\lib\site-packages\skimage\util\dtype.py:141: UserWarning: Possible precision loss when converting from float32 to uint8
  .format(dtypeobj_in, dtypeobj_out))
```

```
41
```

```
C:\Users\zfishlab\AppData\Local\Continuum\anaconda3\envs\test\lib\site-packages\skimage\util\dtype.py:141: UserWarning: Possible precision loss when converting from float32 to uint8
  .format(dtypeobj_in, dtypeobj_out))
```

```
44
```

```
C:\Users\zfishlab\AppData\Local\Continuum\anaconda3\envs\test\lib\site-packages\skimage\util\dtype.py:141: UserWarning: Possible precision loss when converting from float32 to uint8
  .format(dtypeobj_in, dtypeobj_out))
```

```
47
```

```
C:\Users\zfishlab\AppData\Local\Continuum\anaconda3\envs\test\lib\site-packages\skimage\util\dtype.py:141: UserWarning: Possible precision loss when converting from float32 to uint8
  .format(dtypeobj_in, dtypeobj_out))
```

```
4
```

```
C:\Users\zfishlab\AppData\Local\Continuum\anaconda3\envs\test\lib\site-packages\skimage\util\dtype.py:141: UserWarning: Possible precision loss when converting from float32 to uint8
  .format(dtypeobj_in, dtypeobj_out))
```

```
5
```

```
C:\Users\zfishlab\AppData\Local\Continuum\anaconda3\envs\test\lib\site-packages\skimage\util\dtype.py:141: UserWarning: Possible precision loss when converting from float32 to uint8
  .format(dtypeobj_in, dtypeobj_out))
```

```
6
```

```
C:\Users\zfishlab\AppData\Local\Continuum\anaconda3\envs\test\lib\site-packages\skimage\util\dtype.py:141: UserWarning: Possible precision loss when converting from float32 to uint8
  .format(dtypeobj_in, dtypeobj_out))
```

```
7
Wall time: 12min 29s
```

In [12]:

```
def start(k):
    return(ut.start(k,Dbat,[Dbzrf],im=True))
def save_both(k,dfa,dfb):
    ut.save_both(k,dfa,dfb,outdir,'30hpf')
```

In [13]:

```
model = pd.DataFrame({'a':[],'b':[],'c':[]})
def save_model(k,mm,model):
    row = pd.Series({'a':mm[0],'b':mm[1],'c':mm[2]},name=k)
    model = model.append(row)
    return(model)

def fit_model(axi,df,mm=None):
    if mm == None:
        mm = np.polyfit(df.x,df.z,2)
    p = np.poly1d(mm)
    xrange = np.arange(np.min(df.x),np.max(df.x))
    axi.plot(xrange,p(xrange),c='m')
    return(mm)

def pick_pts(x1,z1,vx,vz,x2,z2):
    pts = pd.DataFrame({'x':[x1,vx,x2],'z':[z1,vz,z2]})
    return(pts)
```

# 47¶

Discard because evidence of tear in zrf

In [14]:

```
k,df,Ldf,ax = start('47')
```

# 107¶

In [15]:

```
k,df,Ldf,ax = start('107')
```

In [16]:

```
df1,Ldf1,pts,ax = ut.check_pts(df,Ldf,'z')
```

In [17]:

```
pts.iloc[1].x = 35
pts.iloc[1].z = 12
ax[0,1].scatter(pts.x,pts.z,c='y')
pts
```

Out[17]:

|  | x | z |
| --- | --- | --- |
| 0 | -34.870677 | 17.376143 |
| 1 | 35.000000 | 12.000000 |

In [18]:

```
df2,Ldf2,ax = ut.revise_pts(df,Ldf,'z',pts=pts)
```

In [19]:

```
df3,Ldf3,ax,p = ut.check_yz(df2,Ldf2)
```

In [20]:

```
p = np.poly1d([-3/4,0])
xrange = np.arange(-10,20)
ax[0,2].plot(xrange,p(xrange),c='c')
```

Out[20]:

```
[<matplotlib.lines.Line2D at 0x20ae322a4a8>]
```

In [21]:

```
df4,Ldf4,ax,p = ut.check_yz(df2,Ldf2,mm=[-3/4,0])
```

In [22]:

```
df5,Ldf5,mm,ax = ut.ch_vertex(df4,Ldf4)
```

In [23]:

```
save_both(k,df5,Ldf5[0])
```

```
Write to .\Data\30hpf\Output-02-14-2019\AT_107_30hpf.psi complete
Write to .\Data\30hpf\Output-02-14-2019\ZRF_107_30hpf.psi complete
```

In [24]:

```
model = save_model(k,mm,model)
```

# 102¶

In [25]:

```
k,df,Ldf,ax = start('102')
```

In [26]:

```
mm = fit_model(ax[0,1],df)
```

In [27]:

```
model = save_model(k,mm,model)
```

In [28]:

```
save_both(k,df,Ldf[0])
```

```
Write to .\Data\30hpf\Output-02-14-2019\AT_102_30hpf.psi complete
Write to .\Data\30hpf\Output-02-14-2019\ZRF_102_30hpf.psi complete
```

# 104¶

In [29]:

```
k,df,Ldf,ax = start('104')
```

In [30]:

```
mm = fit_model(ax[0,1],df)
```

In [31]:

```
model = save_model(k,mm,model)
```

In [32]:

```
save_both(k,df,Ldf[0])
```

```
Write to .\Data\30hpf\Output-02-14-2019\AT_104_30hpf.psi complete
Write to .\Data\30hpf\Output-02-14-2019\ZRF_104_30hpf.psi complete
```

# 5¶

Discard because abnormally low amounts of signal

In [33]:

```
k,df,Ldf,ax = start('5')
```

# 105¶

In [34]:

```
k,df,Ldf,ax = start('105')
```

In [35]:

```
df1,Ldf1,pts,ax = ut.check_pts(df,Ldf,'z')
```

In [36]:

```
pts.iloc[1].x = 28
pts.iloc[1].z = 21
pts.iloc[0].z = 24
ax[0,1].scatter(pts.x,pts.z,c='y')
pts
```

Out[36]:

|  | x | z |
| --- | --- | --- |
| 0 | -28.50495 | 24.0 |
| 1 | 28.00000 | 21.0 |

In [37]:

```
df2,Ldf2,ax = ut.revise_pts(df,Ldf,'z',pts=pts)
```

In [38]:

```
df3,Ldf3,mm,ax = ut.ch_vertex(df2,Ldf2)
```

In [39]:

```
save_both(k,df3,Ldf3[0])
```

```
Write to .\Data\30hpf\Output-02-14-2019\AT_105_30hpf.psi complete
Write to .\Data\30hpf\Output-02-14-2019\ZRF_105_30hpf.psi complete
```

In [40]:

```
model = save_model(k,mm,model)
```

# 44¶

In [41]:

```
k,df,Ldf,ax = start('44')
```

In [42]:

```
mm = fit_model(ax[0,1],df)
```

In [43]:

```
pts = pick_pts(-24,13,0,-3,26,14)
ax[0,1].scatter(pts.x,pts.z,c='m',s=50)
```

Out[43]:

```
<matplotlib.collections.PathCollection at 0x20845fc85c0>
```

In [44]:

```
df1,Ldf1,mm,ax = ut.ch_vertex(df,Ldf,pts=pts)
```

In [45]:

```
save_both(k,df1,Ldf1[0])
```

```
Write to .\Data\30hpf\Output-02-14-2019\AT_44_30hpf.psi complete
Write to .\Data\30hpf\Output-02-14-2019\ZRF_44_30hpf.psi complete
```

In [46]:

```
model = save_model(k,mm,model)
```

# 110¶

In [47]:

```
k,df,Ldf,ax = start('110')
```

In [48]:

```
mm = fit_model(ax[0,1],df)
```

In [49]:

```
save_both(k,df,Ldf[0])
```

```
Write to .\Data\30hpf\Output-02-14-2019\AT_110_30hpf.psi complete
Write to .\Data\30hpf\Output-02-14-2019\ZRF_110_30hpf.psi complete
```

In [50]:

```
model = save_model(k,mm,model)
```

# 36¶

In [51]:

```
k,df,Ldf,ax = start('36')
```

In [52]:

```
mm = fit_model(ax[0,1],df)
```

In [53]:

```
save_both(k,df,Ldf[0])
```

```
Write to .\Data\30hpf\Output-02-14-2019\AT_36_30hpf.psi complete
Write to .\Data\30hpf\Output-02-14-2019\ZRF_36_30hpf.psi complete
```

In [54]:

```
model = save_model(k,mm,model)
```

# 103¶

In [55]:

```
k,df,Ldf,ax = start('103')
```

In [56]:

```
df1,Ldf1,pts,ax = ut.check_pts(df,Ldf,'z')
```

In [57]:

```
pts.iloc[1].x = -20
pts.iloc[1].z = 8
ax[0,1].scatter(pts.x,pts.z,c='y')
pts
```

Out[57]:

|  | x | z |
| --- | --- | --- |
| 0 | 27.589957 | 12.466037 |
| 1 | -20.000000 | 8.000000 |

In [58]:

```
df2,Ldf2,ax = ut.revise_pts(df,Ldf,'z',pts=pts)
```

In [59]:

```
df3,Ldf3,mm,ax = ut.ch_vertex(df2,Ldf2)
```

In [60]:

```
pts = pick_pts(-22,10,0,-2,23,10)
ax[0,1].scatter(pts.x,pts.z,c='m',s=50)
```

Out[60]:

```
<matplotlib.collections.PathCollection at 0x20b0c4b42b0>
```

In [61]:

```
df4,Ldf4,mm,ax = ut.ch_vertex(df3,Ldf3,pts=pts)
```

In [62]:

```
save_both(k,df4,Ldf4[0])
```

```
Write to .\Data\30hpf\Output-02-14-2019\AT_103_30hpf.psi complete
Write to .\Data\30hpf\Output-02-14-2019\ZRF_103_30hpf.psi complete
```

In [63]:

```
model = save_model(k,mm,model)
```

# 2¶

In [64]:

```
k,df,Ldf,ax = start('2')
```

In [65]:

```
df1,Ldf1,pts,ax = ut.check_pts(df,Ldf,'z')
```

In [66]:

```
pts.iloc[1].x = -26
pts.iloc[1].z = 10
ax[0,1].scatter(pts.x,pts.z,c='y')
pts
```

Out[66]:

|  | x | z |
| --- | --- | --- |
| 0 | 27.984379 | 14.801871 |
| 1 | -26.000000 | 10.000000 |

In [67]:

```
df2,Ldf2,ax = ut.revise_pts(df,Ldf,'z',pts=pts)
```

In [68]:

```
df2,Ldf2,mm,ax = ut.ch_vertex(df2,Ldf2)
```

In [69]:

```
save_both(k,df2,Ldf2[0])
```

```
Write to .\Data\30hpf\Output-02-14-2019\AT_2_30hpf.psi complete
Write to .\Data\30hpf\Output-02-14-2019\ZRF_2_30hpf.psi complete
```

In [70]:

```
model = save_model(k,mm,model)
```

# 106¶

In [71]:

```
k,df,Ldf,ax = start('106')
```

In [72]:

```
ax = ut.make_graph([Dbat[k].df_thresh,Dbzrf[k].df_thresh])
```

In [73]:

```
df = Dbat[k].df_thresh
Ldf = [Dbzrf[k].df_thresh]
```

In [74]:

```
df1,Ldf1,ax,p = ut.check_yz(Ldf[0],[df])
```

In [75]:

```
df2,Ldf2 = ut.flip(Ldf1[0],[df1])
ax = ut.make_graph([df2]+Ldf2)
```

In [76]:

```
df3,Ldf3,mm,ax = ut.ch_vertex(df2,Ldf2)
```

In [77]:

```
df4,Ldf4,pts,ax = ut.check_pts(df3,Ldf3,'z')
```

In [78]:

```
pts.iloc[1].x = 55
pts.iloc[1].z = 15
ax[0,1].scatter(pts.x,pts.z,c='y')
pts
```

Out[78]:

|  | x | z |
| --- | --- | --- |
| 0 | -2.972424 | 17.200127 |
| 1 | 55.000000 | 15.000000 |

In [79]:

```
df5,Ldf5,ax = ut.revise_pts(df3,Ldf3,'z',pts=pts)
```

In [80]:

```
pts = pick_pts(-2,17,25,0,55,15)
ax[1,1].scatter(pts.x,pts.z,c='m',s=50)
```

Out[80]:

```
<matplotlib.collections.PathCollection at 0x20ae3312ba8>
```

In [81]:

```
df6,Ldf6,mm,ax = ut.ch_vertex(df5,Ldf5,pts=pts)
```

In [82]:

```
save_both(k,df6,Ldf6[0])
```

```
Write to .\Data\30hpf\Output-02-14-2019\AT_106_30hpf.psi complete
Write to .\Data\30hpf\Output-02-14-2019\ZRF_106_30hpf.psi complete
```

In [83]:

```
model = save_model(k,mm,model)
```

# 41¶

In [84]:

```
k,df,Ldf,ax = start('41')
```

In [85]:

```
mm = fit_model(ax[0,1],df)
```

In [86]:

```
save_both(k,df,Ldf[0])
```

```
Write to .\Data\30hpf\Output-02-14-2019\AT_41_30hpf.psi complete
Write to .\Data\30hpf\Output-02-14-2019\ZRF_41_30hpf.psi complete
```

In [87]:

```
model = save_model(k,mm,model)
```

# 1¶

In [88]:

```
k,df,Ldf,ax = start('1')
```

In [89]:

```
mm = fit_model(ax[0,1],df)
```

In [90]:

```
save_both(k,df,Ldf[0])
```

```
Write to .\Data\30hpf\Output-02-14-2019\AT_1_30hpf.psi complete
Write to .\Data\30hpf\Output-02-14-2019\ZRF_1_30hpf.psi complete
```

In [91]:

```
model = save_model(k,mm,model)
```

# 6¶

In [92]:

```
k,df,Ldf,ax = start('6')
```

In [93]:

```
mm = fit_model(ax[0,1],df)
```

In [94]:

```
save_both(k,df,Ldf[0])
```

```
Write to .\Data\30hpf\Output-02-14-2019\AT_6_30hpf.psi complete
Write to .\Data\30hpf\Output-02-14-2019\ZRF_6_30hpf.psi complete
```

In [95]:

```
model = save_model(k,mm,model)
```

# 7¶

In [96]:

```
k,df,Ldf,ax = start('7')
```

In [97]:

```
df1,Ldf1,ax,p = ut.check_yz(df,Ldf)
```

In [98]:

```
df2,Ldf2,mm,ax = ut.ch_vertex(df1,Ldf1)
```

In [99]:

```
pts = pick_pts(-35,14,0,-3,30,12)
ax[0,1].scatter(pts.x,pts.z,c='m',s=50)
```

Out[99]:

```
<matplotlib.collections.PathCollection at 0x20848531ac8>
```

In [100]:

```
df3,Ldf3,mm,ax = ut.ch_vertex(df2,Ldf2,pts=pts)
```

In [101]:

```
save_both(k,df3,Ldf3[0])
```

```
Write to .\Data\30hpf\Output-02-14-2019\AT_7_30hpf.psi complete
Write to .\Data\30hpf\Output-02-14-2019\ZRF_7_30hpf.psi complete
```

In [102]:

```
model = save_model(k,mm,model)
```

# 4¶

In [103]:

```
k,df,Ldf,ax = start('4')
```

In [104]:

```
mm = fit_model(ax[0,1],df)
```

In [105]:

```
save_both(k,df,Ldf[0])
```

```
Write to .\Data\30hpf\Output-02-14-2019\AT_4_30hpf.psi complete
Write to .\Data\30hpf\Output-02-14-2019\ZRF_4_30hpf.psi complete
```

In [106]:

```
model = save_model(k,mm,model)
```

# 108¶

In [107]:

```
k,df,Ldf,ax = start('108')
```

In [108]:

```
df1,Ldf1,pts,ax = ut.check_pts(df,Ldf,'z')
```

In [109]:

```
df2,Ldf2,mm,ax = ut.ch_vertex(df1,Ldf1)
```

In [110]:

```
save_both(k,df2,Ldf2[0])
```

```
Write to .\Data\30hpf\Output-02-14-2019\AT_108_30hpf.psi complete
Write to .\Data\30hpf\Output-02-14-2019\ZRF_108_30hpf.psi complete
```

In [111]:

```
model = save_model(k,mm,model)
```

# 3¶

In [112]:

```
k,df,Ldf,ax = start('3')
```

In [113]:

```
mm = fit_model(ax[0,1],df)
```

In [114]:

```
save_both(k,df,Ldf[0])
```

```
Write to .\Data\30hpf\Output-02-14-2019\AT_3_30hpf.psi complete
Write to .\Data\30hpf\Output-02-14-2019\ZRF_3_30hpf.psi complete
```

In [115]:

```
model = save_model(k,mm,model)
```

### Model¶

In [116]:

```
model.to_csv(os.path.join(outdir,'model.csv'))
```

In [117]:

```
outdir
```

Out[117]:

```
'.\\Data\\30hpf\\Output-02-14-2019'
```

In [118]:

```
model
```

Out[118]:

|  | a | b | c |
| --- | --- | --- | --- |
| 107 | 0.012729 | 1.249706e-16 | -5.578322e-16 |
| 102 | 0.029971 | 2.459886e-16 | 9.524782e-16 |
| 104 | 0.010800 | -5.060717e-17 | 6.924850e-15 |
| 105 | 0.024225 | 6.748456e-17 | -1.605983e-16 |
| 44 | 0.026410 | -9.240142e-18 | -5.125331e-15 |
| 110 | 0.028474 | -2.152950e-16 | -1.120272e-14 |
| 36 | 0.011241 | -1.668182e-16 | 3.357557e-15 |
| 103 | 0.023715 | -5.407316e-17 | -3.076747e-15 |
| 2 | 0.015399 | 4.589006e-17 | 6.541389e-16 |
| 106 | 0.019818 | 4.528415e-17 | -1.949156e-14 |
| 41 | 0.013714 | -1.932264e-17 | 9.528271e-15 |
| 1 | 0.013464 | -1.155351e-17 | -5.103873e-16 |
| 6 | 0.015777 | 8.905672e-17 | -1.046242e-14 |
| 7 | 0.015165 | 6.837824e-17 | -4.953653e-15 |
| 4 | 0.011876 | -2.356489e-16 | -4.532531e-15 |
| 108 | 0.021055 | -1.026399e-16 | 2.558750e-15 |
| 3 | 0.010345 | 2.025744e-17 | -9.583682e-16 |
