## Supplementary material for "ΔSCOPE: A new method to quantify 3D biological structures and identify differences in zebrafish forebrain development": An archived copy of the DeltaSCOPE code repository.: alignment-30hpf.ipynb

```
# %% [code]
import numpy as np
import pandas as pd
import matplotlib.pyplot as plt

import os
import re
from imp import reload
import h5py

# %% [code]
import deltascope as cranium
import deltascope.alignment as ut

# %% [code]
at = ".\\Data\\30hpf\\AT\\Prob"
gfap = ".\\Data\\30hpf\\Gfap\\Prob"

# %% [code]
outdir = ".\\Data\\30hpf\\Output-02-14-2019"

# %% [code]
os.mkdir(outdir)

# %% [code]
Dat = {}
for f in os.listdir(at):
    if 'h5' in f:
        num  = re.findall(r'\d+',f.split('.')[0])[-1]
        Dat[num] = os.path.join(at,f)

# %% [code]
Dzrf = {}
for f in os.listdir(gfap):
    if 'h5' in f:
        num  = re.findall(r'\d+',f.split('.')[0])[-1]
        Dzrf[num] = os.path.join(gfap,f)

# %% [code]
Dbat = {}
Dbzrf = {}

# %% [markdown]
# Data Preprocessing

# %% [code]
klist = Dat.keys()

# %% [code]
param = {
    'gthresh':0.5,
    'scale':[1,1,1],
    'microns':[0.16,0.16,0.21],
    'mthresh':0.5,
    'radius':10,
    'comp_order':[0,2,1],
    'fit_dim':['x','z'],
    'deg':2
}

# %% [code]
%%time
for k in klist:
    if k not in list(Dbat.keys()):
        Dbat[k] = ut.preprocess(Dat[k],param)
        Dbzrf[k] = ut.preprocess(Dzrf[k],param,pca=Dbat[k].pcamed,mm=Dbat[k].mm,vertex=Dbat[k].vertex)
        print(k)
    else:
        print(k,'already processed')

# %% [code]
def start(k):
    return(ut.start(k,Dbat,[Dbzrf],im=True))
def save_both(k,dfa,dfb):
    ut.save_both(k,dfa,dfb,outdir,'30hpf')

# %% [code]
model = pd.DataFrame({'a':[],'b':[],'c':[]})
def save_model(k,mm,model):
    row = pd.Series({'a':mm[0],'b':mm[1],'c':mm[2]},name=k)
    model = model.append(row)
    return(model)

def fit_model(axi,df,mm=None):
    if mm == None:
        mm = np.polyfit(df.x,df.z,2)
    p = np.poly1d(mm)
    xrange = np.arange(np.min(df.x),np.max(df.x))
    axi.plot(xrange,p(xrange),c='m')
    return(mm)

def pick_pts(x1,z1,vx,vz,x2,z2):
    pts = pd.DataFrame({'x':[x1,vx,x2],'z':[z1,vz,z2]})
    return(pts)

# %% [markdown]
# 47

# %% [markdown]
Discard because evidence of tear in zrf

# %% [code]
k,df,Ldf,ax = start('47')

# %% [markdown]
# 107

# %% [code]
k,df,Ldf,ax = start('107')

# %% [code]
df1,Ldf1,pts,ax = ut.check_pts(df,Ldf,'z')

# %% [code]
pts.iloc[1].x = 35
pts.iloc[1].z = 12
ax[0,1].scatter(pts.x,pts.z,c='y')
pts

# %% [code]
df2,Ldf2,ax = ut.revise_pts(df,Ldf,'z',pts=pts)

# %% [code]
df3,Ldf3,ax,p = ut.check_yz(df2,Ldf2)

# %% [code]
p = np.poly1d([-3/4,0])
xrange = np.arange(-10,20)
ax[0,2].plot(xrange,p(xrange),c='c')

# %% [code]
df4,Ldf4,ax,p = ut.check_yz(df2,Ldf2,mm=[-3/4,0])

# %% [code]
df5,Ldf5,mm,ax = ut.ch_vertex(df4,Ldf4)

# %% [code]
save_both(k,df5,Ldf5[0])

# %% [code]
model = save_model(k,mm,model)

# %% [markdown]
# 102

# %% [code]
k,df,Ldf,ax = start('102')

# %% [code]
mm = fit_model(ax[0,1],df)

# %% [code]
model = save_model(k,mm,model)

# %% [code]
save_both(k,df,Ldf[0])

# %% [markdown]
# 104

# %% [code]
k,df,Ldf,ax = start('104')

# %% [code]
mm = fit_model(ax[0,1],df)

# %% [code]
model = save_model(k,mm,model)

# %% [code]
save_both(k,df,Ldf[0])

# %% [markdown]
# 5

# %% [markdown]
Discard because abnormally low amounts of signal

# %% [code]
k,df,Ldf,ax = start('5')

# %% [markdown]
# 105

# %% [code]
k,df,Ldf,ax = start('105')

# %% [code]
df1,Ldf1,pts,ax = ut.check_pts(df,Ldf,'z')

# %% [code]
pts.iloc[1].x = 28
pts.iloc[1].z = 21
pts.iloc[0].z = 24
ax[0,1].scatter(pts.x,pts.z,c='y')
pts

# %% [code]
df2,Ldf2,ax = ut.revise_pts(df,Ldf,'z',pts=pts)

# %% [code]
df3,Ldf3,mm,ax = ut.ch_vertex(df2,Ldf2)

# %% [code]
save_both(k,df3,Ldf3[0])

# %% [code]
model = save_model(k,mm,model)

# %% [markdown]
# 44

# %% [code]
k,df,Ldf,ax = start('44')

# %% [code]
mm = fit_model(ax[0,1],df)

# %% [code]
pts = pick_pts(-24,13,0,-3,26,14)
ax[0,1].scatter(pts.x,pts.z,c='m',s=50)

# %% [code]
df1,Ldf1,mm,ax = ut.ch_vertex(df,Ldf,pts=pts)

# %% [code]
save_both(k,df1,Ldf1[0])

# %% [code]
model = save_model(k,mm,model)

# %% [markdown]
# 110

# %% [code]
k,df,Ldf,ax = start('110')

# %% [code]
mm = fit_model(ax[0,1],df)

# %% [code]
save_both(k,df,Ldf[0])

# %% [code]
model = save_model(k,mm,model)

# %% [markdown]
# 36

# %% [code]
k,df,Ldf,ax = start('36')

# %% [code]
mm = fit_model(ax[0,1],df)

# %% [code]
save_both(k,df,Ldf[0])

# %% [code]
model = save_model(k,mm,model)

# %% [markdown]
# 103

# %% [code]
k,df,Ldf,ax = start('103')

# %% [code]
df1,Ldf1,pts,ax = ut.check_pts(df,Ldf,'z')

# %% [code]
pts.iloc[1].x = -20
pts.iloc[1].z = 8
ax[0,1].scatter(pts.x,pts.z,c='y')
pts

# %% [code]
df2,Ldf2,ax = ut.revise_pts(df,Ldf,'z',pts=pts)

# %% [code]
df3,Ldf3,mm,ax = ut.ch_vertex(df2,Ldf2)

# %% [code]
pts = pick_pts(-22,10,0,-2,23,10)
ax[0,1].scatter(pts.x,pts.z,c='m',s=50)

# %% [code]
df4,Ldf4,mm,ax = ut.ch_vertex(df3,Ldf3,pts=pts)

# %% [code]
save_both(k,df4,Ldf4[0])

# %% [code]
model = save_model(k,mm,model)

# %% [markdown]
# 2

# %% [code]
k,df,Ldf,ax = start('2')

# %% [code]
df1,Ldf1,pts,ax = ut.check_pts(df,Ldf,'z')

# %% [code]
pts.iloc[1].x = -26
pts.iloc[1].z = 10
ax[0,1].scatter(pts.x,pts.z,c='y')
pts

# %% [code]
df2,Ldf2,ax = ut.revise_pts(df,Ldf,'z',pts=pts)

# %% [code]
df2,Ldf2,mm,ax = ut.ch_vertex(df2,Ldf2)

# %% [code]
save_both(k,df2,Ldf2[0])

# %% [code]
model = save_model(k,mm,model)

# %% [markdown]
# 106

# %% [code]
k,df,Ldf,ax = start('106')

# %% [code]
ax = ut.make_graph([Dbat[k].df_thresh,Dbzrf[k].df_thresh])

# %% [code]
df = Dbat[k].df_thresh
Ldf = [Dbzrf[k].df_thresh]

# %% [code]
df1,Ldf1,ax,p = ut.check_yz(Ldf[0],[df])

# %% [code]
df2,Ldf2 = ut.flip(Ldf1[0],[df1])
ax = ut.make_graph([df2]+Ldf2)

# %% [code]
df3,Ldf3,mm,ax = ut.ch_vertex(df2,Ldf2)

# %% [code]
df4,Ldf4,pts,ax = ut.check_pts(df3,Ldf3,'z')

# %% [code]
pts.iloc[1].x = 55
pts.iloc[1].z = 15
ax[0,1].scatter(pts.x,pts.z,c='y')
pts

# %% [code]
df5,Ldf5,ax = ut.revise_pts(df3,Ldf3,'z',pts=pts)

# %% [code]
pts = pick_pts(-2,17,25,0,55,15)
ax[1,1].scatter(pts.x,pts.z,c='m',s=50)

# %% [code]
df6,Ldf6,mm,ax = ut.ch_vertex(df5,Ldf5,pts=pts)

# %% [code]
save_both(k,df6,Ldf6[0])

# %% [code]
model = save_model(k,mm,model)

# %% [markdown]
# 41

# %% [code]
k,df,Ldf,ax = start('41')

# %% [code]
mm = fit_model(ax[0,1],df)

# %% [code]
save_both(k,df,Ldf[0])

# %% [code]
model = save_model(k,mm,model)

# %% [markdown]
# 1

# %% [code]
k,df,Ldf,ax = start('1')

# %% [code]
mm = fit_model(ax[0,1],df)

# %% [code]
save_both(k,df,Ldf[0])

# %% [code]
model = save_model(k,mm,model)

# %% [markdown]
# 6

# %% [code]
k,df,Ldf,ax = start('6')

# %% [code]
mm = fit_model(ax[0,1],df)

# %% [code]
save_both(k,df,Ldf[0])

# %% [code]
model = save_model(k,mm,model)

# %% [markdown]
# 7

# %% [code]
k,df,Ldf,ax = start('7')

# %% [code]
df1,Ldf1,ax,p = ut.check_yz(df,Ldf)

# %% [code]
df2,Ldf2,mm,ax = ut.ch_vertex(df1,Ldf1)

# %% [code]
pts = pick_pts(-35,14,0,-3,30,12)
ax[0,1].scatter(pts.x,pts.z,c='m',s=50)

# %% [code]
df3,Ldf3,mm,ax = ut.ch_vertex(df2,Ldf2,pts=pts)

# %% [code]
save_both(k,df3,Ldf3[0])

# %% [code]
model = save_model(k,mm,model)

# %% [markdown]
# 4

# %% [code]
k,df,Ldf,ax = start('4')

# %% [code]
mm = fit_model(ax[0,1],df)

# %% [code]
save_both(k,df,Ldf[0])

# %% [code]
model = save_model(k,mm,model)

# %% [markdown]
# 108

# %% [code]
k,df,Ldf,ax = start('108')

# %% [code]
df1,Ldf1,pts,ax = ut.check_pts(df,Ldf,'z')

# %% [code]
df2,Ldf2,mm,ax = ut.ch_vertex(df1,Ldf1)

# %% [code]
save_both(k,df2,Ldf2[0])

# %% [code]
model = save_model(k,mm,model)

# %% [markdown]
# 3

# %% [code]
k,df,Ldf,ax = start('3')

# %% [code]
mm = fit_model(ax[0,1],df)

# %% [code]
save_both(k,df,Ldf[0])

# %% [code]
model = save_model(k,mm,model)

# %% [markdown]
# Model

# %% [code]
model.to_csv(os.path.join(outdir,'model.csv'))

# %% [code]
outdir

# %% [code]
model
```
