## Supplementary material for "ΔSCOPE: A new method to quantify 3D biological structures and identify differences in zebrafish forebrain development": An archived copy of the DeltaSCOPE code repository.: transform_after_align.py

```
import deltascope as ds
import time
import glob
import re
import traceback
import os
import multiprocessing as mp
import pandas as pd
from functools import partial

outdirs = ['.\\Data\\22hpf\\Output-02-14-2019',
            '.\\Data\\24hpf\\Output-02-15-2019',
            '.\\Data\\26hpf\\Output-02-14-2019',
            '.\\Data\\28hpf\\Output-02-14-2019-yot-ilastik',
            '.\\Data\\30hpf\\Output-02-14-2019']
deg = 2
fit_dim = ['x','z']

def transform_file(f,model=None):
    
    print(f,'starting')
    tic = time.time()
    
    s = ds.brain()
    df = ds.read_psi(f)
    
    # Transform coordinates if cylindrical column not present
    if 'ac' not in df.columns:
        # Add psi data to brain object
        s.df_align = df[['x','y','z']]
        #snum = re.findall(r'\d+',f.split('.')[0])[0]
        snum = f.split('_')[1]
        num = int(snum)
        
        # Extract math model and add to brain object
        try:
            mm_val = model.loc[num].values
            s.mm = ds.math_model(mm_val)
        except:
            s.mm = s.fit_model(s.df_align,deg,fit_dim)
        
        # Try to transform and report errors
        try:
            s.transform_coordinates()
        except:
            print(f,'failed on transform coordinates',time.time()-tic)
            traceback.print_exc()
        
        # Try to save data and report errors
        try:
            ds.write_data(f,s.df_align)
        except:
            print(f,'failed on write data',time.time()-tic)
            traceback.print_exc()
        
        print(f,'complete',time.time()-tic)
    
    else:
        # Report if file already appears transformed
        print(f,'already transformed')
    
if __name__=='__main__':
    
    for outdir in outdirs:
        
        # Select only psi files from directory
        files = glob.glob(os.path.join(outdir,'*.psi'))
        
        # Extract model from saved csv file
        model = pd.read_csv(os.path.join(outdir,'model.csv'),
                           index_col='Unnamed: 0')
        
        # Create function with a set of inputs
        transform = partial(transform_file,model=model)
        
        # Loop over files and multiprocess in sets of 5
        n = len(files)
        for i in range(0,n,5):
            if i+5>n:
                L = files[i:n]
            else:
                L = files[i:i+5]
                
            pool = mp.Pool()
            pool.map(transform,L)
            pool.close()
            pool.join()
```
