## Supplementary material for "ΔSCOPE: A new method to quantify 3D biological structures and identify differences in zebrafish forebrain development": An archived copy of the DeltaSCOPE code repository.: hss1a_alignment.html

2018-08-13\_hss1a\_alignment-Rerun


In [2]:

```
import numpy as np
import pandas as pd
import matplotlib.pyplot as plt

import os
import re
from imp import reload
import h5py
```

In [3]:

```
import deltascope as cranium
import deltascope.alignment as ut
```

In [4]:

```
at = ".\data\hss1a\AT\Prob"
gfap = ".\data\hss1a\GFAP\Prob"
root = ".\data\hss1a"
```

In [5]:

```
outdir = os.path.join(root,'Output-02-15-2019')
os.mkdir(outdir)
```

In [6]:

```
Dat = {}
for f in os.listdir(at):
    if 'h5' in f:
        num  = re.findall(r'\d+',f.split('.')[0])[-1]
        Dat[num] = os.path.join(at,f)
```

In [7]:

```
Dzrf = {}
for f in os.listdir(gfap):
    if 'h5' in f:
        num  = re.findall(r'\d+',f.split('.')[0])[-1]
        Dzrf[num] = os.path.join(gfap,f)
```

In [8]:

```
Dbat = {}
Dbzrf = {}
```

### Data Processing¶

In [9]:

```
klist = Dat.keys()
```

In [10]:

```
param = {
    'gthresh':0.5,
    'scale':[1,1,1],
    'microns':[0.16,0.16,0.21],
    'mthresh':0.5,
    'radius':10,
    'comp_order':[0,2,1],
    'fit_dim':['x','z'],
    'deg':2
}
```

In [11]:

```
%%time
for k in klist:
    if k not in list(Dbat.keys()):
        Dbat[k] = ut.preprocess(Dat[k],param)
        Dbzrf[k] = ut.preprocess(Dzrf[k],param,pca=Dbat[k].pcamed,mm=Dbat[k].mm,vertex=Dbat[k].vertex)
        #Dbcaax[k] = ut.preprocess(Dcaax[k],param,pca=Dbat[k].pcamed,mm=Dbat[k].mm,vertex=Dbat[k].vertex)
        print(k)
    else:
        print(k,'already processed')
```

```
6
Wall time: 7min 39s
```

### Functions¶

In [12]:

```
def start(k):
    return(ut.start(k,Dbat,[Dbzrf],im=True))
def save_both(k,dfa,dfb):
    ut.save_both(k,dfa,dfb,outdir,'hss1a')
```

In [13]:

```
model = pd.DataFrame({'a':[],'b':[],'c':[]})
def save_model(k,mm,model):
    row = pd.Series({'a':mm[0],'b':mm[1],'c':mm[2]},name=k)
    model = model.append(row)
    return(model)
```

In [14]:

```
def fit_model(axi,df,mm=None):
    if mm == None:
        mm = np.polyfit(df.x,df.z,2)
    p = np.poly1d(mm)
    xrange = np.arange(np.min(df.x),np.max(df.x))
    axi.plot(xrange,p(xrange),c='m')
    return(mm)
```

In [15]:

```
def pick_pts(x1,z1,vx,vz,x2,z2):
    pts = pd.DataFrame({'x':[x1,vx,x2],'z':[z1,vz,z2]})
    return(pts)
```

# 06¶

In [16]:

```
k,df,Ldf,ax = start('06')
```

In [17]:

```
k = '206'
```

In [18]:

```
mm = fit_model(ax[0,1],df)
```

In [19]:

```
model = save_model(k,mm,model)
save_both(k,df,Ldf[0])
```

```
Write to .\data\hss1a\Output-02-15-2019\AT_206_hss1a.psi complete
Write to .\data\hss1a\Output-02-15-2019\ZRF_206_hss1a.psi complete
```

# 20¶

In [20]:

```
k,df,Ldf,ax = start('20')
```

In [21]:

```
df1,Ldf1 = ut.zyswitch(df,Ldf)
ax = ut.make_graph([df1]+Ldf1)
```

In [22]:

```
df2,Ldf2,mm,ax = ut.ch_vertex(df1,Ldf1)
```

In [23]:

```
pts = pick_pts(-36,3,0,-5,42,4)
ax[0,1].scatter(pts.x,pts.z,c='m',s=50)
```

Out[23]:

```
<matplotlib.collections.PathCollection at 0x2e303243e10>
```

In [24]:

```
df3,Ldf3,mm,ax = ut.ch_vertex(df2,Ldf2,pts=pts)
```

In [25]:

```
model = save_model(k,mm,model)
save_both(k,df3,Ldf3[0])
```

```
Write to .\data\hss1a\Output-02-15-2019\AT_20_hss1a.psi complete
Write to .\data\hss1a\Output-02-15-2019\ZRF_20_hss1a.psi complete
```

# 17¶

In [26]:

```
k,df,Ldf,ax = start('17')
```

In [27]:

```
df1,Ldf1,pts,ax = ut.check_pts(df,Ldf,'z')
```

In [28]:

```
pts.iloc[0].x = 35
pts.iloc[0].z = 10
ax[0,1].scatter(pts.x,pts.z,c='y')
pts
```

Out[28]:

|  | x | z |
| --- | --- | --- |
| 0 | 35.000000 | 10.000000 |
| 1 | -38.831946 | 15.116733 |

In [29]:

```
df2,Ldf2,ax = ut.revise_pts(df,Ldf,'z',pts=pts)
```

In [30]:

```
df3,Ldf3,mm,ax = ut.ch_vertex(df2,Ldf2)
```

In [31]:

```
pts = pick_pts(-36,13,0,-4,38,13)
ax[0,1].scatter(pts.x,pts.z,c='m',s=50)
```

Out[31]:

```
<matplotlib.collections.PathCollection at 0x2e301639748>
```

In [32]:

```
df4,Ldf4,mm,ax = ut.ch_vertex(df3,Ldf3,pts=pts)
```

In [33]:

```
model = save_model(k,mm,model)
save_both(k,df4,Ldf4[0])
```

```
Write to .\data\hss1a\Output-02-15-2019\AT_17_hss1a.psi complete
Write to .\data\hss1a\Output-02-15-2019\ZRF_17_hss1a.psi complete
```

# 13¶

In [34]:

```
k,df,Ldf,ax = start('13')
```

In [35]:

```
df1,Ldf1 = ut.zyswitch(df,Ldf)
ax = ut.make_graph([df1]+Ldf1)
```

In [36]:

```
df2,Ldf2,mm,ax = ut.ch_vertex(df1,Ldf1)
```

In [37]:

```
pts = pick_pts(-55,5,0,-3,55,5)
ax[0,1].scatter(pts.x,pts.z,c='m',s=50)
```

Out[37]:

```
<matplotlib.collections.PathCollection at 0x2e3015e3c88>
```

In [38]:

```
df3,Ldf3,mm,ax = ut.ch_vertex(df2,Ldf2,pts=pts)
```

In [39]:

```
model = save_model(k,mm,model)
save_both(k,df3,Ldf3[0])
```

```
Write to .\data\hss1a\Output-02-15-2019\AT_13_hss1a.psi complete
Write to .\data\hss1a\Output-02-15-2019\ZRF_13_hss1a.psi complete
```

In [40]:

```
klist
```

Out[40]:

```
dict_keys(['02', '04', '06', '10', '11', '13', '16', '17', '18', '20', '6'])
```

# 6¶

In [41]:

```
k,df,Ldf,ax = start('6')
```

In [42]:

```
df1,Ldf1 = ut.zyswitch(df,Ldf)
ax = ut.make_graph([df1]+Ldf1)
```

In [43]:

```
df2,Ldf2,mm,ax = ut.ch_vertex(df1,Ldf1)
```

In [44]:

```
model = save_model(k,mm,model)
save_both(k,df2,Ldf2[0])
```

```
Write to .\data\hss1a\Output-02-15-2019\AT_6_hss1a.psi complete
Write to .\data\hss1a\Output-02-15-2019\ZRF_6_hss1a.psi complete
```

# 10¶

In [45]:

```
k,df,Ldf,ax = start('10')
```

In [46]:

```
mm = fit_model(ax[0,1],df)
```

In [47]:

```
pts = pick_pts(-48,15,0,-4,45,13)
ax[0,1].scatter(pts.x,pts.z,c='m',s=50)
```

Out[47]:

```
<matplotlib.collections.PathCollection at 0x2e303397a58>
```

In [48]:

```
df1,Ldf1,mm,ax = ut.ch_vertex(df,Ldf,pts=pts)
```

In [49]:

```
model = save_model(k,mm,model)
save_both(k,df1,Ldf1[0])
```

```
Write to .\data\hss1a\Output-02-15-2019\AT_10_hss1a.psi complete
Write to .\data\hss1a\Output-02-15-2019\ZRF_10_hss1a.psi complete
```

# 04¶

In [50]:

```
k,df,Ldf,ax = start('04')
```

In [51]:

```
k = '204'
```

In [52]:

```
pts = pick_pts(-36,11,0,-3,40,12)
ax[0,1].scatter(pts.x,pts.z,c='m',s=50)
```

Out[52]:

```
<matplotlib.collections.PathCollection at 0x2e303381f60>
```

In [53]:

```
df1,Ldf1,mm,ax = ut.ch_vertex(df,Ldf,pts=pts)
```

In [54]:

```
model = save_model(k,mm,model)
save_both(k,df1,Ldf1[0])
```

```
Write to .\data\hss1a\Output-02-15-2019\AT_204_hss1a.psi complete
Write to .\data\hss1a\Output-02-15-2019\ZRF_204_hss1a.psi complete
```

# 18¶

In [55]:

```
k,df,Ldf,ax = start('18')
```

In [56]:

```
pts = pick_pts(-36,5,-4,-8,30,4)
ax[0,1].scatter(pts.x,pts.z,c='m',s=50)
```

Out[56]:

```
<matplotlib.collections.PathCollection at 0x2e309637dd8>
```

In [57]:

```
df1,Ldf1,mm,ax = ut.ch_vertex(df,Ldf,pts=pts)
```

In [58]:

```
model = save_model(k,mm,model)
save_both(k,df1,Ldf1[0])
```

```
Write to .\data\hss1a\Output-02-15-2019\AT_18_hss1a.psi complete
Write to .\data\hss1a\Output-02-15-2019\ZRF_18_hss1a.psi complete
```

# 11¶

In [59]:

```
k,df,Ldf,ax = start('11')
```

In [60]:

```
mm = fit_model(ax[0,1],df)
```

In [61]:

```
pts = pick_pts(-50,20,0,-3,47,22)
ax[0,1].scatter(pts.x,pts.z,c='m',s=50)
```

Out[61]:

```
<matplotlib.collections.PathCollection at 0x2e30cf92240>
```

In [62]:

```
df1,Ldf1,mm,ax = ut.ch_vertex(df,Ldf,pts=pts)
```

In [63]:

```
model = save_model(k,mm,model)
save_both(k,df1,Ldf1[0])
```

```
Write to .\data\hss1a\Output-02-15-2019\AT_11_hss1a.psi complete
Write to .\data\hss1a\Output-02-15-2019\ZRF_11_hss1a.psi complete
```

# 02¶

Discard due to minimal signal

In [64]:

```
k,df,Ldf,ax = start('02')
```

# 16¶

In [65]:

```
k,df,Ldf,ax = start('16')
```

In [66]:

```
pts = pick_pts(-55,12,0,-4,45,9)
ax[0,1].scatter(pts.x,pts.z,c='m',s=50)
```

Out[66]:

```
<matplotlib.collections.PathCollection at 0x2e34ac40780>
```

In [67]:

```
df1,Ldf1,mm,ax = ut.ch_vertex(df,Ldf,pts=pts)
```

In [68]:

```
model = save_model(k,mm,model)
save_both(k,df1,Ldf1[0])
```

```
Write to .\data\hss1a\Output-02-15-2019\AT_16_hss1a.psi complete
Write to .\data\hss1a\Output-02-15-2019\ZRF_16_hss1a.psi complete
```

### Model¶

In [69]:

```
model
```

Out[69]:

|  | a | b | c |
| --- | --- | --- | --- |
| 206 | 0.009004 | -7.692383e-16 | 2.466602e-14 |
| 20 | 0.005596 | -5.923234e-17 | 2.030734e-16 |
| 17 | 0.012427 | -1.431835e-17 | -2.051165e-15 |
| 13 | 0.002645 | 8.904206e-18 | -5.127900e-16 |
| 6 | 0.001270 | -1.309015e-17 | -1.570219e-15 |
| 10 | 0.008318 | 2.484148e-17 | -6.089154e-15 |
| 204 | 0.010051 | 5.331685e-17 | -1.132164e-15 |
| 18 | 0.011503 | 1.011939e-16 | -3.913462e-15 |
| 11 | 0.010226 | 1.645257e-17 | 1.023585e-14 |
| 16 | 0.005798 | -5.794203e-17 | 4.770577e-15 |

In [70]:

```
model.to_csv(os.path.join(outdir,'model.csv'))
```
