## Supplementary material for "ΔSCOPE: A new method to quantify 3D biological structures and identify differences in zebrafish forebrain development": An archived copy of the DeltaSCOPE code repository.: hss1a_alignment.ipynb

```
# %% [code]
import numpy as np
import pandas as pd
import matplotlib.pyplot as plt

import os
import re
from imp import reload
import h5py

# %% [code]
import deltascope as cranium
import deltascope.alignment as ut 

# %% [code]
at = ".\data\hss1a\AT\Prob"
gfap = ".\data\hss1a\GFAP\Prob"
root = ".\data\hss1a"

# %% [code]
outdir = os.path.join(root,'Output-02-15-2019')
os.mkdir(outdir)

# %% [code]
Dat = {}
for f in os.listdir(at):
    if 'h5' in f:
        num  = re.findall(r'\d+',f.split('.')[0])[-1]
        Dat[num] = os.path.join(at,f)

# %% [code]
Dzrf = {}
for f in os.listdir(gfap):
    if 'h5' in f:
        num  = re.findall(r'\d+',f.split('.')[0])[-1]
        Dzrf[num] = os.path.join(gfap,f)

# %% [code]
Dbat = {}
Dbzrf = {}

# %% [markdown]
# Data Processing

# %% [code]
klist = Dat.keys()

# %% [code]
param = {
    'gthresh':0.5,
    'scale':[1,1,1],
    'microns':[0.16,0.16,0.21],
    'mthresh':0.5,
    'radius':10,
    'comp_order':[0,2,1],
    'fit_dim':['x','z'],
    'deg':2
}

# %% [code]
%%time
for k in klist:
    if k not in list(Dbat.keys()):
        Dbat[k] = ut.preprocess(Dat[k],param)
        Dbzrf[k] = ut.preprocess(Dzrf[k],param,pca=Dbat[k].pcamed,mm=Dbat[k].mm,vertex=Dbat[k].vertex)
        #Dbcaax[k] = ut.preprocess(Dcaax[k],param,pca=Dbat[k].pcamed,mm=Dbat[k].mm,vertex=Dbat[k].vertex)
        print(k)
    else:
        print(k,'already processed')

# %% [markdown]
# Functions

# %% [code]
def start(k):
    return(ut.start(k,Dbat,[Dbzrf],im=True))
def save_both(k,dfa,dfb):
    ut.save_both(k,dfa,dfb,outdir,'hss1a')

# %% [code]
model = pd.DataFrame({'a':[],'b':[],'c':[]})
def save_model(k,mm,model):
    row = pd.Series({'a':mm[0],'b':mm[1],'c':mm[2]},name=k)
    model = model.append(row)
    return(model)

# %% [code]
def fit_model(axi,df,mm=None):
    if mm == None:
        mm = np.polyfit(df.x,df.z,2)
    p = np.poly1d(mm)
    xrange = np.arange(np.min(df.x),np.max(df.x))
    axi.plot(xrange,p(xrange),c='m')
    return(mm)

# %% [code]
def pick_pts(x1,z1,vx,vz,x2,z2):
    pts = pd.DataFrame({'x':[x1,vx,x2],'z':[z1,vz,z2]})
    return(pts)

# %% [markdown]
# 06

# %% [code]
k,df,Ldf,ax = start('06')

# %% [code]
k = '206'

# %% [code]
mm = fit_model(ax[0,1],df)

# %% [code]
model = save_model(k,mm,model)
save_both(k,df,Ldf[0])

# %% [markdown]
# 20

# %% [code]
k,df,Ldf,ax = start('20')

# %% [code]
df1,Ldf1 = ut.zyswitch(df,Ldf)
ax = ut.make_graph([df1]+Ldf1)

# %% [code]
df2,Ldf2,mm,ax = ut.ch_vertex(df1,Ldf1)

# %% [code]
pts = pick_pts(-36,3,0,-5,42,4)
ax[0,1].scatter(pts.x,pts.z,c='m',s=50)

# %% [code]
df3,Ldf3,mm,ax = ut.ch_vertex(df2,Ldf2,pts=pts)

# %% [code]
model = save_model(k,mm,model)
save_both(k,df3,Ldf3[0])

# %% [markdown]
# 17

# %% [code]
k,df,Ldf,ax = start('17')

# %% [code]
df1,Ldf1,pts,ax = ut.check_pts(df,Ldf,'z')

# %% [code]
pts.iloc[0].x = 35
pts.iloc[0].z = 10
ax[0,1].scatter(pts.x,pts.z,c='y')
pts

# %% [code]
df2,Ldf2,ax = ut.revise_pts(df,Ldf,'z',pts=pts)

# %% [code]
df3,Ldf3,mm,ax = ut.ch_vertex(df2,Ldf2)

# %% [code]
pts = pick_pts(-36,13,0,-4,38,13)
ax[0,1].scatter(pts.x,pts.z,c='m',s=50)

# %% [code]
df4,Ldf4,mm,ax = ut.ch_vertex(df3,Ldf3,pts=pts)

# %% [code]
model = save_model(k,mm,model)
save_both(k,df4,Ldf4[0])

# %% [markdown]
# 13

# %% [code]
k,df,Ldf,ax = start('13')

# %% [code]
df1,Ldf1 = ut.zyswitch(df,Ldf)
ax = ut.make_graph([df1]+Ldf1)

# %% [code]
df2,Ldf2,mm,ax = ut.ch_vertex(df1,Ldf1)

# %% [code]
pts = pick_pts(-55,5,0,-3,55,5)
ax[0,1].scatter(pts.x,pts.z,c='m',s=50)

# %% [code]
df3,Ldf3,mm,ax = ut.ch_vertex(df2,Ldf2,pts=pts)

# %% [code]
model = save_model(k,mm,model)
save_both(k,df3,Ldf3[0])

# %% [code]
klist

# %% [markdown]
# 6

# %% [code]
k,df,Ldf,ax = start('6')

# %% [code]
df1,Ldf1 = ut.zyswitch(df,Ldf)
ax = ut.make_graph([df1]+Ldf1)

# %% [code]
df2,Ldf2,mm,ax = ut.ch_vertex(df1,Ldf1)

# %% [code]
model = save_model(k,mm,model)
save_both(k,df2,Ldf2[0])

# %% [markdown]
# 10

# %% [code]
k,df,Ldf,ax = start('10')

# %% [code]
mm = fit_model(ax[0,1],df)

# %% [code]
pts = pick_pts(-48,15,0,-4,45,13)
ax[0,1].scatter(pts.x,pts.z,c='m',s=50)

# %% [code]
df1,Ldf1,mm,ax = ut.ch_vertex(df,Ldf,pts=pts)

# %% [code]
model = save_model(k,mm,model)
save_both(k,df1,Ldf1[0])

# %% [markdown]
# 04

# %% [code]
k,df,Ldf,ax = start('04')

# %% [code]
k = '204'

# %% [code]
pts = pick_pts(-36,11,0,-3,40,12)
ax[0,1].scatter(pts.x,pts.z,c='m',s=50)

# %% [code]
df1,Ldf1,mm,ax = ut.ch_vertex(df,Ldf,pts=pts)

# %% [code]
model = save_model(k,mm,model)
save_both(k,df1,Ldf1[0])

# %% [markdown]
# 18

# %% [code]
k,df,Ldf,ax = start('18')

# %% [code]
pts = pick_pts(-36,5,-4,-8,30,4)
ax[0,1].scatter(pts.x,pts.z,c='m',s=50)

# %% [code]
df1,Ldf1,mm,ax = ut.ch_vertex(df,Ldf,pts=pts)

# %% [code]
model = save_model(k,mm,model)
save_both(k,df1,Ldf1[0])

# %% [markdown]
# 11

# %% [code]
k,df,Ldf,ax = start('11')

# %% [code]
mm = fit_model(ax[0,1],df)

# %% [code]
pts = pick_pts(-50,20,0,-3,47,22)
ax[0,1].scatter(pts.x,pts.z,c='m',s=50)

# %% [code]
df1,Ldf1,mm,ax = ut.ch_vertex(df,Ldf,pts=pts)

# %% [code]
model = save_model(k,mm,model)
save_both(k,df1,Ldf1[0])

# %% [markdown]
# 02

# %% [markdown]
Discard due to minimal signal

# %% [code]
k,df,Ldf,ax = start('02')

# %% [markdown]
# 16

# %% [code]
k,df,Ldf,ax = start('16')

# %% [code]
pts = pick_pts(-55,12,0,-4,45,9)
ax[0,1].scatter(pts.x,pts.z,c='m',s=50)

# %% [code]
df1,Ldf1,mm,ax = ut.ch_vertex(df,Ldf,pts=pts)

# %% [code]
model = save_model(k,mm,model)
save_both(k,df1,Ldf1[0])

# %% [markdown]
# Model

# %% [code]
model

# %% [code]
model.to_csv(os.path.join(outdir,'model.csv'))
```
