## Supplementary material for "ΔSCOPE: A new method to quantify 3D biological structures and identify differences in zebrafish forebrain development": An archived copy of the DeltaSCOPE code repository.: hss1ayot_alignment.html

2018-08-13\_hss1ayot\_alignment-Rerun


In [1]:

```
import numpy as np
import pandas as pd
import matplotlib.pyplot as plt

import os
import re
from imp import reload
import h5py
```

In [2]:

```
import deltascope as cranium
import deltascope.alignment as ut
```

In [3]:

```
gfap = ".\data\hss1ayot\GFAP\Prob"
at = ".\data\hss1ayot\AT\Prob"
root = ".\data\hss1ayot"
```

In [4]:

```
outdir = os.path.join(root,'Output-02-15-2019')
os.mkdir(outdir)
```

In [5]:

```
Dat = {}
for f in os.listdir(at):
    if 'h5' in f:
        num  = re.findall(r'\d+',f.split('.')[0])[-1]
        Dat[num] = os.path.join(at,f)
```

In [6]:

```
Dzrf = {}
for f in os.listdir(gfap):
    if 'h5' in f:
        num  = re.findall(r'\d+',f.split('.')[0])[-1]
        Dzrf[num] = os.path.join(gfap,f)
```

In [7]:

```
Dbat = {}
Dbzrf = {}
```

### Data processing¶

In [8]:

```
klist = Dat.keys()
```

In [9]:

```
param = {
    'gthresh':0.5,
    'scale':[1,1,1],
    'microns':[0.16,0.16,0.21],
    'mthresh':0.5,
    'radius':10,
    'comp_order':[0,2,1],
    'fit_dim':['x','z'],
    'deg':2
}
```

In [10]:

```
%%time
for k in klist:
    if k not in list(Dbat.keys()):
        Dbat[k] = ut.preprocess(Dat[k],param)
        Dbzrf[k] = ut.preprocess(Dzrf[k],param,pca=Dbat[k].pcamed,mm=Dbat[k].mm,vertex=Dbat[k].vertex)
        #Dbcaax[k] = ut.preprocess(Dcaax[k],param,pca=Dbat[k].pcamed,mm=Dbat[k].mm,vertex=Dbat[k].vertex)
        print(k)
    else:
        print(k,'already processed')
```

```
C:\Users\zfishlab\AppData\Local\Continuum\anaconda3\envs\test\lib\site-packages\skimage\util\dtype.py:141: UserWarning: Possible precision loss when converting from float32 to uint8
  .format(dtypeobj_in, dtypeobj_out))
```

```
010
```

```
C:\Users\zfishlab\AppData\Local\Continuum\anaconda3\envs\test\lib\site-packages\skimage\util\dtype.py:141: UserWarning: Possible precision loss when converting from float32 to uint8
  .format(dtypeobj_in, dtypeobj_out))
```

```
012
```

```
C:\Users\zfishlab\AppData\Local\Continuum\anaconda3\envs\test\lib\site-packages\skimage\util\dtype.py:141: UserWarning: Possible precision loss when converting from float32 to uint8
  .format(dtypeobj_in, dtypeobj_out))
```

```
014
```

```
C:\Users\zfishlab\AppData\Local\Continuum\anaconda3\envs\test\lib\site-packages\skimage\util\dtype.py:141: UserWarning: Possible precision loss when converting from float32 to uint8
  .format(dtypeobj_in, dtypeobj_out))
```

```
016
```

```
C:\Users\zfishlab\AppData\Local\Continuum\anaconda3\envs\test\lib\site-packages\skimage\util\dtype.py:141: UserWarning: Possible precision loss when converting from float32 to uint8
  .format(dtypeobj_in, dtypeobj_out))
```

```
02
```

```
C:\Users\zfishlab\AppData\Local\Continuum\anaconda3\envs\test\lib\site-packages\skimage\util\dtype.py:141: UserWarning: Possible precision loss when converting from float32 to uint8
  .format(dtypeobj_in, dtypeobj_out))
```

```
03
```

```
C:\Users\zfishlab\AppData\Local\Continuum\anaconda3\envs\test\lib\site-packages\skimage\util\dtype.py:141: UserWarning: Possible precision loss when converting from float32 to uint8
  .format(dtypeobj_in, dtypeobj_out))
```

```
06
```

```
C:\Users\zfishlab\AppData\Local\Continuum\anaconda3\envs\test\lib\site-packages\skimage\util\dtype.py:141: UserWarning: Possible precision loss when converting from float32 to uint8
  .format(dtypeobj_in, dtypeobj_out))
```

```
07
```

```
C:\Users\zfishlab\AppData\Local\Continuum\anaconda3\envs\test\lib\site-packages\skimage\util\dtype.py:141: UserWarning: Possible precision loss when converting from float32 to uint8
  .format(dtypeobj_in, dtypeobj_out))
```

```
08
```

```
C:\Users\zfishlab\AppData\Local\Continuum\anaconda3\envs\test\lib\site-packages\skimage\util\dtype.py:141: UserWarning: Possible precision loss when converting from float32 to uint8
  .format(dtypeobj_in, dtypeobj_out))
```

```
09
```

```
C:\Users\zfishlab\AppData\Local\Continuum\anaconda3\envs\test\lib\site-packages\skimage\util\dtype.py:141: UserWarning: Possible precision loss when converting from float32 to uint8
  .format(dtypeobj_in, dtypeobj_out))
```

```
7
Wall time: 13min 33s
```

### Functions¶

In [11]:

```
def start(k):
    return(ut.start(k,Dbat,[Dbzrf],im=True))
def save_both(k,dfa,dfb):
    ut.save_both(k,dfa,dfb,outdir,'hss1ayot')
```

In [12]:

```
model = pd.DataFrame({'a':[],'b':[],'c':[]})
def save_model(k,mm,model):
    row = pd.Series({'a':mm[0],'b':mm[1],'c':mm[2]},name=k)
    model = model.append(row)
    return(model)
```

In [13]:

```
def fit_model(axi,df,mm=None):
    if mm == None:
        mm = np.polyfit(df.x,df.z,2)
    p = np.poly1d(mm)
    xrange = np.arange(np.min(df.x),np.max(df.x))
    axi.plot(xrange,p(xrange),c='m')
    return(mm)
```

In [14]:

```
def pick_pts(x1,z1,vx,vz,x2,z2):
    pts = pd.DataFrame({'x':[x1,vx,x2],'z':[z1,vz,z2]})
    return(pts)
```

# 3¶

In [15]:

```
k,df,Ldf,ax = start('3')
```

In [16]:

```
mm = fit_model(ax[0,1],df)
```

In [17]:

```
model = save_model(k,mm,model)
save_both(k,df,Ldf[0])
```

```
Write to .\data\hss1ayot\Output-02-15-2019\AT_3_hss1ayot.psi complete
Write to .\data\hss1ayot\Output-02-15-2019\ZRF_3_hss1ayot.psi complete
```

# 11¶

In [18]:

```
k,df,Ldf,ax = start('11')
```

In [19]:

```
mm = fit_model(ax[0,1],df)
```

In [20]:

```
model = save_model(k,mm,model)
save_both(k,df,Ldf[0])
```

```
Write to .\data\hss1ayot\Output-02-15-2019\AT_11_hss1ayot.psi complete
Write to .\data\hss1ayot\Output-02-15-2019\ZRF_11_hss1ayot.psi complete
```

In [21]:

```
klist
```

Out[21]:

```
dict_keys(['010', '012', '014', '016', '02', '03', '06', '07', '08', '09', '10', '11', '12', '13', '14', '16', '2', '3', '5', '7'])
```

# 010¶

In [22]:

```
k,df,Ldf,ax = start('010')
```

In [23]:

```
df1,Ldf1 = ut.zyswitch(df,Ldf)
ax = ut.make_graph([df1]+Ldf1)
```

In [24]:

```
mm = fit_model(ax[0,1],df1)
```

In [25]:

```
model = save_model(k,mm,model)
save_both(k,df1,Ldf1[0])
```

```
Write to .\data\hss1ayot\Output-02-15-2019\AT_010_hss1ayot.psi complete
Write to .\data\hss1ayot\Output-02-15-2019\ZRF_010_hss1ayot.psi complete
```

# 09¶

Discard; too little signal

In [26]:

```
k,df,Ldf,ax = start('09')
```

# 016¶

discard; limited signal

In [27]:

```
k,df,Ldf,ax = start('016')
```

# 02¶

In [28]:

```
k,df,Ldf,ax = start('02')
```

In [29]:

```
df = Dbat[k].df_thresh
Ldf = [Dbzrf[k].df_thresh]
ax = ut.make_graph([df]+Ldf)
```

In [30]:

```
df1,Ldf1,mm,ax = ut.ch_vertex(df,Ldf)
```

In [31]:

```
model = save_model(k,mm,model)
save_both(k,df1,Ldf1[0])
```

```
Write to .\data\hss1ayot\Output-02-15-2019\AT_02_hss1ayot.psi complete
Write to .\data\hss1ayot\Output-02-15-2019\ZRF_02_hss1ayot.psi complete
```

In [32]:

```
klist
```

Out[32]:

```
dict_keys(['010', '012', '014', '016', '02', '03', '06', '07', '08', '09', '10', '11', '12', '13', '14', '16', '2', '3', '5', '7'])
```

# 14¶

In [33]:

```
k,df,Ldf,ax = start('14')
```

In [34]:

```
df = Dbat[k].df_thresh
Ldf = [Dbzrf[k].df_thresh]
ax = ut.make_graph([df]+Ldf)
```

In [35]:

```
pts = pick_pts(0,12,34,3,72,18)
ax[0,1].scatter(pts.x,pts.z,c='m',s=50)
```

Out[35]:

```
<matplotlib.collections.PathCollection at 0x21813d8e898>
```

In [36]:

```
df1,Ldf1,mm,ax = ut.ch_vertex(df,Ldf,pts=pts)
```

In [37]:

```
df2,Ldf2,pts,ax = ut.check_pts(df1,Ldf1,'z')
```

In [38]:

```
pts.iloc[0].x = 41
pts.iloc[0].z = 15
pts.iloc[1].z = 7
ax[0,1].scatter(pts.x,pts.z,c='y')
pts
```

Out[38]:

|  | x | z |
| --- | --- | --- |
| 0 | 41.000000 | 15.0 |
| 1 | -31.450704 | 7.0 |

In [39]:

```
df3,Ldf3,ax = ut.revise_pts(df1,Ldf1,'z',pts=pts)
```

In [40]:

```
pts = pick_pts(-30,10,5,-1,43,10)
ax[1,1].scatter(pts.x,pts.z,c='m',s=50)
```

Out[40]:

```
<matplotlib.collections.PathCollection at 0x2181547f860>
```

In [41]:

```
df4,Ldf4,mm,ax = ut.ch_vertex(df3,Ldf3,pts=pts)
```

In [42]:

```
model = save_model(k,mm,model)
save_both(k,df4,Ldf4[0])
```

```
Write to .\data\hss1ayot\Output-02-15-2019\AT_14_hss1ayot.psi complete
Write to .\data\hss1ayot\Output-02-15-2019\ZRF_14_hss1ayot.psi complete
```

# 10¶

In [43]:

```
k,df,Ldf,ax = start('10')
```

In [44]:

```
pts = pick_pts(-25,5,3,-5,38,6)
ax[0,1].scatter(pts.x,pts.z,c='m',s=50)
```

Out[44]:

```
<matplotlib.collections.PathCollection at 0x218155c7668>
```

In [45]:

```
df1,Ldf1,mm,ax = ut.ch_vertex(df,Ldf,pts=pts)
```

In [46]:

```
model = save_model(k,mm,model)
save_both(k,df1,Ldf1[0])
```

```
Write to .\data\hss1ayot\Output-02-15-2019\AT_10_hss1ayot.psi complete
Write to .\data\hss1ayot\Output-02-15-2019\ZRF_10_hss1ayot.psi complete
```

# 014¶

In [47]:

```
k,df,Ldf,ax = start('014')
```

In [48]:

```
df = Dbat[k].df_thresh
Ldf = [Dbzrf[k].df_thresh]
ax = ut.make_graph([df]+Ldf)
```

In [49]:

```
pts = pick_pts(0,10,35,2,69,10)
ax[0,1].scatter(pts.x,pts.z,c='m',s=50)
```

Out[49]:

```
<matplotlib.collections.PathCollection at 0x218150f2048>
```

In [50]:

```
df1,Ldf1,mm,ax = ut.ch_vertex(df,Ldf,pts=pts)
```

In [51]:

```
model = save_model(k,mm,model)
save_both(k,df1,Ldf1[0])
```

```
Write to .\data\hss1ayot\Output-02-15-2019\AT_014_hss1ayot.psi complete
Write to .\data\hss1ayot\Output-02-15-2019\ZRF_014_hss1ayot.psi complete
```

# 5¶

In [52]:

```
k,df,Ldf,ax = start('5')
```

In [53]:

```
df = Dbat[k].df_thresh
Ldf = [Dbzrf[k].df_thresh]
ax = ut.make_graph([df]+Ldf)
```

In [54]:

```
pts = pick_pts(0,16,36,1,81,7)
ax[0,1].scatter(pts.x,pts.z,c='m',s=50)
```

Out[54]:

```
<matplotlib.collections.PathCollection at 0x21815bb2a90>
```

In [55]:

```
df1,Ldf1,mm,ax = ut.ch_vertex(df,Ldf,pts=pts)
```

In [56]:

```
model = save_model(k,mm,model)
save_both(k,df1,Ldf1[0])
```

```
Write to .\data\hss1ayot\Output-02-15-2019\AT_5_hss1ayot.psi complete
Write to .\data\hss1ayot\Output-02-15-2019\ZRF_5_hss1ayot.psi complete
```

# 07¶

In [57]:

```
k,df,Ldf,ax = start('07')
```

In [58]:

```
df = Dbat[k].df_thresh
Ldf = [Dbzrf[k].df_thresh]
ax = ut.make_graph([df]+Ldf)
```

In [59]:

```
pts = pick_pts(0,15,30,6,64,16)
ax[0,1].scatter(pts.x,pts.z,c='m',s=50)
```

Out[59]:

```
<matplotlib.collections.PathCollection at 0x2181a7fe2b0>
```

In [60]:

```
df1,Ldf1,mm,ax = ut.ch_vertex(df,Ldf,pts=pts)
```

In [61]:

```
model = save_model(k,mm,model)
save_both(k,df1,Ldf1[0])
```

```
Write to .\data\hss1ayot\Output-02-15-2019\AT_07_hss1ayot.psi complete
Write to .\data\hss1ayot\Output-02-15-2019\ZRF_07_hss1ayot.psi complete
```

# 12¶

In [62]:

```
k,df,Ldf,ax = start('12')
```

In [63]:

```
df = Dbat[k].df_thresh
Ldf = [Dbzrf[k].df_thresh]
ax = ut.make_graph([df]+Ldf)
```

In [64]:

```
df1,Ldf1,pts,ax = ut.check_pts(df,Ldf,'z')
```

In [65]:

```
pts.iloc[0].z = 12
ax[0,1].scatter(pts.x,pts.z,c='y')
pts
```

Out[65]:

|  | x | z |
| --- | --- | --- |
| 0 | 0.00 | 12.00 |
| 1 | 78.72 | 15.33 |

In [66]:

```
df2,Ldf2,ax = ut.revise_pts(df,Ldf,'z',pts=pts)
```

In [67]:

```
df3,Ldf3 = ut.flip(df2,Ldf2)
ax = ut.make_graph([df3]+Ldf3)
```

In [68]:

```
pts = pick_pts(-78,12,-36,0,0,10)
ax[0,1].scatter(pts.x,pts.z,c='m',s=50)
```

Out[68]:

```
<matplotlib.collections.PathCollection at 0x2187eacf908>
```

In [69]:

```
df4,Ldf4,mm,ax = ut.ch_vertex(df3,Ldf3,pts=pts)
```

In [70]:

```
model = save_model(k,mm,model)
save_both(k,df4,Ldf4[0])
```

```
Write to .\data\hss1ayot\Output-02-15-2019\AT_12_hss1ayot.psi complete
Write to .\data\hss1ayot\Output-02-15-2019\ZRF_12_hss1ayot.psi complete
```

# 03¶

In [71]:

```
k,df,Ldf,ax = start('03')
```

In [72]:

```
mm = fit_model(ax[0,1],df)
```

In [73]:

```
pts = pick_pts(-35,11,3,-2,40,11)
ax[0,1].scatter(pts.x,pts.z,c='m',s=50)
```

Out[73]:

```
<matplotlib.collections.PathCollection at 0x2187ed9bc88>
```

In [74]:

```
df1,Ldf1,mm,ax = ut.ch_vertex(df,Ldf,pts=pts)
```

In [75]:

```
model = save_model(k,mm,model)
save_both(k,df1,Ldf1[0])
```

```
Write to .\data\hss1ayot\Output-02-15-2019\AT_03_hss1ayot.psi complete
Write to .\data\hss1ayot\Output-02-15-2019\ZRF_03_hss1ayot.psi complete
```

# 16¶

In [76]:

```
k,df,Ldf,ax = start('16')
```

In [77]:

```
df = Dbat[k].df_thresh
Ldf = [Dbzrf[k].df_thresh]
ax = ut.make_graph([df]+Ldf)
```

In [78]:

```
df1,Ldf1,mm,ax = ut.ch_vertex(df,Ldf)
```

In [79]:

```
model = save_model(k,mm,model)
save_both(k,df1,Ldf1[0])
```

```
Write to .\data\hss1ayot\Output-02-15-2019\AT_16_hss1ayot.psi complete
Write to .\data\hss1ayot\Output-02-15-2019\ZRF_16_hss1ayot.psi complete
```

# 08¶

In [80]:

```
k,df,Ldf,ax = start('08')
```

In [81]:

```
mm = fit_model(ax[0,1],df)
```

In [82]:

```
model = save_model(k,mm,model)
save_both(k,df,Ldf[0])
```

```
Write to .\data\hss1ayot\Output-02-15-2019\AT_08_hss1ayot.psi complete
Write to .\data\hss1ayot\Output-02-15-2019\ZRF_08_hss1ayot.psi complete
```

# 13¶

In [83]:

```
k,df,Ldf,ax = start('13')
```

In [84]:

```
df = Dbat[k].df_thresh
Ldf = [Dbzrf[k].df_thresh]
ax = ut.make_graph([df]+Ldf)
```

In [85]:

```
df1,Ldf1,mm,ax = ut.ch_vertex(df,Ldf)
```

In [86]:

```
model = save_model(k,mm,model)
save_both(k,df1,Ldf1[0])
```

```
Write to .\data\hss1ayot\Output-02-15-2019\AT_13_hss1ayot.psi complete
Write to .\data\hss1ayot\Output-02-15-2019\ZRF_13_hss1ayot.psi complete
```

# 2¶

In [87]:

```
k,df,Ldf,ax = start('2')
```

In [88]:

```
df = Dbat[k].df_thresh
Ldf = [Dbzrf[k].df_thresh]
ax = ut.make_graph([df]+Ldf)
```

In [89]:

```
pts = pick_pts(0,18,45,2,90,15)
ax[0,1].scatter(pts.x,pts.z,c='m',s=50)
```

Out[89]:

```
<matplotlib.collections.PathCollection at 0x2187e9efc50>
```

In [90]:

```
df1,Ldf1,mm,ax = ut.ch_vertex(df,Ldf,pts=pts)
```

In [91]:

```
model = save_model(k,mm,model)
save_both(k,df1,Ldf1[0])
```

```
Write to .\data\hss1ayot\Output-02-15-2019\AT_2_hss1ayot.psi complete
Write to .\data\hss1ayot\Output-02-15-2019\ZRF_2_hss1ayot.psi complete
```

# 7¶

In [92]:

```
k,df,Ldf,ax = start('7')
```

In [93]:

```
df = Dbat[k].df_thresh
Ldf = [Dbzrf[k].df_thresh]
ax = ut.make_graph([df]+Ldf)
```

In [94]:

```
df1,Ldf1,pts,ax = ut.check_pts(df,Ldf,'z')
```

In [95]:

```
pts.iloc[0].z = 4
ax[0,1].scatter(pts.x,pts.z,c='y')
pts
```

Out[95]:

|  | x | z |
| --- | --- | --- |
| 0 | 75.2 | 4.00 |
| 1 | 0.0 | 11.13 |

In [96]:

```
df2,Ldf2,ax = ut.revise_pts(df,Ldf,'z',pts=pts)
```

In [97]:

```
df3,Ldf3,mm,ax = ut.ch_vertex(df2,Ldf2)
```

In [98]:

```
pts = pick_pts(-34,7,4,-1,42,5)
ax[0,1].scatter(pts.x,pts.z,c='m',s=50)
```

Out[98]:

```
<matplotlib.collections.PathCollection at 0x21917835c18>
```

In [99]:

```
df4,Ldf4,mm,ax = ut.ch_vertex(df3,Ldf3,pts=pts)
```

In [100]:

```
model = save_model(k,mm,model)
save_both(k,df4,Ldf4[0])
```

```
Write to .\data\hss1ayot\Output-02-15-2019\AT_7_hss1ayot.psi complete
Write to .\data\hss1ayot\Output-02-15-2019\ZRF_7_hss1ayot.psi complete
```

# 012¶

In [101]:

```
k,df,Ldf,ax = start('012')
```

In [102]:

```
df = Dbat[k].df_thresh
Ldf = [Dbzrf[k].df_thresh]
ax = ut.make_graph([df]+Ldf)
```

In [103]:

```
pts = pick_pts(0,18,44,2,94,19)
ax[0,1].scatter(pts.x,pts.z,c='m',s=50)
```

Out[103]:

```
<matplotlib.collections.PathCollection at 0x2187ed1edd8>
```

In [104]:

```
df1,Ldf1,mm,ax = ut.ch_vertex(df,Ldf,pts=pts)
```

In [105]:

```
model = save_model(k,mm,model)
save_both(k,df1,Ldf1[0])
```

```
Write to .\data\hss1ayot\Output-02-15-2019\AT_012_hss1ayot.psi complete
Write to .\data\hss1ayot\Output-02-15-2019\ZRF_012_hss1ayot.psi complete
```

# 06¶

In [106]:

```
k,df,Ldf,ax = start('06')
```

Discard not enough zrf signal

In [107]:

```
df = Dbat[k].df_thresh
Ldf = [Dbzrf[k].df_thresh]
ax = ut.make_graph([df]+Ldf)
```

In [108]:

```
model
```

Out[108]:

|  | a | b | c |
| --- | --- | --- | --- |
| 3 | 0.005765 | -7.518081e-17 | 5.468862e-15 |
| 11 | 0.010150 | -7.641207e-17 | 2.161441e-15 |
| 010 | 0.000698 | -6.484719e-02 | -2.723168e-01 |
| 02 | 0.004502 | -2.600388e-17 | 3.801242e-15 |
| 14 | 0.008271 | 6.825507e-17 | 3.076827e-15 |
| 10 | 0.010658 | -1.757130e-16 | -1.812101e-14 |
| 014 | 0.006723 | 2.684945e-16 | 1.025566e-15 |
| 5 | 0.006790 | -2.503965e-16 | 2.813754e-15 |
| 07 | 0.009283 | -2.090030e-16 | 1.176538e-17 |
| 12 | 0.007224 | -2.721652e-16 | 4.792220e-15 |
| 03 | 0.009246 | -3.833467e-17 | 1.025582e-15 |
| 16 | 0.006386 | -9.000242e-17 | 9.792146e-16 |
| 08 | 0.007753 | -1.034213e-16 | -1.531882e-15 |
| 13 | 0.009048 | -2.644279e-16 | -4.188198e-15 |
| 2 | 0.007160 | -2.204101e-16 | -7.797574e-15 |
| 7 | 0.004848 | -7.684438e-17 | -9.270975e-17 |
| 012 | 0.007485 | -1.492182e-17 | 4.138972e-15 |

In [109]:

```
model.to_csv(os.path.join(outdir,'model.csv'))
```

In [110]:

```
outdir
```

Out[110]:

```
'.\\data\\hss1ayot\\Output-02-15-2019'
```

In [111]:

```
modelout = pd.read_csv(os.path.join(outdir,'model.csv'))
```

In [112]:

```
modelout
```

Out[112]:

|  | Unnamed: 0 | a | b | c |
| --- | --- | --- | --- | --- |
| 0 | 3 | 0.005765 | -7.518081e-17 | 5.468862e-15 |
| 1 | 11 | 0.010150 | -7.641207e-17 | 2.161441e-15 |
| 2 | 10 | 0.000698 | -6.484719e-02 | -2.723168e-01 |
| 3 | 2 | 0.004502 | -2.600388e-17 | 3.801242e-15 |
| 4 | 14 | 0.008271 | 6.825507e-17 | 3.076827e-15 |
| 5 | 10 | 0.010658 | -1.757130e-16 | -1.812101e-14 |
| 6 | 14 | 0.006723 | 2.684945e-16 | 1.025566e-15 |
| 7 | 5 | 0.006790 | -2.503965e-16 | 2.813754e-15 |
| 8 | 7 | 0.009283 | -2.090030e-16 | 1.176538e-17 |
| 9 | 12 | 0.007224 | -2.721652e-16 | 4.792220e-15 |
| 10 | 3 | 0.009246 | -3.833467e-17 | 1.025582e-15 |
| 11 | 16 | 0.006386 | -9.000242e-17 | 9.792146e-16 |
| 12 | 8 | 0.007753 | -1.034213e-16 | -1.531882e-15 |
| 13 | 13 | 0.009048 | -2.644279e-16 | -4.188198e-15 |
| 14 | 2 | 0.007160 | -2.204101e-16 | -7.797574e-15 |
| 15 | 7 | 0.004848 | -7.684438e-17 | -9.270975e-17 |
| 16 | 12 | 0.007485 | -1.492182e-17 | 4.138972e-15 |

In [113]:

```
model2 = pd.read_csv(os.path.join(outdir,'model.csv'),index_col='Unnamed: 0',dtype={'Unnamed: 0':str})
```

In [114]:

```
model2
```

Out[114]:

|  | a | b | c |
| --- | --- | --- | --- |
| 3 | 0.005765 | -7.518081e-17 | 5.468862e-15 |
| 11 | 0.010150 | -7.641207e-17 | 2.161441e-15 |
| 10 | 0.000698 | -6.484719e-02 | -2.723168e-01 |
| 2 | 0.004502 | -2.600388e-17 | 3.801242e-15 |
| 14 | 0.008271 | 6.825507e-17 | 3.076827e-15 |
| 10 | 0.010658 | -1.757130e-16 | -1.812101e-14 |
| 14 | 0.006723 | 2.684945e-16 | 1.025566e-15 |
| 5 | 0.006790 | -2.503965e-16 | 2.813754e-15 |
| 7 | 0.009283 | -2.090030e-16 | 1.176538e-17 |
| 12 | 0.007224 | -2.721652e-16 | 4.792220e-15 |
| 3 | 0.009246 | -3.833467e-17 | 1.025582e-15 |
| 16 | 0.006386 | -9.000242e-17 | 9.792146e-16 |
| 8 | 0.007753 | -1.034213e-16 | -1.531882e-15 |
| 13 | 0.009048 | -2.644279e-16 | -4.188198e-15 |
| 2 | 0.007160 | -2.204101e-16 | -7.797574e-15 |
| 7 | 0.004848 | -7.684438e-17 | -9.270975e-17 |
| 12 | 0.007485 | -1.492182e-17 | 4.138972e-15 |
