## Supplementary material for "ΔSCOPE: A new method to quantify 3D biological structures and identify differences in zebrafish forebrain development": An archived copy of the DeltaSCOPE code repository.: landmarks.html


In [13]:

```
import deltascope as ds
import deltascope.alignment as ut

import numpy as np
import pandas as pd
import matplotlib.pyplot as plt

from sklearn.preprocessing import normalize
from scipy.optimize import minimize

import os
import tqdm
import json
import time
```

### Import raw data¶

The user needs to specify the directories containing the data of interest. Each sample type should have a key which corresponds to the directory path. Additionally, each object should have a list that includes the channels of interest.

In [2]:

```
# --------------------------------
# -------- User input ------------
# --------------------------------

data = {
    # Specify sample type key
    'wt': {
        # Specify path to data directory
        'path': './../yot_experiment/data/Output_wt_01-23-16-06',
        # Specify which channels are in the directory and are of interest
        'channels': ['AT','ZRF']
    },
    'yot': {
        'path': './../yot_experiment/data/Output_yot_01-24-14-22',
        'channels': ['AT','ZRF']
    },
    'hss1a': {
        'path': './data/hss1a/Output-02-15-2019',
        'channels': ['AT','ZRF']
    },
    'hss1ayot': {
        'path': './data/hss1ayot/Output-02-15-2019',
        'channels': ['AT','ZRF']
    }
}
```

In [3]:

```
data_pairs = []
for s in data.keys():
    for c in data[s]['channels']:
        data_pairs.append((s,c))
```

We can now read in all datafiles specified by the `data` dictionary above.

In [4]:

```
D = {}
for s in data.keys():
    D[s] = {}
    for c in data[s]['channels']:
        D[s][c] = ds.read_psi_to_dict(data[s]['path'],c)
```

```
100%|██████████| 77/77 [00:14<00:00,  5.47it/s]
100%|██████████| 77/77 [00:10<00:00,  2.62it/s]
100%|██████████| 71/71 [00:03<00:00, 18.02it/s]
100%|██████████| 71/71 [00:05<00:00,  7.63it/s]
100%|██████████| 21/21 [00:00<00:00, 47.12it/s]
100%|██████████| 21/21 [00:00<00:00, 23.41it/s]
100%|██████████| 35/35 [00:00<00:00, 83.38it/s]
100%|██████████| 35/35 [00:00<00:00, 44.11it/s]
```

Display the numer of samples for each sample type.

In [6]:

```
len(D['wt']['AT'].keys()),len(D['yot']['AT'].keys()),len(D['hss1a']['AT'].keys()),len(D['hss1ayot']['AT'].keys())
```

Out[6]:

```
(37, 34, 10, 17)
```

### Landmarks¶

Calculate landmark bins based on user input parameters and the previously specified control sample.

In [10]:

```
lm = ds.landmarks(percbins=percbins, rnull=np.nan)
lm.calc_bins(D[s_ctrl][c_ctrl], anum, theta_step)

print('Alpha bins')
print(lm.acbins)
print('Theta bins')
print(lm.tbins)
```

```
Alpha bins
[-83.53412655 -76.57294933 -69.61177212 -62.65059491 -55.6894177
 -48.72824048 -41.76706327 -34.80588606 -27.84470885 -20.88353164
 -13.92235442  -6.96117721   0.           6.96117721  13.92235442
  20.88353164  27.84470885  34.80588606  41.76706327  48.72824048
  55.6894177   62.65059491  69.61177212  76.57294933  83.53412655]
Theta bins
[-3.14159265 -2.35619449 -1.57079633 -0.78539816  0.          0.78539816
  1.57079633  2.35619449  3.14159265]
```

In [14]:

```
lmdf = pd.DataFrame()

# Loop through each pair of stype and channels
for s,c in tqdm.tqdm(data_pairs):
    print(s,c)
    # Calculate landmarks for each sample with this data pair
    for k,df in tqdm.tqdm(D[s][c].items()):
        lmdf = lm.calc_perc(df, k, '-'.join([s,c]), lmdf)
        
# Set timestamp for saving data
tstamp = time.strftime("%m-%d-%H-%M",time.localtime())
        
# Save completed landmarks to a csv file
lmdf.to_csv(tstamp+'_landmarks.csv')

# Save landmark bins to json file
bins = {
    'acbins':list(lm.acbins),
    'tbins':list(lm.tbins)
}
with open(tstamp+'_landmarks_bins.json', 'w') as outfile:
    json.dump(bins, outfile)
```
