## Supplementary material for "ΔSCOPE: A new method to quantify 3D biological structures and identify differences in zebrafish forebrain development": An archived copy of the DeltaSCOPE code repository.: plotting.html


In [1]:

```
import os
import json
import datetime

import tqdm
import glob
from imp import reload

import pandas as pd
import numpy as np
import matplotlib.pyplot as plt
import scipy.stats as stats
from sklearn.preprocessing import normalize
from scipy.optimize import minimize

import deltascope as ds
```

```
C:\Users\zfishlab\AppData\Local\Continuum\anaconda3\envs\deltascope\lib\site-packages\sklearn\ensemble\weight_boosting.py:29: DeprecationWarning: numpy.core.umath_tests is an internal NumPy module and should not be imported. It will be removed in a future NumPy release.
  from numpy.core.umath_tests import inner1d
```

In [2]:

```
# --------------------------------
# -------- User input ------------
# --------------------------------

# Specify path to exported landmark data
lmpath = glob.glob('*landmarks.csv')
binpath = glob.glob('*landmarks_bins.json')
print(lmpath,binpath)

# Pick the correct path from the list
lmpath = lmpath[0]
binpath = binpath[0]
```

```
['03-09-17-14_landmarks.csv'] ['03-09-17-14_landmarks_bins.json']
```

In [3]:

```
# Load landmarks from csv
oldlm = pd.read_csv(lmpath)
```

In [4]:

```
# Load landmark bins 
with open(binpath,'r') as f:
    bins = json.load(f)
acbins = bins['acbins']
tbins = bins['tbins']
```

In [5]:

```
colors = ['#41ab5d','#ef3b2c','#00441b','#67000d']
tarr = np.round(tbins,2)
xarr = np.round(acbins,2)
tpairs = [[tarr[4],tarr[0]],[tarr[5],tarr[1]],[tarr[6],tarr[2]],[tarr[7],tarr[3]]]
```

### Restructure Data¶

We will sort landmark data according to stype and organize it in a two tiered dictionary according to sample type (s) and channel (c).

In [6]:

```
oldlm.stype.unique()
```

Out[6]:

```
array(['wt-AT', 'wt-ZRF', 'yot-AT', 'yot-ZRF', 'hss1a-AT', 'hss1a-ZRF',
       'hss1ayot-AT', 'hss1ayot-ZRF'], dtype=object)
```

In [7]:

```
Dlm = {}
for stype in tqdm.tqdm(oldlm.stype.unique()):
    # These two lines may need to be modified based on stype structure
    s = stype.split('-')[0]
    c = stype.split('-')[-1]
    
    # Add sample type dictionary if not already present
    if s not in Dlm.keys():
        Dlm[s] = {}
    
    # Save sample specific landmark data to dictionary
    Dlm[s][c] = oldlm[oldlm.stype==stype]
```

```
100%|██████████| 8/8 [00:00<00:00, 500.27it/s]
```

### Set up graph data¶

In [8]:

```
comp = [['wt','yot'],['wt','hss1a'],['wt','hss1ayot'],['yot','hss1a'],['yot','hss1ayot'],['hss1a','hss1ayot']]
```

In [9]:

```
qc1 = ['#1b9e77','#d95f02','#7570b3','#e7298a','#66a61e']
qc2 = ['#e41a1c','#377eb8','#4daf4a','#984ea3','#ff7f00']
```

In [10]:

```
gdata = {}
for i,a in enumerate(Dlm.keys()):
    gdata[a] = {}
    
    gdata[a]['AT'] = ds.graphData(Dlm[a]['AT'][Dlm[a]['AT'].stype==a+'-'+'AT'],qc1[i])
    gdata[a]['ZRF'] = ds.graphData(Dlm[a]['ZRF'][Dlm[a]['ZRF'].stype==a+'-'+'ZRF'],qc1[i])
```

In [11]:

```
Lstype = list(Dlm.keys())
Lstype
```

Out[11]:

```
['wt', 'yot', 'hss1a', 'hss1ayot']
```

In [21]:

```
crop = 40 # microns
legend = False
save = True
a = 0.3
pthresh = 0.01
```

#### Individual at/zrf graphs¶

### R¶

In [32]:

```
fig,axr = plt.subplots(4,4,figsize=(16,10),sharey=True,sharex=True)

if crop is not None:
    mask = np.where((xarr>-crop)&(xarr<crop) == True)[0]
    xmin = mask.min()
    xmax = mask.max()
    xarrcr = xarr[xmin:xmax+1]
else:
    xarrcr = xarr

for j in range(4):
    ak = Lstype[j]
    go = gdata[ak]['AT']
    go.prepare_data(xarrcr,tarr,'r')
    
    for i,p in enumerate(tpairs):
        ti1 = np.where(tarr==p[0])[0][0]
        ti2 = np.where(tarr==p[1])[0][0]

        axr[i,j].fill_between(xarrcr,go.avg[:,ti1]+go.sem[:,ti1],go.avg[:,ti1]-go.sem[:,ti1],alpha=a,color='g',zorder=1)
        axr[i,j].fill_between(xarrcr,-go.avg[:,ti2]+go.sem[:,ti2],-go.avg[:,ti2]-go.sem[:,ti2],alpha=a,color='g',zorder=1)

        axr[i,j].plot(xarrcr,go.avg[:,ti1],c='g',zorder=2,label='AT')
        axr[i,j].plot(xarrcr,-go.avg[:,ti2],c='g',zorder=2)
        
    go = gdata[ak]['ZRF']
    go.prepare_data(xarrcr,tarr,'r')
    
    for i,p in enumerate(tpairs):
        ti1 = np.where(tarr==p[0])[0][0]
        ti2 = np.where(tarr==p[1])[0][0]

        axr[i,j].fill_between(xarrcr,go.avg[:,ti1]+go.sem[:,ti1],go.avg[:,ti1]-go.sem[:,ti1],alpha=a,color='r',zorder=1)
        axr[i,j].fill_between(xarrcr,-go.avg[:,ti2]+go.sem[:,ti2],-go.avg[:,ti2]-go.sem[:,ti2],alpha=a,color='r',zorder=1)

        axr[i,j].plot(xarrcr,go.avg[:,ti1],c='r',zorder=2,label='ZRF')
        axr[i,j].plot(xarrcr,-go.avg[:,ti2],c='r',zorder=2)
        
        if legend is not False:
            axr[i,j].legend()
            axr[i,j].set_yticklabels([])
            axr[i,j].set_xticklabels([])
        
        # Add theta labels
        axr[i,0].set_ylabel('top '+str(p[0])+'bottom '+str(p[1]))
        
    axr[0,j].set_title(ak+' '+str(go.arr.shape[-1]))
    
plt.tight_layout()

if legend is not False:
    plt.suptitle('AT/ZRF R distance for each sample type')
    
tstamp = datetime.datetime.now().strftime('%Y-%m-%d')

if save:
    fig.savefig(tstamp+'_s1a-indv-r.pdf')
```

##### Pts¶

In [33]:

```
fig,axr = plt.subplots(4,4,figsize=(16,10),sharey='col',sharex=True)

if crop is not None:
    mask = np.where((xarr>-crop)&(xarr<crop) == True)[0]
    xmin = mask.min()
    xmax = mask.max()
    xarrcr = xarr[xmin:xmax+1]
else:
    xarrcr = xarr

for j in range(4):
    ak = Lstype[j]
    go = gdata[ak]['AT']
    go.prepare_data(xarrcr,tarr,'pts')
    
    for i,p in enumerate(tpairs):
        ti1 = np.where(tarr==p[0])[0][0]
        ti2 = np.where(tarr==p[1])[0][0]

        axr[i,j].fill_between(xarrcr,go.avg[:,ti1]+go.sem[:,ti1],go.avg[:,ti1]-go.sem[:,ti1],alpha=a,color='g',zorder=1)
        axr[i,j].fill_between(xarrcr,-go.avg[:,ti2]+go.sem[:,ti2],-go.avg[:,ti2]-go.sem[:,ti2],alpha=a,color='g',zorder=1)

        axr[i,j].plot(xarrcr,go.avg[:,ti1],c='g',zorder=2,label='AT')
        axr[i,j].plot(xarrcr,-go.avg[:,ti2],c='g',zorder=2)
        
    go = gdata[ak]['ZRF']
    go.prepare_data(xarrcr,tarr,'pts')
    
    for i,p in enumerate(tpairs):
        ti1 = np.where(tarr==p[0])[0][0]
        ti2 = np.where(tarr==p[1])[0][0]

        axr[i,j].fill_between(xarrcr,go.avg[:,ti1]+go.sem[:,ti1],go.avg[:,ti1]-go.sem[:,ti1],alpha=a,color='r',zorder=1)
        axr[i,j].fill_between(xarrcr,-go.avg[:,ti2]+go.sem[:,ti2],-go.avg[:,ti2]-go.sem[:,ti2],alpha=a,color='r',zorder=1)

        axr[i,j].plot(xarrcr,go.avg[:,ti1],c='r',zorder=2,label='ZRF')
        axr[i,j].plot(xarrcr,-go.avg[:,ti2],c='r',zorder=2)
        
        if legend is not False:
            axr[i,j].legend()
            axr[i,j].set_yticklabels([])
            axr[i,j].set_xticklabels([])
        
        # Add theta labels
        axr[i,0].set_ylabel('top '+str(p[0])+'bottom '+str(p[1]))
        
    axr[0,j].set_title(ak+' '+str(go.arr.shape[-1]))
    
plt.tight_layout()

tstamp = datetime.datetime.now().strftime('%Y-%m-%d')

if save:
    fig.savefig(tstamp+'_s1a-indv-pts.pdf')
```

#### AT R comparison¶

In [34]:

```
channel = 'AT'
dtype = 'r'

fig,axr = plt.subplots(4,6,figsize=(24,10),sharey=True,sharex=True)

if crop is not None:
    mask = np.where((xarr>-crop)&(xarr<crop) == True)[0]
    xmin = mask.min()
    xmax = mask.max()
    xarrcr = xarr[xmin:xmax+1]
else:
    xarrcr = xarr

for n,c in enumerate(comp):
    go1 = gdata[c[0]][channel]
    go1.prepare_data(xarrcr,tarr,dtype)
    go2 = gdata[c[1]][channel]
    go2.prepare_data(xarrcr,tarr,dtype)
    
    parr = stats.ttest_ind(go1.arr_masked,go2.arr_masked,axis=2,nan_policy='omit')[1]
    
    parr[parr<pthresh] = 0
    parr[parr>pthresh] = 1
    
    go = go1
    for i,p in enumerate(tpairs):
        ti1 = np.where(tarr==p[0])[0][0]
        ti2 = np.where(tarr==p[1])[0][0]

        axr[i,n].fill_between(xarrcr,go.avg[:,ti1]+go.sem[:,ti1],go.avg[:,ti1]-go.sem[:,ti1],alpha=a,color=go.c,zorder=1)
        axr[i,n].fill_between(xarrcr,-go.avg[:,ti2]+go.sem[:,ti2],-go.avg[:,ti2]-go.sem[:,ti2],alpha=a,color=go.c,zorder=1)

        axr[i,n].plot(xarrcr,go.avg[:,ti1],c=go.c,zorder=2,label='{} {} {} {}'.format(go.arr.shape[-1],c[0],channel,dtype))
        axr[i,n].plot(xarrcr,-go.avg[:,ti2],c=go.c,zorder=2)
        
    go = go2
    for i,p in enumerate(tpairs):
        ti1 = np.where(tarr==p[0])[0][0]
        ti2 = np.where(tarr==p[1])[0][0]

        axr[i,n].fill_between(xarrcr,go.avg[:,ti1]+go.sem[:,ti1],go.avg[:,ti1]-go.sem[:,ti1],alpha=a,color=go.c,zorder=1)
        axr[i,n].fill_between(xarrcr,-go.avg[:,ti2]+go.sem[:,ti2],-go.avg[:,ti2]-go.sem[:,ti2],alpha=a,color=go.c,zorder=1)

        axr[i,n].plot(xarrcr,go.avg[:,ti1],c=go.c,zorder=2,label='{} {} {} {}'.format(go.arr.shape[-1],c[1],channel,dtype))
        axr[i,n].plot(xarrcr,-go.avg[:,ti2],c=go.c,zorder=2)
        
        axr[i,n].scatter(xarrcr,go.avg[:,ti1],c=parr[:,ti1],cmap='Greys_r',zorder=3,vmin=0,vmax=1,edgecolor='k')
        axr[i,n].scatter(xarrcr,-go.avg[:,ti2],c=parr[:,ti2],cmap='Greys_r',zorder=3,vmin=0,vmax=1,edgecolor='k')
        
        axr[0,n].legend(bbox_to_anchor=(0., 1.02, 1., .102), loc=3,ncol=2, mode="expand", borderaxespad=0.)

        if legend is not False:
            axr[i,j].legend()
            axr[i,j].set_yticklabels([])
            axr[i,j].set_xticklabels([])
        
        # Add theta labels
        axr[i,0].set_ylabel('top '+str(p[0])+'bottom '+str(p[1]))
    
plt.tight_layout()

tstamp = datetime.datetime.now().strftime('%Y-%m-%d')

if save:
    fig.savefig(tstamp+'_s1a-comparison-{}-{}.pdf'.format(channel,dtype))
```

```
C:\Users\zfishlab\AppData\Local\Continuum\anaconda3\envs\deltascope\lib\site-packages\ipykernel_launcher.py:22: RuntimeWarning: invalid value encountered in less
C:\Users\zfishlab\AppData\Local\Continuum\anaconda3\envs\deltascope\lib\site-packages\ipykernel_launcher.py:23: RuntimeWarning: invalid value encountered in greater
```

#### AT Pts comparison¶

In [35]:

```
channel = 'AT'
dtype = 'pts'

fig,axr = plt.subplots(4,6,figsize=(25,10),sharey='col',sharex=True)

if crop is not None:
    mask = np.where((xarr>-crop)&(xarr<crop) == True)[0]
    xmin = mask.min()
    xmax = mask.max()
    xarrcr = xarr[xmin:xmax+1]
else:
    xarrcr = xarr

for n,c in enumerate(comp):
    go1 = gdata[c[0]][channel]
    go1.prepare_data(xarrcr,tarr,dtype)
    go2 = gdata[c[1]][channel]
    go2.prepare_data(xarrcr,tarr,dtype)
    
    parr = stats.ttest_ind(go1.arr_masked,go2.arr_masked,axis=2,nan_policy='omit')[1]
    
    parr[parr<pthresh] = 0
    parr[parr>pthresh] = 1
    
    go = go1
    for i,p in enumerate(tpairs):
        ti1 = np.where(tarr==p[0])[0][0]
        ti2 = np.where(tarr==p[1])[0][0]

        axr[i,n].fill_between(xarrcr,go.avg[:,ti1]+go.sem[:,ti1],go.avg[:,ti1]-go.sem[:,ti1],alpha=a,color=go.c,zorder=1)
        axr[i,n].fill_between(xarrcr,-go.avg[:,ti2]+go.sem[:,ti2],-go.avg[:,ti2]-go.sem[:,ti2],alpha=a,color=go.c,zorder=1)

        axr[i,n].plot(xarrcr,go.avg[:,ti1],c=go.c,zorder=2,label='{} {} {} {}'.format(go.arr.shape[-1],c[0],channel,dtype))
        axr[i,n].plot(xarrcr,-go.avg[:,ti2],c=go.c,zorder=2)
        
    go = go2
    for i,p in enumerate(tpairs):
        ti1 = np.where(tarr==p[0])[0][0]
        ti2 = np.where(tarr==p[1])[0][0]

        axr[i,n].fill_between(xarrcr,go.avg[:,ti1]+go.sem[:,ti1],go.avg[:,ti1]-go.sem[:,ti1],alpha=a,color=go.c,zorder=1)
        axr[i,n].fill_between(xarrcr,-go.avg[:,ti2]+go.sem[:,ti2],-go.avg[:,ti2]-go.sem[:,ti2],alpha=a,color=go.c,zorder=1)

        axr[i,n].plot(xarrcr,go.avg[:,ti1],c=go.c,zorder=2,label='{} {} {} {}'.format(go.arr.shape[-1],c[1],channel,dtype))
        axr[i,n].plot(xarrcr,-go.avg[:,ti2],c=go.c,zorder=2)
        
        axr[i,n].scatter(xarrcr,go.avg[:,ti1],c=parr[:,ti1],cmap='Greys_r',zorder=3,vmin=0,vmax=1,edgecolor='k')
        axr[i,n].scatter(xarrcr,-go.avg[:,ti2],c=parr[:,ti2],cmap='Greys_r',zorder=3,vmin=0,vmax=1,edgecolor='k')
        
        axr[0,n].legend(bbox_to_anchor=(0., 1.02, 1., .102), loc=3,ncol=2, mode="expand", borderaxespad=0.)

        if legend is not False:
            axr[i,j].legend()
            axr[i,j].set_yticklabels([])
            axr[i,j].set_xticklabels([])
        
        # Add theta labels
        axr[i,0].set_ylabel('top '+str(p[0])+'bottom '+str(p[1]))
    
plt.tight_layout()

tstamp = datetime.datetime.now().strftime('%Y-%m-%d')

if save:
    fig.savefig(tstamp+'_s1a-comparison-{}-{}.pdf'.format(channel,dtype))
```

```
C:\Users\zfishlab\AppData\Local\Continuum\anaconda3\envs\deltascope\lib\site-packages\ipykernel_launcher.py:22: RuntimeWarning: invalid value encountered in less
C:\Users\zfishlab\AppData\Local\Continuum\anaconda3\envs\deltascope\lib\site-packages\ipykernel_launcher.py:23: RuntimeWarning: invalid value encountered in greater
```

#### Comparison zrf R¶

In [36]:

```
channel = 'ZRF'
dtype = 'r'

fig,axr = plt.subplots(4,6,figsize=(26,10),sharey=True,sharex=True)

if crop is not None:
    mask = np.where((xarr>-crop)&(xarr<crop) == True)[0]
    xmin = mask.min()
    xmax = mask.max()
    xarrcr = xarr[xmin:xmax+1]
else:
    xarrcr = xarr

for n,c in enumerate(comp):
    go1 = gdata[c[0]][channel]
    go1.prepare_data(xarrcr,tarr,dtype)
    go2 = gdata[c[1]][channel]
    go2.prepare_data(xarrcr,tarr,dtype)
    
    parr = stats.ttest_ind(go1.arr_masked,go2.arr_masked,axis=2,nan_policy='omit')[1]
    
    parr[parr<pthresh] = 0
    parr[parr>pthresh] = 1
    
    go = go1
    for i,p in enumerate(tpairs):
        ti1 = np.where(tarr==p[0])[0][0]
        ti2 = np.where(tarr==p[1])[0][0]

        axr[i,n].fill_between(xarrcr,go.avg[:,ti1]+go.sem[:,ti1],go.avg[:,ti1]-go.sem[:,ti1],alpha=a,color=go.c,zorder=1)
        axr[i,n].fill_between(xarrcr,-go.avg[:,ti2]+go.sem[:,ti2],-go.avg[:,ti2]-go.sem[:,ti2],alpha=a,color=go.c,zorder=1)

        axr[i,n].plot(xarrcr,go.avg[:,ti1],c=go.c,zorder=2,label='{} {} {} {}'.format(go.arr.shape[-1],c[0],channel,dtype))
        axr[i,n].plot(xarrcr,-go.avg[:,ti2],c=go.c,zorder=2)
        
    go = go2
    for i,p in enumerate(tpairs):
        ti1 = np.where(tarr==p[0])[0][0]
        ti2 = np.where(tarr==p[1])[0][0]

        axr[i,n].fill_between(xarrcr,go.avg[:,ti1]+go.sem[:,ti1],go.avg[:,ti1]-go.sem[:,ti1],alpha=a,color=go.c,zorder=1)
        axr[i,n].fill_between(xarrcr,-go.avg[:,ti2]+go.sem[:,ti2],-go.avg[:,ti2]-go.sem[:,ti2],alpha=a,color=go.c,zorder=1)

        axr[i,n].plot(xarrcr,go.avg[:,ti1],c=go.c,zorder=2,label='{} {} {} {}'.format(go.arr.shape[-1],c[1],channel,dtype))
        axr[i,n].plot(xarrcr,-go.avg[:,ti2],c=go.c,zorder=2)
        
        axr[i,n].scatter(xarrcr,go.avg[:,ti1],c=parr[:,ti1],cmap='Greys_r',zorder=3,vmin=0,vmax=1,edgecolor='k')
        axr[i,n].scatter(xarrcr,-go.avg[:,ti2],c=parr[:,ti2],cmap='Greys_r',zorder=3,vmin=0,vmax=1,edgecolor='k')
        
        axr[0,n].legend(bbox_to_anchor=(0., 1.02, 1., .102), loc=3,ncol=2, mode="expand", borderaxespad=0.)
        
        if legend is not False:
            axr[i,j].legend()
            axr[i,j].set_yticklabels([])
            axr[i,j].set_xticklabels([])
        
        # Add theta labels
        axr[i,0].set_ylabel('top '+str(p[0])+'bottom '+str(p[1]))
    
plt.tight_layout()

tstamp = datetime.datetime.now().strftime('%Y-%m-%d')

if save:
    fig.savefig(tstamp+'_s1a-comparison-{}-{}.pdf'.format(channel,dtype))
```

```
C:\Users\zfishlab\AppData\Local\Continuum\anaconda3\envs\deltascope\lib\site-packages\ipykernel_launcher.py:22: RuntimeWarning: invalid value encountered in less
C:\Users\zfishlab\AppData\Local\Continuum\anaconda3\envs\deltascope\lib\site-packages\ipykernel_launcher.py:23: RuntimeWarning: invalid value encountered in greater
```

#### comparison zrf pts¶

In [37]:

```
channel = 'ZRF'
dtype = 'pts'

fig,axr = plt.subplots(4,6,figsize=(28,11),sharey='col',sharex=True)

if crop is not None:
    mask = np.where((xarr>-crop)&(xarr<crop) == True)[0]
    xmin = mask.min()
    xmax = mask.max()
    xarrcr = xarr[xmin:xmax+1]
else:
    xarrcr = xarr

for n,c in enumerate(comp):
    go1 = gdata[c[0]][channel]
    go1.prepare_data(xarrcr,tarr,dtype)
    go2 = gdata[c[1]][channel]
    go2.prepare_data(xarrcr,tarr,dtype)
    
    parr = stats.ttest_ind(go1.arr_masked,go2.arr_masked,axis=2,nan_policy='omit')[1]
    
    parr[parr<pthresh] = 0
    parr[parr>pthresh] = 1
    
    go = go1
    for i,p in enumerate(tpairs):
        ti1 = np.where(tarr==p[0])[0][0]
        ti2 = np.where(tarr==p[1])[0][0]

        axr[i,n].fill_between(xarrcr,go.avg[:,ti1]+go.sem[:,ti1],go.avg[:,ti1]-go.sem[:,ti1],alpha=a,color=go.c,zorder=1)
        axr[i,n].fill_between(xarrcr,-go.avg[:,ti2]+go.sem[:,ti2],-go.avg[:,ti2]-go.sem[:,ti2],alpha=a,color=go.c,zorder=1)

        axr[i,n].plot(xarrcr,go.avg[:,ti1],c=go.c,zorder=2,label='{} {} {} {}'.format(go.arr.shape[-1],c[0],channel,dtype))
        axr[i,n].plot(xarrcr,-go.avg[:,ti2],c=go.c,zorder=2)
        
    go = go2
    for i,p in enumerate(tpairs):
        ti1 = np.where(tarr==p[0])[0][0]
        ti2 = np.where(tarr==p[1])[0][0]

        axr[i,n].fill_between(xarrcr,go.avg[:,ti1]+go.sem[:,ti1],go.avg[:,ti1]-go.sem[:,ti1],alpha=a,color=go.c,zorder=1)
        axr[i,n].fill_between(xarrcr,-go.avg[:,ti2]+go.sem[:,ti2],-go.avg[:,ti2]-go.sem[:,ti2],alpha=a,color=go.c,zorder=1)

        axr[i,n].plot(xarrcr,go.avg[:,ti1],c=go.c,zorder=2,label='{} {} {} {}'.format(go.arr.shape[-1],c[1],channel,dtype))
        axr[i,n].plot(xarrcr,-go.avg[:,ti2],c=go.c,zorder=2)
        
        axr[i,n].scatter(xarrcr,go.avg[:,ti1],c=parr[:,ti1],cmap='Greys_r',zorder=3,vmin=0,vmax=1,edgecolor='k')
        axr[i,n].scatter(xarrcr,-go.avg[:,ti2],c=parr[:,ti2],cmap='Greys_r',zorder=3,vmin=0,vmax=1,edgecolor='k')
        
        axr[0,n].legend(bbox_to_anchor=(0., 1.02, 1., .102), loc=3,ncol=2, mode="expand", borderaxespad=0.)

        if legend is not False:
            axr[i,j].legend()
            axr[i,j].set_yticklabels([])
            axr[i,j].set_xticklabels([])
        
        # Add theta labels
        axr[i,0].set_ylabel('top '+str(p[0])+'bottom '+str(p[1]))
    
plt.tight_layout()

tstamp = datetime.datetime.now().strftime('%Y-%m-%d')

if save:
    fig.savefig(tstamp+'_s1a-comparison-{}-{}.pdf'.format(channel,dtype))
```

```
C:\Users\zfishlab\AppData\Local\Continuum\anaconda3\envs\deltascope\lib\site-packages\ipykernel_launcher.py:22: RuntimeWarning: invalid value encountered in less
C:\Users\zfishlab\AppData\Local\Continuum\anaconda3\envs\deltascope\lib\site-packages\ipykernel_launcher.py:23: RuntimeWarning: invalid value encountered in greater
```
