## Supplementary material for "ΔSCOPE: A new method to quantify 3D biological structures and identify differences in zebrafish forebrain development": An archived copy of the DeltaSCOPE code repository.: TEMP-alignment.ipynb

````
# %% [markdown]
# Introduction
This notebook facilitates the manual curation of sample alignment.

# %% [code]
import deltascope.alignment as ut

import numpy as np
import pandas as pd
import matplotlib.pyplot as plt
import h5py

import os
import re
import time
import tqdm

# %% [markdown]
# Setup

# %% [markdown]
## Parameters

# %% [code]
# --------------------------------
# -------- User input ------------
# --------------------------------

param = {
    'gthresh':0.5,
    'scale':[1,1,1],
    'microns':[0.16,0.16,0.21],
    'mthresh':0.5,
    'radius':10,
    'comp_order':[0,2,1],
    'fit_dim':['x','z'],
    'deg':2,
    
    # Don't forget to modify this with a specific sample name
    'expname':'expname'
}

# %% [markdown]
## Directories

# %% [code]
# --------------------------------
# -------- User input ------------
# --------------------------------

# Specify file paths to directories containing probability files
# after processing by ilastik
gfap = os.path.abspath('..\expname\GFAP\Prob')
at = os.path.abspath('..\expname\AT\Prob')

# Specify root directory where output will be saved
root = os.path.abspath('..\expname')

# %% [code]
# Output directory with timestamp
outname = 'Output_'+time.strftime("%m-%d-%H-%M",
                                 time.localtime())

# Create output directory
outdir = os.path.join(root,outname)
os.mkdir(outdir)

# %% [markdown]
## Extract list of files

# %% [code]
Dat = {}
for f in os.listdir(at):
    if 'h5' in f:
        num  = re.findall(r'\d+',f.split('.')[0])[-1]
        Dat[num] = os.path.join(at,f)

# %% [code]
Dzrf = {}
for f in os.listdir(gfap):
    if 'h5' in f:
        num  = re.findall(r'\d+',f.split('.')[0])[-1]
        Dzrf[num] = os.path.join(gfap,f)

# %% [code]
# Extract list of filename keys
klist = Dat.keys()

# %% [code]
# Create dictionaries to contain the deltascope brain object for each sample
Dbat = {}
Dbzrf = {}

# %% [markdown]
# Import raw data and perform preprocessing

# %% [code]
%%time
for k in tqdm.tqdm(klist):
    if k not in list(Dbat.keys()):
        Dbat[k] = ut.preprocess(Dat[k],param)
        Dbzrf[k] = ut.preprocess(Dzrf[k],param,pca=Dbat[k].pcamed,
                                 mm=Dbat[k].mm,vertex=Dbat[k].vertex)
    else:
        print(k,'already processed')

# %% [markdown]
# Example alignment

The `start` function initializes a sample for alignment. It displays a plot of the data with three projections (XY, XZ, and ZY) that should give a general sense of sample alignment. Note, that the `start` function attempts to align the sample using PCA. In some really bad cases, it is better to work on a sample without PCA alignment, in which case we can use the following code to initialize:
```python
k = klist[0]
df,Ldf = get_dfs(k)
ax = ut.make_graph([df]+Ldf)
```

In most cases, the following approach will be appropriate.

```python
k,df,Ldf,ax = start(klist[0])
```

`k`: The dictionary key that identifies this sample. It should be the sample number extracted from the filename.  
`df`: The dataframe containing datapoints associated with the primary alignment channel, in this case, AT.  
`Ldf`: A list of additional dataframes corresponding to other channels in the collection. In this template, we are assuming only one additional channel, ZRF.  
`ax`: The array containing subplots for this sample. There should be three projections for each channel and a row per channel. In this example the dimensions of the array are 2x3.

## Correction options

![alignment-correction](alignment-correction.jpg)

### A: Rotation around the X axis
This rotation error can best be identified in the YZ projection where the line of the sample does not fall on either axis. In order to correct this error, we will select two points in YZ that will be used to calculate a line that fits the sample. We can then use this line to calculate the angle of rotation needed to align the sample with the Z axis.

To perform this rotation, we will use the `ut.check_yz` function, which will fit a line in the YZ plane to use for rotation. This function takes only `df` and `Ldf` as required inputs.

```python
df1,Ldf1,ax,p = ut.check_yz(df,Ldf)
```

This function returns updated verions of `df` and `Ldf` following the rotation. I typically define new variables `df1` and `Ldf1` to keep track of the changes. It also returns a plotting object `ax` that will display a before and after comparison of the alignment with the points used for alignment plotted on the before plot for reference. Finally, it returns the `np.poly1d` object which contains the line that was used for alignment.

If the results of this alignment are good, we can proceed with `df1` and `Ldf1`. Alternatively, we can try to manually assign an improved line and pass the resulting `np.poly1d` object to `ut.check_yz` as an optional arguement `ut.check_yz(df,Ldf,mm=np.poly1d)`.

### B: Rotation around the Y axis
This error can be seen in the XZ projection where the parabola of the sample is tilted towards one side or the other. In order to correct this error, we will select two points that mark the endpoints of the parabola. The line between these two points will be used to calculate the angle of rotation to correct the parabola.  

To perform this rotation, we will use the `check_pts` function, which will perform a rotation either in the XY or XZ plane. It requires three parameters: `df`, `Ldf`, and the second dimension of the plane of interest (`'y'` or `'z'`).

```python
# Attempt rotation based on alignment points calculated from the data
df1,Ldf1,pts,ax = ut.check_pts(df,Ldf,'z')
```

In addition to typical outputs, this function returns `pts`, which is a pandas dataframe specifying two points in the XZ plane that were used for alignment.

If we are unhappy with the results of the alignment, we can manually adjust the anchor points and then recalculate the rotation.

```python
# Assign new values to pts
pts.iloc[0].x = 10
pts.iloc[1].z = 50

# Replot the updated pts to check choices
ax[0,1].scatter(pts.x,pts.z,c='y')
```

If these `pts` look good, then we can use `ut.revise_pts` to recalculate the rotation.

```python
df2,Ldf2,ax = ut.revise_pts(df,Ldf,'z',pts=pts)
```

If we are happy with these results, we could use `df2` and `Ldf2` to save as our final result.

### C: Mismatched Y and Z axes
Here the parabola appears in the XY projection when we expect it in the XZ projection. We can correct this by simply switching the Y and Z axes. The `ut.zyswitch` function facilitates this process.

```python
df1,Ldf1 = zyswitch(df,Ldf)
# Plot data to check correction result
ax = ut.make_graph(df1,Ldf1)
```

### D: Upside down parabola
We expect the parabola to lie in the positive half of the Z dimension. To correct an upside down parabola, we rotate the sample by 180$^\circ$ around X axis. Here, we will use the `ut.flip` function.

```python
df1,Ldf1 = ut.flip(df,Ldf)
# Plot data to check correction result
ax = ut.make_graph(df1,Ldf1)
```

## Correct vertex 
After performing the corrections described in the previous section, the vertex of the parabola may no longer be positioned at the origin. The function `ut.ch_vertex` attempts to reposition the vertex to the origin. This function also returns the math model `mm`, which describes the parabola of the data. We will save `mm` for future reference.

For this example, we will assume that we are happy with the alignment in `df1` and `Ldf1`.

```python
df2,Ldf2,mm,ax = ut.ch_vertex(df1,Ldf1)
```

If we disagree with the assignment of the vertex, we can manually pick three points that will be used to recalculate the vertex. These points should mark the two ends of the parabola (x1,z1) and (x2,z2) as well as the approximate vertex (vx,vz). We can specify and check these points before using them to shift the sample.

```python
pts = pick_pts(-78,12,-36,0,0,10) #these are essentially random numbers for example
ax[0,1].scatter(pts.x,pts.z,c='m',s=50)
```

Finally if we like these points, we can run `ut.ch_vertex` again with these specific points.

```python
df3,Ldf3,mm,ax = ut.ch_vertex(df2,Ldf2,pts=pts)
```

In order to complete the alignment process, we need to add the math model `mm` to dataframe for future reference and save the aligned data.

```python
model = save_model(k,mm,model)
save_both(k,df3,Ldf3[0])
```

### Mack's Notes on Minimum one must do to each sample
```python
k,df,Ldf,ax = start(klist[0])
df,Ldf,mm,ax = ut.ch_vertex(df,Ldf)
model = save_model(k,mm,model)
save_both(k,df,Ldf[0])
```


# %% [markdown]
# Define experiment specific functions

# %% [code]
''' Define wrapper functions for starting and saving to minimize the number 
of inputs that the user needs to type for each call of the function.'''
def start(k):
    return(ut.start(k,Dbat,[Dbzrf],im=True))
def save_both(k,dfa,dfb):
    ut.save_both(k,dfa,dfb,outdir,param.expname)

'''Save model parameters for each file to a dataframe that can be 
exported for later reference.'''
model = pd.DataFrame({'a':[],'b':[],'c':[]})
def save_model(k,mm,model):
    row = pd.Series({'a':mm[0],'b':mm[1],'c':mm[2]},name=k)
    model = model.append(row)
    return(model)

'''Define a function that can both fit a model and plot it on an existing plot'''
def fit_model(axi,df,mm=None):
    if mm == None:
        mm = np.polyfit(df.x,df.z,2)
    p = np.poly1d(mm)
    xrange = np.arange(np.min(df.x),np.max(df.x))
    axi.plot(xrange,p(xrange),c='m')
    return(mm)

'''Take a set of points and transform to a dataframe format for ease of access.'''
def pick_pts(x1,z1,vx,vz,x2,z2):
    pts = pd.DataFrame({'x':[x1,vx,x2],'z':[z1,vz,z2]})
    return(pts)

# %% [markdown]
# Process remaining samples
The workflow presented in the section above needs to be applied to all samples in `klist`. It is up to the user to decide which corrections are appropriate for each individual sample. I recommend that this review is also used as an opportunity to exclude samples with issues in the raw data, such as tears in the commissure or limited signal. The reason for rejecting a sample can be recorded in this notebook for future reference.

# %% [code]
klist

# %% [markdown]
# Wrapping up: after all samples are processed
Once all of the data has been processed, we want to save the model data we collected to a file for future reference.

# %% [code]
model.to_csv(os.path.join(outdir,'model.csv'))

# %% [markdown]
Additionally, it can be helpful to export a html or pdf version of the notebook that preserves all of the plots generated during the alignment process. To do so, use the Jupyter Lab menu interface: File > Export Notebook As > HTML or PDF.
````
