## Supplementary material for "ΔSCOPE: A new method to quantify 3D biological structures and identify differences in zebrafish forebrain development": An archived copy of the DeltaSCOPE code repository.: TEMP-landmarks.ipynb

```
# %% [markdown]
# Introduction: Landmarks

# %% [code]
import deltascope as ds
import deltascope.alignment as ut

import numpy as np
import pandas as pd
import matplotlib.pyplot as plt

from sklearn.preprocessing import normalize
from scipy.optimize import minimize

import os
import tqdm
import json
import time

# %% [markdown]
# Import raw data
The user needs to specify the directories containing the data of interest. Each sample type should have a key which corresponds to the directory path. Additionally, each object should have a list that includes the channels of interest.

# %% [code]
# --------------------------------
# -------- User input ------------
# --------------------------------

data = {
    # Specify sample type key
    'wt': {
        # Specify path to data directory
        'path': 'path\to\data\directory\sample1',
        # Specify which channels are in the directory and are of interest
        'channels': ['AT','ZRF']
    },
    'stype2': {
        'path': 'path\to\data\directory\sample2',
        'channels': ['AT','ZRF']
    }
}

# %% [markdown]
We'll generate a list of pairs of stypes and channels for ease of use.

# %% [code]
data_pairs = []
for s in data.keys():
    for c in data[s]['channels']:
        data_pairs.append((s,c))

# %% [markdown]
We can now read in all datafiles specified by the `data` dictionary above.

# %% [code]
D = {}
for s in data.keys():
    D[s] = {}
    for c in data[s]['channels']:
        D[s][c] = ds.read_psi_to_dict(data[s]['path'],c)

# %% [markdown]
# Calculate landmark bins

# %% [code]
# --------------------------------
# -------- User input ------------
# --------------------------------

# Pick an integer value for bin number based on results above
anum = 25

# Specify the percentiles which will be used to calculate landmarks
percbins = [50]

# %% [markdown]
Calculate landmark bins based on user input parameters and the previously specified control sample.

# %% [code]
lm = ds.landmarks(percbins=percbins, rnull=np.nan)
lm.calc_bins(D[s_ctrl][c_ctrl], anum, theta_step)

print('Alpha bins')
print(lm.acbins)
print('Theta bins')
print(lm.tbins)

# %% [markdown]
# Calculate landmarks

# %% [code]
lmdf = pd.DataFrame()

# Loop through each pair of stype and channels
for s,c in tqdm.tqdm(data_pairs):
    print(s,c)
    # Calculate landmarks for each sample with this data pair
    for k,df in tqdm.tqdm(D[s][c].items()):
        lmdf = lm.calc_perc(df, k, '-'.join([s,c]), lmdf)
        
# Set timestamp for saving data
tstamp = time.strftime("%m-%d-%H-%M",time.localtime())
        
# Save completed landmarks to a csv file
lmdf.to_csv(tstamp+'_landmarks.csv')

# Save landmark bins to json file
bins = {
    'acbins':list(lm.acbins),
    'tbins':list(lm.tbins)
}
with open(tstamp+'_landmarks_bins.json', 'w') as outfile:
    json.dump(bins, outfile)
```
