## Supplementary material for "ΔSCOPE: A new method to quantify 3D biological structures and identify differences in zebrafish forebrain development": An archived copy of the DeltaSCOPE code repository.: TEMP-transformation.py

```
import deltascope as ds
import multiprocessing as mp
import time
import os
from functools import partial
from sys import argv
import re

# Define parameters or config file
# An input from a file will override dictionary input from dparams

# Specify path if parameters are in json file
config_path = None # or transformation_config.json

# Fill out dictionary to specify parameters here
dparams = {
			"rootdir": "",
			"expname": "",
			"c1-dir": "",
			"c1-key": "",
			"c2-dir": "",
			"c2-key": "",
			"c3-dir": "",
			"c3-key": "",
			"c4-dir": "",
			"c4-key": "",
			"genthresh": None,
			"microns": [None,None,None],
			"scale": [1,1,1],
			"medthresh": None,
			"radius": None,
			"comporder": [None,None,None],
			"fitdim": ["",""],
			"deg": None,
			"twoD": None
		}

# -----------------------------------------
# No modifications needed below this line
# -----------------------------------------

def check_nums(P):
	'''
	Check that the numbers of files selected by the same list index match
	Raises an error if file numbers are mismatched

	:param :class:`paramClass` P: Object containing all variables from config file
	'''
	Lnums = []

	for i,f in enumerate(P.c1_files):
		n = re.findall(r'\d+',f.split('.')[0])[0]
		for Lf in P.Lcfiles:
			if n not in Lf[i]:
				print('File numbers are mismatched between channel directories.')
				raise
		Lnums.append(i)

	return(Lnums)

def process(num,P=None):
	'''
	Run through the processing steps for a single sample through saving psi files

	:param int num: Index of the file that is currently being processed
	:param :class:`paramClass` P: Object containing all variables from config file
	'''

	tic = time.time()
	print(num,'Starting sample')

	#Extract sample number from c1 filename
	snum = re.findall(r'\d+',P.c1_files[num].split('.')[0])[0]

	e = ds.embryo(P.expname,snum,P.outdir)

	#Add channels and preprocess data
	try:
		e.add_channel(os.path.join(P.c1_dir,P.c1_files[num]),P.c1_key)
		e.chnls[P.c1_key].preprocess_data(P.genthresh,P.scale,P.microns)
		for i in range(len(P.Lcdir)):
			e.add_channel(os.path.join(P.Lcdir[i],P.Lcfiles[i][num]),P.Lckey[i])
			e.chnls[P.Lckey[i]].preprocess_data(P.genthresh,P.scale,P.microns)
	except:
		print(num,'failed on preprocess_data',time.time()-tic)

	#Calculate PCA transformation for structural channel, c1
	try:
		if P.twoD == True:
			e.chnls[P.c1_key].calculate_pca_median_2d(e.chnls[P.c1_key].raw_data,P.medthresh,P.radius,P.microns)
			pca = e.chnls[P.c1_key].pcamed
			e.chnls[P.c1_key].pca_transform_2d(e.chnls[P.c1_key].df_thresh,pca,P.comporder,P.fitdim,deg=P.deg)

			#Transform additional channels
			for i in range(len(P.Lcdir)):
				e.chnls[P.Lckey[i]].pca_transform_2d(e.chnls[P.Lckey[i]].df_thresh,pca,P.comporder,P.fitdim,deg=P.deg)

		else:
			e.chnls[P.c1_key].calculate_pca_median(e.chnls[P.c1_key].raw_data,P.medthresh,P.radius,P.microns)
			pca = e.chnls[P.c1_key].pcamed
			e.chnls[P.c1_key].pca_transform_3d(e.chnls[P.c1_key].df_thresh,pca,P.comporder,P.fitdim,deg=P.deg)

			#Transform additional channels
			for i in range(len(P.Lcdir)):
				e.chnls[P.Lckey[i]].pca_transform_3d(e.chnls[P.Lckey[i]].df_thresh,pca,P.comporder,P.fitdim,deg=P.deg)
	except:
		print(num,'failed on pca',time.time()-tic)

	print(num,'Starting coordinate transformation')
	try:
		e.chnls[P.c1_key].transform_coordinates()
		for i in range(len(P.Lcdir)):
			e.chnls[P.Lckey[i]].transform_coordinates()
	except:
		print(num,'failed on coordinate transform',time.time()-tic)

	try:
		e.save_psi()
	except:
		print(num,'failed on save_psi',time.time()-tic)

	toc = time.time()
	print(num,'Complete',toc-tic)

if __name__=='__main__':

	f,config_path = argv
	P = paramsClass(path=config_path, 
					dparams=dparams)

	#Create out directory stamped with current date and time
	outdir = os.path.join(P.rootdir,'Output'+time.strftime("%m-%d-%H-%M",time.localtime()))
	os.mkdir(outdir)
	P.add_outdir(outdir)
	print('outdir',outdir)

	processfxn = partial(process,P=P)

	Lnums = check_nums(P)
	n = len(Lnums)
	# Initiate map pools in sets of 5 samples
	for i in range(0,n,5):
		if i+5>n:
			L = Lnums[i:n]
		else:
			L = Lnums[i:i+5]

		pool = mp.Pool()
		pool.map(processfxn,L)
		pool.close()
		pool.join()
```
