## Supplementary material for "ΔSCOPE: A new method to quantify 3D biological structures and identify differences in zebrafish forebrain development": An archived copy of the DeltaSCOPE code repository.: alignment-wt.html


### Introduction¶

This notebook facilitates the manual curation of sample alignment.

In [1]:

```
import deltascope.alignment as ut

import numpy as np
import pandas as pd
import matplotlib.pyplot as plt
import h5py

import os
import re
import time
import tqdm
```

### Setup¶

#### Parameters¶

In [2]:

```
# --------------------------------
# -------- User input ------------
# --------------------------------

param = {
    'gthresh':0.5,
    'scale':[1,1,1],
    'microns':[0.16,0.16,0.21],
    'mthresh':0.5,
    'radius':20,
    'comp_order':[0,2,1],
    'fit_dim':['x','z'],
    'deg':2,
    
    # Don't forget to modify this with a specific sample name
    'expname':'wt'
}
```

#### Directories¶

In [3]:

```
# --------------------------------
# -------- User input ------------
# --------------------------------

# Specify file paths to directories containing probability files
# after processing by ilastik
gfap = '.\data\ZRF1 Wild Type HZ H5 files\Prob_7-29'
at = '.\data\AT Wild Type HZ H5 files\Prob'

# Specify root directory where output will be saved
root = '.\data'

root
```

Out[3]:

```
'.\\data'
```

In [4]:

```
# Output directory with timestamp
outname = 'Output_'+param['expname']+time.strftime("%m-%d-%H-%M",
                                 time.localtime())

# Create output directory
outdir = os.path.join(root,outname)
os.mkdir(outdir)
```

#### Extract list of files¶

In [12]:

```
Dat = {}
for f in os.listdir(at):
    if 'h5' in f:
        num  = re.findall(r'\d+',f.split('.')[0])[-1]
        Dat[num] = os.path.join(at,f)
```

In [13]:

```
Dzrf = {}
for f in os.listdir(gfap):
    if 'h5' in f:
        num  = re.findall(r'\d+',f.split('.')[0])[-1]
        Dzrf[num] = os.path.join(gfap,f)
```

In [14]:

```
# Create dictionaries to contain the deltascope brain object for each sample
Dbat = {}
Dbzrf = {}
```

In [15]:

```
klist = Dzrf.keys()
klist
```

Out[15]:

```
dict_keys(['101', '102', '103', '104', '105', '106', '107', '108', '109', '110', '111', '112', '113', '114', '115', '116', '117', '118', '119', '120', '121', '122', '123', '124', '125', '126', '127', '128', '129', '130', '131', '132', '133', '134', '135', '136', '137', '138', '139', '140', '141', '142', '143'])
```

### PCA: Import raw data and perform preprocessing¶

In [16]:

```
%%time
for k in tqdm.tqdm(klist):
    if k not in list(Dbat.keys()):
        try:
            Dbat[k] = ut.preprocess(Dat[k],param)
            Dbzrf[k] = ut.preprocess(Dzrf[k],param,pca=Dbat[k].pcamed,
                                 mm=Dbat[k].mm,vertex=Dbat[k].vertex)
        except:
            print(k,'caused an error')
    else:
        print(k,'already processed')
```

```
  0%|          | 0/43 [00:00<?, ?it/s]C:\Users\zfishlab\AppData\Local\Continuum\anaconda3\envs\test\lib\site-packages\skimage\util\dtype.py:141: UserWarning: Possible precision loss when converting from float32 to uint8
  .format(dtypeobj_in, dtypeobj_out))
  2%|▏         | 1/43 [01:37<1:08:05, 97.28s/it]C:\Users\zfishlab\AppData\Local\Continuum\anaconda3\envs\test\lib\site-packages\skimage\util\dtype.py:141: UserWarning: Possible precision loss when converting from float32 to uint8
  .format(dtypeobj_in, dtypeobj_out))
  5%|▍         | 2/43 [03:10<1:05:43, 96.18s/it]C:\Users\zfishlab\AppData\Local\Continuum\anaconda3\envs\test\lib\site-packages\skimage\util\dtype.py:141: UserWarning: Possible precision loss when converting from float32 to uint8
  .format(dtypeobj_in, dtypeobj_out))
  7%|▋         | 3/43 [04:53<1:05:21, 98.03s/it]C:\Users\zfishlab\AppData\Local\Continuum\anaconda3\envs\test\lib\site-packages\skimage\util\dtype.py:141: UserWarning: Possible precision loss when converting from float32 to uint8
  .format(dtypeobj_in, dtypeobj_out))
  9%|▉         | 4/43 [06:02<58:02, 89.29s/it]  C:\Users\zfishlab\AppData\Local\Continuum\anaconda3\envs\test\lib\site-packages\skimage\util\dtype.py:141: UserWarning: Possible precision loss when converting from float32 to uint8
  .format(dtypeobj_in, dtypeobj_out))
 12%|█▏        | 5/43 [07:44<59:05, 93.29s/it]C:\Users\zfishlab\AppData\Local\Continuum\anaconda3\envs\test\lib\site-packages\skimage\util\dtype.py:141: UserWarning: Possible precision loss when converting from float32 to uint8
  .format(dtypeobj_in, dtypeobj_out))
 14%|█▍        | 6/43 [08:55<53:24, 86.60s/it]C:\Users\zfishlab\AppData\Local\Continuum\anaconda3\envs\test\lib\site-packages\skimage\util\dtype.py:141: UserWarning: Possible precision loss when converting from float32 to uint8
  .format(dtypeobj_in, dtypeobj_out))
 16%|█▋        | 7/43 [10:19<51:24, 85.68s/it]C:\Users\zfishlab\AppData\Local\Continuum\anaconda3\envs\test\lib\site-packages\skimage\util\dtype.py:141: UserWarning: Possible precision loss when converting from float32 to uint8
  .format(dtypeobj_in, dtypeobj_out))
 19%|█▊        | 8/43 [11:50<50:55, 87.30s/it]C:\Users\zfishlab\AppData\Local\Continuum\anaconda3\envs\test\lib\site-packages\skimage\util\dtype.py:141: UserWarning: Possible precision loss when converting from float32 to uint8
  .format(dtypeobj_in, dtypeobj_out))
 21%|██        | 9/43 [13:43<53:47, 94.93s/it]C:\Users\zfishlab\AppData\Local\Continuum\anaconda3\envs\test\lib\site-packages\skimage\util\dtype.py:141: UserWarning: Possible precision loss when converting from float32 to uint8
  .format(dtypeobj_in, dtypeobj_out))
 23%|██▎       | 10/43 [15:00<49:16, 89.61s/it]C:\Users\zfishlab\AppData\Local\Continuum\anaconda3\envs\test\lib\site-packages\skimage\util\dtype.py:141: UserWarning: Possible precision loss when converting from float32 to uint8
  .format(dtypeobj_in, dtypeobj_out))
 26%|██▌       | 11/43 [16:08<44:23, 83.25s/it]C:\Users\zfishlab\AppData\Local\Continuum\anaconda3\envs\test\lib\site-packages\skimage\util\dtype.py:141: UserWarning: Possible precision loss when converting from float32 to uint8
  .format(dtypeobj_in, dtypeobj_out))
 28%|██▊       | 12/43 [17:37<43:54, 84.97s/it]C:\Users\zfishlab\AppData\Local\Continuum\anaconda3\envs\test\lib\site-packages\skimage\util\dtype.py:141: UserWarning: Possible precision loss when converting from float32 to uint8
  .format(dtypeobj_in, dtypeobj_out))
 30%|███       | 13/43 [19:43<48:32, 97.09s/it]C:\Users\zfishlab\AppData\Local\Continuum\anaconda3\envs\test\lib\site-packages\skimage\util\dtype.py:141: UserWarning: Possible precision loss when converting from float32 to uint8
  .format(dtypeobj_in, dtypeobj_out))
 33%|███▎      | 14/43 [21:09<45:26, 94.00s/it]C:\Users\zfishlab\AppData\Local\Continuum\anaconda3\envs\test\lib\site-packages\skimage\util\dtype.py:141: UserWarning: Possible precision loss when converting from float32 to uint8
  .format(dtypeobj_in, dtypeobj_out))
 35%|███▍      | 15/43 [23:06<46:58, 100.67s/it]C:\Users\zfishlab\AppData\Local\Continuum\anaconda3\envs\test\lib\site-packages\skimage\util\dtype.py:141: UserWarning: Possible precision loss when converting from float32 to uint8
  .format(dtypeobj_in, dtypeobj_out))
 37%|███▋      | 16/43 [25:04<47:44, 106.10s/it]C:\Users\zfishlab\AppData\Local\Continuum\anaconda3\envs\test\lib\site-packages\skimage\util\dtype.py:141: UserWarning: Possible precision loss when converting from float32 to uint8
  .format(dtypeobj_in, dtypeobj_out))
 40%|███▉      | 17/43 [27:05<47:49, 110.35s/it]C:\Users\zfishlab\AppData\Local\Continuum\anaconda3\envs\test\lib\site-packages\skimage\util\dtype.py:141: UserWarning: Possible precision loss when converting from float32 to uint8
  .format(dtypeobj_in, dtypeobj_out))
 42%|████▏     | 18/43 [28:08<40:07, 96.30s/it] C:\Users\zfishlab\AppData\Local\Continuum\anaconda3\envs\test\lib\site-packages\skimage\util\dtype.py:141: UserWarning: Possible precision loss when converting from float32 to uint8
  .format(dtypeobj_in, dtypeobj_out))
 44%|████▍     | 19/43 [29:16<35:05, 87.73s/it]C:\Users\zfishlab\AppData\Local\Continuum\anaconda3\envs\test\lib\site-packages\skimage\util\dtype.py:141: UserWarning: Possible precision loss when converting from float32 to uint8
  .format(dtypeobj_in, dtypeobj_out))
 47%|████▋     | 20/43 [30:33<32:22, 84.45s/it]C:\Users\zfishlab\AppData\Local\Continuum\anaconda3\envs\test\lib\site-packages\skimage\util\dtype.py:141: UserWarning: Possible precision loss when converting from float32 to uint8
  .format(dtypeobj_in, dtypeobj_out))
 49%|████▉     | 21/43 [31:34<28:26, 77.59s/it]C:\Users\zfishlab\AppData\Local\Continuum\anaconda3\envs\test\lib\site-packages\skimage\util\dtype.py:141: UserWarning: Possible precision loss when converting from float32 to uint8
  .format(dtypeobj_in, dtypeobj_out))
 51%|█████     | 22/43 [32:12<22:56, 65.56s/it]C:\Users\zfishlab\AppData\Local\Continuum\anaconda3\envs\test\lib\site-packages\skimage\util\dtype.py:141: UserWarning: Possible precision loss when converting from float32 to uint8
  .format(dtypeobj_in, dtypeobj_out))
 53%|█████▎    | 23/43 [33:00<20:07, 60.39s/it]C:\Users\zfishlab\AppData\Local\Continuum\anaconda3\envs\test\lib\site-packages\skimage\util\dtype.py:141: UserWarning: Possible precision loss when converting from float32 to uint8
  .format(dtypeobj_in, dtypeobj_out))
 56%|█████▌    | 24/43 [34:23<21:15, 67.11s/it]C:\Users\zfishlab\AppData\Local\Continuum\anaconda3\envs\test\lib\site-packages\skimage\util\dtype.py:141: UserWarning: Possible precision loss when converting from float32 to uint8
  .format(dtypeobj_in, dtypeobj_out))
 58%|█████▊    | 25/43 [35:27<19:51, 66.19s/it]C:\Users\zfishlab\AppData\Local\Continuum\anaconda3\envs\test\lib\site-packages\skimage\util\dtype.py:141: UserWarning: Possible precision loss when converting from float32 to uint8
  .format(dtypeobj_in, dtypeobj_out))
 60%|██████    | 26/43 [37:39<24:22, 86.05s/it]C:\Users\zfishlab\AppData\Local\Continuum\anaconda3\envs\test\lib\site-packages\skimage\util\dtype.py:141: UserWarning: Possible precision loss when converting from float32 to uint8
  .format(dtypeobj_in, dtypeobj_out))
 63%|██████▎   | 27/43 [38:38<20:44, 77.76s/it]C:\Users\zfishlab\AppData\Local\Continuum\anaconda3\envs\test\lib\site-packages\skimage\util\dtype.py:141: UserWarning: Possible precision loss when converting from float32 to uint8
  .format(dtypeobj_in, dtypeobj_out))
 65%|██████▌   | 28/43 [40:34<22:21, 89.40s/it]C:\Users\zfishlab\AppData\Local\Continuum\anaconda3\envs\test\lib\site-packages\skimage\util\dtype.py:141: UserWarning: Possible precision loss when converting from float32 to uint8
  .format(dtypeobj_in, dtypeobj_out))
 67%|██████▋   | 29/43 [41:47<19:41, 84.42s/it]C:\Users\zfishlab\AppData\Local\Continuum\anaconda3\envs\test\lib\site-packages\skimage\util\dtype.py:141: UserWarning: Possible precision loss when converting from float32 to uint8
  .format(dtypeobj_in, dtypeobj_out))
 70%|██████▉   | 30/43 [43:45<20:30, 94.62s/it]C:\Users\zfishlab\AppData\Local\Continuum\anaconda3\envs\test\lib\site-packages\skimage\util\dtype.py:141: UserWarning: Possible precision loss when converting from float32 to uint8
  .format(dtypeobj_in, dtypeobj_out))
 72%|███████▏  | 31/43 [45:33<19:43, 98.59s/it]C:\Users\zfishlab\AppData\Local\Continuum\anaconda3\envs\test\lib\site-packages\skimage\util\dtype.py:141: UserWarning: Possible precision loss when converting from float32 to uint8
  .format(dtypeobj_in, dtypeobj_out))
 74%|███████▍  | 32/43 [47:41<19:41, 107.40s/it]C:\Users\zfishlab\AppData\Local\Continuum\anaconda3\envs\test\lib\site-packages\skimage\util\dtype.py:141: UserWarning: Possible precision loss when converting from float32 to uint8
  .format(dtypeobj_in, dtypeobj_out))
 77%|███████▋  | 33/43 [49:11<17:01, 102.14s/it]C:\Users\zfishlab\AppData\Local\Continuum\anaconda3\envs\test\lib\site-packages\skimage\util\dtype.py:141: UserWarning: Possible precision loss when converting from float32 to uint8
  .format(dtypeobj_in, dtypeobj_out))
 79%|███████▉  | 34/43 [50:19<13:45, 91.73s/it] C:\Users\zfishlab\AppData\Local\Continuum\anaconda3\envs\test\lib\site-packages\skimage\util\dtype.py:141: UserWarning: Possible precision loss when converting from float32 to uint8
  .format(dtypeobj_in, dtypeobj_out))
 81%|████████▏ | 35/43 [51:00<10:13, 76.64s/it]C:\Users\zfishlab\AppData\Local\Continuum\anaconda3\envs\test\lib\site-packages\skimage\util\dtype.py:141: UserWarning: Possible precision loss when converting from float32 to uint8
  .format(dtypeobj_in, dtypeobj_out))
 84%|████████▎ | 36/43 [52:55<10:17, 88.17s/it]C:\Users\zfishlab\AppData\Local\Continuum\anaconda3\envs\test\lib\site-packages\skimage\util\dtype.py:141: UserWarning: Possible precision loss when converting from float32 to uint8
  .format(dtypeobj_in, dtypeobj_out))
 86%|████████▌ | 37/43 [54:18<08:39, 86.65s/it]C:\Users\zfishlab\AppData\Local\Continuum\anaconda3\envs\test\lib\site-packages\skimage\util\dtype.py:141: UserWarning: Possible precision loss when converting from float32 to uint8
  .format(dtypeobj_in, dtypeobj_out))
 88%|████████▊ | 38/43 [55:34<06:56, 83.27s/it]C:\Users\zfishlab\AppData\Local\Continuum\anaconda3\envs\test\lib\site-packages\skimage\util\dtype.py:141: UserWarning: Possible precision loss when converting from float32 to uint8
  .format(dtypeobj_in, dtypeobj_out))
 91%|█████████ | 39/43 [57:17<05:57, 89.32s/it]C:\Users\zfishlab\AppData\Local\Continuum\anaconda3\envs\test\lib\site-packages\skimage\util\dtype.py:141: UserWarning: Possible precision loss when converting from float32 to uint8
  .format(dtypeobj_in, dtypeobj_out))
 93%|█████████▎| 40/43 [58:52<04:32, 90.95s/it]C:\Users\zfishlab\AppData\Local\Continuum\anaconda3\envs\test\lib\site-packages\skimage\util\dtype.py:141: UserWarning: Possible precision loss when converting from float32 to uint8
  .format(dtypeobj_in, dtypeobj_out))
 95%|█████████▌| 41/43 [1:00:45<03:15, 97.60s/it]C:\Users\zfishlab\AppData\Local\Continuum\anaconda3\envs\test\lib\site-packages\skimage\util\dtype.py:141: UserWarning: Possible precision loss when converting from float32 to uint8
  .format(dtypeobj_in, dtypeobj_out))
 98%|█████████▊| 42/43 [1:03:03<01:49, 109.66s/it]C:\Users\zfishlab\AppData\Local\Continuum\anaconda3\envs\test\lib\site-packages\skimage\util\dtype.py:141: UserWarning: Possible precision loss when converting from float32 to uint8
  .format(dtypeobj_in, dtypeobj_out))
100%|██████████| 43/43 [1:05:27<00:00, 119.93s/it]
```

```
Wall time: 1h 5min 27s
```

### Define experiment specific functions¶

In [17]:

```
''' Define wrapper functions for starting and saving to minimize the number 
of inputs that the user needs to type for each call of the function.'''
def start(k):
    return(ut.start(k,Dbat,[Dbzrf],im=True))
def save_both(k,dfa,dfb):
    ut.save_both(k,dfa,dfb,outdir,param['expname'])

'''Save model parameters for each file to a dataframe that can be 
exported for later reference.'''
model = pd.DataFrame({'a':[],'b':[],'c':[]})
def save_model(k,mm,model):
    row = pd.Series({'a':mm[0],'b':mm[1],'c':mm[2]},name=k)
    model = model.append(row)
    return(model)

'''Define a function that can both fit a model and plot it on an existing plot'''
def fit_model(axi,df,mm=None):
    if mm == None:
        mm = np.polyfit(df.x,df.z,2)
    p = np.poly1d(mm)
    xrange = np.arange(np.min(df.x),np.max(df.x))
    axi.plot(xrange,p(xrange),c='m')
    return(mm)

'''Take a set of points and transform to a dataframe format for ease of access.'''
def pick_pts(x1,z1,vx,vz,x2,z2):
    pts = pd.DataFrame({'x':[x1,vx,x2],'z':[z1,vz,z2]})
    return(pts)
```

### Align Samples¶

In [18]:

```
klist
```

Out[18]:

```
dict_keys(['101', '102', '103', '104', '105', '106', '107', '108', '109', '110', '111', '112', '113', '114', '115', '116', '117', '118', '119', '120', '121', '122', '123', '124', '125', '126', '127', '128', '129', '130', '131', '132', '133', '134', '135', '136', '137', '138', '139', '140', '141', '142', '143'])
```

In [19]:

```
k,df,Ldf,ax = start('101')
```

In [20]:

```
df1,Ldf1,pts,ax = ut.check_pts(df,Ldf,'z')
```

In [21]:

```
df2,Ldf2,mm,ax = ut.ch_vertex(df1,Ldf1)
```

In [22]:

```
model = save_model(k,mm,model)
save_both(k,df2,Ldf2[0])
```

```
Write to .\data\Output_wt03-09-21-29\AT_101_wt.psi complete
Write to .\data\Output_wt03-09-21-29\ZRF_101_wt.psi complete
```

In [23]:

```
k,df,Ldf,ax = start('102')
```

In [24]:

```
df1,Ldf1,pts,ax = ut.check_pts(df,Ldf,'z')
```

In [25]:

```
df2,Ldf2,ax,p = ut.check_yz(df1,Ldf1)
```

In [26]:

```
p
```

Out[26]:

```
poly1d([-0.01371249,  0.09442082])
```

[a,b] $y=ax+b$

In [27]:

```
p
```

Out[27]:

```
poly1d([-0.01371249,  0.09442082])
```

In [29]:

```
p1 = [5/30,0]
df3,Ldf3,ax,p = ut.check_yz(df1,Ldf1,mm=p1)
```

In [30]:

```
df4,Ldf4,mm,ax = ut.ch_vertex(df3,Ldf3)
```

In [31]:

```
model = save_model(k,mm,model)
save_both(k,df4,Ldf4[0])
```

```
Write to .\data\Output_wt03-09-21-29\AT_102_wt.psi complete
Write to .\data\Output_wt03-09-21-29\ZRF_102_wt.psi complete
```

In [32]:

```
k,df,Ldf,ax = start('103')
```

Sample 103 appears warped in both xy and yz projections and will be discarded from processing

In [33]:

```
klist
```

Out[33]:

```
dict_keys(['101', '102', '103', '104', '105', '106', '107', '108', '109', '110', '111', '112', '113', '114', '115', '116', '117', '118', '119', '120', '121', '122', '123', '124', '125', '126', '127', '128', '129', '130', '131', '132', '133', '134', '135', '136', '137', '138', '139', '140', '141', '142', '143'])
```

In [34]:

```
k,df,Ldf,ax = start('104')
```

In [ ]:

In [35]:

```
df1,Ldf1,pts,ax = ut.check_pts(df,Ldf,'z')
```

In [36]:

```
df2,Ldf2,mm,ax = ut.ch_vertex(df1,Ldf1)
```

In [37]:

```
model = save_model(k,mm,model)
save_both(k,df2,Ldf2[0])
```

```
Write to .\data\Output_wt03-09-21-29\AT_104_wt.psi complete
Write to .\data\Output_wt03-09-21-29\ZRF_104_wt.psi complete
```

In [38]:

```
k,df,Ldf,ax = start('105')
```

In [39]:

```
p
```

Out[39]:

```
poly1d([0.16666667, 0.        ])
```

In [40]:

```
df1,Ldf1,ax,p = ut.check_yz(df,Ldf)
```

In [41]:

```
p
```

Out[41]:

```
poly1d([ 0.01671887, -0.12012197])
```

In [42]:

```
p1=[.0167,-3.0]
```

In [43]:

```
p1
```

Out[43]:

```
[0.0167, -3.0]
```

In [44]:

```
df2,Ldf2,ax,p1 = ut.check_yz(df1,Ldf1)
```

In [45]:

```
model = save_model(k,mm,model)
save_both(k,df2,Ldf2[0])
```

```
Write to .\data\Output_wt03-09-21-29\AT_105_wt.psi complete
Write to .\data\Output_wt03-09-21-29\ZRF_105_wt.psi complete
```

In [46]:

```
k,df,Ldf,ax = start('106')
```

In [47]:

```
model = save_model(k,mm,model)
save_both(k,df,Ldf[0])
```

```
Write to .\data\Output_wt03-09-21-29\AT_106_wt.psi complete
Write to .\data\Output_wt03-09-21-29\ZRF_106_wt.psi complete
```

In [49]:

```
k,df,Ldf,ax = start('107')
save_both(k,df3,Ldf3[0])
```

```
Write to .\data\Output_wt03-09-21-29\AT_107_wt.psi complete
Write to .\data\Output_wt03-09-21-29\ZRF_107_wt.psi complete
```

In [50]:

```
model = save_model(k,mm,model)
save_both(k,df,Ldf[0])
```

```
Write to .\data\Output_wt03-09-21-29\AT_107_wt.psi complete
Write to .\data\Output_wt03-09-21-29\ZRF_107_wt.psi complete
```

In [51]:

```
k,df,Ldf,ax = start('108')
```

In [52]:

```
df1,Ldf1,pts,ax = ut.check_pts(df,Ldf,'z')
```

In [53]:

```
pts.iloc[0].x = 50
pts.iloc[1].z = 38
```

In [54]:

```
pts
```

Out[54]:

|  | x | z |
| --- | --- | --- |
| 0 | 50.000000 | 44.1625 |
| 1 | 52.855046 | 38.0000 |

In [55]:

```
pts.iloc[0].x = -50
```

In [ ]:

In [56]:

```
ax[0,1].scatter(pts.x,pts.z,c='y')
```

Out[56]:

```
<matplotlib.collections.PathCollection at 0x27a3b07bb00>
```

In [57]:

```
df2,Ldf2,ax = ut.revise_pts(df,Ldf,'z',pts=pts)
```

In [58]:

```
df3,Ldf3,mm,ax = ut.ch_vertex(df2,Ldf2,pts=pts)
```

```
c:\users\zfishlab\code\deltascope\deltascope\alignment.py:236: RankWarning: Polyfit may be poorly conditioned
  cf = np.polyfit(pts.x,pts.z,2)
c:\users\zfishlab\code\deltascope\deltascope\alignment.py:257: RankWarning: Polyfit may be poorly conditioned
  cfout = np.polyfit(pts.x-x,pts.z-z,2)
```

In [59]:

```
df3,Ldf3,mm,ax = ut.ch_vertex(df2,Ldf2,pts=pts)
```

```
c:\users\zfishlab\code\deltascope\deltascope\alignment.py:236: RankWarning: Polyfit may be poorly conditioned
  cf = np.polyfit(pts.x,pts.z,2)
c:\users\zfishlab\code\deltascope\deltascope\alignment.py:257: RankWarning: Polyfit may be poorly conditioned
  cfout = np.polyfit(pts.x-x,pts.z-z,2)
```

In [60]:

```
pts = pick_pts(-55,40,-8,-2,50,35) #these are essentially random numbers for example
```

In [61]:

```
df3,Ldf3,mm,ax = ut.ch_vertex(df2,Ldf2,pts=pts)
```

In [62]:

```
model = save_model(k,mm,model)
save_both(k,df3,Ldf3[0])
```

```
Write to .\data\Output_wt03-09-21-29\AT_108_wt.psi complete
Write to .\data\Output_wt03-09-21-29\ZRF_108_wt.psi complete
```

k,df,Ldf,ax = start('109')df1,Ldf1,pts,ax = ut.check\_pts(df,Ldf,'z')

In [63]:

```
df2,Ldf2,mm,ax = ut.ch_vertex(df1,Ldf1)
```

In [64]:

```
pts = pick_pts(-53,40,-9,0,60,40)
```

df3,Ldf3,mm,ax = ut.ch\_vertex(df2,Ldf2,pts=pts)

In [65]:

```
model = save_model(k,mm,model)
save_both(k,df3,Ldf3[0])
```

```
Write to .\data\Output_wt03-09-21-29\AT_108_wt.psi complete
Write to .\data\Output_wt03-09-21-29\ZRF_108_wt.psi complete
```

k,df,Ldf,ax= start('110')

In [66]:

```
model = save_model(k,mm,model)
save_both(k,df,Ldf[0])
```

```
Write to .\data\Output_wt03-09-21-29\AT_108_wt.psi complete
Write to .\data\Output_wt03-09-21-29\ZRF_108_wt.psi complete
```

k,df,Ldf,ax = start('111')df1,Ldf1,pts,ax = ut.check\_pts(df,Ldf,'z')df2,Ldf2,mm,ax = ut.ch\_vertex(df1,Ldf1)

In [67]:

```
model = save_model(k,mm,model)
save_both(k,df2,Ldf2[0])
```

```
Write to .\data\Output_wt03-09-21-29\AT_108_wt.psi complete
Write to .\data\Output_wt03-09-21-29\ZRF_108_wt.psi complete
```

k,df,Ldf,ax = start('112')df1,Ldf1,pts,ax = ut.check\_pts(df,Ldf,'z')df2,Ldf2,mm,ax = ut.ch\_vertex(df1,Ldf1)

In [68]:

```
model = save_model(k,mm,model)
save_both(k,df2,Ldf2[0])
```

```
Write to .\data\Output_wt03-09-21-29\AT_108_wt.psi complete
Write to .\data\Output_wt03-09-21-29\ZRF_108_wt.psi complete
```

k,df,Ldf,ax = start('113')df1,Ldf1,pts,ax = ut.check\_pts(df,Ldf,'z')df2,Ldf2,mm,ax = ut.ch\_vertex(df1,Ldf1)

In [69]:

```
model = save_model(k,mm,model)
save_both(k,df2,Ldf2[0])
```

```
Write to .\data\Output_wt03-09-21-29\AT_108_wt.psi complete
Write to .\data\Output_wt03-09-21-29\ZRF_108_wt.psi complete
```

k,df,Ldf,ax = start('114')

sample rejected for being truncated at midline. likely improper collection that was not superficial enough to capture POC.

k,df,Ldf,ax = start('115')

In [70]:

```
model = save_model(k,mm,model)
save_both(k,df,Ldf[0])
```

```
Write to .\data\Output_wt03-09-21-29\AT_108_wt.psi complete
Write to .\data\Output_wt03-09-21-29\ZRF_108_wt.psi complete
```

k,df,Ldf,ax = start('116')df1,Ldf1,ax,p = ut.check\_yz(df,Ldf)

In [71]:

```
model = save_model(k,mm,model)
save_both(k,df1,Ldf1[0])
```

```
Write to .\data\Output_wt03-09-21-29\AT_108_wt.psi complete
Write to .\data\Output_wt03-09-21-29\ZRF_108_wt.psi complete
```

k,df,Ldf,ax = start('117')

In [72]:

```
model = save_model(k,mm,model)
save_both(k,df1,Ldf1[0])
```

```
Write to .\data\Output_wt03-09-21-29\AT_108_wt.psi complete
Write to .\data\Output_wt03-09-21-29\ZRF_108_wt.psi complete
```

k,df,Ldf,ax = start('118')df1,Ldf1,ax,p = ut.check\_yz(df,Ldf)df2,Ldf2,pts,ax = ut.check\_pts(df1,Ldf1,'z')df3,Ldf3,mm,ax = ut.ch\_vertex(df2,Ldf2)

In [73]:

```
model = save_model(k,mm,model)
save_both(k,df3,Ldf3[0])
```

```
Write to .\data\Output_wt03-09-21-29\AT_108_wt.psi complete
Write to .\data\Output_wt03-09-21-29\ZRF_108_wt.psi complete
```

k,df,Ldf,ax =start('119')df1,Ldf1,pts,ax = ut.check\_pts(df,Ldf,'z')df2,Ldf2,mm,ax = ut.ch\_vertex(df1,Ldf1)

In [74]:

```
model = save_model(k,mm,model)
save_both(k,df2,Ldf2[0])
```

```
Write to .\data\Output_wt03-09-21-29\AT_108_wt.psi complete
Write to .\data\Output_wt03-09-21-29\ZRF_108_wt.psi complete
```

In [75]:

```
klist
```

Out[75]:

```
dict_keys(['101', '102', '103', '104', '105', '106', '107', '108', '109', '110', '111', '112', '113', '114', '115', '116', '117', '118', '119', '120', '121', '122', '123', '124', '125', '126', '127', '128', '129', '130', '131', '132', '133', '134', '135', '136', '137', '138', '139', '140', '141', '142', '143'])
```

k,df,Ldf,ax = start('120')

In [76]:

```
df1,Ldf1,pts,ax = ut.check_pts(df,Ldf,'z')
```

df2,Ldf2,mm,ax = ut.ch\_vertex(df1,Ldf1)

In [77]:

```
model = save_model(k,mm,model)
save_both(k,df2,Ldf2[0])
```

```
Write to .\data\Output_wt03-09-21-29\AT_108_wt.psi complete
Write to .\data\Output_wt03-09-21-29\ZRF_108_wt.psi complete
```

k,df,Ldf,ax = start('121')

Commissure is obviously physically deformed due to pressure from coverslip and will not take parabolic curve well. skipping sample.

In [78]:

```
k,df,Ldf,ax = start('122')
```

In [79]:

```
model = save_model(k,mm,model)
save_both(k,df,Ldf[0])
```

```
Write to .\data\Output_wt03-09-21-29\AT_122_wt.psi complete
Write to .\data\Output_wt03-09-21-29\ZRF_122_wt.psi complete
```

In [80]:

```
k,df,Ldf,ax = start('123')
```

In [81]:

```
model = save_model(k,mm,model)
save_both(k,df,Ldf[0])
```

```
Write to .\data\Output_wt03-09-21-29\AT_123_wt.psi complete
Write to .\data\Output_wt03-09-21-29\ZRF_123_wt.psi complete
```

In [82]:

```
k,df,Ldf,ax = start('124')
```

In [83]:

```
model = save_model(k,mm,model)
save_both(k,df,Ldf[0])
```

```
Write to .\data\Output_wt03-09-21-29\AT_124_wt.psi complete
Write to .\data\Output_wt03-09-21-29\ZRF_124_wt.psi complete
```

In [84]:

```
k,df,Ldf,ax = start('125')
```

In [85]:

```
df1,Ldf1,pts,ax = ut.check_pts(df,Ldf,'z')
```

In [86]:

```
df2,Ldf2,mm,ax = ut.ch_vertex(df1,Ldf1)
```

In [87]:

```
model = save_model(k,mm,model)
save_both(k,df2,Ldf2[0])
```

```
Write to .\data\Output_wt03-09-21-29\AT_125_wt.psi complete
Write to .\data\Output_wt03-09-21-29\ZRF_125_wt.psi complete
```

In [88]:

```
k,df,Ldf,ax = start('126')
```

In [89]:

```
df1,Ldf1,ax,p = ut.check_yz(df,Ldf)
```

In [90]:

```
p
```

Out[90]:

```
poly1d([-0.02718542,  0.25853679])
```

In [91]:

```
p1 = [-5/20, 2]
```

In [92]:

```
df1,Ldf1,ax,p1 = ut.check_yz(df,Ldf)
```

In [93]:

```
df1,Ldf1,ax,p = ut.check_yz(df,Ldf,mm=p1)
```

In [94]:

```
p1
```

Out[94]:

```
poly1d([-0.02718542,  0.25853679])
```

In [95]:

```
p2 =[-.5, 2]
```

In [96]:

```
df1,Ldf1,ax,p = ut.check_yz(df,Ldf,mm=p2)
```

k,df,Ldf,ax=start('128')df,Ldf,mm,ax = ut.ch\_vertex(df,Ldf)

In [97]:

```
model = save_model(k,mm,model)
save_both(k,df,Ldf[0])
```

```
Write to .\data\Output_wt03-09-21-29\AT_126_wt.psi complete
Write to .\data\Output_wt03-09-21-29\ZRF_126_wt.psi complete
```

In [98]:

```
k,df,Ldf,ax=start('129')
```

In [99]:

```
df,Ldf,mm,ax = ut.ch_vertex(df,Ldf)
model = save_model(k,mm,model)
save_both(k,df,Ldf[0])
```

```
Write to .\data\Output_wt03-09-21-29\AT_129_wt.psi complete
Write to .\data\Output_wt03-09-21-29\ZRF_129_wt.psi complete
```

In [100]:

```
k,df,Ldf,ax=start('130')
```

In [101]:

```
df,Ldf,mm,ax = ut.ch_vertex(df,Ldf)
```

In [102]:

```
model = save_model(k,mm,model)
save_both(k,df,Ldf[0])
```

```
Write to .\data\Output_wt03-09-21-29\AT_130_wt.psi complete
Write to .\data\Output_wt03-09-21-29\ZRF_130_wt.psi complete
```

In [103]:

```
k,df,Ldf,ax=start('131')
```

In [104]:

```
df,Ldf,pts,ax = ut.check_pts(df,Ldf,'z')
```

In [105]:

```
df2,Ldf2,mm,ax = ut.ch_vertex(df,Ldf)
```

In [106]:

```
model = save_model(k,mm,model)
save_both(k,df2,Ldf2[0])
```

```
Write to .\data\Output_wt03-09-21-29\AT_131_wt.psi complete
Write to .\data\Output_wt03-09-21-29\ZRF_131_wt.psi complete
```

In [107]:

```
k,df,Ldf,ax=start('128')
```

In [108]:

```
df1,Ldf1,pts,ax = ut.check_pts(df,Ldf,'z')
```

In [109]:

```
df2,Ldf2,mm,ax = ut.ch_vertex(df1,Ldf1)
```

In [110]:

```
model = save_model(k,mm,model)
save_both(k,df2,Ldf2[0])
```

```
Write to .\data\Output_wt03-09-21-29\AT_128_wt.psi complete
Write to .\data\Output_wt03-09-21-29\ZRF_128_wt.psi complete
```

In [111]:

```
model
```

Out[111]:

|  | a | b | c |
| --- | --- | --- | --- |
| 101 | 0.010452 | -4.985302e-17 | -1.016496e-14 |
| 102 | 0.007950 | -3.772889e-17 | 6.879950e-15 |
| 104 | 0.013104 | -7.465448e-16 | -1.424932e-14 |
| 105 | 0.013104 | -7.465448e-16 | -1.424932e-14 |
| 106 | 0.013104 | -7.465448e-16 | -1.424932e-14 |
| 107 | 0.013104 | -7.465448e-16 | -1.424932e-14 |
| 108 | 0.014586 | -2.110834e-16 | -1.755301e-14 |
| 108 | 0.012598 | -6.073178e-16 | 7.742279e-15 |
| 108 | 0.012598 | -6.073178e-16 | 7.742279e-15 |
| 108 | 0.012598 | -6.073178e-16 | 7.742279e-15 |
| 108 | 0.012598 | -6.073178e-16 | 7.742279e-15 |
| 108 | 0.012598 | -6.073178e-16 | 7.742279e-15 |
| 108 | 0.012598 | -6.073178e-16 | 7.742279e-15 |
| 108 | 0.012598 | -6.073178e-16 | 7.742279e-15 |
| 108 | 0.012598 | -6.073178e-16 | 7.742279e-15 |
| 108 | 0.012598 | -6.073178e-16 | 7.742279e-15 |
| 108 | 0.012598 | -6.073178e-16 | 7.742279e-15 |
| 108 | 0.012598 | -6.073178e-16 | 7.742279e-15 |
| 122 | 0.012598 | -6.073178e-16 | 7.742279e-15 |
| 123 | 0.012598 | -6.073178e-16 | 7.742279e-15 |
| 124 | 0.012598 | -6.073178e-16 | 7.742279e-15 |
| 125 | 0.021211 | 8.045999e-16 | 8.068820e-16 |
| 126 | 0.021211 | 8.045999e-16 | 8.068820e-16 |
| 129 | 0.013999 | -2.511842e-17 | -2.442675e-15 |
| 130 | 0.005348 | 5.544343e-18 | 1.539247e-15 |
| 131 | 0.009818 | 3.432638e-16 | 3.693030e-15 |
| 128 | 0.009238 | -7.245532e-16 | 4.018019e-15 |

In [112]:

```
k,df,Ldf,ax=start('127')
```

In [113]:

```
df1,Ldf1,pts,ax = ut.check_pts(df,Ldf,'z')
```

In [114]:

```
k,df,Ldf,ax=start('132')
```

In [115]:

```
df1,Ldf1,pts,ax = ut.check_pts(df,Ldf,'z')
```

In [116]:

```
df2,Ldf2,mm,ax = ut.ch_vertex(df1,Ldf1)
```

In [117]:

```
model = save_model(k,mm,model)
save_both(k,df2,Ldf2[0])
```

```
Write to .\data\Output_wt03-09-21-29\AT_132_wt.psi complete
Write to .\data\Output_wt03-09-21-29\ZRF_132_wt.psi complete
```

In [118]:

```
k,df,Ldf,ax=start('133')
```

In [119]:

```
df1,Ldf1,pts,ax = ut.check_pts(df,Ldf,'z')
```

In [120]:

```
df2,Ldf2,mm,ax = ut.ch_vertex(df1,Ldf1)
```

In [121]:

```
model = save_model(k,mm,model)
save_both(k,df2,Ldf2[0])
```

```
Write to .\data\Output_wt03-09-21-29\AT_133_wt.psi complete
Write to .\data\Output_wt03-09-21-29\ZRF_133_wt.psi complete
```

In [122]:

```
k,df,Ldf,ax=start('134')
```

In [123]:

```
df2,Ldf2,mm,ax = ut.ch_vertex(df,Ldf)
```

In [124]:

```
model = save_model(k,mm,model)
save_both(k,df2,Ldf2[0])
```

```
Write to .\data\Output_wt03-09-21-29\AT_134_wt.psi complete
Write to .\data\Output_wt03-09-21-29\ZRF_134_wt.psi complete
```

In [125]:

```
k,df,Ldf,ax=start('135')
```

In [126]:

```
df1,Ldf1,pts,ax = ut.check_pts(df,Ldf,'z')
```

In [127]:

```
df2,Ldf2,mm,ax = ut.ch_vertex(df1,Ldf1)
```

In [128]:

```
df3,Ldf3,ax,p = ut.check_yz(df2,Ldf2)
```

In [129]:

```
model = save_model(k,mm,model)
save_both(k,df2,Ldf2[0])
```

```
Write to .\data\Output_wt03-09-21-29\AT_135_wt.psi complete
Write to .\data\Output_wt03-09-21-29\ZRF_135_wt.psi complete
```

In [ ]:

### Wrapping up: after all samples are processed¶

Once all of the data has been processed, we want to save the model data we collected to a file for future reference.

In [ ]:

In [130]:

```
model.to_csv(os.path.join(outdir,'model.csv'))
```

Additionally, it can be helpful to export a html or pdf version of the notebook that preserves all of the plots generated during the alignment process. To do so, use the Jupyter Lab menu interface: File > Export Notebook As > HTML or PDF.

In [131]:

```
klist
```

Out[131]:

```
dict_keys(['101', '102', '103', '104', '105', '106', '107', '108', '109', '110', '111', '112', '113', '114', '115', '116', '117', '118', '119', '120', '121', '122', '123', '124', '125', '126', '127', '128', '129', '130', '131', '132', '133', '134', '135', '136', '137', '138', '139', '140', '141', '142', '143'])
```

In [132]:

```
k,df,Ldf,ax = start('136')
```

In [133]:

```
df2,Ldf2,mm,ax = ut.ch_vertex(df1,Ldf1)
```

In [134]:

```
model = save_model(k,mm,model)
save_both(k,df2,Ldf2[0])
```

```
Write to .\data\Output_wt03-09-21-29\AT_136_wt.psi complete
Write to .\data\Output_wt03-09-21-29\ZRF_136_wt.psi complete
```

In [135]:

```
k,df,Ldf,ax = start('137')
```

In [136]:

```
df1,Ldf1,mm,ax = ut.ch_vertex(df,Ldf)
```

In [137]:

```
model = save_model(k,mm,model)
save_both(k,df1,Ldf1[0])
```

```
Write to .\data\Output_wt03-09-21-29\AT_137_wt.psi complete
Write to .\data\Output_wt03-09-21-29\ZRF_137_wt.psi complete
```

In [138]:

```
k,df,Ldf,ax = start('138')
```

In [139]:

```
df1,Ldf1,mm,ax = ut.ch_vertex(df,Ldf)
```

In [140]:

```
model = save_model(k,mm,model)
save_both(k,df1,Ldf1[0])
```

```
Write to .\data\Output_wt03-09-21-29\AT_138_wt.psi complete
Write to .\data\Output_wt03-09-21-29\ZRF_138_wt.psi complete
```

In [141]:

```
k,df,Ldf,ax = start('139')
```

In [142]:

```
df1,Ldf1,mm,ax = ut.ch_vertex(df,Ldf)
```

In [143]:

```
df1,Ldf1,pts,ax = ut.check_pts(df,Ldf,'z')
```

In [144]:

```
df2,Ldf2,mm,ax = ut.ch_vertex(df1,Ldf1)
```

In [145]:

```
model = save_model(k,mm,model)
save_both(k,df2,Ldf2[0])
```

```
Write to .\data\Output_wt03-09-21-29\AT_139_wt.psi complete
Write to .\data\Output_wt03-09-21-29\ZRF_139_wt.psi complete
```

In [146]:

```
k,df,Ldf,ax = start('140')
```

sample highly disturbed in imaging structure, possibly due to tear on left of the image causing loss of data, Vertex will be highly questionable.

In [147]:

```
k,df,Ldf,ax = start('141')
```

In [148]:

```
df1,Ldf1,mm,ax = ut.ch_vertex(df,Ldf)
```

In [149]:

```
model = save_model(k,mm,model)
save_both(k,df1,Ldf1[0])
```

```
Write to .\data\Output_wt03-09-21-29\AT_141_wt.psi complete
Write to .\data\Output_wt03-09-21-29\ZRF_141_wt.psi complete
```

In [150]:

```
k,df,Ldf,ax = start('142')
```

In [151]:

```
df1,Ldf1,mm,ax = ut.ch_vertex(df,Ldf)
```

In [152]:

```
model = save_model(k,mm,model)
save_both(k,df1,Ldf1[0])
```

```
Write to .\data\Output_wt03-09-21-29\AT_142_wt.psi complete
Write to .\data\Output_wt03-09-21-29\ZRF_142_wt.psi complete
```

In [153]:

```
k,df,Ldf,ax = start('143')
```

In [154]:

```
df1,Ldf1,mm,ax = ut.ch_vertex(df,Ldf)
```

In [155]:

```
model = save_model(k,mm,model)
save_both(k,df1,Ldf1[0])
```

```
Write to .\data\Output_wt03-09-21-29\AT_143_wt.psi complete
Write to .\data\Output_wt03-09-21-29\ZRF_143_wt.psi complete
```

In [156]:

```
klist
```

Out[156]:

```
dict_keys(['101', '102', '103', '104', '105', '106', '107', '108', '109', '110', '111', '112', '113', '114', '115', '116', '117', '118', '119', '120', '121', '122', '123', '124', '125', '126', '127', '128', '129', '130', '131', '132', '133', '134', '135', '136', '137', '138', '139', '140', '141', '142', '143'])
```

In [157]:

```
k,df,Ldf,ax = start('105')
```

In [158]:

```
df1,Ldf1,mm,ax = ut.ch_vertex(df,Ldf)
```

In [159]:

```
model = save_model(k,mm,model)
save_both(k,df1,Ldf1[0])
```

```
Write to .\data\Output_wt03-09-21-29\AT_105_wt.psi complete
Write to .\data\Output_wt03-09-21-29\ZRF_105_wt.psi complete
```

In [160]:

```
k,df,Ldf,ax = start('106')
```

In [161]:

```
df1,Ldf1,mm,ax = ut.ch_vertex(df,Ldf)
```

In [162]:

```
model = save_model(k,mm,model)
save_both(k,df1,Ldf1[0])
```

```
Write to .\data\Output_wt03-09-21-29\AT_106_wt.psi complete
Write to .\data\Output_wt03-09-21-29\ZRF_106_wt.psi complete
```

In [163]:

```
k,df,Ldf,ax = start('107')
```

In [164]:

```
df1,Ldf1,mm,ax = ut.ch_vertex(df,Ldf)
```

In [165]:

```
model = save_model(k,mm,model)
save_both(k,df1,Ldf1[0])
```

```
Write to .\data\Output_wt03-09-21-29\AT_107_wt.psi complete
Write to .\data\Output_wt03-09-21-29\ZRF_107_wt.psi complete
```

In [166]:

```
klist
```

Out[166]:

```
dict_keys(['101', '102', '103', '104', '105', '106', '107', '108', '109', '110', '111', '112', '113', '114', '115', '116', '117', '118', '119', '120', '121', '122', '123', '124', '125', '126', '127', '128', '129', '130', '131', '132', '133', '134', '135', '136', '137', '138', '139', '140', '141', '142', '143'])
```

In [167]:

```
k,df,Ldf,ax = start('110')
```

In [168]:

```
df1,Ldf1,mm,ax = ut.ch_vertex(df,Ldf)
```

In [169]:

```
model = save_model(k,mm,model)
save_both(k,df1,Ldf1[0])
```

```
Write to .\data\Output_wt03-09-21-29\AT_110_wt.psi complete
Write to .\data\Output_wt03-09-21-29\ZRF_110_wt.psi complete
```

In [170]:

```
k,df,Ldf,ax = start('115')
```

In [171]:

```
df1,Ldf1,mm,ax = ut.ch_vertex(df,Ldf)
```

In [172]:

```
model = save_model(k,mm,model)
save_both(k,df1,Ldf1[0])
```

```
Write to .\data\Output_wt03-09-21-29\AT_115_wt.psi complete
Write to .\data\Output_wt03-09-21-29\ZRF_115_wt.psi complete
```

In [173]:

```
klist
```

Out[173]:

```
dict_keys(['101', '102', '103', '104', '105', '106', '107', '108', '109', '110', '111', '112', '113', '114', '115', '116', '117', '118', '119', '120', '121', '122', '123', '124', '125', '126', '127', '128', '129', '130', '131', '132', '133', '134', '135', '136', '137', '138', '139', '140', '141', '142', '143'])
```

In [174]:

```
k,df,Ldf,ax = start('116')
```

In [175]:

```
df1,Ldf1,mm,ax = ut.ch_vertex(df,Ldf)
```

In [176]:

```
model = save_model(k,mm,model)
save_both(k,df1,Ldf1[0])
```

```
Write to .\data\Output_wt03-09-21-29\AT_116_wt.psi complete
Write to .\data\Output_wt03-09-21-29\ZRF_116_wt.psi complete
```

In [177]:

```
k,df,Ldf,ax = start('117')
```

In [178]:

```
df1,Ldf1,mm,ax = ut.ch_vertex(df,Ldf)
```

In [179]:

```
model = save_model(k,mm,model)
save_both(k,df1,Ldf1[0])
```

```
Write to .\data\Output_wt03-09-21-29\AT_117_wt.psi complete
Write to .\data\Output_wt03-09-21-29\ZRF_117_wt.psi complete
```

In [180]:

```
k,df,Ldf,ax = start('122')
```

In [181]:

```
df1,Ldf1,mm,ax = ut.ch_vertex(df,Ldf)
```

In [182]:

```
model = save_model(k,mm,model)
save_both(k,df1,Ldf1[0])
```

```
Write to .\data\Output_wt03-09-21-29\AT_122_wt.psi complete
Write to .\data\Output_wt03-09-21-29\ZRF_122_wt.psi complete
```

In [183]:

```
k,df,Ldf,ax = start('123')
```

In [184]:

```
df1,Ldf1,mm,ax = ut.ch_vertex(df,Ldf)
```

In [185]:

```
model = save_model(k,mm,model)
save_both(k,df1,Ldf1[0])
```

```
Write to .\data\Output_wt03-09-21-29\AT_123_wt.psi complete
Write to .\data\Output_wt03-09-21-29\ZRF_123_wt.psi complete
```

In [186]:

```
k,df,Ldf,ax = start('124')
```

In [187]:

```
df1,Ldf1,mm,ax = ut.ch_vertex(df,Ldf)
```

In [188]:

```
model = save_model(k,mm,model)
save_both(k,df1,Ldf1[0])
```

```
Write to .\data\Output_wt03-09-21-29\AT_124_wt.psi complete
Write to .\data\Output_wt03-09-21-29\ZRF_124_wt.psi complete
```

In [189]:

```
k,df,Ldf,ax = start('126')
```

In [190]:

```
df1,Ldf1,mm,ax = ut.ch_vertex(df,Ldf)
```

In [191]:

```
model = save_model(k,mm,model)
save_both(k,df1,Ldf1[0])
```

```
Write to .\data\Output_wt03-09-21-29\AT_126_wt.psi complete
Write to .\data\Output_wt03-09-21-29\ZRF_126_wt.psi complete
```

In [193]:

```
model.to_csv(os.path.join(outdir,'models.csv'))
```
