## Supplementary material for "ΔSCOPE: A new method to quantify 3D biological structures and identify differences in zebrafish forebrain development": An archived copy of the DeltaSCOPE code repository.: alignment-wt.ipynb

```
# %% [markdown]
# Introduction
This notebook facilitates the manual curation of sample alignment.

# %% [code]
import deltascope.alignment as ut

import numpy as np
import pandas as pd
import matplotlib.pyplot as plt
import h5py

import os
import re
import time
import tqdm

# %% [markdown]
# Setup

# %% [markdown]
## Parameters

# %% [code]
# --------------------------------
# -------- User input ------------
# --------------------------------

param = {
    'gthresh':0.5,
    'scale':[1,1,1],
    'microns':[0.16,0.16,0.21],
    'mthresh':0.5,
    'radius':20,
    'comp_order':[0,2,1],
    'fit_dim':['x','z'],
    'deg':2,
    
    # Don't forget to modify this with a specific sample name
    'expname':'wt'
}

# %% [markdown]
## Directories

# %% [code]
# --------------------------------
# -------- User input ------------
# --------------------------------

# Specify file paths to directories containing probability files
# after processing by ilastik
gfap = '.\data\ZRF1 Wild Type HZ H5 files\Prob_7-29'
at = '.\data\AT Wild Type HZ H5 files\Prob'

# Specify root directory where output will be saved
root = '.\data'

root

# %% [code]
# Output directory with timestamp
outname = 'Output_'+param['expname']+time.strftime("%m-%d-%H-%M",
                                 time.localtime())

# Create output directory
outdir = os.path.join(root,outname)
os.mkdir(outdir)

# %% [markdown]
## Extract list of files

# %% [code]
Dat = {}
for f in os.listdir(at):
    if 'h5' in f:
        num  = re.findall(r'\d+',f.split('.')[0])[-1]
        Dat[num] = os.path.join(at,f)

# %% [code]
Dzrf = {}
for f in os.listdir(gfap):
    if 'h5' in f:
        num  = re.findall(r'\d+',f.split('.')[0])[-1]
        Dzrf[num] = os.path.join(gfap,f)

# %% [code]
# Create dictionaries to contain the deltascope brain object for each sample
Dbat = {}
Dbzrf = {}

# %% [code]
klist = Dzrf.keys()
klist

# %% [markdown]
# PCA: Import raw data and perform preprocessing

# %% [code]
%%time
for k in tqdm.tqdm(klist):
    if k not in list(Dbat.keys()):
        try:
            Dbat[k] = ut.preprocess(Dat[k],param)
            Dbzrf[k] = ut.preprocess(Dzrf[k],param,pca=Dbat[k].pcamed,
                                 mm=Dbat[k].mm,vertex=Dbat[k].vertex)
        except:
            print(k,'caused an error')
    else:
        print(k,'already processed')

# %% [markdown]
# Define experiment specific functions

# %% [code]
''' Define wrapper functions for starting and saving to minimize the number 
of inputs that the user needs to type for each call of the function.'''
def start(k):
    return(ut.start(k,Dbat,[Dbzrf],im=True))
def save_both(k,dfa,dfb):
    ut.save_both(k,dfa,dfb,outdir,param['expname'])

'''Save model parameters for each file to a dataframe that can be 
exported for later reference.'''
model = pd.DataFrame({'a':[],'b':[],'c':[]})
def save_model(k,mm,model):
    row = pd.Series({'a':mm[0],'b':mm[1],'c':mm[2]},name=k)
    model = model.append(row)
    return(model)

'''Define a function that can both fit a model and plot it on an existing plot'''
def fit_model(axi,df,mm=None):
    if mm == None:
        mm = np.polyfit(df.x,df.z,2)
    p = np.poly1d(mm)
    xrange = np.arange(np.min(df.x),np.max(df.x))
    axi.plot(xrange,p(xrange),c='m')
    return(mm)

'''Take a set of points and transform to a dataframe format for ease of access.'''
def pick_pts(x1,z1,vx,vz,x2,z2):
    pts = pd.DataFrame({'x':[x1,vx,x2],'z':[z1,vz,z2]})
    return(pts)

# %% [markdown]
# Align Samples

# %% [code]
klist

# %% [code]
k,df,Ldf,ax = start('101')

# %% [code]
df1,Ldf1,pts,ax = ut.check_pts(df,Ldf,'z')

# %% [code]
df2,Ldf2,mm,ax = ut.ch_vertex(df1,Ldf1)

# %% [code]
model = save_model(k,mm,model)
save_both(k,df2,Ldf2[0])

# %% [code]
k,df,Ldf,ax = start('102')

# %% [code]
df1,Ldf1,pts,ax = ut.check_pts(df,Ldf,'z')

# %% [code]
df2,Ldf2,ax,p = ut.check_yz(df1,Ldf1)

# %% [code]
p

# %% [markdown]
\[a,b] $y=ax+b$

# %% [code]
p

# %% [code]
p1 = [5/30,0]
df3,Ldf3,ax,p = ut.check_yz(df1,Ldf1,mm=p1)

# %% [code]
df4,Ldf4,mm,ax = ut.ch_vertex(df3,Ldf3)

# %% [code]
model = save_model(k,mm,model)
save_both(k,df4,Ldf4[0])

# %% [code]
k,df,Ldf,ax = start('103')

# %% [markdown]
Sample 103 appears warped in both xy and yz projections and will be discarded from processing

# %% [code]
klist

# %% [code]
k,df,Ldf,ax = start('104')

# %% [code]


# %% [code]
df1,Ldf1,pts,ax = ut.check_pts(df,Ldf,'z')

# %% [code]
df2,Ldf2,mm,ax = ut.ch_vertex(df1,Ldf1)

# %% [code]
model = save_model(k,mm,model)
save_both(k,df2,Ldf2[0])

# %% [code]
k,df,Ldf,ax = start('105')

# %% [code]
p

# %% [code]
df1,Ldf1,ax,p = ut.check_yz(df,Ldf)

# %% [code]
p

# %% [code]
p1=[.0167,-3.0]

# %% [code]
p1

# %% [code]
df2,Ldf2,ax,p1 = ut.check_yz(df1,Ldf1)

# %% [code]
model = save_model(k,mm,model)
save_both(k,df2,Ldf2[0])

# %% [code]
k,df,Ldf,ax = start('106')

# %% [raw]


# %% [code]
model = save_model(k,mm,model)
save_both(k,df,Ldf[0])

# %% [code]
k,df,Ldf,ax = start('107')
save_both(k,df3,Ldf3[0])

# %% [code]
model = save_model(k,mm,model)
save_both(k,df,Ldf[0])

# %% [code]
k,df,Ldf,ax = start('108')

# %% [code]
df1,Ldf1,pts,ax = ut.check_pts(df,Ldf,'z')

# %% [code]
pts.iloc[0].x = 50
pts.iloc[1].z = 38

# %% [code]
pts

# %% [code]
pts.iloc[0].x = -50

# %% [code]


# %% [code]
ax[0,1].scatter(pts.x,pts.z,c='y')

# %% [code]

df2,Ldf2,ax = ut.revise_pts(df,Ldf,'z',pts=pts)

# %% [code]
df3,Ldf3,mm,ax = ut.ch_vertex(df2,Ldf2,pts=pts)

# %% [code]
df3,Ldf3,mm,ax = ut.ch_vertex(df2,Ldf2,pts=pts)

# %% [code]
pts = pick_pts(-55,40,-8,-2,50,35) #these are essentially random numbers for example


# %% [code]

df3,Ldf3,mm,ax = ut.ch_vertex(df2,Ldf2,pts=pts)

# %% [code]
model = save_model(k,mm,model)
save_both(k,df3,Ldf3[0])

# %% [raw]
k,df,Ldf,ax = start('109')

# %% [raw]
df1,Ldf1,pts,ax = ut.check_pts(df,Ldf,'z')

# %% [code]
df2,Ldf2,mm,ax = ut.ch_vertex(df1,Ldf1)

# %% [code]
pts = pick_pts(-53,40,-9,0,60,40)

# %% [raw]
df3,Ldf3,mm,ax = ut.ch_vertex(df2,Ldf2,pts=pts)

# %% [code]
model = save_model(k,mm,model)
save_both(k,df3,Ldf3[0])

# %% [raw]
k,df,Ldf,ax= start('110')

# %% [code]
model = save_model(k,mm,model)
save_both(k,df,Ldf[0])

# %% [raw]
k,df,Ldf,ax = start('111')

# %% [raw]
df1,Ldf1,pts,ax = ut.check_pts(df,Ldf,'z')

# %% [raw]
df2,Ldf2,mm,ax = ut.ch_vertex(df1,Ldf1)

# %% [code]
model = save_model(k,mm,model)
save_both(k,df2,Ldf2[0])

# %% [raw]
k,df,Ldf,ax = start('112')

# %% [raw]
df1,Ldf1,pts,ax = ut.check_pts(df,Ldf,'z')

# %% [raw]
df2,Ldf2,mm,ax = ut.ch_vertex(df1,Ldf1)

# %% [code]
model = save_model(k,mm,model)
save_both(k,df2,Ldf2[0])

# %% [raw]
k,df,Ldf,ax = start('113')

# %% [raw]
df1,Ldf1,pts,ax = ut.check_pts(df,Ldf,'z')

# %% [raw]
df2,Ldf2,mm,ax = ut.ch_vertex(df1,Ldf1)

# %% [code]
model = save_model(k,mm,model)
save_both(k,df2,Ldf2[0])

# %% [raw]
k,df,Ldf,ax = start('114')

# %% [markdown]
sample rejected for being truncated at midline. likely improper collection that was not superficial enough to capture POC.

# %% [raw]
k,df,Ldf,ax = start('115')

# %% [code]
model = save_model(k,mm,model)
save_both(k,df,Ldf[0])

# %% [raw]
k,df,Ldf,ax = start('116')

# %% [raw]
df1,Ldf1,ax,p = ut.check_yz(df,Ldf)

# %% [code]
model = save_model(k,mm,model)
save_both(k,df1,Ldf1[0])

# %% [raw]
k,df,Ldf,ax = start('117')

# %% [code]
model = save_model(k,mm,model)
save_both(k,df1,Ldf1[0])

# %% [raw]
k,df,Ldf,ax = start('118')

# %% [raw]
df1,Ldf1,ax,p = ut.check_yz(df,Ldf)

# %% [raw]
df2,Ldf2,pts,ax = ut.check_pts(df1,Ldf1,'z')

# %% [raw]
df3,Ldf3,mm,ax = ut.ch_vertex(df2,Ldf2)

# %% [code]
model = save_model(k,mm,model)
save_both(k,df3,Ldf3[0])

# %% [raw]
k,df,Ldf,ax =start('119')

# %% [raw]
df1,Ldf1,pts,ax = ut.check_pts(df,Ldf,'z')

# %% [raw]
df2,Ldf2,mm,ax = ut.ch_vertex(df1,Ldf1)

# %% [code]
model = save_model(k,mm,model)
save_both(k,df2,Ldf2[0])

# %% [code]
klist


# %% [raw]
k,df,Ldf,ax = start('120')

# %% [code]
df1,Ldf1,pts,ax = ut.check_pts(df,Ldf,'z')

# %% [raw]
df2,Ldf2,mm,ax = ut.ch_vertex(df1,Ldf1)

# %% [code]
model = save_model(k,mm,model)
save_both(k,df2,Ldf2[0])

# %% [raw]
k,df,Ldf,ax = start('121')

# %% [markdown]
Commissure is obviously physically deformed due to pressure from coverslip and will not take parabolic curve well. skipping sample.

# %% [code]
k,df,Ldf,ax = start('122')

# %% [code]
model = save_model(k,mm,model)
save_both(k,df,Ldf[0])

# %% [code]
k,df,Ldf,ax = start('123')

# %% [code]
model = save_model(k,mm,model)
save_both(k,df,Ldf[0])

# %% [code]
k,df,Ldf,ax = start('124')

# %% [code]
model = save_model(k,mm,model)
save_both(k,df,Ldf[0])

# %% [code]
k,df,Ldf,ax = start('125')

# %% [code]
df1,Ldf1,pts,ax = ut.check_pts(df,Ldf,'z')

# %% [code]
df2,Ldf2,mm,ax = ut.ch_vertex(df1,Ldf1)

# %% [code]
model = save_model(k,mm,model)
save_both(k,df2,Ldf2[0])

# %% [code]
k,df,Ldf,ax = start('126')

# %% [code]
df1,Ldf1,ax,p = ut.check_yz(df,Ldf)

# %% [code]
p

# %% [code]
p1 = [-5/20, 2]

# %% [code]
df1,Ldf1,ax,p1 = ut.check_yz(df,Ldf)

# %% [code]
df1,Ldf1,ax,p = ut.check_yz(df,Ldf,mm=p1)

# %% [code]
p1

# %% [code]
p2 =[-.5, 2]

# %% [code]
df1,Ldf1,ax,p = ut.check_yz(df,Ldf,mm=p2)

# %% [raw]
k,df,Ldf,ax=start('128')

# %% [raw]
df,Ldf,mm,ax = ut.ch_vertex(df,Ldf)

# %% [code]
model = save_model(k,mm,model)
save_both(k,df,Ldf[0])

# %% [code]
k,df,Ldf,ax=start('129')

# %% [code]
df,Ldf,mm,ax = ut.ch_vertex(df,Ldf)
model = save_model(k,mm,model)
save_both(k,df,Ldf[0])

# %% [code]
k,df,Ldf,ax=start('130')

# %% [code]
df,Ldf,mm,ax = ut.ch_vertex(df,Ldf)

# %% [code]
model = save_model(k,mm,model)
save_both(k,df,Ldf[0])

# %% [code]
k,df,Ldf,ax=start('131')

# %% [code]
df,Ldf,pts,ax = ut.check_pts(df,Ldf,'z')

# %% [code]
df2,Ldf2,mm,ax = ut.ch_vertex(df,Ldf)

# %% [code]
model = save_model(k,mm,model)
save_both(k,df2,Ldf2[0])

# %% [code]
k,df,Ldf,ax=start('128')

# %% [code]
df1,Ldf1,pts,ax = ut.check_pts(df,Ldf,'z')

# %% [code]
df2,Ldf2,mm,ax = ut.ch_vertex(df1,Ldf1)

# %% [code]
model = save_model(k,mm,model)
save_both(k,df2,Ldf2[0])

# %% [code]
model

# %% [code]
k,df,Ldf,ax=start('127')

# %% [code]
df1,Ldf1,pts,ax = ut.check_pts(df,Ldf,'z')

# %% [code]
k,df,Ldf,ax=start('132')

# %% [code]
df1,Ldf1,pts,ax = ut.check_pts(df,Ldf,'z')

# %% [code]
df2,Ldf2,mm,ax = ut.ch_vertex(df1,Ldf1)

# %% [code]
model = save_model(k,mm,model)
save_both(k,df2,Ldf2[0])

# %% [code]
k,df,Ldf,ax=start('133')

# %% [code]
df1,Ldf1,pts,ax = ut.check_pts(df,Ldf,'z')

# %% [code]
df2,Ldf2,mm,ax = ut.ch_vertex(df1,Ldf1)

# %% [code]
model = save_model(k,mm,model)
save_both(k,df2,Ldf2[0])

# %% [code]
k,df,Ldf,ax=start('134')

# %% [code]
df2,Ldf2,mm,ax = ut.ch_vertex(df,Ldf)

# %% [code]
model = save_model(k,mm,model)
save_both(k,df2,Ldf2[0])

# %% [code]
k,df,Ldf,ax=start('135')

# %% [code]
df1,Ldf1,pts,ax = ut.check_pts(df,Ldf,'z')

# %% [code]
df2,Ldf2,mm,ax = ut.ch_vertex(df1,Ldf1)

# %% [code]
df3,Ldf3,ax,p = ut.check_yz(df2,Ldf2)

# %% [code]
model = save_model(k,mm,model)
save_both(k,df2,Ldf2[0])

# %% [code]


# %% [markdown]
# Wrapping up: after all samples are processed
Once all of the data has been processed, we want to save the model data we collected to a file for future reference.

# %% [code]


# %% [code]
model.to_csv(os.path.join(outdir,'model.csv'))

# %% [markdown]
Additionally, it can be helpful to export a html or pdf version of the notebook that preserves all of the plots generated during the alignment process. To do so, use the Jupyter Lab menu interface: File > Export Notebook As > HTML or PDF.

# %% [code]
klist

# %% [code]
k,df,Ldf,ax = start('136')

# %% [code]
df2,Ldf2,mm,ax = ut.ch_vertex(df1,Ldf1)

# %% [code]
model = save_model(k,mm,model)
save_both(k,df2,Ldf2[0])

# %% [code]
k,df,Ldf,ax = start('137')

# %% [code]
df1,Ldf1,mm,ax = ut.ch_vertex(df,Ldf)

# %% [code]
model = save_model(k,mm,model)
save_both(k,df1,Ldf1[0])

# %% [code]
k,df,Ldf,ax = start('138')

# %% [code]
df1,Ldf1,mm,ax = ut.ch_vertex(df,Ldf)

# %% [code]
model = save_model(k,mm,model)
save_both(k,df1,Ldf1[0])

# %% [code]
k,df,Ldf,ax = start('139')

# %% [code]
df1,Ldf1,mm,ax = ut.ch_vertex(df,Ldf)

# %% [code]
df1,Ldf1,pts,ax = ut.check_pts(df,Ldf,'z')

# %% [code]
df2,Ldf2,mm,ax = ut.ch_vertex(df1,Ldf1)

# %% [code]
model = save_model(k,mm,model)
save_both(k,df2,Ldf2[0])

# %% [code]
k,df,Ldf,ax = start('140')

# %% [markdown]
sample highly disturbed in imaging structure, possibly due to tear on left of the image causing loss of data, Vertex will be highly questionable.

# %% [code]
k,df,Ldf,ax = start('141')

# %% [code]
df1,Ldf1,mm,ax = ut.ch_vertex(df,Ldf)

# %% [code]
model = save_model(k,mm,model)
save_both(k,df1,Ldf1[0])

# %% [code]
k,df,Ldf,ax = start('142')

# %% [code]
df1,Ldf1,mm,ax = ut.ch_vertex(df,Ldf)

# %% [code]
model = save_model(k,mm,model)
save_both(k,df1,Ldf1[0])

# %% [code]
k,df,Ldf,ax = start('143')

# %% [code]
df1,Ldf1,mm,ax = ut.ch_vertex(df,Ldf)

# %% [code]
model = save_model(k,mm,model)
save_both(k,df1,Ldf1[0])

# %% [code]
klist

# %% [code]
k,df,Ldf,ax = start('105')

# %% [code]
df1,Ldf1,mm,ax = ut.ch_vertex(df,Ldf)

# %% [code]
model = save_model(k,mm,model)
save_both(k,df1,Ldf1[0])

# %% [code]
k,df,Ldf,ax = start('106')

# %% [code]
df1,Ldf1,mm,ax = ut.ch_vertex(df,Ldf)

# %% [code]
model = save_model(k,mm,model)
save_both(k,df1,Ldf1[0])

# %% [code]
k,df,Ldf,ax = start('107')

# %% [code]
df1,Ldf1,mm,ax = ut.ch_vertex(df,Ldf)

# %% [code]
model = save_model(k,mm,model)
save_both(k,df1,Ldf1[0])

# %% [code]
klist

# %% [code]
k,df,Ldf,ax = start('110')

# %% [code]
df1,Ldf1,mm,ax = ut.ch_vertex(df,Ldf)

# %% [code]
model = save_model(k,mm,model)
save_both(k,df1,Ldf1[0])

# %% [code]
k,df,Ldf,ax = start('115')

# %% [code]
df1,Ldf1,mm,ax = ut.ch_vertex(df,Ldf)

# %% [code]
model = save_model(k,mm,model)
save_both(k,df1,Ldf1[0])

# %% [code]
klist

# %% [code]
k,df,Ldf,ax = start('116')

# %% [code]
df1,Ldf1,mm,ax = ut.ch_vertex(df,Ldf)

# %% [code]
model = save_model(k,mm,model)
save_both(k,df1,Ldf1[0])

# %% [code]
k,df,Ldf,ax = start('117')

# %% [code]
df1,Ldf1,mm,ax = ut.ch_vertex(df,Ldf)

# %% [code]
model = save_model(k,mm,model)
save_both(k,df1,Ldf1[0])

# %% [code]
k,df,Ldf,ax = start('122')

# %% [code]
df1,Ldf1,mm,ax = ut.ch_vertex(df,Ldf)

# %% [code]
model = save_model(k,mm,model)
save_both(k,df1,Ldf1[0])

# %% [code]
k,df,Ldf,ax = start('123')

# %% [code]
df1,Ldf1,mm,ax = ut.ch_vertex(df,Ldf)

# %% [code]
model = save_model(k,mm,model)
save_both(k,df1,Ldf1[0])

# %% [code]
k,df,Ldf,ax = start('124')

# %% [code]
df1,Ldf1,mm,ax = ut.ch_vertex(df,Ldf)

# %% [code]
model = save_model(k,mm,model)
save_both(k,df1,Ldf1[0])

# %% [code]
k,df,Ldf,ax = start('126')

# %% [code]
df1,Ldf1,mm,ax = ut.ch_vertex(df,Ldf)

# %% [code]
model = save_model(k,mm,model)
save_both(k,df1,Ldf1[0])

# %% [code]
model.to_csv(os.path.join(outdir,'models.csv'))
```
