## Supplementary material for "ΔSCOPE: A new method to quantify 3D biological structures and identify differences in zebrafish forebrain development": An archived copy of the DeltaSCOPE code repository.: alignment-yot.ipynb

```
# %% [markdown]
# Introduction
This notebook facilitates the manual curation of sample alignment.

# %% [code]
import deltascope.alignment as ut

import numpy as np
import pandas as pd
import matplotlib.pyplot as plt
import h5py

import os
import re
import time
import tqdm

# %% [markdown]
# Setup

# %% [markdown]
## Parameters

# %% [code]
# --------------------------------
# -------- User input ------------
# --------------------------------

param = {
    'gthresh':0.5,
    'scale':[1,1,1],
    'microns':[0.16,0.16,0.21],
    'mthresh':0.5,
    'radius':10,
    'comp_order':[0,2,1],
    'fit_dim':['x','z'],
    'deg':2,
    
    # Don't forget to modify this with a specific sample name
    'expname':'yot'
}

# %% [markdown]
## Directories

# %% [code]
# --------------------------------
# -------- User input ------------
# --------------------------------

# Specify file paths to directories containing probability files
# after processing by ilastik
gfap = '.\data\ZRF1 You-Too HZ H5 files\Prob_7-29'
at = os.path.abspath('.\data\AT You-Too HZ Mutant H5 files\Prob_7-26')

# Specify root directory where output will be saved
root = '.\data'

root

# %% [code]
# Output directory with timestamp
outname = 'Output_'+param['expname']+time.strftime("%m-%d-%H-%M",
                                 time.localtime())

# Create output directory
outdir = os.path.join(root,outname)
os.mkdir(outdir)

# %% [markdown]
## Extract list of files

# %% [code]
Dat = {}
for f in os.listdir(at):
    if 'h5' in f:
        num  = re.findall(r'\d+',f.split('.')[0])[-1]
        Dat[num] = os.path.join(at,f)

Dat

# %% [code]
Dzrf = {}
for f in os.listdir(gfap):
    if 'h5' in f:
        num  = re.findall(r'\d+',f.split('.')[0])[-1]
        Dzrf[num] = os.path.join(gfap,f)

# %% [code]
# Extract list of filename keys
klist = list(Dat.keys())

# %% [code]
# Create dictionaries to contain the deltascope brain object for each sample
Dbat = {}
Dbzrf = {}

# %% [markdown]
# PCA: Import raw data and perform preprocessing

# %% [markdown]
You-too processing fo samples 19 21, and 172, with radius of 5, vs 10, to deal with the loss of all data in the array caused by radius 10 median filter causing the small sparse axon crossing to be comletely lost to the filter.

# %% [code]
%%time
for k in tqdm.tqdm(klist):
    if k not in list(Dbat.keys()):
        if k in ['19','21','172']:
            param['radius'] = 5
        else:
            param['radius'] = 10
            
        try:
            Dbat[k] = ut.preprocess(Dat[k],param)
            Dbzrf[k] = ut.preprocess(Dzrf[k],param,pca=Dbat[k].pcamed,
                                 mm=Dbat[k].mm,vertex=Dbat[k].vertex)
        except:
            print(k,'caused an error')
    else:
        print(k,'already processed')

# %% [markdown]
# Define experiment specific functions

# %% [code]
''' Define wrapper functions for starting and saving to minimize the number 
of inputs that the user needs to type for each call of the function.'''
def start(k):
    return(ut.start(k,Dbat,[Dbzrf],im=True))
def save_both(k,dfa,dfb):
    ut.save_both(k,dfa,dfb,outdir,param['expname'])

'''Save model parameters for each file to a dataframe that can be 
exported for later reference.'''
model = pd.DataFrame({'a':[],'b':[],'c':[]})
def save_model(k,mm,model):
    row = pd.Series({'a':mm[0],'b':mm[1],'c':mm[2]},name=k)
    model = model.append(row)
    return(model)

'''Define a function that can both fit a model and plot it on an existing plot'''
def fit_model(axi,df,mm=None):
    if mm == None:
        mm = np.polyfit(df.x,df.z,2)
    p = np.poly1d(mm)
    xrange = np.arange(np.min(df.x),np.max(df.x))
    axi.plot(xrange,p(xrange),c='m')
    return(mm)

'''Take a set of points and transform to a dataframe format for ease of access.'''
def pick_pts(x1,z1,vx,vz,x2,z2):
    pts = pd.DataFrame({'x':[x1,vx,x2],'z':[z1,vz,z2]})
    return(pts)

# %% [markdown]
You-too processing fo samples 19 21, and 172, with radius of 5, vs 20, to deal with the loss of all data in the array caused by radius 20 median filter causing the small sparse axon crossing to be comletely lost to the filter.

# %% [code]
k,df,Ldf,ax = start('19')

# %% [code]
df1,Ldf1,pts,ax = ut.check_pts(df,Ldf,'z')

# %% [code]

pts.iloc[0].x=40

# %% [code]
pts.iloc[0].z=15

# %% [code]
df1,Ldf1,ax = ut.revise_pts(df,Ldf,'z',pts=pts)

# %% [code]
df2,Ldf2,mm,ax = ut.ch_vertex(df1,Ldf1)

# %% [code]
model = save_model(k,mm,model)
save_both(k,df2,Ldf2[0])

# %% [code]
k,df,Ldf,ax = start('21')

# %% [code]
df1,Ldf1,pts,ax = ut.check_pts(df,Ldf,'z')

# %% [code]
df2,Ldf2,mm,ax = ut.ch_vertex(df1,Ldf1)

# %% [code]
model = save_model(k,mm,model)
save_both(k,df2,Ldf2[0])

# %% [code]
k,df,Ldf,ax = start('172')

# %% [code]
df1,Ldf1,mm,ax = ut.ch_vertex(df,Ldf)

# %% [code]
model = save_model(k,mm,model)
save_both(k,df1,Ldf1[0])

# %% [markdown]
Processing of remaining you-too samples with 10 median filter

# %% [code]
klist

# %% [code]
k,df,Ldf,ax = start('01')

# %% [code]
df1,Ldf1,mm,ax = ut.ch_vertex(df,Ldf)

# %% [code]
model = save_model(k,mm,model)
save_both(k,df1,Ldf1[0])

# %% [code]
klist

# %% [code]
k,df,Ldf,ax = start('02')

# %% [code]
df1,Ldf1,mm,ax = ut.ch_vertex(df,Ldf)

# %% [code]
model = save_model(k,mm,model)
save_both(k,df1,Ldf1[0])

# %% [code]
k,df,Ldf,ax = start('03')

# %% [code]
df1,Ldf1,mm,ax = ut.ch_vertex(df,Ldf)

# %% [code]
model = save_model(k,mm,model)
save_both(k,df1,Ldf1[0])

# %% [code]
k,df,Ldf,ax = start('04')

# %% [code]
df1,Ldf1,ax,p = ut.check_yz(df,Ldf)

# %% [code]
pts = pick_pts(-60,8,-18,-10,25,8)

# %% [code]
df2,Ldf2,mm,ax = ut.ch_vertex(df1,Ldf1,pts=pts)

# %% [code]
model = save_model(k,mm,model)
save_both(k,df2,Ldf2[0])

# %% [code]
klist

# %% [code]
k,df,Ldf,ax = start('06')

# %% [code]
df1,Ldf1,ax,p = ut.check_yz(df,Ldf)

# %% [code]
df2,Ldf2,mm,ax = ut.ch_vertex(df1,Ldf1)

# %% [code]
model = save_model(k,mm,model)
save_both(k,df2,Ldf2[0])

# %% [code]
klist

# %% [code]
k,df,Ldf,ax = start('112')

# %% [code]
df1,Ldf1,ax,p = ut.check_yz(df,Ldf)

# %% [code]
df2,Ldf2,mm,ax = ut.ch_vertex(df1,Ldf1)

# %% [code]
model = save_model(k,mm,model)
save_both(k,df2,Ldf2[0])

# %% [code]
k,df,Ldf,ax = start('24')

# %% [code]
df2,Ldf2,ax,p = ut.check_yz(df1,Ldf1)

# %% [code]
df1,Ldf1,pts,ax = ut.check_pts(df,Ldf,'z')

# %% [code]
df3,Ldf3,mm,ax = ut.ch_vertex(df2,Ldf2)

# %% [code]
model = save_model(k,mm,model)
save_both(k,df3,Ldf3[0])

# %% [code]
klist

# %% [code]
k,df,Ldf,ax = start('152')

# %% [code]
df1,Ldf1,ax,p = ut.check_yz(df,Ldf)

# %% [code]
df2,Ldf2,mm,ax = ut.ch_vertex(df1,Ldf1)

# %% [code]
model = save_model(k,mm,model)
save_both(k,df2,Ldf2[0])

# %% [code]
klist

# %% [code]
k,df,Ldf,ax = start('317')

# %% [code]
df1,Ldf1,ax,p = ut.check_yz(df,Ldf)

# %% [code]
p1=1,0

# %% [code]
p1

# %% [code]
df1,Ldf1,ax,p1= ut.check_yz(df,Ldf, mm=p1)

# %% [code]
pts.iloc[1].x=30

# %% [code]
pts.iloc[1].z=9

# %% [code]
df2,Ldf2,ax = ut.revise_pts(df1,Ldf1,'z',pts=pts)

# %% [code]
pts = pick_pts(-45,9,-8,-5,30,9)

# %% [code]
df3,Ldf3,mm,ax = ut.ch_vertex(df2,Ldf2,pts=pts)

# %% [code]
p1=-.3,0

# %% [code]
df1,Ldf1,ax,p1 = ut.check_yz(df,Ldf,mm=p1)

# %% [code]
model = save_model(k,mm,model)
save_both(k,df3,Ldf3[0])

# %% [code]
klist

# %% [code]
k,df,Ldf,ax = start('318')

# %% [code]
p1=.3,0

# %% [code]
df1,Ldf1,ax,p1 = ut.check_yz(df,Ldf,mm=p1)

# %% [code]
df2,Ldf2,pts,ax = ut.check_pts(df1,Ldf1,'z')

# %% [code]
pts.iloc[1].x = -40

# %% [code]
pts.iloc[1].z=30

# %% [code]
pts.iloc[0].x

# %% [code]
pts.iloc[0].z=25

# %% [code]
df2,Ldf2,ax = ut.revise_pts(df1,Ldf1,'z',pts=pts)

# %% [code]
df3,Ldf3,mm,ax = ut.ch_vertex(df2,Ldf2)

# %% [code]
pts = pick_pts(-42,30,-5,-5,41,30)

# %% [code]
df3,Ldf3,mm,ax = ut.ch_vertex(df2,Ldf2,pts=pts)

# %% [code]
model = save_model(k,mm,model)
save_both(k,df3,Ldf3[0])

# %% [code]
klist

# %% [code]
k,df,Ldf,ax = start('319')

# %% [code]
p1=-.4,-5

# %% [code]
df1,Ldf1,ax,p1 =ut.check_yz(df,Ldf,mm=p1)

# %% [code]
pts = pick_pts(-35,22,0,-2,35,22)

# %% [code]
df2,Ldf2,mm,ax = ut.ch_vertex(df1,Ldf1,pts=pts)

# %% [code]
model = save_model(k,mm,model)
save_both(k,df2,Ldf2[0])

# %% [code]
klist

# %% [code]
k,df,Ldf,ax = start('320')

# %% [code]
p1=-.5,-10

# %% [code]
df1,Ldf1,ax,p1 =ut.check_yz(df,Ldf,mm=p1)

# %% [code]
pts = pick_pts(-38,28,0,-10,45,28)

# %% [code]
df2,Ldf2,mm,ax = ut.ch_vertex(df1,Ldf1,pts=pts)

# %% [code]
model = save_model(k,mm,model)
save_both(k,df2,Ldf2[0])

# %% [code]
klist

# %% [code]
k,df,Ldf,ax= start('321') 

# %% [code]
df1,Ldf1,mm,ax = ut.ch_vertex(df,Ldf)

# %% [code]
model = save_model(k,mm,model)
save_both(k,df1,Ldf1[0])

# %% [code]
k,df,Ldf,ax = start('323')

# %% [code]
df1,Ldf1,mm,ax = ut.ch_vertex(df,Ldf)

# %% [code]
model = save_model(k,mm,model)
save_both(k,df1,Ldf1[0])

# %% [code]
k,df,Ldf,ax = start('324')

# %% [code]
p1=.4,10

# %% [code]
df1,Ldf1,ax,p1 =ut.check_yz(df,Ldf,mm=p1)

# %% [code]
df2,Ldf2,mm,ax = ut.ch_vertex(df1,Ldf1)

# %% [code]
pts = pick_pts(-33,18,0,-4,33,18) 

# %% [code]
df3,Ldf3,mm,ax = ut.ch_vertex(df1,Ldf1,pts=pts)

# %% [code]
model = save_model(k,mm,model)
save_both(k,df3,Ldf3[0])

# %% [code]
k,df,Ldf,ax = start('325')

# %% [code]
p1=.5,10

# %% [code]
df1,Ldf1,ax,p1 =ut.check_yz(df,Ldf,mm=p1)

# %% [code]
pts = pick_pts(-30,12,0,-8,34,12) 

# %% [code]
df3,Ldf3,mm,ax = ut.ch_vertex(df1,Ldf1,pts=pts)

# %% [code]
model = save_model(k,mm,model)
save_both(k,df3,Ldf3[0])

# %% [code]
klist

# %% [code]
k,df,Ldf,ax = start('326')

# %% [code]
pts = pick_pts(-35,20,-7,-7,38,20) 

# %% [code]
df3,Ldf3,mm,ax = ut.ch_vertex(df1,Ldf1,pts=pts)

# %% [code]
model = save_model(k,mm,model)
save_both(k,df3,Ldf3[0])

# %% [code]
k,df,Ldf,ax = start('327')

# %% [code]
pts = pick_pts(-37,35,-2,-5,34,33) 

# %% [code]
df3,Ldf3,mm,ax = ut.ch_vertex(df,Ldf,pts=pts)

# %% [code]
model = save_model(k,mm,model)
save_both(k,df3,Ldf3[0])

# %% [code]
k,df,Ldf,ax= start('328')

# %% [code]
pts = pick_pts(-37,25,-8,-6,36,25) 

# %% [code]
df3,Ldf3,mm,ax = ut.ch_vertex(df,Ldf,pts=pts)

# %% [code]
model = save_model(k,mm,model)
save_both(k,df3,Ldf3[0])

# %% [code]
k,df,Ldf,ax = start('329')

# %% [code]
p1=.4,0

# %% [code]
df1,Ldf1,ax,p1 =ut.check_yz(df,Ldf,mm=p1)

# %% [code]
pts = pick_pts(-43,40,-4,-6,41,40) 

# %% [code]
df3,Ldf3,mm,ax = ut.ch_vertex(df1,Ldf1,pts=pts)

# %% [code]
model = save_model(k,mm,model)
save_both(k,df3,Ldf3[0])

# %% [code]
k,df,Ldf,ax = start('330')

# %% [code]
df2,Ldf2,mm,ax = ut.ch_vertex(df,Ldf)

# %% [code]
model = save_model(k,mm,model)
save_both(k,df2,Ldf2[0])

# %% [code]
k,df,Ldf,ax = start('331')

# %% [code]
pts.iloc[0].z=33

# %% [code]
df2,Ldf2,ax = ut.revise_pts(df,Ldf,'z',pts=pts)

# %% [code]
pts = pick_pts(-45,30,-12,-10,38,30) 

# %% [code]
df3,Ldf3,mm,ax = ut.ch_vertex(df2,Ldf2,pts=pts)

# %% [code]
model = save_model(k,mm,model)
save_both(k,df3,Ldf3[0])

# %% [code]
k,df,Ldf,ax = start('332')

# %% [code]
p1=1.9,5

# %% [code]
df1,Ldf1,ax,p1 =ut.check_yz(df,Ldf,mm=p1)

# %% [code]
pts = pick_pts(-35,20,0,-9,40,20)

# %% [code]
df3,Ldf3,mm,ax = ut.ch_vertex(df1,Ldf1,pts=pts)

# %% [code]
model = save_model(k,mm,model)
save_both(k,df3,Ldf3[0])

# %% [code]
k,df,Ldf,ax= start('333')

# %% [code]
df2,Ldf2,mm,ax = ut.ch_vertex(df,Ldf)

# %% [code]
model = save_model(k,mm,model)
save_both(k,df2,Ldf2[0])

# %% [code]
k,df,Ldf,ax = start('334')

# %% [code]
p1=1.5,0

# %% [code]
df1,Ldf1,ax,p1 =ut.check_yz(df,Ldf,mm=p1)

# %% [code]
df2,Ldf2,mm,ax = ut.ch_vertex(df1,Ldf1)

# %% [code]
model = save_model(k,mm,model)
save_both(k,df2,Ldf2[0])

# %% [code]
k,df,Ldf,ax = start('335')

# %% [code]
p1=1.1,0

# %% [code]
df1,Ldf1,ax,p1 =ut.check_yz(df,Ldf,mm=p1)

# %% [code]
df2,Ldf2,mm,ax = ut.ch_vertex(df1,Ldf1)

# %% [code]
model = save_model(k,mm,model)
save_both(k,df2,Ldf2[0])

# %% [code]
k,df,Ldf,ax = start('336')

# %% [code]
pts = pick_pts(-47,30,0,-10,47,30)

# %% [code]
df3,Ldf3,mm,ax = ut.ch_vertex(df,Ldf,pts=pts)

# %% [code]
model = save_model(k,mm,model)
save_both(k,df3,Ldf3[0])

# %% [code]
k,df,Ldf,ax= start('337')

# %% [code]


# %% [code]
p1=2,0

# %% [code]
df1,Ldf1,ax,p1 =ut.check_yz(df,Ldf,mm=p1)

# %% [code]
df2,Ldf2,mm,ax = ut.ch_vertex(df1,Ldf1)

# %% [code]
model = save_model(k,mm,model)
save_both(k,df2,Ldf2[0])

# %% [code]
k,df,Ldf,ax = start('338')

# %% [code]
p1=1.3,0

# %% [code]
df1,Ldf1,ax,p1 =ut.check_yz(df,Ldf,mm=p1)

# %% [code]
df2,Ldf2,mm,ax = ut.ch_vertex(df1,Ldf1)

# %% [code]
model = save_model(k,mm,model)
save_both(k,df2,Ldf2[0])

# %% [code]
k,df,Ldf,ax = start('339')

# %% [code]
p1=2.5,0

# %% [code]
df1,Ldf1,ax,p1 =ut.check_yz(df,Ldf,mm=p1)

# %% [code]
df2,Ldf2,mm,ax = ut.ch_vertex(df1,Ldf1)

# %% [code]
model = save_model(k,mm,model)
save_both(k,df2,Ldf2[0])

# %% [code]
k,df,Ldf,ax = start('340')

# %% [code]
p1=-.6,0

# %% [code]
df1,Ldf1,ax,p1 =ut.check_yz(df,Ldf,mm=p1)

# %% [code]
pts = pick_pts(-42,25,0,-7,45,25) 

# %% [code]
df3,Ldf3,mm,ax = ut.ch_vertex(df1,Ldf1,pts=pts)

# %% [code]
model = save_model(k,mm,model)
save_both(k,df3,Ldf3[0])

# %% [code]
model

# %% [code]
model.to_csv(os.path.join(outdir,'model.csv'))

# %% [code]
import re

# %% [code]
f = 'AT_4_wt.psi'

# %% [code]
re.findall(r'\d+',f.split('.')[0])[0]
```
