## Supplementary material for "ΔSCOPE: A new method to quantify 3D biological structures and identify differences in zebrafish forebrain development": An archived copy of the DeltaSCOPE code repository.: plotting.html


In [2]:

```
import os
import json
import time
import datetime

import tqdm
import glob
from imp import reload

import pandas as pd
import numpy as np
import matplotlib.pyplot as plt
import scipy.stats as stats
from sklearn.preprocessing import normalize
from scipy.optimize import minimize

import deltascope as ds
```

In [3]:

```
# --------------------------------
# -------- User input ------------
# --------------------------------

# Specify path to exported landmark data
lmpath = glob.glob('03*landmarks.csv')[0]
binpath = glob.glob('03*landmarks_bins.json')[0]
lmpath,binpath
```

Out[3]:

```
('03-10-03-20_landmarks.csv', '03-10-03-20_landmarks_bins.json')
```

In [4]:

```
# Load landmarks from csv
oldlm = pd.read_csv(lmpath)
```

In [5]:

```
oldlm.head()
```

Out[5]:

|  | Unnamed: 0 | -13.61\_-6.8\_-0.79\_0.0\_50\_pts | -13.61\_-6.8\_-0.79\_0.0\_50\_r | -13.61\_-6.8\_-1.57\_-0.79\_50\_pts | -13.61\_-6.8\_-1.57\_-0.79\_50\_r | -13.61\_-6.8\_-2.36\_-1.57\_50\_pts | -13.61\_-6.8\_-2.36\_-1.57\_50\_r | -13.61\_-6.8\_-3.14\_-2.36\_50\_pts | -13.61\_-6.8\_-3.14\_-2.36\_50\_r | -13.61\_-6.8\_0.0\_0.79\_50\_pts | ... | 74.84\_81.64\_-3.14\_-2.36\_50\_r | 74.84\_81.64\_0.0\_0.79\_50\_pts | 74.84\_81.64\_0.0\_0.79\_50\_r | 74.84\_81.64\_0.79\_1.57\_50\_pts | 74.84\_81.64\_0.79\_1.57\_50\_r | 74.84\_81.64\_1.57\_2.36\_50\_pts | 74.84\_81.64\_1.57\_2.36\_50\_r | 74.84\_81.64\_2.36\_3.14\_50\_pts | 74.84\_81.64\_2.36\_3.14\_50\_r | stype |
| --- | --- | --- | --- | --- | --- | --- | --- | --- | --- | --- | --- | --- | --- | --- | --- | --- | --- | --- | --- | --- | --- |
| 0 | 101 | 0.0 | NaN | 0.0 | NaN | 0.0 | NaN | 0.0 | NaN | 0.0 | ... | NaN | 0.0 | NaN | 0.0 | NaN | 0.0 | NaN | 0.0 | NaN | wt-AT |
| 1 | 102 | 12.0 | 0.591255 | 4.0 | 0.356970 | 43.0 | 1.563061 | 202.0 | 1.139003 | 4.0 | ... | NaN | 0.0 | NaN | 0.0 | NaN | 0.0 | NaN | 0.0 | NaN | wt-AT |
| 2 | 104 | 953.0 | 2.394736 | 1491.0 | 3.236693 | 367.0 | 3.908196 | 0.0 | NaN | 318.0 | ... | NaN | 0.0 | NaN | 0.0 | NaN | 0.0 | NaN | 0.0 | NaN | wt-AT |
| 3 | 105 | 153.0 | 3.092621 | 1224.0 | 3.664182 | 143.0 | 3.798781 | 0.0 | NaN | 0.0 | ... | NaN | 0.0 | NaN | 0.0 | NaN | 0.0 | NaN | 0.0 | NaN | wt-AT |
| 4 | 106 | 4.0 | 14.989111 | 33.0 | 3.444925 | 624.0 | 2.077761 | 389.0 | 1.135013 | 0.0 | ... | NaN | 0.0 | NaN | 0.0 | NaN | 0.0 | NaN | 0.0 | NaN | wt-AT |

5 rows × 386 columns

In [6]:

```
# Load landmark bins 
with open(binpath,'r') as f:
    bins = json.load(f)
acbins = bins['acbins']
tbins = bins['tbins']
```

In [7]:

```
colors = ['#41ab5d','#ef3b2c','#00441b','#67000d']
tarr = np.round(tbins,2)
xarr = np.round(acbins,2)
tpairs = [[tarr[0],tarr[4]],[tarr[1],tarr[5]],[tarr[2],tarr[6]],[tarr[3],tarr[7]]]
```

### Restructure Data¶

We will sort landmark data according to stype and organize it in a two tiered dictionary according to sample type (s) and channel (c).

In [8]:

```
oldlm.stype.unique()
```

Out[8]:

```
array(['wt-AT', 'wt-ZRF', 'you-too-AT', 'you-too-ZRF'], dtype=object)
```

In [9]:

```
Dlm = {}
for stype in tqdm.tqdm(oldlm.stype.unique()):
    # These two lines may need to be modified based on stype structure
    s = stype.split('-')[0]
    c = stype.split('-')[-1]
    
    # Add sample type dictionary if not already present
    if s not in Dlm.keys():
        Dlm[s] = {}
    
    # Save sample specific landmark data to dictionary
    Dlm[s][c] = oldlm[oldlm.stype==stype]
```

```
100%|██████████| 4/4 [00:00<00:00, 571.78it/s]
```

#### Set up graph data¶

In [11]:

```
gdata = {}
gdata['you'] = {}
gdata['wt'] = {}

gdata['you']['ZRF'] = ds.graphData(Dlm['you']['ZRF'],colors[3])
gdata['wt']['ZRF'] = ds.graphData(Dlm['wt']['ZRF'],colors[1])
gdata['you']['AT'] = ds.graphData(Dlm['you']['AT'],colors[2])
gdata['wt']['AT'] = ds.graphData(Dlm['wt']['AT'],colors[0])
```

### Graphs¶

In [12]:

```
crop = 40 # microns
legend = False
save = True
a = 0.3
pthresh = 0.01
```

In [35]:

```
channel = 'ZRF'

fig,axr = plt.subplots(2,4,figsize=(12,5),sharey='row',sharex=True)

if crop is not None:
    mask = np.where((xarr>-crop)&(xarr<crop) == True)[0]
    xmin = mask.min()
    xmax = mask.max()
    xarrcr = xarr[xmin:xmax+1]
else:
    xarrcr = xarr

for I,dtype in enumerate(['r','pts']):
    go1 = gdata['wt'][channel]
    go1.prepare_data(xarrcr,tarr,dtype)
    go2 = gdata['you'][channel]
    go2.prepare_data(xarrcr,tarr,dtype)
    
    parr = stats.ttest_ind(go1.arr_masked,go2.arr_masked,axis=2,nan_policy='omit')[1]

    parr[parr<pthresh] = 0
    parr[parr>pthresh] = 1
    
    # Plot wildtype data
    go = go1
    for i,p in enumerate(tpairs):
        n = i
        i = I
        
        ti1 = np.where(tarr==p[0])[0][0]
        ti2 = np.where(tarr==p[1])[0][0]

        axr[i,n].fill_between(xarrcr,go.avg[:,ti1]+go.sem[:,ti1],go.avg[:,ti1]-go.sem[:,ti1],alpha=a,color=go.c,zorder=1)
        axr[i,n].fill_between(xarrcr,-go.avg[:,ti2]+go.sem[:,ti2],-go.avg[:,ti2]-go.sem[:,ti2],alpha=a,color=go.c,zorder=1)

        axr[i,n].plot(xarrcr,go.avg[:,ti1],c=go.c,zorder=2,label='{} {}'.format(go.arr.shape[-1],'wt'))
        axr[i,n].plot(xarrcr,-go.avg[:,ti2],c=go.c,zorder=2)
        
    # Plot mutant data
    go = go2
    for i,p in enumerate(tpairs):
        n = i
        i = I
        
        ti1 = np.where(tarr==p[0])[0][0]
        ti2 = np.where(tarr==p[1])[0][0]

        axr[i,n].fill_between(xarrcr,go.avg[:,ti1]+go.sem[:,ti1],go.avg[:,ti1]-go.sem[:,ti1],alpha=a,color=go.c,zorder=1)
        axr[i,n].fill_between(xarrcr,-go.avg[:,ti2]+go.sem[:,ti2],-go.avg[:,ti2]-go.sem[:,ti2],alpha=a,color=go.c,zorder=1)

        axr[i,n].plot(xarrcr,go.avg[:,ti1],c=go.c,zorder=2,label='{} {}'.format(go.arr.shape[-1],'yot'))
        axr[i,n].plot(xarrcr,-go.avg[:,ti2],c=go.c,zorder=2)
        
        axr[i,n].scatter(xarrcr,go.avg[:,ti1],c=parr[:,ti1],cmap='Greys_r',zorder=3,vmin=0,vmax=1,edgecolor='k')
        axr[i,n].scatter(xarrcr,-go.avg[:,ti2],c=parr[:,ti2],cmap='Greys_r',zorder=3,vmin=0,vmax=1,edgecolor='k')
        
        axr[0,n].legend(bbox_to_anchor=(0., 1.02, 1., .102), loc=3,ncol=2, mode="expand", borderaxespad=0.)

plt.tight_layout()
        
tstamp = datetime.datetime.now().strftime('%Y-%m-%d')

if save:
    fig.savefig(tstamp+'_yot-wt-{}.pdf'.format(channel))
```

```
C:\Users\zfishlab\AppData\Local\Continuum\anaconda3\envs\test\lib\site-packages\ipykernel_launcher.py:21: RuntimeWarning: invalid value encountered in less
C:\Users\zfishlab\AppData\Local\Continuum\anaconda3\envs\test\lib\site-packages\ipykernel_launcher.py:22: RuntimeWarning: invalid value encountered in greater
```

In [34]:

```
channel = 'AT'

fig,axr = plt.subplots(2,4,figsize=(12,5),sharey='row',sharex=True)

if crop is not None:
    mask = np.where((xarr>-crop)&(xarr<crop) == True)[0]
    xmin = mask.min()
    xmax = mask.max()
    xarrcr = xarr[xmin:xmax+1]
else:
    xarrcr = xarr

for I,dtype in enumerate(['r','pts']):
    go1 = gdata['wt'][channel]
    go1.prepare_data(xarrcr,tarr,dtype)
    go2 = gdata['you'][channel]
    go2.prepare_data(xarrcr,tarr,dtype)
    
    parr = stats.ttest_ind(go1.arr_masked,go2.arr_masked,axis=2,nan_policy='omit')[1]

    parr[parr<pthresh] = 0
    parr[parr>pthresh] = 1
    
    # Plot wildtype data
    go = go1
    for i,p in enumerate(tpairs):
        n = i
        i = I
        
        ti1 = np.where(tarr==p[0])[0][0]
        ti2 = np.where(tarr==p[1])[0][0]

        axr[i,n].fill_between(xarrcr,go.avg[:,ti1]+go.sem[:,ti1],go.avg[:,ti1]-go.sem[:,ti1],alpha=a,color=go.c,zorder=1)
        axr[i,n].fill_between(xarrcr,-go.avg[:,ti2]+go.sem[:,ti2],-go.avg[:,ti2]-go.sem[:,ti2],alpha=a,color=go.c,zorder=1)

        axr[i,n].plot(xarrcr,go.avg[:,ti1],c=go.c,zorder=2,label='{} {}'.format(go.arr.shape[-1],'wt'))
        axr[i,n].plot(xarrcr,-go.avg[:,ti2],c=go.c,zorder=2)
        
    # Plot mutant data
    go = go2
    for i,p in enumerate(tpairs):
        n = i
        i = I
        
        ti1 = np.where(tarr==p[0])[0][0]
        ti2 = np.where(tarr==p[1])[0][0]

        axr[i,n].fill_between(xarrcr,go.avg[:,ti1]+go.sem[:,ti1],go.avg[:,ti1]-go.sem[:,ti1],alpha=a,color=go.c,zorder=1)
        axr[i,n].fill_between(xarrcr,-go.avg[:,ti2]+go.sem[:,ti2],-go.avg[:,ti2]-go.sem[:,ti2],alpha=a,color=go.c,zorder=1)

        axr[i,n].plot(xarrcr,go.avg[:,ti1],c=go.c,zorder=2,label='{} {}'.format(go.arr.shape[-1],'yot'))
        axr[i,n].plot(xarrcr,-go.avg[:,ti2],c=go.c,zorder=2)
        
        axr[i,n].scatter(xarrcr,go.avg[:,ti1],c=parr[:,ti1],cmap='Greys_r',zorder=3,vmin=0,vmax=1,edgecolor='k')
        axr[i,n].scatter(xarrcr,-go.avg[:,ti2],c=parr[:,ti2],cmap='Greys_r',zorder=3,vmin=0,vmax=1,edgecolor='k')
        
        axr[0,n].legend(bbox_to_anchor=(0., 1.02, 1., .102), loc=3,ncol=2, mode="expand", borderaxespad=0.)

plt.tight_layout()

tstamp = datetime.datetime.now().strftime('%Y-%m-%d')

if save:
    fig.savefig(tstamp+'_yot-wt-{}.pdf'.format(channel))
```

```
C:\Users\zfishlab\AppData\Local\Continuum\anaconda3\envs\test\lib\site-packages\ipykernel_launcher.py:21: RuntimeWarning: invalid value encountered in less
C:\Users\zfishlab\AppData\Local\Continuum\anaconda3\envs\test\lib\site-packages\ipykernel_launcher.py:22: RuntimeWarning: invalid value encountered in greater
```
