## Supplementary material for "ΔSCOPE: A new method to quantify 3D biological structures and identify differences in zebrafish forebrain development": An archived copy of the DeltaSCOPE code repository.: appClass.py

```
#!/usr/bin/env python

import tkinter as tk
from tkinter import filedialog, messagebox
from tkinter import ttk

import os
import yaml
import copy
import cranium
import numpy as np
import time
import pandas as pd

import matplotlib
matplotlib.use('TkAgg')
from matplotlib.backends.backend_tkagg import FigureCanvasTkAgg, NavigationToolbar2TkAgg
import matplotlib.pyplot as plt
from matplotlib.figure import Figure

class channelMaster:
	def __init__(self,parent,text,n):
		self.name = tk.Entry(parent)
		self.name.grid(row=n,column=1)
		self.name.insert(0,text)
		self.dir = None
		self.dButton = ttk.Button(parent,text='Select Folder',command=self.open_dir)
		self.dButton.grid(row=n,column=2)
		self.files = None

		self.key = n

	def open_dir(self):
		self.dir = filedialog.askdirectory()
		self.dButton['text'] = self.dir

	def return_data_dict(self):
		out = {}
		#out['files'] = self.files
		out['dir'] = self.dir
		out['name'] = self.name
		print(out)

		return(out)

class Sample:
	def __init__(self,key):
		self.key = key
		self.fs = ['','','','']
		self.outpaths = ['','','','']

		self.cbVar = tk.IntVar()
		self.degVar = tk.StringVar()
		self.directVar = tk.StringVar()

	def tab2row(self,tab,row):

		self.checkB = tk.Checkbutton(tab,text=self.key,variable=self.cbVar)
		self.checkB.select()
		self.checkB.grid(row=row,column=0)

		for i,f in enumerate(self.fs):
			ttk.Label(tab,text=f).grid(row=row,column=i+1)

	def tab3row(self,tab,r,app):

		#Sample number label
		tk.Label(tab,text=self.key).grid(row=r,column=0)

		self.plotB = tk.Button(tab,text='PCA 2D',command=lambda:self.first_alignment(app.p,app.Lc,2))
		self.plotB.grid(row=r,column=1)

		self.plotB3 = tk.Button(tab,text='PCA 3D',command=lambda:self.first_alignment(app.p,app.Lc,3))
		self.plotB3.grid(row=r,column=2)

		rotDegrees = ('-180','-90','0','90','180')
		rotDirection = ('X','Y','Z')

		self.degVar.set(rotDegrees[2])
		self.degM = tk.OptionMenu(tab,self.degVar,*rotDegrees)
		self.degM.grid(row=r,column=3)

		self.directVar.set(rotDirection[0])
		self.directM = tk.OptionMenu(tab,self.directVar,*rotDirection)
		self.directM.grid(row=r,column=4)

		self.rotB = tk.Button(tab,text='Rotate',command=self.rotate_data)
		self.rotB.grid(row=r,column=5)

		self.doneVar = tk.IntVar()
		self.doneCb = tk.Checkbutton(tab,variable=self.doneVar)
		self.doneCb.grid(row=r,column=6)

	def first_alignment(self,p,Lc,D):
		tic = time.time()

		for i,c in enumerate(Lc):
			print(i,c.name,c.dir)

		pc = self.check_settings(p)

		e = cranium.embryo(pc['expname'],self.key,pc['outdir'])
		for i,c in enumerate(Lc):
			print(i,c.dir,c.key)
			if c.dir != None:
				print(i,'data processing')
				e.add_channel(os.path.join(c.dir,self.fs[i]),c.key)
				e.chnls[c.key].preprocess_data(pc['genthresh'],[1,1,1],pc['microns'])

				#Processing for the structural channel
				if i==0:
					if D == 3:
						print('comporder',pc['comporder'])
						e.chnls[c.key].calculate_pca_median(e.chnls[c.key].raw_data,pc['medthresh'],
							pc['radius'],pc['microns'])
						pca = e.chnls[c.key].pcamed
						e.chnls[c.key].pca_transform_3d(e.chnls[c.key].df_thresh,pca,
							pc['comporder'],pc['fitdim'],deg=pc['deg'])
					elif D == 2:
						e.chnls[c.key].calculate_pca_median_2d(e.chnls[c.key].raw_data,pc['medthresh'],
							pc['radius'],pc['microns'])
						pca = e.chnls[c.key].pcamed
						e.chnls[c.key].pca_transform_2d(e.chnls[c.key].df_thresh,pca,
							pc['comporder'],pc['fitdim'],deg=pc['deg'])

					mm = e.chnls[c.key].mm
					vertex = e.chnls[c.key].vertex

				#Secondary channel processing
				if D == 3:
					e.chnls[c.key].pca_transform_3d(e.chnls[c.key].df_thresh,pca,
						pc['comporder'],pc['fitdim'],
						deg=pc['deg'],mm=mm,vertex=vertex)
				if D == 2:
					e.chnls[c.key].pca_transform_2d(e.chnls[c.key].df_thresh,pca,
						pc['comporder'],pc['fitdim'],
						deg=pc['deg'],mm=mm,vertex=vertex)

			print('sample alignment complete',time.time()-tic)

		#Save data
		for i,c in enumerate(Lc):
			if c.dir != None:
				print(i,'data saving')
				cname = c.name.get()
				for p in ':,./\\[]{};() ':
					cname = cname.replace(p,'')

				fpath = os.path.join(pc['outdir'],'_'.join([cname,pc['expname'],self.key])+'.psi')
				cranium.write_data(fpath,e.chnls[c.key].df_align)
				self.outpaths[i] = fpath

		self.plot_projections([e.chnls[Lc[0].key].df_thresh, e.chnls[Lc[0].key].df_align],mm)

	def rotate_data(self):
		deg = np.deg2rad(int(self.degVar.get()))
		print(deg)
		dM = {
			'X':np.array([[1,0,0],
					[0,np.cos(deg),-np.sin(deg)],
					[0,np.sin(deg),np.cos(deg)]]),
			'Y':np.array([[np.cos(deg),0,np.sin(deg)],
					[0,1,0],
					[-np.sin(deg),0,np.cos(deg)]]),
			'Z':np.array([[np.cos(deg),-np.sin(deg),0],
					[np.sin(deg),np.cos(deg),0],
					[0,0,1]])
		}
		M = dM[self.directVar.get()]
		print(M)

		for fp in self.outpaths:
			if fp != '':
				try:
					df = cranium.read_psi(fp)
				except:
					messagebox.showerror('Error',fp+' does not exist for rotation.')
					return

				rot = np.dot(np.array(df[['x','y','z']]),M)
				dfr = pd.DataFrame({'x':rot[:,0],'y':rot[:,1],'z':rot[:,2]})

				cranium.write_data(fp,dfr)
				print('rotating',fp)

		self.plot_projections([cranium.read_psi(self.outpaths[0])],None)
		print('plotting done')

	def plot_projections(self,Ldf,mm):
		n = len(Ldf)
		if n == 1: n=2
		fig, axarr = plt.subplots(n,3,num='Sample '+self.key) #(rows, columns)

		for i,df in enumerate(Ldf):
			df = df.sample(frac=0.01)
			axarr[i,0].scatter(df.x,df.z)
			axarr[i,1].scatter(df.x,df.y)
			axarr[i,2].scatter(df.z,df.y)

			##### To plot the model I need to account for which dimensions the model is supposed to be fit in 

			axarr[i,0].set_title('Y projection')
			axarr[i,1].set_title('Z projection')
			axarr[i,2].set_title('X projection')
			axarr[i,0].set_xlabel('X')
			axarr[i,0].set_ylabel('Z')
			axarr[i,1].set_xlabel('X')
			axarr[i,1].set_ylabel('Y')
			axarr[i,2].set_xlabel('Z')
			axarr[i,2].set_ylabel('Y')

		fig.show()

	def check_settings(self,p):

		pc = {}

		#Validate experiment name and outdir
		v = p['expname']['entry'].get()
		if v == '':
			messagebox.showerror('Error','Experiment name is not defined')
			return
		else:
			pc['expname'] = v

		if p['outdir'] == None:
			messagebox.showerror('Error','Output directory has not been selected')
			return
		else:
			pc['outdir'] = p['outdir']

		#Check median threshold value
		s = p['medthresh']['entry'].get()
		try:
			v = float(s)
			if v <= 1 and v >= 0:
				pc['medthresh'] = v
			else:
				messagebox.showerror('Error','Median threshold input must be a number between 0 and 1')
		except ValueError:
			messagebox.showerror('Error','Median threshold input must be a number between 0 and 1')
			return

		#Check radius must be integer
		s = p['radius']['entry'].get()
		try:
			pc['radius'] = int(s)
		except ValueError:
			messagebox.showerror('Error','Radius input must be an integer')
			return

		#Check genthresh float between 0 and 1
		s = p['genthresh']['entry'].get()
		try:
			v = float(s)
			if v <= 1 and v >= 0:
				pc['genthresh'] = v
			else:
				messagebox.showerror('Error','General threshold input must be a number between 0 and 1')
				return
		except ValueError:
			messagebox.showerror('Error','General threshold input must be a number between 0 and 1')
			return

		#Check microns
		lv = []
		for key in ['entry_x','entry_y','entry_z']:
			s = p['microns'][key].get()
			try:
				v = float(s)
				lv.append(v)
			except ValueError:
				messagebox.showerror('Error','Micron inputs must be numeric values')
				return
				
			pc['microns'] = lv

		#Check degree
		s = p['deg']['entry'].get()
		try:
			pc['deg'] = int(s)
		except ValueError:
			messagebox.showerror('Error','Degree input must be an integer')
			return

		#Check component order
		lv = []
		for key in ['entry_x','entry_y','entry_z']:
			s = p['comporder'][key].get()
			try:
				v = int(s)
				lv.append(v)
			except ValueError:
				messagebox.showerror('Error','Component order inputs must be numeric values')
				return
			pc['comporder'] = lv

		#Get fit dimensions
		s = p['fitdim']['entry'].get()
		if s == 'XZ Plane':
			pc['fitdim'] = ['x','z']
		elif s == 'XY Plane':
			pc['fitdim'] = ['x','y']
		elif s == 'YZ Plane':
			pc['fitdim'] = ['y','z']

		for key in pc.keys():
			print(key,pc[key])

		return(pc)
```
