## Supplementary material for "ΔSCOPE: A new method to quantify 3D biological structures and identify differences in zebrafish forebrain development": An archived copy of the DeltaSCOPE code repository.: gui_dev.py

```
from tkinter import Tk, Label, Button, filedialog
from tkinter import ttk


class Gui:

	def __init__(self,master):
		self.master = master
		master.title('simple gui')

		ttk.Style().configure("TNotebook", background='b')
		ttk.Style().map("TNotebook.tab",background=[('selected','r')])
		# Style().configure('TNotebook.Tab')

		note = ttk.Notebook(root)
		self.tab1 = page(self.master,note,'1')
		self.tab1.set_instruction('Select directories with raw data')
		Button(self.tab1.tab,text='Select Channel 1',command=self.open_dir1).grid(row=2,column=1)
		Button(self.tab1.tab,text='Select Channel 2',command=self.open_dir2).grid(row=2,column=2)

		self.tab2 = page(self.master,note,'2')
		note.grid(row=1,columnspan=2)

		Button(self.tab2.tab,text='Identify sample numbers',command=self.calc_num).grid(row=3,columnspan=2)

		# self.greet_button = Button(master,text="greet", command = self.greet)
		# self.greet_button.pack()

		# self.close_button = Button(master, text="close", command=master.quit)
		# self.close_button.pack()

		# self.open_button = Button(master, text='Open', command=self.open_dir)
		# self.open_button.pack()

		# tkvar = StringVar(root)
		# choices = {'a','b','c'}
		# tkvar.set('a')

		# popupMenu = OptionMenu(self.master, tkvar, *choices)
		# popupMenu.pack()

	def open_dir1(self):
		self.c1dir = filedialog.askdirectory()
		Label(self.tab2.tab,text='Channel 1:').grid(row=1,column=1)
		Label(self.tab2.tab,text=self.c1dir).grid(row=2,column=1)

	def open_dir2(self):
		self.c2dir = filedialog.askdirectory()
		Label(self.tab2.tab,text='Channel 2:').grid(row=1,column=2)
		Label(self.tab2.tab,text=self.c2dir).grid(row=2,column=2)

	def calc_num(self):
		files = os.listdir(self.c1dir)
		self.s_num = []

		for f in files:
			if 'h5' in f and 'Probabilities' in f:
				self.s_num.append(f.split('_')[1])
				


class page():

	def __init__(self,master,note,label):
		self.master = master
		self.note = note

		self.tab = ttk.Frame(note)
		self.note.add(self.tab,text=label)

	def set_instruction(self,text):

		self.instr = Label(self.tab,text=text)
		self.instr.grid(row=1)

	def activate(self):
		self.note.tab(self.tab,state='normal')

	def deactivate(self):
		self.note.tab(self.tab,state='disabled')

root = Tk()
my_gui = Gui(root)
root.mainloop()
```
