## Supplementary material for "ΔSCOPE: A new method to quantify 3D biological structures and identify differences in zebrafish forebrain development": An archived copy of the DeltaSCOPE code repository.: sample_app.py

```
#!/usr/bin/env python

import tkinter as tk
from tkinter import filedialog, messagebox
from tkinter import ttk

import os
import yaml
import copy
import cranium
import numpy as np
import time
from appClass import *

import matplotlib
matplotlib.use('TkAgg')
from matplotlib.backends.backend_tkagg import FigureCanvasTkAgg, NavigationToolbar2TkAgg
import matplotlib.pyplot as plt
from matplotlib.figure import Figure

class Application(tk.Frame):
	def __init__(self,master=None):
		tk.Frame.__init__(self,master)

		self.grid(sticky=tk.N+tk.S+tk.E+tk.W)
		self.createNotebook()
		self.s = ttk.Style()

		#Create none variables so that app can always be saved
		self.fnums, self.Lnums = None,None

	def createNotebook(self):
		'''Create notebook to contain different tabs with data processing steps'''
		self.note = ttk.Notebook(self)
		self.note.grid()

		self.tab1 = ttk.Frame(self.note)
		self.note.add(self.tab1,text='Channels')
		self.tab2 = ttk.Frame(self.note)
		self.note.add(self.tab2,text='Files',state='disabled')
		self.tab25 = ttk.Frame(self.note)
		self.note.add(self.tab25,text='Settings',state='normal')
		self.tab3 = ttk.Frame(self.note)
		self.note.add(self.tab3,text='Alignment',state='disabled')

		self.createTab1()
		self.createTab2()
		self.createTab25()
		self.createTab3()

	def createTab1(self):
		'''Create tab that allows user to define the source directory for data'''

		#Deactivated next button that turns on after user adds data to channel 1
		self.nextB = tk.Button(self.tab1,text='Next',command=lambda:self.note.select(self.tab2),state=tk.DISABLED)
		self.nextB.grid(row=5,column=2)

		#Initialize channel objects to support selection of source directories
		self.cs = channelMaster(self.tab1,'Structural channel: ',0)
		self.cs.dButton.bind('<Button-1>',lambda e:self.activate(self.nextB,self.note,self.tab2))
		self.c2 = channelMaster(self.tab1,'Channel 2: ',1)
		self.c3 = channelMaster(self.tab1,'Channel 3: ',2)
		self.c4 = channelMaster(self.tab1,'Channel 4: ',3)
		self.c5 = channelMaster(self.tab1,'Channel 5: ',4)
		self.Lc = [self.cs,self.c2,self.c3,self.c4,self.c5]

		#Loads saved files
		self.loadB = tk.Button(self.tab1,text='Load saved file',command=self.load_status)
		self.loadB.grid()

	def createTab2(self):
		'''Tab 2 displays a list of files found in source directory
		and has check boxes for selection'''
		
		#Generate file list button
		self.filegenB = tk.Button(self.tab2,text='Generate file list',command=self.filegen)
		self.filegenB.grid(row=0)

		#Save button
		self.saveB = tk.Button(self.tab2,text='Save',command=self.save_status)
		self.saveB.grid(row=0,column=1)

		#Go to next page button
		self.nextB2 = tk.Button(self.tab2,text='Next',command=lambda:self.next_tab(self.tab25))
		self.nextB2.grid(row=0,column=2)

	def createTab25(self):
		'''Text entry objects for all parameters with examples already entered'''

		##### Tab 2.5 #####
		r = 0
		self.outdir = None
		self.outdirE = tk.Button(self.tab25,text='Final Output Folder: ',command=self.select_outdir)
		self.outdirE.grid(row=r,column=0)

		self.nextB25 = tk.Button(self.tab25,text='Next',command=lambda:self.next_tab(self.tab3))
		self.nextB25.grid(row=0,column=2)

		#Info in dict: row,title,example,help text
		self.p = {
			'expname': {
				'row': 1,
				'title': 'Experiment name: ',
				'example': 'experiment'
			},
			'medthresh': {
				'row': 2,
				'title': 'Median filter threshold: ',
				'example': '0.25'
			},
			'radius': {
				'row': 3,
				'title': 'Median filter radius: ',
				'example': '20',
			},
			'genthresh': {
				'row': 4,
				'title': 'General threshold: ',
				'example': '0.5'
			},
			'microns': {
				'row': 5,
				'title': 'Micron dimensions of voxel: ',
				'example': ['0.16','0.16','0.21']
			},
			'deg': {
				'row': 6,
				'title': 'Degree of fitted function: ',
				'example': '2'
			},
			'comporder': {
				'row': 7,
				'title': 'Assignment order of principle components: ',
				'example': ['0','2','1']
			},
			'fitdim': {
				'row': 8,
				'title': 'Dimensions to fit function: ',
				'example': 'x,z'
			}
		}

		#Create text entry objects
		for key in self.p.keys():
			#Triple entry
			if key in ['microns','comporder']:
				tk.Label(self.tab25,text=self.p[key]['title']).grid(row=self.p[key]['row'],column=0)

				tk.Label(self.tab25,text='X:').grid(row=self.p[key]['row'],column=1)
				self.p[key]['entry_x'] = tk.Entry(self.tab25)
				self.p[key]['entry_x'].grid(row=self.p[key]['row'],column=2)
				self.p[key]['entry_x'].insert(0,self.p[key]['example'][0])

				tk.Label(self.tab25,text='Y:').grid(row=self.p[key]['row'],column=3)
				self.p[key]['entry_y'] = tk.Entry(self.tab25)
				self.p[key]['entry_y'].grid(row=self.p[key]['row'],column=4)
				self.p[key]['entry_y'].insert(0,self.p[key]['example'][1])

				tk.Label(self.tab25,text='Z:').grid(row=self.p[key]['row'],column=5)
				self.p[key]['entry_z'] = tk.Entry(self.tab25)
				self.p[key]['entry_z'].grid(row=self.p[key]['row'],column=6)
				self.p[key]['entry_z'].insert(0,self.p[key]['example'][2])

			#Drop down fit dim
			elif key == 'fitdim':
				tk.Label(self.tab25,text=self.p[key]['title']).grid(row=self.p[key]['row'],column=0)

				options = ('XZ Plane','XY Plane', 'YZ Plane')
				self.p[key]['entry'] = tk.StringVar()
				
				self.p[key]['entry'].set(options[0])

				self.p[key]['menu'] = tk.OptionMenu(self.tab25,self.p[key]['entry'],*options)
				self.p[key]['menu'].grid(row=self.p[key]['row'],column=1)

			#Single entry
			else:
				tk.Label(self.tab25,text=self.p[key]['title']).grid(row=self.p[key]['row'],column=0)
				self.p[key]['entry'] = tk.Entry(self.tab25)
				self.p[key]['entry'].grid(row=self.p[key]['row'],column=1)
				self.p[key]['entry'].insert(0,self.p[key]['example'])

		self.p['outdir'] = None

	def createTab3(self):

		##### Tab 3 #####
		# self.plotB = tk.Button(self.tab3,text='plot',command=lambda:self.plot_projection('06'))
		# self.plotB.grid(row=1)

		self.pBar = ttk.Progressbar(self.tab3)
		self.pBar.grid(row=0,column=1)

		tk.Label(self.tab3,text='Rotate by ___ degrees:').grid(row=0,column=3)
		tk.Label(self.tab3,text='around ___ axis:').grid(row=0,column=4)
		tk.Label(self.tab3,text='Done?').grid(row=0,column=6)

		
	def activate(self,button,nb,ntab):
		button['state'] = tk.NORMAL
		button['bg'] = 'green'
		nb.tab(ntab,state='normal')

	def next_tab(self,ntab):
		self.note.tab(ntab,state='normal')
		self.note.select(ntab)

	def select_outdir(self):
		self.p['outdir'] = filedialog.askdirectory()
		tk.Label(self.tab25,text=self.p['outdir']).grid(row=0,column=1)

	def filegen(self):
		self.fnums = {}
		self.Lnums = []
		
		for i,c in enumerate(self.Lc):
			if c.dir != None and c.dir != '':
				files = os.listdir(c.dir)
			else:
				files = []

			c.files = []
			for f in files:
				if 'Probabilities' in f and 'h5' in f:
					key = f.split('_')[-2]
					if i == 0:
						self.fnums[key] = Sample(key)
						self.fnums[key].fs[i] = f
						self.Lnums.append(int(key))
					else:
						self.fnums[key].fs[i] = f

		self.Lnums.sort()
		self.display_files()

	def display_files(self):

		#Header 
		ttk.Label(self.tab2,text='Use?').grid(row=1,column=0)
		#ttk.Label(self.tab2,text='Number').grid(row=1,column=2)
		ttk.Label(self.tab2,text='Structural').grid(row=1,column=1)
		ttk.Label(self.tab2,text='Channel 2').grid(row=1,column=2)
		ttk.Label(self.tab2,text='Channel 3').grid(row=1,column=3)
		ttk.Label(self.tab2,text='Channel 4').grid(row=1,column=4)

		for i,n in enumerate(self.Lnums):
			key = str(n)
			if len(key) == 1:
				key = '0'+ key
			self.fnums[key].tab2row(self.tab2,i+2)
			self.fnums[key].tab3row(self.tab3,i+2,self)


	def out_fnums(self):
		for key in self.fnums.keys():
			print(self.fnums[key][-1].get())
		print('******')

	def save_status(self):

		fpath = filedialog.asksaveasfilename(defaultextension=".yml")
		print(fpath)

		out = {}
		out['curtab'] = self.note.tab(self.note.select(),'text')

		for i,c in enumerate([self.cs,self.c2,self.c3,self.c4]):
			if c == None:
				out['c'+str(i)] = None
			else:
				out['c'+str(i)] = c.return_data_dict()

		out['Lnums'] = self.Lnums

		out['fnums'] = {}
		for key in self.fnums.keys():
			line = copy.deepcopy(self.fnums[key][:-1])
			line.append(self.fnums[key][-1].get())
			out['fnums'][key] = line

		print(out)
		print(self.fnums)

		f = open(fpath,'w')
		yaml.dump(out,f)
		f.close()

	def load_status(self):

		fpath = filedialog.askopenfilename()
		print(fpath)
		s = open(fpath,'r').read()
		d = yaml.load(s)

		varbind = {
			'curtab': {'Channels':self.tab1,'Files':self.tab2,'Alignment':self.tab3},
			'fnums': self.fnums,
			'Lnums': self.Lnums,
			'c0': self.cs,
			'c1': self.c2,
			'c2': self.c3,
			'c3': self.c4
		}

		n = 0
		for key in d.keys():
			if key == 'Lnums':
				self.Lnums = d[key]
			elif key == 'fnums':
				if d['fnums'] != None:
					self.fnums = {}
					for key in d['fnums'].keys():
						self.fnums[key] = d['fnums'][key][:-1]
						self.fnums[key].append(tk.IntVar().set(d['fnums'][key][-1]))
			elif key != 'curtab':
				if d[key] == None:
					varbind[key] = d[key]
				else:
					varbind[key].dir = d[key]['dir']
					varbind[key].dButton['text'] = d[key]['dir']
					varbind[key].name = d[key]['name']


		if d['curtab'] in ['Files','Alignment']:
			self.activate(self.nextB,self.note,self.tab2)
		if self.fnums != None:
			self.display_files()
		self.note.select(varbind['curtab'][d['curtab']])
		print('load complete')

if __name__=='__main__':
	app = Application()
	app.master.title('Sample Application')
	app.mainloop()
```
