## Supplementary material for "ΔSCOPE: A new method to quantify 3D biological structures and identify differences in zebrafish forebrain development": An archived copy of the DeltaSCOPE code repository.: README.rst

deltascope
===========

This package is designed to quantify biological structures in 3D image data. 

Features
+++++++++

- Compare sets of 3D biological images to identify differences
- Automatically align the structure in the image to correct for variation introduced during mounting and imaging 
- Generate descriptive graphs that quantify both the average and variation of the data
- Use machine learning techniques to classify samples and identify regions of statistically significant difference

Installation
++++++++++++

Package hosted on `PyPI <https://pypi.python.org/pypi/deltascope>`_. ::

	$ pip install deltascope

Support
++++++++

- Complete documentation is available on `Read the Docs <http://deltascope.readthedocs.io/en/latest/>`_.
- Check out the `Frequently Asked Question <faq>`_ page.
- Submit an issue describing a problem or question on the project's Github `issue tracker <https://github.com/msschwartz21/deltascope/issues>`_.

Contribute
+++++++++++

- Issue Tracker: https://github.com/msschwartz21/deltascope/issues
- Source Code: https://github.com/msschwartz21/deltascope

License
++++++++

This project is licensed under the GNU General Public License.
