## Supplementary material for "ΔSCOPE: A new method to quantify 3D biological structures and identify differences in zebrafish forebrain development": An archived copy of the DeltaSCOPE code repository.: ipynb_generator.py

````
from __future__ import print_function
from builtins import zip
from builtins import str
from builtins import range
__author__ = 'Hans Petter Langtangen <>'

#example
#python .\ipynb_generator.py datareport.aipynb test.ipynb WTDIR="C:\\Users\\zfishlab\\Desktop\\zrf1wt13umyot21um\\wt\\Output69semifinal" MUTDIR="C:\\Users\\zfishlab\\Desktop\\zrf1wt13umyot21um\\yot\\Output92semifinal" DTYPE='AT' MYVAR=4 GRADE='excellent'
# C:\Users\zfishlab\Desktop\zrf1wt13umyot21um\Correct_alignment

import sys, os, re, logging

# Mapping of shortnames like py to full language
# name like python used by markdown/pandoc
shortname2language = dict(
    py='Python', ipy='Python', pyshell='Python', cy='Python',
    c='C', cpp='Cpp', f='Fortran', f95='Fortran95',
    rb='Ruby', pl='Perl', sh='Shell', js='JavaScript', html='HTML',
    tex='Tex', sys='Bash',
    )

def read(text, argv=sys.argv[2:]):
    lines = text.splitlines()
    # First read all include statements
    for i in range(len(lines)):
        if lines[i].startswith('#include "'):
            filename = lines[i].split('"')[1]
            with open(filename, 'r') as f:
                include_text = f.read()
            lines[i] = include_text
    text = '\n'.join(lines)
    logging.info('******* text after include:\n%s' % text)

    # Run Mako
    mako_kwargs = {}
    for arg in argv:
        key, value = arg.split('=')
        mako_kwargs[key] = value

    encoding = 'utf-8'
    try:
        import mako
        has_mako = True
    except ImportError:
        print('Cannot import mako - mako is not run')
        has_mako = False

    if has_mako:
        from mako.template import Template
        from mako.lookup import TemplateLookup
        lookup = TemplateLookup(directories=[os.curdir])

        text = str(text)
        temp = Template(text=text, lookup=lookup,
                        strict_undefined=True)
        logging.info('******* mako_kwargs: %s' % str(mako_kwargs))
        text = temp.render(**mako_kwargs)

    logging.info('******* text after mako:\n%s' % text)

    # Parse the cells
    lines = text.splitlines()
    cells = []
    inside = None    # indicates which type of cell we are inside
    fullname = None  # full language name in code cells
    for line in lines:
        if line.startswith('-----'):
            # New cell, what type?
            m = re.search(r'-----([a-z0-9-]+)?', line)
            if m:
                shortname = m.group(1)
                if shortname:
                    # Check if code is to be typeset as static
                    # Markdown code (e.g., shortname=py-t)
                    logging.info('******* found shortname %s'
                                 % shortname)
                    astext = shortname[-2:] == '-t'
                    logging.info(
                        '******* cell: astext=%s shortname=%s'
                        % (astext, shortname))
                    if astext:
                        # Markdown
                        shortname = shortname[:-2]
                        inside = 'markdown'
                        cells.append(['markdown', 'code', ['\n']])
                        cells[-1][2].append('```%s\n' % fullname)
                    else:
                        # Code cell
                        if shortname in shortname2language:
                            fullname = shortname2language[shortname]
                        inside = 'codecell'
                        cells.append(['codecell', fullname, []])
                else:
                    logging.info('******* cell: markdown')
                    # Markdown cell
                    inside = 'markdown'
                    cells.append(['markdown', 'text', ['\n']])
            else:
                raise SyntaxError(
                    'Wrong syntax of cell delimiter:\n%s'
                    % repr(line))
        else:
            # Ordinary line in a cell
            if inside in ('markdown', 'codecell'):
                cells[-1][2].append(line)
            else:
                raise SyntaxError(
                    'line\n  %s\nhas not beginning cell delimiter'
                    % line)
    # Merge the lines in each cell to a string
    for i in range(len(cells)):
        if cells[i][0] == 'markdown' and cells[i][1] == 'code':
            # Add an ending ``` of code
            cells[i][2].append('```\n')
        cells[i][2] = '\n'.join(cells[i][2])
    import pprint
    logging.info('******* cell data structure:\b%s'
                 % pprint.pformat(cells))
    return cells

def write(cells, nb_version=4):
    """Turn cells list into valid IPython notebook code."""
    # Use IPython.nbformat functionality for writing the notebook
    
    Dmetadata = {
        "anaconda-cloud": {},
        "kernalspec": {
            "display_name": "Python [Root]",
            "language": "python",
            "name": "Python [Root]"
        },
        "language_info": {
            "codemirror_mode": {
            "name": "ipython",
            "version": 3
            }
        },
        "file_extension": ".py",
        "mimetype": "test/x-python",
        "name": "python",
        "nbconvert_exporter": "python",
        "pygments_lexer": "ipython3",
        "version": "3.5.2"
    }

    if nb_version == 3:
        from nbformat.v3 import (
            new_code_cell, new_text_cell, new_worksheet,
            new_notebook, new_metadata, new_author)
        nb = new_worksheet()

    elif nb_version == 4:
        from nbformat.v4 import (
            new_code_cell, new_markdown_cell, new_notebook)
        nb_cells = []

    for cell_tp, language, block in cells:
        if cell_tp == 'markdown':
            if nb_version == 3:
                nb.cells.append(
                    new_text_cell(u'markdown', source=block))
            elif nb_version == 4:
                nb_cells.append(
                    new_markdown_cell(source=block))
        elif cell_tp == 'codecell':
            if nb_version == 3:
                nb.cells.append(new_code_cell(input=block))
            elif nb_version == 4:
                nb_cells.append(new_code_cell(source=block))

    if nb_version == 3:
        nb = new_notebook(worksheets=[nb], metadata=new_metadata())
        # Let us make v4 notebook here by upgrading
        from nbformat.v4 import upgrade
        nb = upgrade(nb)
        import nbformat.v4.nbjson as nbjson
        # Convert nb to json format
        filestr = nbjson.writes(nb)
    elif nb_version == 4:
        nb = new_notebook(cells=nb_cells, metadata=Dmetadata)
        from nbformat import writes
        filestr = writes(nb, version=4)
    return filestr

def driver():
    """Compile a document and its variables."""
    try:
        filename = sys.argv[1]
        with open(filename, 'r') as f:
            text = f.read()
    except (IndexError, IOError) as e:
        print('Usage: %s filename' % (sys.argv[0]))
        print(e)
        sys.exit(1)
    cells = read(text, argv=sys.argv[3:])
    filestr = write(cells, 3)
    outname = sys.argv[2]
    with open(outname, 'w') as f:
        f.write(filestr)


def test_notebook_generator():
    """Unit test for nosetests or py.test."""
    argv = ['NAME=Core Dump',
            'ADDRESS=Seg. Fault Ltd and Univ. of C. Space',
            'IC=2']
    with open('.test1.aipynb', 'r') as f:
        text = f.read()
    logging.info('******* original text:\n%s' % text)
    cells = read(text, argv=argv)
    computed = write(cells, 3).strip()
    print(computed)

    with open('.test1.ipynb', 'r') as f:
        expected = f.read().strip()

    for c1, c2, n in zip(computed, expected, list(range(len(expected)))):
        if c1 != c2:
            print('character no.%d differ: %s vs %s' % (n, c1, c2))
    assert computed == expected

if __name__ == '__main__':
    logfile = 'tmp.log'
    if os.path.isfile(logfile):
        os.remove(logfile)
    logging.basicConfig(format='%(message)s', level=logging.DEBUG,
                        filename=logfile)
    driver()
````
