## Supplementary material for "ΔSCOPE: A new method to quantify 3D biological structures and identify differences in zebrafish forebrain development": An archived copy of the DeltaSCOPE code repository.: mp_create_csv.py

```
#import plotly

#plotly.tools.set_credentials_file(username='msschwartz21', api_key='OrM0hMDBvseeT6SCjxNb')

def process_data(path):

	tic = time.time()

	s = cranium.brain()
	s.read_data(path)
	s.create_dataframe()
	s.pca_double_transform(0.5,['x','z'])
	s.transform_coordinates()

	columns = ['x','y','z','ac','r','theta']
	cranium.write_data(path[:-3]+'.psi',s.df_align[columns])

	toc = time.time()
	print(toc-tic)

if __name__=='__main__':

	root = 'C:\\Users\\zfishlab\\Desktop\\zrf1wt13umyot21um'
	stypes = ['wt']#,'yot'] #['wt','yot']
	chs = ['AT2\\Prob','ZRF1']
	prefixes = ['AT','ZRF1']

	#Create outdir
	outdir = os.path.join(root,'Output'+str(randint(10,99)))
	os.mkdir(outdir)
	print('outdir',outdir)

	for stype in stypes:
		d = os.path.join(root,stype,chs[0])
		print(stype,'starting')

		files = os.listdir(d)
		#Get sample numbers
		nums = []
		for file in files:
			if 'h5' in file and 'Probabilities' in file:
				nums.append(os.path.join(d,file))

		print(nums)

		processfxn = partial(cranium.process_sample,
			root=os.path.join(root,stype),
			outdir=outdir,
			name=stype,
			chs=chs,
			prefixes=prefixes,
			threshold=0.5,
			scale=[100000,1,1000],
			deg=2,
			primary_key='at',
			comp_order=[0,2,1],
			fit_dim=['x','z'],
			flip_dim='z')

		n = len(nums)
		for i in range(0,n,5):
			if i+5 > n:
				L = nums[i:n]
			else:
				L = nums[i:i+5]

			pool = mp.Pool()
			pool.map(process_data,L)
			pool.close()
			pool.join()

		print(stype, 'complete')


	# for d in dirs:
	# 	print(d)
	# 	files = os.listdir(d)
		
	# 	#Check that all files are hdf5
	# 	hfiles = []
	# 	for file in files:
	# 		if 'h5' in file and 'Probabilities' in file:
	# 			hfiles.append(os.path.join(d,file))

	# 	pool = mp.Pool()
	# 	pool.map(cranium.process_sample,hfiles)
	# 	pool.close()
	# 	pool.join()

	# 	print(d,'complete')
```
