## Supplementary material for "ΔSCOPE: A new method to quantify 3D biological structures and identify differences in zebrafish forebrain development": An archived copy of the DeltaSCOPE code repository.: mp_modchannel.py

```
import cranium
import multiprocessing as mp
import time
import os
import sys
from pushbullet import Pushbullet
from functools import partial
from random import randint

def process_data(num,name=None,microns=None,outdir=None,adir=None,zdir=None,gtype=None):

	tic = time.time()

	e = cranium.embryo(name,num,outdir)
	e.add_channel(os.path.join(adir,'AT_'+num+'_Probabilities.h5'),'AT')
	e.add_channel(os.path.join(zdir,gtype+'_'+num+'_Probabilities.h5'),gtype)
	e.process_channels(0.25,0.5,7,[1,1,1],microns,2,'AT',[0,2,1],['x','z'])

	e.save_psi()

	toc = time.time()
	print(num,toc-tic)

if __name__=='__main__':

	ywt = [
		"C:\\Users\\zfishlab\\Desktop\\zrf1wt13umyot21um\\wt\\AT2\\prob",
		"C:\\Users\\zfishlab\\Desktop\\zrf1wt13umyot21um\\wt\\ZRF1",
		"C:\\Users\\zfishlab\\Desktop\\zrf1wt13umyot21um\\wt"
	]

	yot = [
		"C:\\Users\\zfishlab\\Desktop\\zrf1wt13umyot21um\\yot\\AT2\\prob",
		"C:\\Users\\zfishlab\\Desktop\\zrf1wt13umyot21um\\yot\\ZRF1",
		"C:\\Users\\zfishlab\\Desktop\\zrf1wt13umyot21um\\yot"
	]

	yotdirset = [ywt,yot]
	yotnames = ['wt_yot','yot']
	yotmicrons = [[0.16,0.16,0.13],[0.16,0.16,0.21]]

	#S2s3 gfap dirs
	wt = [
		"D:\\s2s3 and zrf data 6.28.17\\slit2slit3\\WT\\AT", #adir
		"D:\\s2s3 and zrf data 6.28.17\\slit2slit3\\WT\\GFAP", #ZDIR
		"D:\\s2s3 and zrf data 6.28.17\\slit2slit3\\WT" #root
	]

	mut = [
		"D:\\s2s3 and zrf data 6.28.17\\slit2slit3\\mutant\\AT",
		"D:\\s2s3 and zrf data 6.28.17\\slit2slit3\\mutant\\GFAP",
		"D:\\s2s3 and zrf data 6.28.17\\slit2slit3\\mutant"
	]

	# dirset=[ywt,yot,wt,mut]
	# names=['wt_yot','yot','wt_s2s3','s2s3']
	# s2s3microns = [[0.16,0.16,0.13],[0.16,0.16,0.21],[0.16,0.16,0.21],[0.16,0.16,0.21]]
	# Lgtype = ['ZRF1','ZRF1','GFAP','GFAP']


	zrf2 = [
		"D:\\zrfs\\ZRF2\\wt\\AT",
		"D:\\zrfs\\ZRF2\\wt\\ZRF2",
		"D:\\zrfs\\ZRF2\\wt"
	]
	
	zrf3 = [
		"D:\\zrfs\\ZRF3\\AT",
		"D:\\zrfs\\ZRF3\\ZRF3",
		"D:\\zrfs\\ZRF3"
	]

	zrf4 = [
		"D:\\zrfs\\ZRF4\\wt\\AT",
		"D:\\zrfs\\ZRF4\\wt\\ZRF4",
		"D:\\zrfs\\ZRF4\\wt"
	]

	# dirset = [zrf2,zrf3,zrf4,ywt,yot,wt,mut]
	# names = ['zrf2_wt','zrf3_wt','zrf4_wt''wt_yot','yot','wt_s2s3','s2s3']
	# microns = [[.16,.16,.21],[.16,.16,.21],[.16,.16,.21],
	# [0.16,0.16,0.13],[0.16,0.16,0.21],[0.16,0.16,0.21],[0.16,0.16,0.21]]
	# Lgtype = ['zrf2','zrf3','zrf4','ZRF1','ZRF1','GFAP','GFAP']

	hss1a = [
		"C:\\Users\\zfishlab\\Desktop\\HSs1a_yot_expt\\zrf1\\hss1a\\AT",
		"C:\\Users\\zfishlab\\Desktop\\HSs1a_yot_expt\\zrf1\\hss1a\\GFAP",
		"C:\\Users\\zfishlab\\Desktop\\HSs1a_yot_expt\\zrf1\\hss1a"
	]

	hss1ayot = [
		"C:\\Users\\zfishlab\\Desktop\\HSs1a_yot_expt\\zrf1\\hss1ayot\\AT",
		"C:\\Users\\zfishlab\\Desktop\\HSs1a_yot_expt\\zrf1\\hss1ayot\\GFAP",
		"C:\\Users\\zfishlab\\Desktop\\HSs1a_yot_expt\\zrf1\\hss1ayot"
	]

	dirset = [hss1ayot]
	names = ['hss1ayot']
	microns = [[.16,.16,.31]]
	Lgtype = ['gfap']

	for i,dirs in enumerate(dirset):

		micron = microns[i]
		name = names[i]
		adir,zdir,root = dirs[0],dirs[1],dirs[2]

		#Create outdir
		outdir = os.path.join(root,'Output'+str(randint(10,99))+'thetafix')
		os.mkdir(outdir)
		print('outdir',outdir)

		processfxn = partial(process_data,name=name,microns=micron,outdir=outdir,adir=adir,zdir=zdir,gtype=Lgtype[i])

		nums = []
		for f in os.listdir(adir):
			if 'Probabilities' in f and 'h5' in f:
				print(f)
				nums.append(f.split('_')[1])

		print(nums)

		n = len(nums)
		for i in range(0,n,5):
			if i+5>n:
				L = nums[i:n]
			else:
				L = nums[i:i+5]

			# if i >= 5: break

			pool = mp.Pool()
			pool.map(processfxn,L)
			pool.close()
			pool.join()


		print('complete',root)
```
