## Supplementary material for "ΔSCOPE: A new method to quantify 3D biological structures and identify differences in zebrafish forebrain development": An archived copy of the DeltaSCOPE code repository.: multiprocess_11-9.py

```
import os
import numpy as np
import cranium
import multiprocessing as mp

genthresh = 0.5
medthresh = 0.25
radius = 20
microns = [0.16,0.16,0.21]
deg = 2
comporder = [0,2,1]
fitdim = ['x','z']
scale = [1,1,1]

columns = ['x','y','z','ac','r','theta']

outdir = "D:\\Wildtype_for_modeling\\PSI"

def process(filepath):
	s = cranium.brain()
	print(filepath)
	s.read_data(filepath)

	s.preprocess_data(genthresh,[1,1,1],microns)
	s.calculate_pca_median(s.raw_data,medthresh,radius,microns)

	s.pca_transform_3d(s.df_thresh,s.pcamed,comporder,fitdim)
	s.transform_coordinates()

	fname = filepath.split('\\')[-1].split('_')
	name = fname[0]+'_'+fname[1]+'_'+fname[2]+'.psi'

	cranium.write_data(os.path.join(outdir,name),s.df_align[columns])

	print(name)

ddir = "D:\\Wildtype_for_modeling\\Prob"

if __name__=='__main__':

	num=[]
	for f in os.listdir(ddir):
		if 'Probabilities' in f and 'h5' in f:
			num.append(os.path.join(ddir,f))

	for i in range(0,len(num),5):
		if i+5>len(num):
			L = num[i:len(num)]
		else:
			L = num[i:i+5]

		pool = mp.Pool()
		pool.map(process,L)
		pool.close()
		pool.join()
```
