## Supplementary material for "ΔSCOPE: A new method to quantify 3D biological structures and identify differences in zebrafish forebrain development": An archived copy of the DeltaSCOPE code repository.: multiprocess_11-20.py

```
import cranium
import multiprocessing as mp
import time
import os
import sys
from pushbullet import Pushbullet
from functools import partial
from random import randint


genthresh = 0.5
medthresh = 0.25
radius = 20
microns = [0.16,0.16,0.21]
deg = 2
comporder = [0,2,1]
fitdim = ['x','z']
scale = [1,1,1]

columns = ['x','y','z','ac','r','theta']

def process(num,expname=None,outdir=None,gtype=None,adir=None,zdir=None):
	tic = time.time()

	e = cranium.embryo(expname,num,outdir)

	e.add_channel(os.path.join(adir,'AT_'+num+'_Probabilities.h5'),'AT')
	e.add_channel(os.path.join(zdir,gtype+'_'+num+'_Probabilities.h5'),gtype)
	e.chnls['AT'].preprocess_data(genthresh,[1,1,1],microns)
	e.chnls[gtype].preprocess_data(genthresh,[1,1,1],microns)

	e.chnls['AT'].calculate_pca_median(e.chnls['AT'].raw_data,medthresh,radius,microns)
	pca = e.chnls['AT'].pcamed
	e.chnls['AT'].pca_transform_3d(e.chnls['AT'].df_thresh,pca,comporder,fitdim,deg=2)

	mm = e.chnls['AT'].mm
	vertex = e.chnls['AT'].vertex

	e.chnls[gtype].pca_transform_3d(e.chnls[gtype].df_thresh,pca,comporder,fitdim,deg=2,mm=mm,vertex=vertex)

	e.chnls['AT'].transform_coordinates()
	e.chnls[gtype].transform_coordinates()

	e.save_psi()

	toc = time.time()
	print(num,toc-tic)

if __name__=='__main__':

	zrf2 = [
		"D:\\zrfs\\ZRF2\\wt\\AT",
		"D:\\zrfs\\ZRF2\\wt\\ZRF2",
		"D:\\zrfs\\ZRF2\\wt"
	]
	
	zrf3 = [
		"D:\\zrfs\\ZRF3\\AT",
		"D:\\zrfs\\ZRF3\\ZRF3",
		"D:\\zrfs\\ZRF3"
	]

	zrf4 = [
		"D:\\zrfs\\ZRF4\\wt\\AT",
		"D:\\zrfs\\ZRF4\\wt\\ZRF4",
		"D:\\zrfs\\ZRF4\\wt"
	]

	Lgtype = ['zrf2','zrf3','zrf4']
	names = ['wtzrf2','wtzrf3','wtzrf4']
	dirset = [zrf2,zrf3,zrf4]

	for i,dirs in enumerate(dirset):
		name = names[i]
		adir,zdir,root = dirs[0],dirs[1],dirs[2]

		#Create outdir
		outdir = os.path.join(root,'11-20_Output'+str(randint(10,99)))
		os.mkdir(outdir)
		print('outdir',outdir)

		processfxn = partial(process,expname=name,outdir=outdir,gtype=Lgtype[i],adir=adir,zdir=zdir)

		nums = []
		for f in os.listdir(adir):
			if 'Probabilities' in f and 'h5' in f:
				print(f)
				nums.append(f.split('_')[1])

		print(nums)

		n = len(nums)
		for i in range(0,n,5):
			if i+5>n:
				L = nums[i:n]
			else:
				L = nums[i:i+5]

			# if i >= 5: break

			pool = mp.Pool()
			pool.map(processfxn,L)
			pool.close()
			pool.join()


		print('complete',root)
```
