## Supplementary material for "ΔSCOPE: A new method to quantify 3D biological structures and identify differences in zebrafish forebrain development": An archived copy of the DeltaSCOPE code repository.: setup.py

```
import setuptools


def readme():
    with open('README.rst') as f:
        return (f.read())


def requirements():
    with open('requirements.txt') as f:
        return (f.read().split('\n'))


setuptools.setup(name='deltascope',
                 version='1.0.0',
                 description='Python package to quantify biological structure',
                 long_description=readme(),
                 keywords='biology image analysis quantification',
                 url='https://github.com/msschwartz21/deltascope',
                 author='Morgan Schwartz',
                 author_email='',
                 license='GNU GPL',
                 packages=['deltascope'],
                 install_requires=requirements(),
                 include_package_data=True,
                 classifiers=[
                     "License :: OSI Approved :: GNU General Public License (GPL)",
                     "Programming Language :: Python :: 3.7"
                 ]
                 )
```
