## Supplementary figures and images for "ΔSCOPE: A new method to quantify 3D biological structures and identify differences in zebrafish forebrain development"

### ajax-loader.gif

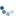

### alignment-correction.jpg

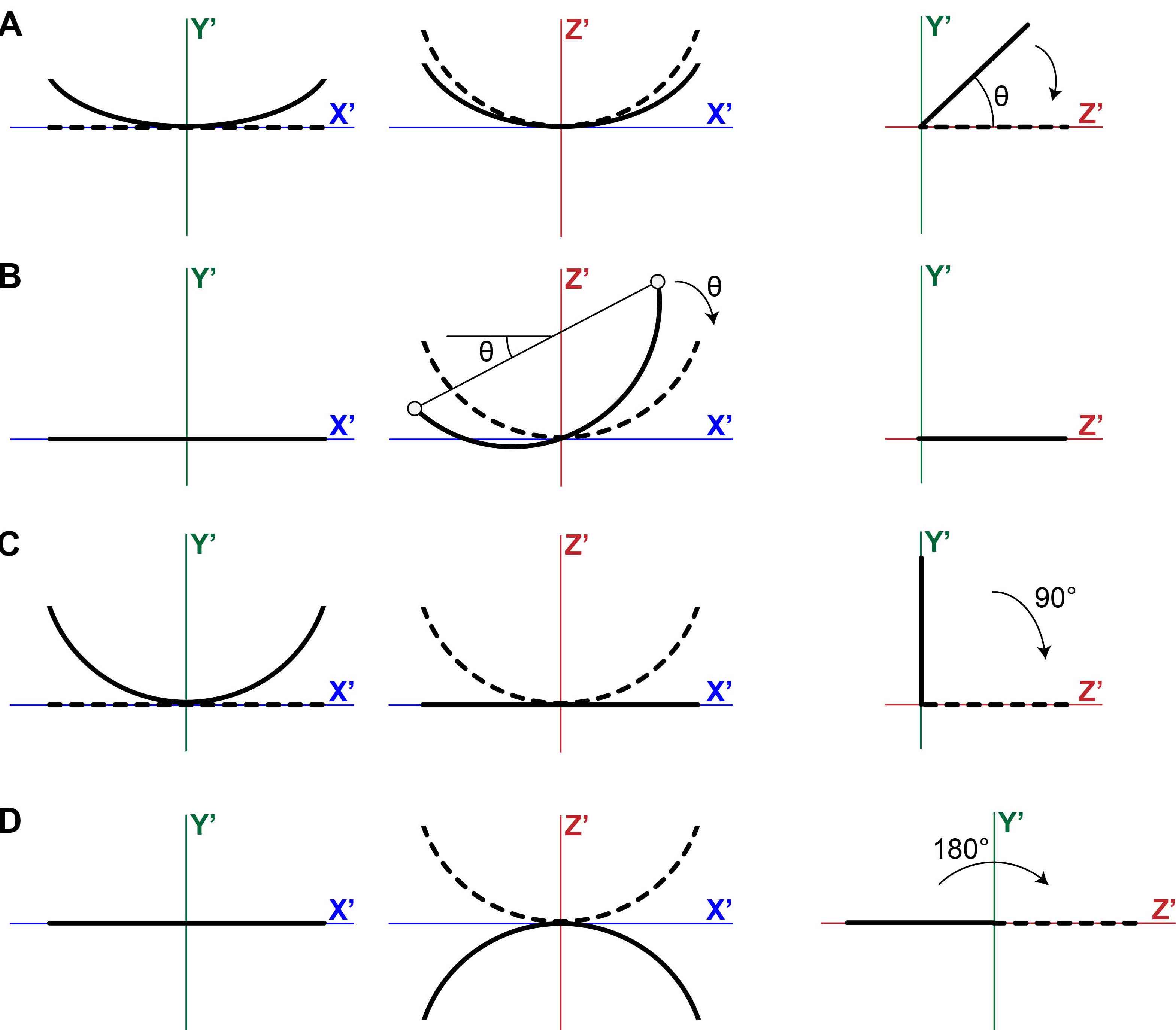

### anumOpt.png

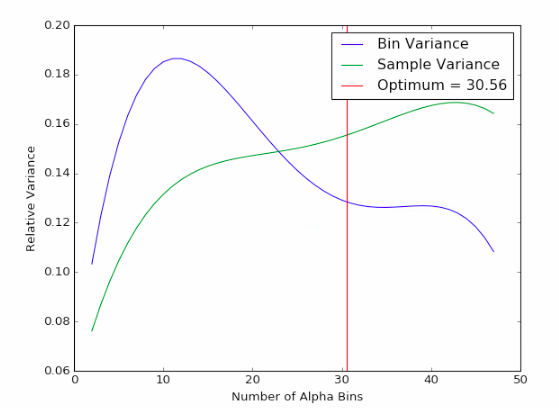

### cordsystem.jpg

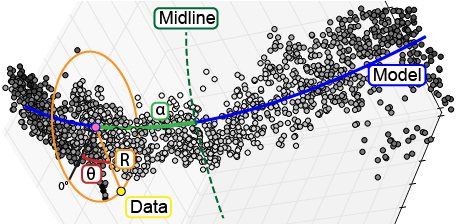

### landmarks.png

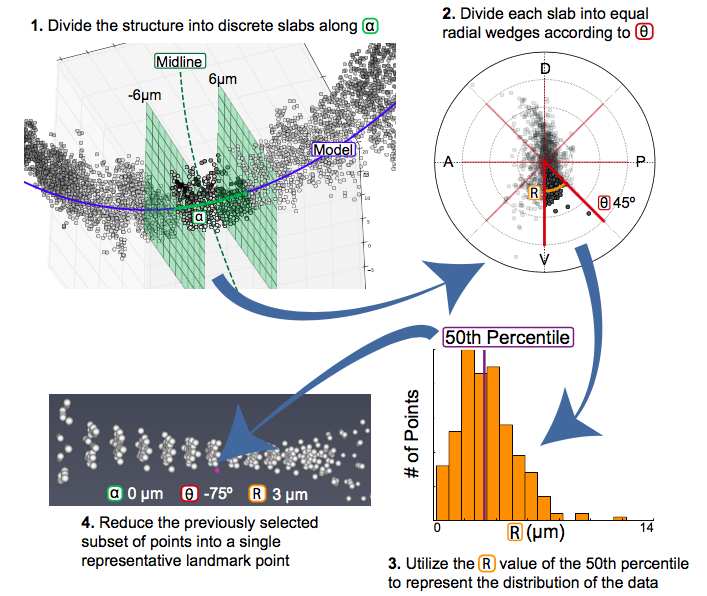

### pca.jpg

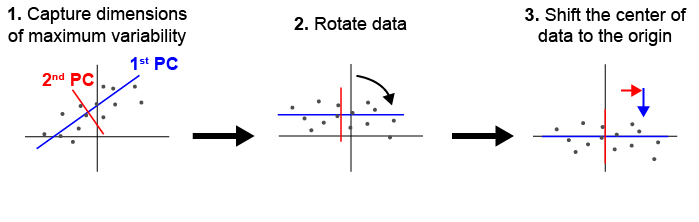

### pcafix.jpg

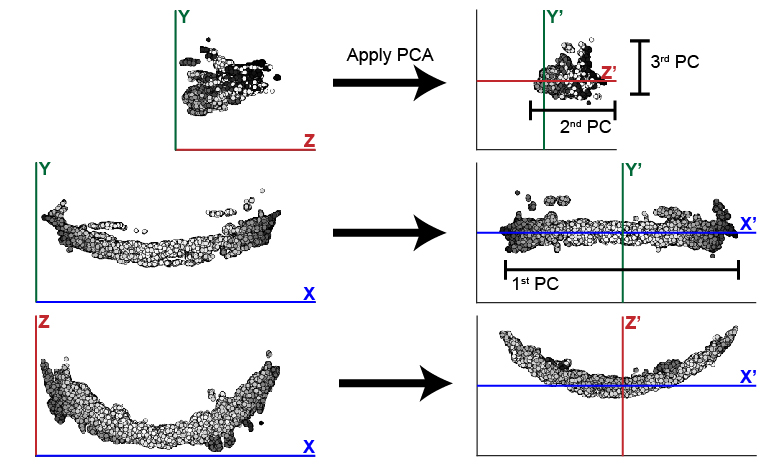
