## Supplementary material for "ΔSCOPE: A new method to quantify 3D biological structures and identify differences in zebrafish forebrain development": A pdf copy of documentation that accompanies the DeltaSCOPE code repository.

### deltascope Documentation

***Release 1.0.0***

**Morgan Schwartz**

**Jul 08, 2019**

### Contents

**1 Features** **1**

**2 Installation** **3**

**3 Support** **5**

**4 Contribute** **7**

**5 License** **9**

**6 Contents** **11**

**7 Indices and tables** **39**

**Python Module Index** **41**

**Index** **43**

CHAPTER 1

### Features

- Compare sets of 3D biological images to identify differences
- Automatically align the structure in the image to correct for variation introduced during mounting and imaging
- Generate descriptive graphs that quantify both the average and variation of the data
- Use machine learning techniques to classify samples and identify regions of statistically significant difference

Check out Start Here to find out if deltascope is right for you.

CHAPTER 2

### Installation

Package hosted on PyPI. See Start Here for more information.

```
$ pip install deltascope
```

### CHAPTER 3

#### Support

- Complete documentation is available on Read the Docs.
- Check out the Frequently Asked Question page.
- Submit an issue describing a problem or question on the project's Github issue tracker.

CHAPTER 4

### Contribute

- Issue Tracker: https://github.com/msschwartz21/deltascope/issues
- Source Code: https://github.com/msschwartz21/deltascope

### CHAPTER 5

#### License

This project is licensed under the GNU General Public License.

CHAPTER 6

### Contents

#### 6.1 Start Here

##### 6.1.1 Is deltascope right for you?

deltascope may be able to help you if:

- your data consists of a set of 3D image stacks
- your data contains a clear structure and shape that has consistent gross morphology between control and experimental samples
- you want to identify extreme or subtle differences in the structure between your experimental groups
- you have up to 4 different channels to compare

deltascope cannot help if:

- your data was collected using cryosections that need to be aligned after imaging
- your experiment changes the gross anatomy of the structure between control and experimental samples

##### 6.1.2 Installation

deltascope can be installed using pip, Python's package installer. Some deltascope dependencies are not available through pip, so we recommend that you install Anaconda which automatically includes and installs the remaining dependencies. See Setting up a Python environment for more details.

```
$ pip install deltascope
```

**Note:** If you are unfamiliar with the command line, check out this tutorial. Additional resources regarding the command line are available here.

**Warning:** Packages required for deltascope depend on Visual C++ Build Tools, which can be downloaded at build tools.

##### 6.1.3 Setting up a Python environment

If you're new to scientific computing with Python, we recommend that you install Anaconda to manage your Python installation. Anaconda is a framework for scientific computing with Python that will install important packages (numpy, scipy, and matplotlib).

**Warning:** deltascope is written in Python 3, and requires the installation the Python 3 version of Anaconda.

#### 6.2 Data Preparation

##### 6.2.1 Biological Questions

deltascope compares biological structures in three dimensions in order to preserve spatial relationship data that is lost in maximum intensity projections (MIPs). In order to apply deltascope to a new biological structure, certain conditions must be satisfied:

1. The gross morphology of the structure must be roughly consistent between samples.
2. The x, y, and z dimensions of the structure cannot be too similar in extent. For example, in the spinal cord, the anterior-posterior axis is the longest and the medial-lateral axis is the smallest with the dorsal-ventral axis falling between these two dimensions. The different proportional sizes of these axes enables us to consistently align the structure in 3D space regardless of the sample's orientation during image collection.
3. The gross morphology of the structure can be described with a simple polynomial equation. For example, the spinal cord can be described by a line that falls at the midline of the medial-lateral axis: y = mx + b.

##### 6.2.2 File Format

Most microscopes save data in their own proprietary data format: for example, Zeiss, `.czi`; Leica, `.lif`. In order to ensure that image data is legible to all components of the workflow, files need to be converted to the HDF5 (`.h5`) format specified by ImageJ. This conversion can be easily executed in Fiji using the BioFormats plugin to import proprietary file formats and the HDF5 plugin to export HDF5 files. Multichannel collections need to be split into individual channels before being saved as HDF5 files, with coherent file names utilized to preserve file history.

##### 6.2.3 Signal Normalization

Biological fluorescence microscopy data contains variation in signal intensity due to both biological and technical error. For example, the top of the sample is frequently brighter than the bottom because it is closer to the objective and also has not been as bleached by the collection of previous optical sections. If we were to try to select the set of points that represent 'true signal' by applying a single intensity threshold, points that represent background at the top of the stack may have the same intensity as points of true signal at the bottom of the stack.

We have implemented an adaptive thresholding protocol that avoids these challenges, utilizing the open-source software, Ilastik. This software uses machine learning principles in order to predict the likelihood that a particular pixel contains true signal. The probability is calculated based on user annotation of images, in which regions of true signal and background are labeled. This protocol allows the user to apply their domain knowledge of the sample in order to best distinguish signal from background. Tutorials describing how to install and implement Ilastiks pixel classification workflows are available on Ilastiks website.

#### 6.3 deltascope Data Processing

**Note:** This guide will describe a set of parameters that the user needs to specify when running deltascope. A complete list of all parameters if available on the Parameter Reference page.

##### 6.3.1 Intensity Thresholding

###### Goal

At this point, each sample/channel should have been processed by Ilastik to create a new `c1_10_Probabilities.h5` file. If you open this file in Fiji utilizing the fiji HDF5 plugin, it should contain three dimensions and two channels (signal and background, but for our purposes the two are interchangeable). Each pixel should have a value ranging between 0 and 1. If we are inspecting the signal channel, pixels with a value close to 0 (low p value) are highly likely to be true signal. Correspondingly, pixels with a value close to 1 are likely to be background. The `c1_10_Probabilities.h5` file by Ilastik contains two channels (signal and background), which are inverse images. For a given pixel, the background intensity value is 1 minus the signal intensity value. In order to simplify our data, we will apply a threshold that will divide the data into two sets of pixels: signal and background. In the steps that follow, we will only use the set of pixels that correspond to true signal. This set of pixels may be also referred to as a set of points or a point cloud. In order to avoid keeping track of which channel is the signal channel, we assume that after applying a threshold there will be more points will fall in the background group than in the signal group. If this is not true, better Ilastik training or a stricter threshold is reccomended, or deltascope will instead be examining the structure of your background.

**Warning:** If your data contains more points of signal than background, deltascope will select your background channel as the signal channel. In order to correct this assumption, the inequality in `brain.read_data()` will need to be changed to equate a large number of points with signal instead of background.

###### Setting Parameters

In order to divide the image into two sets of pixels (signal and background), we use a parameter, `genthresh`, to determine which group the pixels fall into. We have found that a value of 0.5 is sufficient to divide the data and have not found that the results vary greatly if it is changed; however, if your data contains a lot of intermediate background values (0.4-0.7), you may benefit from a smaller threshold, e.g. 0.3.

The following code requires a parameter `micron`, which specifies the dimensions of the image voxel in microns. It is a list of the form `[x,y,z]`. This information can typically be found in the metadata of a microscope collection file.

This section of the code also includes a deprecated parameter `scale`, which must be set to `[1,1,1]`.

###### Code Instructions

The following instructions apply to processing a single sample. Details regarding function parameters can be found under `embryo` and `brain.preprocess_data()`. Raw image data will be imported by `embryo.add_channel()`.

```
import deltascope

#Create an embryo object that facilitates data processing
e = deltascope.embryo(experiment-name,sample-number,directory)

#For each channel in your sample, add a channel with a unique name, e.g. 'c1' or 'c2'
e.add_channel(c1-filepath,c1-name)
e.add_channel(c2-filepath,c2-name)

#Threshold each channel and scale points according to voxel dimension in microns
e.chnls[c1-name].preprocess_data(genthresh,scale,microns)
e.chnls[c2-name].preprocess_data(genthresh,scale,microns)
```

##### 6.3.2 Sample alignment using principle component analysis (PCA)

###### Goal

###### Principle Component Analysis

During typical collections of biological samples, each sample will be oriented slightly differently in relation to the microscope due to variations in the shape and size of the sample as well as human error during the mounting process. As a result of this variation, we cannot directly compare samples in 3D space without realigning them. We have implemented principle component analysis in order to automate the process of alignment without a need for human supervision. Fig. 6.1 shown below illustrates how PCA can be used to align a set of points in 2D. deltascope uses the same process in 3D to identify biologically meaningful axes present in biological structures.

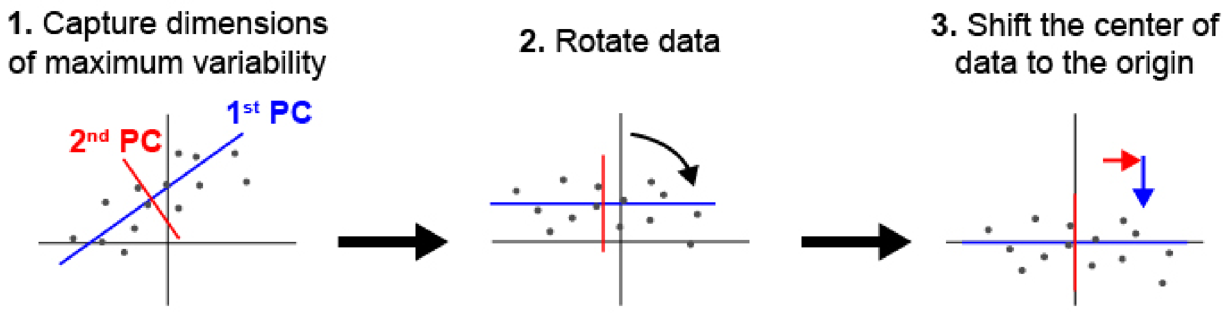

Fig. 6.1: Principle component analysis (PCA) can be used to identify and align samples along consistent axes. (1) In 2D, PCA first selects the axis that captures the most variability in the data (1st PC). The 2nd PC is selected in a position orthogonal to the 1st PC that captures the remaining variation in the data. (2) The data are then rotated so that the 1st and 2nd PCs correspond with x and y axes respectively. (3) Finally, we have added a step of identifying the center of the data and shifting it to the origin.

###### Structural Channel Processing

We designate one channel, the `structural channel`, which we will use for PCA to align samples. Since we are interested in the gross morphology of this channel, we apply two data preprocessing steps to reduce the data down to only essential points. First, we return to *brain.raw_data* and apply a new threshold, *medthresh*, which is typically more stringent than *genthresh*. This step ensures we are only considering points of signal with extremely high likelihood of being real. Second, we apply a median filter to the data twice, which smooths out the structure and eliminates small points of variation that may interfere with the alignment process of the gross structure.

PCA outputs three new dimensions, the 1st, 2nd, and 3rd PCs. These components will be reassigned to the X, Y, and Z axes to help the user maintain orientation in regards to their data. In the case of the zebrafish post-optic commissure shown below (Fig. 6.2), the 1st PC is reassigned to the X axis and the 2nd and 3rd PCs are assigned to Z and Y respectively. These assignments honor the user's expectation of the sample's alignment in 3D space. The assignment of components to axes can be modified using the parameter `comporder`.

> **Warning:** In order for PCA to consistently align your samples in the same orientation, we are assuming that the three dimensions of your structure are of different relative sizes. Since PCA looks for the axes that capture the most variation in your data, a sample that has axes of the same relative size will not have any distinguishing characteristics that PCA can use to identify and separate different axes.

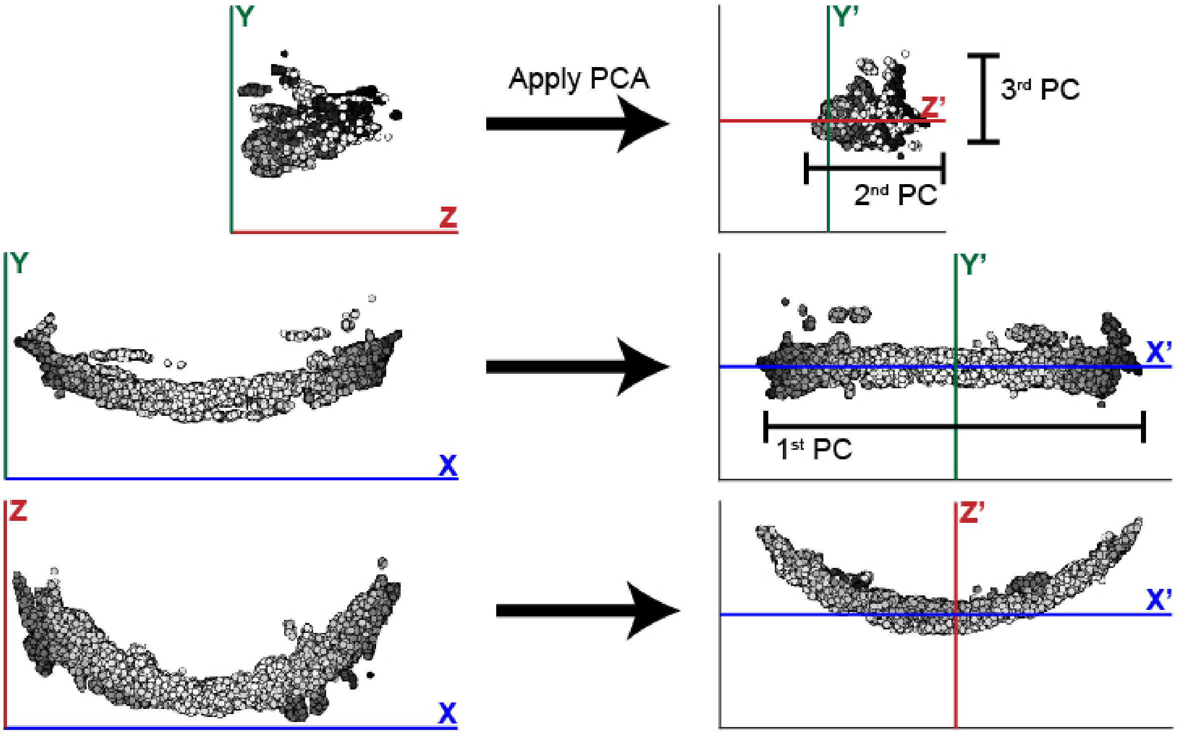

Fig. 6.2: This example illustrates the efficacy of PCA at changing the orientation of the zebrafish post optic commissure. In this case, the 1st PC is significantly longer than the 2nd and 3rd. While these two remaining components are similar in size, the typically longer depth of the 2nd PC distinguishes it from the 3rd PC.

##### Model Fitting

In addition to rotating the orientation of the data in 3D space, we also want to align the center of all samples at the origin. In order to determine the center of the data, we fit a simple mathematical model to the data that will also be used later in the analysis. The zebrafish post optic commissure shown above forms a parabolic structure, which can be described by y = ax^2 + bx + c. For simplicity, we fit the model in two dimensions, while holding one dimension constant. In the case of the POC, the parabolic structure lies flat in the XZ plane, which means that the structure can be described using exclusively the X and Z dimensions. The dimensions, which will be used to fit the 2D model, are specified in the parameter *fitdim*. Additionally the *deg* parameter specifies the degree of the function that fits to the data.

##### Setting Parameters

*medthresh* is typically set to 0.25, in comparison to a value of 0.5 for *genthresh*. If your data contains aberrant signal that does not contribute to the gross morphology of the structure, an even lower *medthresh* may help limit the negative influence of noisy signal. Additionally, the *radius* of the median filter can also be tuned to eliminate noisy signal. The typical value for *radius* is 20, which refers to the number of neighboring points that are considered in the median filter. A smaller value for *radius* will preserve small variation in signal, while a larger value will cause even more blunting and smoothing of the data. Prior to running the median threshold, it is reccomended that the user load several HDF5 files containing the structure of interest into FIJI and test several median thresholds to determine which best resolves their structure. Utilize the best median threshold radius in *radius*.

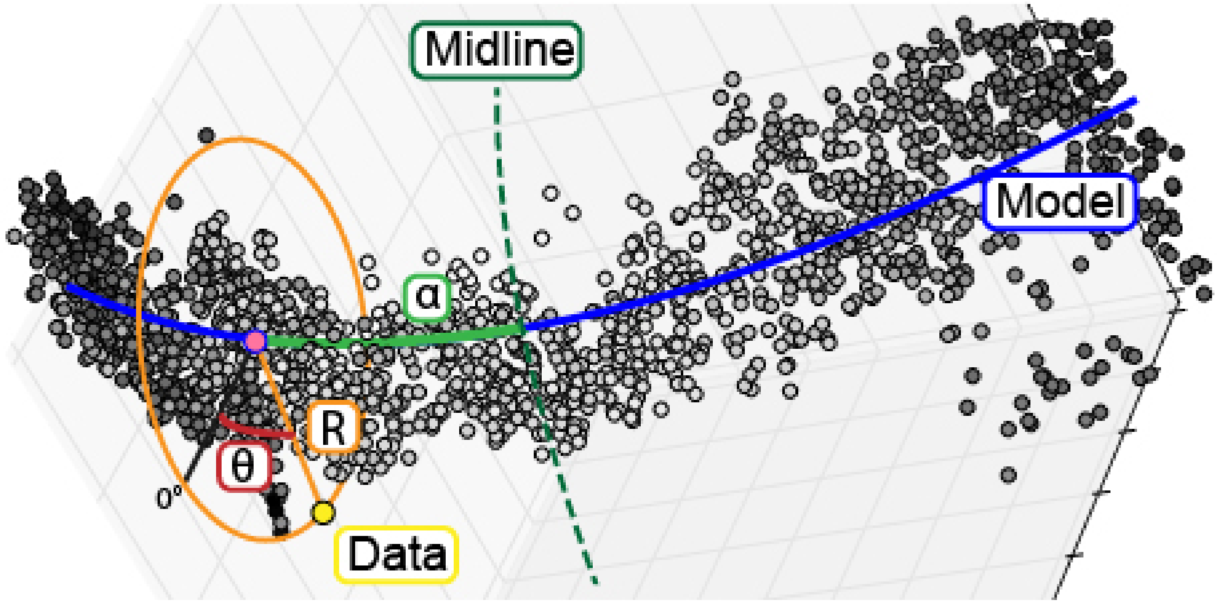

Fig. 6.3: To enable analysis of data point relative to a biological structure, points are transformed from a Cartesian coordinate system (x,y,z) into a cylindrical coordinate system ($alpha$,$theta$,R) defined relative to the structure.

##### Code Instructions

This transformation does not require defining any parameters; however, it assumes the data has already been thresholded and aligned using PCA.

```
#Transform each channel to cylindrical coordinates
e.chnls['c1'].transform_coordinates()
e.chnls['c2'].transform_coordinates()

#Save processed data to .psi file
e.save_psi()
```

**Warning:** This processing step is time consuming. We recommend running multiple samples in parallel in order to reduce the total amount of computational time required.

##### 6.3.4 Batch Processing

In order to reduce processing time, we have implemented a basic multiprocessing tool that runs 5 samples in parallel at a time. For more information, see *Batch Processing: Transformation*.

#### 6.4 Landmark Calculation

Following the steps described in *deltascope Data Processing*, your data should be saved as a set of `.psi` files containing 6 values for each point: x,y,z,alpha,r,theta. Depending on the size and resolution of your original image, you will have thousands of points, which are unwieldy when you are trying to compare sample sets with several images. In order to reduce the size of the data and facilitate direct comparison, we calculate a set of landmarks that describe the data.

##### 6.4.1 What is a landmark?

Landmark points are frequently used in the study of morphology to describe and compare structures. Classically, an individual with expert knowledge of the structure would define a set of points that are present in all structures, but subject to variation. For example, in the human face, landmarks might be placed at the corners of the eyes and mouth as well as the tip of the nose. If we were to compare many different faces, we could use the difference in the position of the landmarks to describe how the faces varied.

##### 6.4.2 Unbiased Landmarks

The challenge with the classical approach to landmark analysis lies in the step of assigning landmarks. If an expert user is selecting regions of the structure to assign landmarks to, they are projecting their own expectations as to where they expect to see variation. We have developed a method of automatically calculating landmarks that describe the structure without bias and allow the user to discover new regions of interest.

##### 6.4.3 How are landmarks calculated?

The calculation of unbiased landmarks relies on *Cylindrical Coordinates* that were previously defined (Fig. 6.3). First, the data is divided into equally sized sections along the alpha axis (Fig. 6.4.1). The user specifies the number of divisions *`anum`* and the data is divided accordingly. Next, each alpha subdivision is divided into radial wedges (Fig. 6.4.2) according to the parameter *`tsize`*, which specifies the size of each wedge. Finally, the distribution of points in the r axis is calculated according to the percentiles specified by the user in *`percbins`* (Fig. 6.4.3). Following these three steps, each subdivision can be represented by a single point that describes the distribution of the data in all three dimensions (Fig. 6.4.4) For more information on these parameters, see *Landmark Calculation*.

###### Code Sample

```
import deltascope
import numpy as np

anum = 30
tstep = np.pi/4

#Create a landmark object
lm = deltascope.landmarks(percbins=[50],rnull=15)
lm.calc_bins(dfs,anum,tstep)
```

(continues on next page)

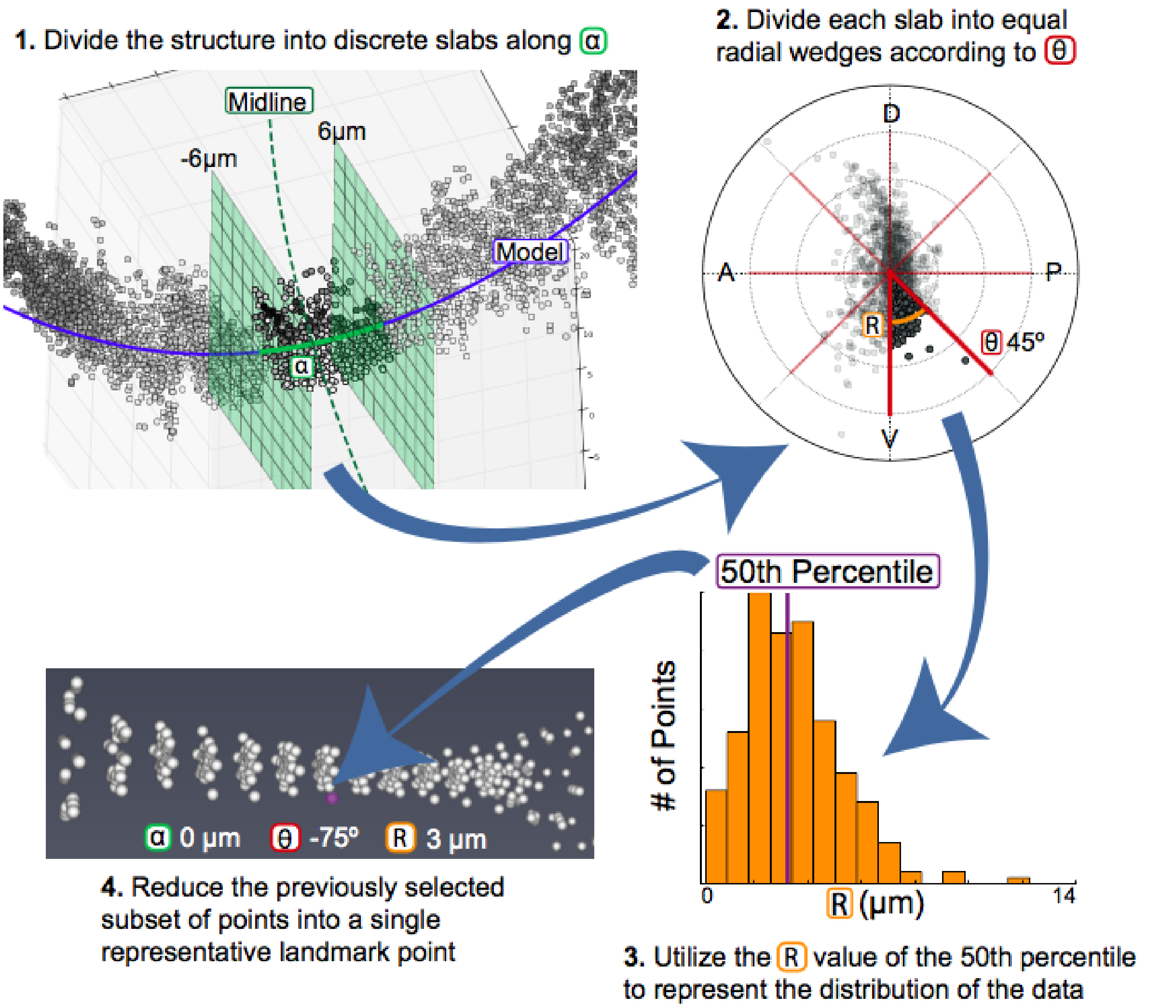

Fig. 6.4: In order to calculate landmarks, we will subdivide the data along the alpha and theta axes before calculating the r value that describes the distribution of the data.

(continued from previous page)

```
#Calculate landmarks for each sample and append to a single dataframe
outlm = pd.DataFrame()
for k in dfs.keys():
        outlm = lm.calc_perc(dfs[k],k,'stype',outlm)
```

##### 6.4.4 Selecting `anum`

The `anumSelect` can be used to identify the optimum number of sections along alpha. We use two measure of variance to test a range of *anum*. The first test compares the variance of adjacent landmark wedges. The second test compares the variability of samples in a landmark. As shown in Fig. 6.5, the optimum value of *anum* minimizes the variance of both tests.

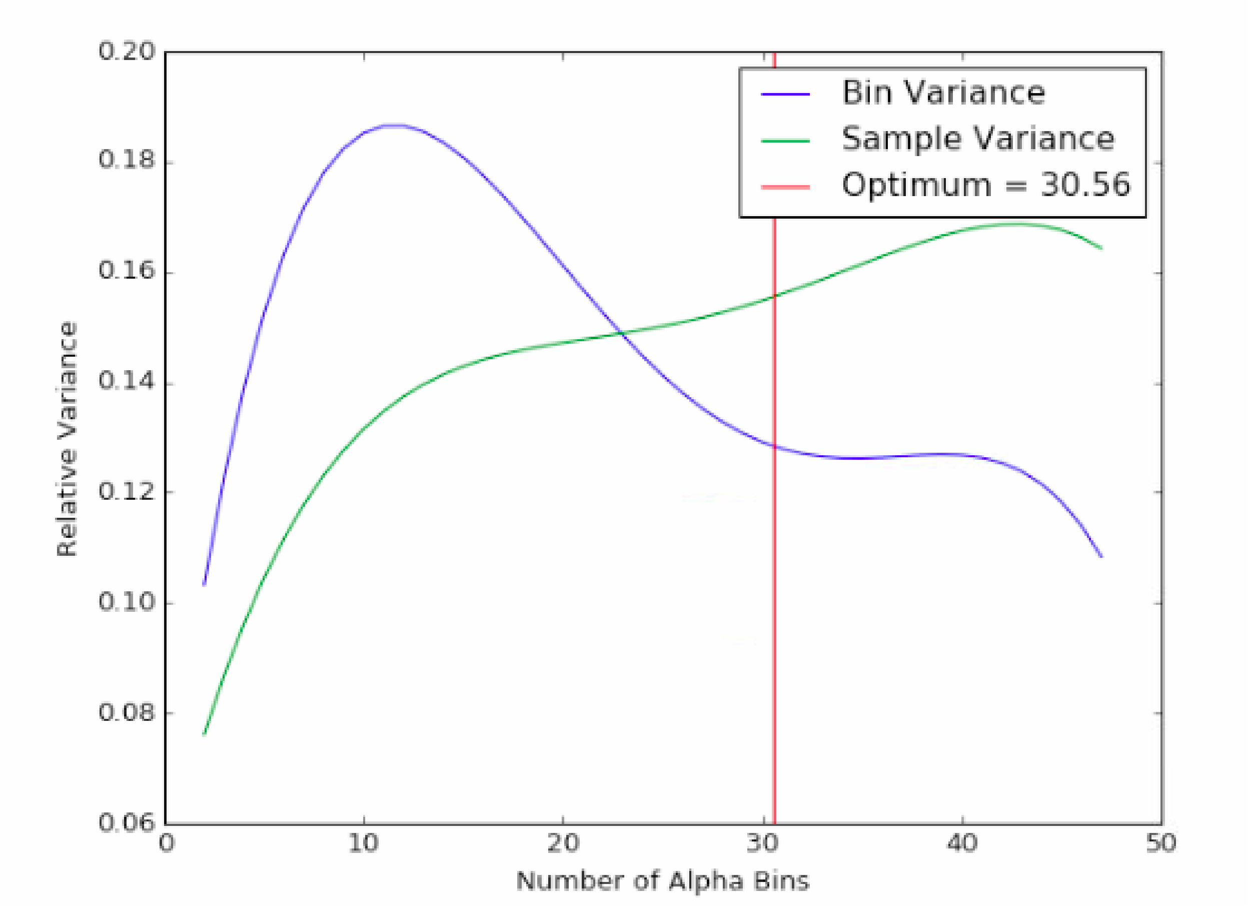

Fig. 6.5: We select the value of *anum* that minimizes both the bin variance and the sample variance.

###### Code Sample

```
import deltascope

#Create a optimization object
```

(continues on next page)

(continued from previous page)

```
opt = deltascope.anumSelect(dfs)

tstep = np.pi/4

#Initiate parameter sweep
opt.param_sweep(tstep,amn=2,amx=50,step=1,percbins=[50],rnull=15)

#Plot raw data
opt.plot_rawdata()

poly_degree = 4

#Test polynomial fit
opt.plot_fitted(poly_degree)

best_guess = 30

#Find the optimum value of anum
opt.find_optimum_anum(poly_degree,best_guess)
```

##### 6.4.5 Graphing Landmark Data

In order to facilitate easy visualization, the `graphSet` and `graphData` classes manage graphing commands and any necessary data transformation.

**Todo:** Code sample for graphing functions

#### 6.5 Sample Classification

##### 6.5.1 Goal

Previously we went through the process of calculating *landmarks* in order to identify a reduced set of point that are comparable between samples. In order to learn more about which points distinguish control samples from experimental samples, we can build a statistical model that will use landmarks to classify control and experimental. After we have developed a classification model, we can look at which landmarks were most important for classification and use this information to identify regions of biological difference.

##### 6.5.2 Dimensionality Reduction

Currently, your dataset may have several hundred landmark points that describe the shape and variability of the data. However, it is likely that the number of landmark points outnumbers the number of samples. Statistical modeling techniques are most effective when the number of points is smaller than the number samples that are training the model. In this situation we can think of each landmark point as a dimension of the data. In order to reduce the dimensionality of the data (e.g. the number of landmark points), we will use principle component analysis (PCA) to identify a reduced set of components (new dimensions) that capture all of the variability that is present in the landmark points. Each component is a mixture of landmark points with some points having greater influence than others.

##### 6.5.3 Coding Instructions

```
import deltascope
import pandas as pd

#Read csv file that contains landmark data for both sample groups
df = pd.read_csv(landmark_file)

#Create the tree classifier object
tc = deltascope.treeClassifier(df)

#Apply pca to automatically reduce the dimensionality to the optimum number of␣
↪dimensions
tc.apply_pca()

#Fit the classifier based on landmarks
tc.fit_classifier()

#Visualize the landmarks that had the highest impact on the classifier
tc.plot_top_components(index=10)
```

#### 6.6 Checklist

1. Convert data to HDF5 files containing a single sample/channel per file
2. Train Ilastik on each channel and process all files to create '_Probability.h5' files

#### 6.7 Parameter Reference

##### 6.7.1 Transformation

Default: `2`

> **Warning:** The infrastructure to support degrees other than 2 is not currently in place. Check here for updates.

##### 6.7.2 Landmark Calculation

**anum**

This integer specifies the number of divisions along the alpha axis when calculating landmarks. See *Selecting anum* for guidance on setting this parameter.

Example: `20`

**tsize**

This parameter sets the size of each radial wedge in the landmark calculation. The program works in radians so this parameter should be a float that can evenly divide into 2Pi. We have found that Pi/4 (45º) is a biologically appropriate division for our typical structures.

Example: `0.79`

**percbins**

This parameter is a list of integers that specifies what percentile should be used to calculate the distribution of points along r.

Example: `[50]`

#### 6.8 Useful Resources

##### 6.8.1 Command line tutorials

- A basic introduction to the command line interface
- An introduction to the command line interface for windows Powershell
- An introduction to the command line interface for Linux and MacOS Bash

##### 6.8.2 Pip package installer

- An introduction to pip pythons package installer
- A discussion of the differences of pip and conda the package manager that accompanies Anaconda

##### 6.8.3 Version control with Git and Github

- A very basic introduction to git and version control
- A discussion of why and how to use Github
- A cheat sheet for running Git from the command line

##### 6.8.4 Jupyter Notebook

- A cheat sheet for using Jupyter Notebook

#### 6.9 Frequently Asked Questions

##### 6.9.1 My file paths are causing errors?

Try changing each individual slash in your path to have two slashes.

##### 6.9.2 What is a `.psi` file?

A `.psi` file is similar in structure to a comma separated value (`.csv`) file with the addition of header text that defines metadata for the file. For example:

```
# PSI Format 1.0
#
# column[0] = "Id"
# column[1] = "x"
# column[2] = "y"
# column[3] = "z"
# column[4] = "ac"
# symbol[4] = "A"
# type[4] = float
# column[5] = "r"
# symbol[5] = "R"
# type[5] = float
# column[6] = "theta"
# symbol[6] = "T"
# type[6] = float
52337 0 0
1 0 0
0 1 0
0 0 1
```

#### 6.10 Batch Processing: Transformation

`mp-transformation.py` can be run from the command line after setting parameters in `mp-transformation-config.json`. When you run the script, you only need to provide the path to the config file as an argument

```
$ python mp-transformation.py "C:\\path\\to\\mp-transformation-config.json"
```

In addition to the parameters described in Parameter Reference, `mp-transformation-config.json` requires a set of additional parameters:

**rootdir**
: **Required**: String specifying the complete path to a directory where an output folder should be created

**expname**
: **Required**: String specifying the experiment name that will be incorporated into output files

**c1-dir**
: **Required**: String specifying the path to the directory containing the _`Probabilities.files` for the structural channel

**c1-key**
: **Required**: String that will serve as a key for the structural channel and will name the output files

**c2-dir**
: *Optional*: Same as *c1-dir*, but for an additional channel

**c2-key**
: *Optional*: Same as *c1-key*, but for an additional channel

**c3-dir**
: *Optional*: Same as *c1-dir*, but for an additional channel

**c3-key**
: *Optional*: Same as *c1-key*, but for an additional channel

**c4-dir**
: *Optional*: Same as *c1-dir*, but for an additional channel

**c4-key**
*Optional*: Same as *c1-key*, but for an additional channel

**twoD**
A boolean value that specifies, which PCA transformation functions will be used.

`True`: `brain.calculate_pca_median_2d()` and `brain.pca_transform_2d()` will be used to hold one axis constant, while the other two are realigned with PCA

`False`: `brain.calculate_pca_median()` and `brain.pca_transform_3d()` will be used to transform and realign samples in all three dimensions

##### 6.10.1 API

`deltascope.mpTransformation.`**check_nums**(*P*)
Check that the numbers of files selected by the same list index match Raises an error if file numbers are mismatched

:param `paramClass` P: Object containing all variables from config file

**class** `deltascope.mpTransformation.`**paramsClass**(*path*)
A class to read and validate parameters for multiprocessing transformation. Validated parameters can be read as attributes of the object

**add_outdir**(*path*)
Add out directory as an attribute of the class

**Parameters** **path** (*str*) – Complete path to the output directory

**check_config**(*D*, *path*)
Check that each parameter in the config file is correct and raise an error if it isn't

**Parameters**

- **D** (*dict*) – Dictionary containing parameters from the config file
- **path** (*str*) – Complete filepath to the config file

`deltascope.mpTransformation.`**process**(*num*, *P=None*)
Run through the processing steps for a single sample through saving psi files

**Parameters** **num** (*int*) – Index of the file that is currently being processed

:param `paramClass` P: Object containing all variables from config file

#### 6.11 API

**class** `deltascope.`**brain**
Object to manage biological data and associated functions.

**Returns** Pandas DataFrame with xyz and probability value for each point

**find_arclength**(*xc*)
Calculate arclength by integrating the derivative of the math model in xy plane

$$\int_{vertex}^{point} \sqrt{1 + (2ax + b)^2}$$

**Parameters** **row** (*float*) – Postion in the x axis along the curve

**Returns** Length of the arc along the curve between the row and the vertex

**Return type** float

**find_distance**(*t*, *point*)
Find euclidean distance between math model(t) and data point in the xy plane

**Parameters**

- **t** (*float*) – float value defining point on the line
- **point** (*array*) – array [x,y] defining data point

**Returns** distance between the two points

**Return type** float

**find_min_distance**(*row*)
Find the point on the curve that produces the minimum distance between the point and the data point using scipy.optimize.minimize(`brain.find_distance()`)

**Parameters** **row** (*pd.Series*) – row from dataframe in the form of a pandas Series

**Returns** point in the curve (xc, yc, zc) and r

**Return type** floats

**find_r** (*row*, *zc*, *yc*, *xc*)
Calculate r using the Pythagorean theorem

**Parameters**

- **row** (*pd.Series*) – row from dataframe in the form of a pandas Series
- **yc** (*float*) – Y position of the closest point in the curve to the data point
- **zc** (*float*) – Z position of the closest point in the curve to the data point
- **xc** (*float*) – X position of hte closest point in the curve to the data point

**Returns** r, distance between the point and the model

**Return type** float

**find_theta** (*row*, *zc*, *yc*)
Calculate theta for a row containing data point in relationship to the xz plane

**Parameters**

- **row** (*pd.Series*) – row from dataframe in the form of a pandas Series
- **yc** (*float*) – Y position of the closest point in the curve to the data point
- **zc** (*float*) – Z position of the closest point in the curve to the data point

**Returns** theta, angle between point and the model plane

**Return type** float

**fit_model** (*df*, *deg*, *fit_dim*)
Fit model to dataframe

**Parameters**

- **df** (*pd.DataFrame*) – Dataframe containing at least x,y,z
- **deg** (*int*) – Degree of the function that should be fit to the model
- **fit_dim** (*array*) – Array of length two containing two strings describing the first and second axis for fitting the model, e.g. ['x','z']

**Returns** math model

**Return type** *math_model*

**flip_data** (*df*)
Rotate data by 180 degrees

**Parameters** **df** (*dataframe*) – Pandas dataframe containing x,y,z data

**Returns** Rotated dataframe

**integrand** (*x*)
Function to integrate to calculate arclength

**Parameters** **filepath** (*str*) – Filepath to hdf5 probability file

**Returns** Creates the variable `brain.raw_data`

**raw_data**
Array of shape [z,y,x] containing raw probability data

**setup_test_data** (*size=None, gthresh=0.5, scale=[1, 1, 1], microns=[0.16, 0.16, 0.21], mthresh=0.2, radius=20, comp_order=[0, 2, 1], fit_dim=['x', 'z'], deg=2*)
Setup a test dataset to use for testing transform coordinates :param int size: Number of points to sample for the test dataset

**subset_data** (*df*, *sample_frac=0.5*)
Takes a random sample of the data based on the value between 0 and 1 defined for sample_frac

Creates the variable `brain.subset`

**Parameters**

- **pd.DataFrame** – Dataframe which will be sampled
- **sample_frac** (*float*) – (or None) Value between 0 and 1 specifying proportion of the dataset that should be randomly sampled for plotting

**subset**
Random sample of the input dataframe

**transform_coordinates** ()
Transform coordinate system so that each point is defined relative to math model by (alpha,theta,r) (only applied to `brain.df_align`)

**Returns** appends columns r, xc, yc, zc, ac, theta to `brain.df_align`

deltascope.**calc_variance** (*anum*, *dfs*)
Calculate the variance between samples according to bin position and variance between adjacent bins

**Parameters**

- **anum** (*int*) – Number of bins which the arclength axis should be divided into
- **dfs** (*dict*) – Dictionary of dfs which are going to be processed

**Returns** Two arrays: svar (anum,tnum) and bvar (anum*tnum,snum)

**Return type** np.array

deltascope.**calculate_area_error** (*pdf*, *Lkde*, *x*)
Calculate area between PDF and each kde in Lkde

**Parameters**

- **pdf** (*array*) – Array of probability distribution function that is the same shape as kdes in Lkde
- **Lkde** (*list*) – List of arrays of Kdes
- **x** (*array*) – Array of datapoints used to generate pdf and kdes

**Returns** List of error values for each kde in Lkde

deltascope.**calculate_models** (*Ldf*)
Calculate model for each dataframe in list and add to new dataframe

**Parameters** **Ldf** (*list*) – List of dataframes containing aligned data

**Returns** pd.Dataframe with a,b,c values for parabolic model

deltascope.**convert_to_arr** (*xarr*, *tarr*, *DT*, *mdf*, *Ldf=[]*)
Convert a pandas dataframe containing landmarks as columns and samples as rows into a 3D numpy array

The columns of *mdf* determine which landmarks will be saved into the array. Any additional dataframes that need to be converted can be included in Ldf

**Parameters**

- **xarr** (*np.array*) – Array containing all unique x values of landmarks in the dataset
- **tarr** (*np.array*) – Array containing all unique t values of landmarks in the dataset
- **DT** (*str*) – Either `r` or `pts` indicating which data type should be saved to the array

- **mdf** (*pd.DataFrame*) – Main landmark dataframe containing landmarks as columns and samples as rows
- **Ldf** (*list*) – List of additional pd.DataFrames that should also be converted to arrays

**Returns** Array of the main dataframe and list of arrays converted from Ldf

**save_psi**()
Save all channels into psi files following the naming scheme [`embryo.name`]_[`embryo.number`]_[*channel name*].psi

deltascope.**find_anchors**(*df*, *dim*)

**Parameters** **dim** (*str*) – either y or z

deltascope.**generate_kde**(*data*, *var*, *x*, *absv=False*)
Generate list of KDEs from either dictionary or list of data

**Parameters**

- **data** – pd.DataFrames to convert
- **var** (*str*) – Name of column to select from df
- **x** (*array*) – Array of datapoints to evaluate KDE on
- **absv** (*bool*) – (or None) Set to True to use absolute value of selected data for KDE calculation

**Type** dict or list

**Returns** List of KDE arrays

**class** deltascope.**landmarks**(*percbins=[10, 50, 90]*, *rnull=15*)
Class to handle calculation of landmarks to describe structural data

**Parameters**

- **percbins** (*list*) – (or None) Must be a list of integers between 0 and 100
- **rnull** (*int*) – (or None) When the r value cannot be calculated it will be set to this value

brain.**lm_wt_rf**
pd.DataFrame, which wildtype landmarks will be added to

brain.**lm_mt_rf**
pd.DataFrame, which mutant landmarks will be added to

brain.**rnull**
Integer specifying the value which null landmark calculations will be set to

brain.**percbins**
Integer specifying the percentiles which will be used to calculate landmarks

**calc_bins** (*Ldf*, *ac_num*, *tstep*)
Calculates alpha and theta bins based on ac_num and tstep

Creates `landmarks.acbins` and `landmarks.tbins`

> **Warning:** *tstep* does not handle scenarios where 2pi is not evenly divisible by tstep

**Parameters**

- **Ldf** (*dict*) – Dict dataframes that are being used for the analysis
- **ac_num** (*int*) – Integer indicating the number of divisions that should be made along alpha
- **tstep** (*float*) – The size of each bin used for alpha

**acbins**
List containing the boundaries of each bin along alpha based on *ac_num*

**tbins**
List containing the boundaries of each bin along theta based on *tstep*

**calc_mt_landmarks** (*df*, *snum*, *wt*)

> **Warning:** Deprecated function, but attempted to calculate mutant landmarks based on the number of points found in the wildtype standard

**calc_perc** (*df*, *snum*, *dtype*, *out*)
Calculate landmarks for a dataframe based on the bins and percentiles that have been previously defined

**Parameters**

- **df** (*pd.DataFrame*) – Dataframe containing columns x,y,z,alpha,r,theta
- **snum** (*str*) – String containing a sample identifier that can be converted to an integer
- **dtype** (*str*) – String describing the sample group to which the sample belongs, e.g. control or experimental

**Returns** pd.DataFrame with new landmarks appended

**calc_wt_reformat** (*df*, *snum*)

> **Warning:** Deprecated function, but includes code pertaining to calculating point based data

**class** deltascope.**math_model** (*model*)
Object to contain attributes associated with the math model of a sample

**Parameters** **model** (*array*) – Array of coefficients calculated by np.polyfit

**cf**
Array of coefficients for the math model

**p**
Poly1d function for the math model to allow calculation and plotting of the model

**class** deltascope.**paramsClass**(*path=None*, *dparams=None*)

A class to read and validate parameters for multiprocessing transformation. Validated parameters can be read as attributes of the object

**add_outdir**(*path*)

Add out directory as an attribute of the class

**Parameters** **path** (*str*) – Complete path to the output directory

**check_config**(*D*, *path*)

Check that each parameter in the config file is correct and raise an error if it isn't

**Parameters**

- **D** (*dict*) – Dictionary containing parameters from the config file
- **path** (*str*) – Complete filepath to the config file

deltascope.**process_sample**(*num*, *root*, *outdir*, *name*, *chs*, *prefixes*, *threshold*, *scale*, *deg*, *primary_key*, *comp_order*, *fit_dim*, *flip_dim*)

Process single sample through *brain* class and saves df to csv

> **Warning:** Out of date and will probably fail

**Parameters**

- **num** (*str*) – Sample number
- **root** (*str*) – Complete path to the root directory for this sample set
- **name** (*str*) – Name describing this sample set
- **outdir** (*str*) – Complete path to output directory
- **chs** (*array*) – Array containing strings specifying the directories for each channel
- **prefixes** (*array*) – Array containing strings specifying the file prefix for each channel
- **threshold** (*float*) – Value between 0 and 1 to use as a cutoff for minimum pixel value
- **scale** (*array*) – Array with three values representing the constant by which to multiply x,y,z respectively
- **deg** (*int*) – Degree of the function that should be fit to the model
- **primary_key** (*str*) – Key for the primary structural channel which PCA and the model should be fit too

deltascope.**read_psi**(*filepath*)

Reads psi file at the given filepath and returns data in a pandas DataFrame

**Parameters** **filepath** (*str*) – Complete filepath to file

**Returns** pd.Dataframe containing data

deltascope.**read_psi_to_dict**(*directory*, *dtype*)

Read psis from directory into dictionary of dfs with filtering based on dtype

**Parameters**

- **directory** (*str*) – Directory to get psis from
- **dtype** (*str*) – Usually 'AT' or 'ZRF1'

**Returns** Dictionary of pd.DataFrame

deltascope.**reformat_to_cart**(*df*)
Take a dataframe in which columns contain the bin parameters and convert to a cartesian coordinate system

**Parameters** **df** (*pd.DataFrame*) – Dataframe containing columns with string names that contain the bin parameter

**Returns** pd.DataFrame with each landmark as a row and columns: x,y,z,r,r_std,t,pts

deltascope.**rescale_variable**(*Ddfs*, *var*, *newvar*)
Rescale variable from -1 to 1 and save in newvar column on original dataframe

**Parameters**

- **Ddfs** (*dict*) – Dictionary of pd.DataFrames
- **var** (*str*) – Name of column to select from dfs
- **newvar** (*str*) – Name to use for new data in appended column

**Returns** Dictionary of dataframes containing column of rescaled data

deltascope.**subplot_lmk**(*ax*, *p*, *avg*, *sem*, *parr*, *xarr*, *tarr*, *dtype*, *Pn={'alpha': 0.3, 'cmap': 'Greys_r', 'mtc': 'r', 'tarr': None, 'wtc': 'b', 'xarr': None, 'zfb': 1, 'zln': 2, 'zpt': 3}*)
Plot a ribbon of average and standard error of the mean onto the subplot, *ax*

**Parameters**

- **ax** (*plt.Subplot*) – Matplotlib subplot onto which the data should be plotted
- **p** (*list*) – List of two theta values that should be plotted
- **avg** (*np.array*) – Array of shape (xvalues,tvalues) containing the average values of the data
- **sem** (*np.array*) – Array of shape (xvalues,tvalues) containing the standard error of the mean values of the data
- **parr** (*np.array*) – Array of shape (xvalues,tvalues) containing the p values for the data
- **dtype** (*str*) – String describing sample type
- **Pn** – Dictionary containing the following values: 'zln':2,'zpt':3,'zfb':1,'wtc':'b','mtc':'r','alpha':0.3,'cmap':'Greys_r'

**Type** dict or None

deltascope.**write_data**(*filepath*, *df*)
Writes data in PSI format to file after writing header using `write_header()`. Closes file at the conclusion of writing data.

**Parameters**

- **filepath** (*str*) – Complete filepath to output file
- **df** (*pd.DataFrame*) – dataframe containing columns x,y,z,ac,r,theta

deltascope.**write_header**(*f*)
Writes header for PSI file with columns Id,x,y,z,ac,r,theta

**Parameters** **f** (*file*) – file object created by 'open(filename,'w')'

CHAPTER 7

### Indices and tables

- genindex
- modindex
- search

### Python Module Index

## d

deltascope, 26
deltascope.mpTransformation, 26

### Index

## A

acbins (*deltascope.landmarks attribute*), 35
add_aligned_df() (*deltascope.brain method*), 26
add_channel() (*deltascope.embryo method*), 33
add_outdir() (*deltascope.mpTransformation.paramsClass method*), 26
add_outdir() (*deltascope.paramsClass method*), 36
add_psi_data() (*deltascope.embryo method*), 33
add_thresh_df() (*deltascope.brain method*), 27
align_data() (*deltascope.brain method*), 27
anum, 20
anum, 18, 20

## B

brain (*class in deltascope*), 26

## C

c1-dir, 25
c1-key, 25, 26
calc_bins() (*deltascope.landmarks method*), 34
calc_coord() (*deltascope.brain method*), 27
calc_mt_landmarks() (*deltascope.landmarks method*), 35
calc_perc() (*deltascope.landmarks method*), 35
calc_variance() (*in module deltascope*), 32
calc_wt_reformat() (*deltascope.landmarks method*), 35
calculate_area_error() (*in module deltascope*), 32
calculate_models() (*in module deltascope*), 32
calculate_pca_median() (*deltascope.brain method*), 27
calculate_pca_median_2d() (*deltascope.brain method*), 28
cf (*deltascope.math_model attribute*), 35
check_config() (*deltascope.mpTransformation.paramsClass method*), 26
check_config() (*deltascope.paramsClass method*), 36
check_nums() (*in module deltascope.mpTransformation*), 26
chnls (*deltascope.embryo attribute*), 33
comporder, 15, 16, 23
convert_to_arr() (*in module deltascope*), 32
create_dataframe() (*deltascope.brain method*), 28

## D

deg, 16
deltascope (*module*), 26
deltascope.mpTransformation (*module*), 26
df_align (*deltascope.brain attribute*), 27
df_scl (*deltascope.brain attribute*), 31
df_thresh (*deltascope.brain attribute*), 31

## E

embryo (*class in deltascope*), 33
environment variable
    anum, 20
environment variable
    anum, 18, 20, 23
    c1-dir, 25
    c1-key, 25, 26
    c2-dir, 25
    c2-key, 25
    c3-dir, 25
    c3-key, 25
    c4-dir, 25
    c4-key, 26
    comporder, 15, 16, 23
    deg, 16, 23
    expname, 25
    fitdim, 16, 23
    genthresh, 13, 14, 16, 22, 23
    medthresh, 14, 16, 22
    micron, 13
    microns, 22

percbins, 18, 24
radius, 16, 23
rootdir, 25
scale, 13, 22
tsize, 18, 23
twoD, 26

## F

find_anchors() (*in module deltascope*), 34
find_arclength() (*deltascope.brain method*), 28
find_distance() (*deltascope.brain method*), 28
find_min_distance() (*deltascope.brain method*), 28
find_r() (*deltascope.brain method*), 29
find_theta() (*deltascope.brain method*), 29
fit_model() (*deltascope.brain method*), 29
fitdim, 16, 23
flip_data() (*deltascope.brain method*), 29

## G

generate_kde() (*in module deltascope*), 34
genthresh, 13, 14, 16, 23

## I

integrand() (*deltascope.brain method*), 29

## L

landmarks (*class in deltascope*), 34
lm_mt_rf (*deltascope.landmarks.brain attribute*), 34
lm_wt_rf (*deltascope.landmarks.brain attribute*), 34

## M

math_model (*class in deltascope*), 35
median (*deltascope.brain attribute*), 27
medthresh, 14, 16
micron, 13
microns, 22
mm (*deltascope.brain attribute*), 27

## N

name (*deltascope.embryo attribute*), 33
number (*deltascope.embryo attribute*), 33

## O

outdir (*deltascope.embryo attribute*), 33

## P

p (*deltascope.math_model attribute*), 35
paramsClass (*class in deltascope*), 35
paramsClass (*class in deltascope.mpTransformation*), 26
pca_transform_2d() (*deltascope.brain method*), 29
pca_transform_3d() (*deltascope.brain method*), 30
pcamed (*deltascope.brain attribute*), 27
percbins, 18
percbins (*deltascope.landmarks.brain attribute*), 34
plot_projections() (*deltascope.brain method*), 30
preprocess_data() (*deltascope.brain method*), 31
process() (*in module deltascope.mpTransformation*), 26
process_alignment_data() (*deltascope.brain method*), 31
process_channels() (*deltascope.embryo method*), 33
process_sample() (*in module deltascope*), 36

## R

radius, 16, 23
raw_data (*deltascope.brain attribute*), 31
read_data() (*deltascope.brain method*), 31
read_psi() (*in module deltascope*), 36
read_psi_to_dict() (*in module deltascope*), 36
reformat_to_cart() (*in module deltascope*), 37
rescale_variable() (*in module deltascope*), 37
rnull (*deltascope.landmarks.brain attribute*), 34

## S

save_projections() (*deltascope.embryo method*), 34
save_psi() (*deltascope.embryo method*), 34
scale, 13
setup_test_data() (*deltascope.brain method*), 31
subplot_lmk() (*in module deltascope*), 37
subset (*deltascope.brain attribute*), 32
subset_data() (*deltascope.brain method*), 31

## T

tbins (*deltascope.landmarks attribute*), 35
threshold (*deltascope.brain attribute*), 31
transform_coordinates() (*deltascope.brain method*), 32
tsize, 18

## W

write_data() (*in module deltascope*), 37
write_header() (*in module deltascope*), 37
